# Comprehensive patient-level classification and quantification of driver events in TCGA PanCanAtlas cohorts

**DOI:** 10.1101/2021.03.17.435828

**Authors:** Alexey D. Vyatkin, Danila V. Otnyukov, Sergey V. Leonov, Aleksey V. Belikov

**Author notes:** In memory of Alexey D. Vyatkin, 13.07.1997-14.02.2021.

## Abstract

There is a growing need to develop novel therapeutics for targeted treatment of cancer. The prerequisite to success is the knowledge about which types of molecular alterations are predominantly driving tumorigenesis. To shed light onto this subject, we have utilized the largest database of human cancer mutations – TCGA PanCanAtlas, multiple established algorithms for cancer driver prediction (2020plus, CHASMplus, CompositeDriver, dNdScv, DriverNet, HotMAPS, OncodriveCLUSTL, OncodriveFML) and developed four novel computational pipelines: SNADRIF (Single Nucleotide Alteration DRIver Finder), GECNAV (Gene Expression-based Copy Number Alteration Validator), ANDRIF (ANeuploidy DRIver Finder) and PALDRIC (PAtient-Level DRIver Classifier). A unified workflow integrating all these pipelines, algorithms and datasets at cohort and patient levels was created.

We have found that there are on average 12 driver events per tumour, of which 0.6 are single nucleotide alterations (SNAs) in oncogenes, 1.5 are amplifications of oncogenes, 1.2 are SNAs in tumour suppressors, 2.1 are deletions of tumour suppressors, 1.5 are driver chromosome losses, 1 is a driver chromosome gain, 2 are driver chromosome arm losses, and 1.5 are driver chromosome arm gains. The average number of driver events per tumour increases with age (from 7 to 15) and cancer stage (from 10 to 15) and varies strongly between cancer types (from 1 to 24). Patients with 1 and 7 driver events per tumour are the most frequent, and there are very few patients with more than 40 events. In tumours having only one driver event, this event is most often an SNA in an oncogene. However, with increasing number of driver events per tumour, the contribution of SNAs decreases, whereas the contribution of copy-number alterations and aneuploidy events increases.

**Author Summary:** By analysing genomic and transcriptomic data from 10000 cancer patients through our custom-built computational pipelines and previously established third-party algorithms, we have found that half of all driver events in a patient’s tumour appear to be gains and losses of chromosomal arms and whole chromosomes. We therefore suggest that future therapeutics development efforts should be focused on targeting aneuploidy. We have also found that approximately a third of driver events in a patient are whole gene amplifications and deletions. Thus, therapies aimed at copy-number alterations also appear very promising. On the other hand, drugs aiming at point mutations are predicted to be less successful, as these alterations are responsible for just a couple of drivers per tumour. One notable exception are patients having only one driver event in their tumours, as this event is almost always a single nucleotide alteration in an oncogene.

## Introduction

Driver events are the molecular and cellular events that drive cancer progression, often called driver mutations [1]. The knowledge about the spectrum and quantity of driver events in individual patients and groups of patients is crucial to inform design and selection of targeted therapeutics. Historically, most attention has been devoted to point mutations or single nucleotide alterations (SNAs), as most driver prediction algorithms work only with this class of driver events. As the average numbers of driver SNAs per tumour in various cancer types have been estimated before and shown to vary within 1-10 range, with the average across all cancer types of 4 [2], this defined the current thinking that the number of driver events per tumour is relatively low.

However, SNA drivers represent only the tip of the iceberg, and are likely not the most crucial contribution to cancer progression. SNAs are very well tolerated by cancer cells [2]. It is known that cancer cells contain large numbers of deletions and amplifications (often called copy number alterations, or CNAs), translocations, inversions, full chromosome and chromosomal arm gains and losses (aneuploidy), as well as epigenetic modifications [3–5], but the driver potential of these alterations has been left almost unexplored due to the scarcity of driver prediction algorithms for these classes of events. Here, we set the goal to predict, classify and quantify as many different classes of driver events as possible, using clear and straightforward principles, and developed custom computational pipelines for this purpose.

Another shortcoming of the majority of existing driver prediction algorithms is that they work at the cohort level, i.e. they predict driver mutations for large groups of patients, usually having a particular cancer type. This does not allow to look at the composition of driver events in individual patients. We wrote specific scripts to convert cohort-level predictions into patient-level events, which also allowed seamless integration of the results from various third-party algorithms, including 2020plus [6], CHASMplus [7], CompositeDriver [8], dNdScv [2], DriverNet [9], HotMAPS [10], OncodriveCLUSTL [11], and OncodriveFML [12].. This is useful, as each individual driver prediction algorithm has its own strengths and shortcomings, and combining results from multiple algorithms allows to obtain more complete and balanced picture, ensuring that less driver mutations have been missed. Having patient-level data is also convenient for analysing driver event composition in various demographic and clinical groups (e.g. patients grouped by age, sex or cancer stage) without the need to rerun the driver prediction algorithms on subgroups of patients, which would otherwise not only consume additional time and resources but, most importantly, reduce the predictive power. Patient-level data also allow to study heterogeneity of cancer patient cohorts, e.g. by using distribution histograms.

In addition to these existing driver prediction algorithms, we decided to create our own, using clear and simple rules to have an internal reference standard. We called this algorithm SNADRIF – SNA DRIver Finder. It is a Python 3.7 software package that predicts cancer driver genes from the TCGA PanCanAtlas SNA data and classifies them into oncogenes and tumour suppressors. Driver prediction is based on calculating the ratio of nonsynonymous SNAs to silent SNAs [2], whereas driver classification is based on calculating the ratio of hyperactivating SNAs to inactivating SNAs [13]. Bootstrapping is used to calculate statistical significance and Benjamini– Hochberg procedure is used to keep false discovery rate under 5%.

Copy-number alterations (CNA) usually involve large chunks of DNA containing tens or hundreds of genes, which makes CNA data not very useful for uncovering individual driver genes. Nevertheless, it is an important source of information about amplifications and deletions of driver genes predicted from SNA data. However, due to CNA data coarseness, we wanted to clarify the actual copy number status of individual genes using mRNA and miRNA expression data available at TCGA PanCanAtlas. For this purpose, we created another Python 3.7 software package called GECNAV - Gene Expression-based CNA Validator. CNA validation is based on comparing the CNA status of a given gene in a given patient to expression of this gene in this patient relative to the median expression of this gene across all patients.

Aneuploidy – chromosome arm and full chromosome gains and losses – makes a substantial contribution to the number of driver alterations per tumour, however, there are no existing algorithms to differentiate driver aneuploidies from passenger ones. Therefore, we built our own pipeline called ANDRIF - ANeuploidy DRIver Finder. It is a Python 3.7 software package that predicts driver chromosomal arm or full chromosome gains or losses from the TCGA PanCanAtlas aneuploidy data. Driver prediction is based on calculating the average alteration status for each arm or chromosome in each cancer type. Bootstrapping is used to obtain the realistic distribution of the average alteration statuses under the null hypothesis. Benjamini– Hochberg procedure is performed to keep the false discovery rate under 5%.

Finally, we needed an algorithm to integrate all data on driver alterations from different algorithms - our own and third-party. We called this algorithm PALDRIC - PAtient-Level DRIver Classifier. It is a Python 3.7 software package that translates cohort-level lists of driver genes or mutations to the patient level, classifies driver events according to the molecular causes and functional consequences, and presents comprehensive statistics on various kinds of driver events in various demographic and clinical groups of patients. PALDRIC allows to natively combine outputs of various third-party algorithms to investigate optimal combinations. Our overall workflow can be seen in **Fig 1**.

**Fig 1.**
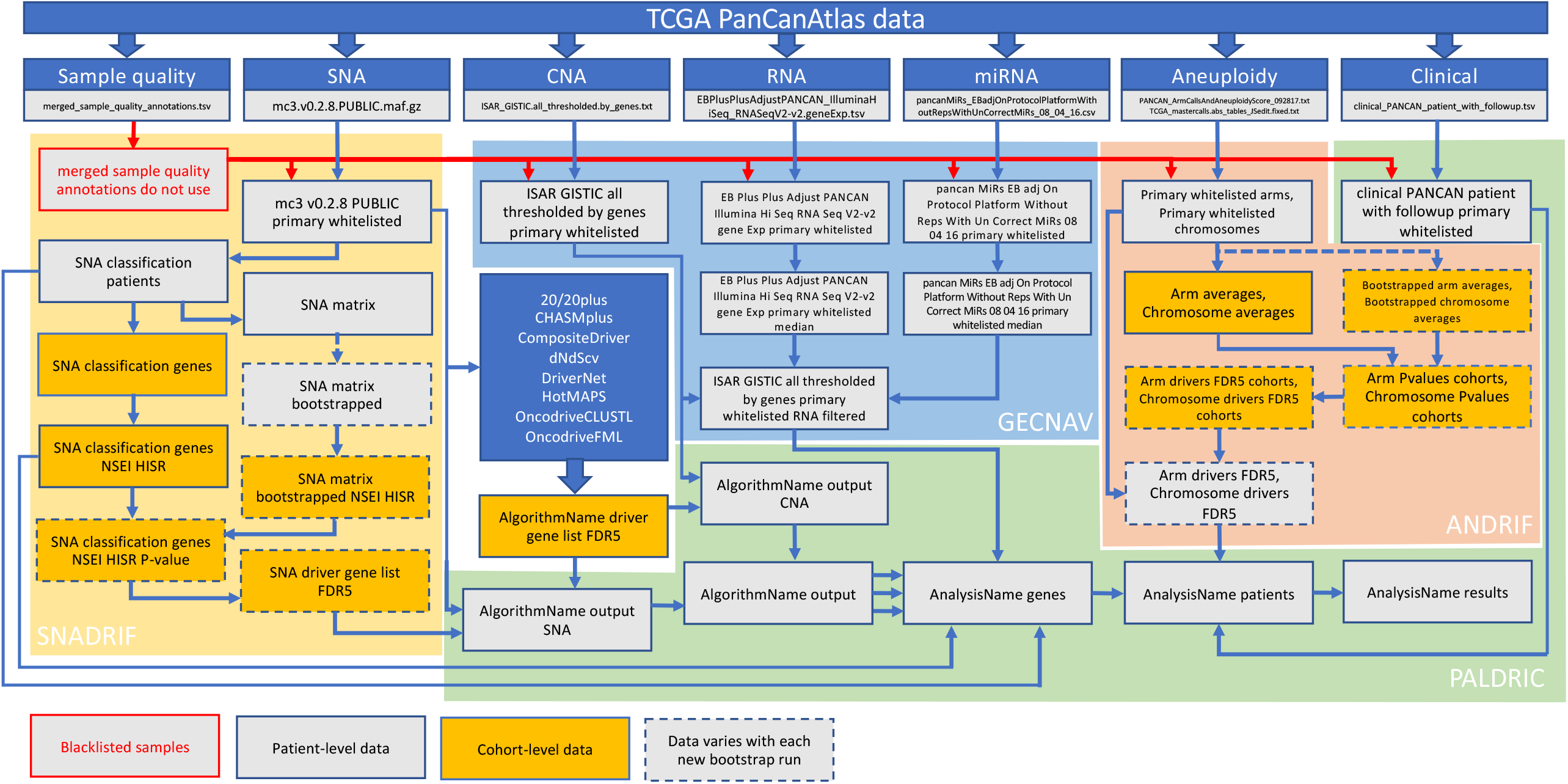
Workflow. Actual names of the files generated and used in the workflow are shown.

## Results

To get two different perspectives on the number and composition of driver events, we performed two different analyses. In the first one, we used the combination of results from SNADRIF, several third-party algorithms - 2020plus, CHASMplus, CompositeDriver, HotMAPS, OncodriveFML, and a consensus driver gene list from 26 algorithms [8], each of them applied to the whole TCGA PanCanAtlas dataset. We also used a list of COSMIC Cancer Gene Census Tier 1 genes affected by somatic SNAs and CNAs, as it represents the current gold standard of verified cancer drivers [14]. To minimize false positives, we used only genes that were predicted as drivers by at least two of our sources, including CGC and the list from Bailey et al. [8]. Benchmarking (see **S1 Files**) showed that such combinatory approach allows to identify more high-confidence driver genes than CGC or Bailey et al. lists contain (244 vs 182 vs 164, **Fig 2A**) and to recover more CGC genes (62%) than any individual algorithm or even Bailey et al list (**Fig 2B**). The overlap of three sources recovered fewer CGC genes (45%, Fig 2B), therefore we concluded that the overlap of two sources is optimal. SNADRIF sensitivity was comparable to CompositeDriver (**Fig 2B**), and it contributed 16 driver genes (6.5%) to the overall consensus, i.e. it validated 16 genes that were found only in one other source, including two genes (MYCL and TBL1XR1) that were present only in CGC and not predicted by any third-party algorithm (**Table 1**). Unfortunately, application of driver gene lists discovered through pancancer analysis equally to every cancer type results in unrealistically high numbers of driver events per patient, which is to be expected as this approach ignores tissue specificity of driver genes. Therefore, we present this analysis only as **S2 Files**.

**Fig 2.**
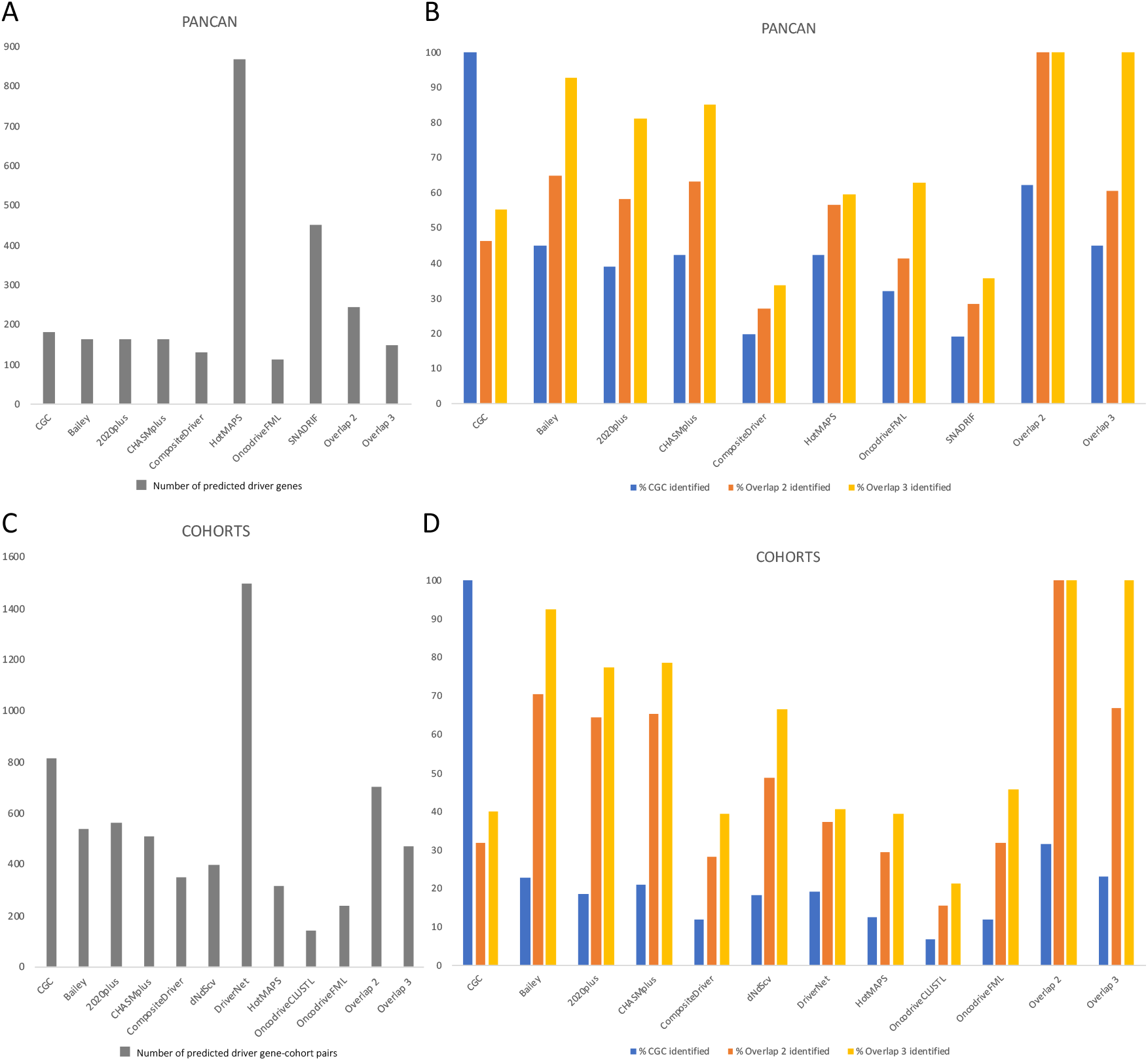
Benchmarking of driver prediction algorithms. (**A**) Number of driver genes predicted by each algorithm applied to pancancer dataset. CGC – COSMIC Cancer Gene Census Tier 1 genes affected by somatic SNAs and CNAs; Overlap 2 and Overlap 3 – genes predicted by at least 2 or 3 of the listed algorithms, respectively. (**B**) Sensitivity of algorithms applied to pancancer dataset, estimated as the percentage of genes from positive control lists identified. Positive control lists include CGC (blue), Overlap 2 (orange) and Overlap 3 (yellow), see legend for (A). (**C**) Number of driver gene-cohort pairs predicted by each algorithm applied individually to each cancer type (cohort). CGC - COSMIC Cancer Gene Census Tier 1 gene-cohort pairs affected by somatic SNAs and CNAs; Overlap 2 and Overlap 3 – gene-cohort pairs predicted by at least 2 or 3 of the listed algorithms, respectively. (**D**) Sensitivity of algorithms applied individually to each cancer type (cohort), estimated as the percentage of gene-cohort pairs from positive control lists identified. Positive control lists include CGC (blue), Overlap 2 (orange) and Overlap 3 (yellow), see legend for (C).

**Table 1.** Driver genes uniquely predicted by SNADRIF and one other source.

| Gene | Entrez Gene ID | Also predicted by |
| --- | --- | --- |
| AMD1 | 262 | HotMAPS |
| BMPR1A | 657 | 2020plus |
| CMTR2 | 55783 | OncodriveFML |
| ELL2 | 22936 | HotMAPS |
| HNRNPDL | 9987 | 2020plus |
| LYRM4 | 57128 | OncodriveFML |
| METTL14 | 57721 | HotMAPS |
| MYCL | 4610 | CGC |
| PPP3R1 | 5534 | HotMAPS |
| PTMA | 5757 | Bailey et al, 2018 |
| RPL22 | 6146 | Bailey et al, 2018 |
| SAMHD1 | 25939 | HotMAPS |
| TBL1XR1 | 79718 | CGC |
| TDG | 6996 | HotMAPS |
| TNNI3K | 51086 | HotMAPS |
| TTK | 7272 | HotMAPS |

In the second analysis we used the combination of results from 2020plus, CHASMplus, CompositeDriver, dNdScv, DriverNet, HotMAPS, OncodriveCLUSTL, OncodriveFML, the consensus list from [8] and a list of Cancer Gene Census Tier 1 genes affected by somatic SNAs and CNAs, applied separately to each cancer cohort of TCGA PanCanAtlas. Applying algorithms to individual cohorts allows to discover cancer type-specific drivers and avoid contamination by false positives, i.e. driver genes discovered during pancancer analysis that do not in reality play any role in a given cancer type. On the other hand, much fewer patients are available for cohort-specific analysis, and this decreases statistical power to discover new driver genes. Of note, our SNADRIF algorithm works best for pancancer analysis and struggles with small cohorts, due to scarcity of point mutations. However, when a combination of driver prediction algorithms is used, there are lower chances of missing an important driver gene even in a cohort-specific analysis, as algorithms based on differing principles complement each other. To minimize false positives, we used only genes that were predicted as drivers in the same cancer type by at least two of our sources, including CGC and the list from Bailey et al [8]. CGC attribution of driver genes to various cancer types is not very precise or comprehensive, therefore benchmarking of cohort-specific algorithms on CGC likely underestimates their true performance. Anyhow, benchmarking (see **S1 Files**) showed that our combinatory approach allows to recover more CGC gene-cohort pairs (32%) than any individual algorithm or even Bailey et al list (**Fig 2D**). Similarly to pancan analysis, the overlap of three sources recovered fewer CGC gene-cohort pairs (23%, **Fig 2D**), therefore we chose the overlap of two sources as optimal. Benchmarking also showed that our combinatory approach allows to identify more high-confidence driver gene-cohort pairs than are present on the Bailey et al. list (705 vs 537, **Fig 2C**). The results of this analysis would be presented in the following paragraphs, whereas the underlying data and additional graphs could be found in **S3 Files**.

**Fig 3** shows the average number of driver events of different classes in males and females. It could be seen that half of all driver events are chromosome- and arm-level gains and losses. Moreover, a third of all driver events are amplifications of oncogenes and deletions of tumour suppressors. Only one sixth of all driver events are SNAs. Although the overall difference between males and females in the average number of driver events is not significant (P=0.462, two-tailed heteroscedastic t-test, see **S4 Files**), there are significant differences in particular types of driver events. Males have significantly less SNAs in oncogenes (P=3.62*10^−15^) and in tumour suppressors (P=1.06*10^−16^), but significantly more amplifications of oncogenes (P=1.97*10^−6^), simultaneous occurrences of SNAs in one allele and deletions of the other allele in tumour suppressors (P=1.57*10^−5^), driver chromosome losses (P=0.01) and driver chromosome gains (P=3.5*10^−10^). High number of SNAs in females might be at least partially explained by the contribution from UCEC cohort (see **Fig 7**).

**Fig 3.**
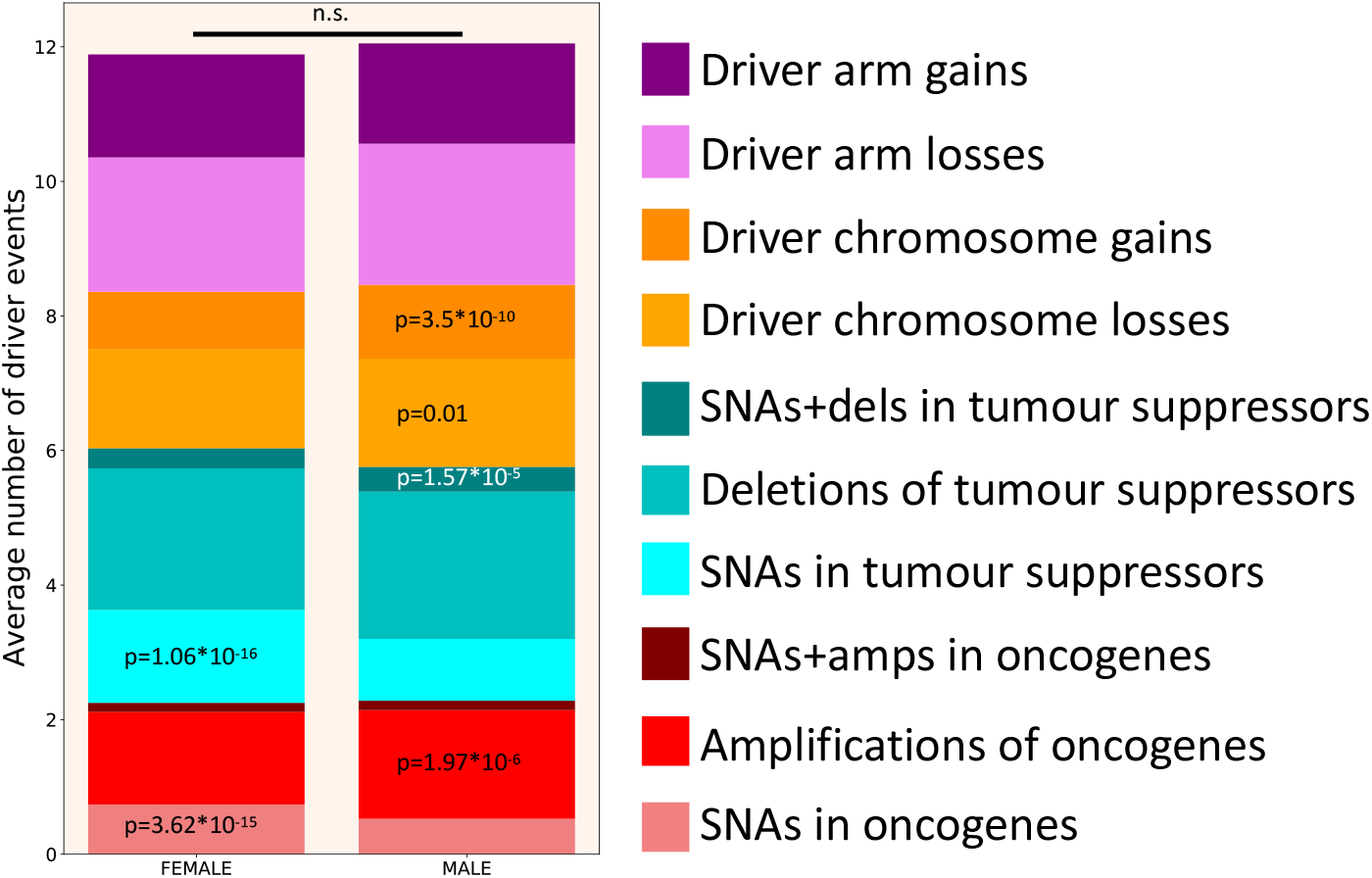
Driver event distribution by patient’s gender. Two-tailed heteroscedastic t-test was performed. p-values indicate a significant increase in the average number of driver events of a given type compared to the opposite gender. n.s. - difference not significant.

**Fig 4** demonstrates an increase in the number of driver events with age in males and females. It can be seen that in female patients the number of driver events increases dramatically from the earliest age until approximately 50 y.o., i.e. the age of menopause. After this, the growth of the number of driver events slows down. In males the picture is different. There is a high number of driver events in the earliest age group, due to the frequent occurrence of testicular cancer (TGCT) in this age group and a relatively high number of driver events in this cancer type (see **Fig 7**). The next age group has much lower number of drivers (P=0.041, one-tailed heteroscedastic t-test, see **S4 Files**), and like in females, their number increases with age, but until an older age, approximately 70 y.o., when their growth slows down. Surprisingly, the number of SNAs in oncogenes does not increase with age, at least since 30 y.o.

**Fig 4.**
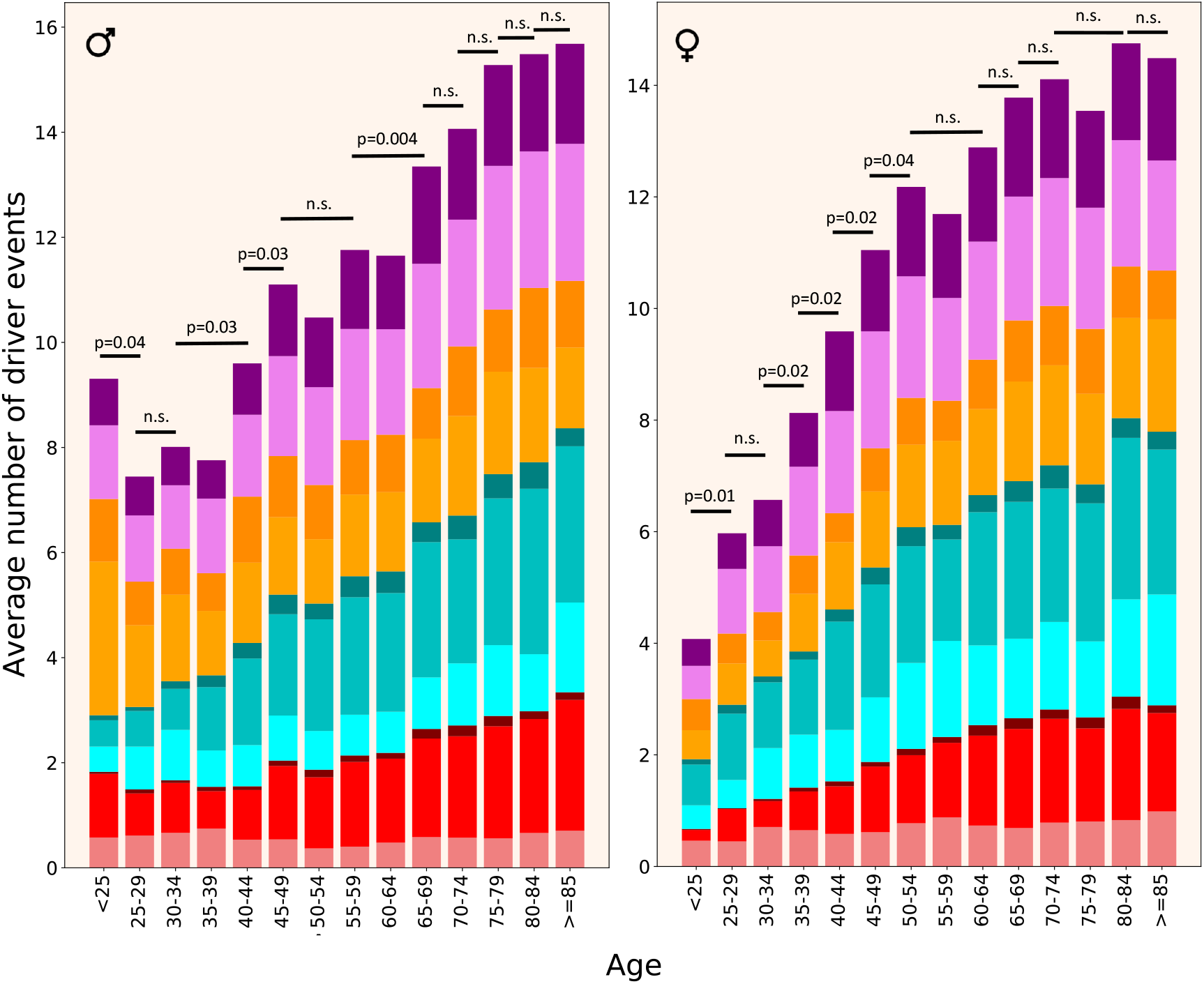
Driver event distribution by patient’s age. Colour coding as in Fig 3. One-tailed heteroscedastic t-test was performed. p-values indicate a significant difference in the average number of driver events between indicated age groups. n.s. - difference not significant.

**Fig 5** demonstrates that in females, the number of driver events increases dramatically from Stage I to Stage II (P=9*10^−15^, two-tailed heteroscedastic t-test, see **S4 Files**), after which stays more or less constant. It is interesting that this increase is due to CNAs and aneuploidy events but not due to point mutations. The number of SNAs actually decreases. In males, the increase in driver events with progression from Stage I to Stage IV is more gradual and chaotic. As in females, there is a prominent increase from Stage I to Stage II (P=4*10^−5^) but, for unclear reason, there appears to be less driver events in Stage III than in Stage II tumours (P=0.003). However, Stage IV tumours have more events than Stage II (P=3.3*10^−9^) and Stage III (P=4.9*10^−17^) tumours. Unlike in females, the increase is mediated also by SNAs, in addition to CNAs and aneuploidy events. In males, there are no significant changes in the number of driver chromosome losses between stages. This complicated picture might be explained by unequal representation of patients with different cancer stages amongst cancer types. Indeed, a closer look at the data (see **S3 Files**) shows that there are twice as many PRAD patients with Stage III than Stage II cancer, and because PRAD patients typically have a lower than average number of driver events (see **Fig 7**), this leads to the counterintuitive result of Stage III tumours having less driver events than Stage II tumours in the pooled analysis of male patients. Likewise, there are six times as many UCEC patients with Stage I than Stage II cancer (see **S3 Files**), and because UCEC patients typically have a higher than average number of driver SNAs (see **Fig 7**), this leads to the counterintuitive result of Stage II tumours having less SNA driver events than Stage I tumours in the pooled analysis of female patients.

**Fig 5.**
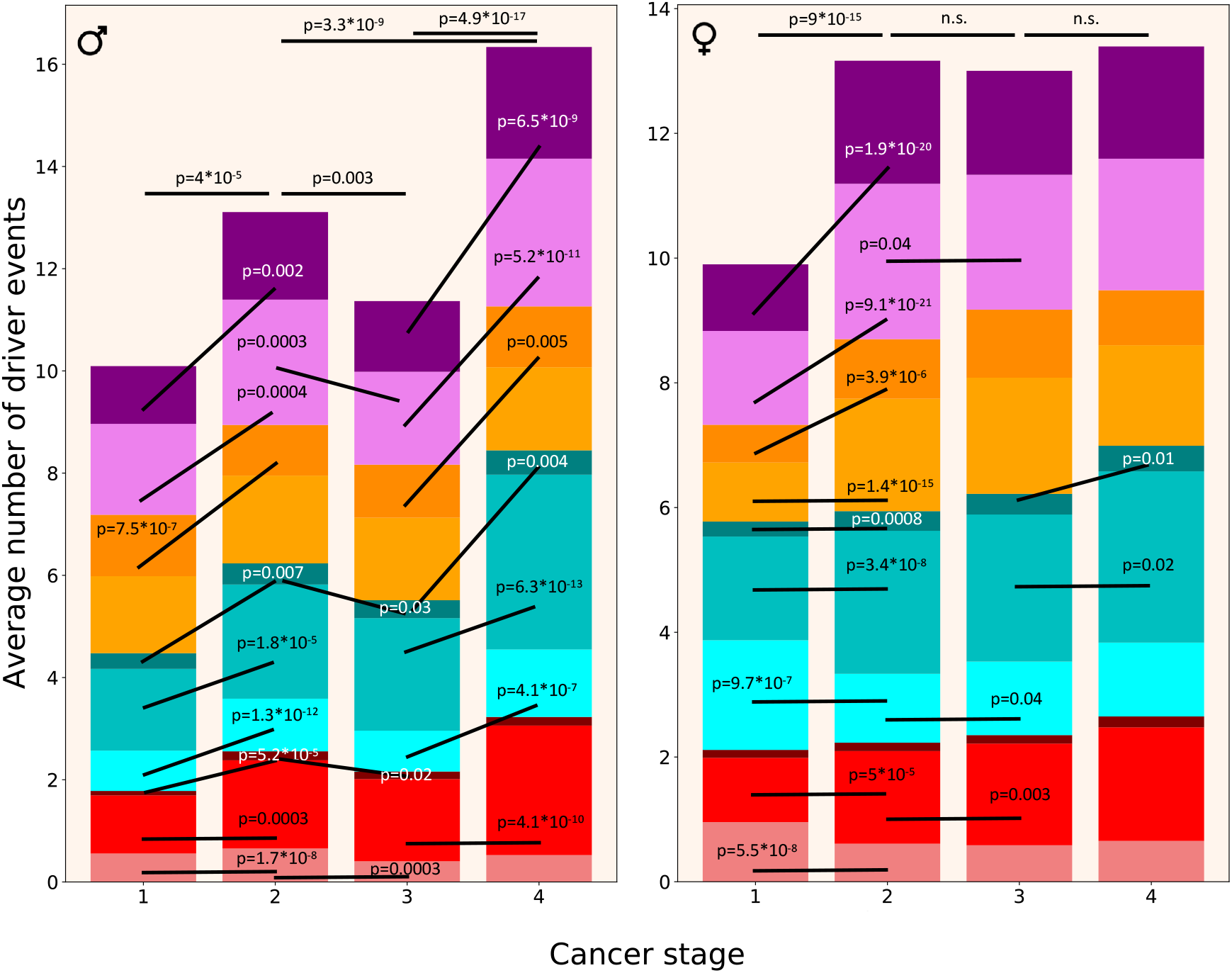
Driver event distribution by cancer stage. Colour coding as in Fig 3. Two-tailed heteroscedastic t-test was performed. p-values indicate a significant difference in the average number of driver events of a given type between the indicated stages. n.s. - difference not significant.

In **Fig 6** we aimed to show how different classes of driver events increase proportionally to the total number of driver events in a patient. It is striking that the number of SNAs in oncogenes does not increase, considering that if a tumour has only one driver event it is almost certain to be an SNA in an oncogene. There are very few patients with more than 30 driver events (see **Fig 8**), therefore the results for those groups become unreliable.

**Fig 6.**
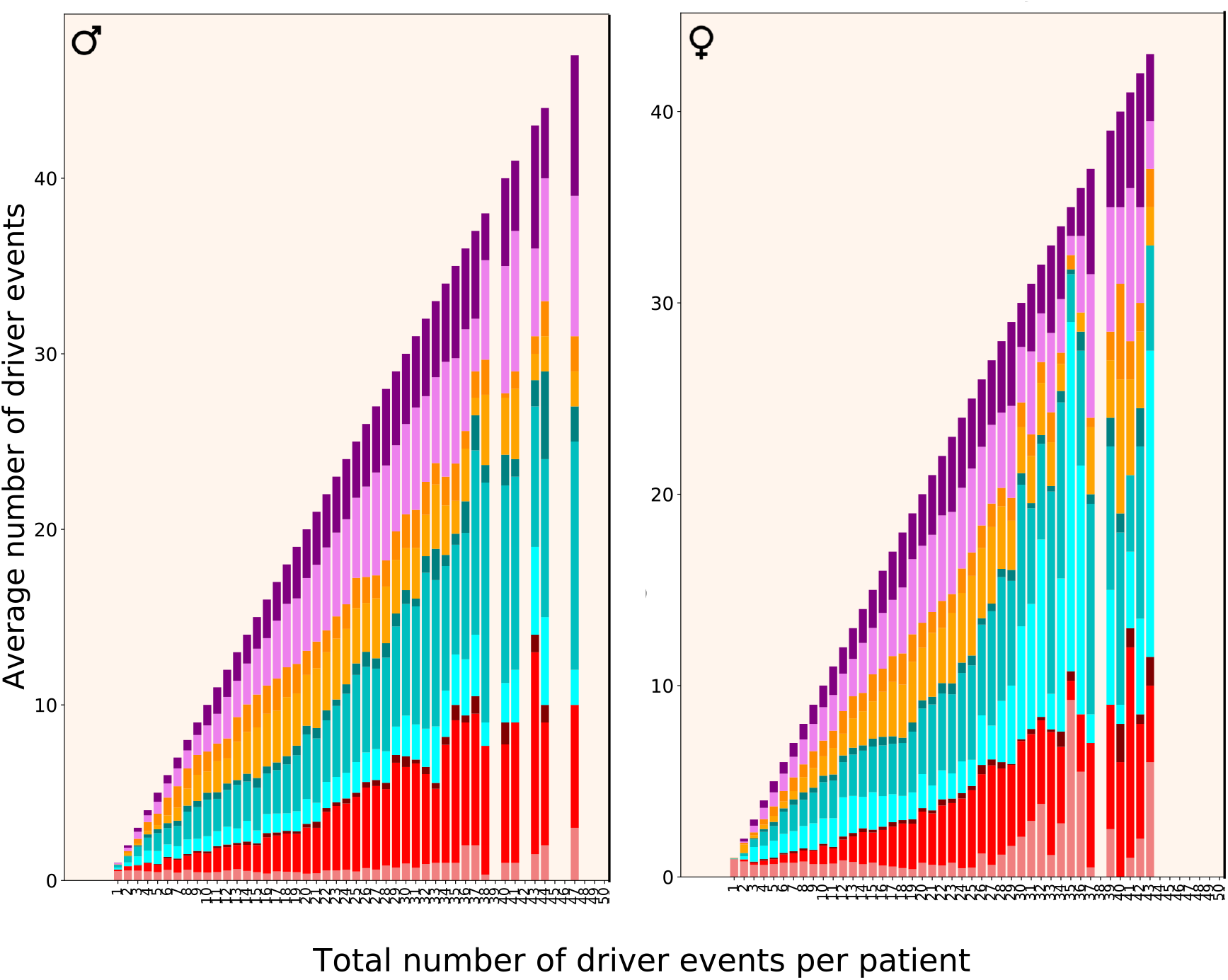
Driver event distribution by total number of driver events per patient. Colour coding as in Fig 3.

**Fig 7.**
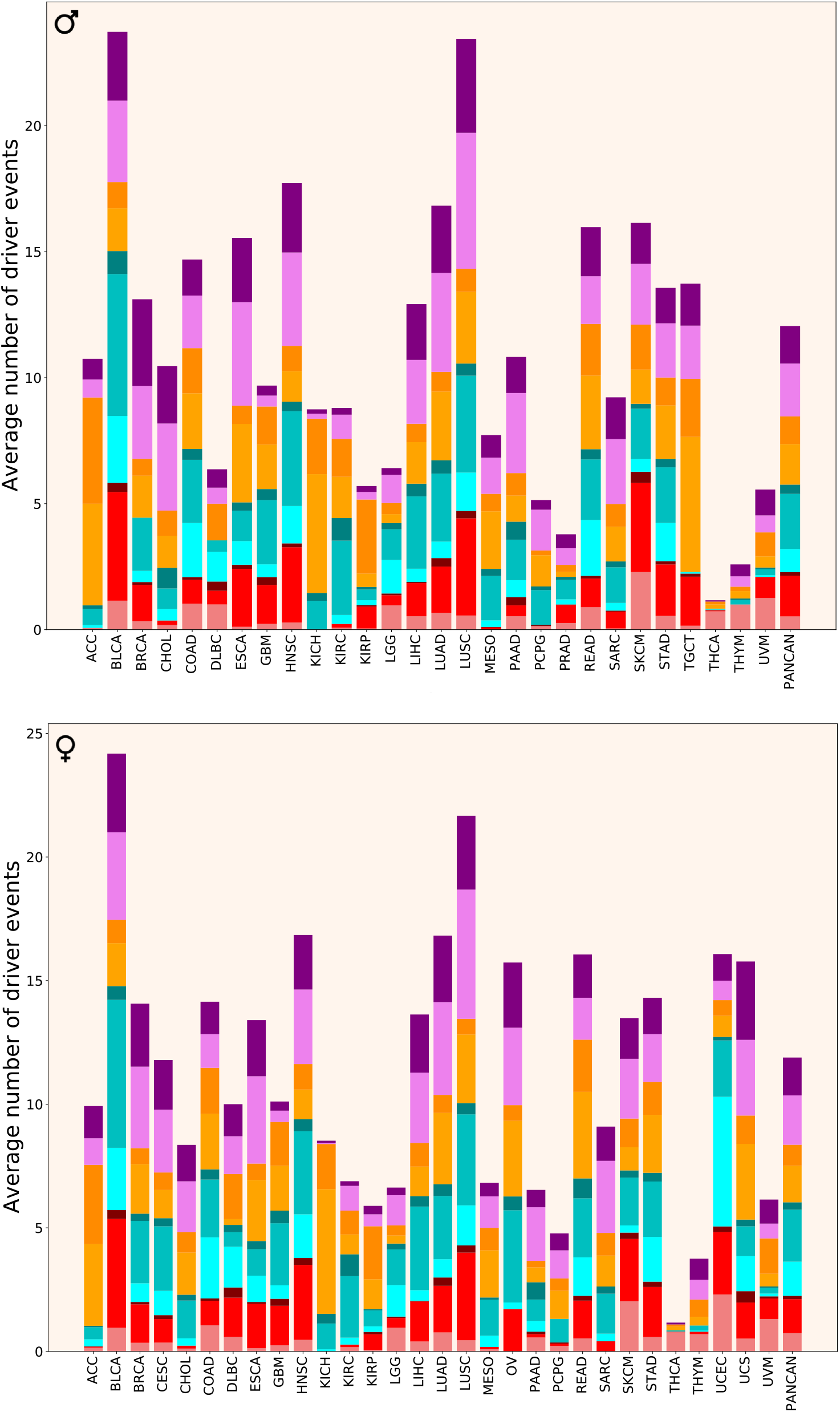
Driver event distribution by cancer type. Colour coding as in Fig 3.

**Fig 8.**
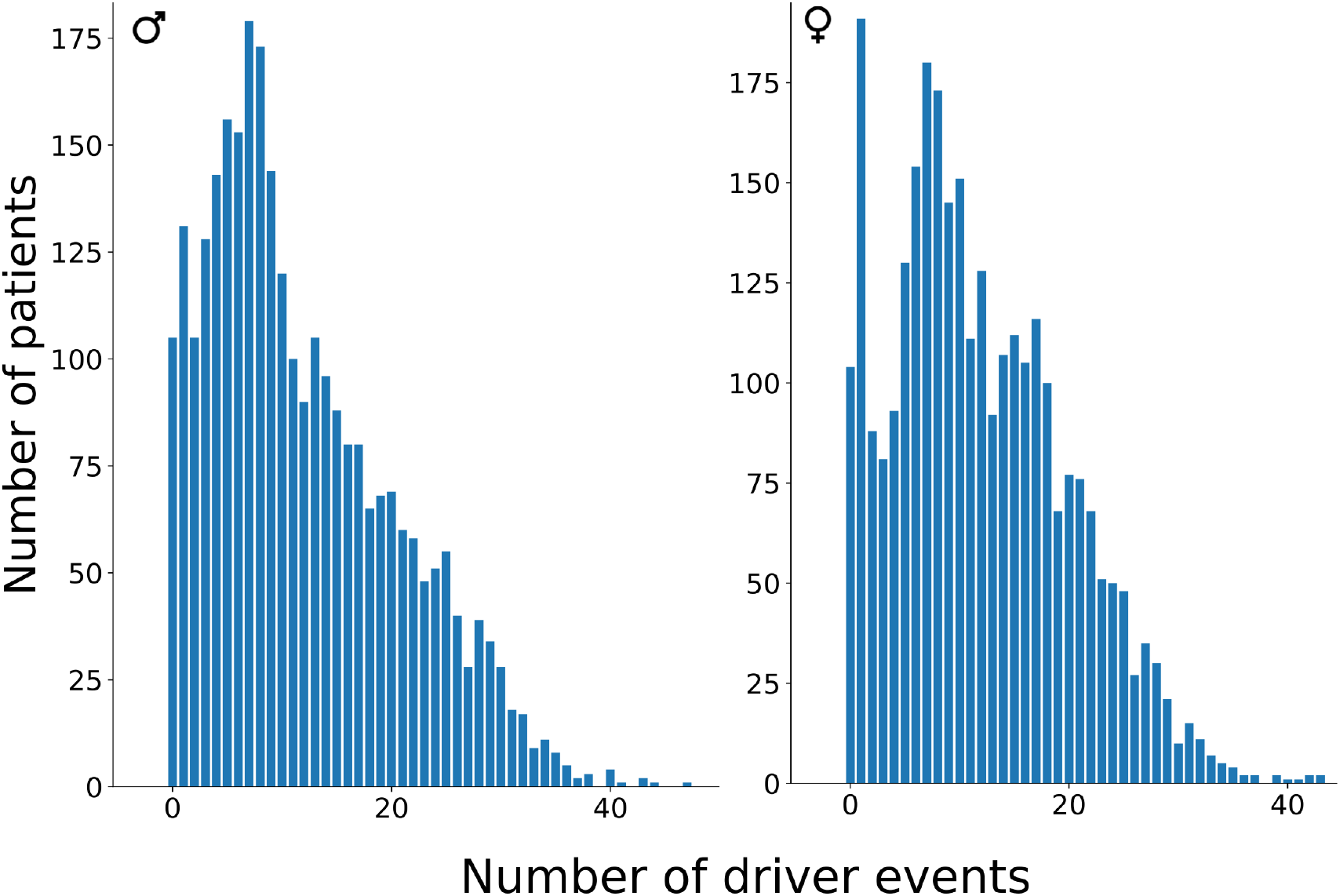
Patient distribution by total number of driver events per patient.

**Fig 7** shows the average number of driver events of different classes in different cancer cohorts. It can be immediately seen that there is a dramatic difference in the number and distribution of driver events per patient between cancer types. Some cancer types (ACC, CHOL, KICH, KIRP, MESO, PCPG, SARC, TGCT) are dominated by aneuploidy events, whereas in others aneuploidy events contribute half or less drivers. In ACC, KICH, KIRP and TGCT, the absolute majority of driver events are chromosome gains and losses. ACC, CHOL, KICH, KIRC, MESO and PCPG have almost no driver alterations in oncogenes. Additionally, BRCA, ESCA, GBM, HNSC, KIRP, SARC and TGCT have almost no SNAs in oncogenes, whereas THCA and THYM have no amplifications of oncogenes. Moreover, TGCT, THCA, THYM and UVM have almost no driver alterations in tumour suppressors, whereas KICH and PCPG are missing only SNAs in tumour suppressors. Tumour suppressors in DLBC and UCEC and oncogenes in DLBC, LGG, PAAD, THCA, THYM and UVM are predominantly affected by SNAs, which is not seen in other cancer types.

**Fig. 8** shows the actual distributions of driver event quantities amongst patients of all cancer types combined. It can be seen that these distributions have a sharp peak at one driver event per tumour, which is the dominant peak in females, and a smoother peak at 7 events, which is the dominant peak in males. THCA is the major contributor to the one-driver-event peak, but also PRAD and THYM. There are almost no patients with more than 40 driver events per tumour.

## Discussion

Our results show that half of all driver events are chromosome- and arm-level gains and losses, a third are amplifications of oncogenes and deletions of tumour suppressors and only one sixth are SNAs. Hopefully, this finding will help to shift the current perception that there are very few driver events per tumour and they are mostly SNAs to the greater appreciation of copy-number alterations and especially aneuploidy events. This may stimulate efforts in development of novel therapeutics targeting these types of driver alterations.

It is interesting to speculate why the number of driver events steadily increases in women until 50 y.o. and in men until 70 y.o. As 50 y.o. is the average age of menopause whereas males are fertile until much older ages, it is logical to suggest that driver accumulation is somehow promoted by high levels of sex hormones. If we concur with the theory that driver events accumulate in stem cells in large part due to errors in DNA replication and chromosome segregation [15], and combine it with the knowledge that sex hormones increase the rate of stem cell division [16–18], the link becomes clear. It is known that individuals with higher levels of sex hormones have higher rates of cancer [19].

In tumours with one driver event, this event is almost always an SNA in an oncogene, and with increasing number of total driver events the number of CNAs and aneuploidy events increases dramatically, whereas the number of SNAs in tumour suppressors increases only slightly. Most strikingly, the number of SNAs in oncogenes does not increase at all. Thus, it is tempting to suggest that a hyperactivating SNA in an oncogene is the seed which initiates cancer and enables later appearance of CNAs and aneuploidy events, the latter two driving the progression of cancer. However, it cannot be excluded that in tumours having more than one driver event the initiating event is CNA or aneuploidy, as these are cross-sectional and not longitudinal data.

Our results on the number and composition of driver events amongst various cancer types and individual patients pose several interesting questions. First, why there is such a great difference in the number and composition of driver events between cancer types? Why only one driver event is sufficient to initiate thyroid cancer but two dozens are observed in bladder carcinomas? Why some cancer types are dominated by aneuploidy drivers and others by SNA drivers? Why some cancers have no alterations in oncogenes whereas others have none in tumour suppressors? Are these differences explained by different tissue microenvironments to which these tumours have to adapt? Why than for some patients within the same cancer type one driver event is sufficient to develop a detectable tumour whereas the others are not diagnosed until dozens of events are accumulated?

One possible explanation is that in patients with low number of predicted driver events some actually important drivers have not been discovered due to technical reasons, from sequencing to bioinformatic analysis. Some of them might even be epigenetic [20], a class of drivers we have not analysed in this study. Another possible explanation is that in patients with high numbers of predicted drivers some of those drivers are actually false positives due to imperfections of driver prediction algorithms. Although both of these explanations are likely true to some degree, their contribution does not seem to be so large as to dramatically affect the results. The most probable explanation is the inherent stochasticity of carcinogenesis processes [21,22] combined with unequal strength of different driver events [23]. For example, patients having tumours with only one driver event were unlucky that this event happened (by chance) in one of the most crucial genes such as BRAF. On the other hand, other patients did not develop a detectable tumour until they have accumulated 30 events because those events involved only weak drivers. It would be interesting to develop a bioinformatic approach to determine the strength of driver genes, e.g. based on how often a given gene is found amongst a few sufficient ones.

Overall, our study brought some clarity to the distributions of driver events of various classes in various demographic and clinical groups of cancer patients and highlighted the importance of aneuploidy events and copy-number alterations. Many crucial questions, such as the reason behind the dramatic variability in the number and composition of driver events between cancer types and patients, remain to be answered.

## Methods

### Source files and initial filtering

TCGA PanCanAtlas data were used. Files “Analyte level annotations - merged_sample_quality_annotations.tsv”, “ABSOLUTE purity/ploidy file - TCGA_mastercalls.abs_tables_JSedit.fixed.txt”, “Aneuploidy scores and arm calls file - PANCAN_ArmCallsAndAneuploidyScore_092817.txt”, “Public mutation annotation file - mc3.v0.2.8.PUBLIC.maf.gz”, “gzipped ISAR-corrected GISTIC2.0 all_thresholded.by_genes file - ISAR_GISTIC.all_thresholded.by_genes.txt”, “RNA batch corrected matrix - EBPlusPlusAdjustPANCAN_IlluminaHiSeq_RNASeqV2.geneExp.tsv”, “miRNA batch corrected matrix - pancanMiRs_EBadjOnProtocolPlatformWithoutRepsWithUnCorrectMiRs_08_04_16.csv”, were downloaded from https://gdc.cancer.gov/about-data/publications/PanCan-CellOfOrigin.

Using TCGA barcodes (see https://docs.gdc.cancer.gov/Encyclopedia/pages/TCGA_Barcode/ and https://gdc.cancer.gov/resources-tcga-users/tcga-code-tables/sample-type-codes), all samples except primary tumours (barcoded 01, 03, 09) were removed from all files. Based on the information in the column “Do_not_use” in the file “Analyte level annotations - merged_sample_quality_annotations.tsv”, all samples with “True” value were removed from all files. All samples with “Cancer DNA fraction” <0.5 or unknown or with “Subclonal genome fraction” >0.5 or unknown in the file “TCGA_mastercalls.abs_tables_JSedit.fixed.txt” were removed from the file “PANCAN_ArmCallsAndAneuploidyScore_092817.txt”. Moreover, all samples without “PASS” value in the column “FILTER” were removed from the file “mc3.v0.2.8.PUBLIC.maf.gz” and zeros in the column “Entrez_Gene_Id” were replaced with actual Entrez gene IDs, determined from the corresponding ENSEMBL gene IDs in the column “Gene” and external database ftp://ftp.ncbi.nih.gov/gene/DATA/GENE_INFO/Mammalia/Homo_sapiens.gene_info.gz. Filtered files were saved as “Primary_whitelisted_arms.tsv”, “mc3.v0.2.8.PUBLIC_primary_whitelisted_Entrez.tsv”, “ISAR_GISTIC.all_thresholded.by_genes_primary_whitelisted.tsv”, “EBPlusPlusAdjustPANCAN_IlluminaHiSeq_RNASeqV2-v2.geneExp_primary_whitelisted.tsv”, “pancanMiRs_EBadjOnProtocolPlatformWithoutRepsWithUnCorrectMiRs_08_04_16_primary_whit elisted.tsv”.

### RNA filtering of CNAs

Using the file “EBPlusPlusAdjustPANCAN_IlluminaHiSeq_RNASeqV2-v2.geneExp_primary_whitelisted.tsv”, the median expression level for each gene across patients was determined. If the expression for a given gene in a given patient was below 0.05x median value, it was encoded as “-2”, if between 0.05x and 0.75x median value, it was encoded as “-1”, if between 1.25x and 1.75x median value, it was encoded as “1”, if above 1.75x median value, it was encoded as “2”, otherwise it was encoded as “0”. The file was saved as “EBPlusPlusAdjustPANCAN_IlluminaHiSeq_RNASeqV2-v2.geneExp_primary_whitelisted_median.tsv.” The same operations were performed with the file “pancanMiRs_EBadjOnProtocolPlatformWithoutRepsWithUnCorrectMiRs_08_04_16_primary_ whitelisted.tsv”, which was saved as “pancanMiRs_EBadjOnProtocolPlatformWithoutRepsWithUnCorrectMiRs_08_04_16_primary_ whitelisted_median.tsv” Next, the file “ISAR_GISTIC.all_thresholded.by_genes_primary_whitelisted.tsv” was processed according to the following rules: if the gene CNA status in a given patient was not zero and had the same sign as the gene expression status in the same patient (file “EBPlusPlusAdjustPANCAN_IlluminaHiSeq_RNASeqV2-v2.geneExp_primary_whitelisted_median.tsv” or “pancanMiRs_EBadjOnProtocolPlatformWithoutRepsWithUnCorrectMiRs_08_04_16_primary_ whitelisted_median.tsv” for miRNA genes), then the CNA status value was replaced with the gene expression status value, otherwise it was replaced by zero. If the corresponding expression status for a given gene was not found then its CNA status was not changed. The resulting file was saved as “ISAR_GISTIC.all_thresholded.by_genes_primary_whitelisted_RNAfiltered.tsv”

We named this algorithm GECNAV (Gene Expression-based CNA Validator) and created a Github repository: https://github.com/belikov-av/GECNAV. The package used to generate data in this article is available as **S5 Files**.

### Aneuploidy driver prediction

Using the file “Primary_whitelisted_arms.tsv”, the average alteration status of each chromosomal arm was calculated for each cancer type and saved as a matrix file “Arm_averages.tsv”. By drawing statuses randomly with replacement (bootstrapping) from any cell of “Primary_whitelisted_arms.tsv”, for each cancer type the number of statuses corresponding to the number of patients in that cancer type were generated and their average was calculated. The procedure was repeated 10000 times, the median for each cancer type was calculated and the results were saved as a matrix file “Bootstrapped_arm_averages.tsv”.

P-value for each arm alteration status was calculated for each cancer type. To do this, first the alteration status for a given cancer type and a given arm in “Arm_averages.tsv” was compared to the median bootstrapped arm alteration status for this cancer type in “Bootstrapped_arm_averages.tsv”. If the status in “Arm_averages.tsv” was higher than zero and the median in “Bootstrapped_arm_averages.tsv”, the number of statuses for this cancer type in “Bootstrapped_arm_averages.tsv” that are higher than the status in “Arm_averages.tsv” was counted and divided by 5000. If the status in “Arm_averages.tsv” was lower than zero and the median in “Bootstrapped_arm_averages.tsv”, the number of statuses for this cancer type in “Bootstrapped_arm_averages.tsv” that are lower than the status in “Arm_averages.tsv” was counted and divided by 5000, and marked with minus to indicate arm loss. Other values were ignored (cells left empty). The results were saved as a matrix file “Arm_Pvalues_cohorts.tsv”.

For each cancer type, Benjamini–Hochberg procedure with FDR=5% was applied to P-values in “Arm_Pvalues_cohorts.tsv” and passing P-values were encoded as “DAG” (Driver arm gain) or “DAL” (Driver arm loss) if marked with minus. The other cells were made empty and the results were saved as a matrix file “Arm_drivers_FDR5_cohorts.tsv”.

Alterations were classified according to the following rules: if the arm status in a given patient (file “Primary_whitelisted_arms.tsv”) was “-1” and the average alteration status of a given arm in the same cancer type (file “Arm_drivers_FDR5_cohorts.tsv”) was “DAL”, then the alteration in the patient was classified as “DAL”. If the arm status in a given patient was “1” and the average alteration status of a given arm in the same cancer type was “DAG”, then the alteration in the patient was classified as “DAG”. In all other cases an empty cell was written. The total number of DALs and DAGs was calculated, patients with zero drivers were removed, and the results were saved as a matrix file “Arm_drivers_FDR5.tsv”.

Using the file “Primary_whitelisted_arms.tsv”, the values for the whole chromosomes were calculated using the following rules: if both p- and q-arm statuses were “1” then the chromosome status was written as “1”; if both p- and q-arm statuses were “-1” then the chromosome status was written as “-1”; if at least one arm status was not known (empty cell) then the chromosome status was written as empty cell; in all other cases the chromosome status was written as “0”. For one-arm chromosomes (13, 14, 15, 21, 22), their status equals the status of the arm. The resulting file was saved as “Primary_whitelisted_chromosomes.tsv”.

The same procedures as described above for chromosomal arms were repeated for the whole chromosomes, with the resulting file “Chromosome_drivers_FDR5.tsv”. Chromosome drivers were considered to override arm drivers, so if a chromosome had “DCL” (Driver chromosome loss) or “DCG” (Driver chromosome gain), no alterations were counted on the arm level, to prevent triple counting of the same event.

We named this algorithm ANDRIF (ANeuploidy DRIver Finder) and created a Github repository: https://github.com/belikov-av/ANDRIF. The package used to generate data in this article is available as **S6 Files**.

### SNA driver prediction

Using the file “mc3.v0.2.8.PUBLIC_primary_whitelisted_Entrez.tsv” all SNAs were classified according to the column “Variant_Classification”. “Frame_Shift_Del”, “Frame_Shift_Ins”, “Nonsense_Mutation”, “Nonstop_Mutation” and “Translation_Start_Site” were considered potentially inactivating; “De_novo_Start_InFrame”, “In_Frame_Del”, “In_Frame_Ins” and “Missense_Mutation” were considered potentially hyperactivating; “De_novo_Start_OutOfFrame” and “Silent” were considered passengers; the rest were considered unclear. The classification results were saved as the file “SNA_classification_patients.tsv”, with columns “Tumor_Sample_Barcode”, “Hugo_Symbol”, “Entrez_Gene_Id”, “Gene”, “Number of hyperactivating SNAs”, “Number of inactivating SNAs”, “Number of SNAs with unclear role”, “Number of passenger SNAs”.

Using this file, the sum of all alterations in all patients was calculated for each gene. Genes containing only SNAs with unclear role (likely, noncoding genes) were removed, also from “SNA_classification_patients.tsv”. Next, the Nonsynonymous SNA Enrichment Index (NSEI) was calculated for each gene as

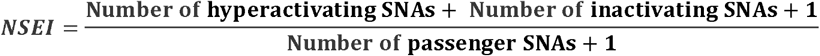

and the Hyperactivating to Inactivating SNA Ratio (HISR) was calculated for each gene as

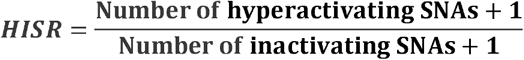

Genes for which the sum of hyperactivating, inactivating and passenger SNAs was less than 10 were removed to ensure sufficient precision of NSEI and HISR calculation, and the results were saved as “SNA_classification_genes_NSEI_HISR.tsv”.

Using the file “SNA_classification_patients.tsv”, the gene-patient matrix “SNA_matrix.tsv” was constructed, encoding the “Number of hyperactivating SNAs”, “Number of inactivating SNAs”, “Number of SNAs with unclear role” and “Number of passenger SNAs” as one number separated by dots (e.g. “2.0.1.1”). If data for a given gene were absent in a given patient, it was encoded as “0.0.0.0”. By drawing statuses randomly with replacement (bootstrapping) from any cell of “SNA_matrix.tsv” 10000 times for each patient, the matrix file “SNA_matrix_bootstrapped.tsv” was created. The sums of statuses in “SNA_matrix_bootstrapped.tsv” were calculated for each iteration separately, and then the corresponding NSEI and HISR indices were calculated and the results were saved as “SNA_bootstrapped_NSEI_HISR.tsv”. Null hypothesis P-values were calculated for each iteration as the number of NSEI values higher than a given iteration’s NSEI value and divided by 10000. The histogram of bootstrapped p-values was plotted to check for the uniformity of the null hypothesis p-value distribution.

P-value for each gene was calculated as the number of NSEI values in “SNA_bootstrapped_NSEI_HISR.tsv” more extreme than its NSEI value in “SNA_classification_genes_NSEI_HISR.tsv” and divided by 10000. The results were saved as “SNA_classification_genes_NSEI_HISR_Pvalues.tsv”. Benjamini–Hochberg procedure with FDR(Q)=5% was applied to P-values in “SNA_classification_genes_NSEI_HISR_Pvalues.tsv”, and genes that pass were saved as “SNA_driver_gene_list_FDR5.tsv”.

We named this algorithm SNADRIF (SNA DRIver Finder) and created a Github repository: https://github.com/belikov-av/SNADRIF. The package used to generate data in this article is available as **S7 Files**.

### Driver prediction algorithms sources and benchmarking

Lists of driver genes and mutations predicted by various algorithms (**Table 2**) applied to PanCanAtlas data were downloaded from https://gdc.cancer.gov/about-data/publications/pancan-driver (2020plus, CompositeDriver, DriverNet, HotMAPS, OncodriveFML), https://karchinlab.github.io/CHASMplus (CHASMplus), as well as received by personal communication from Francisco Martínez-Jiménez, Institute for Research in Biomedicine, Barcelona, (dNdScv, OncodriveCLUSTL, OncodriveFML). All genes and mutations with q-value > 0.05 were removed. Additionally, a consensus driver gene list from 26 algorithms applied to PanCanAtlas data [8] was downloaded from https://www.cell.com/cell/fulltext/S0092-8674(18)30237-X and a COSMIC Cancer Gene Census (CGC) Tier 1 gene list [14] was downloaded from https://cancer.sanger.ac.uk/cosmic/census?tier=1. Only genes affected by somatic SNAs and CNAs present in the TCGA cancer types were used for further analyses from the CGC list. Cancer type names in the CGC list were manually converted to the closest possible TCGA cancer type abbreviation. Entrez Gene IDs were identified for each gene using HUGO Symbol and external database ftp://ftp.ncbi.nih.gov/gene/DATA/GENE_INFO/Mammalia/Homo_sapiens.gene_info.gz.

**Table 2.**
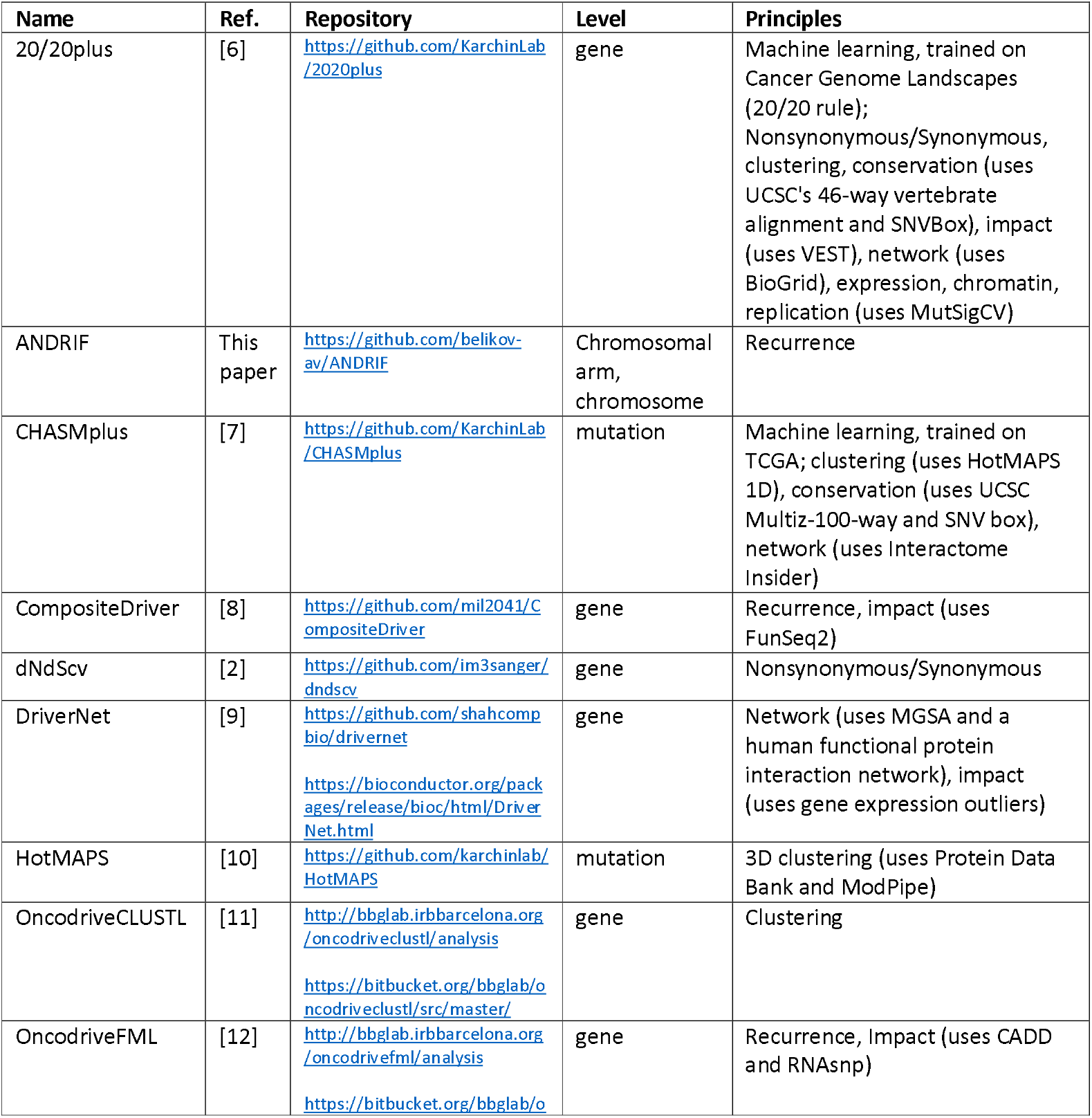
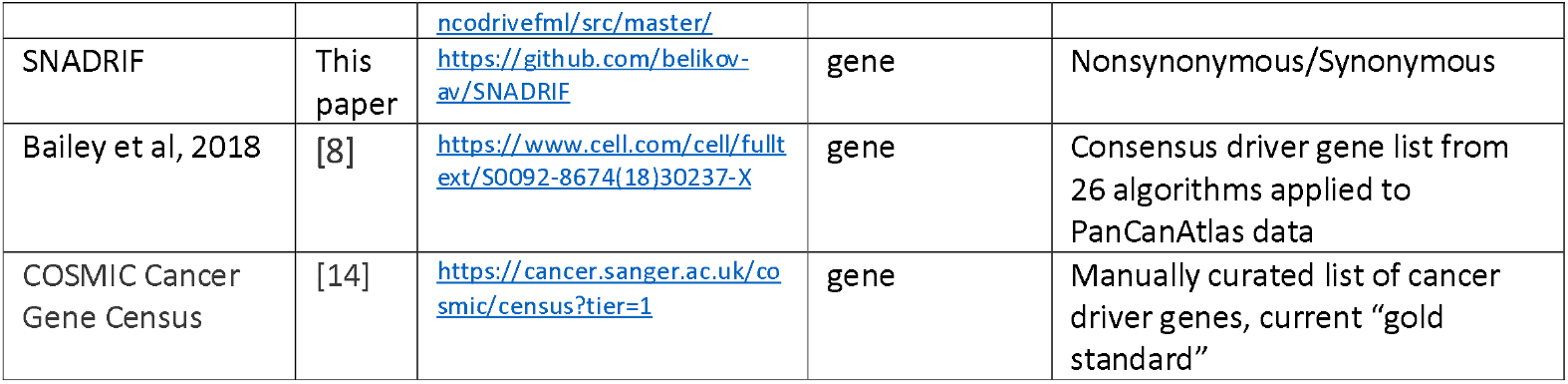
Driver prediction algorithms.

The sensitivity of algorithms was assessed as the percentage of genes in a positive control list that were predicted as drivers by an algorithm, because Sensitivity=True positives/(True positives + False negatives). Three positive control lists were used – CGC Tier 1 genes affected by somatic SNAs or CNAs in TCGA cancer types, a list of genes identified by at least two of all our sources (including CGC and Bailey), and a list of genes identified by at least three of all our sources (including CGC and Bailey). Sensitivity was assessed separately for algorithms applied to individual cancer types as the percentage of gene-cohort pairs in a positive control list that were matched by gene-cohort pairs predicted by an algorithm.

### Conversion of population-level data to patient-level data

For lists of driver genes, all entries from the file “mc3.v0.2.8.PUBLIC_primary_whitelisted_Entrez.tsv” were removed except those that satisfied the following conditions simultaneously: “Entrez Gene ID” matches the one in the driver list; cancer type (identified by matching “Tumor_Sample_Barcode” with “bcr_patient_barcode” and “acronym” in “clinical_PANCAN_patient_with_followup.tsv”) matches “cohort” in the driver list or the driver list is for pancancer analysis; “Variant_Classification” column contains one of the following values: “De_novo_Start_InFrame”, “Frame_Shift_Del”, “Frame_Shift_Ins”, “In_Frame_Del”, “In_Frame_Ins”, “Missense_Mutation”, “Nonsense_Mutation”, “Nonstop_Mutation”, “Translation_Start_Site”.

For lists of driver mutations, the procedures were the same, except that Ensembl Transcript ID and nucleotide/amino acid substitution were used for matching instead of Entrez Gene ID. These data (only columns “TCGA Barcode”, “HUGO Symbol”, “Entrez Gene ID”) were saved as “AlgorithmName_output_SNA.tsv”.

Additionally, all entries from the file “ISAR_GISTIC.all_thresholded.by_genes_primary_whitelisted.tsv” were removed except those that satisfied the following conditions simultaneously: “Locus ID” matches “Entrez Gene ID” in the driver list; cancer type (identified by matching Tumor Sample Barcode with “bcr_patient_barcode” and “acronym” in “clinical_PANCAN_patient_with_followup.tsv”) matches “cohort” in the driver list or the driver list is for pancancer analysis; CNA values are “2”, “1”, “-1” or “-2”. These data were converted from the matrix to a list format (with columns “TCGA Barcode”, “HUGO Symbol”, “Entrez Gene ID”) and saved as “AlgorithmName_output_CNA.tsv”.

Finally, the files “AlgorithmName_output_SNA.tsv” and “AlgorithmName_output_CNA.tsv” were combined, duplicate TCGA Barcode-Entrez Gene ID pairs were removed, and the results saved as “AlgorithmName_output.tsv”.

### Driver event classification and analysis

The file “Clinical with Follow-up - clinical_PANCAN_patient_with_followup.tsv” was downloaded from https://gdc.cancer.gov/node/905/. All patients with “icd_o_3_histology” different from XXXX/3 (primary malignant neoplasm) were removed, as well as all patients not simultaneously present in the following three files: “mc3.v0.2.8.PUBLIC_primary_whitelisted_Entrez.tsv”, “ISAR_GISTIC.all_thresholded.by_genes_primary_whitelisted.tsv” and “Primary_whitelisted_arms.tsv”. The resulting file was saved as “clinical_PANCAN_patient_with_followup_primary_whitelisted.tsv”.

Several chosen “AlgorithmName_output.tsv” files were combined and all TCGA Barcode-Entrez Gene ID pairs not present in at least two output files were removed. Entries with TCGA Barcodes not present in “clinical_PANCAN_patient_with_followup_primary_whitelisted.tsv” were removed as well. Matching “Number of hyperactivating SNAs” and “Number of inactivating SNAs” for each TCGA Barcode-Entrez Gene ID pair were taken from the “SNA_classification_patients.tsv” file, in case of no match zeros were written. Matching HISR value was taken from “SNA_classification_genes_NSEI_HISR.tsv” for each Entrez Gene ID, in case of no match empty cell was left. Matching CNA status was taken from “ISAR_GISTIC.all_thresholded.by_genes_primary_whitelisted_RNAfiltered.tsv” for each TCGA Barcode-Entrez Gene ID pair, in case of no match zero was written.

Each TCGA Barcode-Entrez Gene ID pair was classified according to the **Table 3:**

**Table 3.**
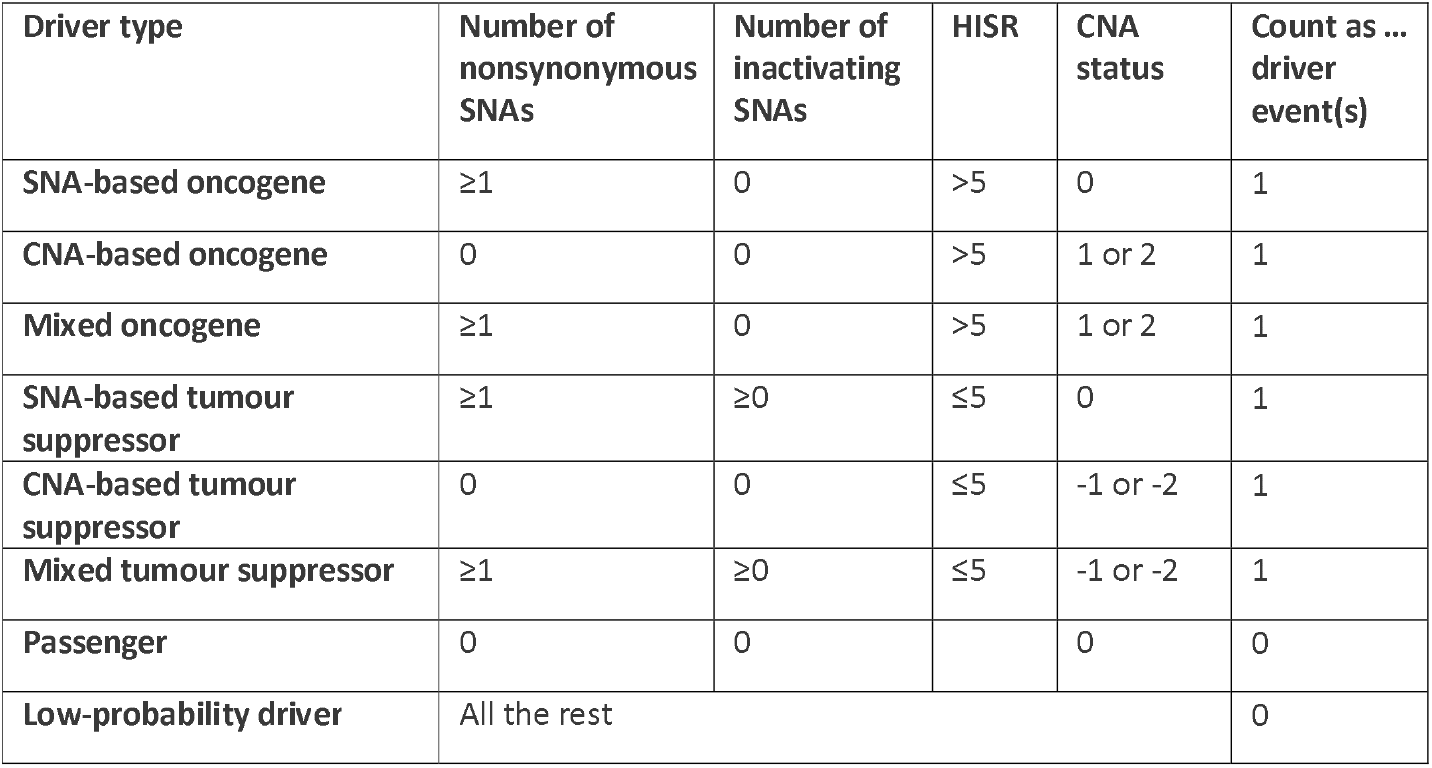
Driver event classification rules.

| Driver type | Number of nonsynonymous SNAs | Number of inactivating SNAs | HISR | CNA status | Count as ... driver event(s) |
| --- | --- | --- | --- | --- | --- |
| SNA-based oncogene | ≥1 | 0 | >5 | 0 | 1 |
| CNA-based oncogene | 0 | 0 | >5 | 1 or 2 | 1 |
| Mixed oncogene | ≥1 | 0 | >5 | 1 or 2 | 1 |
| SNA-based tumour suppressor | ≥1 | ≥0 | ≤5 | 0 | 1 |
| CNA-based tumour suppressor | 0 | 0 | ≤5 | -1 or -2 | 1 |
| Mixed tumour suppressor | ≥1 | ≥0 | ≤5 | -1 or -2 | 1 |
| Passenger | 0 | 0 |  | 0 | 0 |
| Low-probability driver | All the rest |  |  |  | 0 |

Results of this classification were saved as “AnalysisName_genes_level2.tsv”.

Using this file, the number of driver events of each type was counted for each patient. Information on the number of driver chromosome and arm losses and gains for each patient was taken from the files “Chromosome_drivers_FDR5.tsv” and “Arm_drivers_FDR5.tsv”. All patients not present in the files “AnalysisName_genes_level2.tsv”, “Chromosome_drivers_FDR5.tsv” and “Arm_drivers_FDR5.tsv”, but present in the file “clinical_PANCAN_patient_with_followup_primary_whitelisted.tsv”, were added with zero values for the numbers of driver events. Information on the cancer type (“acronym”), gender (“gender”), age (“age_at_initial_pathologic_diagnosis”) and tumor stage (“pathologic_stage”, if no data then “clinical_stage”, if no data then “pathologic_T”, if no data then “clinical_T”) was taken from the file “clinical_PANCAN_patient_with_followup_primary_whitelisted.tsv”. The results were saved as “AnalysisName_patients.tsv”.

Using the file “AnalysisName_patients.tsv”, the number of patients with each integer total number of driver events from 0 to 100 was counted for each cancer type, also for males and females separately, and histograms were plotted for each cancer type-gender combination. Using the same file “AnalysisName_patients.tsv”, the average number of various types of driver events was calculated for each cancer type, tumour stage, age group, as well as for patients with each integer total number of driver events from 1 to 100. Analyses were performed for total population and for males and females separately, and cumulative histograms were plotted for each file.

We named this algorithm PALDRIC (PAtient-Level DRIver Classifier) and created a Github repository: https://github.com/belikov-av/PALDRIC. The package used to generate data in this article is available as **S8 Files**.

## Acknowledgements

AVB acknowledges MIPT 5-100 program support for early career researchers.

---

**S1 Files. Benchmarking of driver prediction algorithms**.

**S2 Files. Output from PALDRIC algorithm applied to pancancer dataset**.

**S3 Files. Output from PALDRIC algorithm applied to individual cohorts**.

**S4 Files. Statistical significance testing for Figures 3, 4, 5**.

**S5 Files. GECNAV package**.

**S6 Files. ANDRIF package**.

**S7 Files. SNADRIF package**.

**S8 Files. PALDRIC package**.

## References

1. Pon JR, Marra MA. Driver and Passenger Mutations in Cancer. Annual Review of Pathology: Mechanisms of Disease. 2015;10: 25–50. doi:10.1146/annurev-pathol-012414-040312

2. Martincorena I, Raine KM, Gerstung M, Dawson KJ, Haase K, Van Loo P, et al. Universal Patterns of Selection in Cancer and Somatic Tissues. Cell. 2017;171: 1029-1041.e21. doi:10.1016/j.cell.2017.09.042

3. Zack TI, Schumacher SE, Carter SL, Cherniack AD, Saksena G, Tabak B, et al. Pan-cancer patterns of somatic copy number alteration. Nature Genetics. 2013;45: 1134–1140. doi:10.1038/ng.2760

4. Ben-David U, Amon A. Context is everything: aneuploidy in cancer. Nature Reviews Genetics. 2020;21: 44–62. doi:10.1038/s41576-019-0171-x

5. Dawson MA, Kouzarides T. Cancer Epigenetics: From Mechanism to Therapy. Cell. 2012;150: 12–27. doi:10.1016/j.cell.2012.06.013

6. Tokheim CJ, Papadopoulos N, Kinzler KW, Vogelstein B, Karchin R. Evaluating the evaluation of cancer driver genes. Proceedings of the National Academy of Sciences. 2016;113: 14330LP–14335. doi:10.1073/pnas.1616440113

7. Tokheim C, Karchin R. CHASMplus Reveals the Scope of Somatic Missense Mutations Driving Human Cancers. Cell Systems. 2019;9: 9-23.e8. doi:10.1016/j.cels.2019.05.005

8. Bailey MH, Tokheim C, Porta-Pardo E, Sengupta S, Bertrand D, Weerasinghe A, et al. Comprehensive Characterization of Cancer Driver Genes and Mutations. Cell. 2018;173: 371-385.e18. doi:10.1016/j.cell.2018.02.060

9. Bashashati A, Haffari G, Ding J, Ha G, Lui K, Rosner J, et al. DriverNet: uncovering the impact of somatic driver mutations on transcriptional networks in cancer. Genome Biology. 2012;13: R124. doi:10.1186/gb-2012-13-12-r124

10. Tokheim C, Bhattacharya R, Niknafs N, Gygax DM, Kim R, Ryan M, et al. Exome-Scale Discovery of Hotspot Mutation Regions in Human Cancer Using 3D Protein Structure. Cancer Research. 2016;76: 3719LP–3731. doi:10.1158/0008-5472.CAN-15-3190

11. Arnedo-Pac C, Mularoni L, Muiños F, Gonzalez-Perez A, Lopez-Bigas N. OncodriveCLUSTL: a sequence-based clustering method to identify cancer drivers. Bioinformatics. 2019;35: 4788–4790. doi:10.1093/bioinformatics/btz501

12. Mularoni L, Sabarinathan R, Deu-Pons J, Gonzalez-Perez A, López-Bigas N. OncodriveFML: a general framework to identify coding and non-coding regions with cancer driver mutations. Genome Biology. 2016;17: 128. doi:10.1186/s13059-016-0994-0

13. Vogelstein B, Papadopoulos N, Velculescu VE, Zhou S, Diaz LA, Kinzler KW. Cancer Genome Landscapes. Science. 2013;339: 1546LP–1558. doi:10.1126/science.1235122

14. Sondka Z, Bamford S, Cole CG, Ward SA, Dunham I, Forbes SA. The COSMIC Cancer Gene Census: describing genetic dysfunction across all human cancers. Nature Reviews Cancer. 2018;18: 696–705. doi:10.1038/s41568-018-0060-1

15. Tomasetti C, Li L, Vogelstein B. Stem cell divisions, somatic mutations, cancer etiology, and cancer prevention. Science. 2017;355: 1330LP–1334. doi:10.1126/science.aaf9011

16. Nakada D, Oguro H, Levi BP, Ryan N, Kitano A, Saitoh Y, et al. Oestrogen increases haematopoietic stem-cell self-renewal in females and during pregnancy. Nature. 2014;505: 555–558. doi:10.1038/nature12932

17. Asselin-Labat M-L, Vaillant F, Sheridan JM, Pal B, Wu D, Simpson ER, et al. Control of mammary stem cell function by steroid hormone signalling. Nature. 2010;465: 798–802. doi:10.1038/nature09027

18. Joshi PA, Jackson HW, Beristain AG, Di Grappa MA, Mote PA, Clarke CL, et al. Progesterone induces adult mammary stem cell expansion. Nature. 2010;465: 803–807. doi:10.1038/nature09091

19. Folkerd EJ, Dowsett M. Influence of Sex Hormones on Cancer Progression. Journal of Clinical Oncology. 2010;28: 4038–4044. doi:10.1200/JCO.2009.27.4290

20. Lyu J, Li JJ, Su J, Peng F, Chen YE, Ge X, et al. DORGE: Discovery of Oncogenes and tumoR suppressor genes using Genetic and Epigenetic features. Science Advances. 2020;6: eaba6784. doi:10.1126/sciadv.aba6784

21. Belikov AV. The number of key carcinogenic events can be predicted from cancer incidence. Scientific Reports. 2017;7. doi:10.1038/s41598-017-12448-7

22. Belikov AV, Vyatkin A, Leonov SV. The Erlang distribution approximates the age distribution of incidence of childhood and young adulthood cancers. Coates P, editor. PeerJ. 2021;9: e11976. doi:10.7717/peerj.11976

23. Zhang M, Jang H, Nussinov R. PI3K Driver Mutations: A Biophysical Membrane-Centric Perspective. Cancer Research. 2021;81: 237LP–247. doi:10.1158/0008-5472.CAN-20-0911

