## Supplementary figures and images for "Comprehensive patient-level classification and quantification of driver events in TCGA PanCanAtlas cohorts"

### 2021_8_22_13_29_ACC_FEMALE.pdf

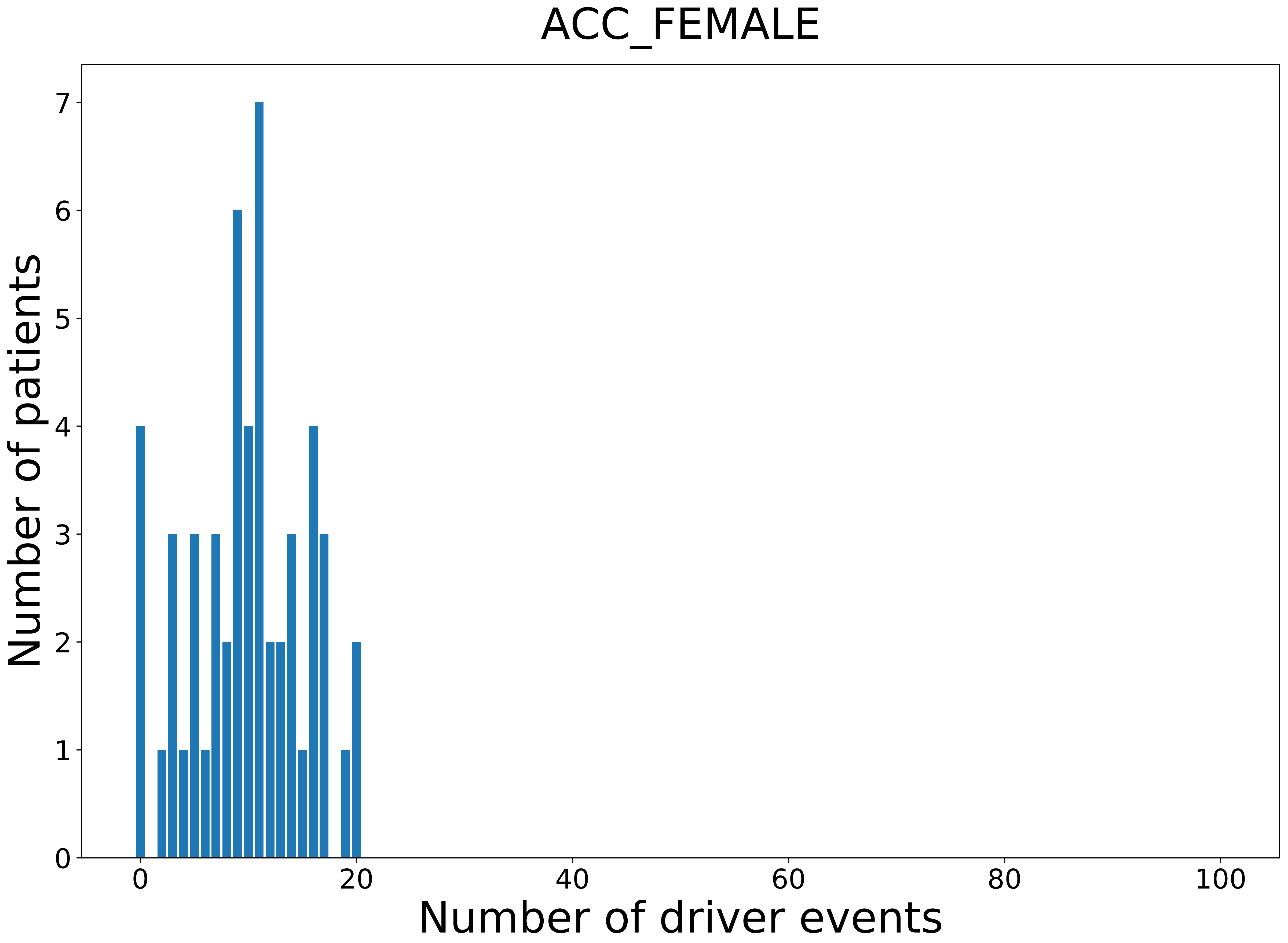

### 2021_8_22_13_29_ACC_MALE.pdf

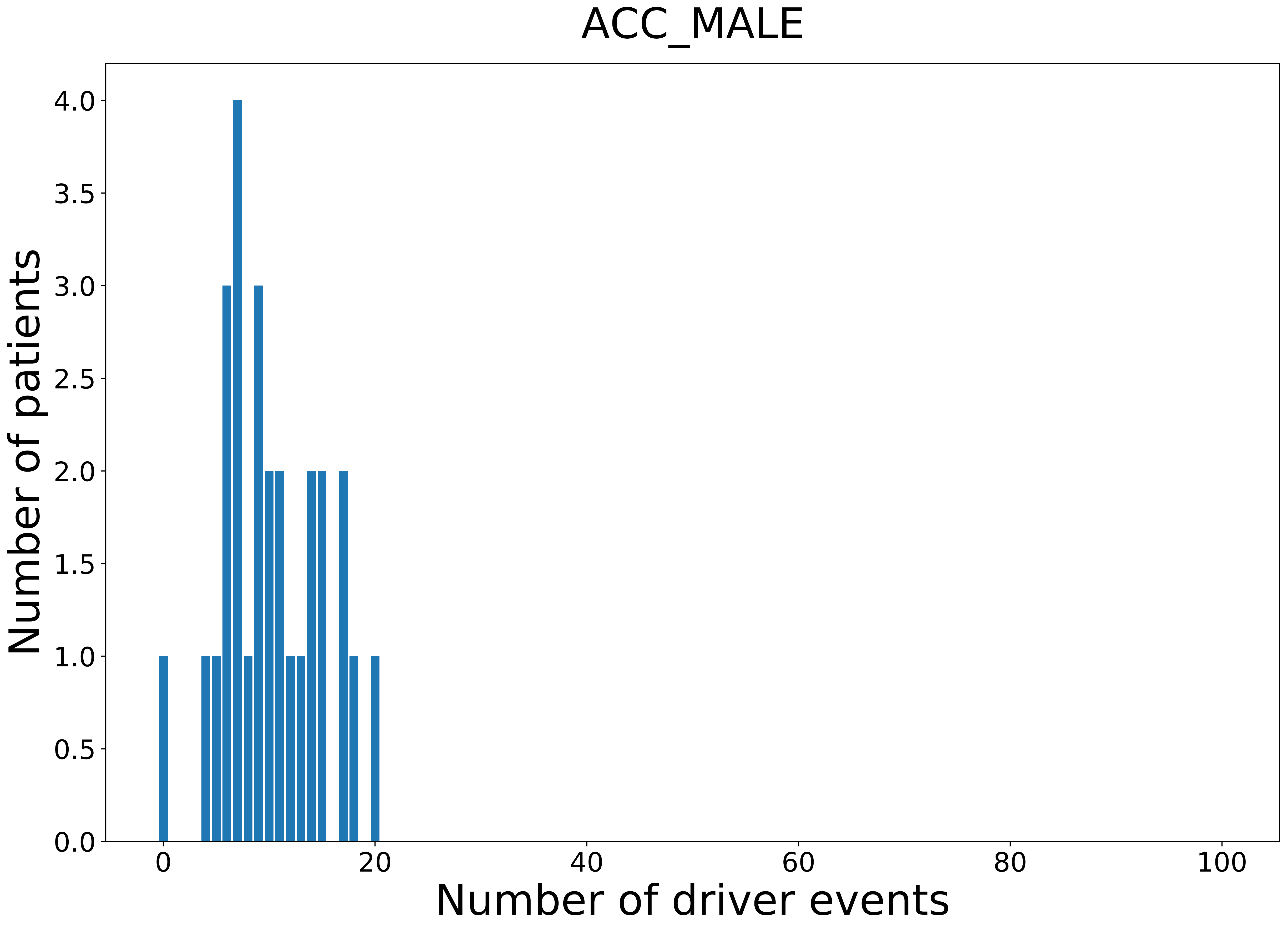

### 2021_8_22_13_29_BLCA.pdf

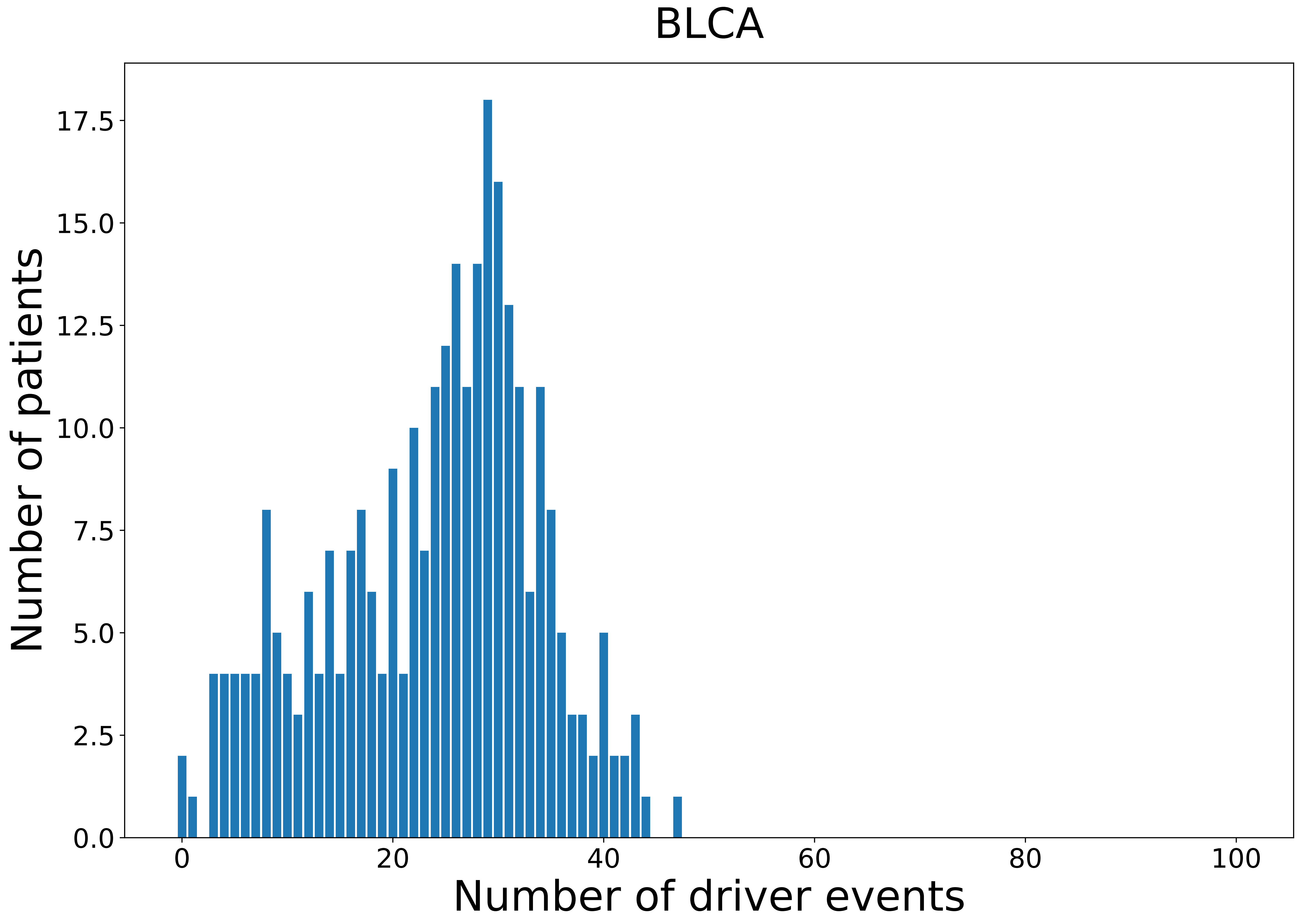

### 2021_8_22_13_29_BLCA_FEMALE.pdf

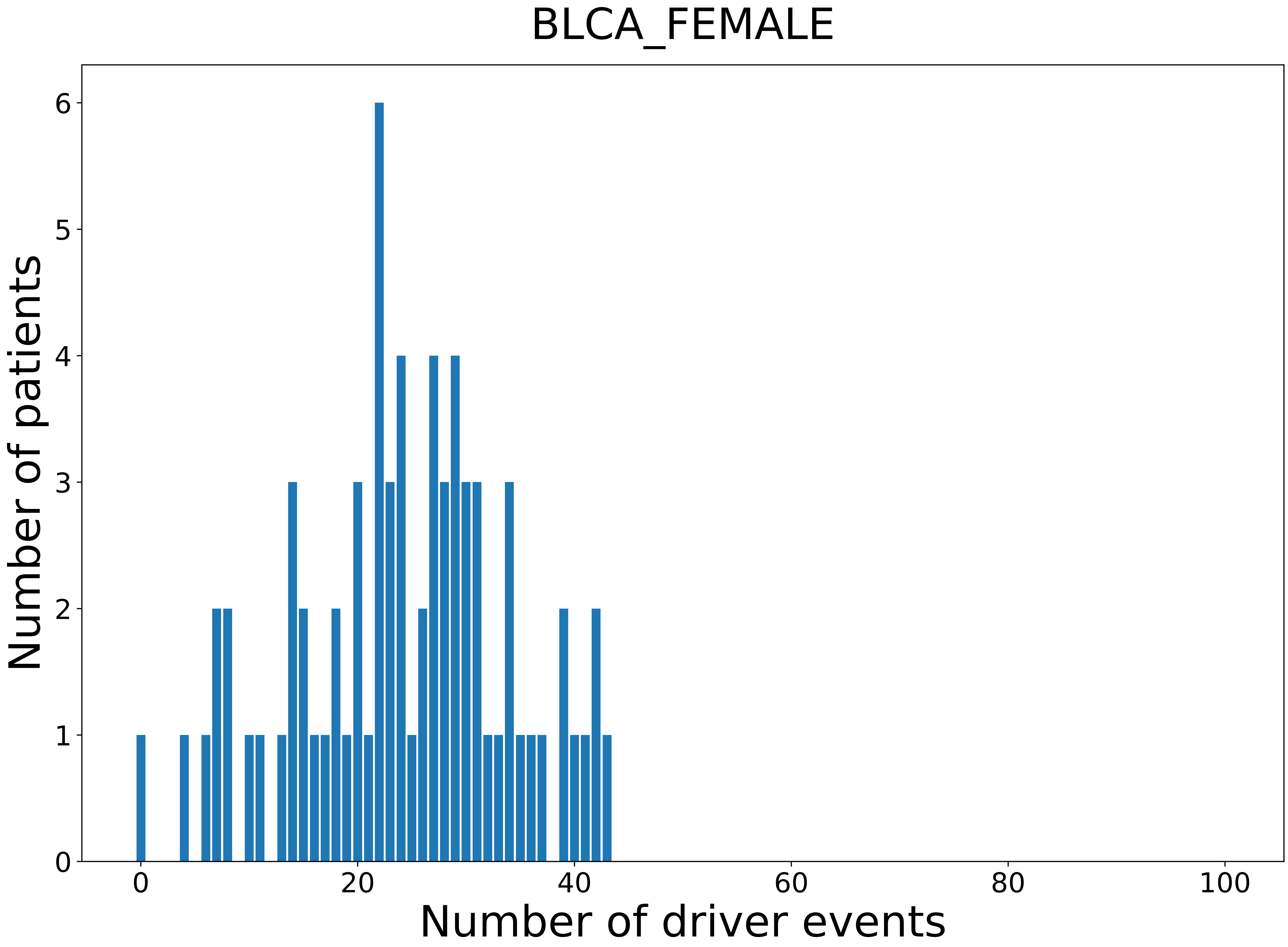

### 2021_8_22_13_29_BLCA_MALE.pdf

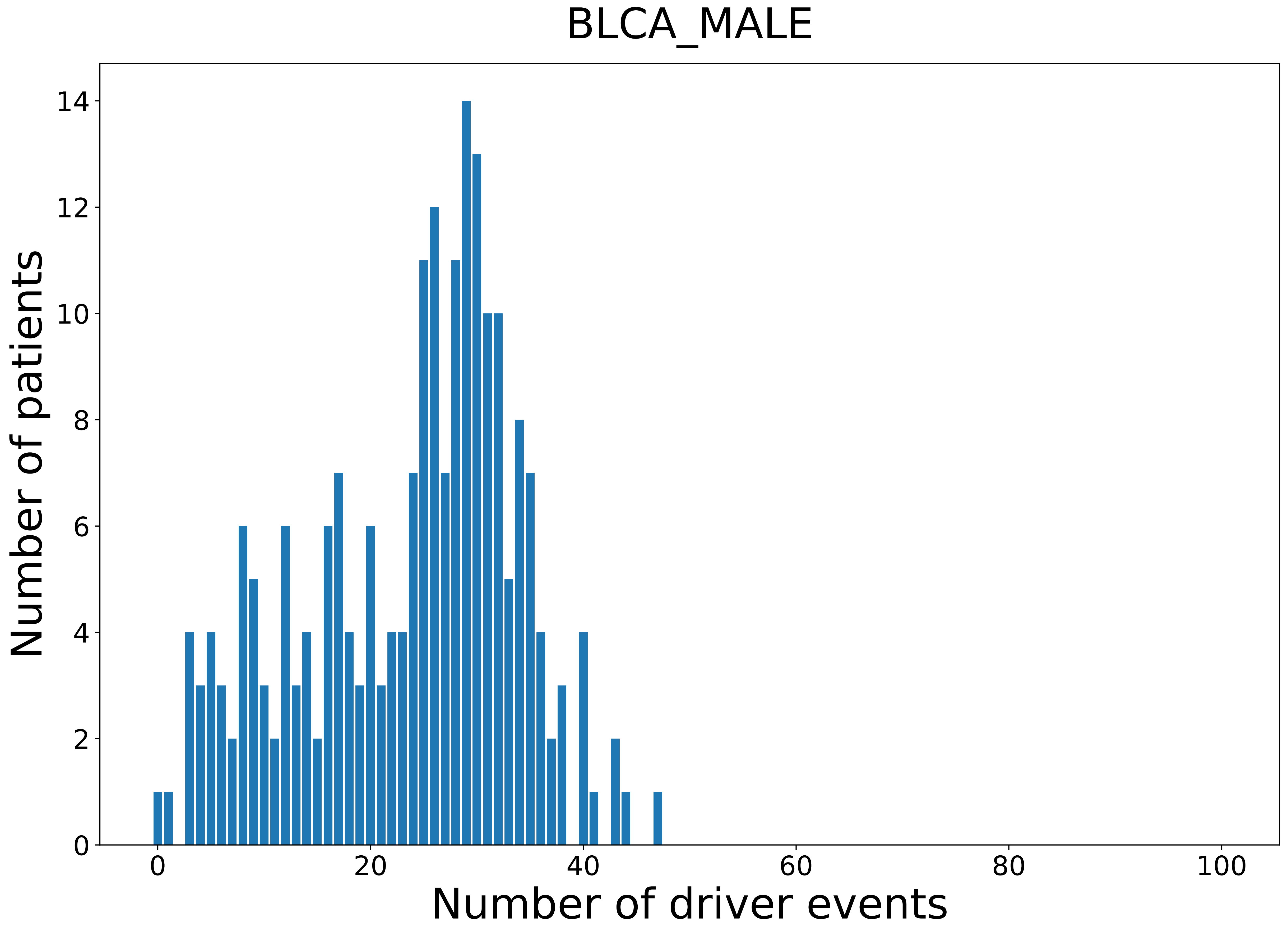

### 2021_8_22_13_29_BRCA.pdf

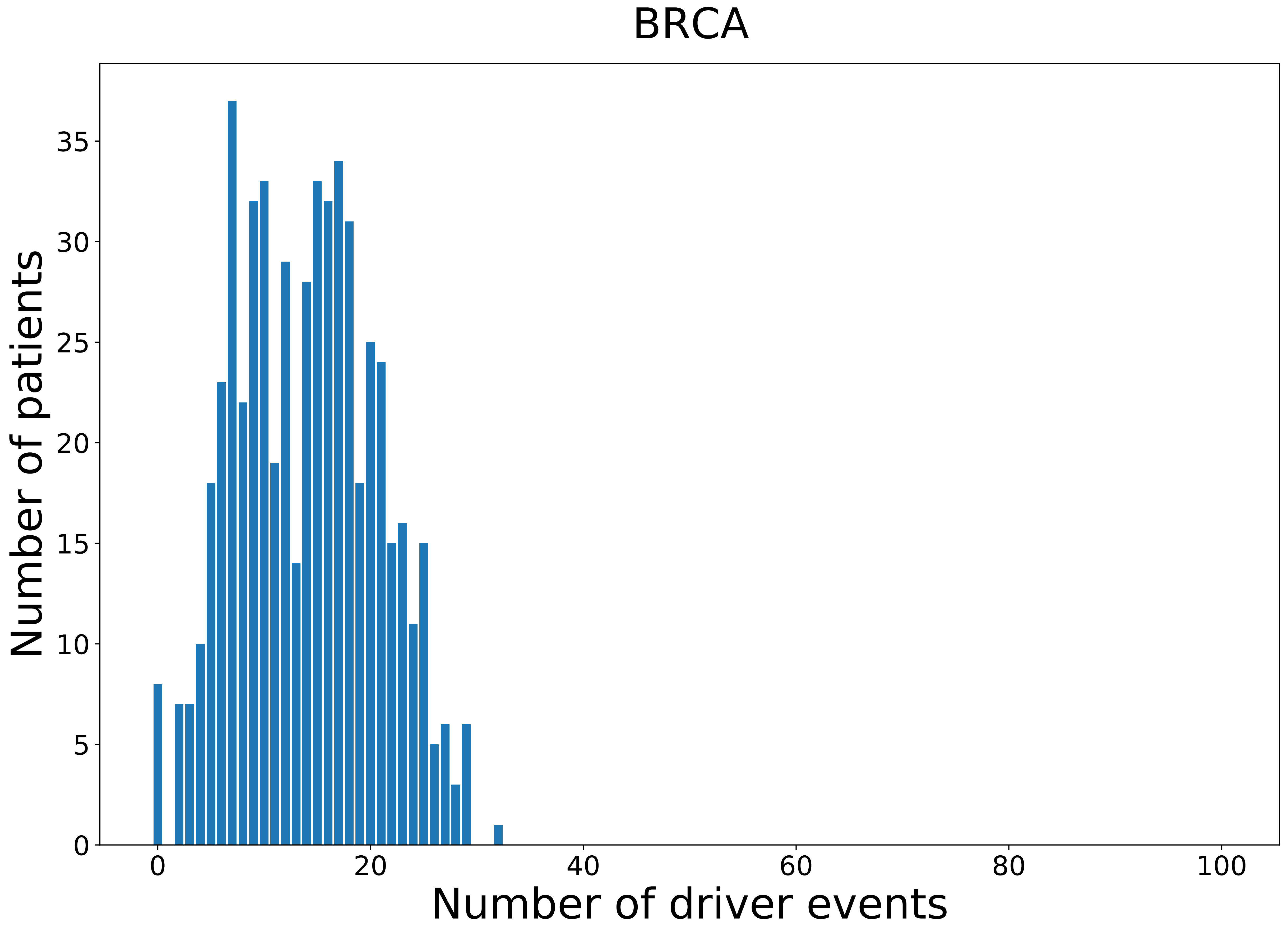

### 2021_8_22_13_29_BRCA_FEMALE.pdf

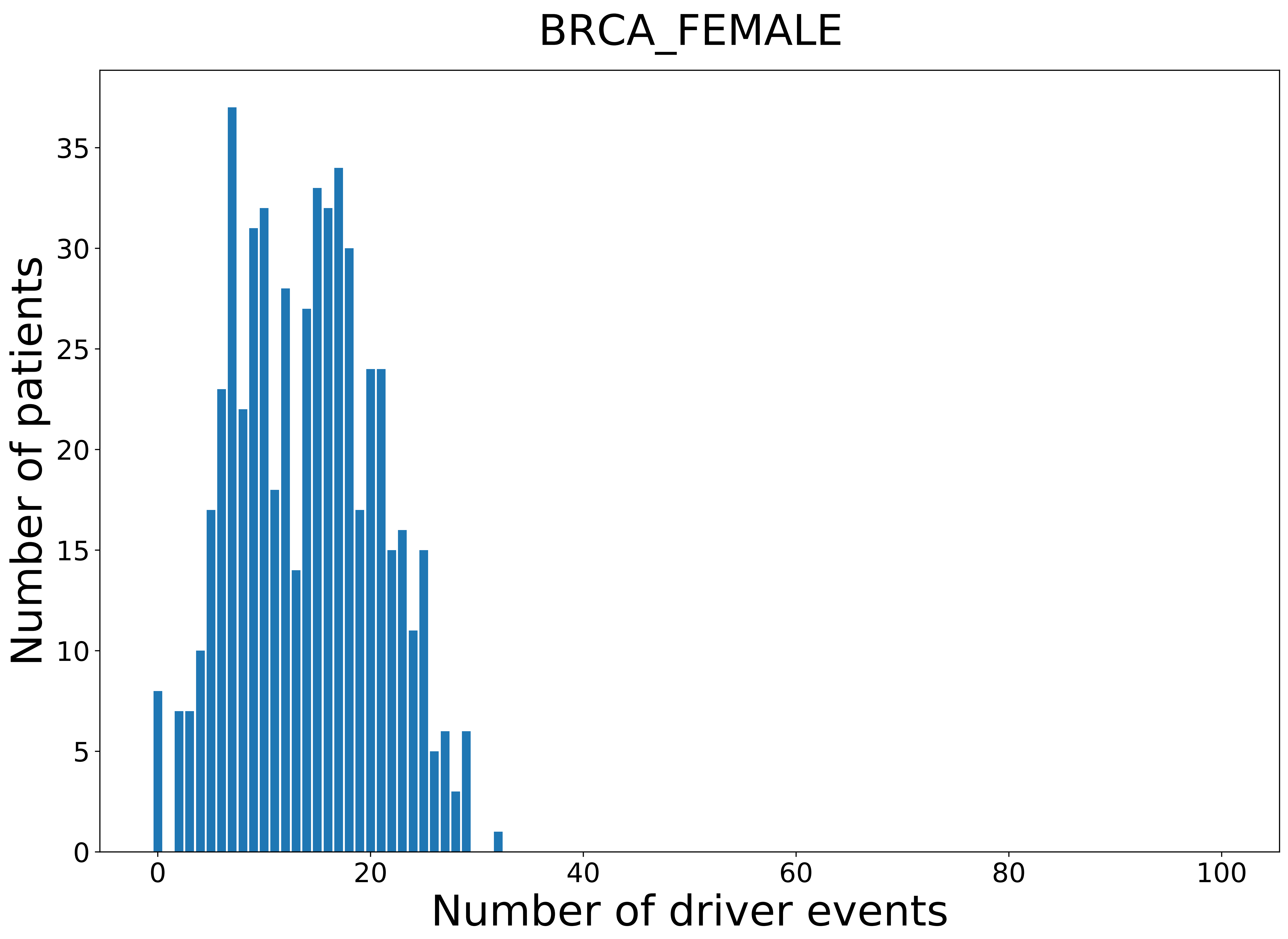

### 2021_8_22_13_29_CESC.pdf

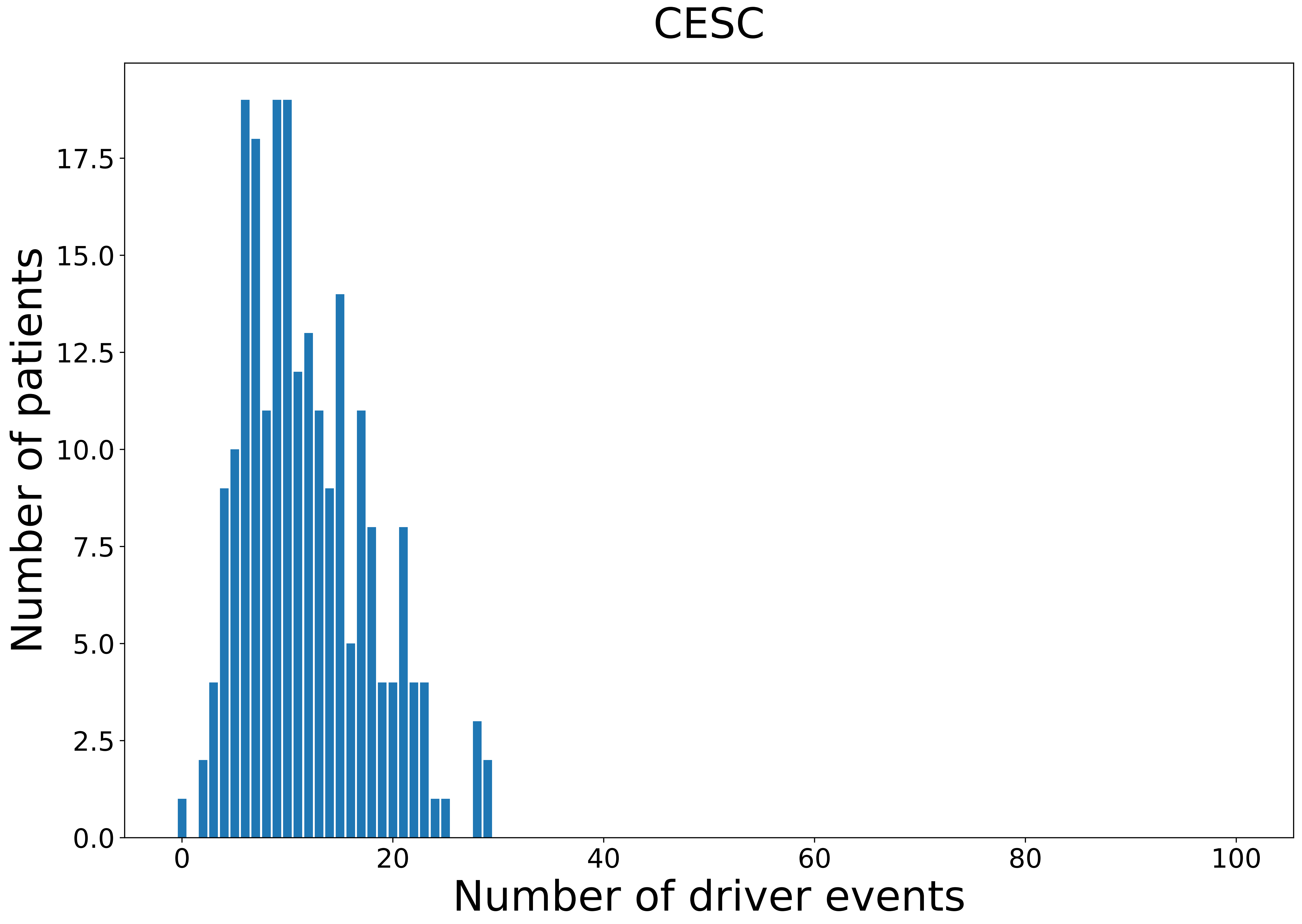

### 2021_8_22_13_29_CESC_FEMALE.pdf

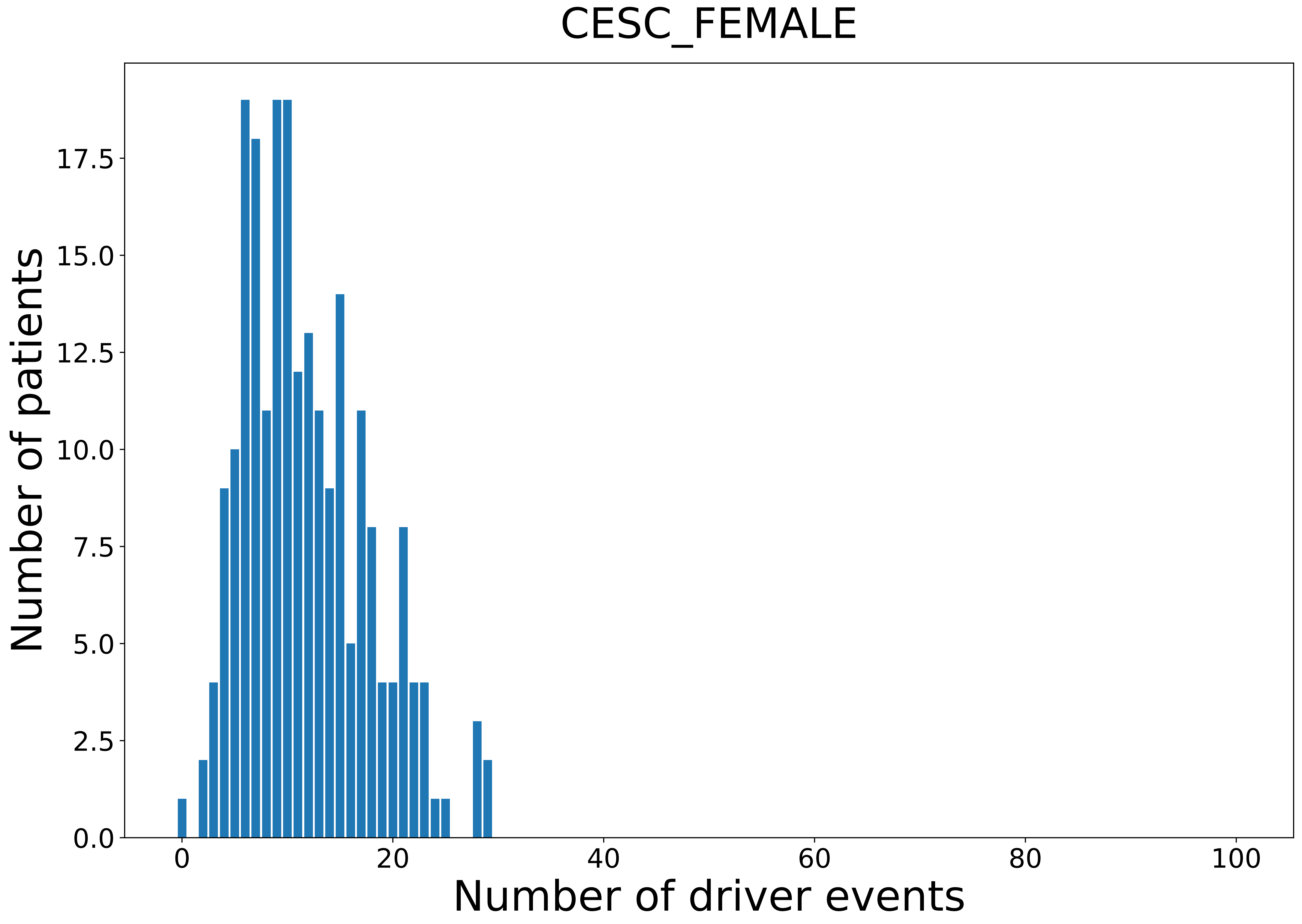

### 2021_8_22_13_29_CHOL.pdf

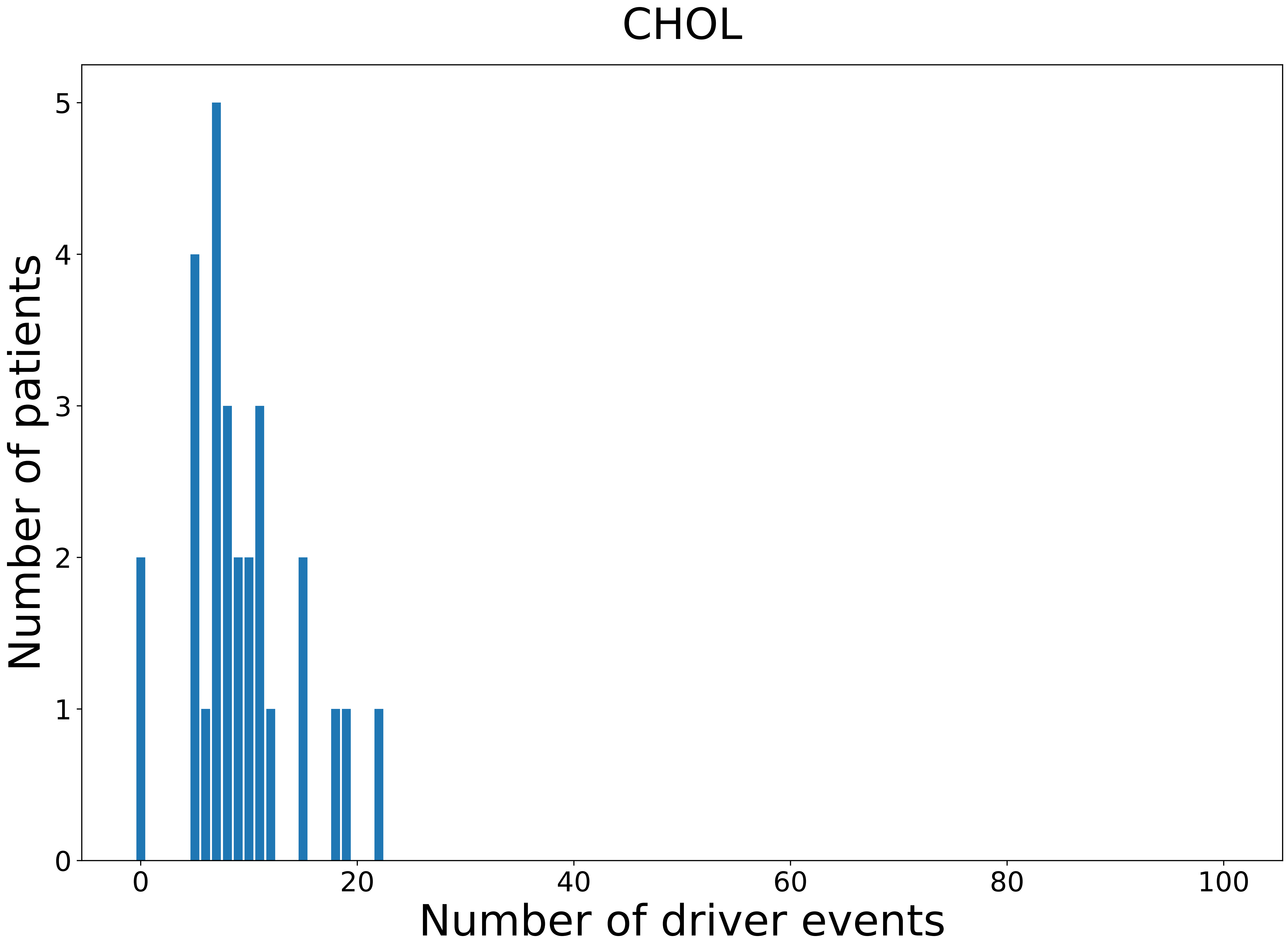

### 2021_8_22_13_29_CHOL_FEMALE.pdf

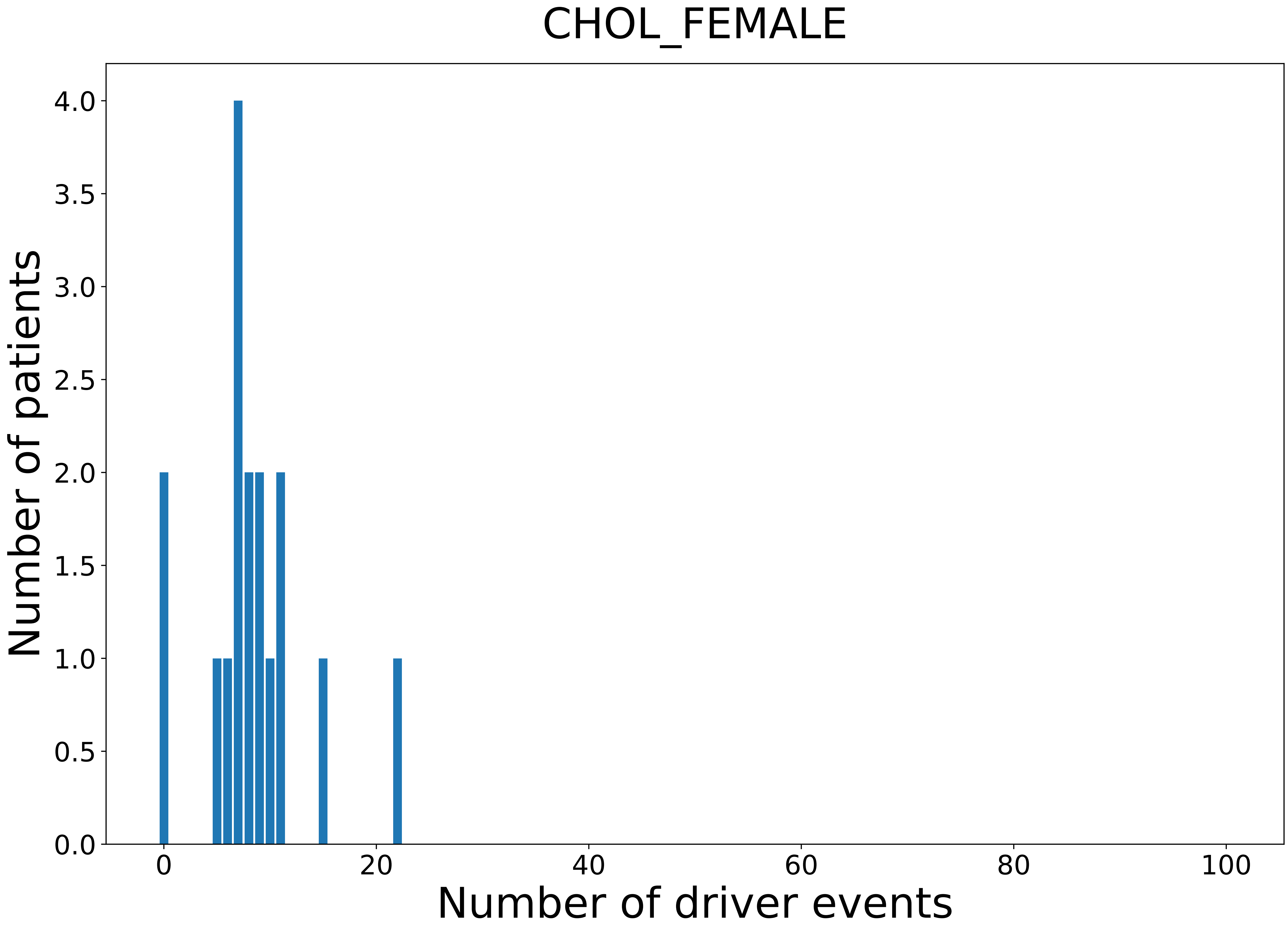

### 2021_8_22_13_29_CHOL_MALE.pdf

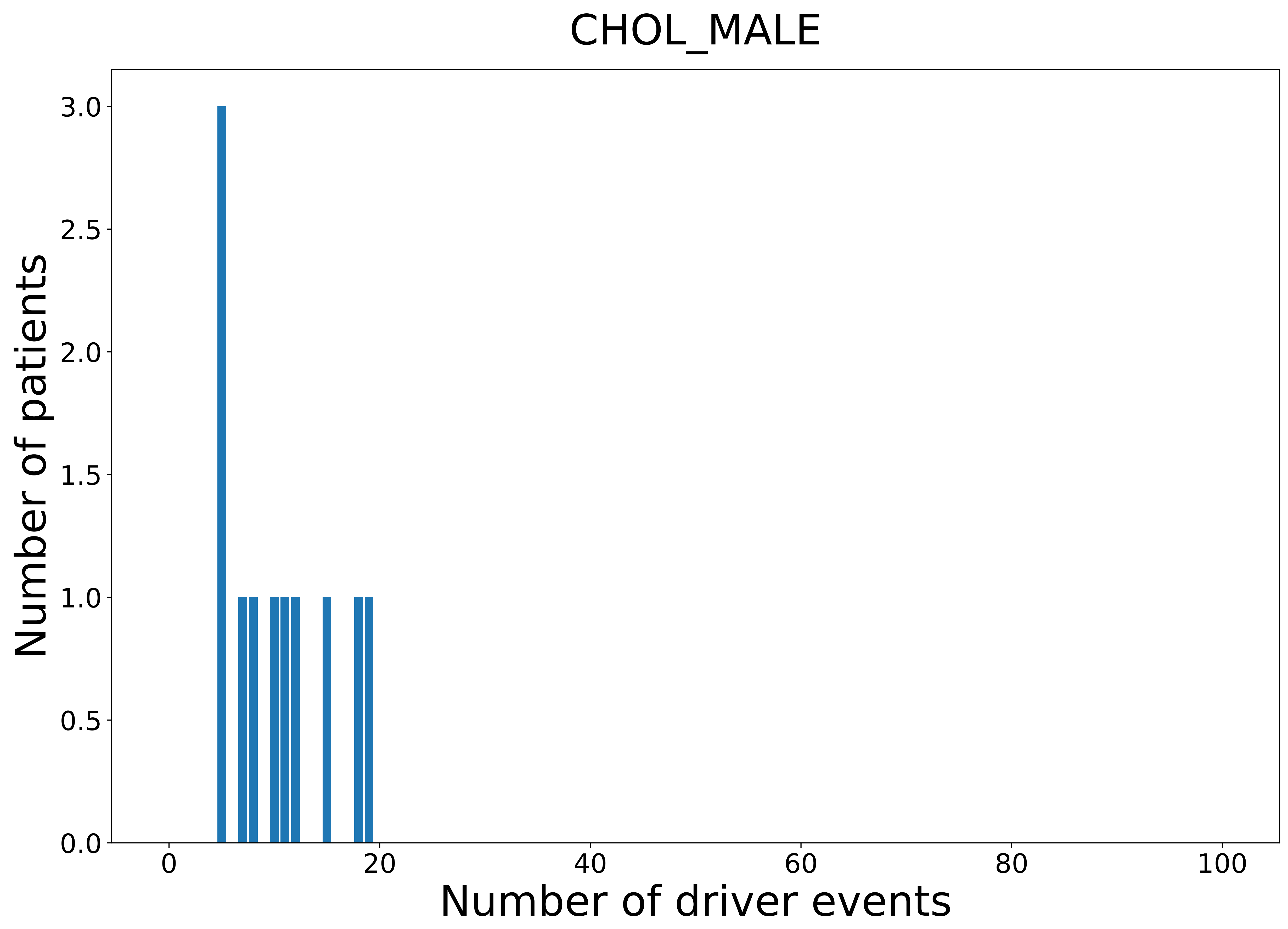

### 2021_8_22_13_29_COAD.pdf

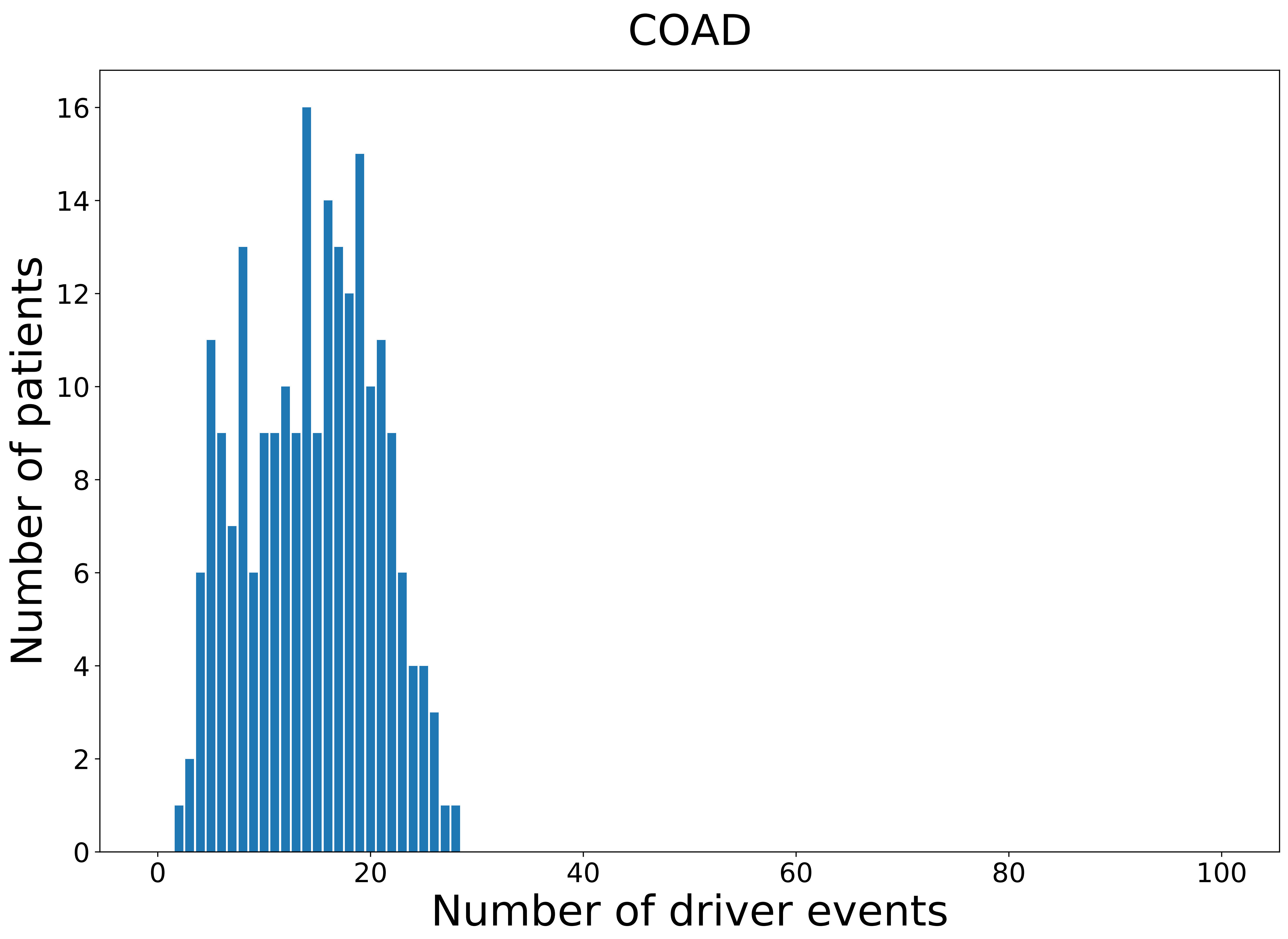

### 2021_8_22_13_29_COAD_MALE.pdf

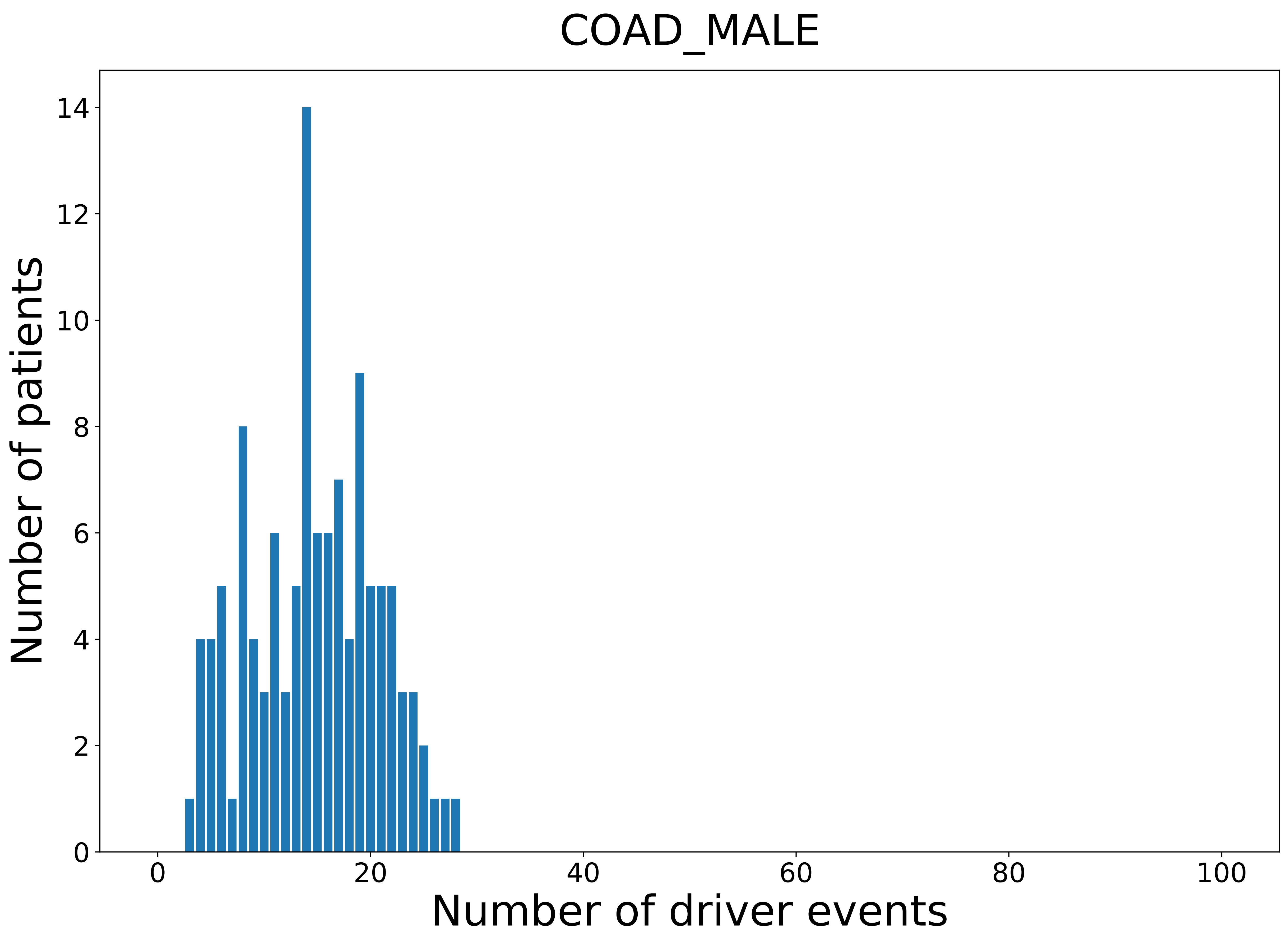

### 2021_8_22_13_29_distribution_age.pdf

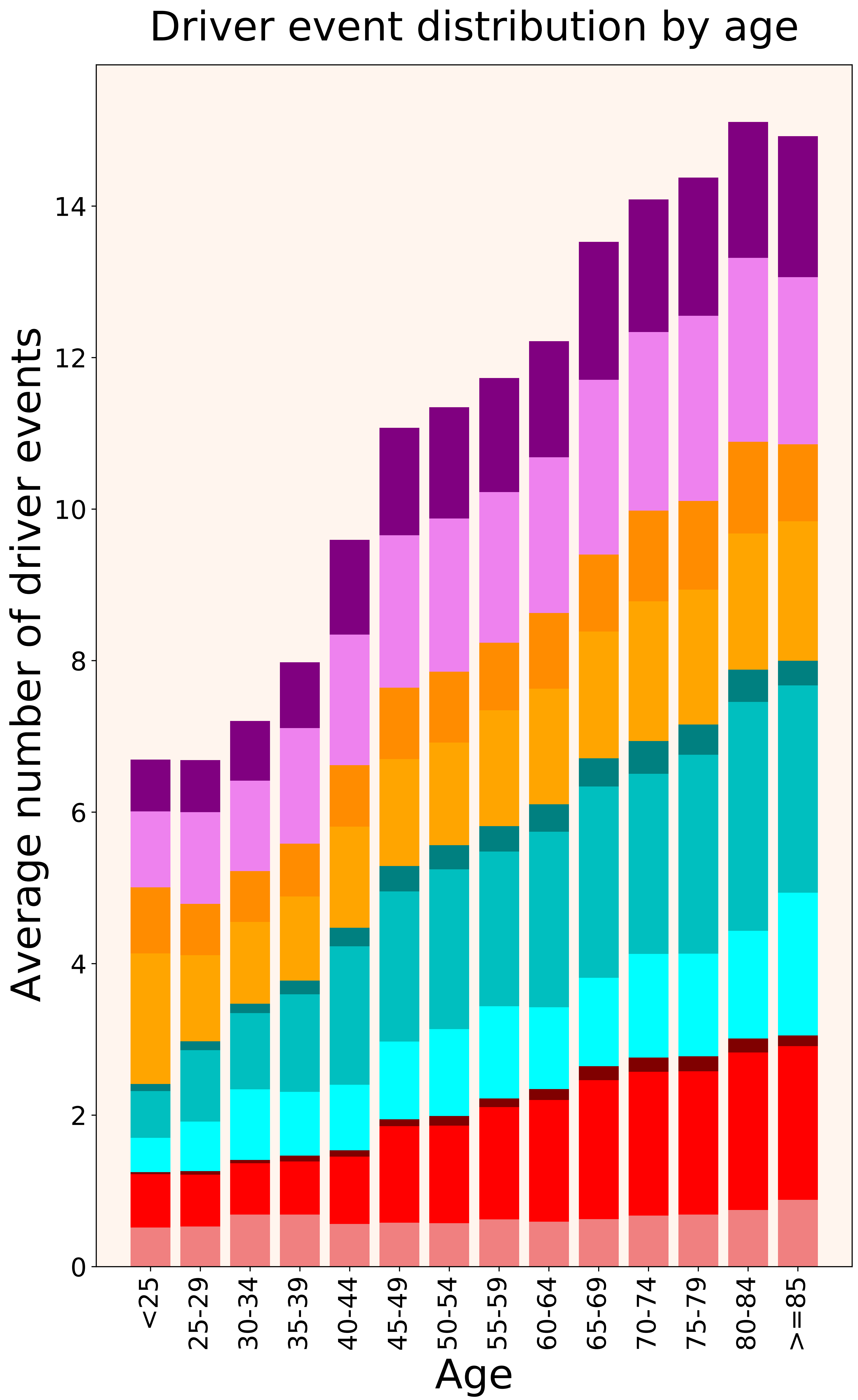

### 2021_8_22_13_29_distribution_age_females.pdf

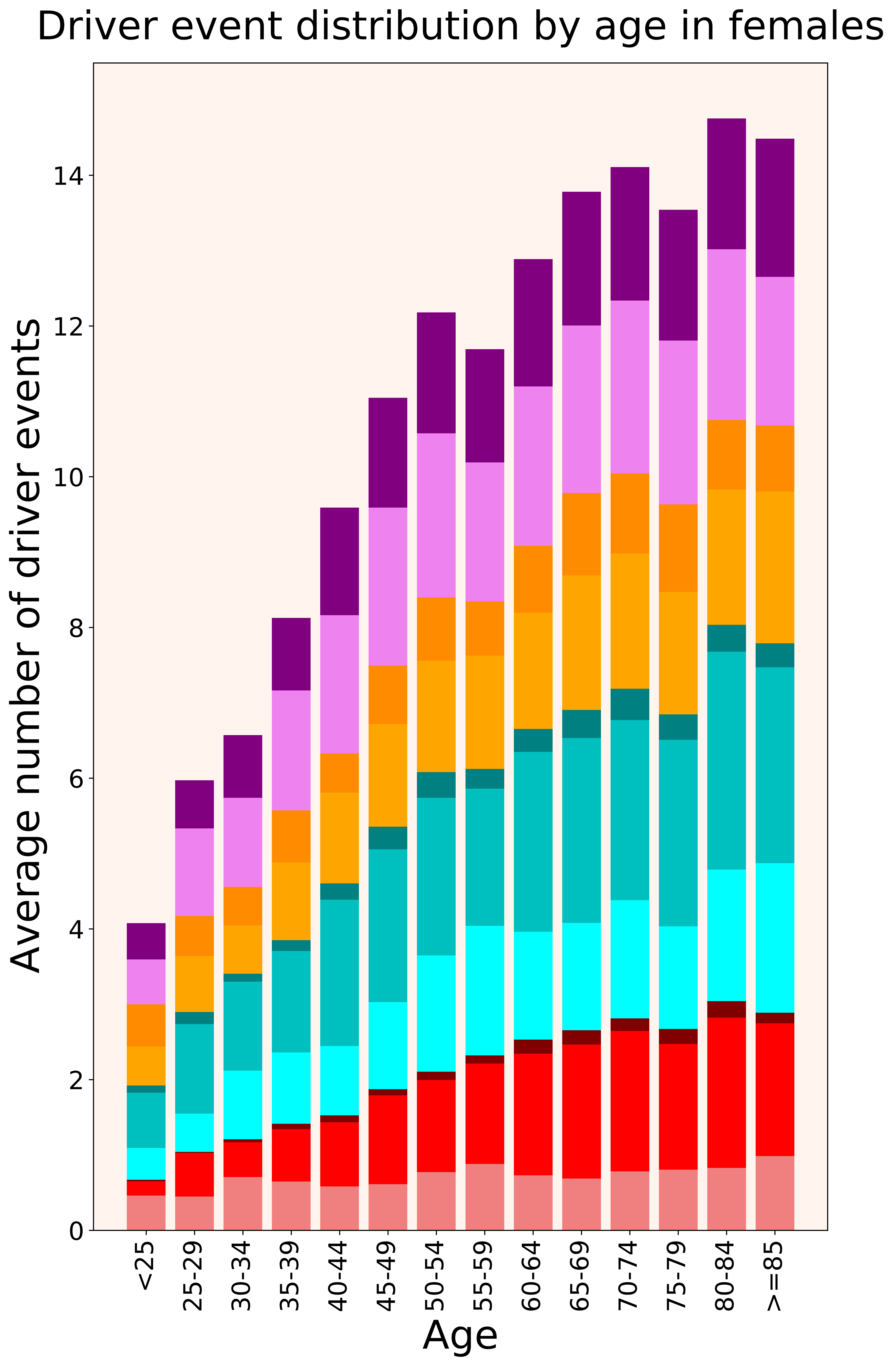

### 2021_8_22_13_29_distribution_age_males.pdf

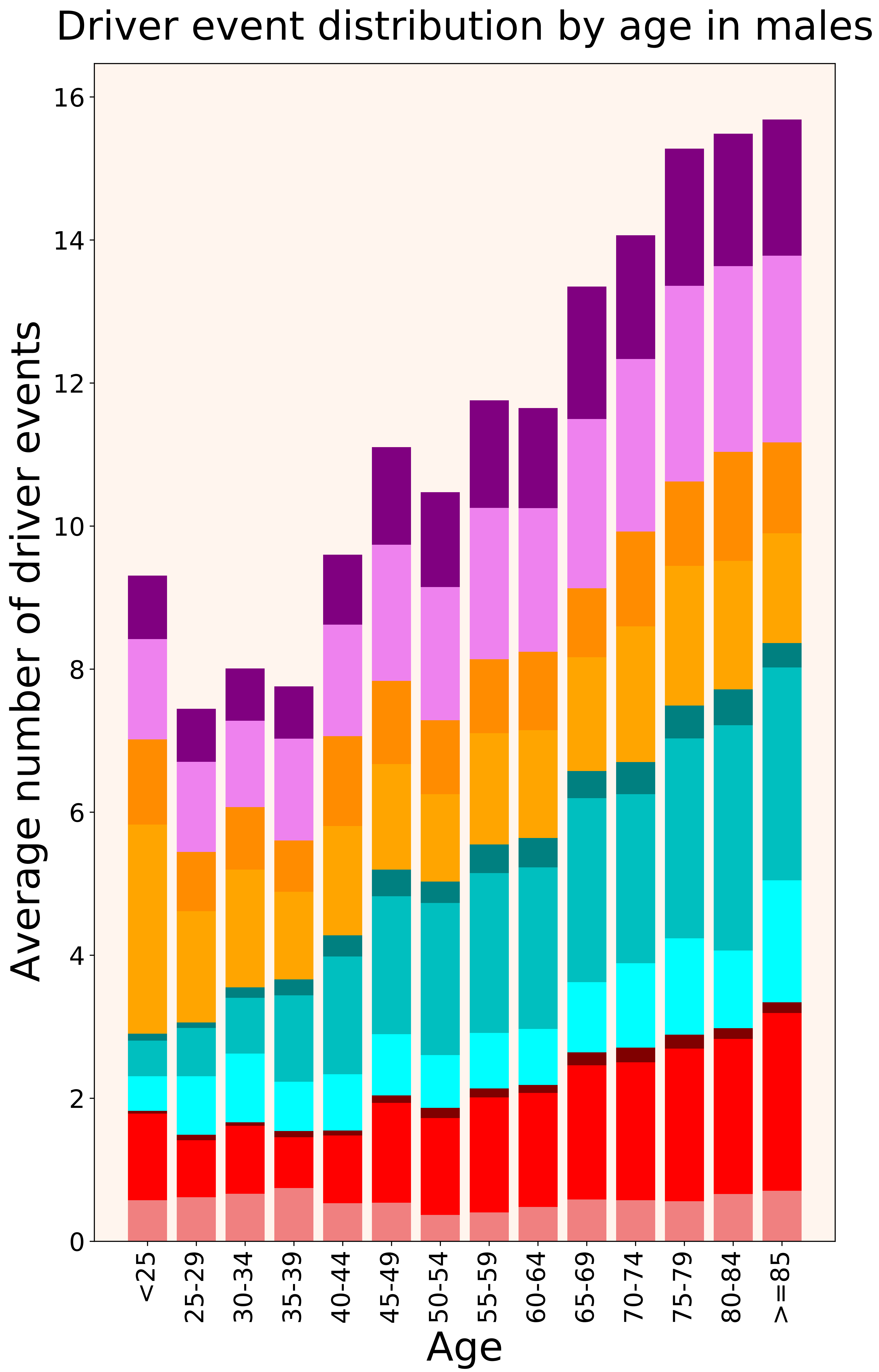

### 2021_8_22_13_29_distribution_cohorts.pdf

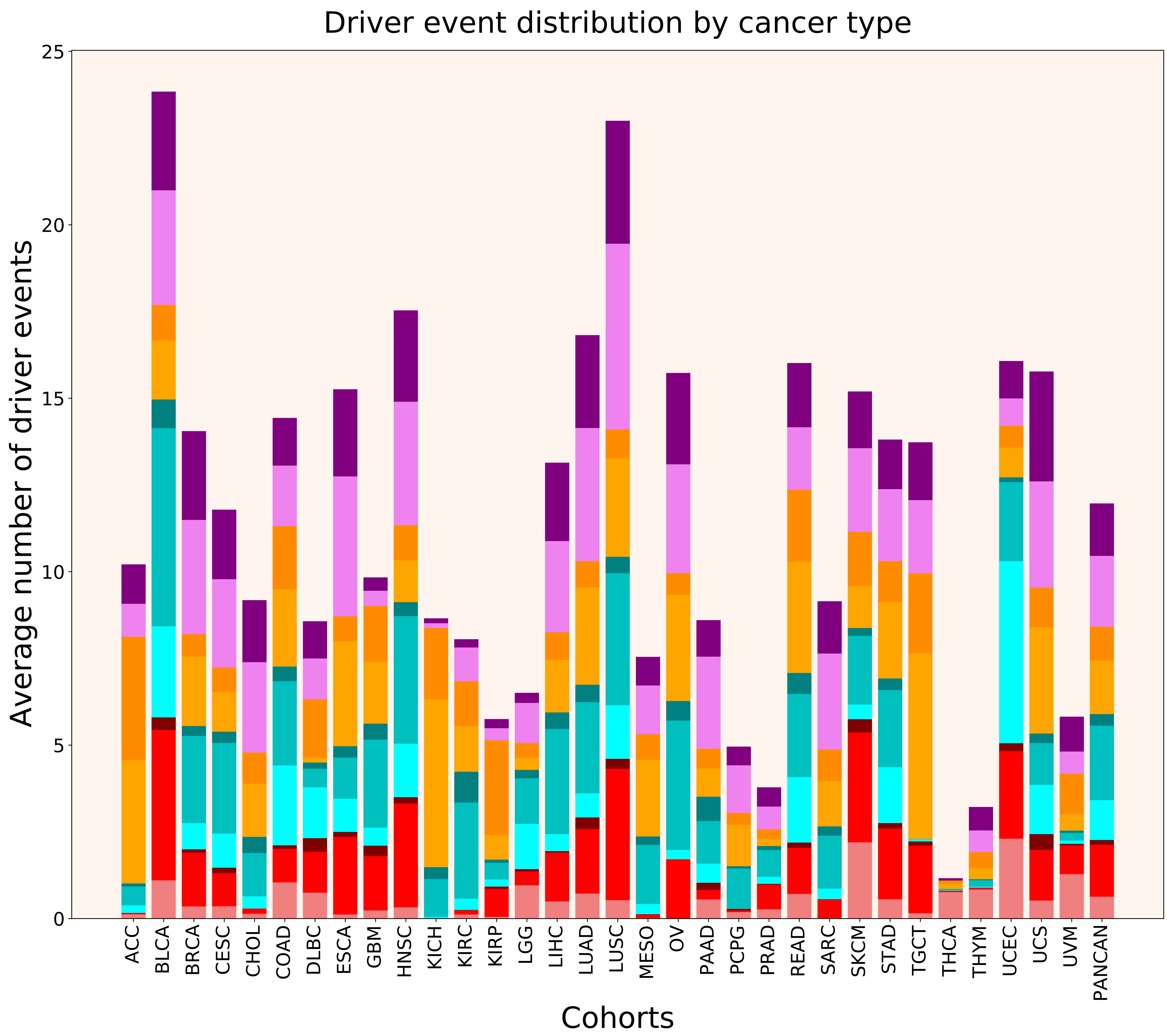

### 2021_8_22_13_29_distribution_cohorts_females.pdf

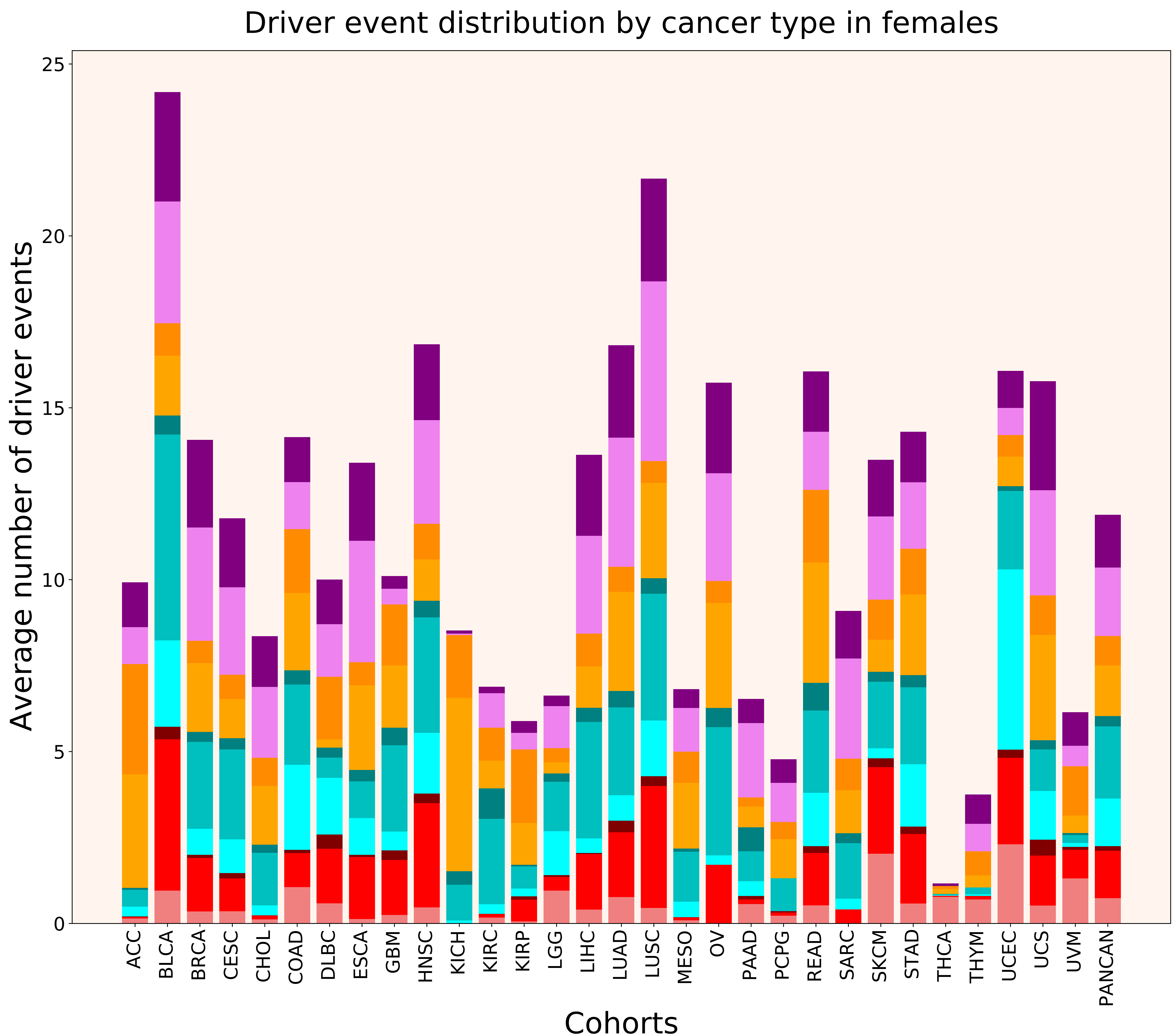

### 2021_8_22_13_29_distribution_cohorts_males.pdf

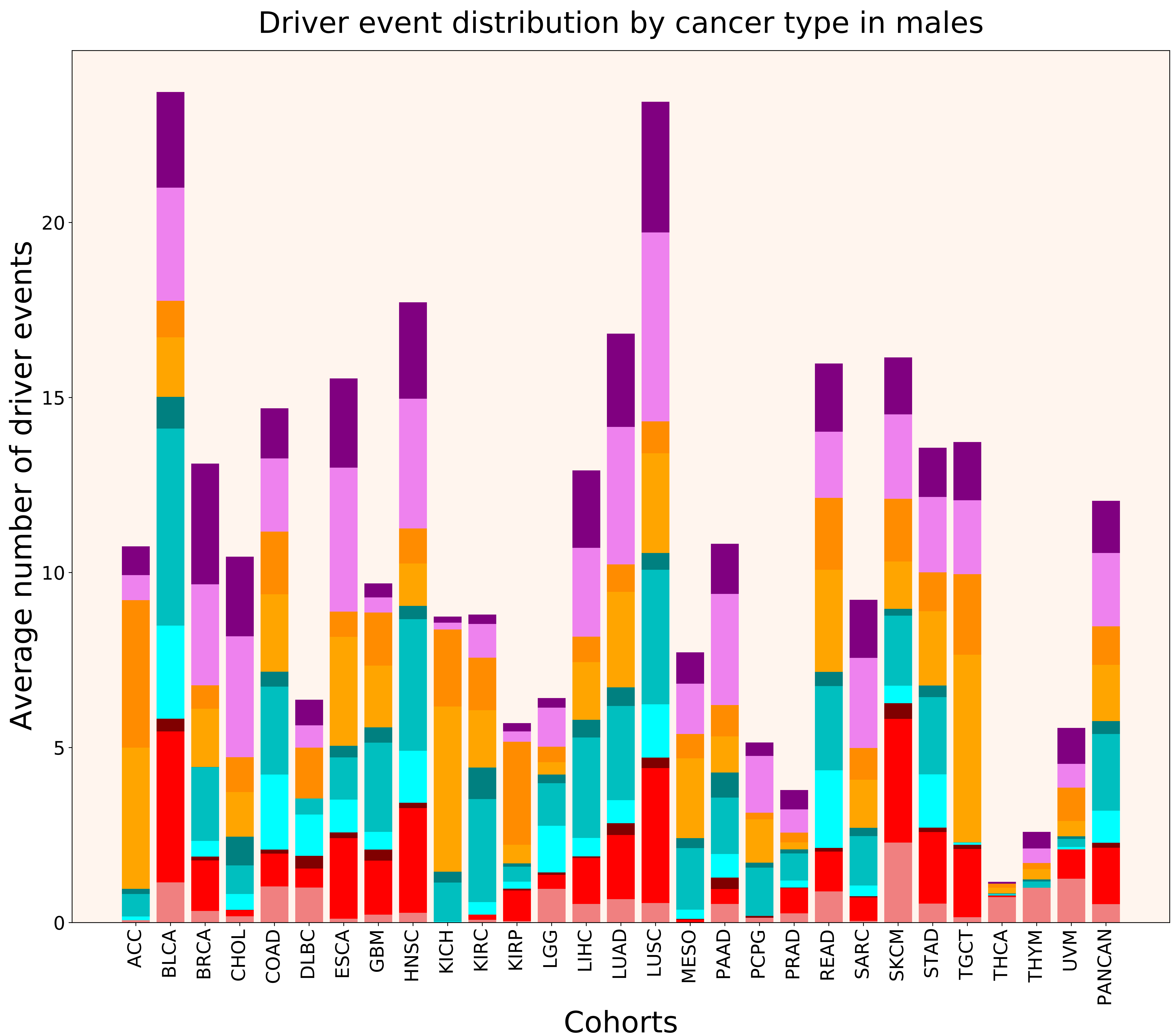

### 2021_8_22_13_29_distribution_events_detailed.pdf

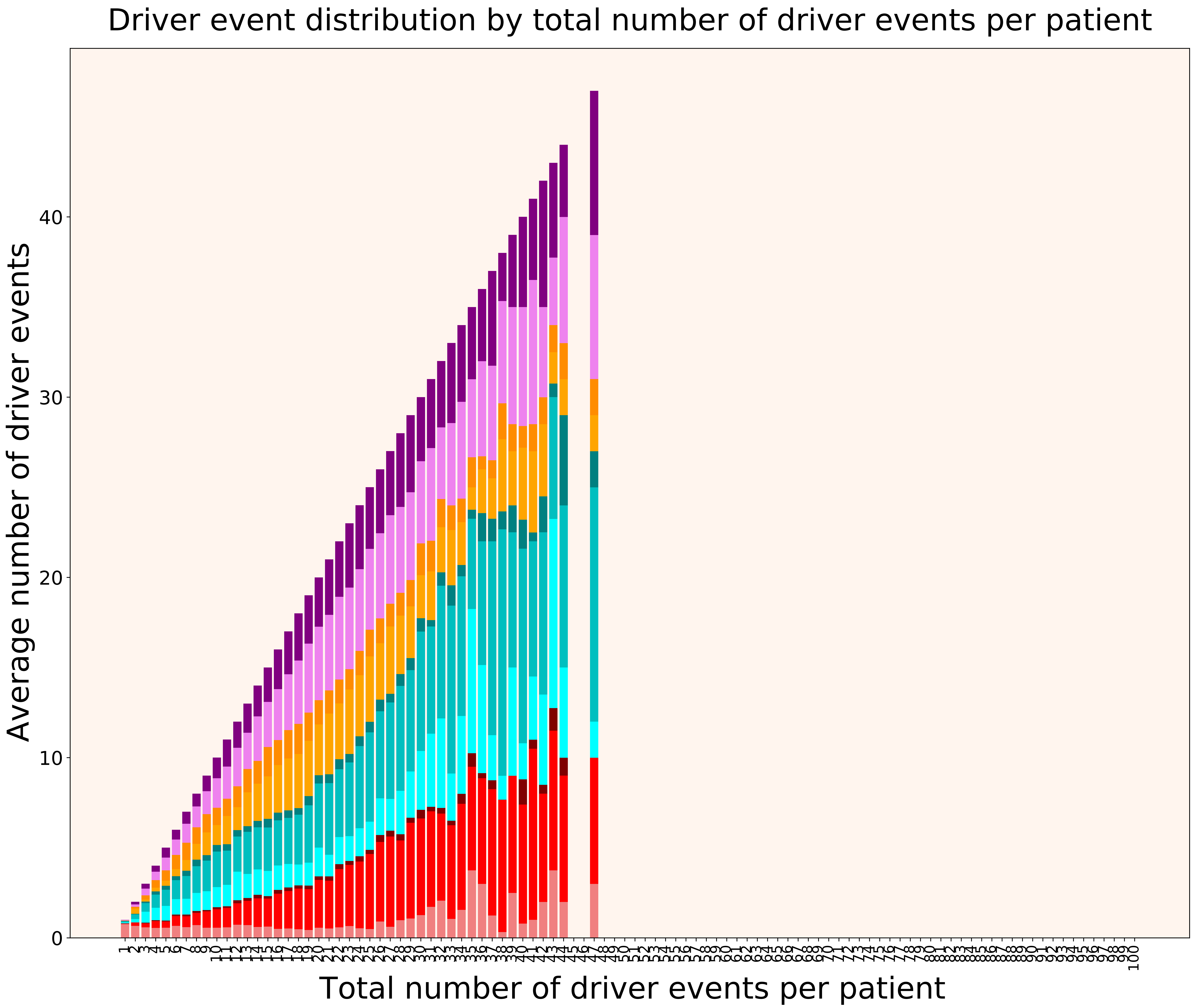

### 2021_8_22_13_29_distribution_events_detailed_females.pdf

#### Driver event distribution by total number of driver events per patient in females

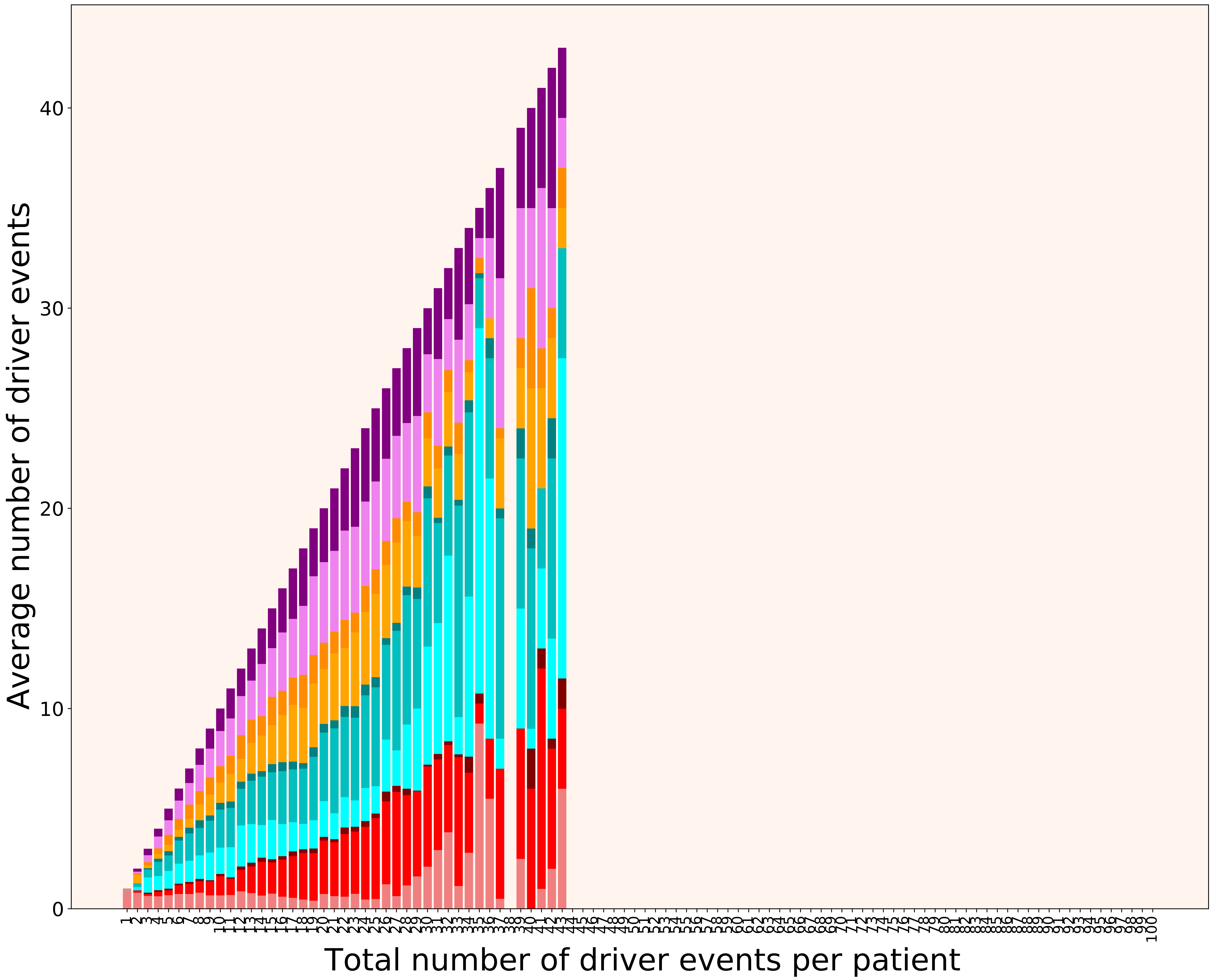

### 2021_8_22_13_29_distribution_events_detailed_males.pdf

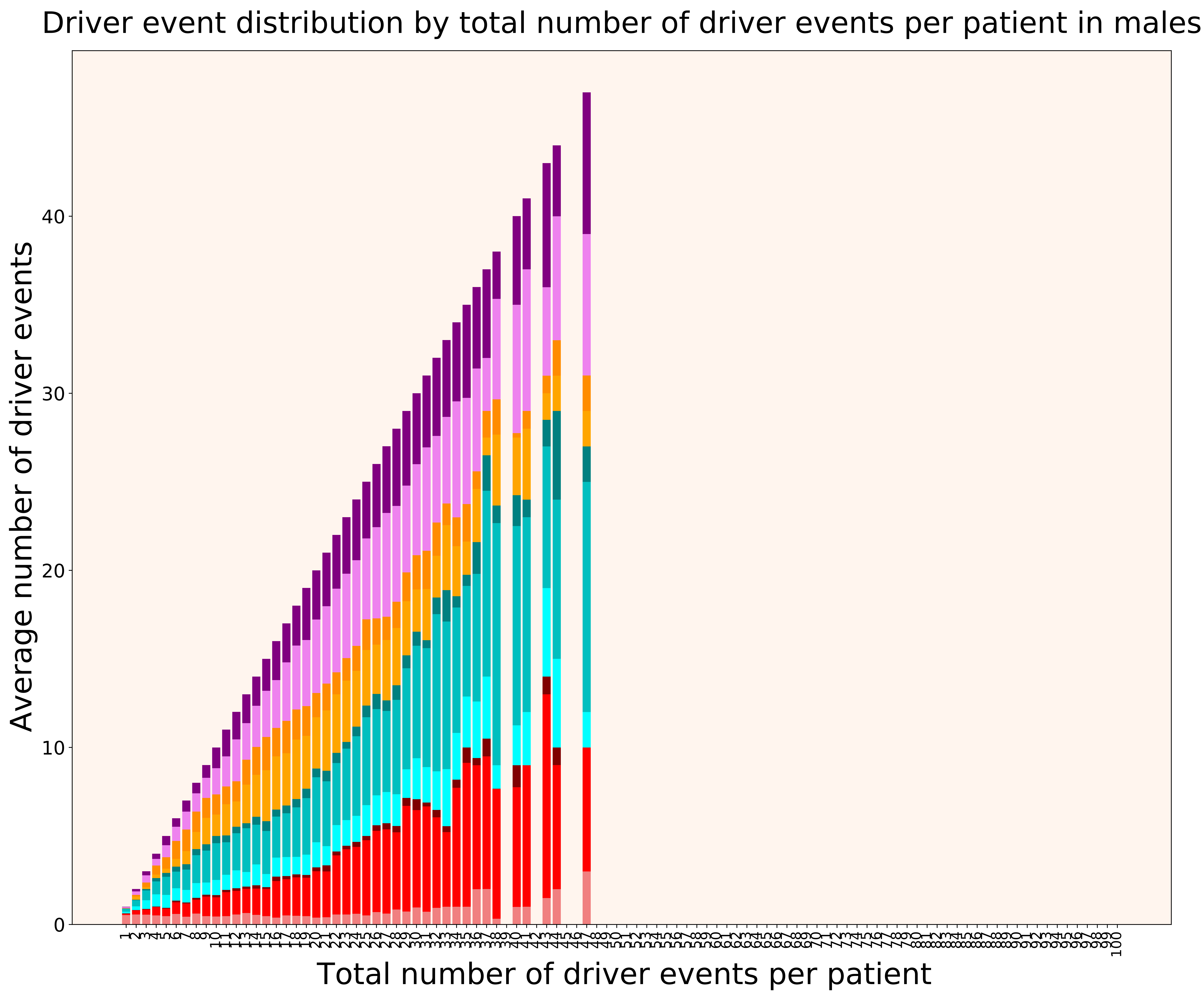

### 2021_8_22_13_29_distribution_gender.pdf

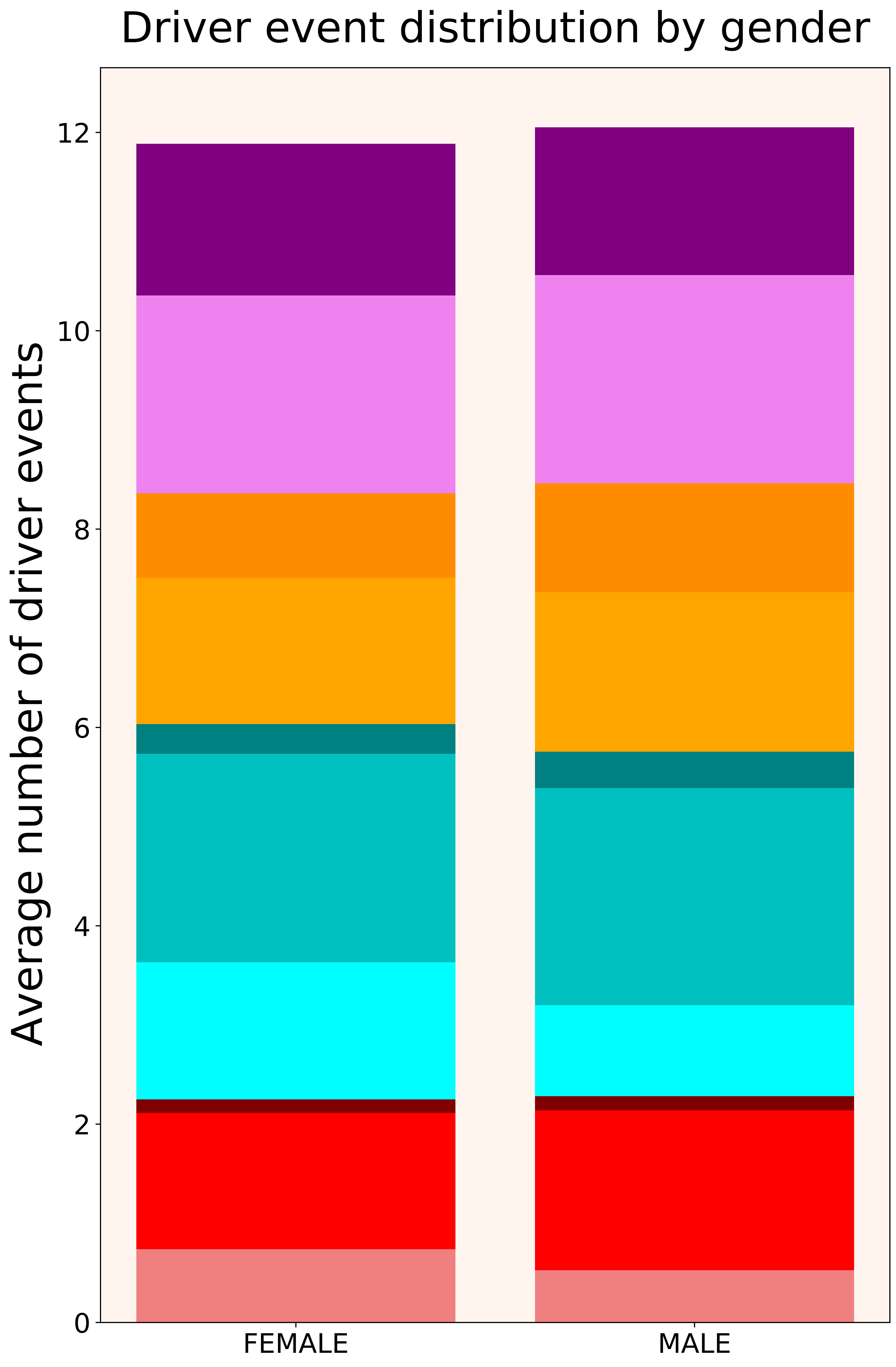

### 2021_8_22_13_29_distribution_stages.pdf

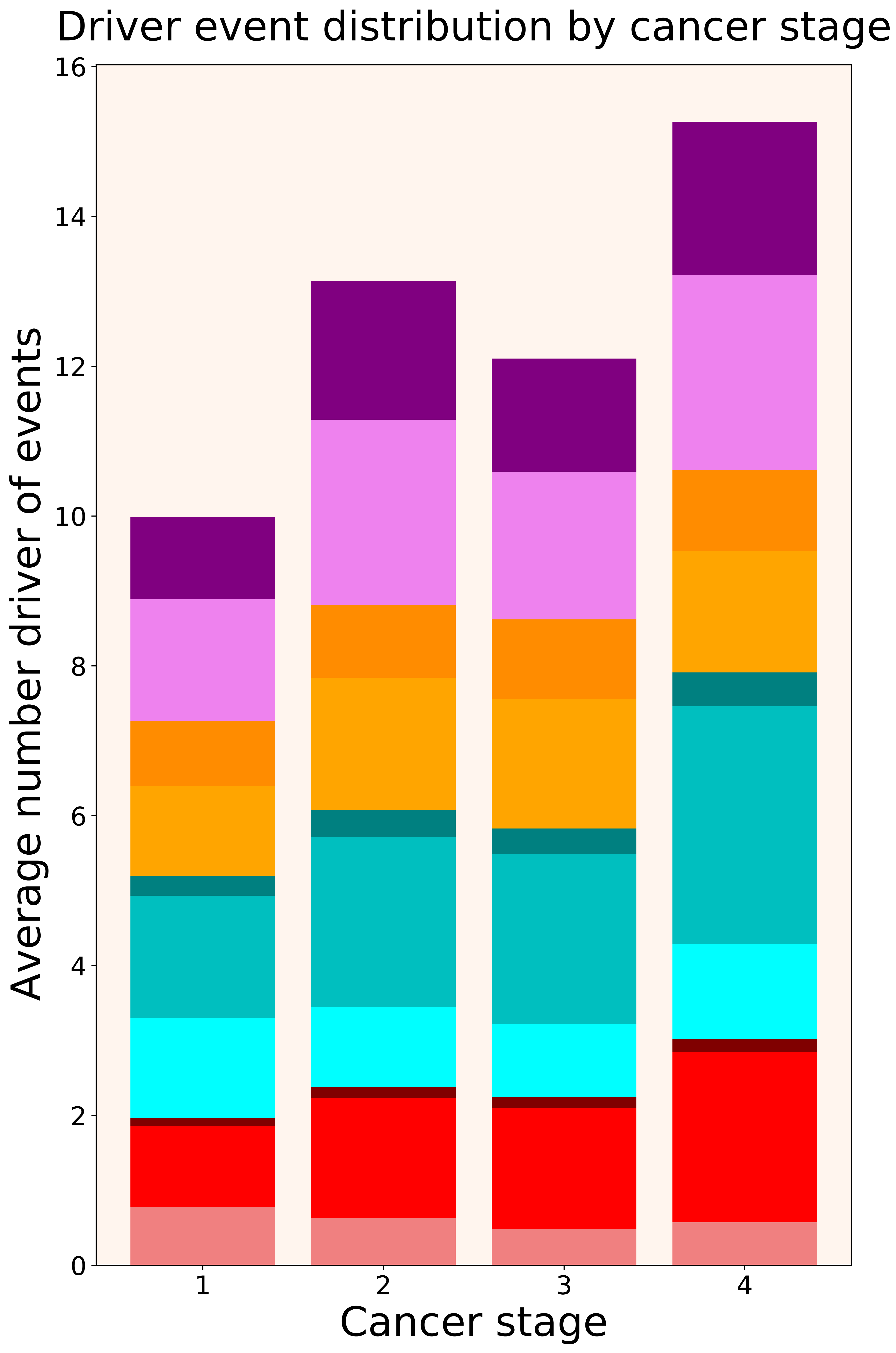

### 2021_8_22_13_29_distribution_stages_females.pdf

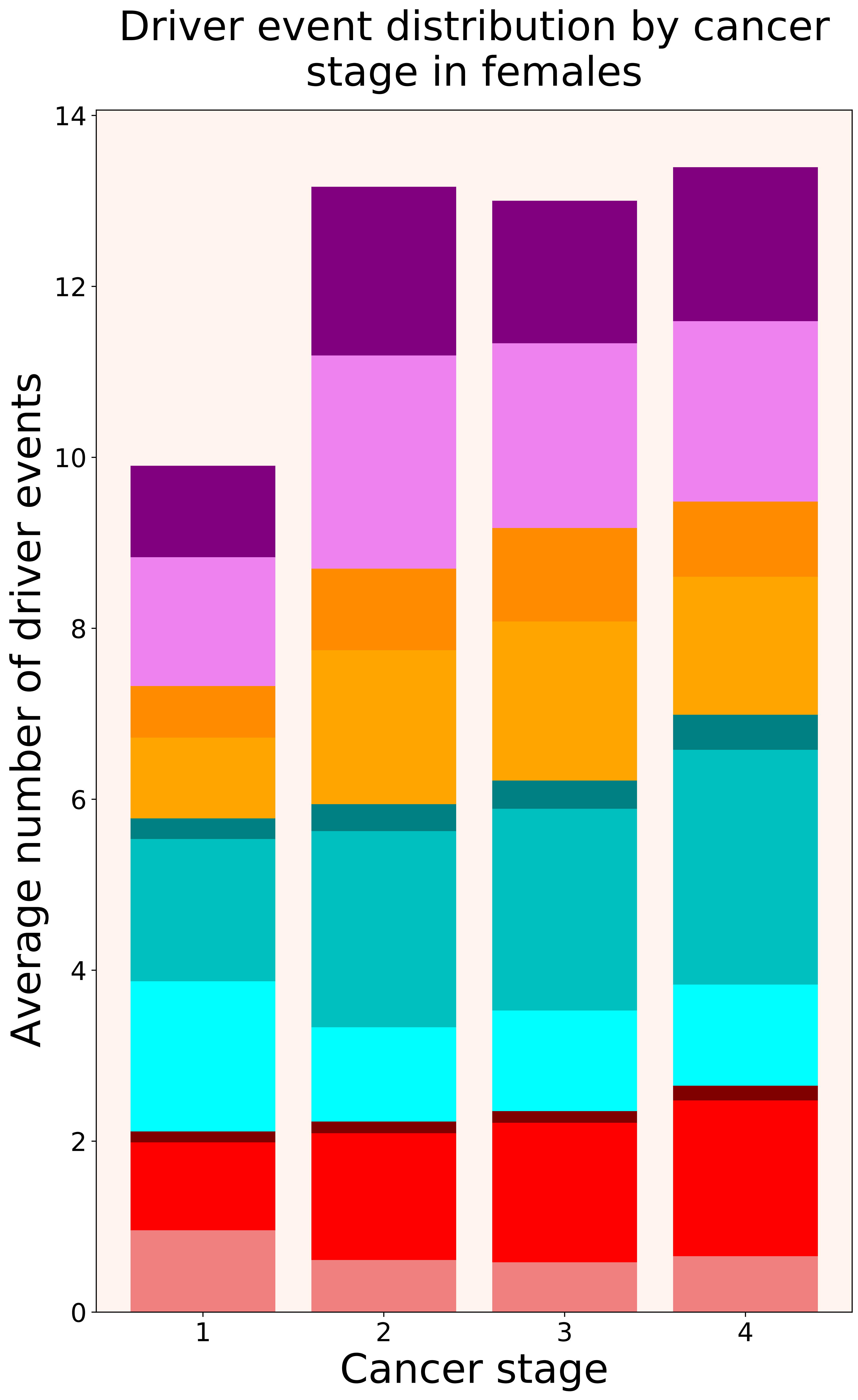

### 2021_8_22_13_29_distribution_stages_males.pdf

#### Driver event distribution by cancer stage in males

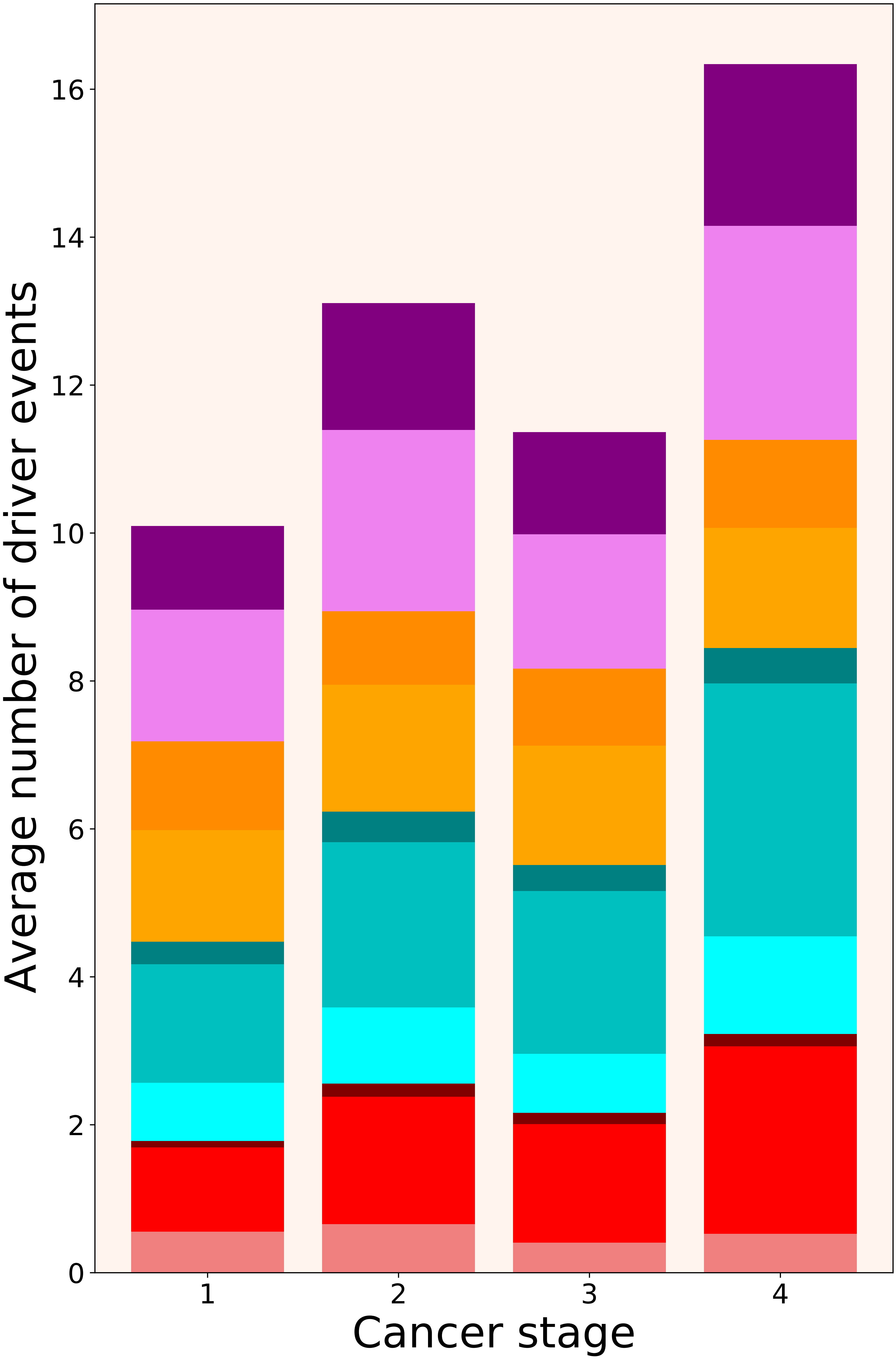

### 2021_8_22_13_29_DLBC.pdf

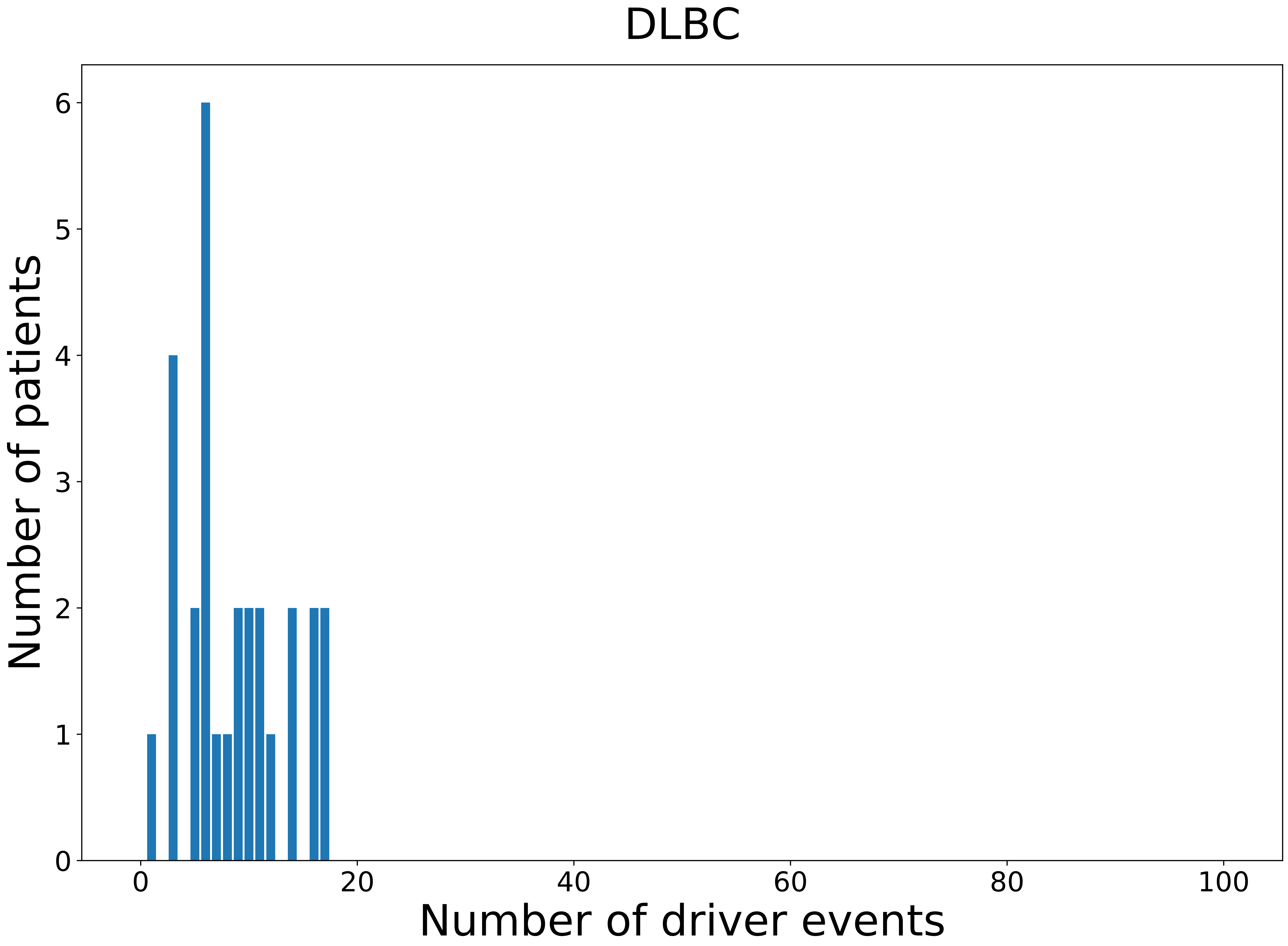

### 2021_8_22_13_29_DLBC_FEMALE.pdf

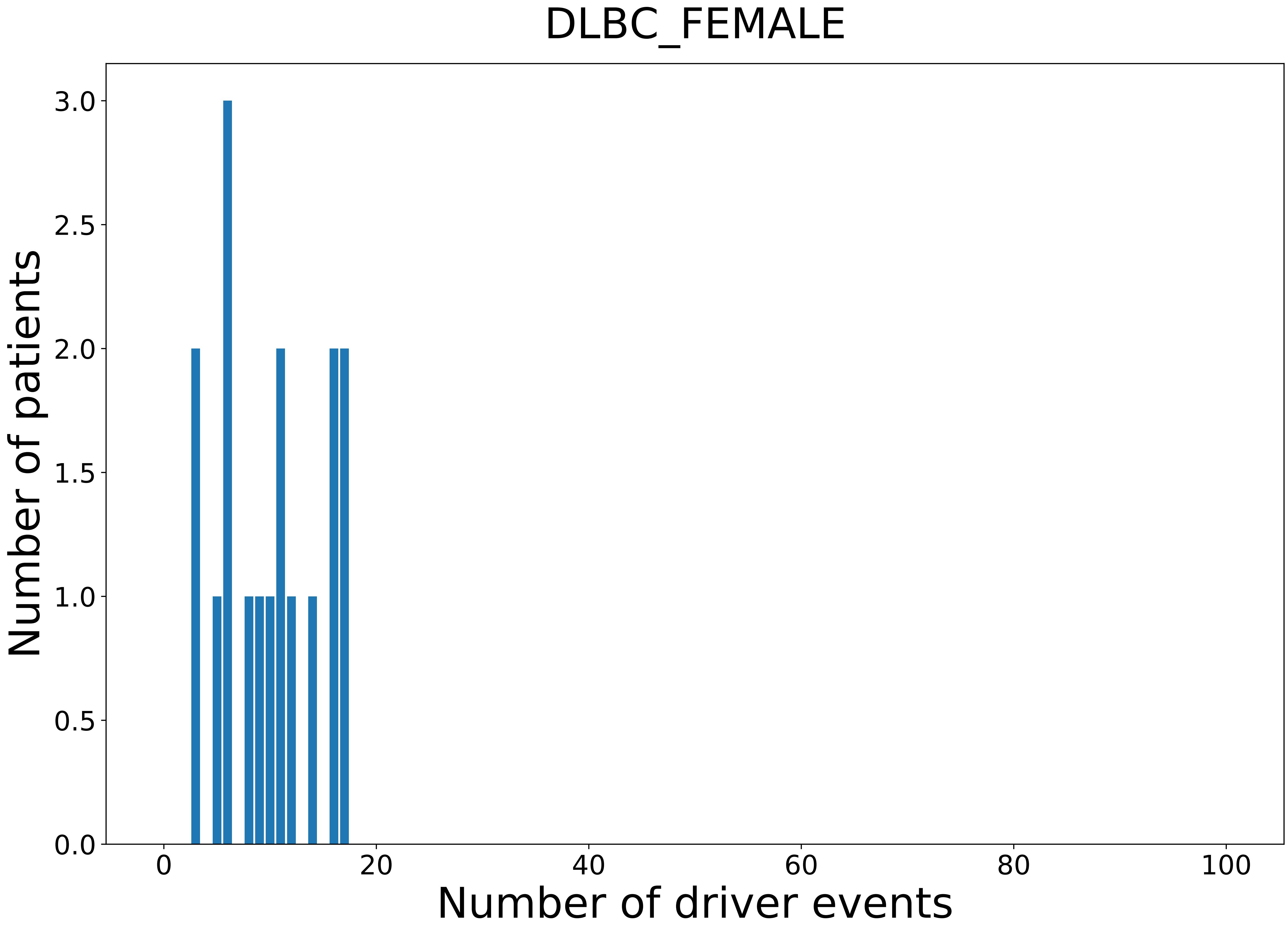

### 2021_8_22_13_29_DLBC_MALE.pdf

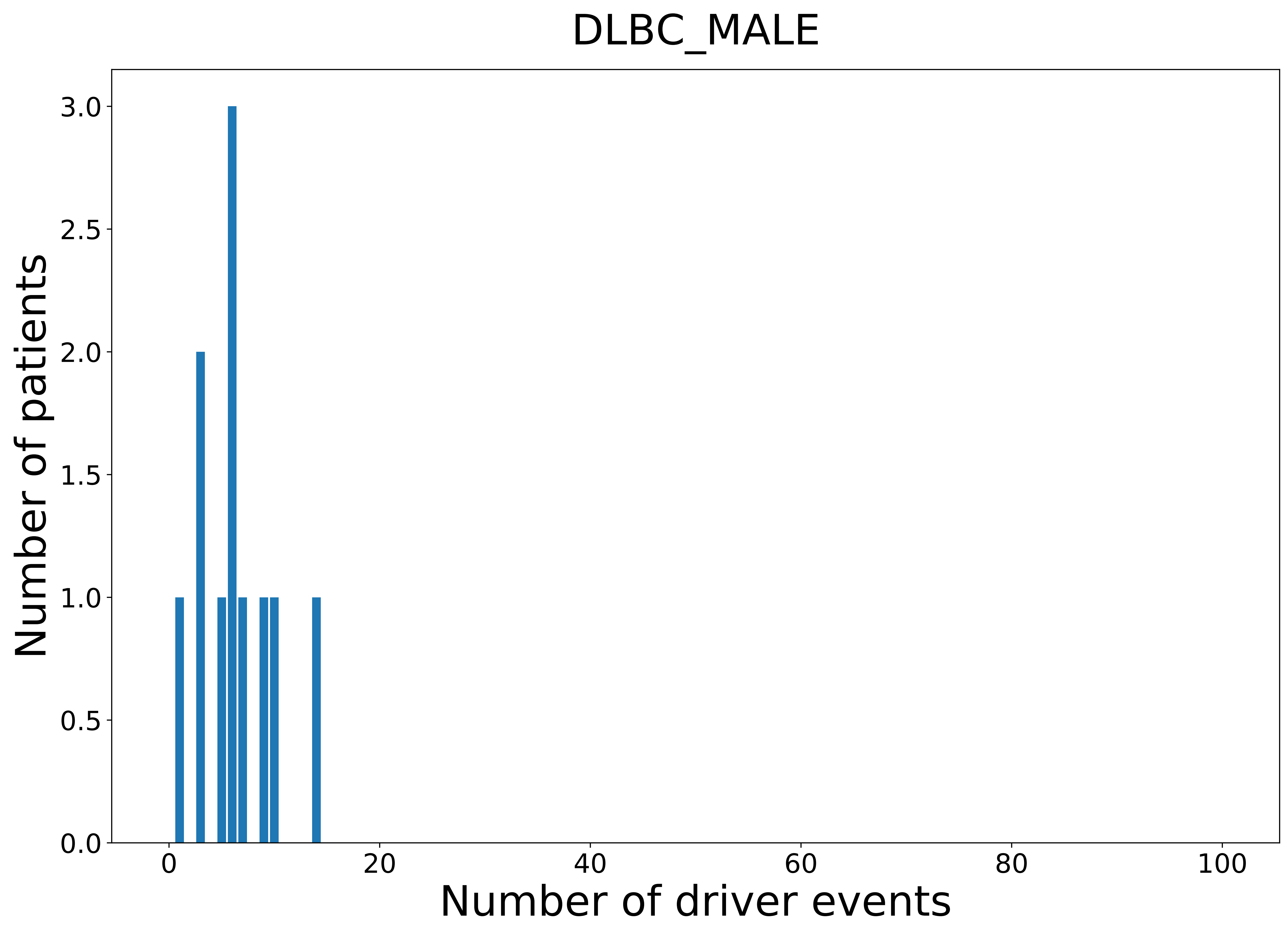

### 2021_8_22_13_29_ESCA.pdf

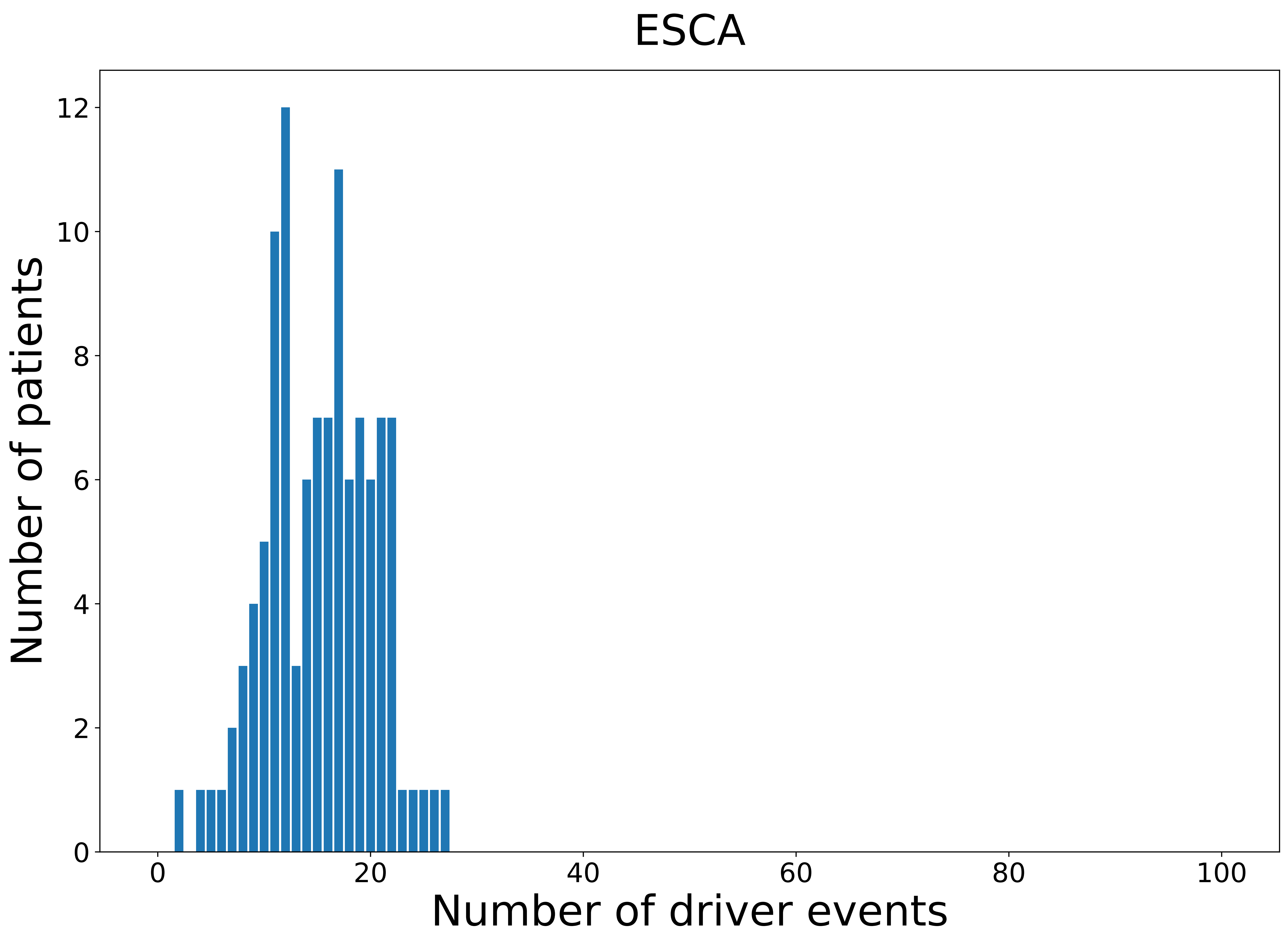

### 2021_8_22_13_29_ESCA_FEMALE.pdf

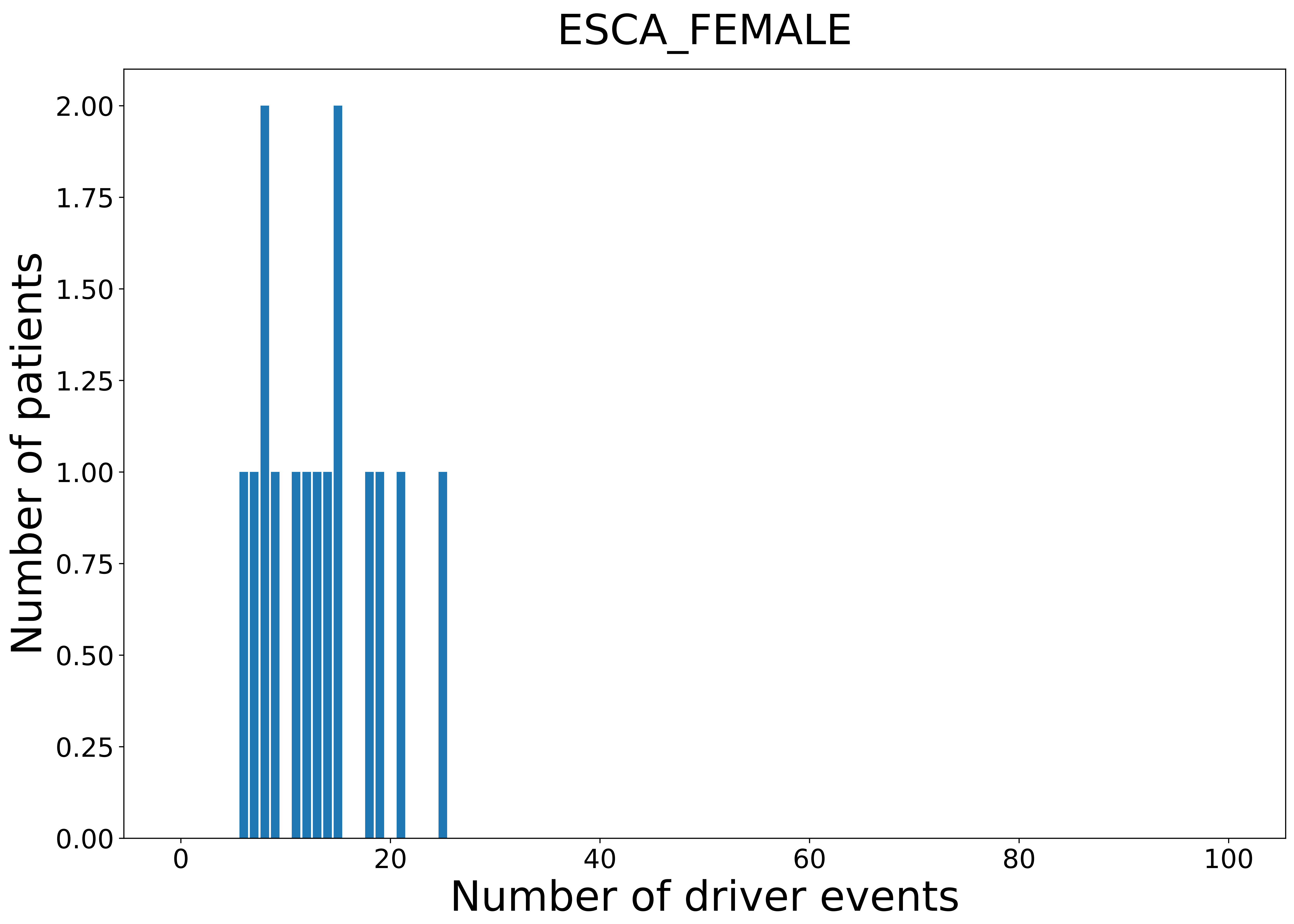

### 2021_8_22_13_29_ESCA_MALE.pdf

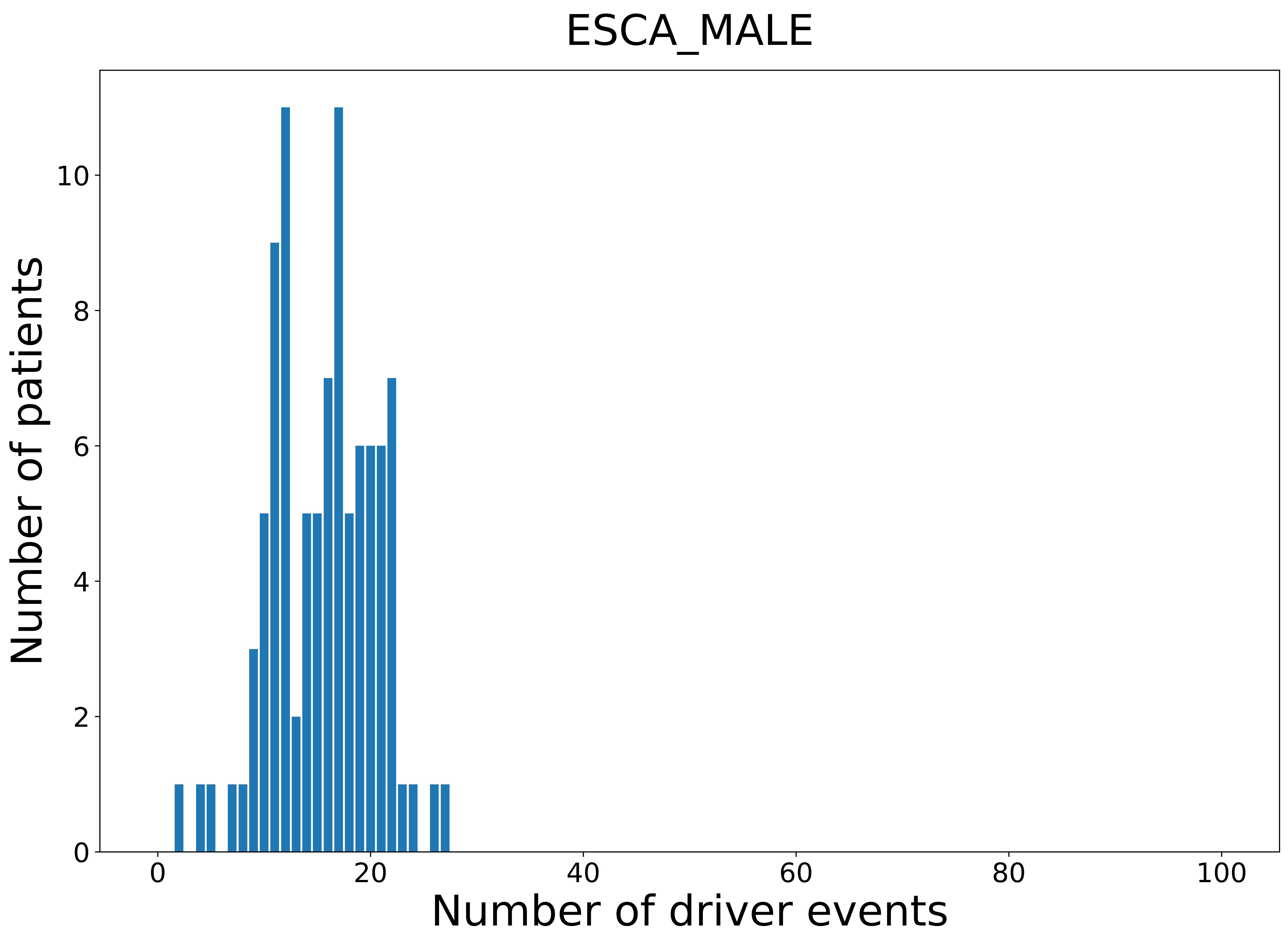

### 2021_8_22_13_29_GBM.pdf

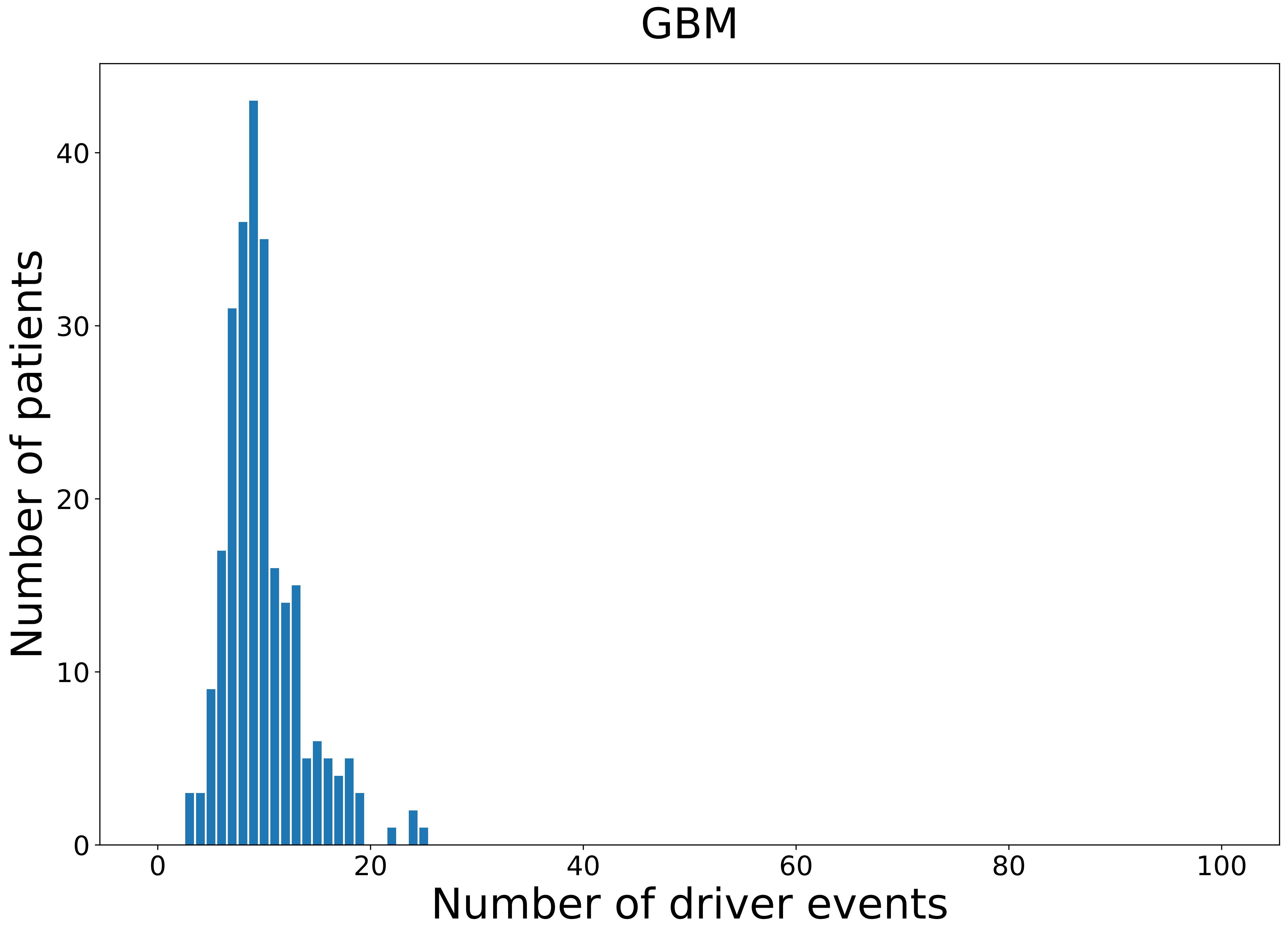

### 2021_8_22_13_29_GBM_FEMALE.pdf

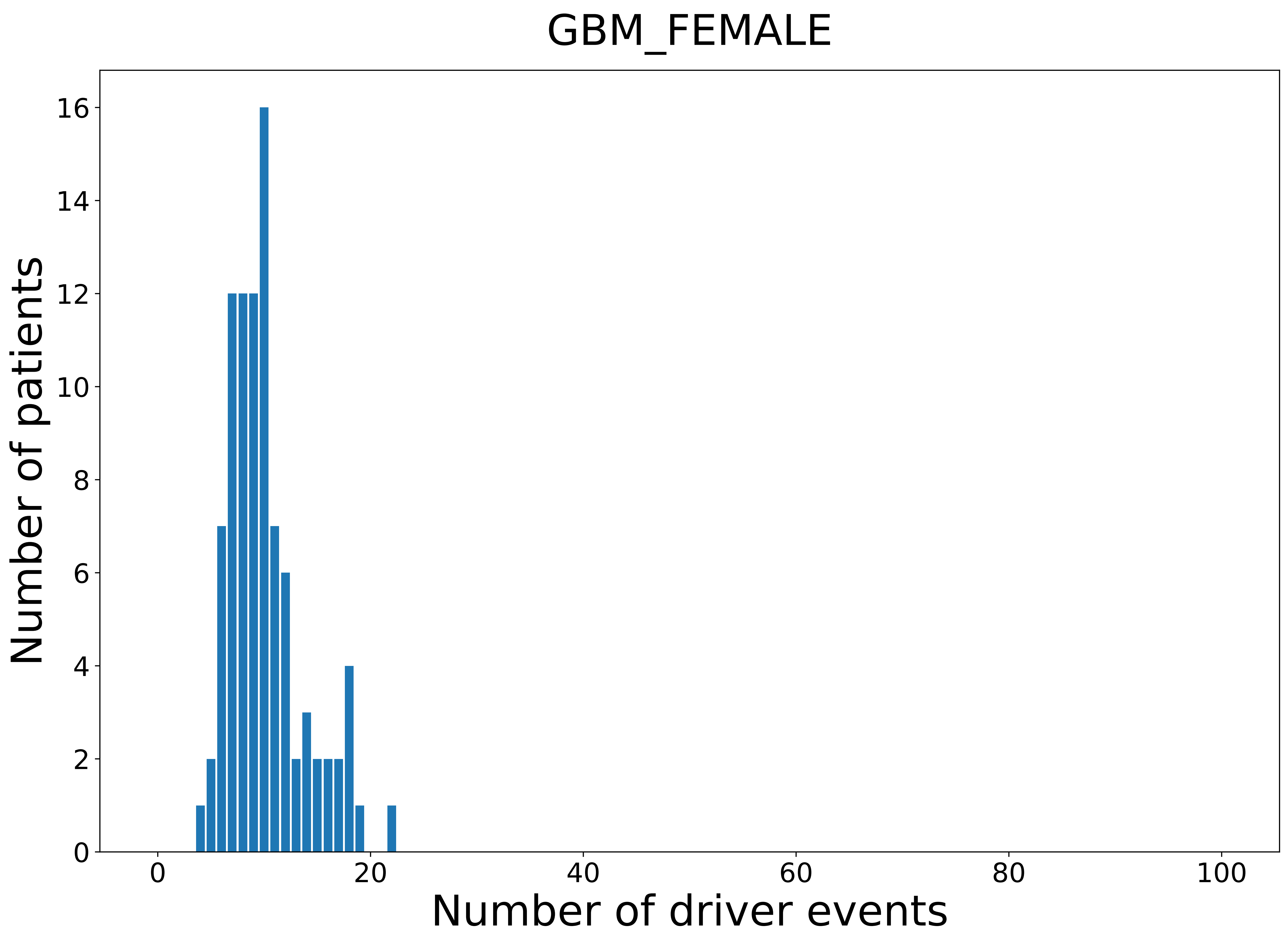

### 2021_8_22_13_29_GBM_MALE.pdf

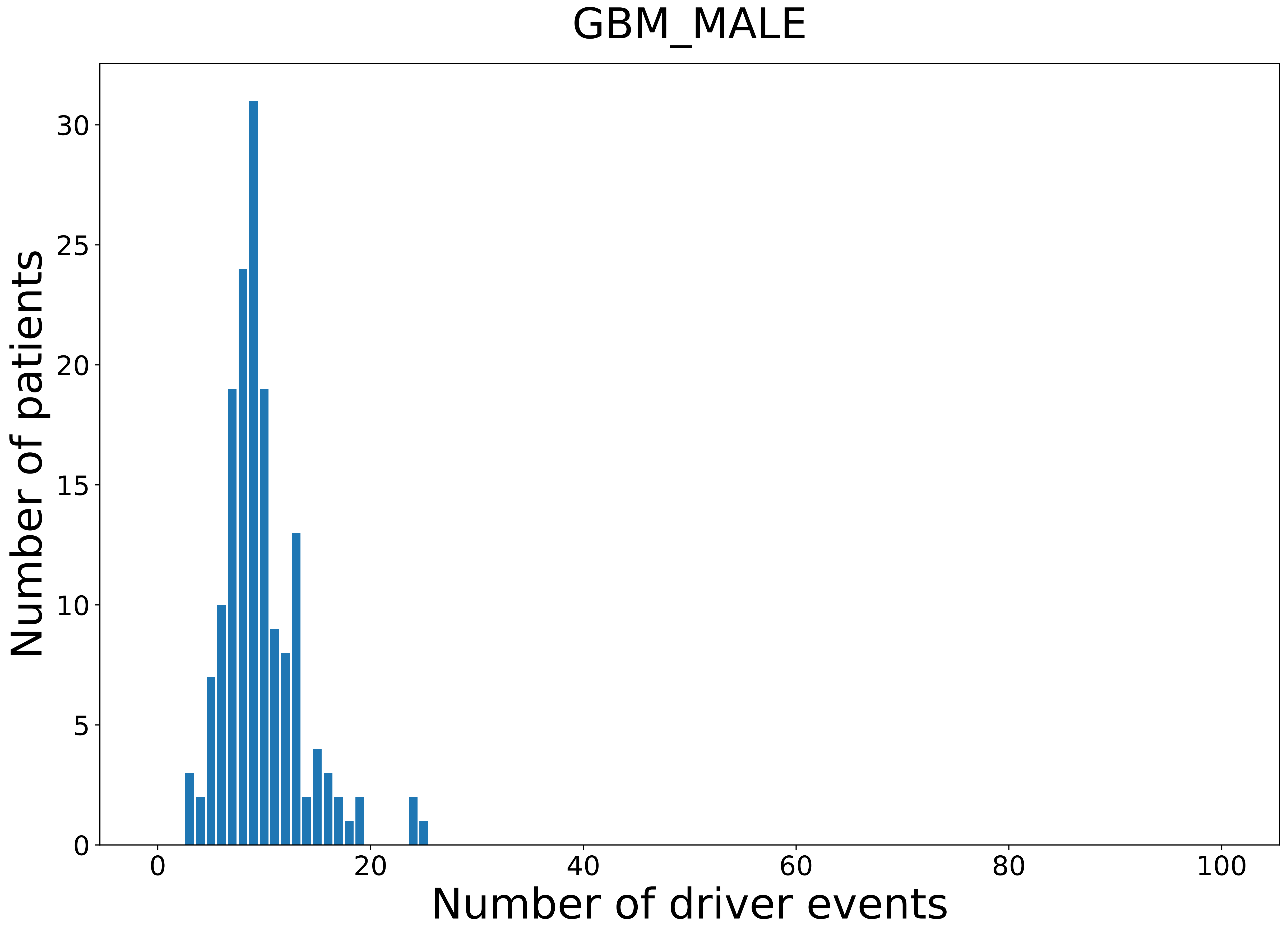

### 2021_8_22_13_29_HNSC.pdf

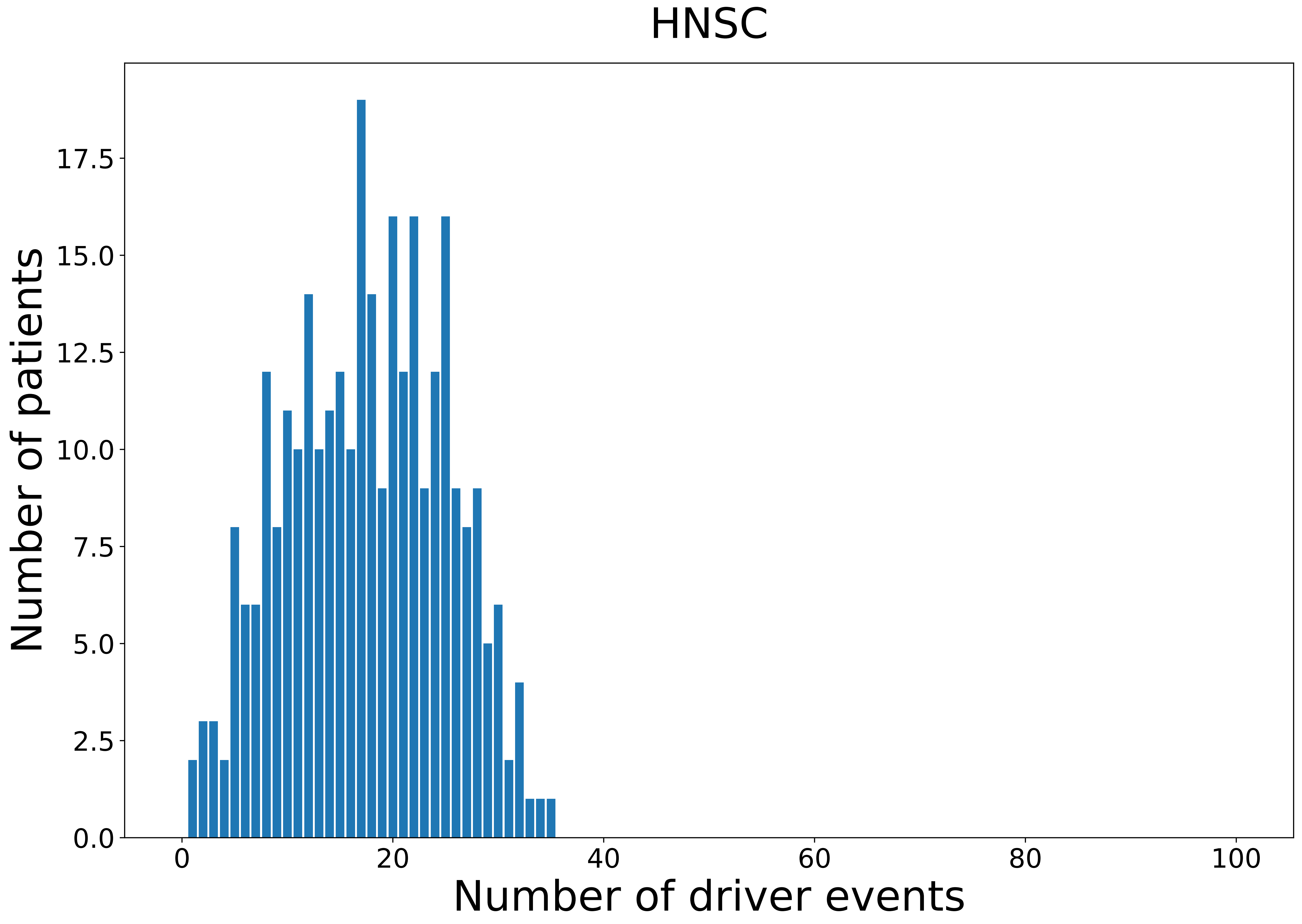

### 2021_8_22_13_29_HNSC_FEMALE.pdf

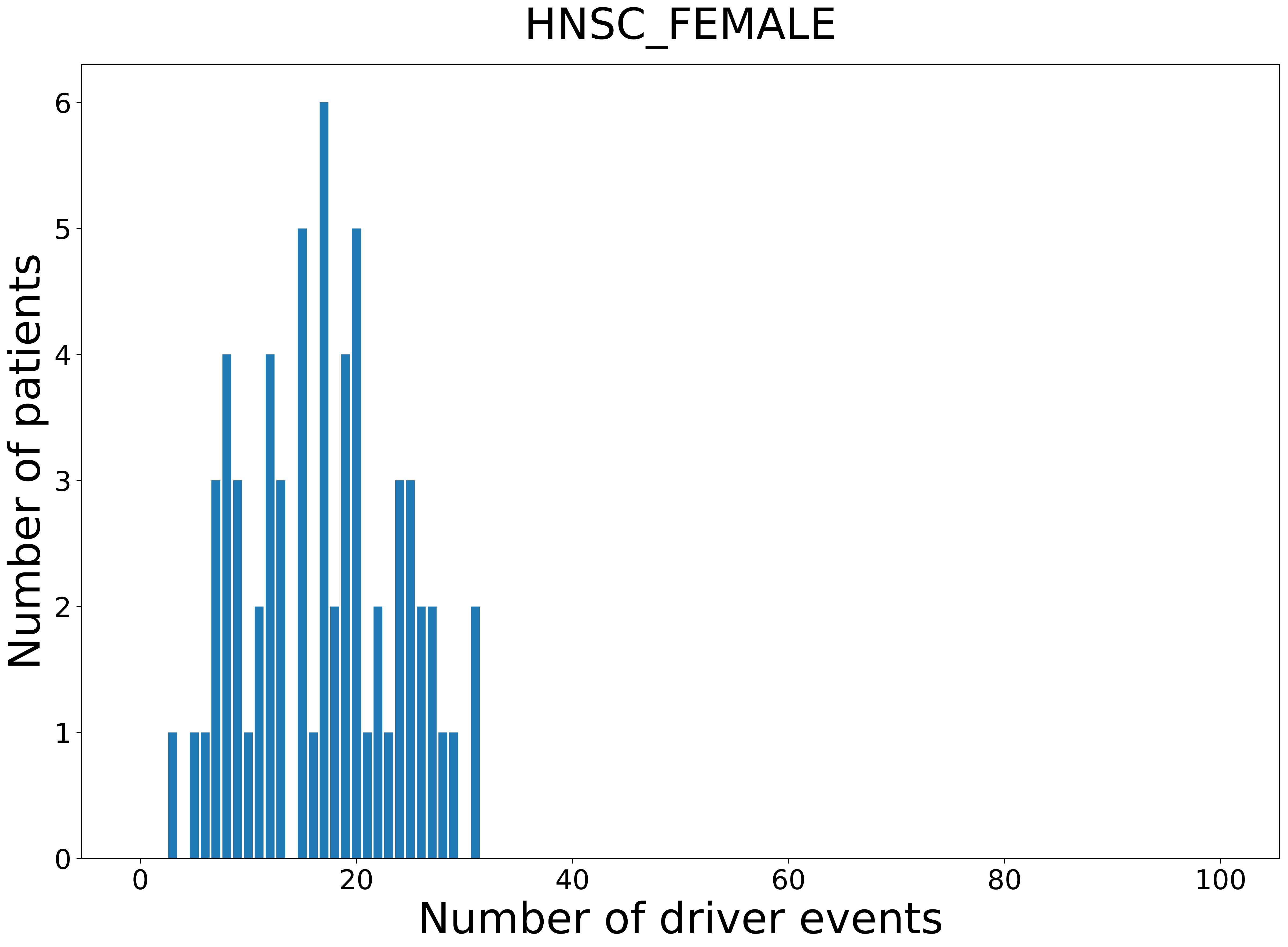

### 2021_8_22_13_29_HNSC_MALE.pdf

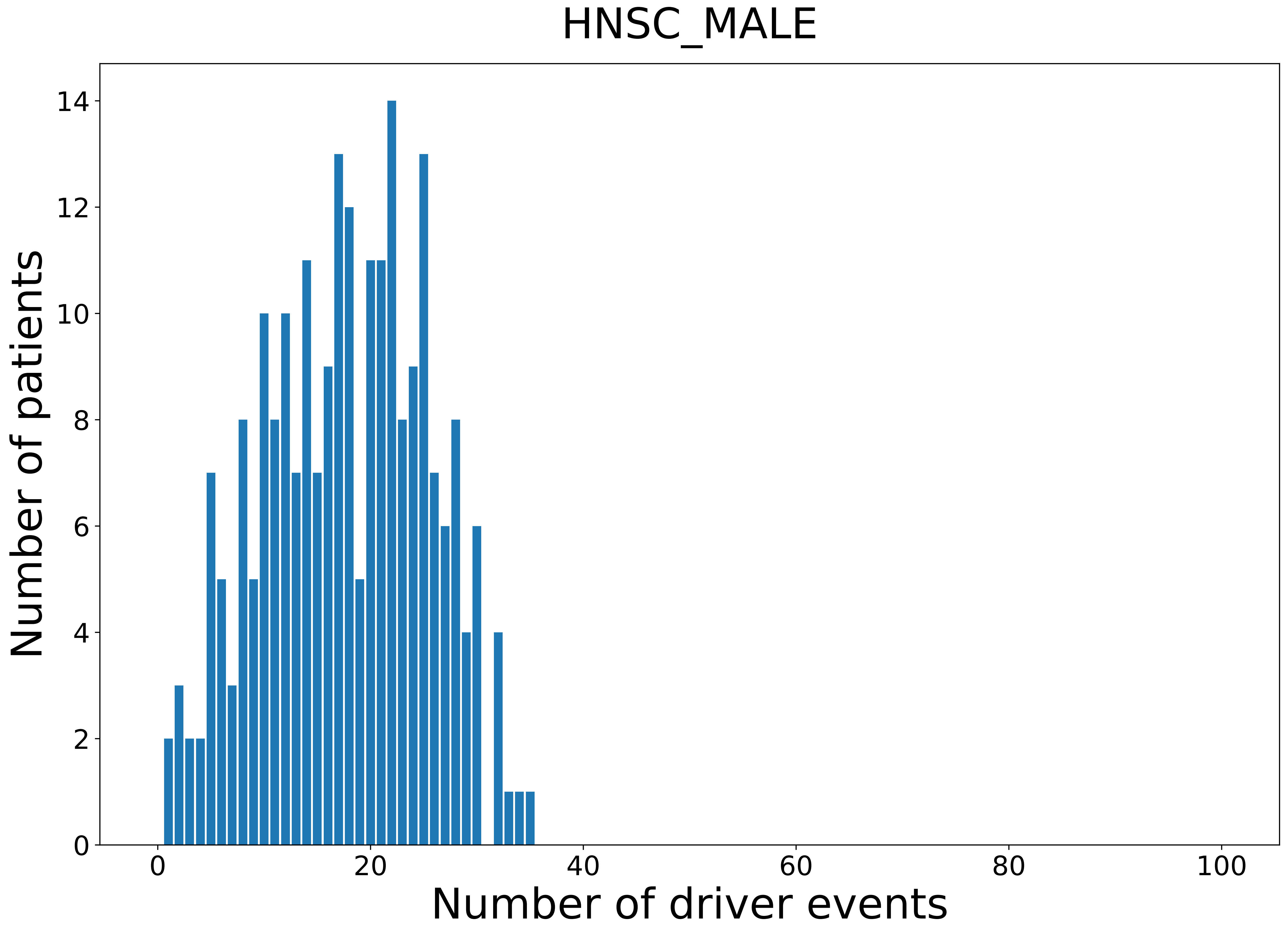

### 2021_8_22_13_29_KICH.pdf

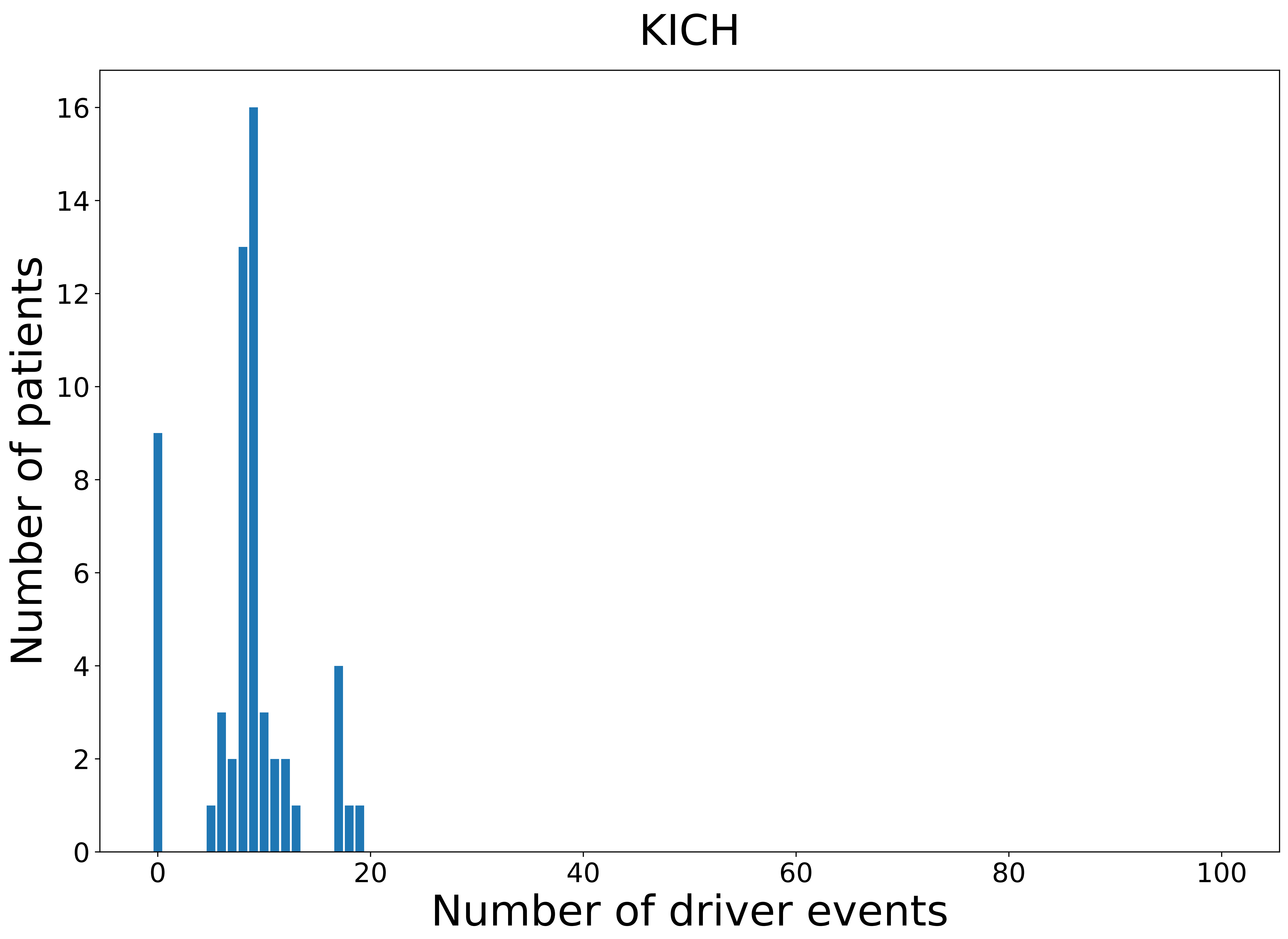

### 2021_8_22_13_29_KICH_FEMALE.pdf

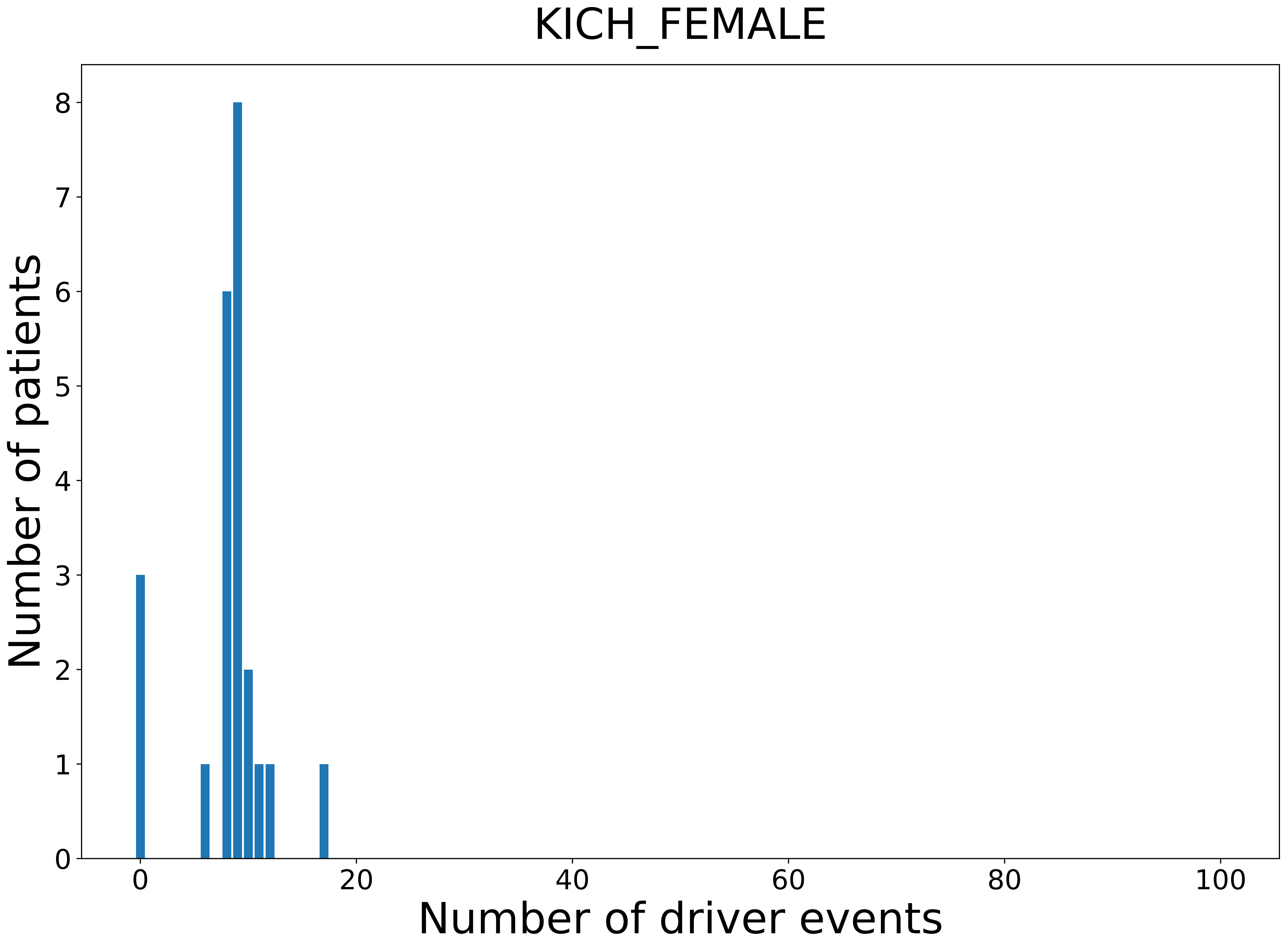

### 2021_8_22_13_29_KICH_MALE.pdf

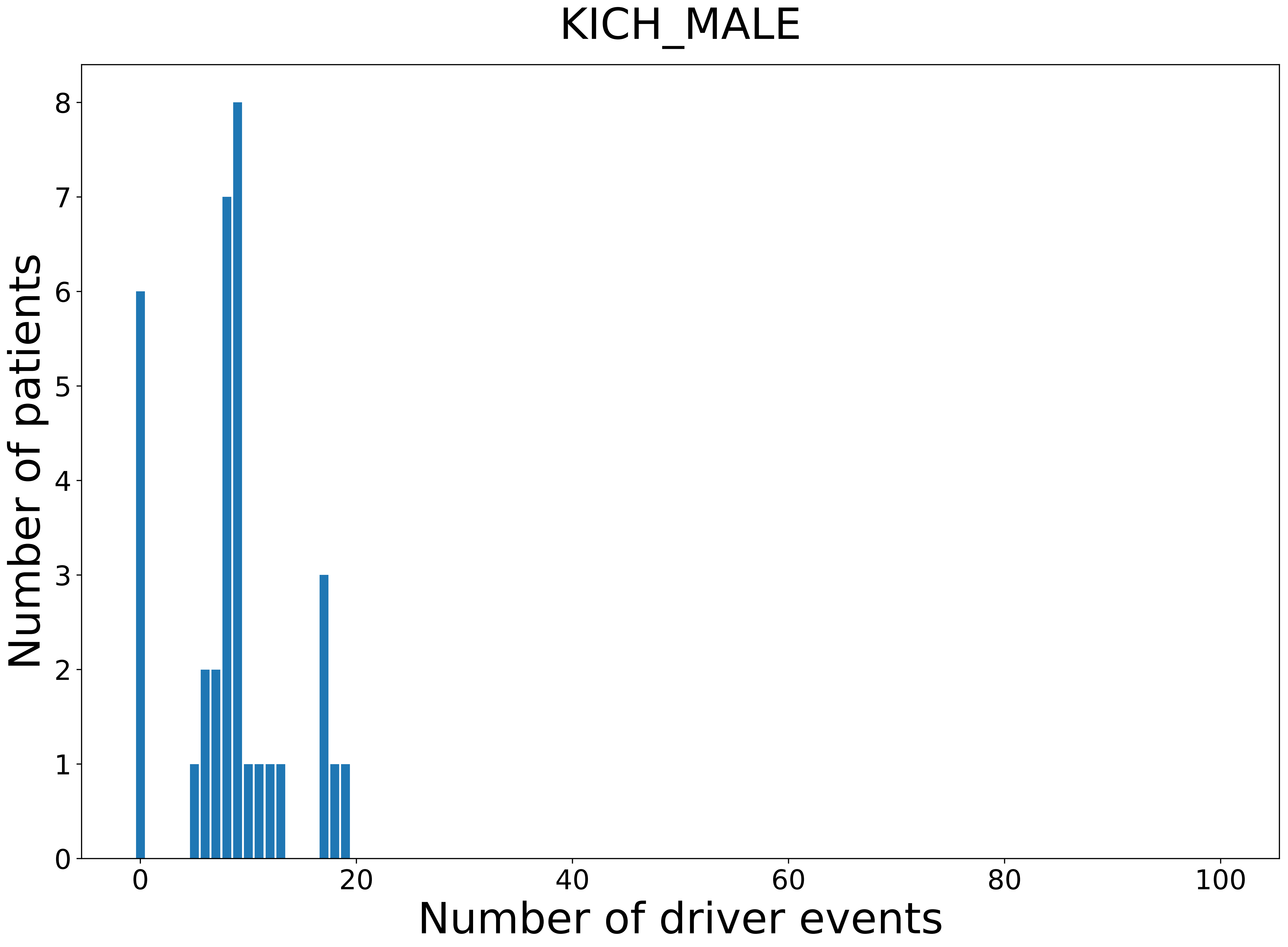

### 2021_8_22_13_29_KIRC.pdf

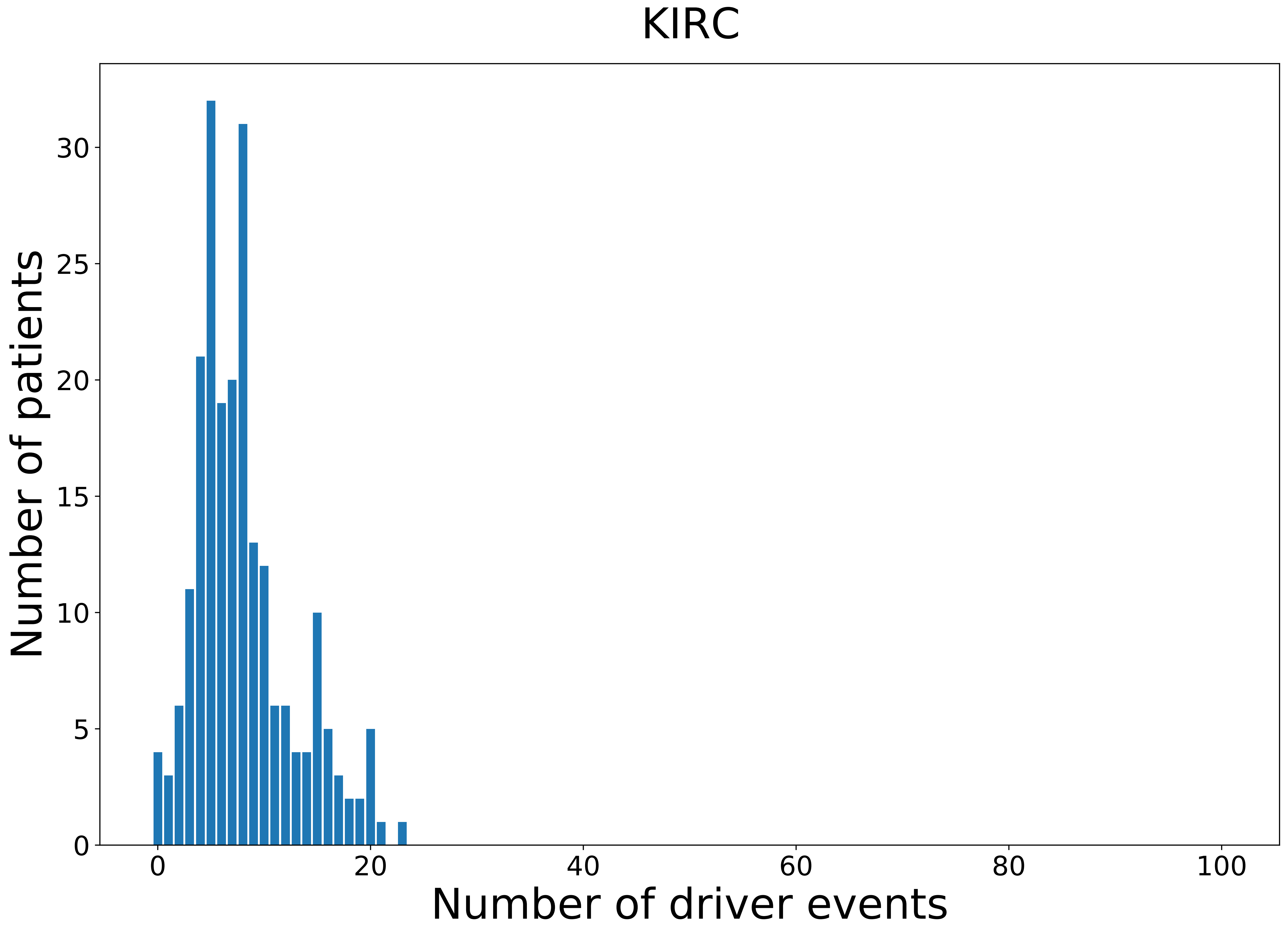

### 2021_8_22_13_29_KIRC_FEMALE.pdf

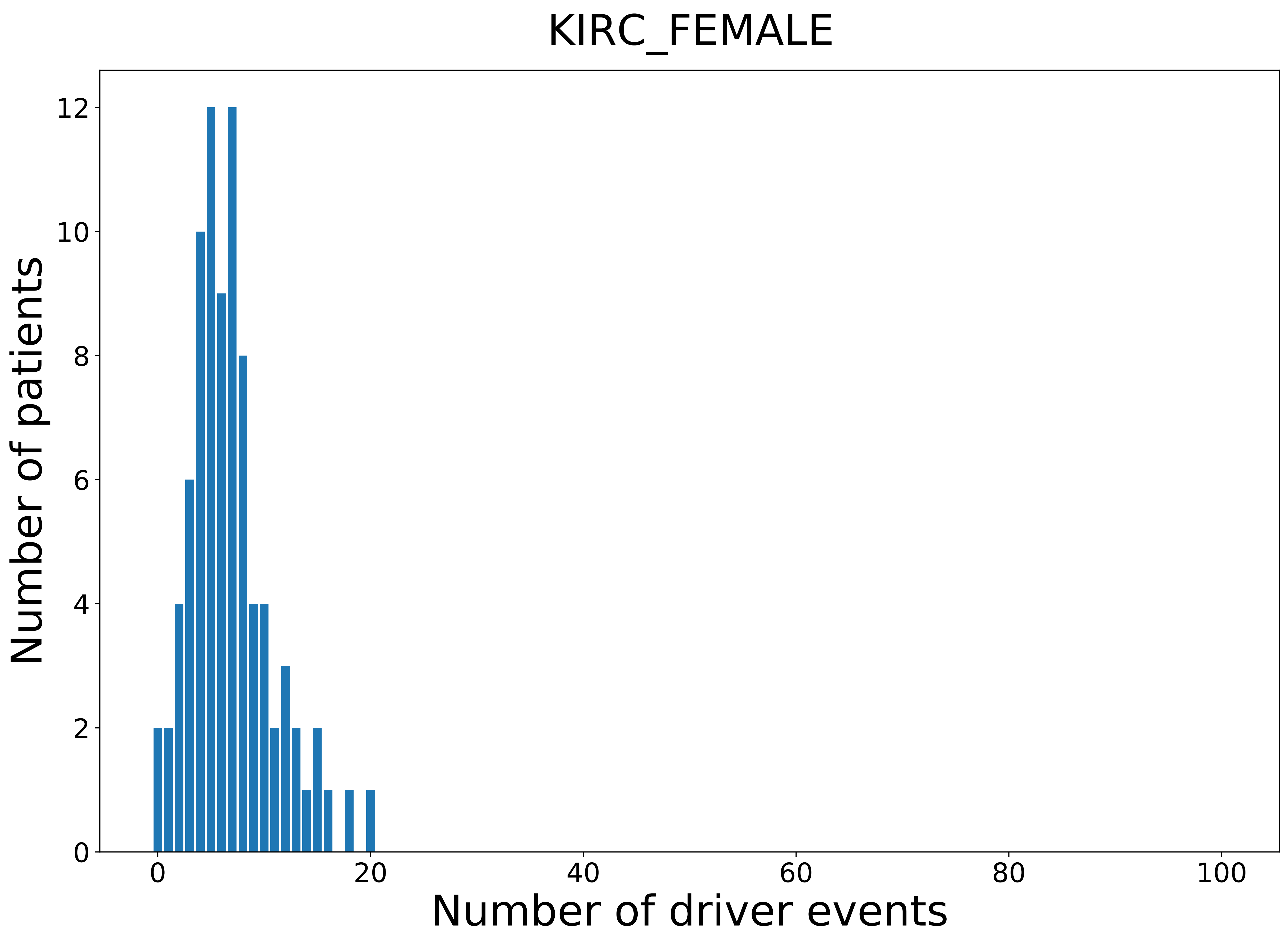

### 2021_8_22_13_29_KIRP.pdf

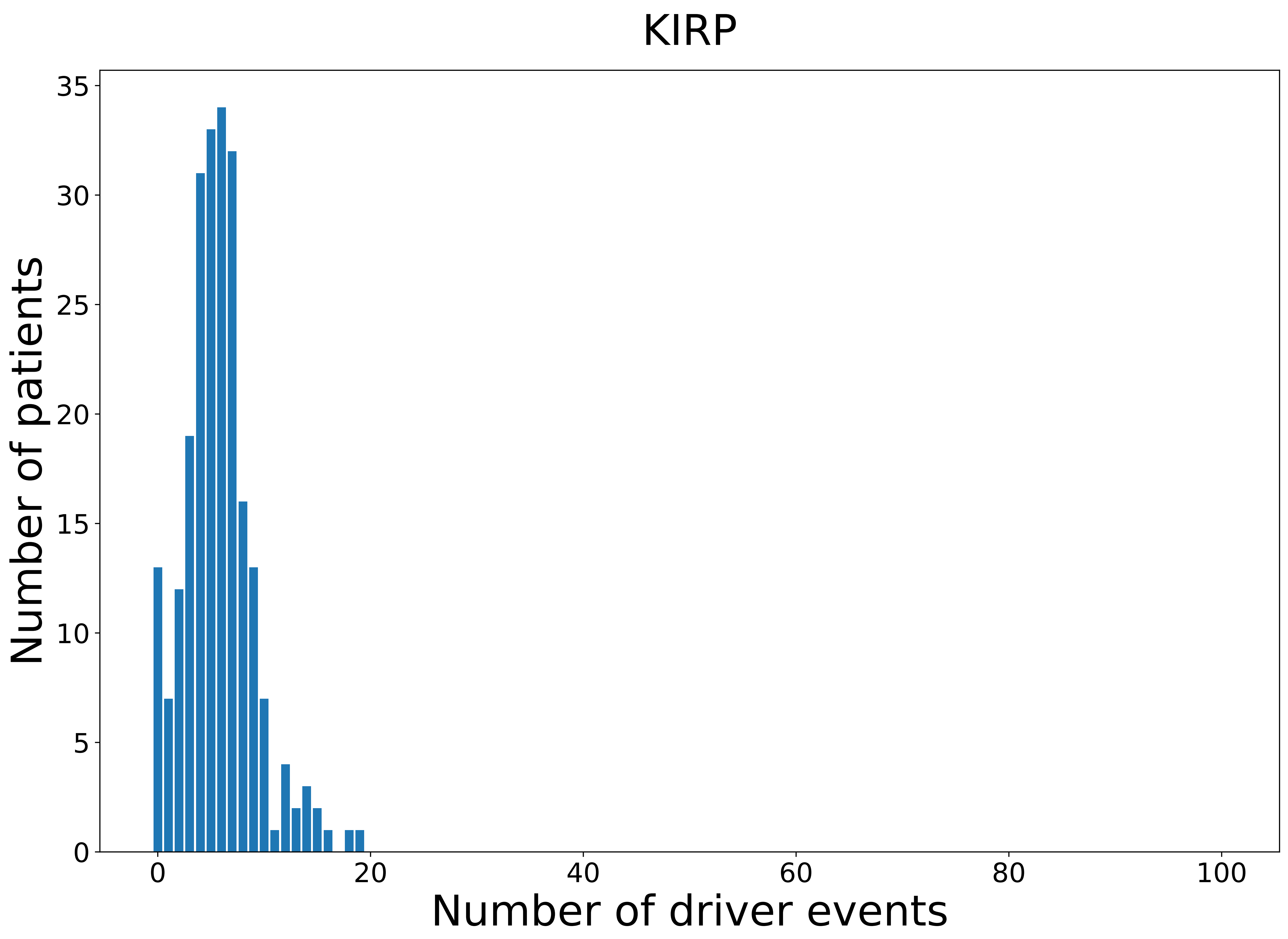

### 2021_8_22_13_29_KIRP_FEMALE.pdf

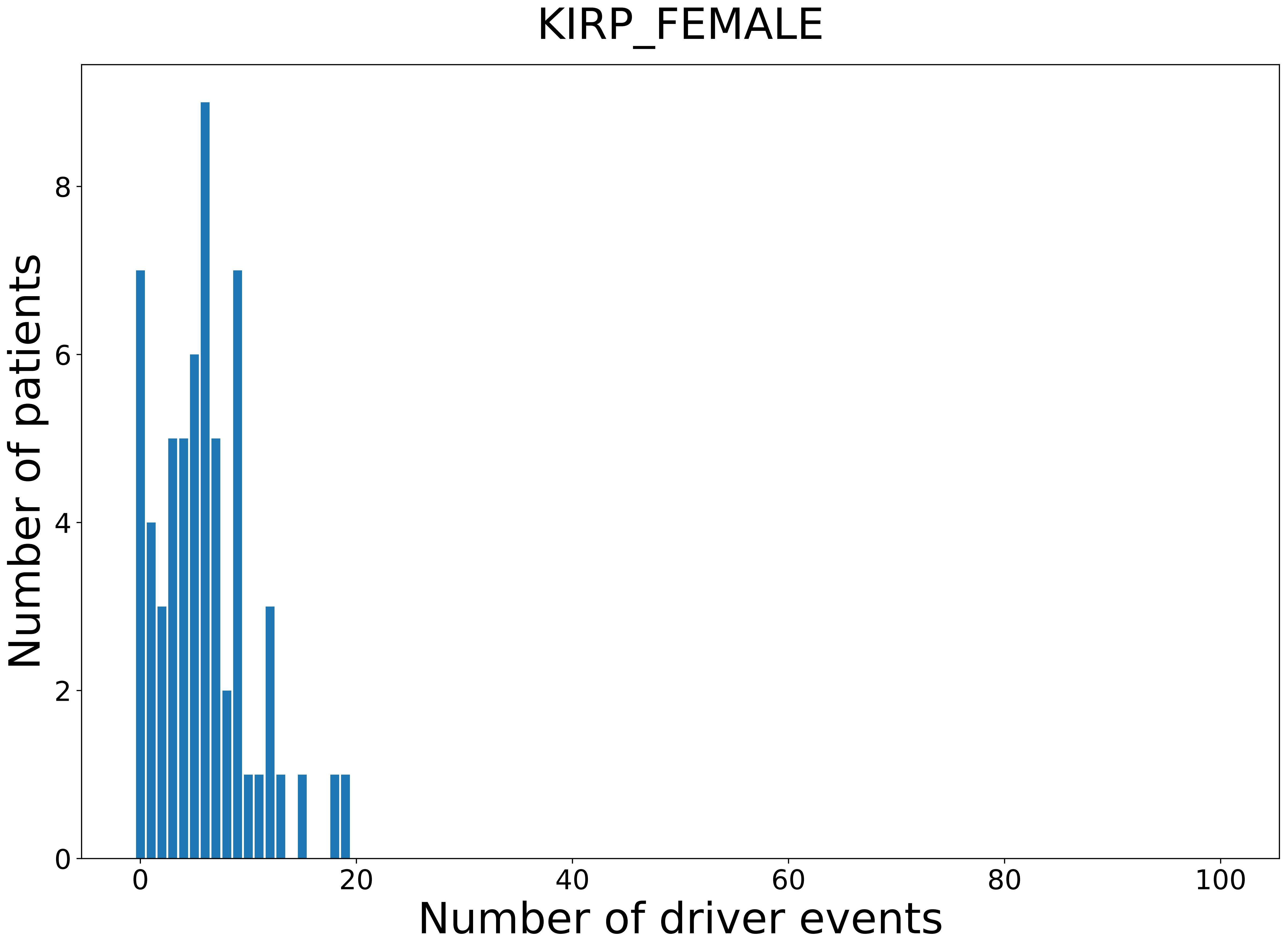

### 2021_8_22_13_29_KIRP_MALE.pdf

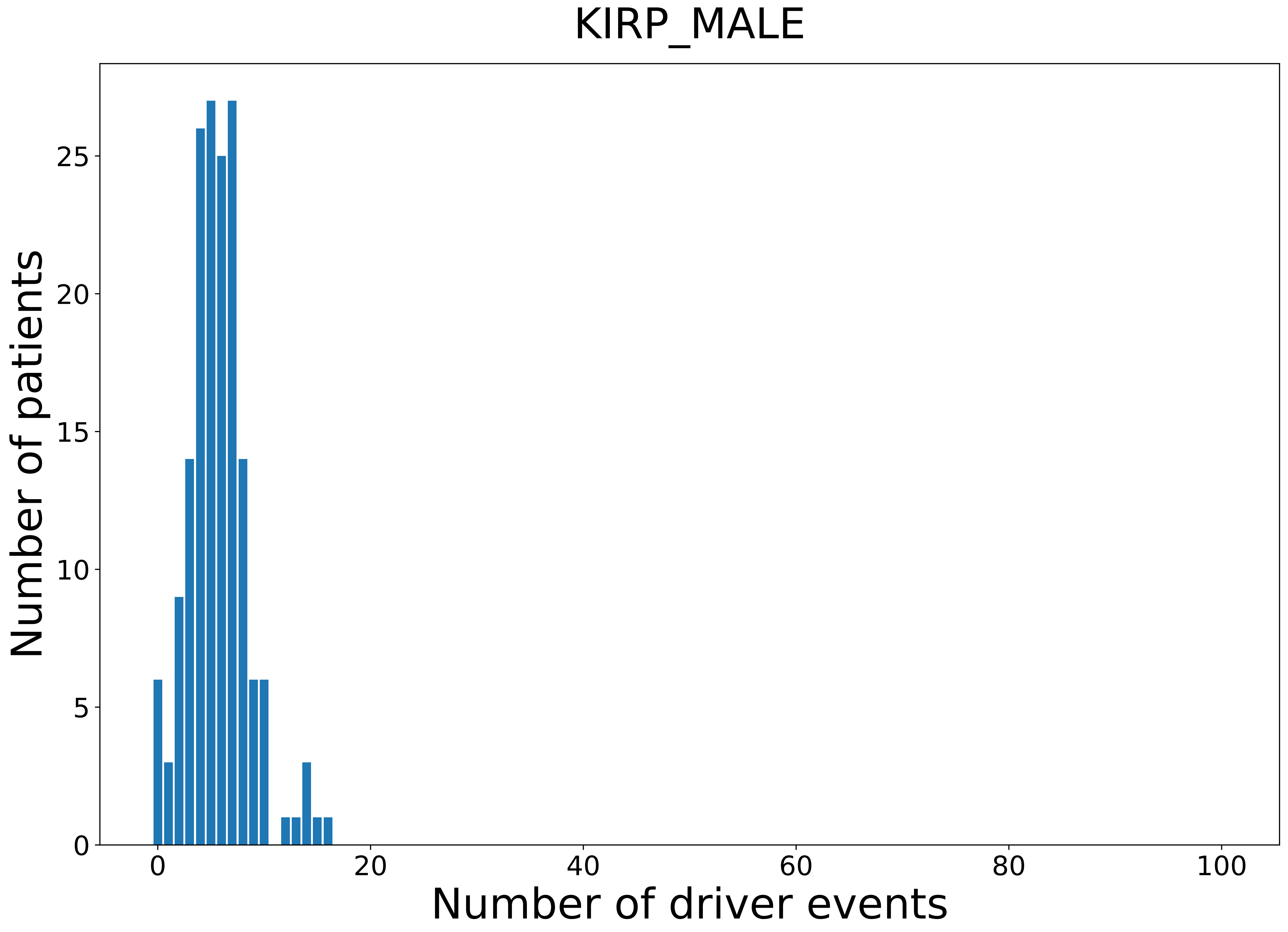

### 2021_8_22_13_29_LGG_FEMALE.pdf

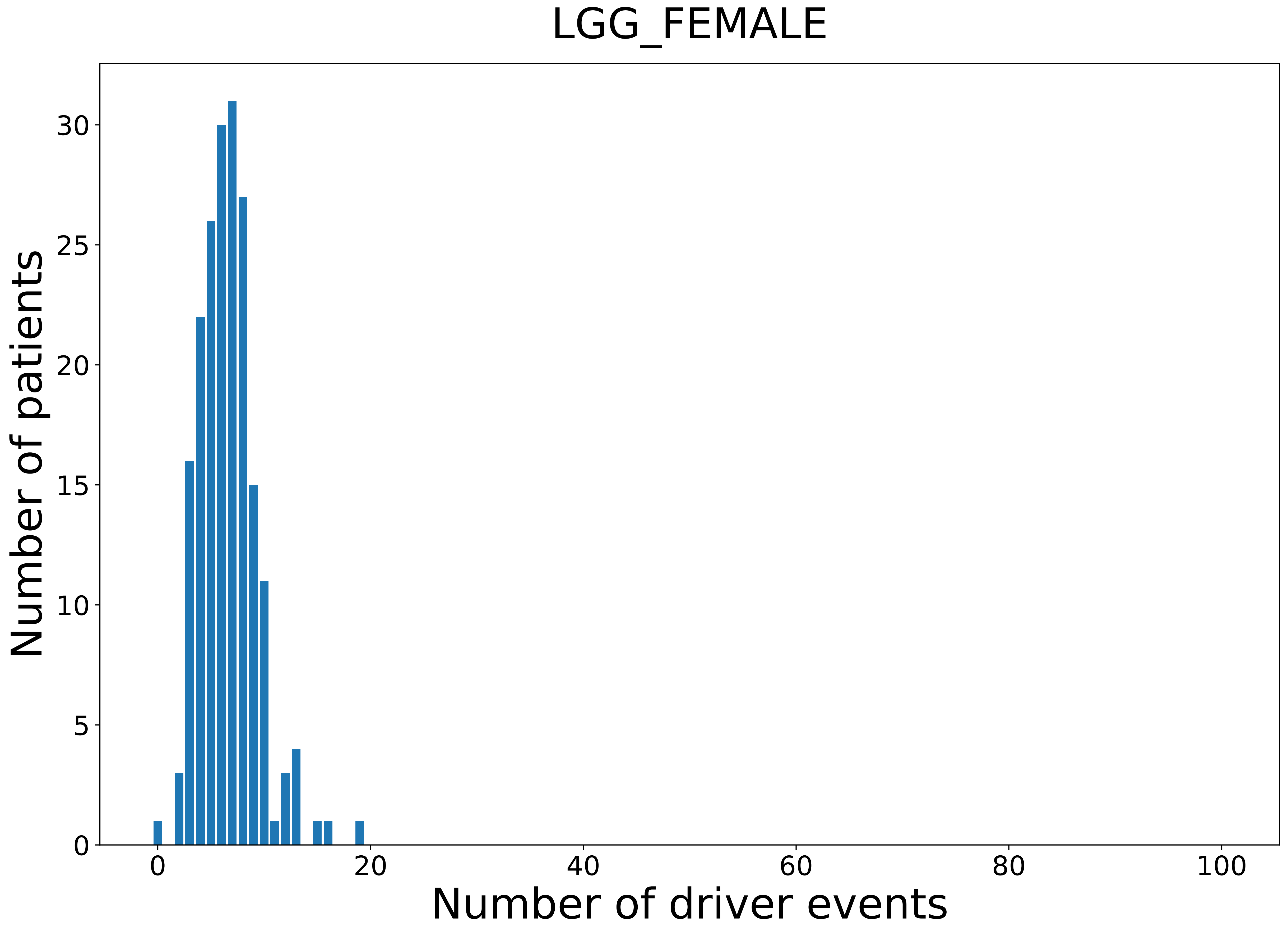

### 2021_8_22_13_29_LGG_MALE.pdf

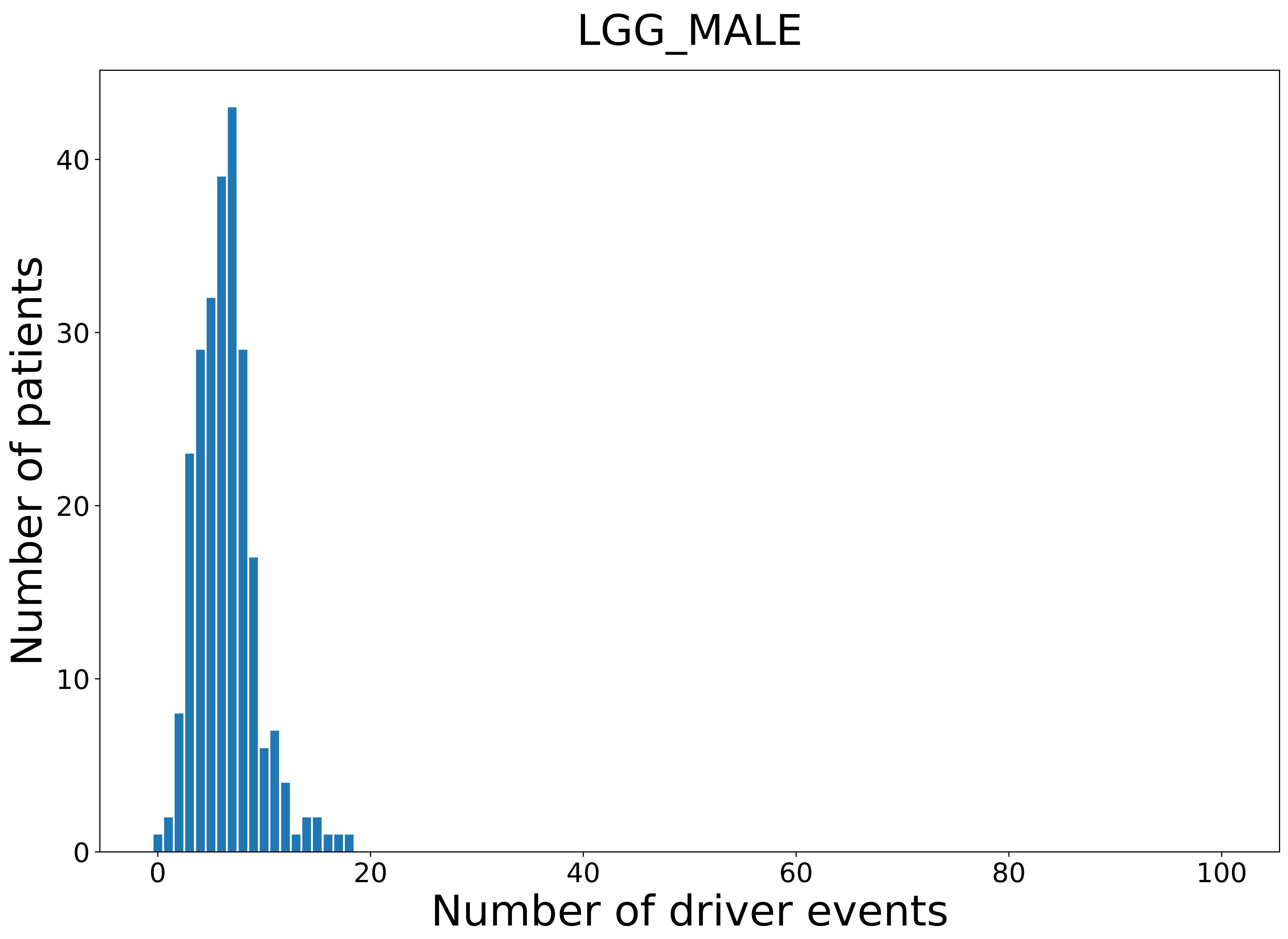

### 2021_8_22_13_29_LIHC.pdf

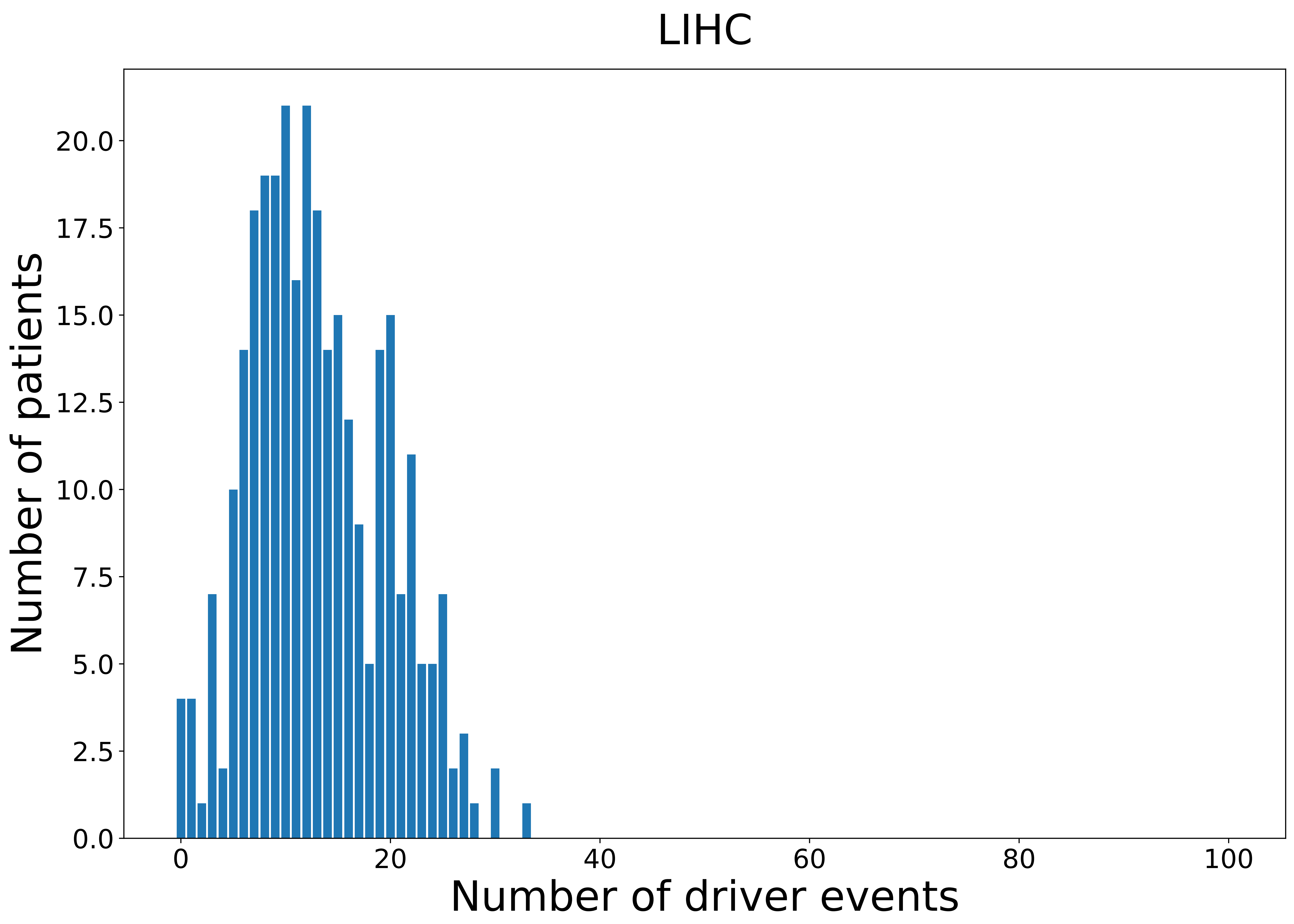

### 2021_8_22_13_29_LIHC_FEMALE.pdf

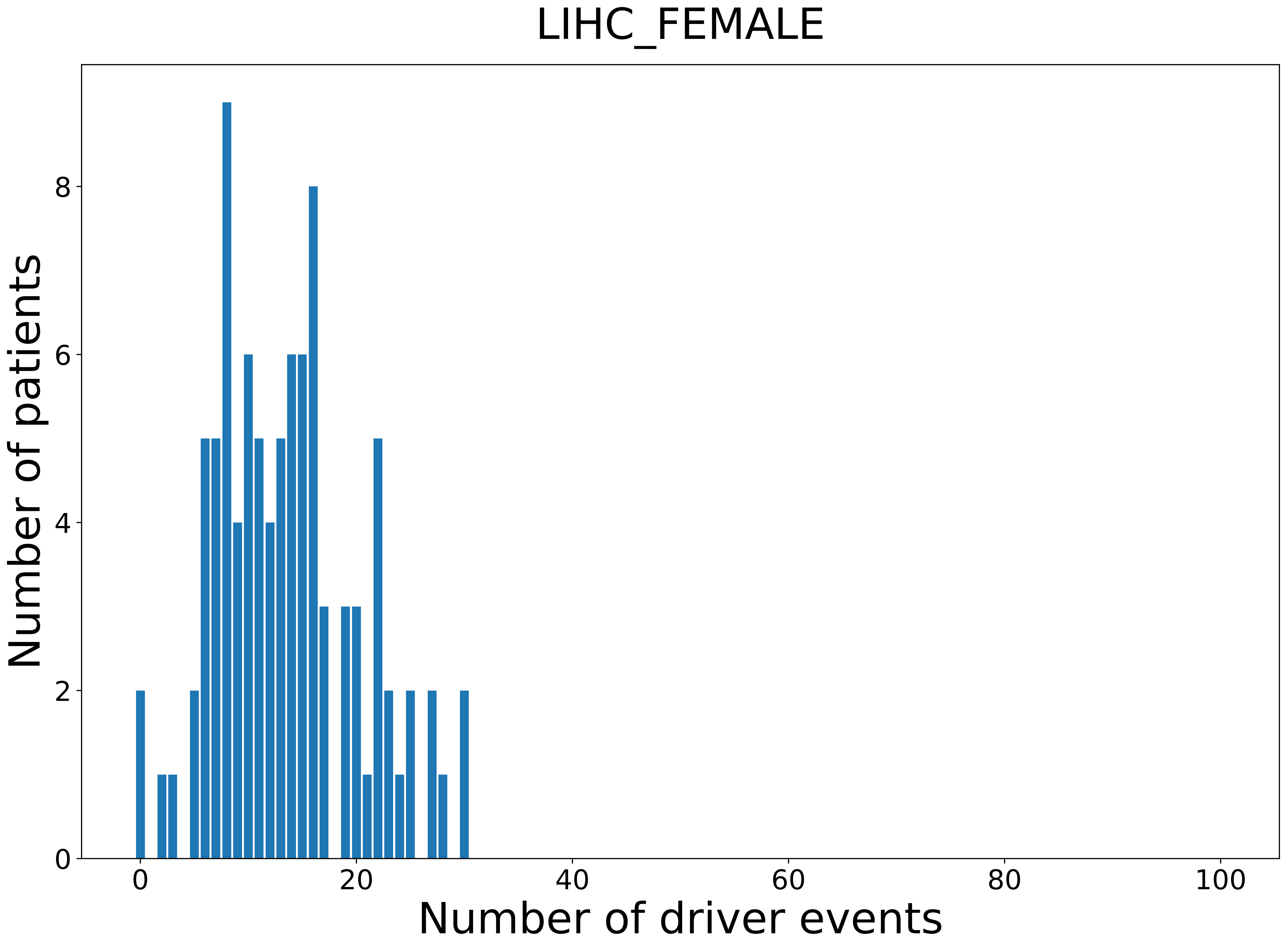

### 2021_8_22_13_29_LIHC_MALE.pdf

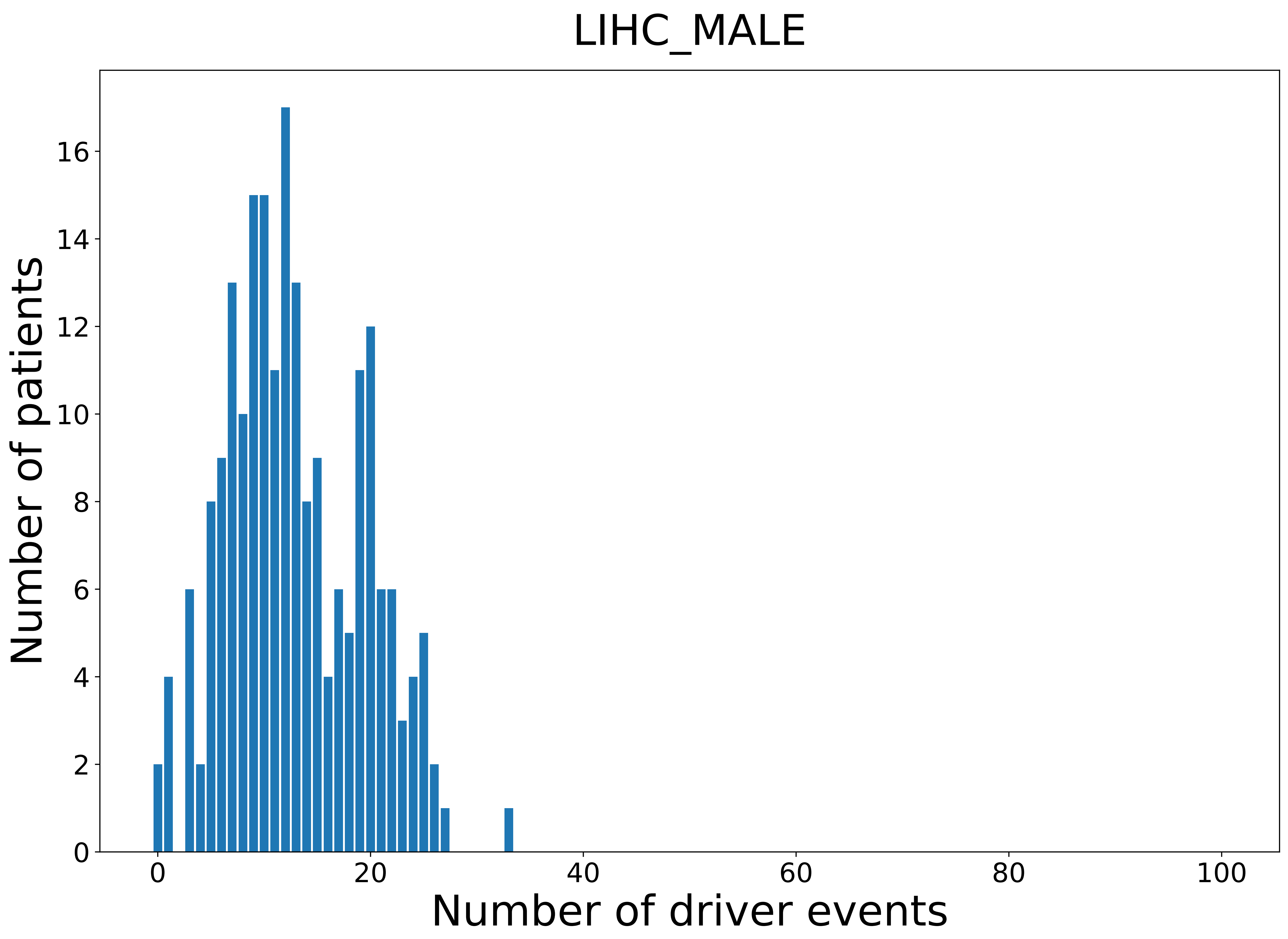

### 2021_8_22_13_29_LUAD.pdf

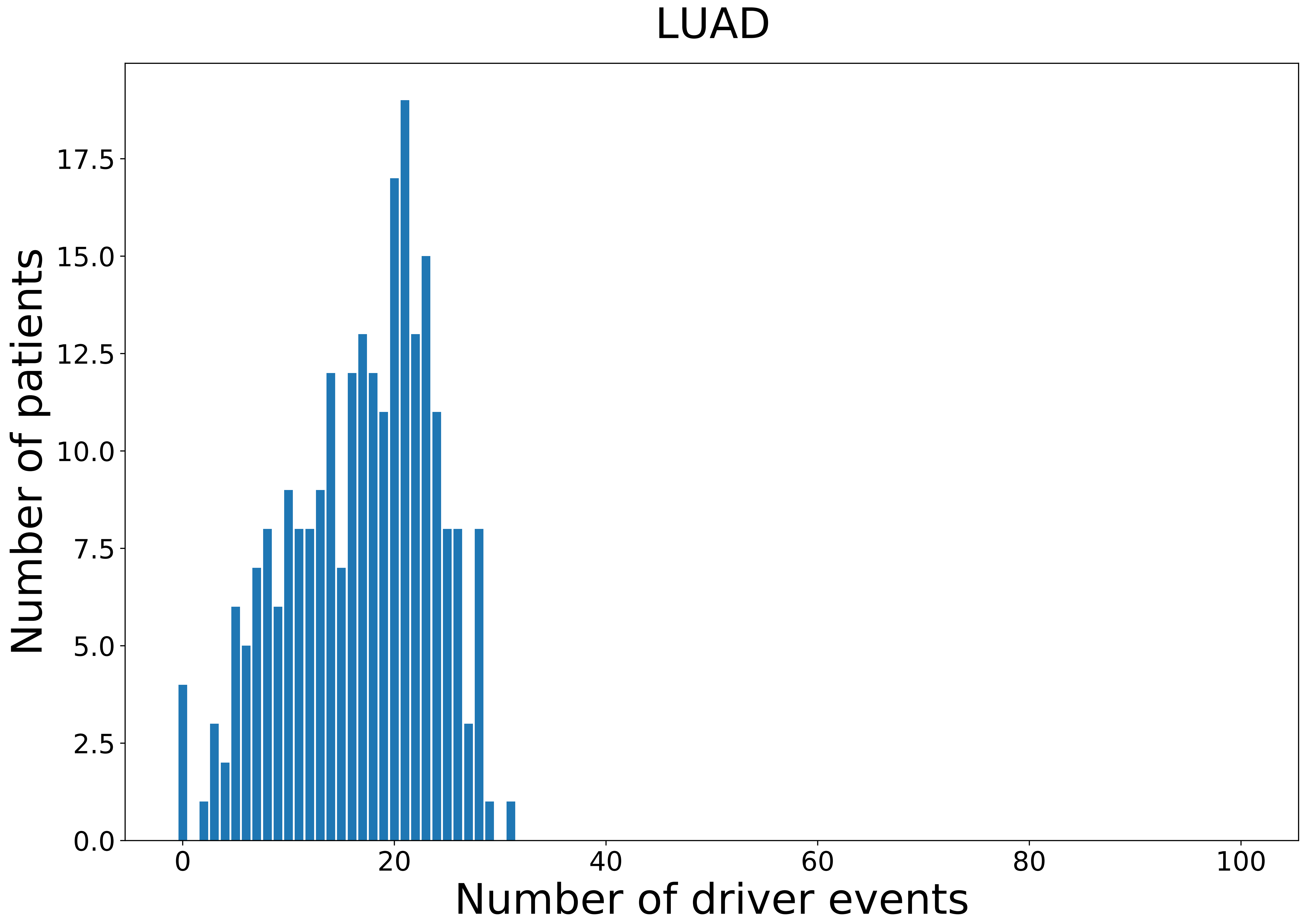

### 2021_8_22_13_29_LUAD_FEMALE.pdf

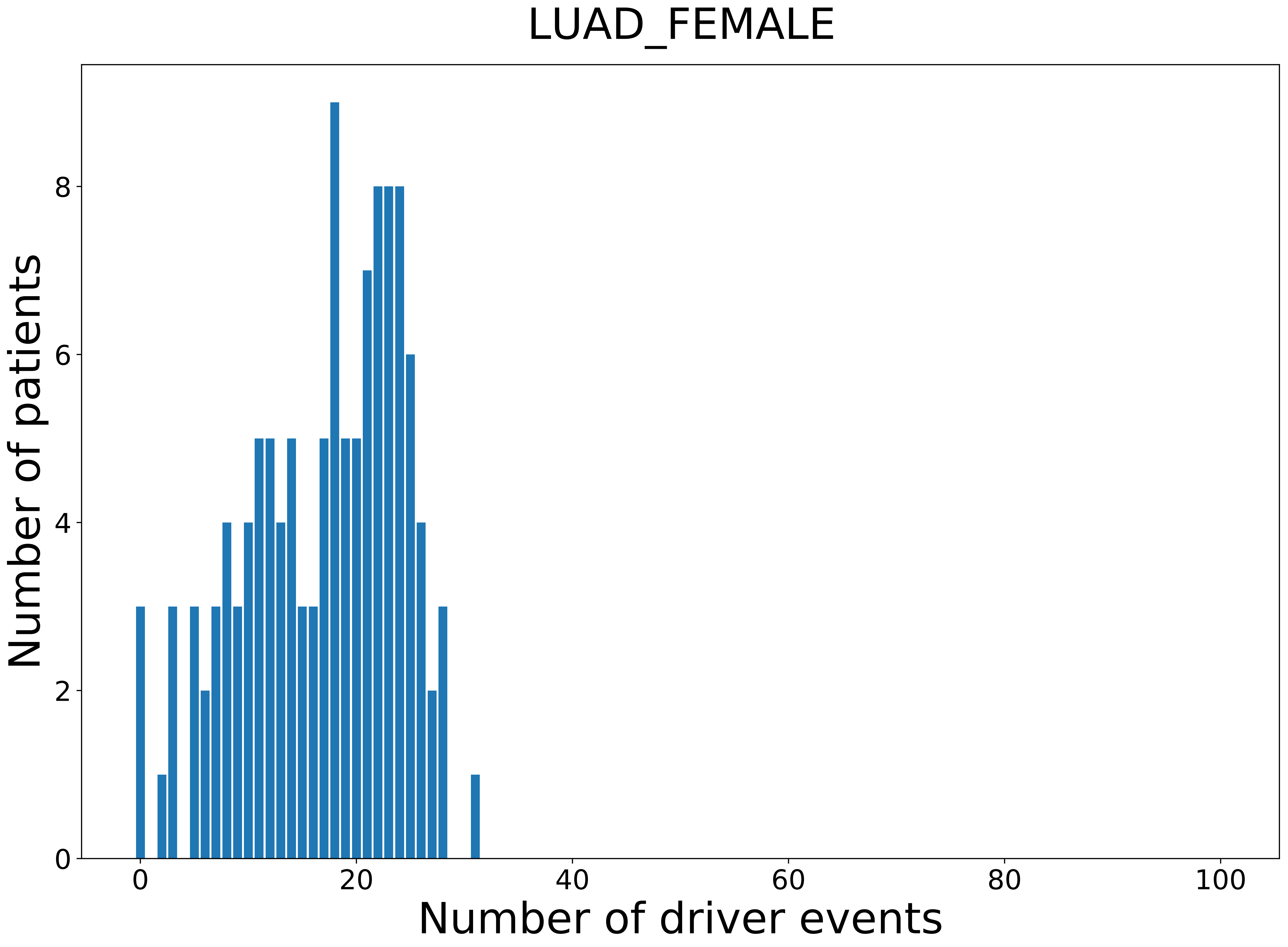

### 2021_8_22_13_29_LUAD_MALE.pdf

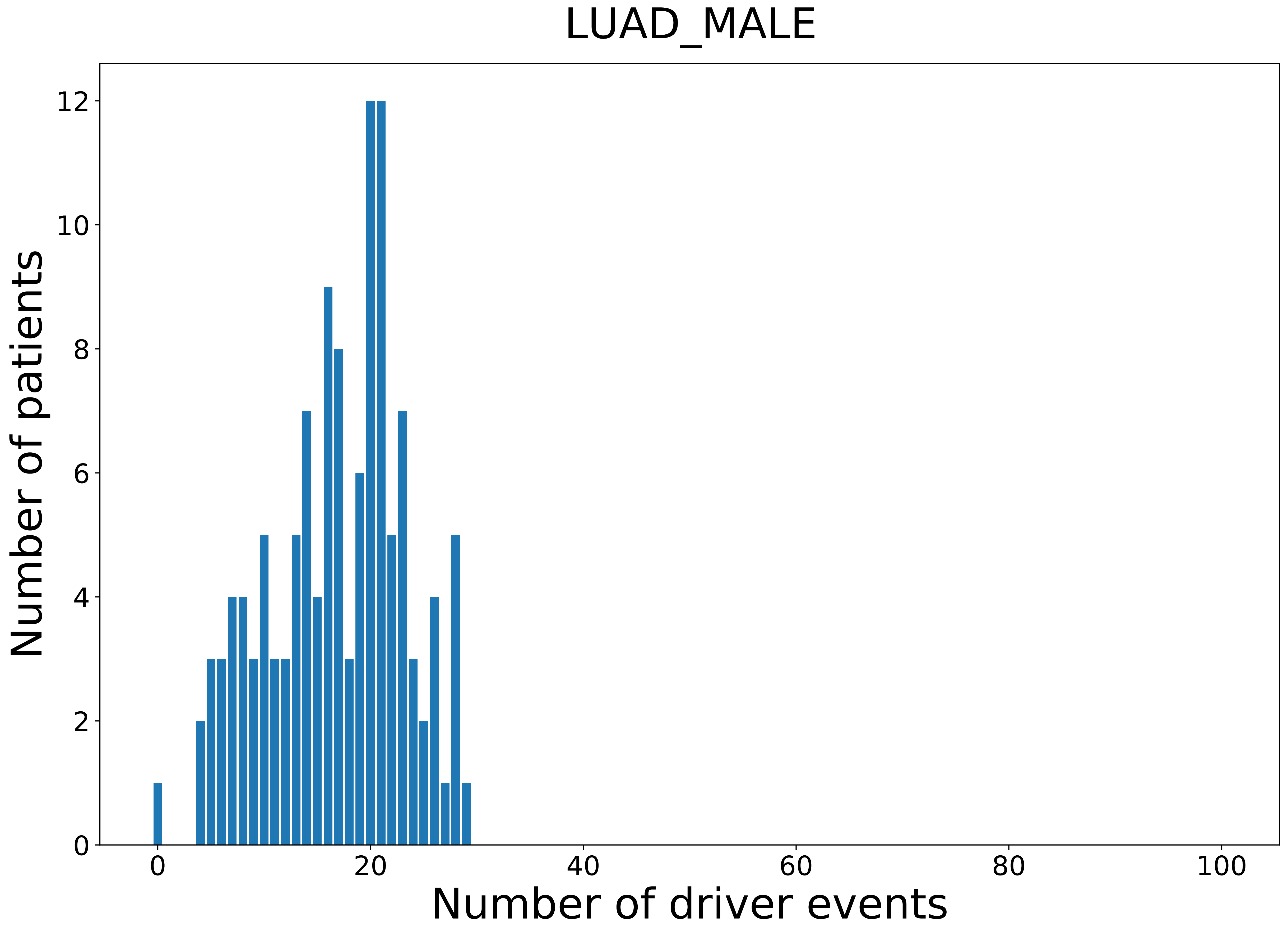

### 2021_8_22_13_29_LUSC_FEMALE.pdf

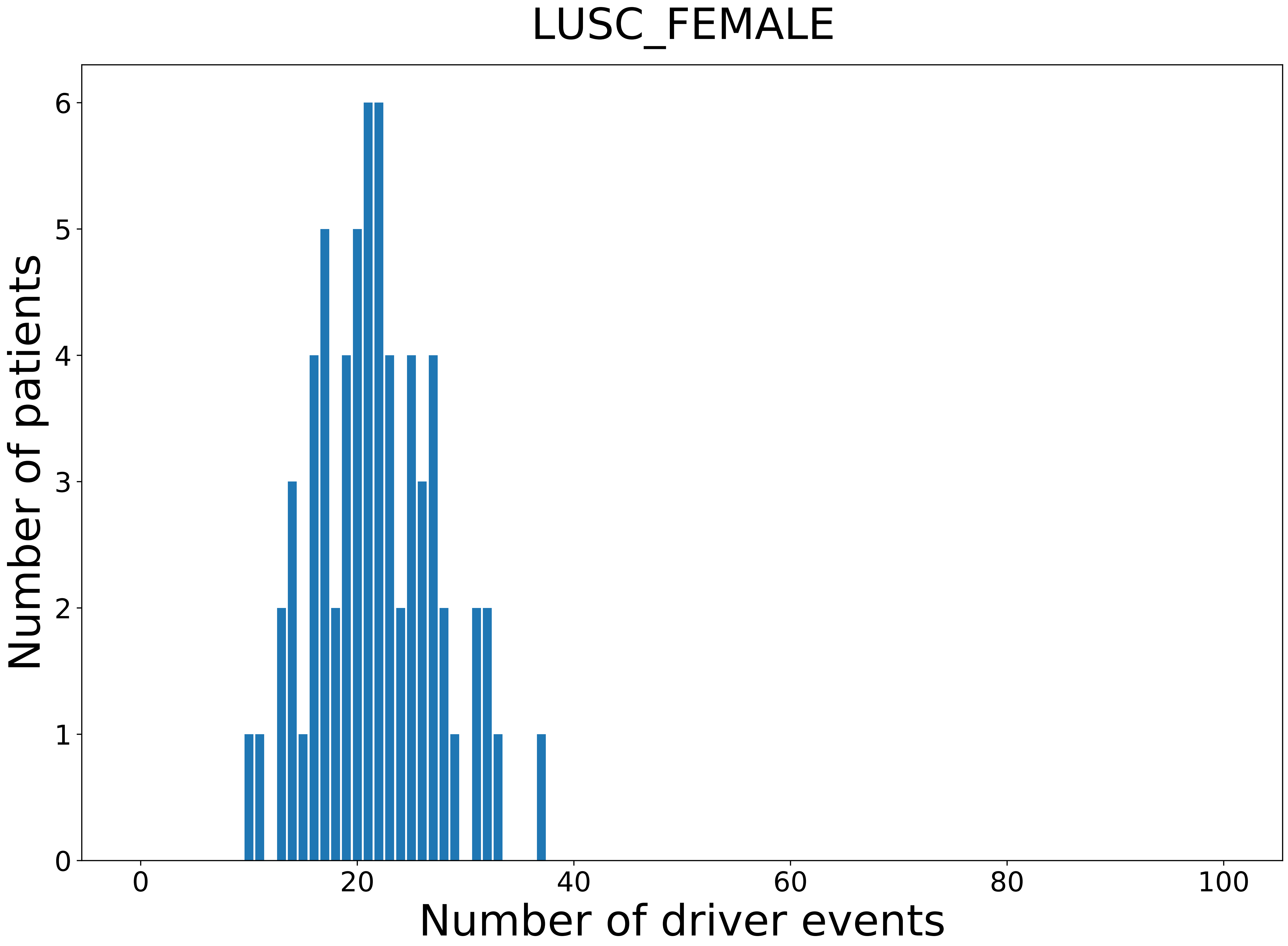

### 2021_8_22_13_29_LUSC_MALE.pdf

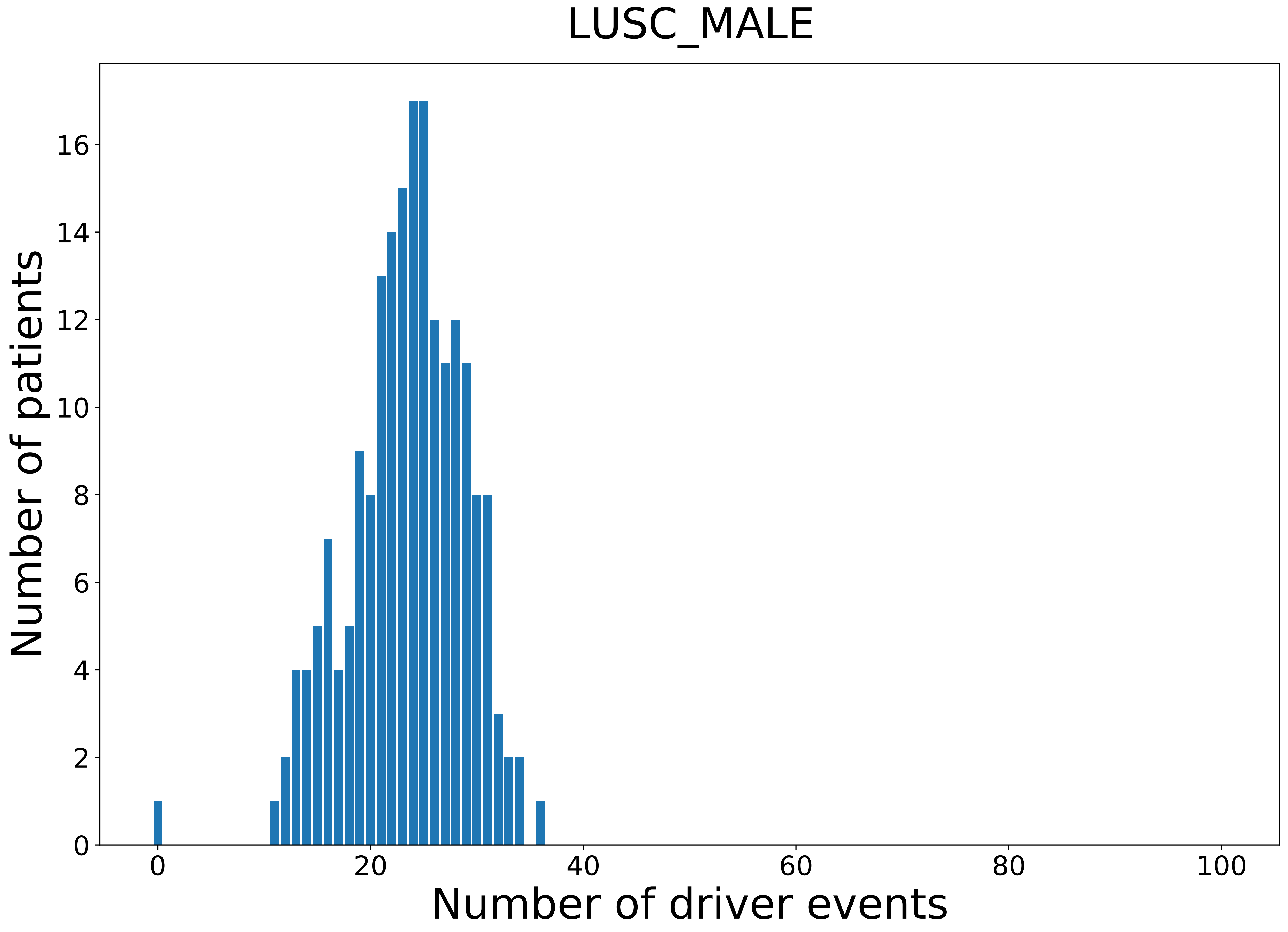

### 2021_8_22_13_29_MESO.pdf

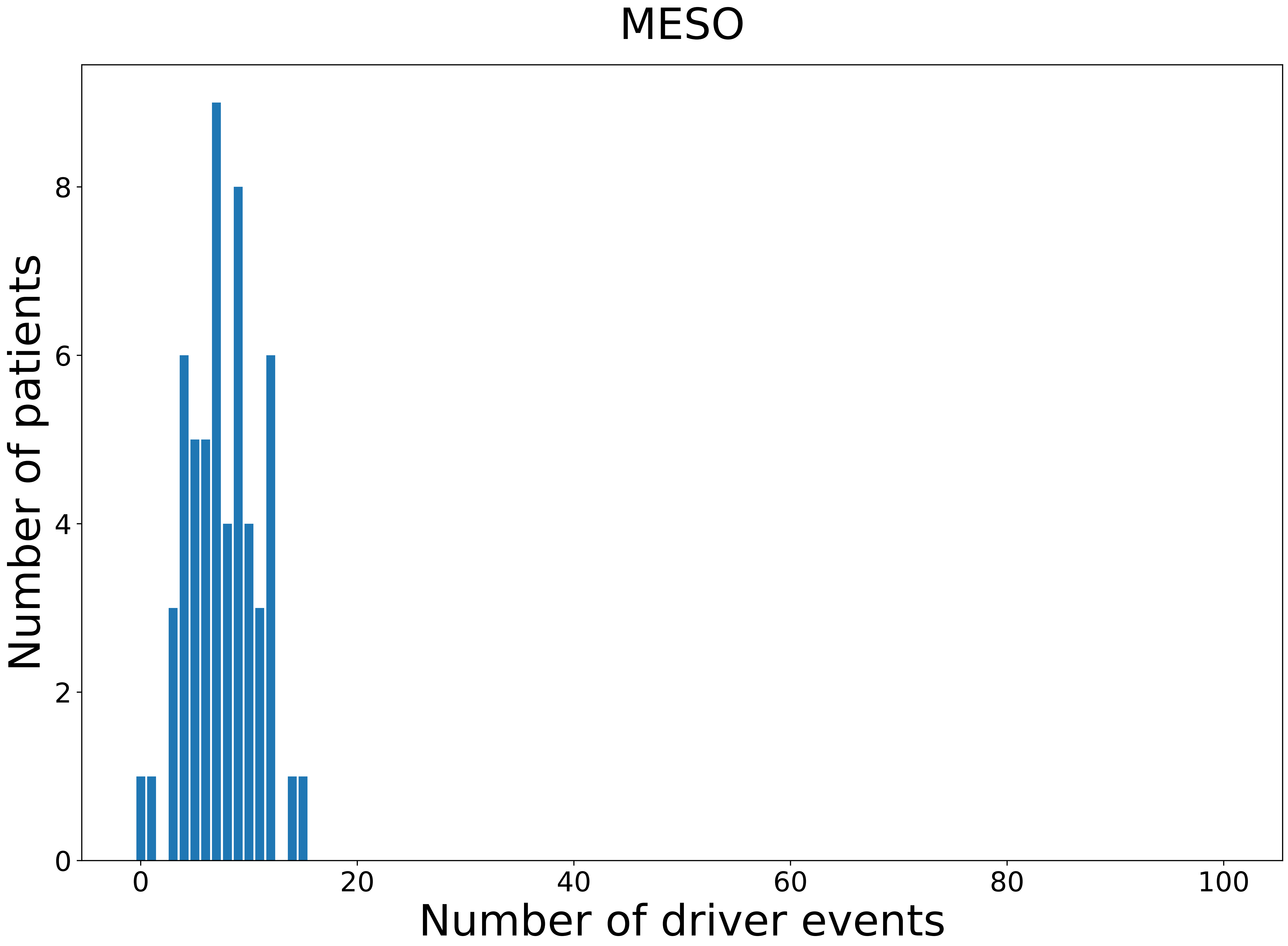

### 2021_8_22_13_29_MESO_FEMALE.pdf

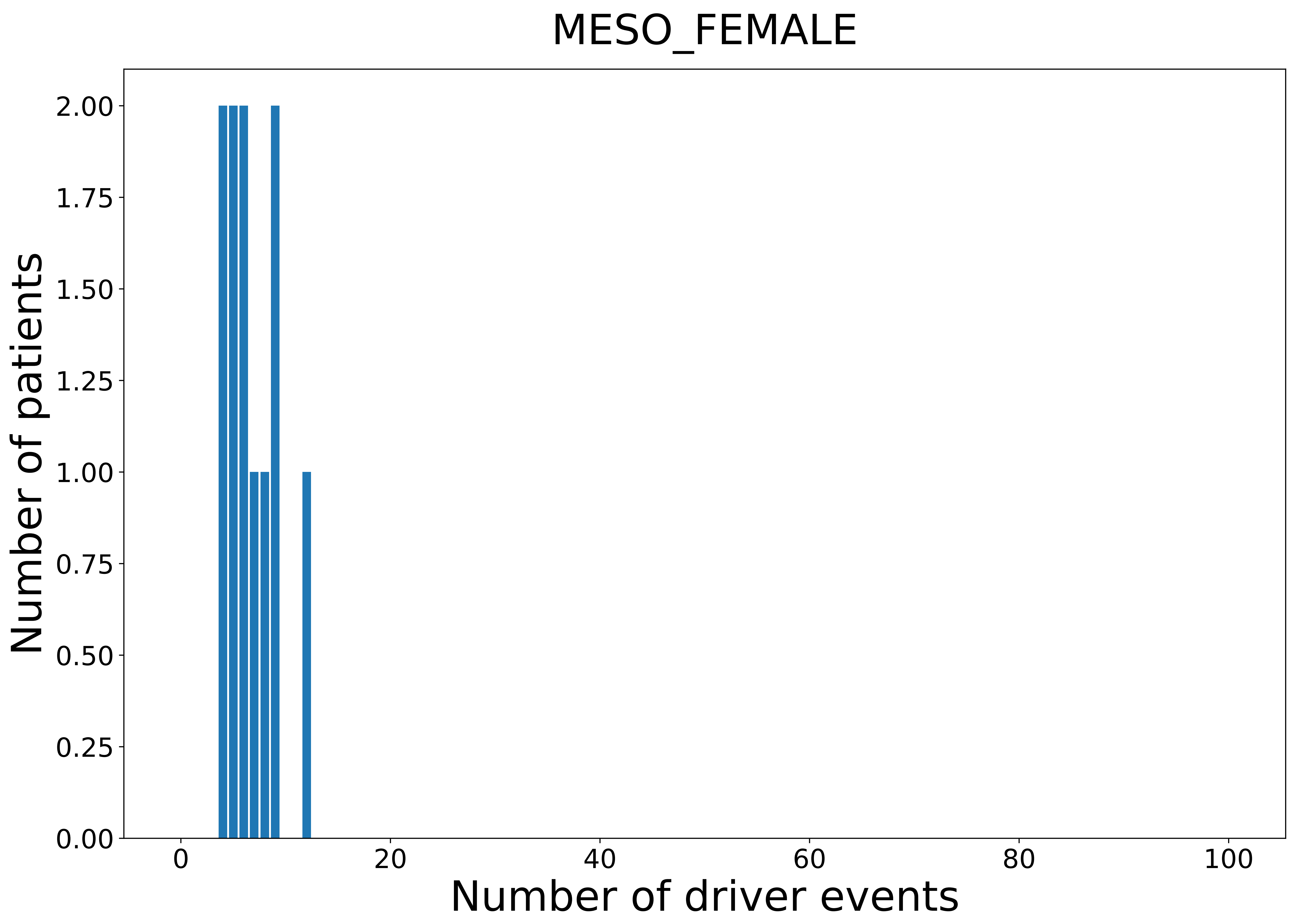

### 2021_8_22_13_29_MESO_MALE.pdf

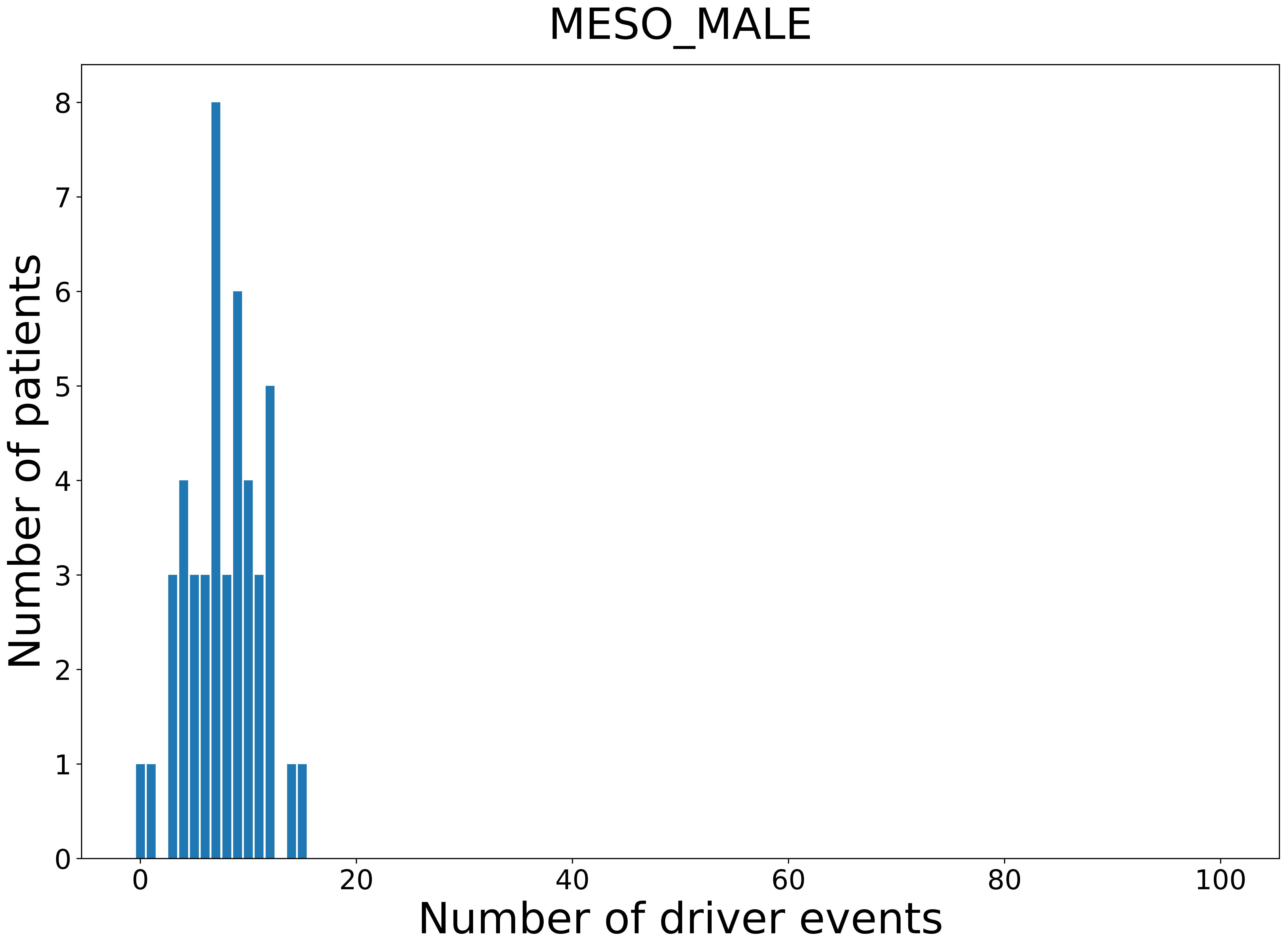

### 2021_8_22_13_29_OV.pdf

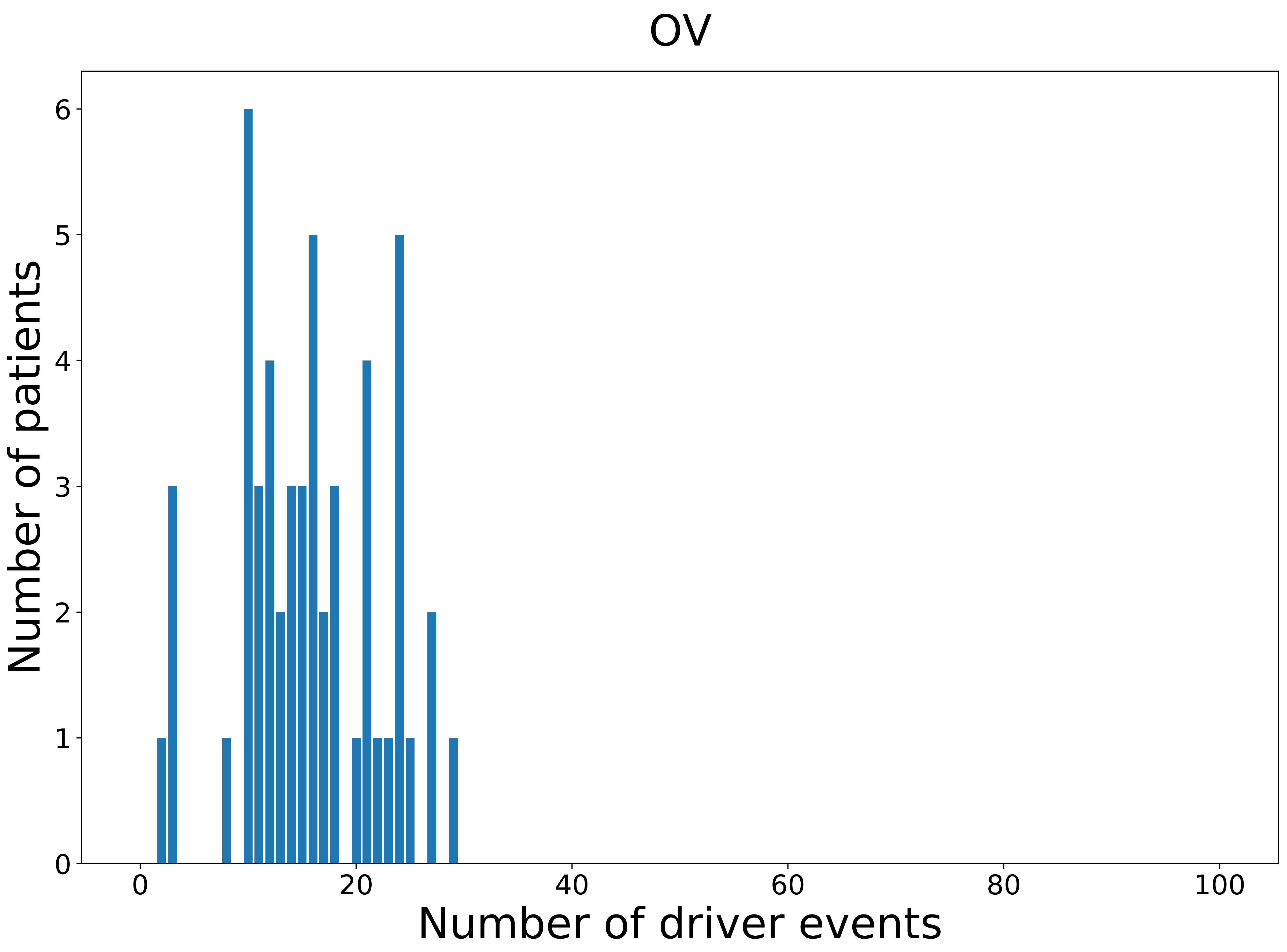

### 2021_8_22_13_29_OV_FEMALE.pdf

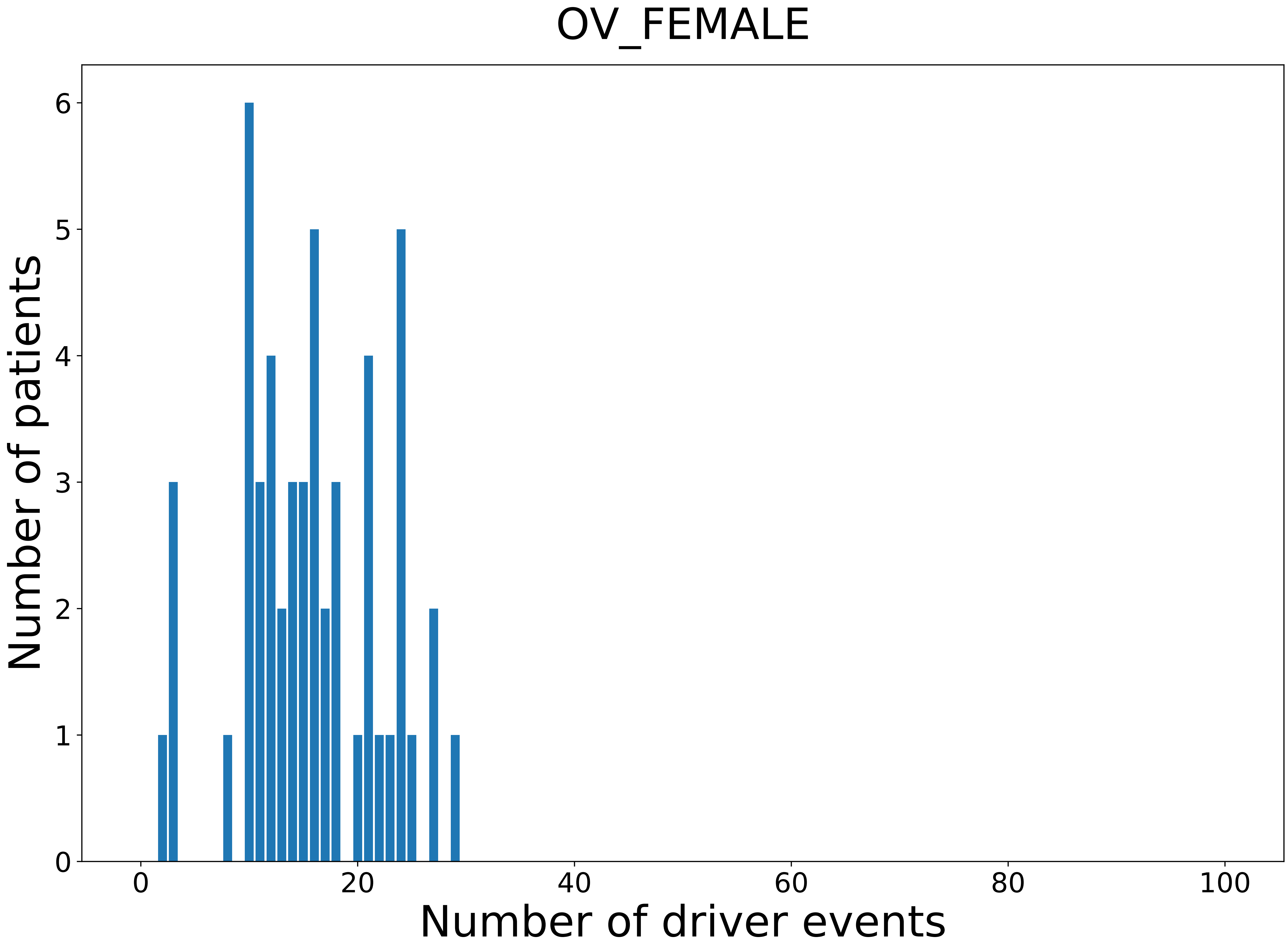

### 2021_8_22_13_29_PAAD.pdf

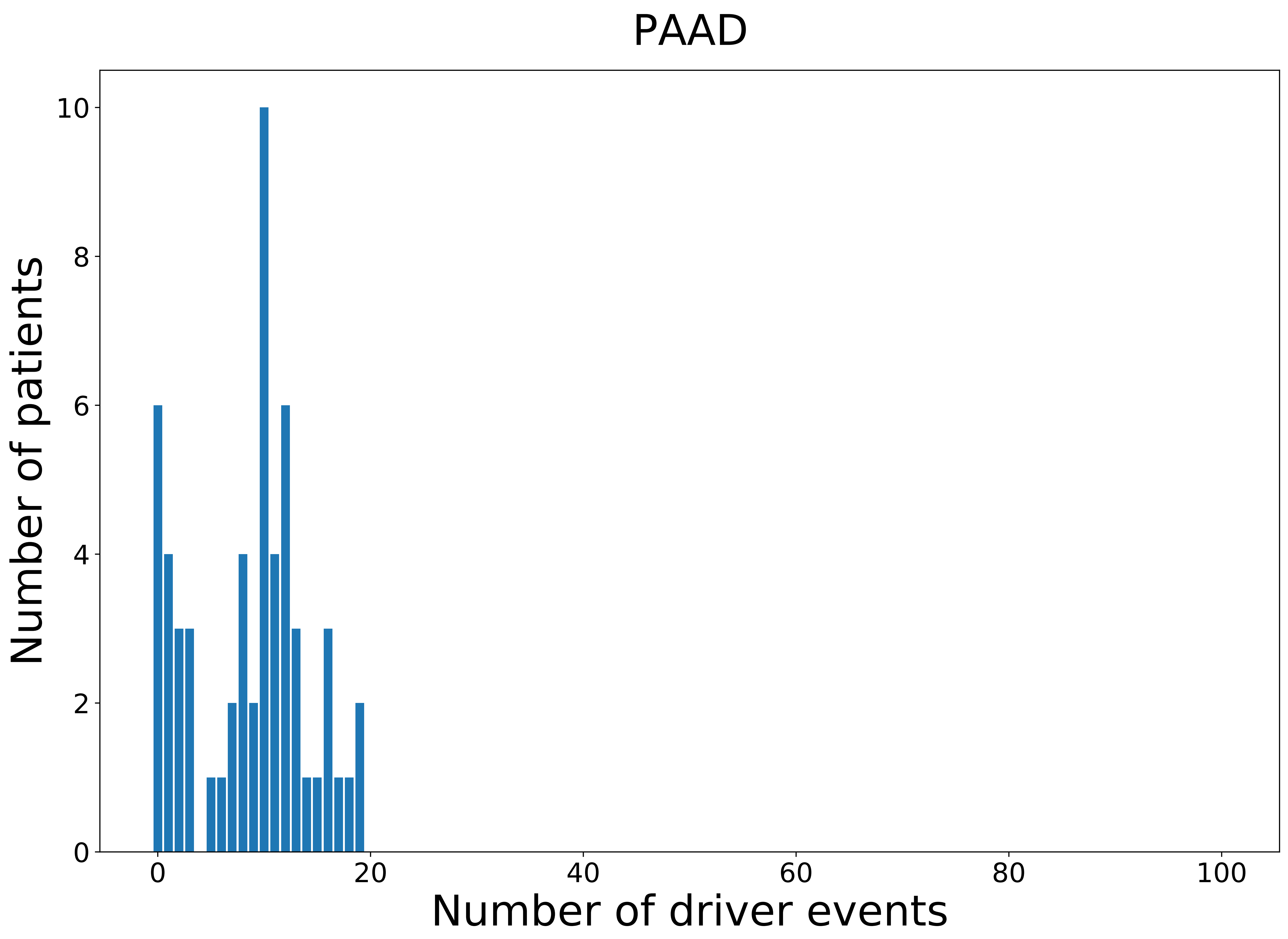

### 2021_8_22_13_29_PAAD_FEMALE.pdf

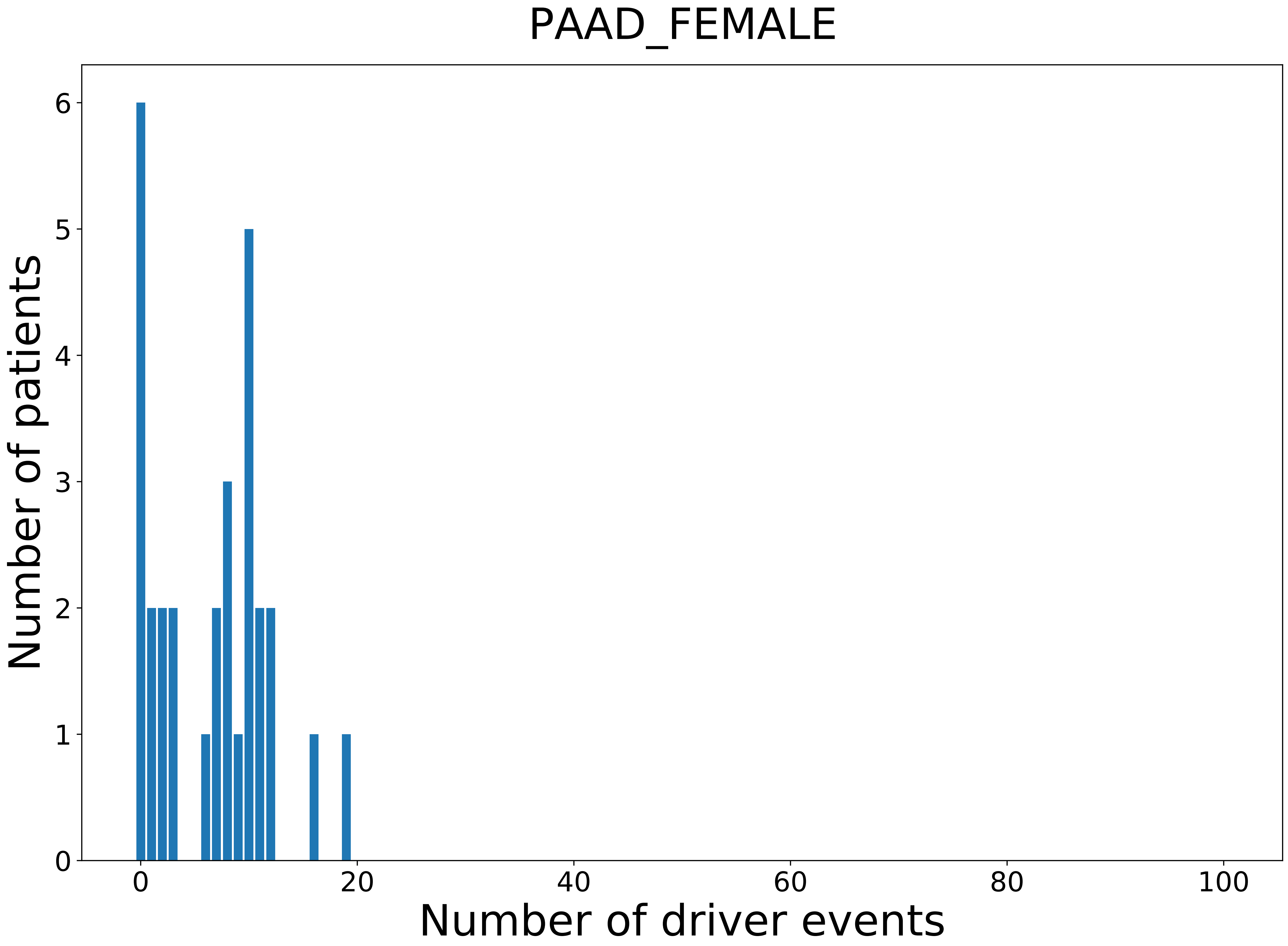

### 2021_8_22_13_29_PAAD_MALE.pdf

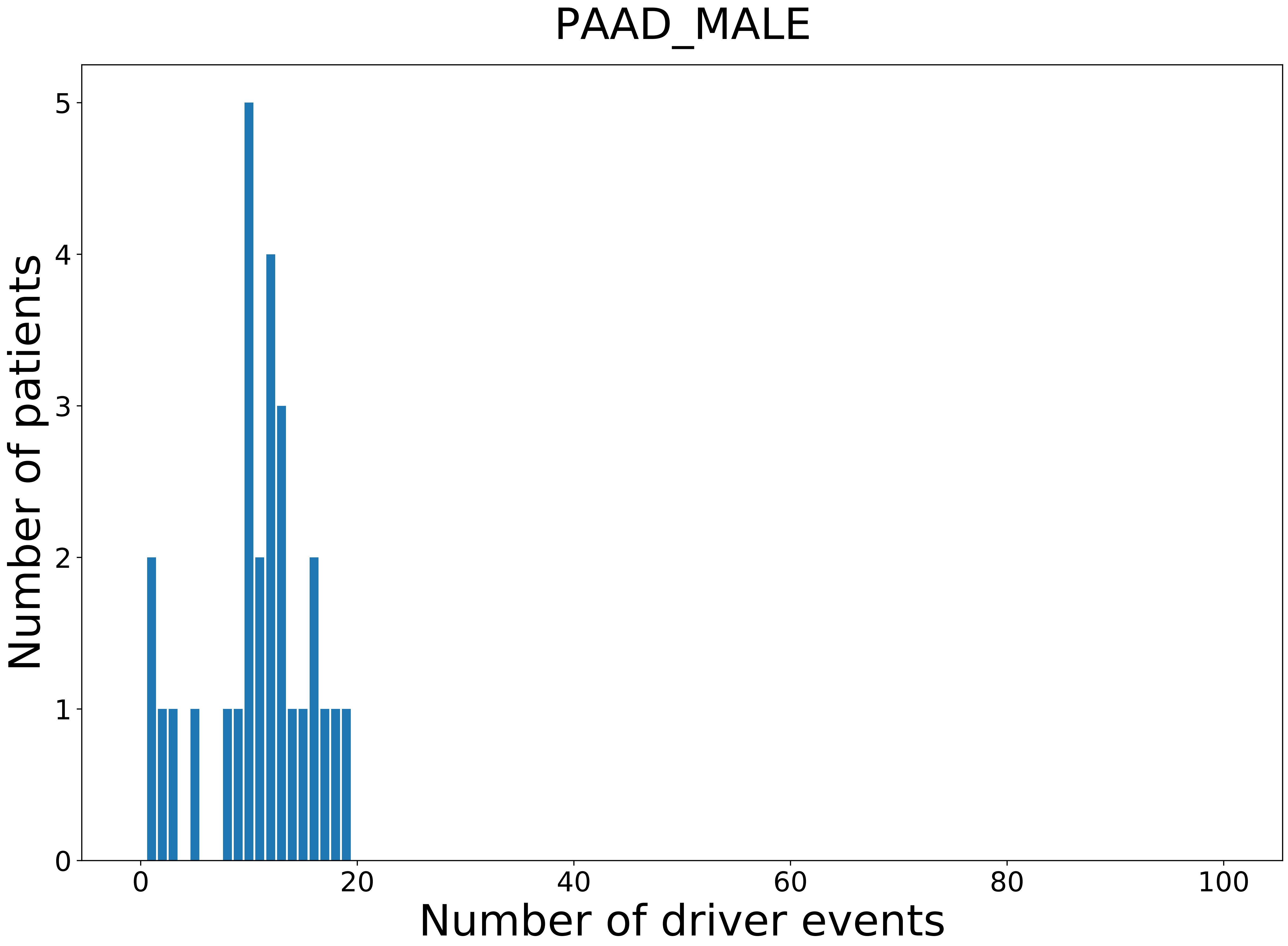

### 2021_8_22_13_29_PANCAN.pdf

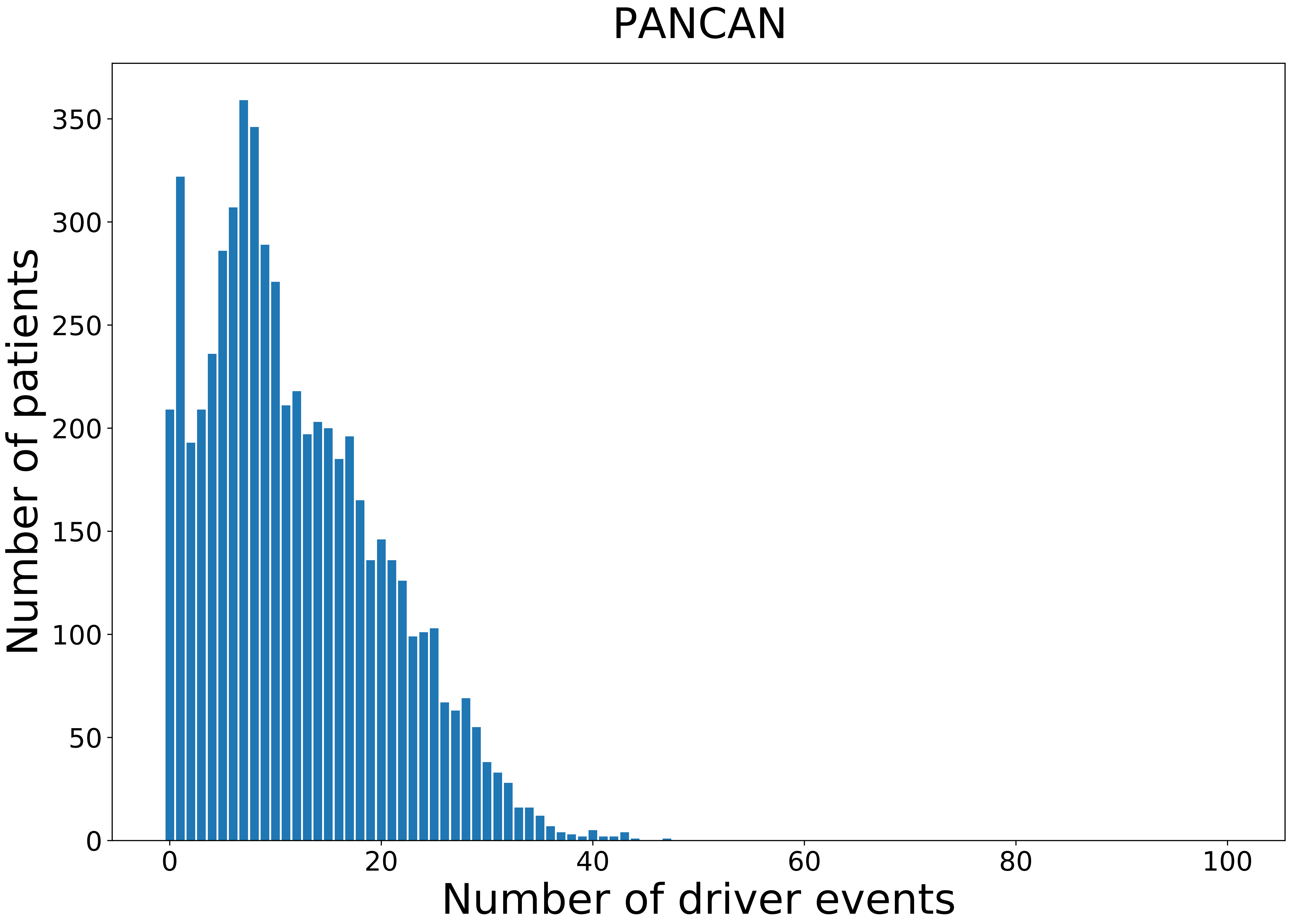

### 2021_8_22_13_29_PANCAN_FEMALE.pdf

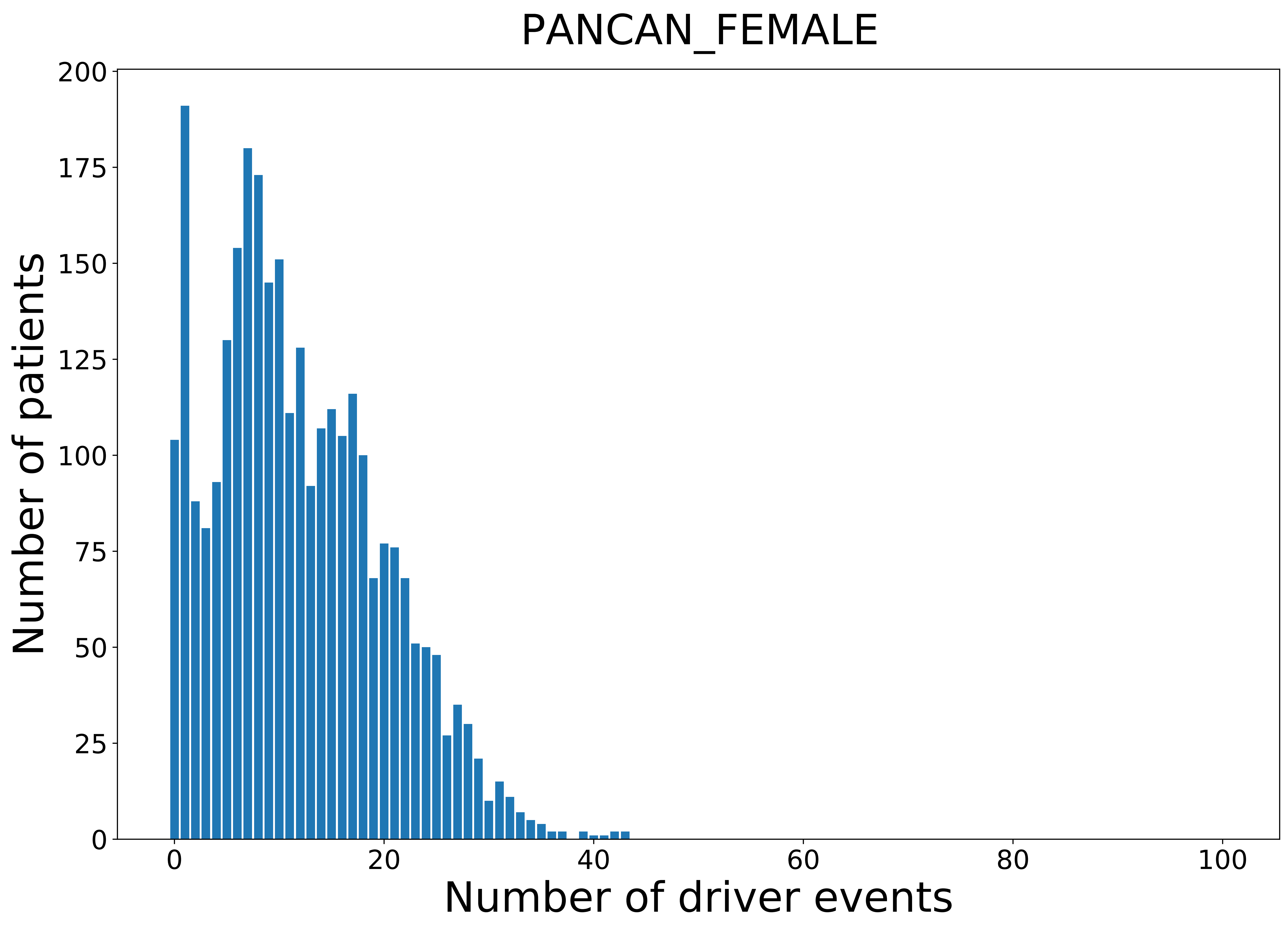

### 2021_8_22_13_29_PANCAN_MALE.pdf

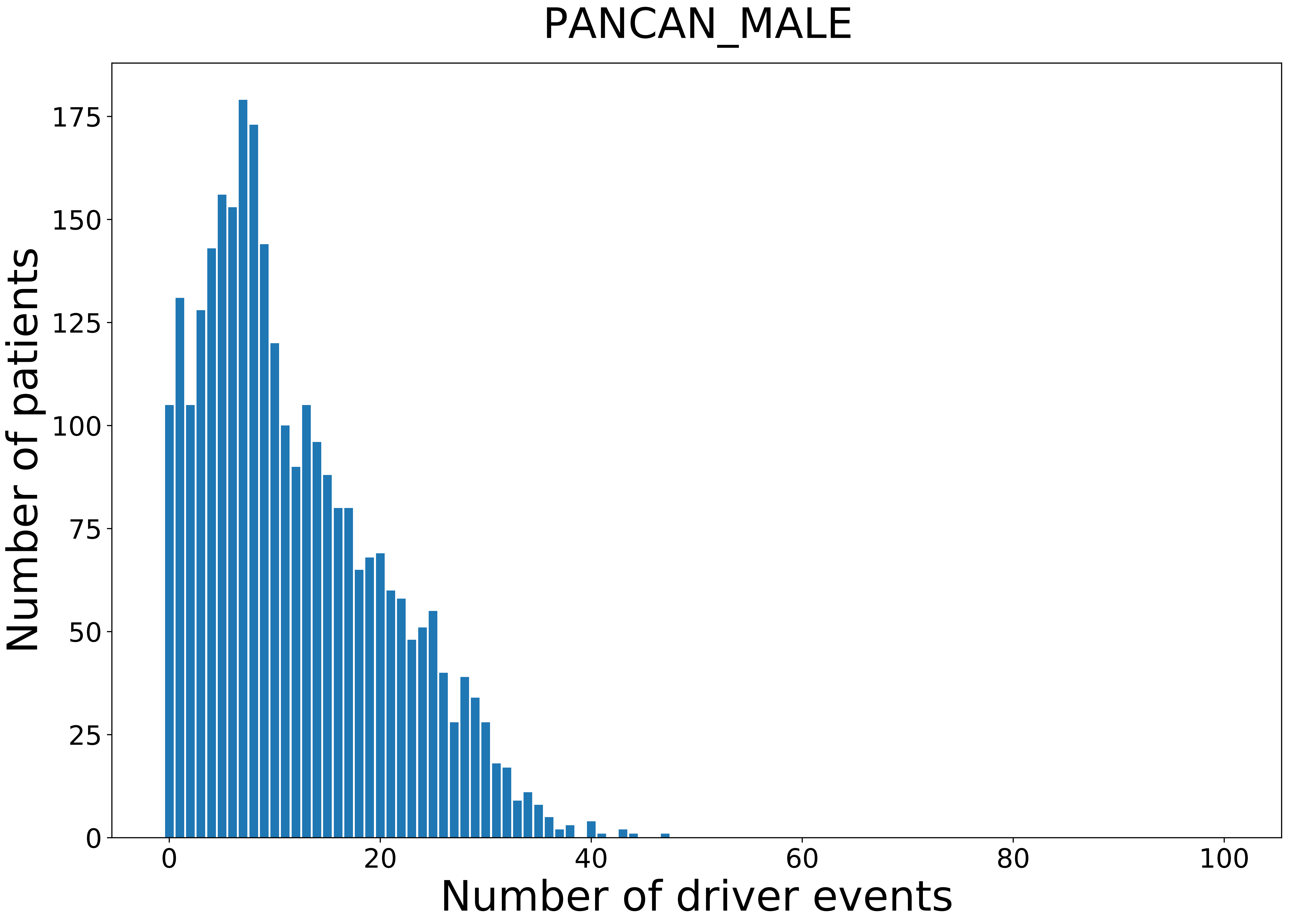

### 2021_8_22_13_29_PCPG_FEMALE.pdf

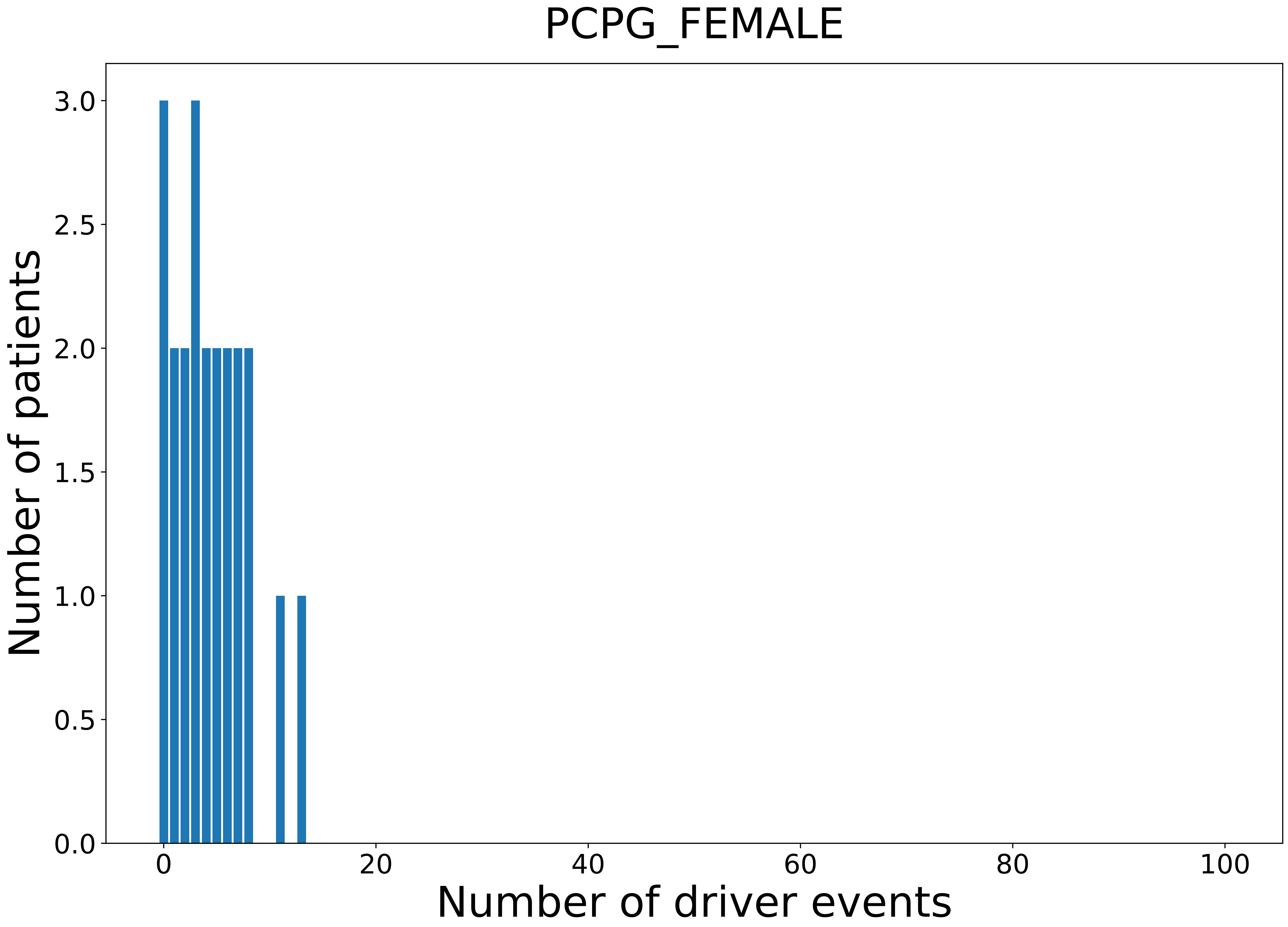

### 2021_8_22_13_29_PCPG_MALE.pdf

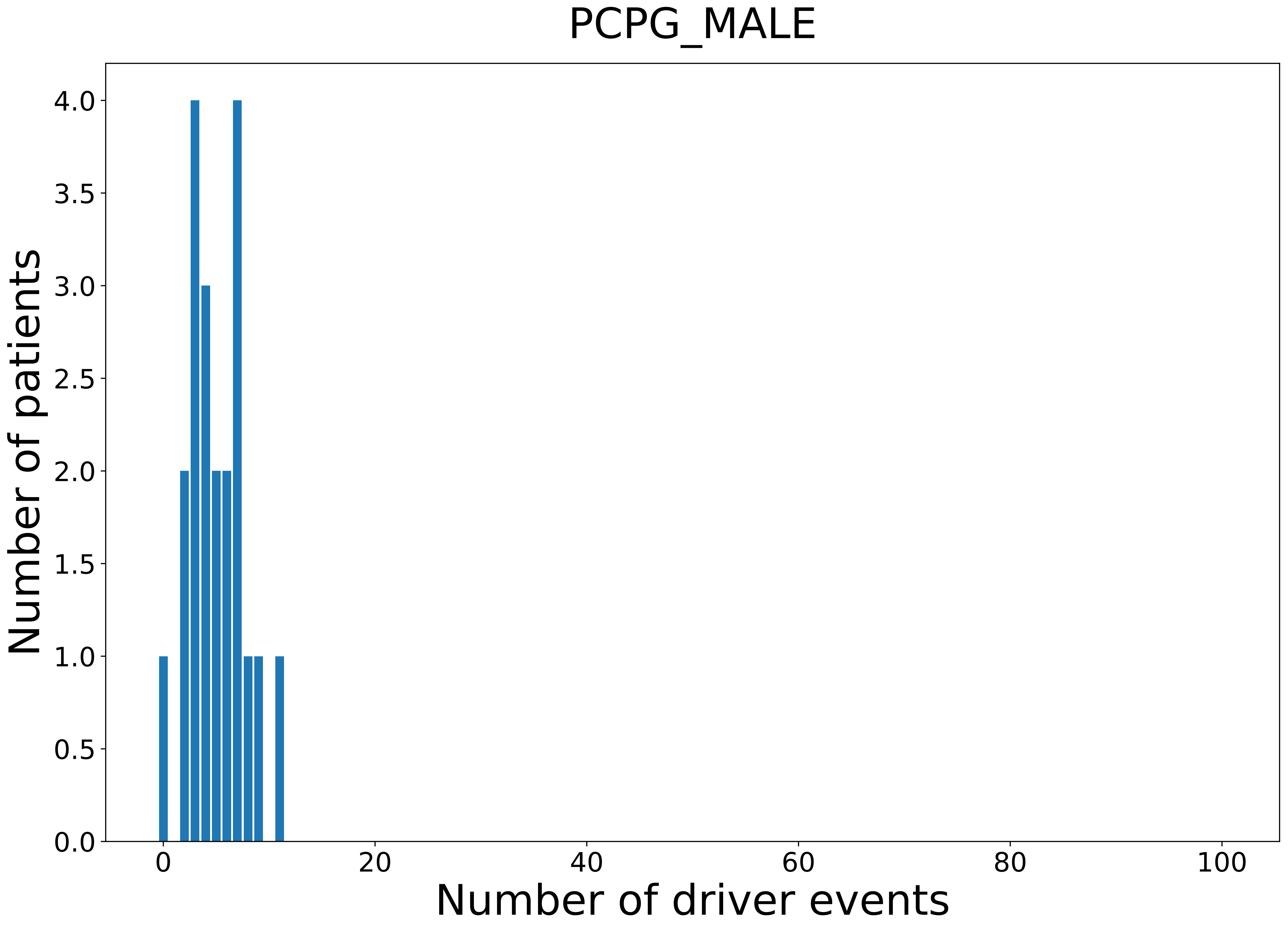

### 2021_8_22_13_29_PRAD.pdf

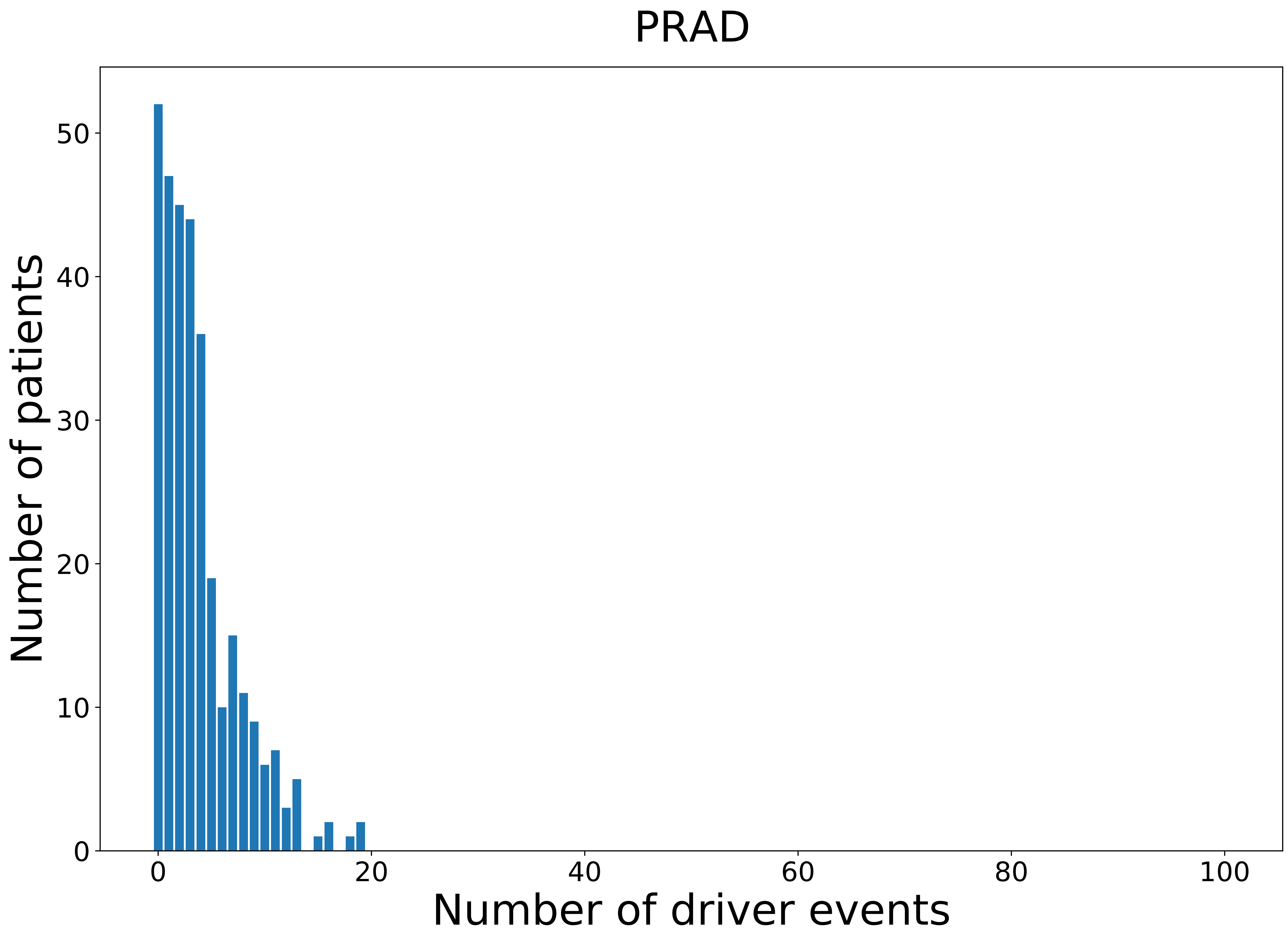

### 2021_8_22_13_29_PRAD_MALE.pdf

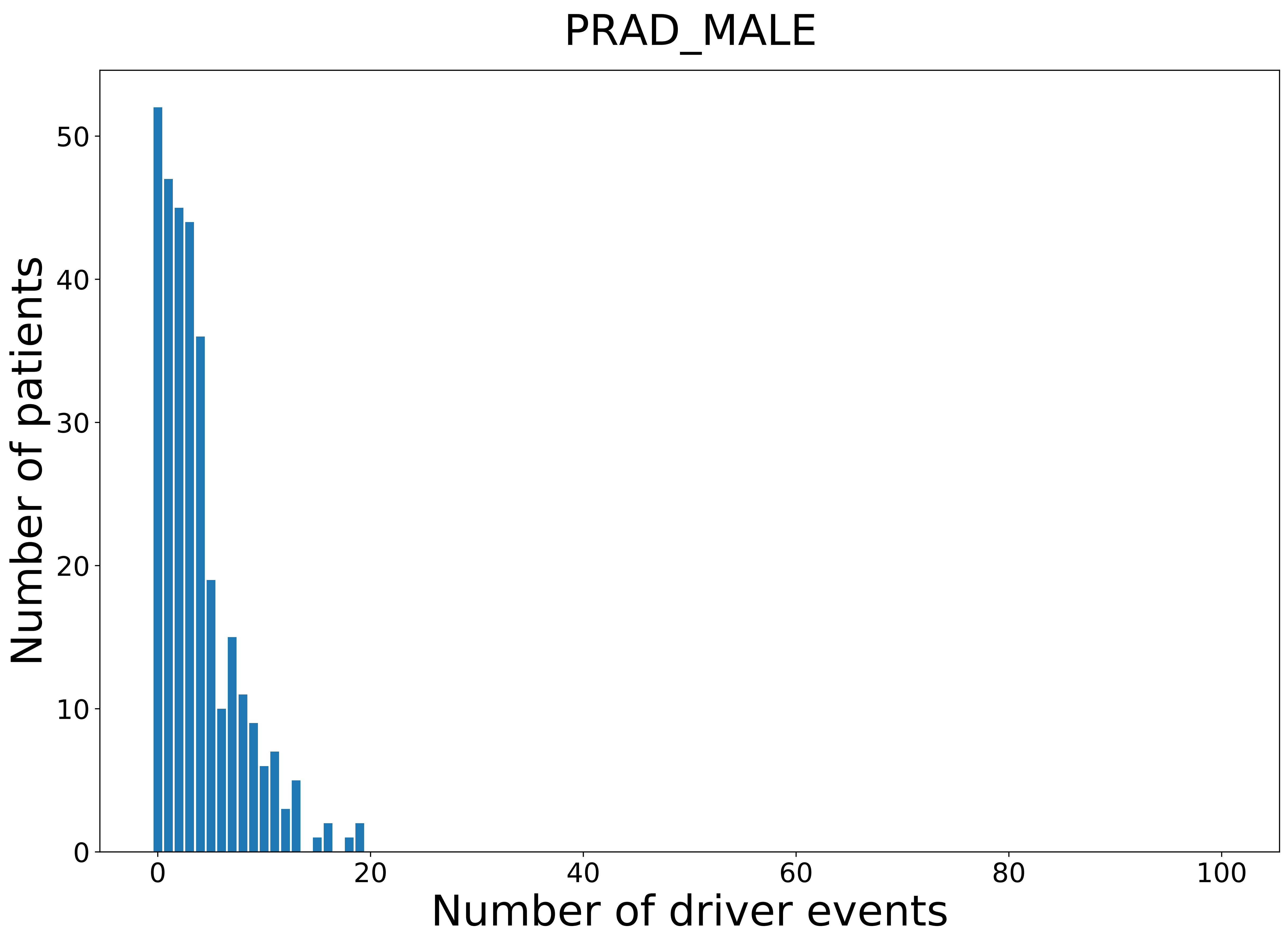

### 2021_8_22_13_29_READ.pdf

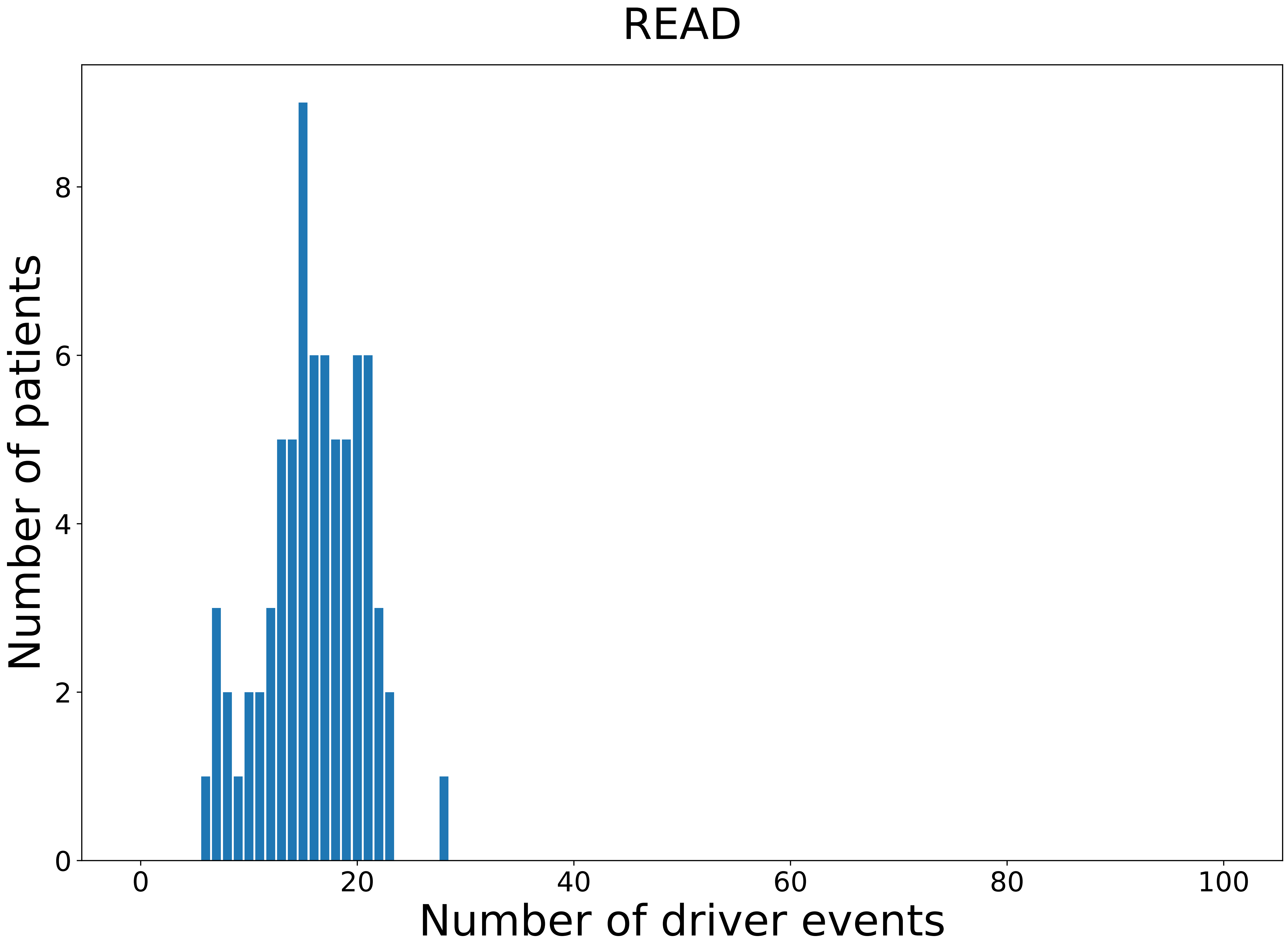

### 2021_8_22_13_29_READ_FEMALE.pdf

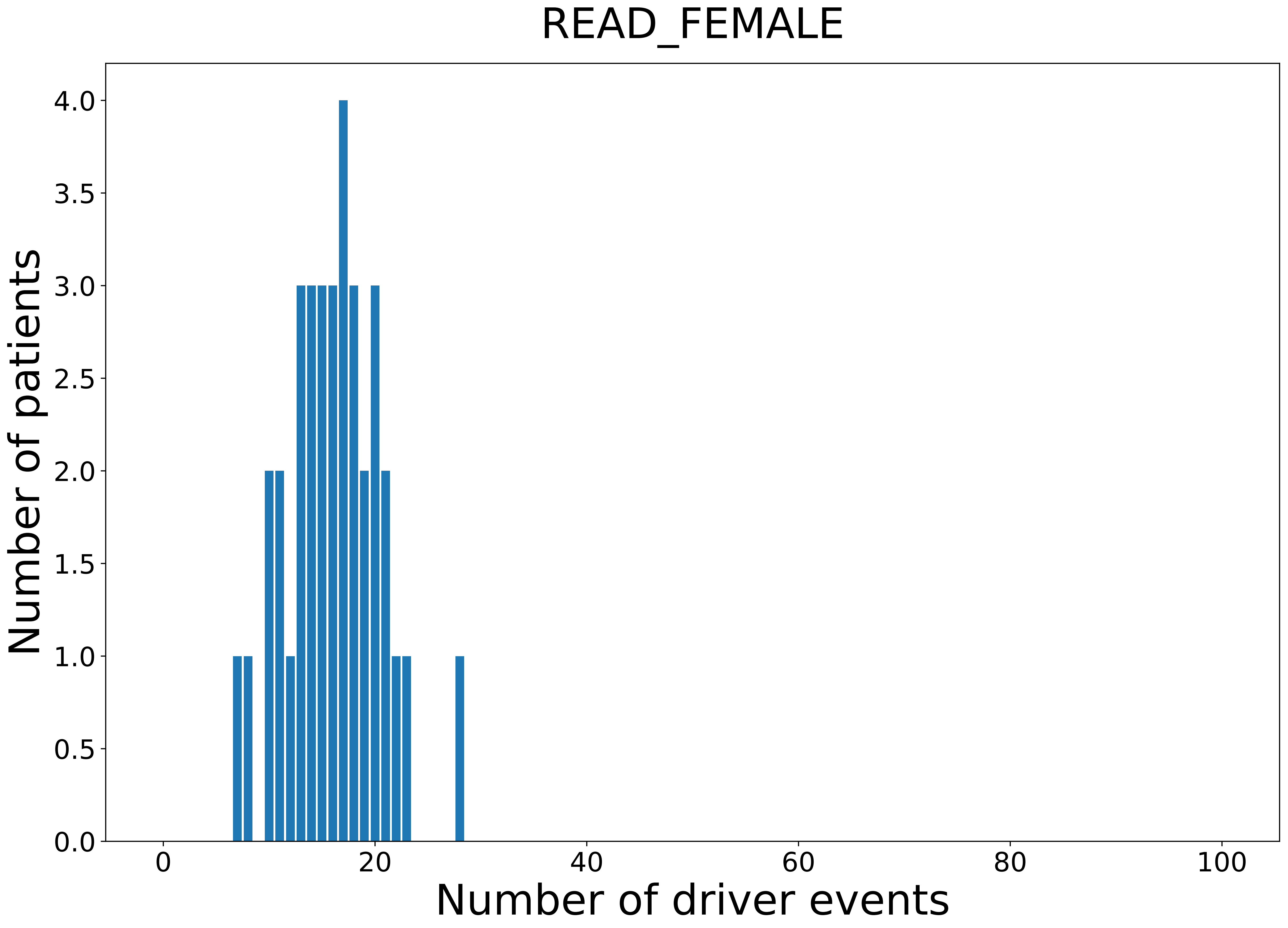

### 2021_8_22_13_29_READ_MALE.pdf

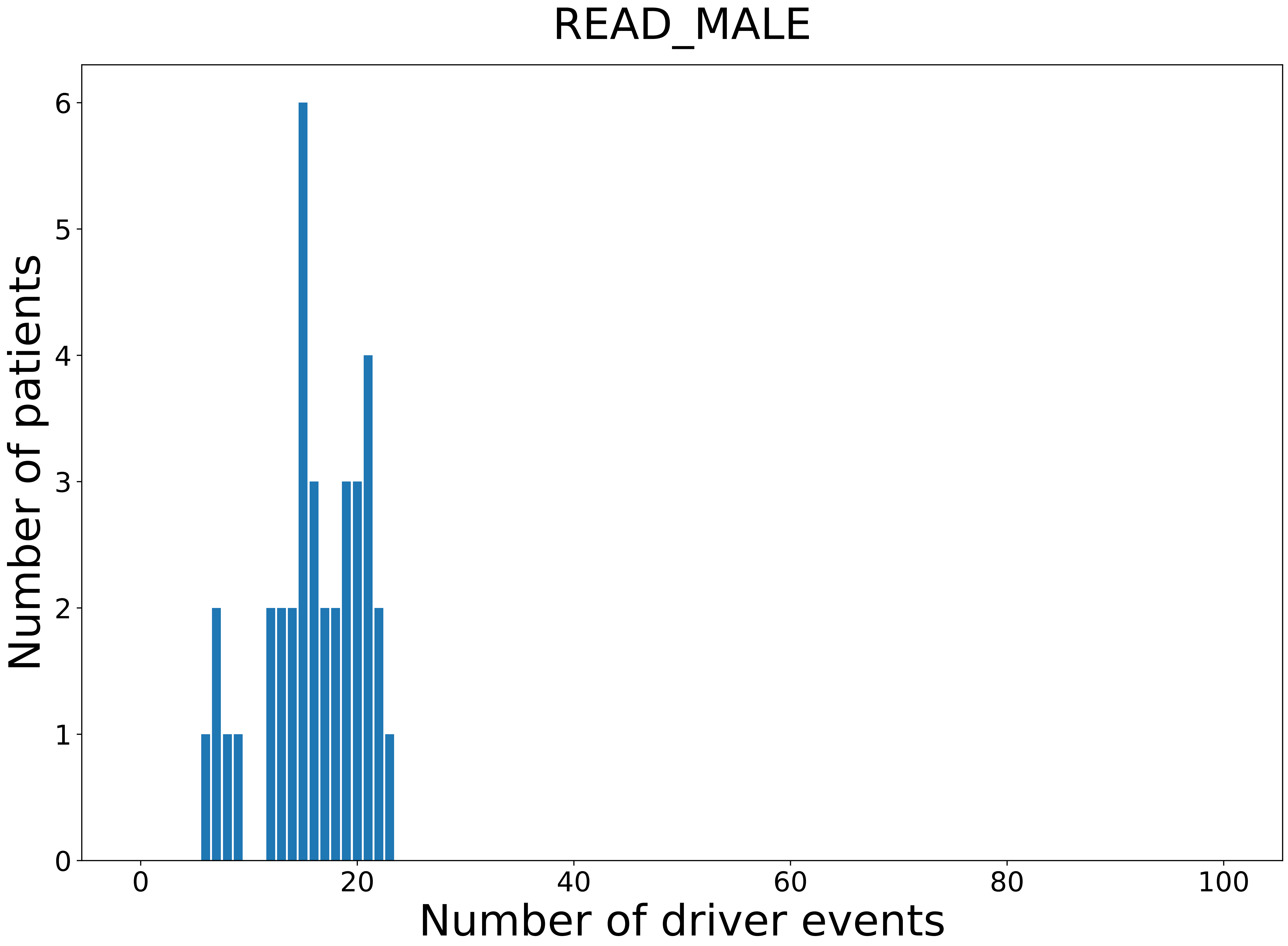

### 2021_8_22_13_29_SARC.pdf

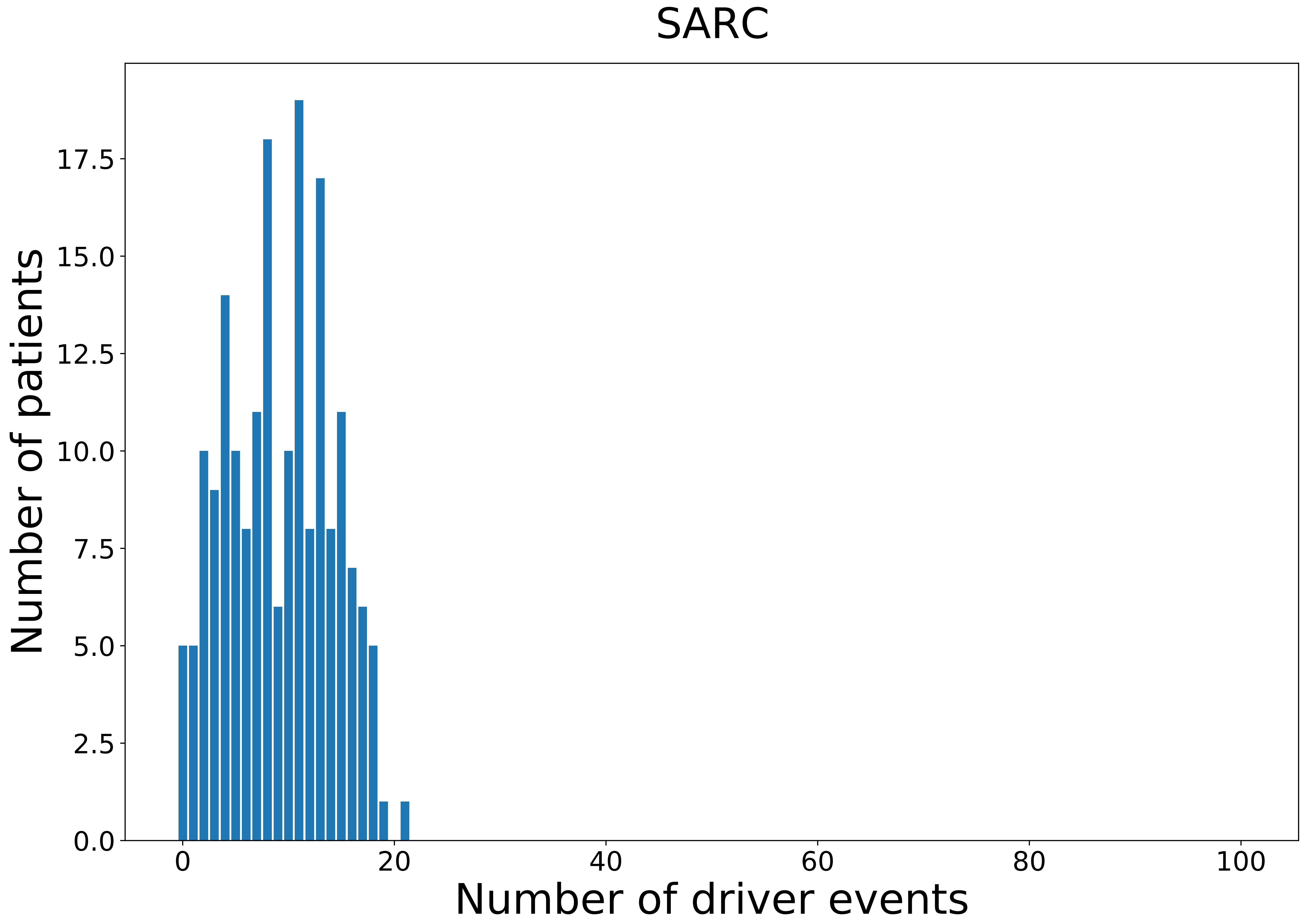

### 2021_8_22_13_29_SARC_MALE.pdf

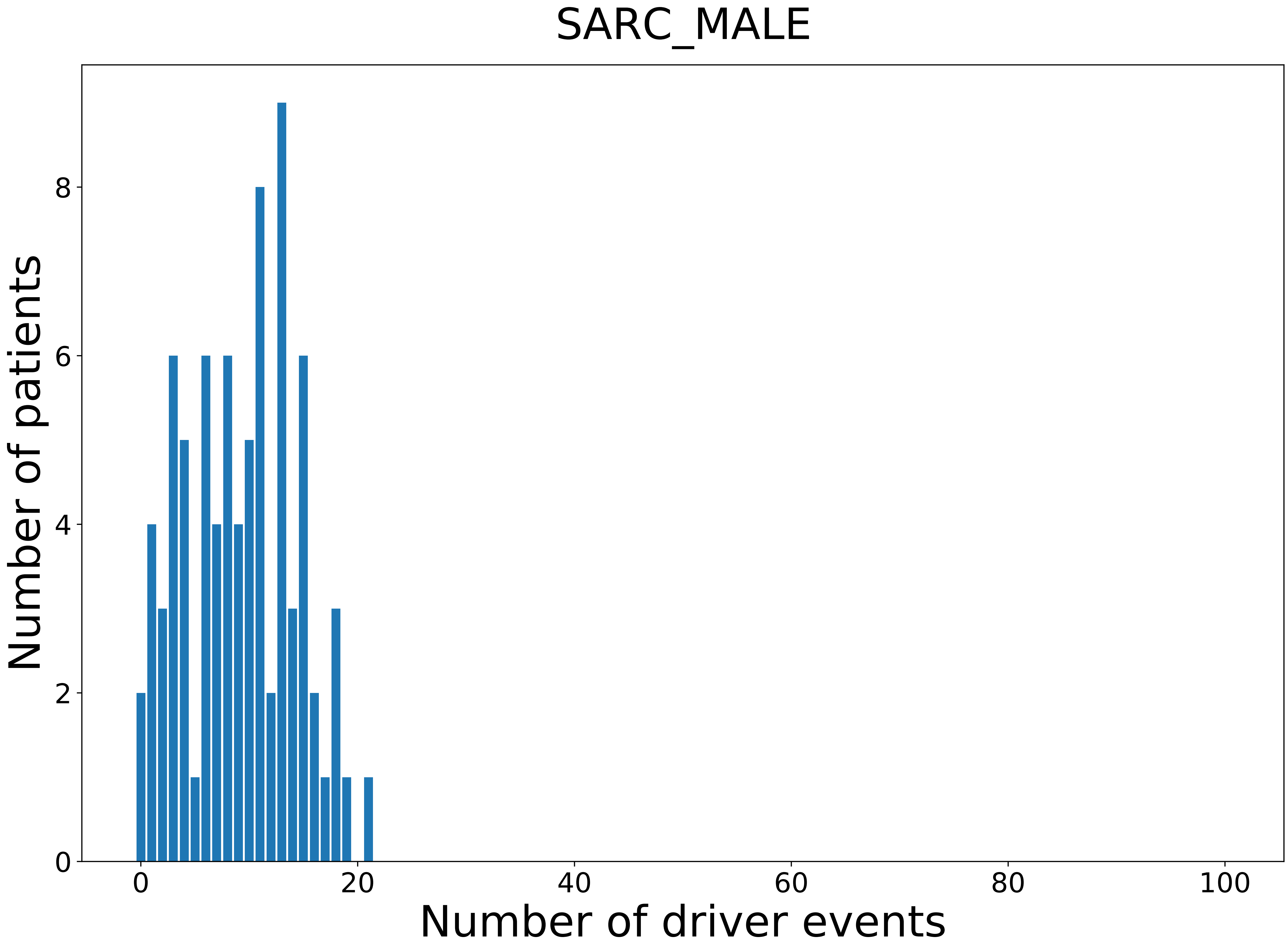

### 2021_8_22_13_29_SKCM.pdf

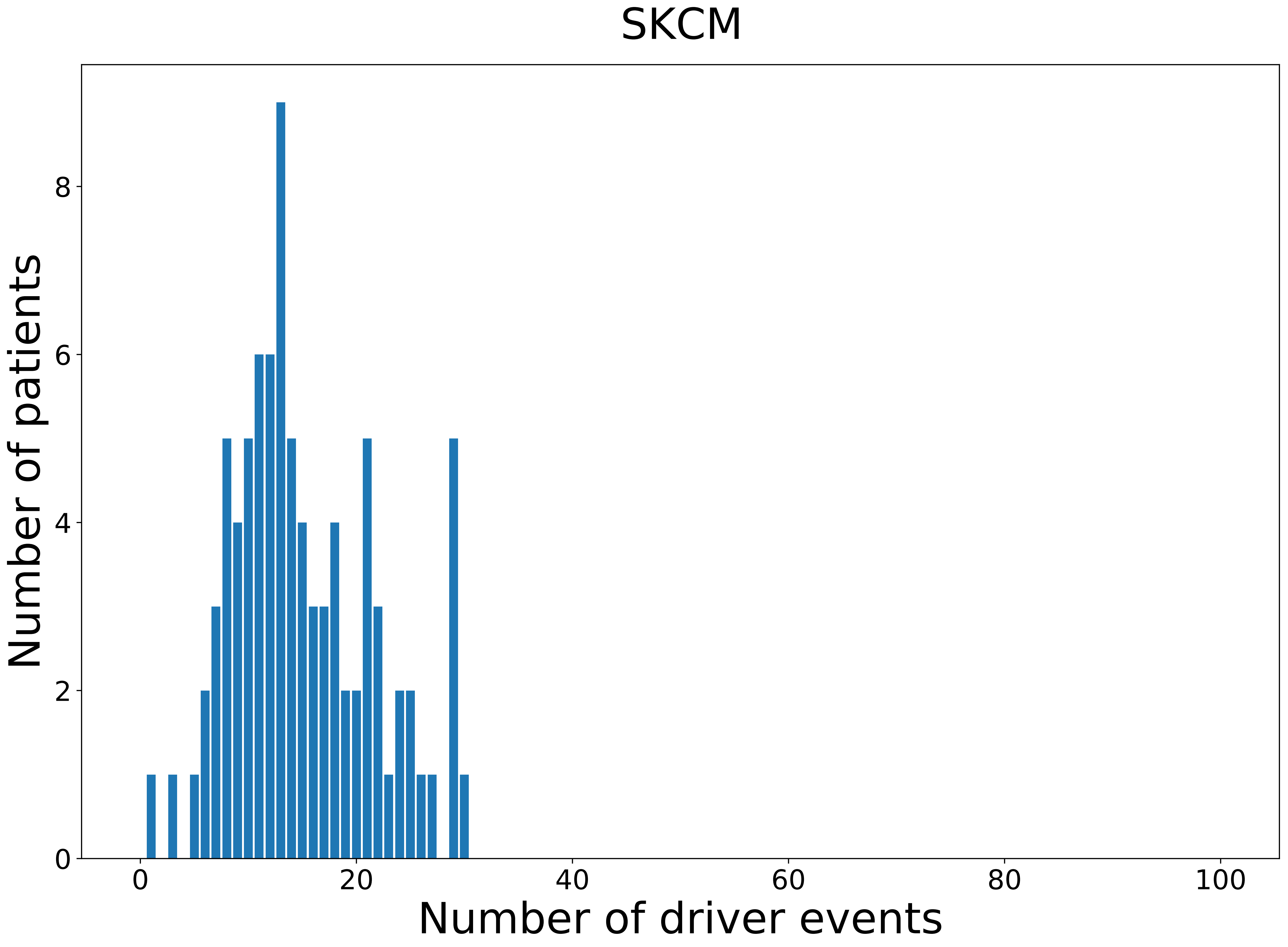

### 2021_8_22_13_29_SKCM_FEMALE.pdf

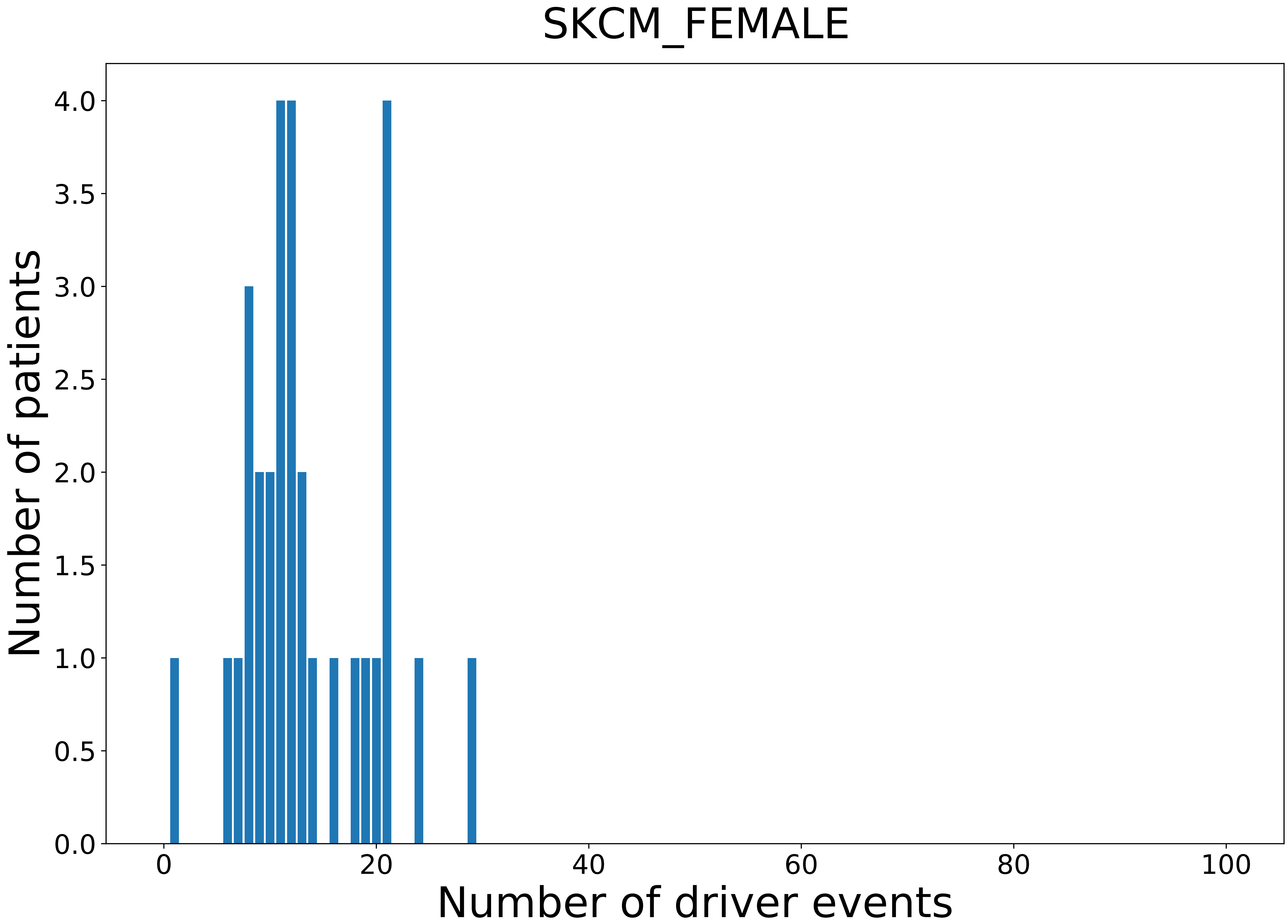

### 2021_8_22_13_29_SKCM_MALE.pdf

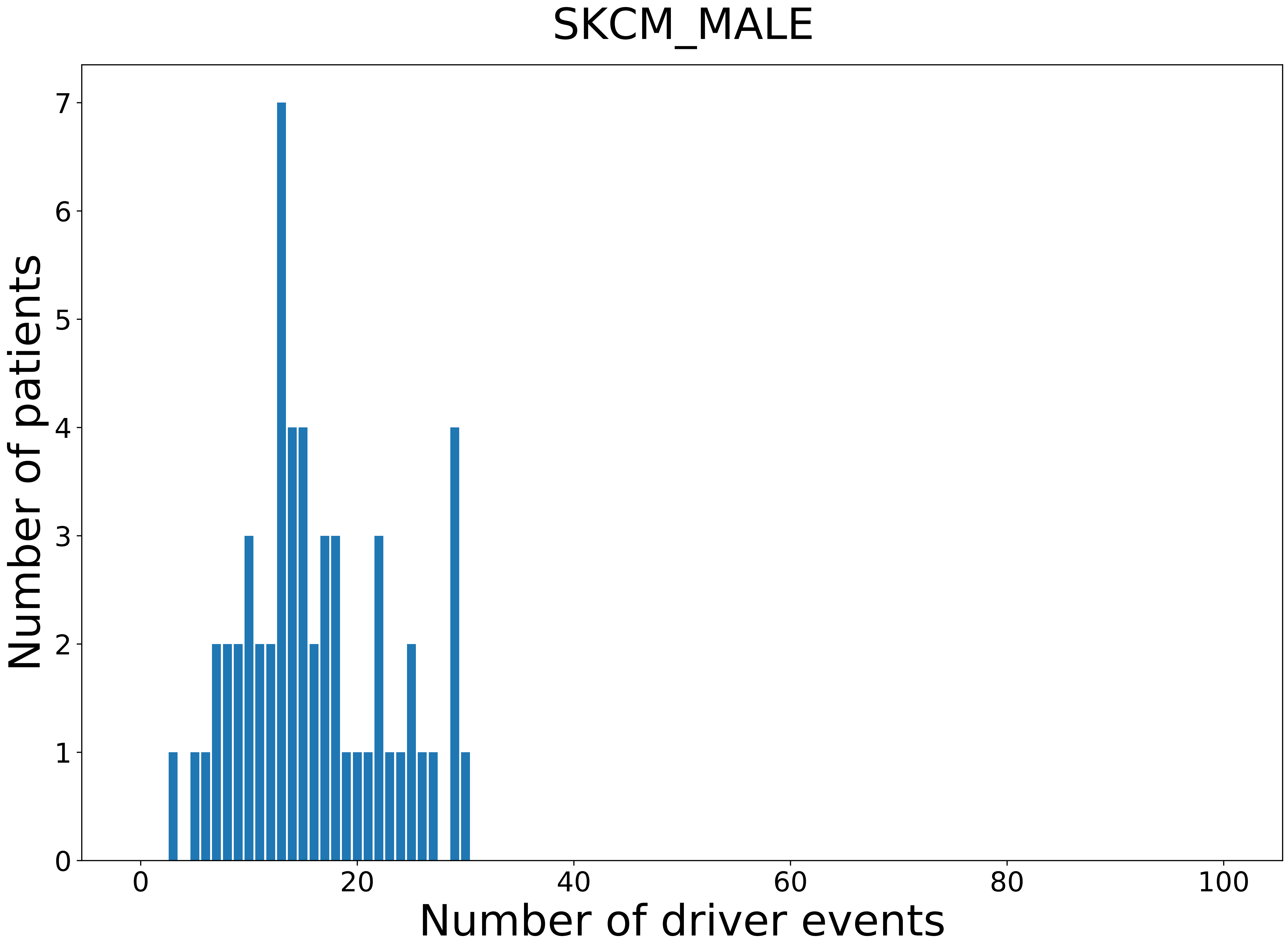

### 2021_8_22_13_29_STAD.pdf

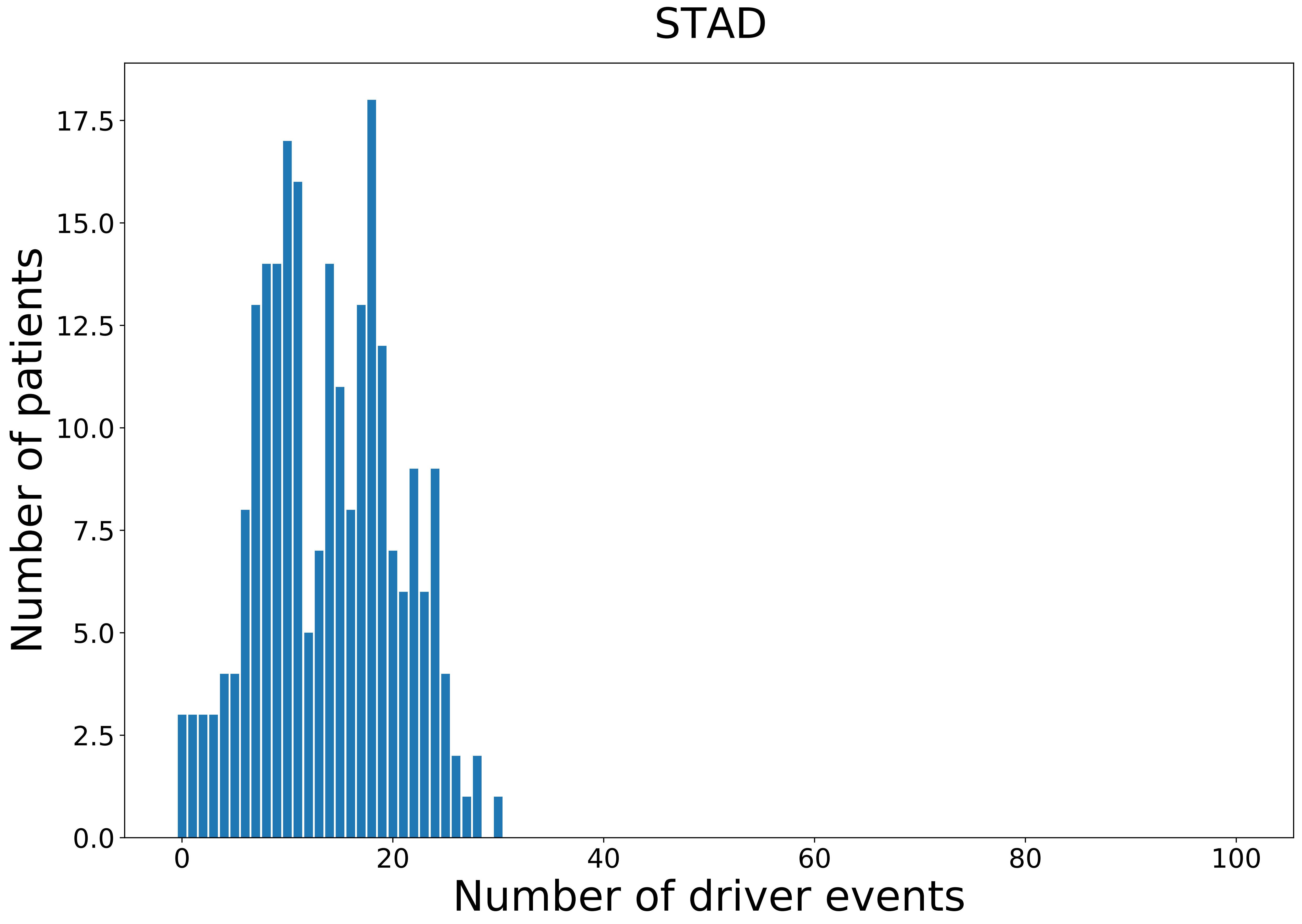

### 2021_8_22_13_29_STAD_MALE.pdf

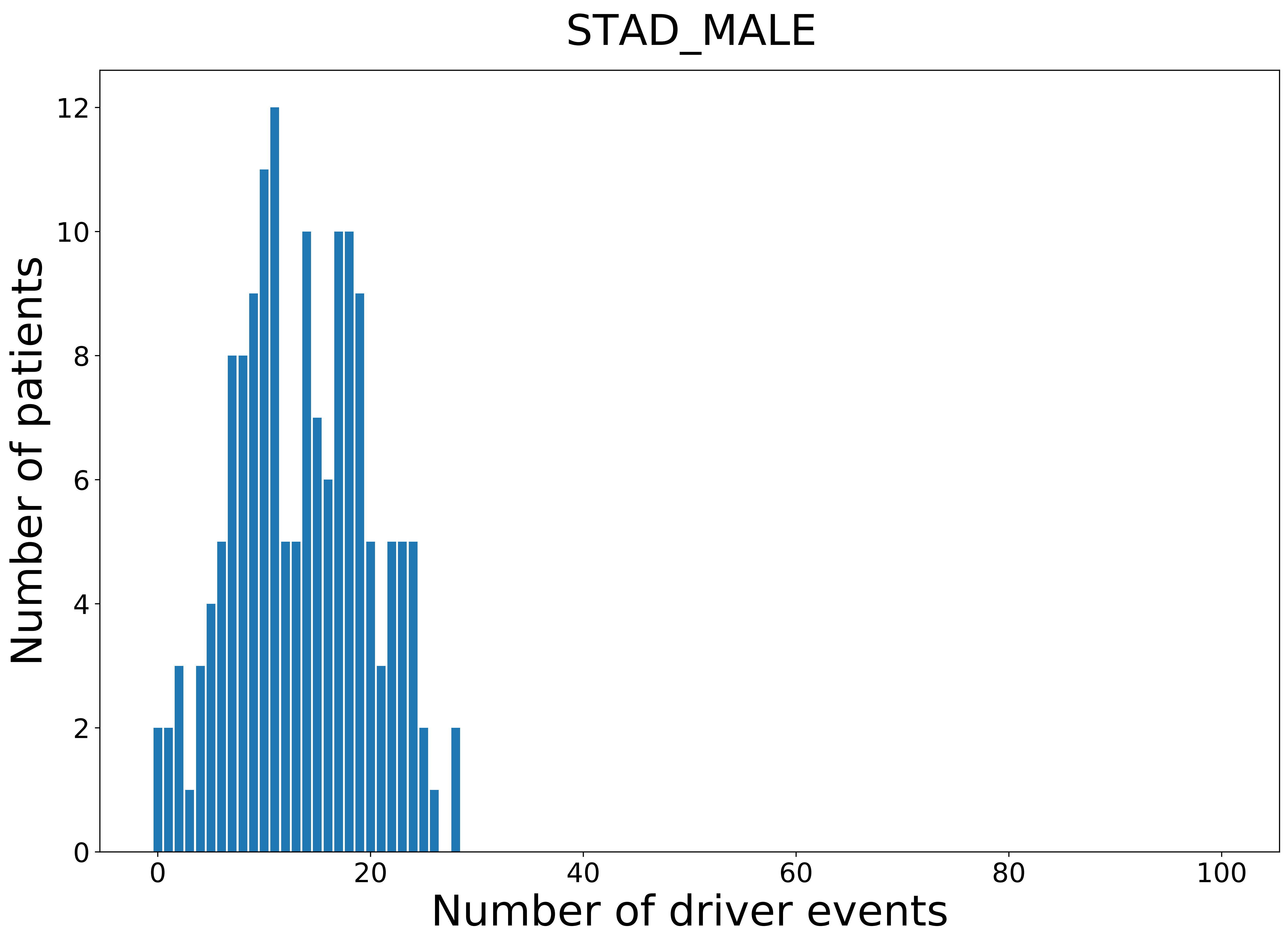

### 2021_8_22_13_29_TGCT_MALE.pdf

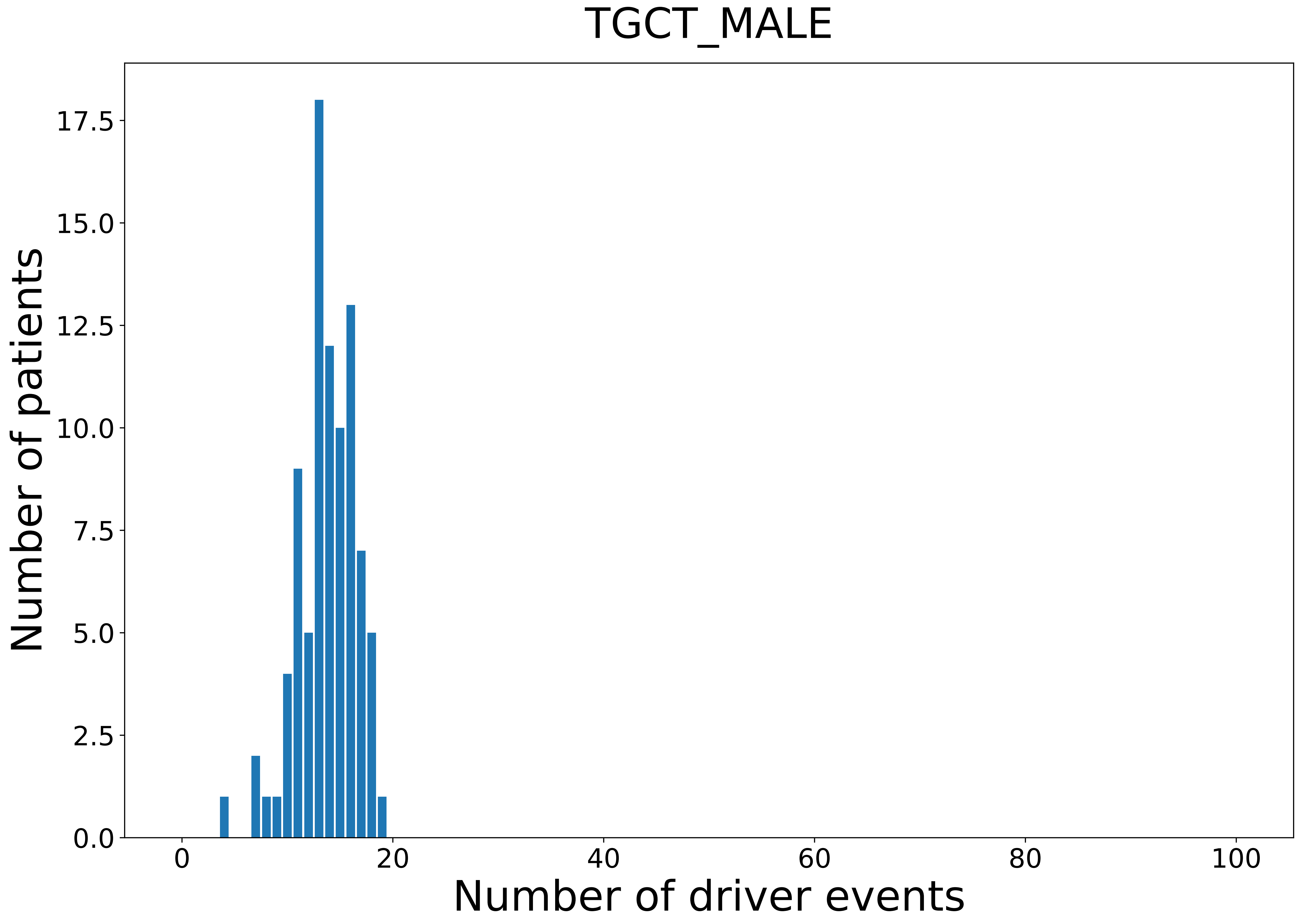

### 2021_8_22_13_29_THCA.pdf

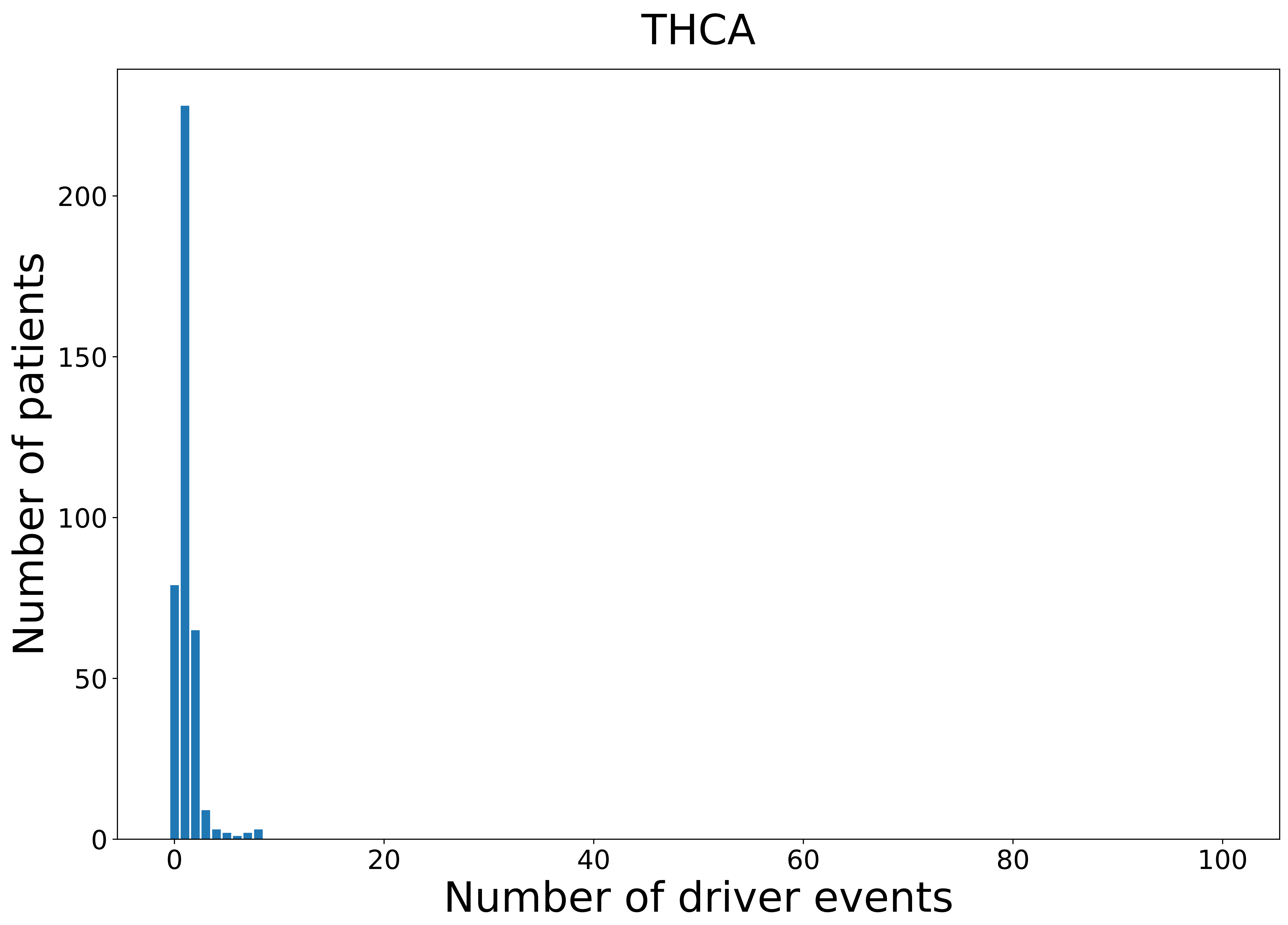

### 2021_8_22_13_29_THCA_FEMALE.pdf

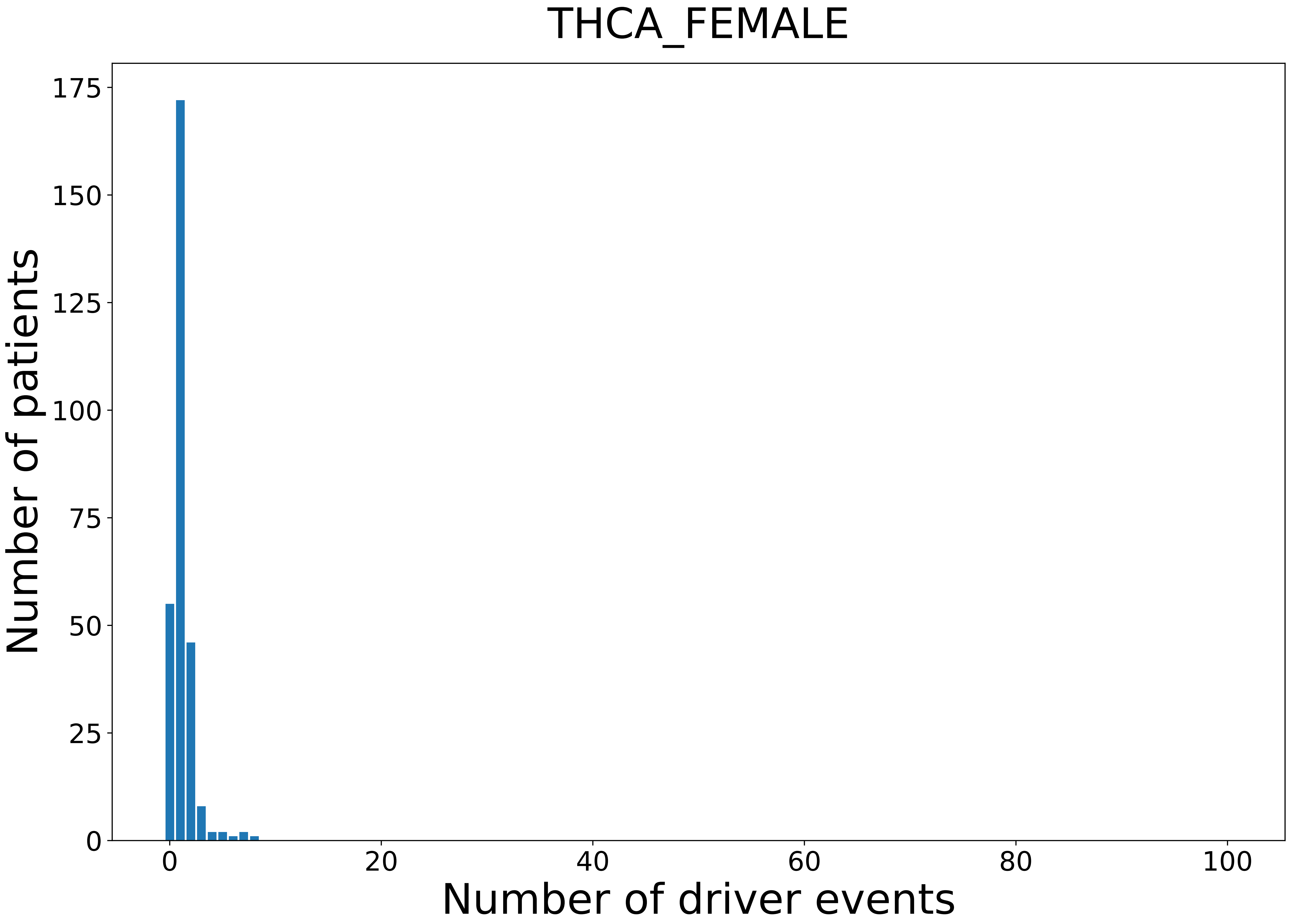

### 2021_8_22_13_29_THCA_MALE.pdf

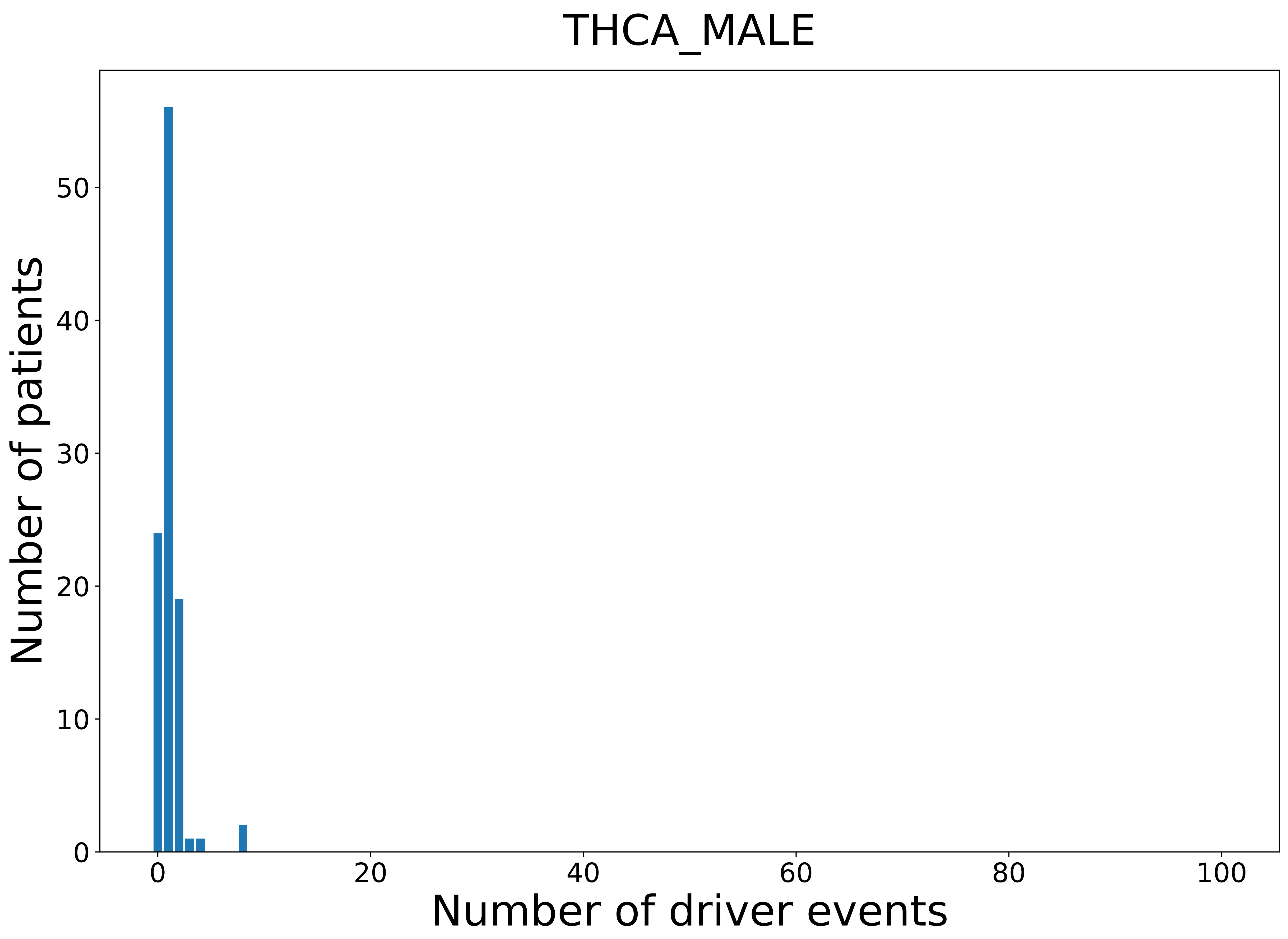

### 2021_8_22_13_29_THYM.pdf

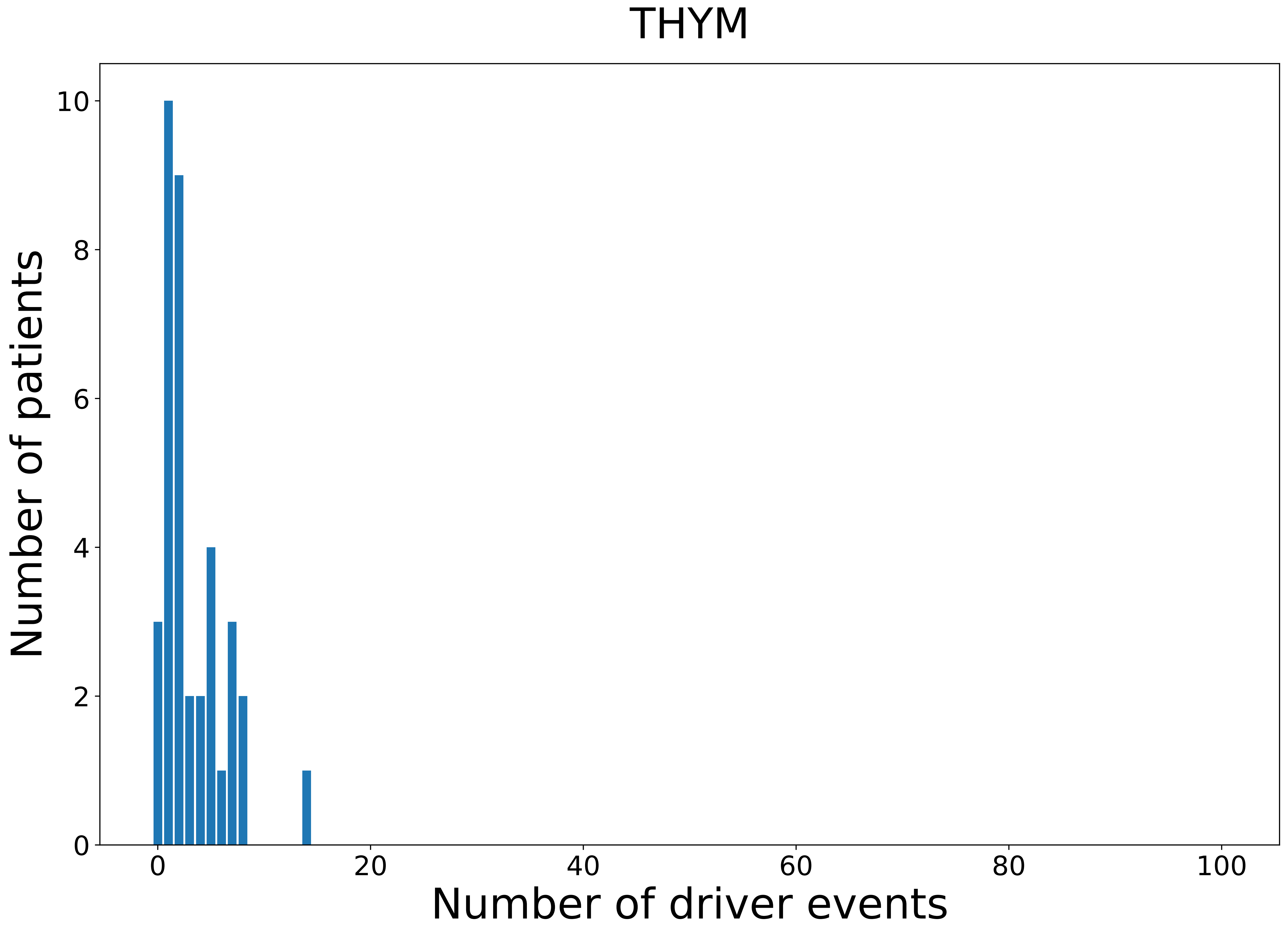

### 2021_8_22_13_29_THYM_FEMALE.pdf

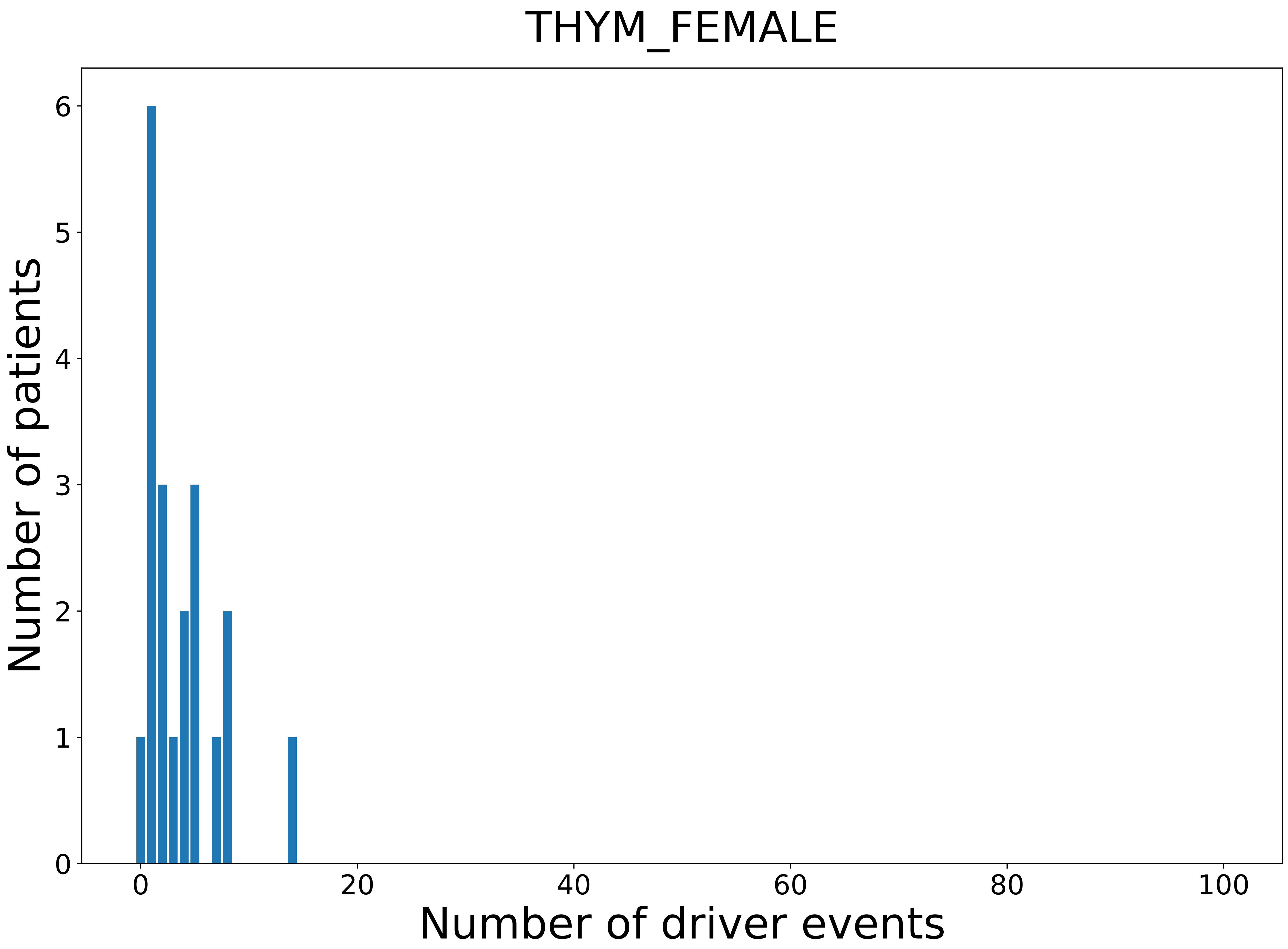

### 2021_8_22_13_29_THYM_MALE.pdf

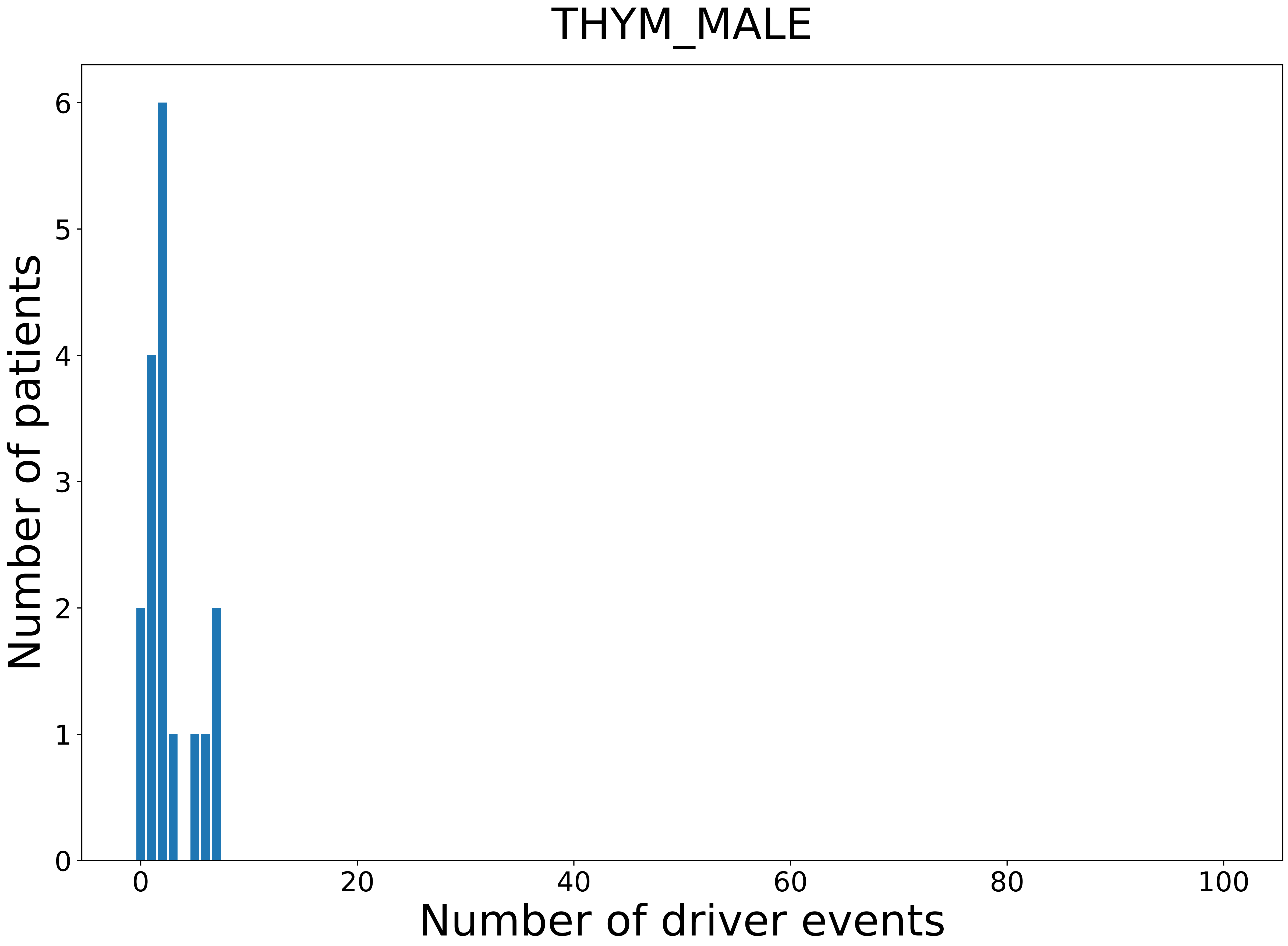

### 2021_8_22_13_29_UCEC.pdf

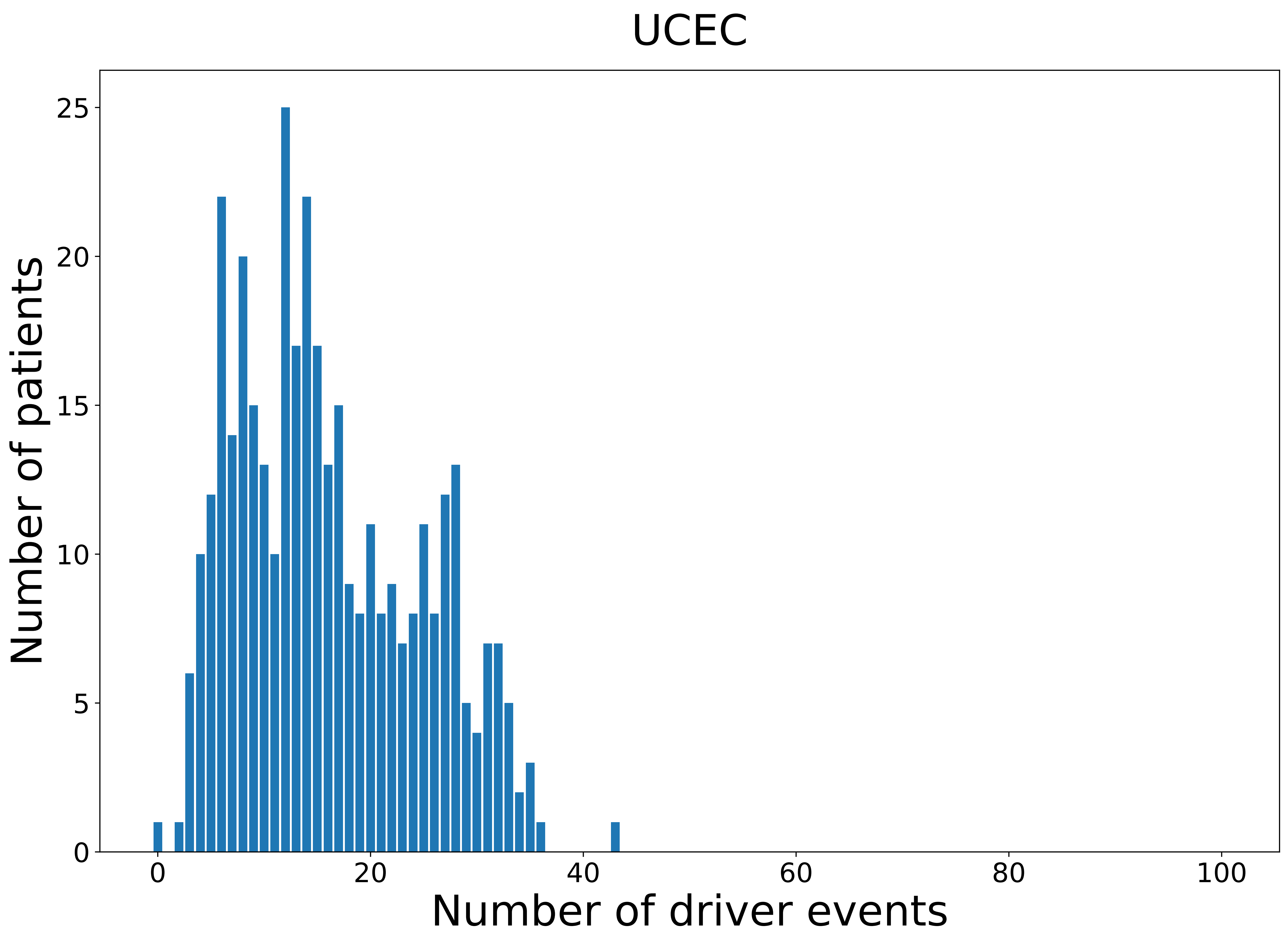

### 2021_8_22_13_29_UCEC_FEMALE.pdf

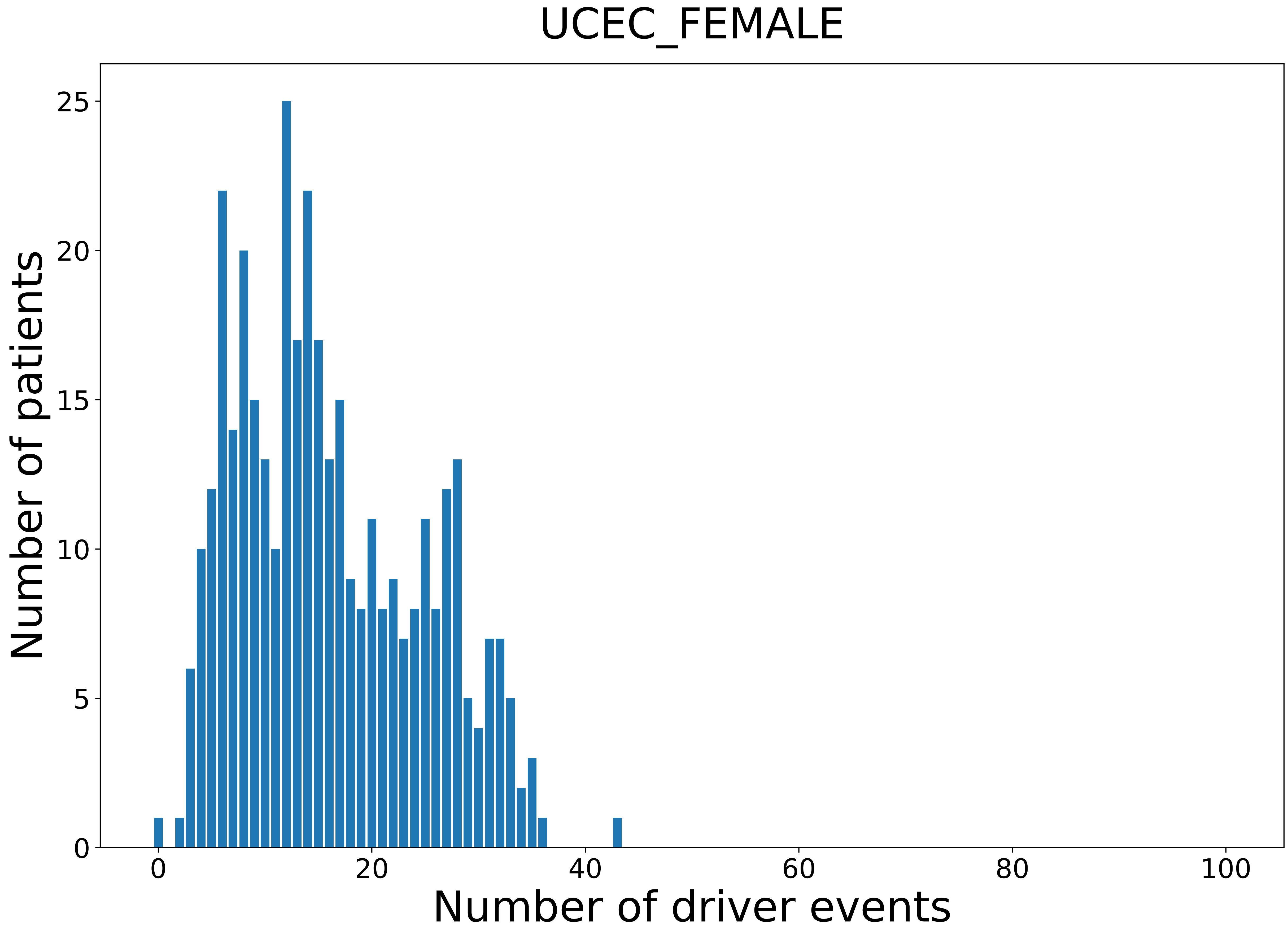

### 2021_8_22_13_29_UCS.pdf

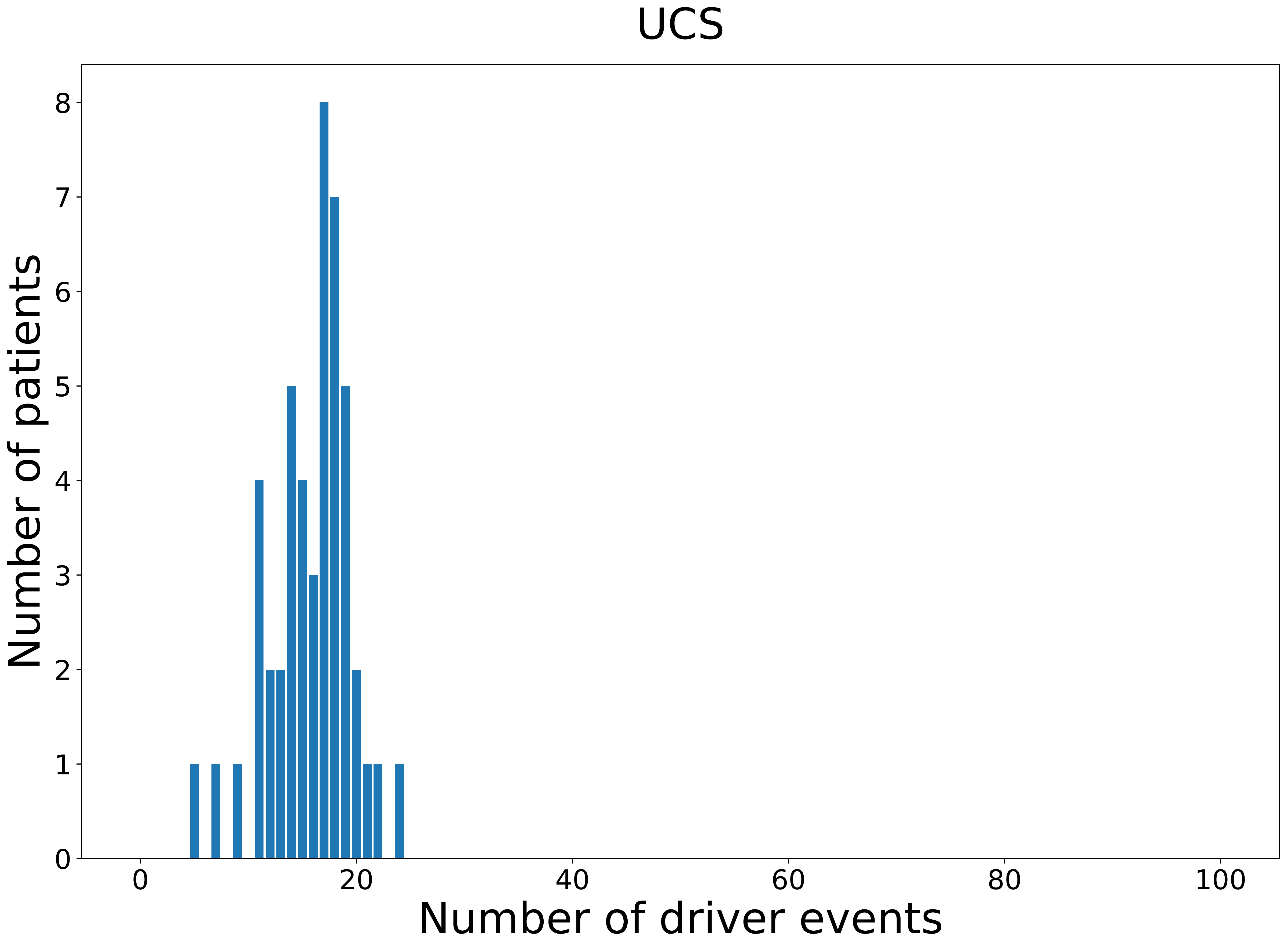

### 2021_8_22_13_29_UCS_FEMALE.pdf

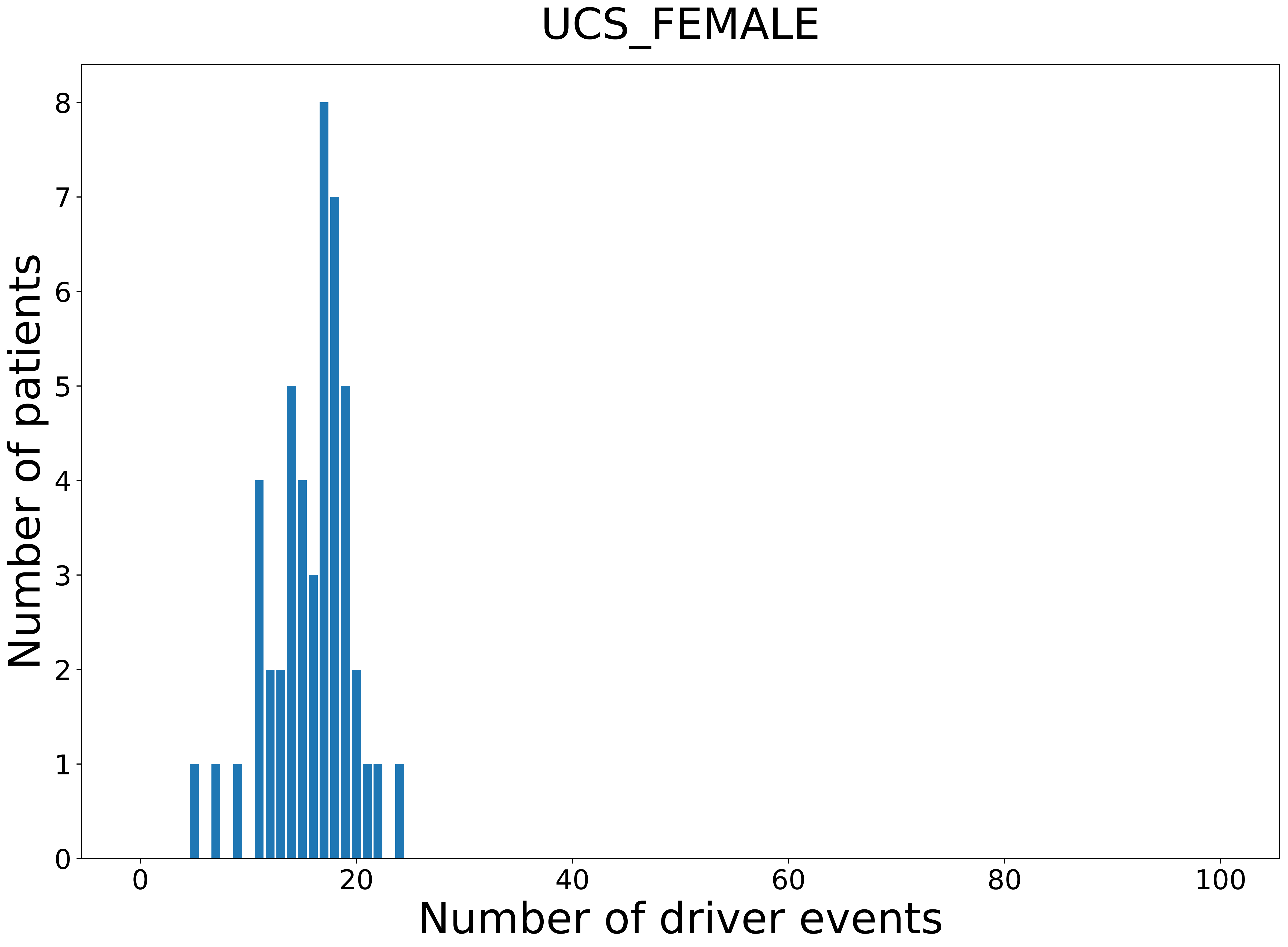

### 2021_8_22_13_29_UVM.pdf

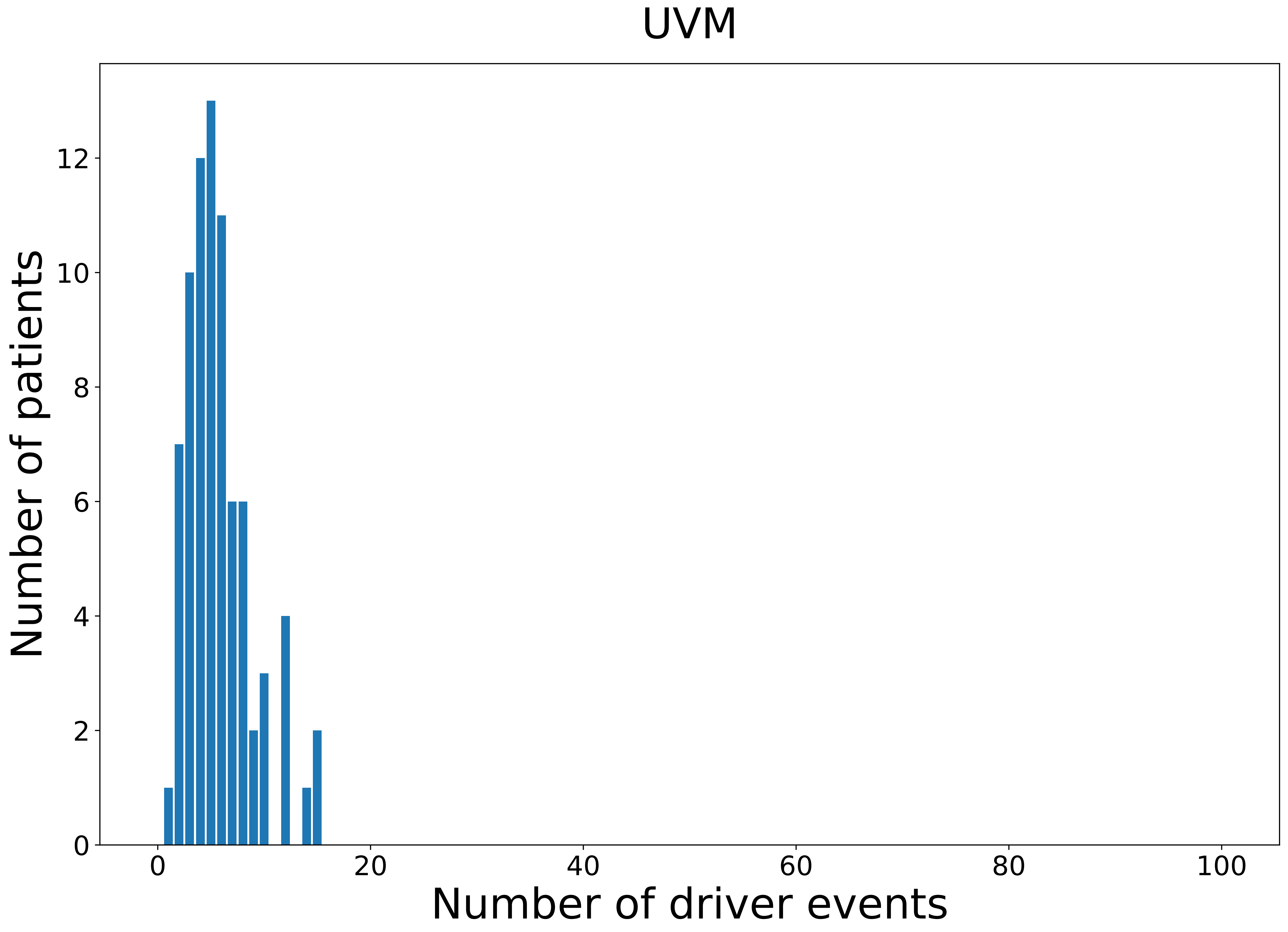

### 2021_8_22_13_42_CESC.pdf

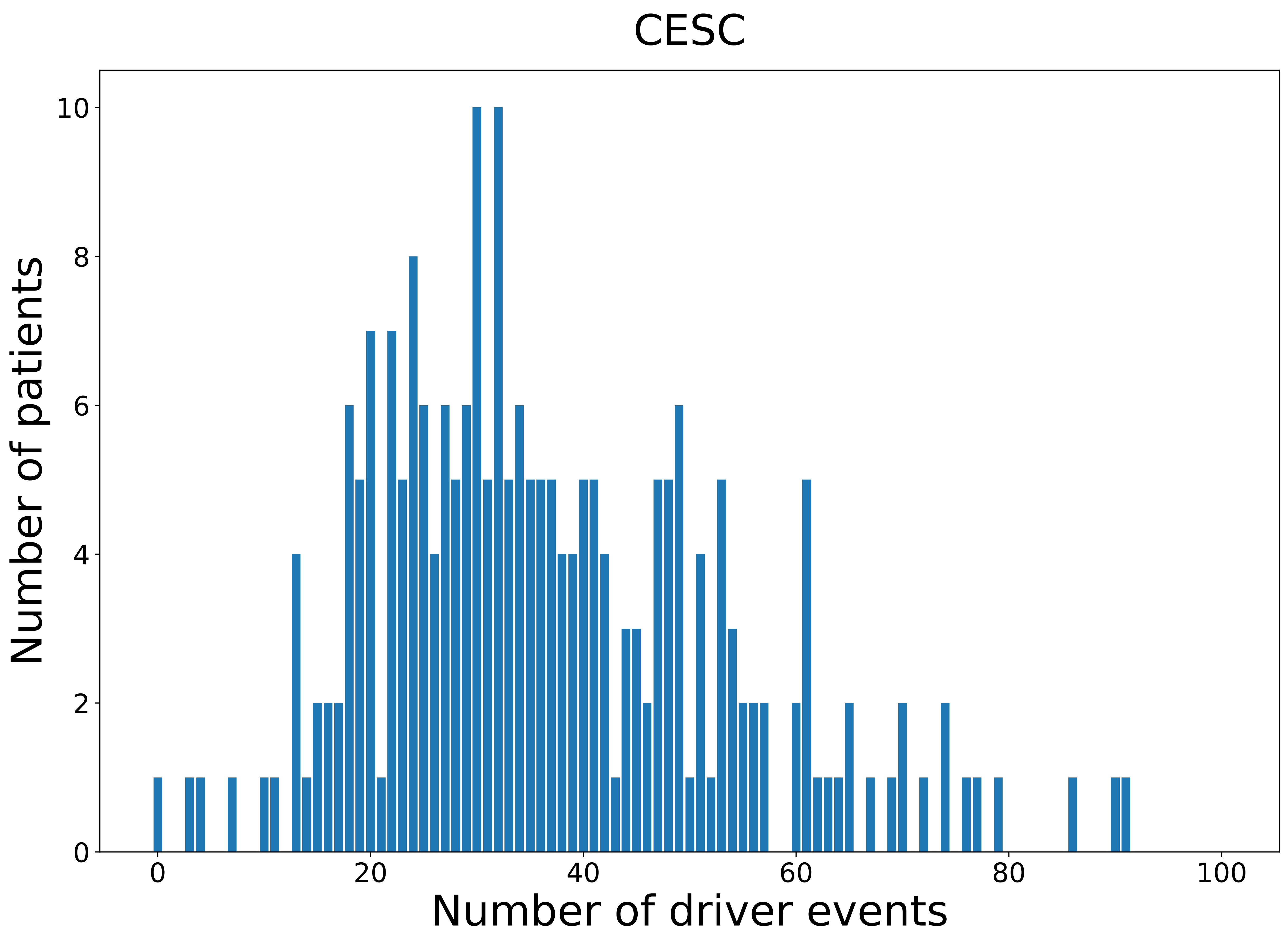

### 2021_8_22_13_42_KICH_MALE.pdf

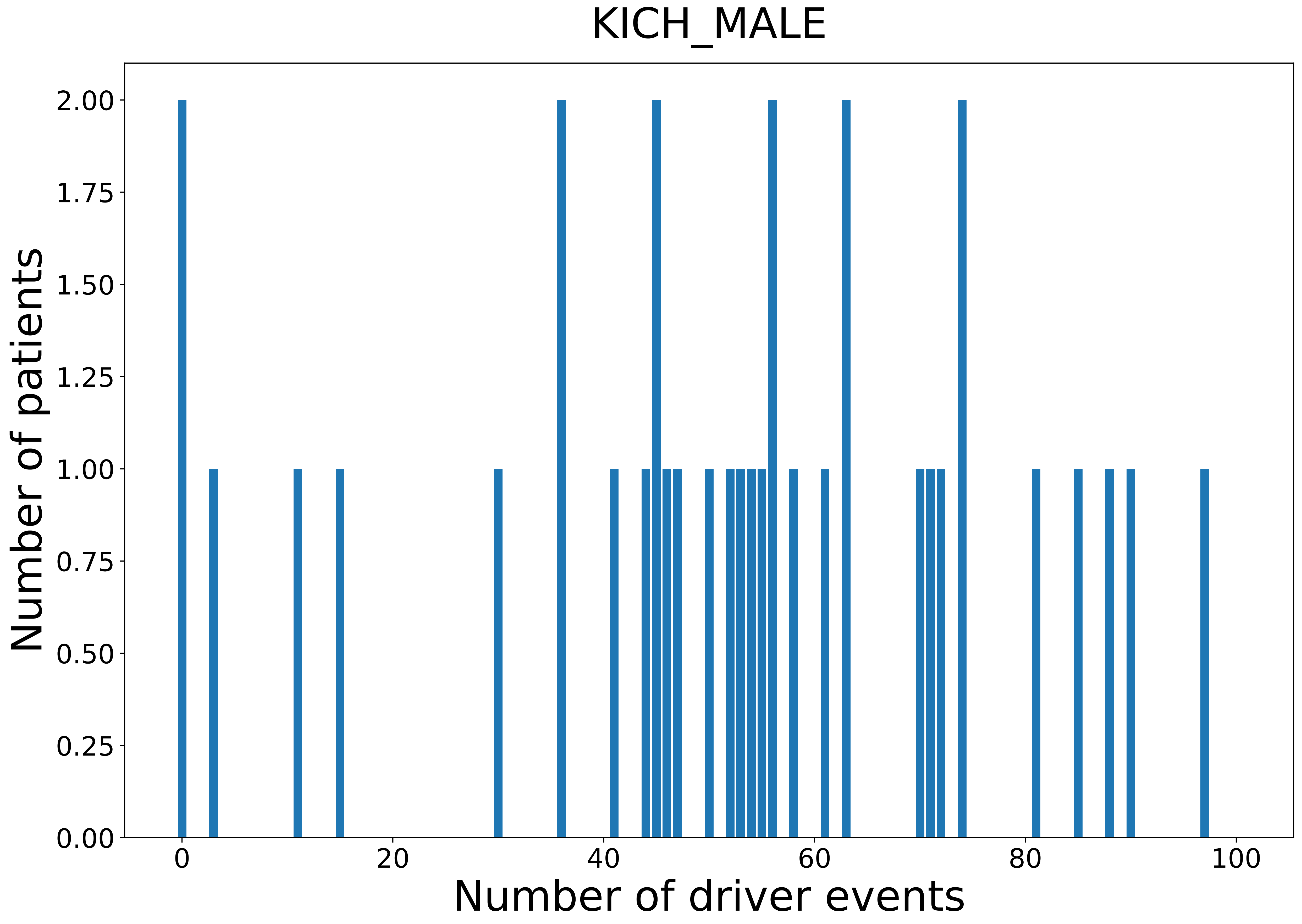

### 2021_8_22_13_42_PCPG_MALE.pdf

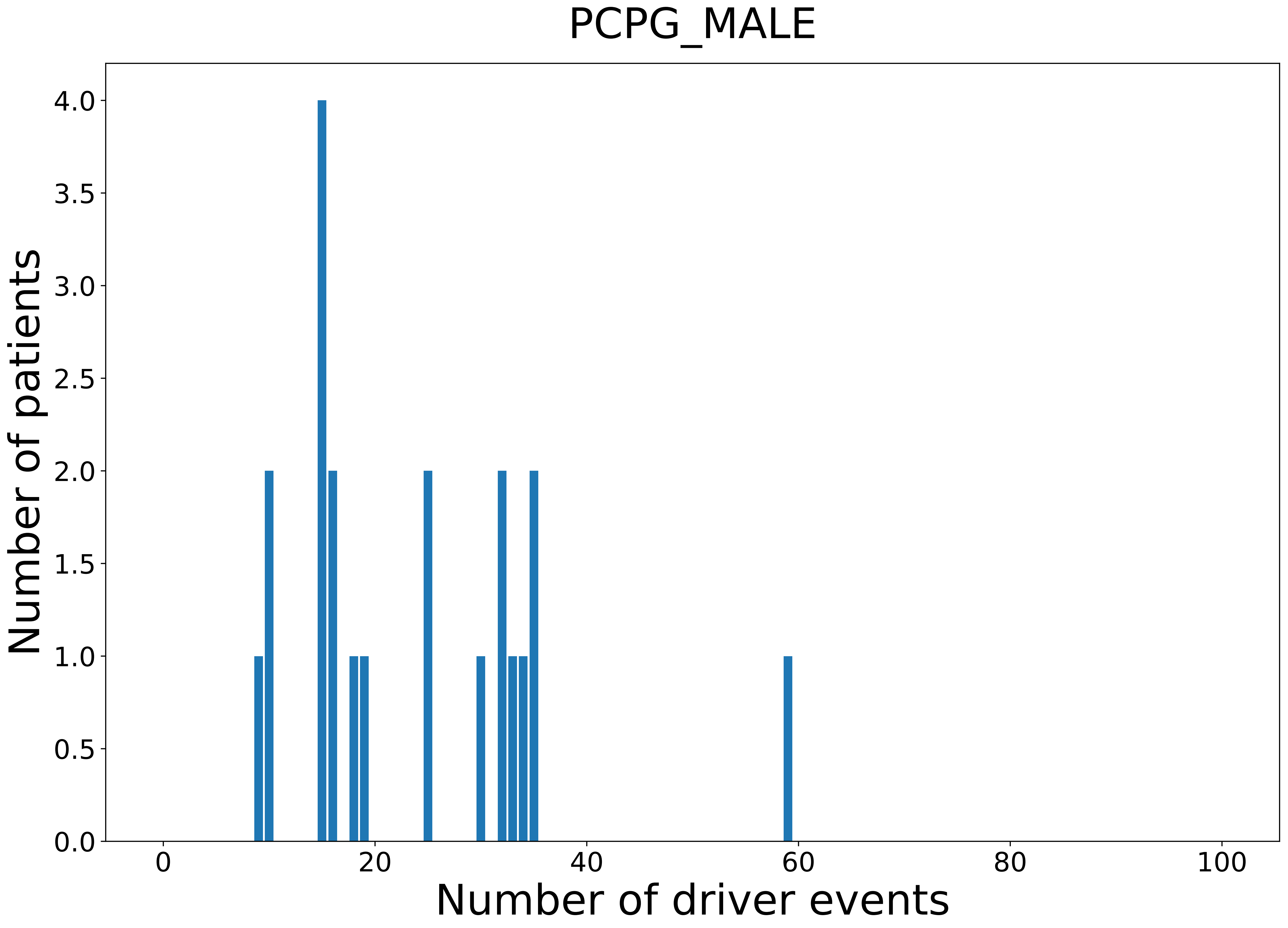

### 2021_8_22_13_42_READ_MALE.pdf

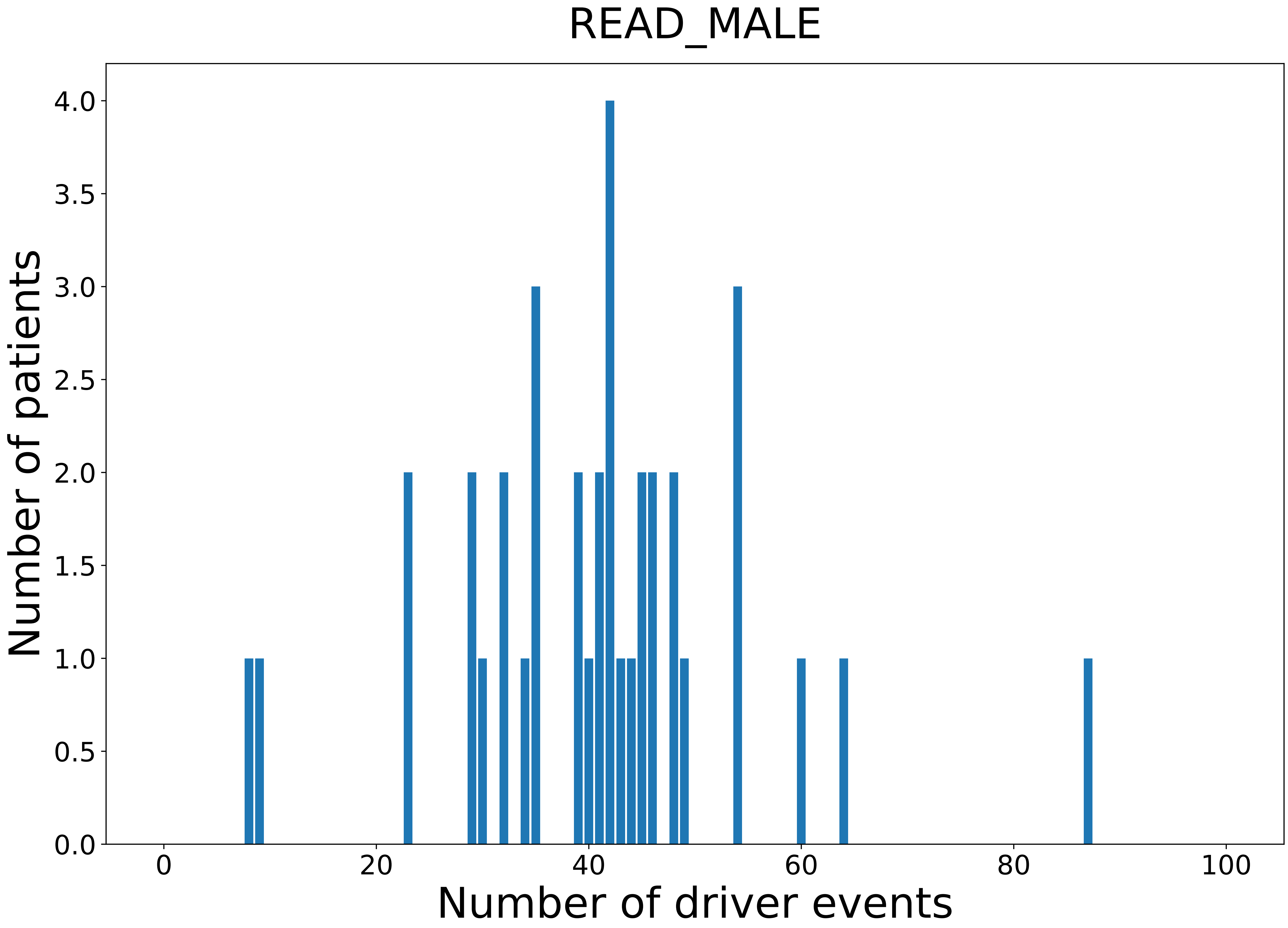

### 2021_8_22_13_42_STAD_FEMALE.pdf

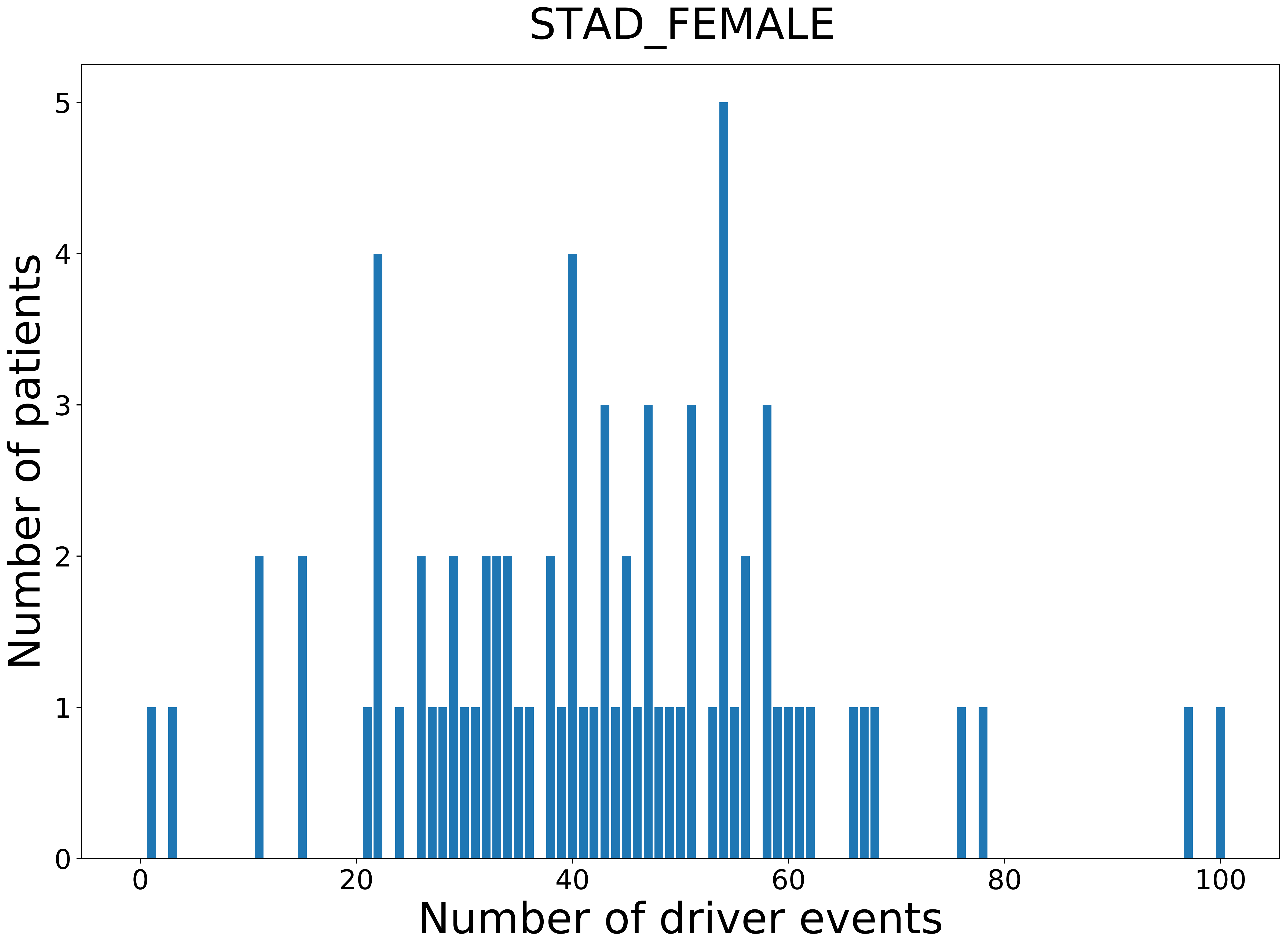

### 2021_8_22_13_42_THCA_MALE.pdf

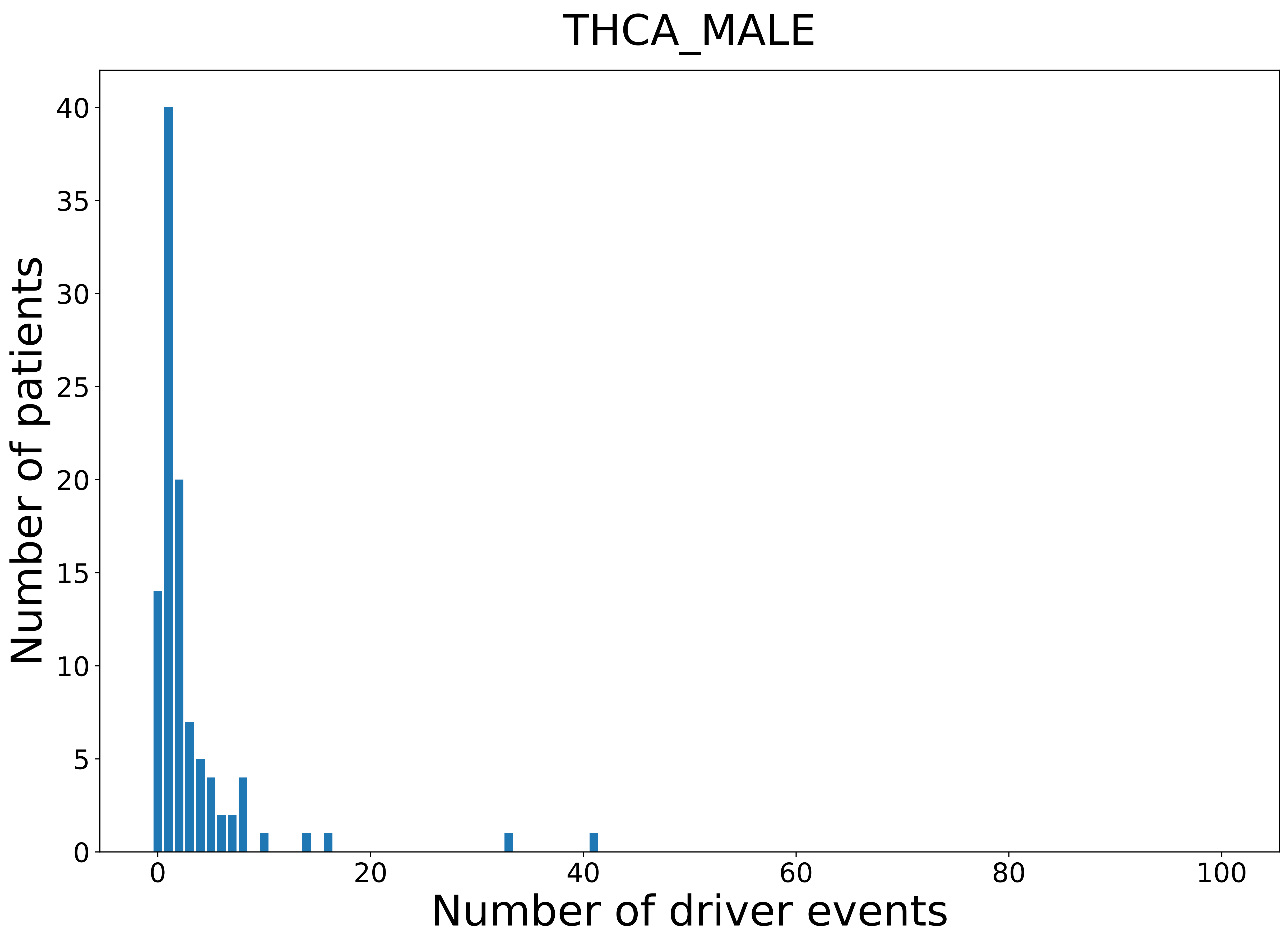
