## Supplementary figures and images for "Comprehensive patient-level classification and quantification of driver events in TCGA PanCanAtlas cohorts"

### 2021_8_22_13_42_ACC.pdf

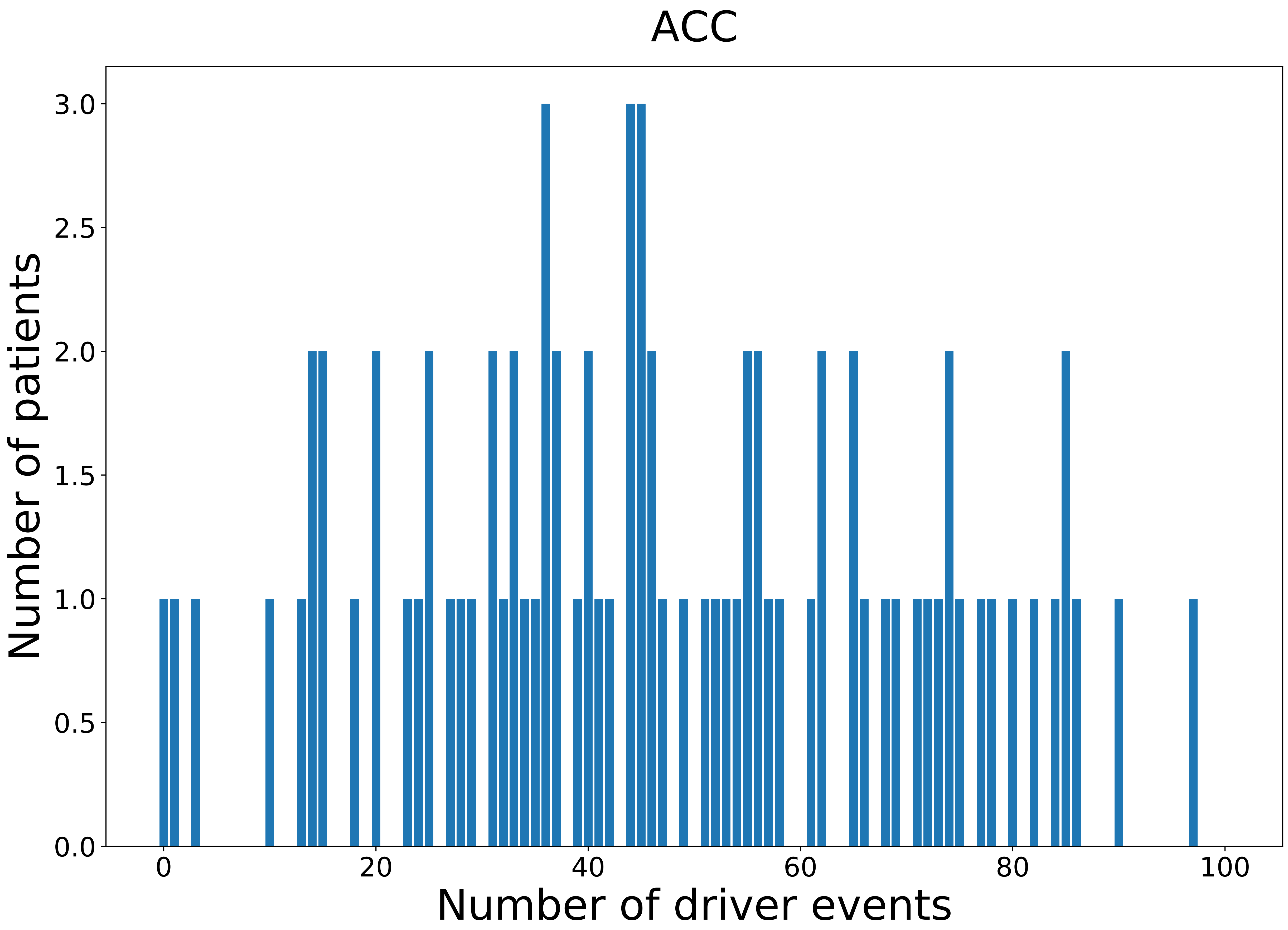

### 2021_8_22_13_42_ACC_FEMALE.pdf

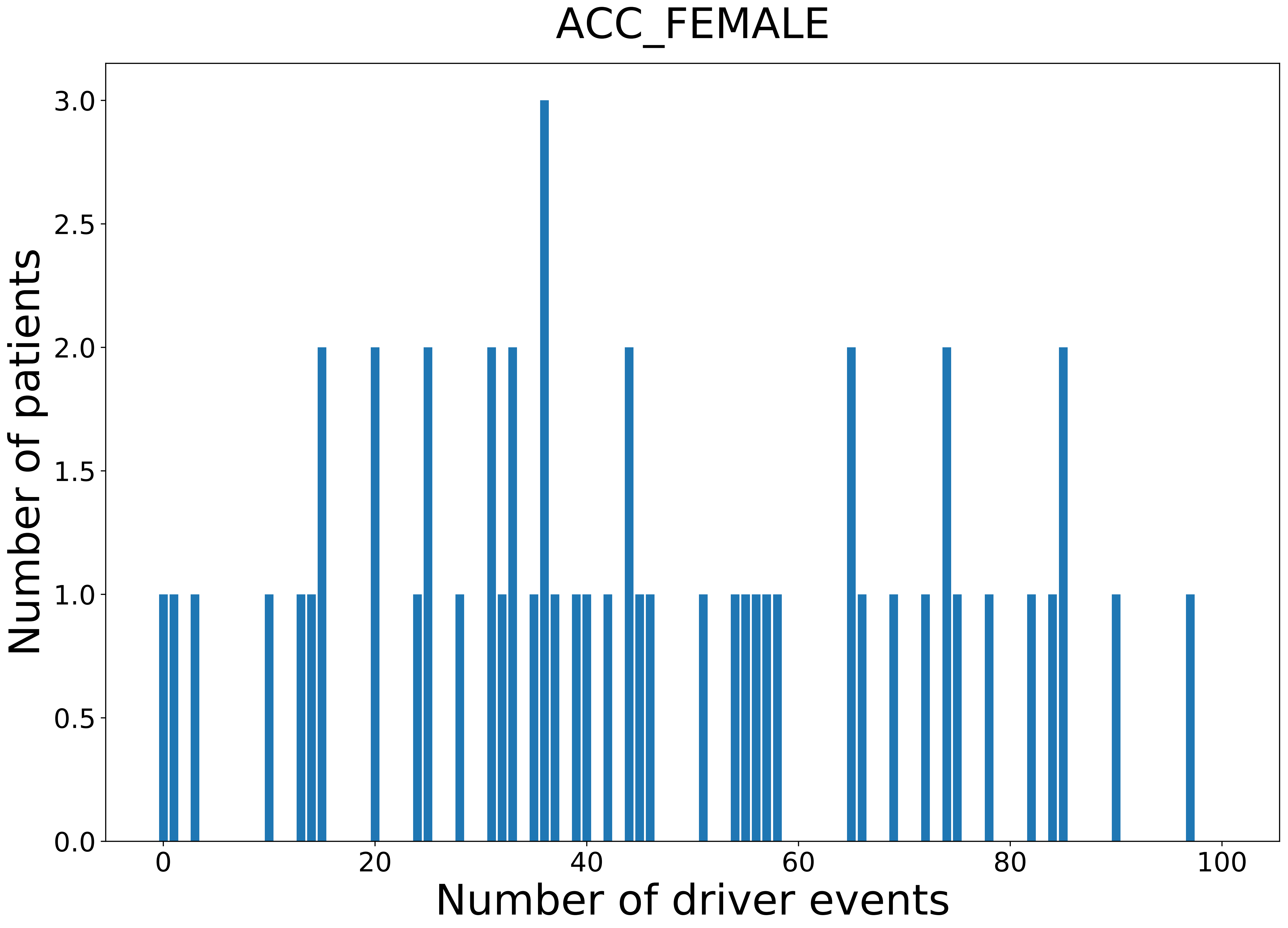

### 2021_8_22_13_42_ACC_MALE.pdf

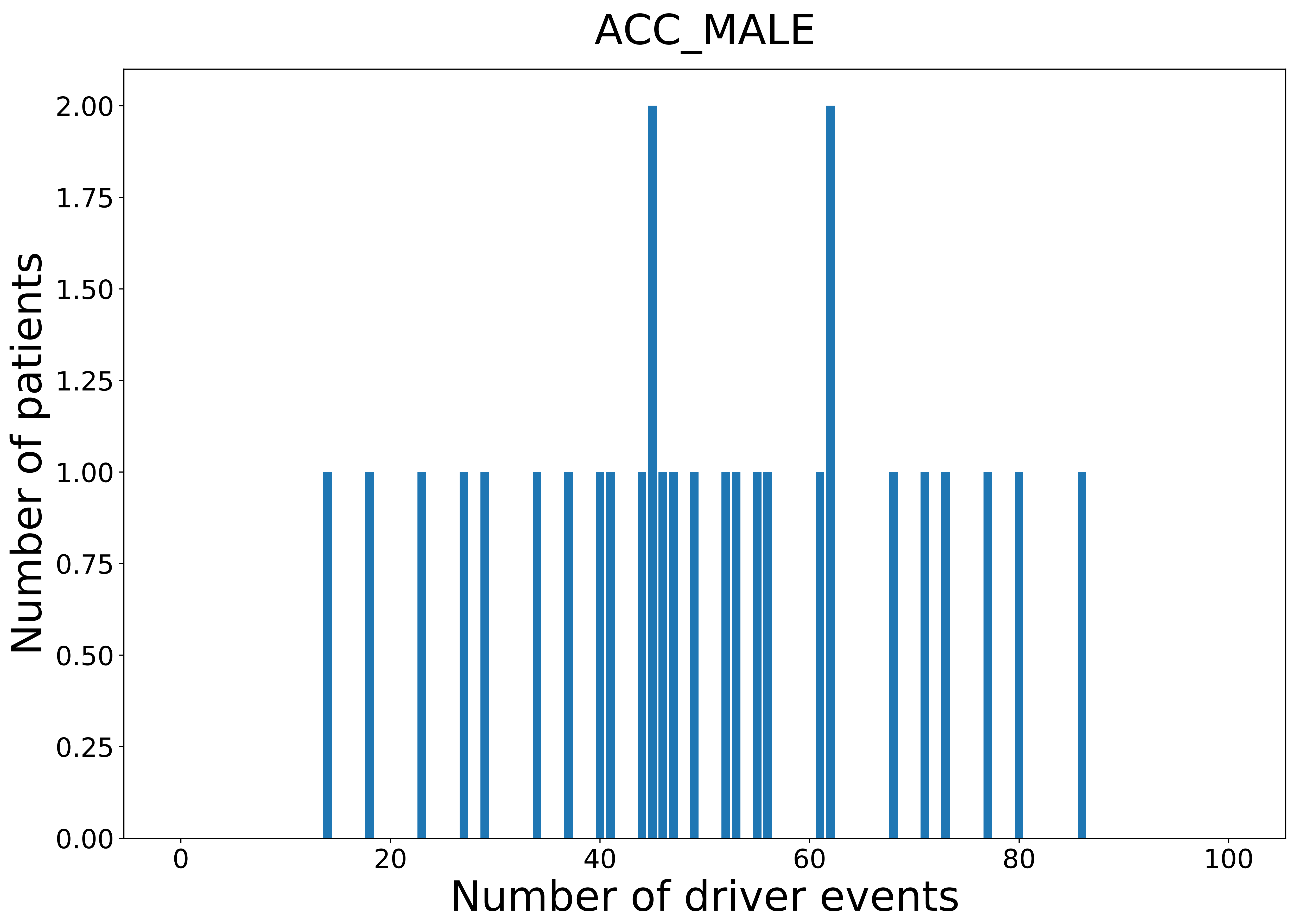

### 2021_8_22_13_42_BLCA.pdf

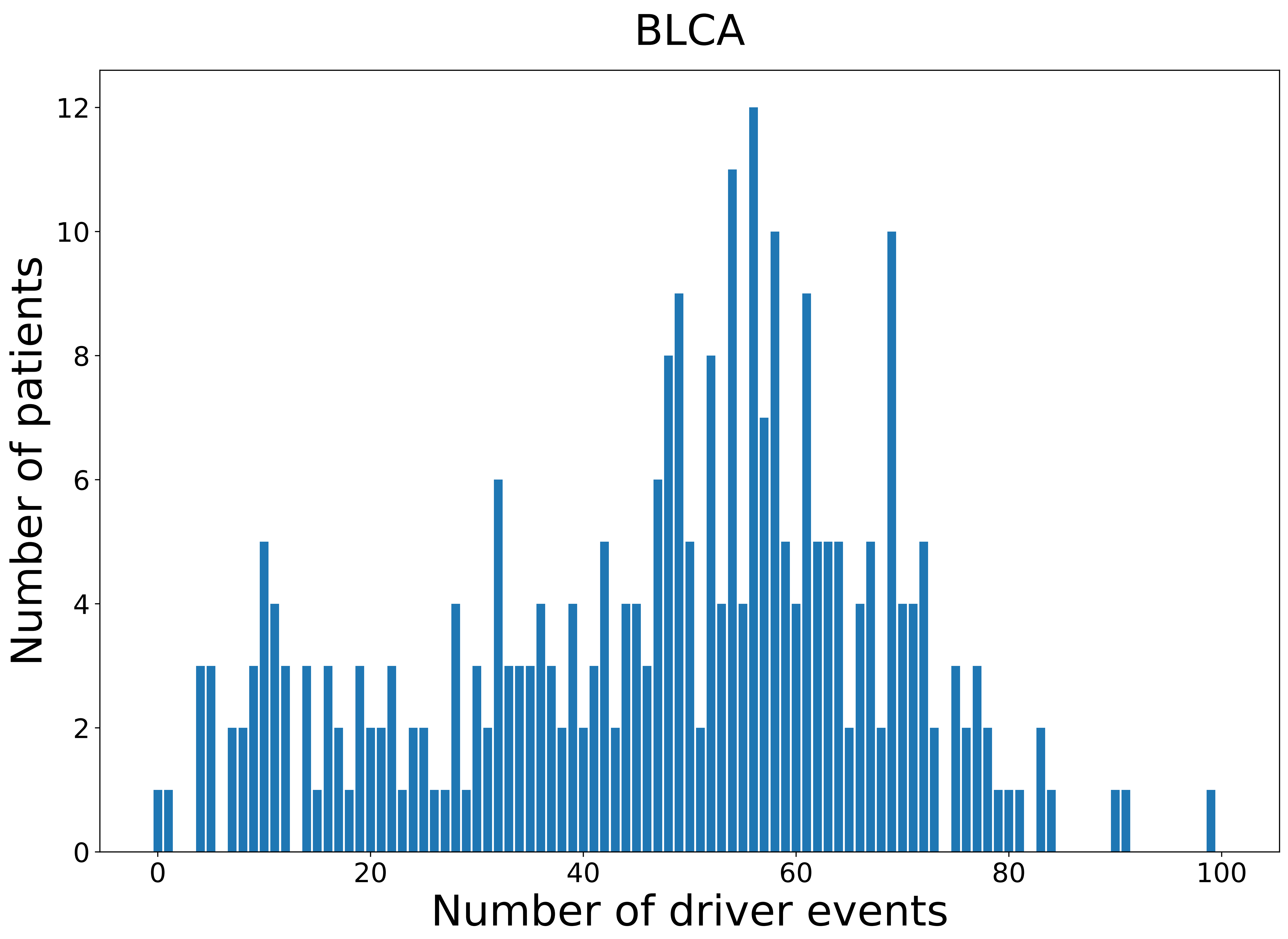

### 2021_8_22_13_42_BLCA_FEMALE.pdf

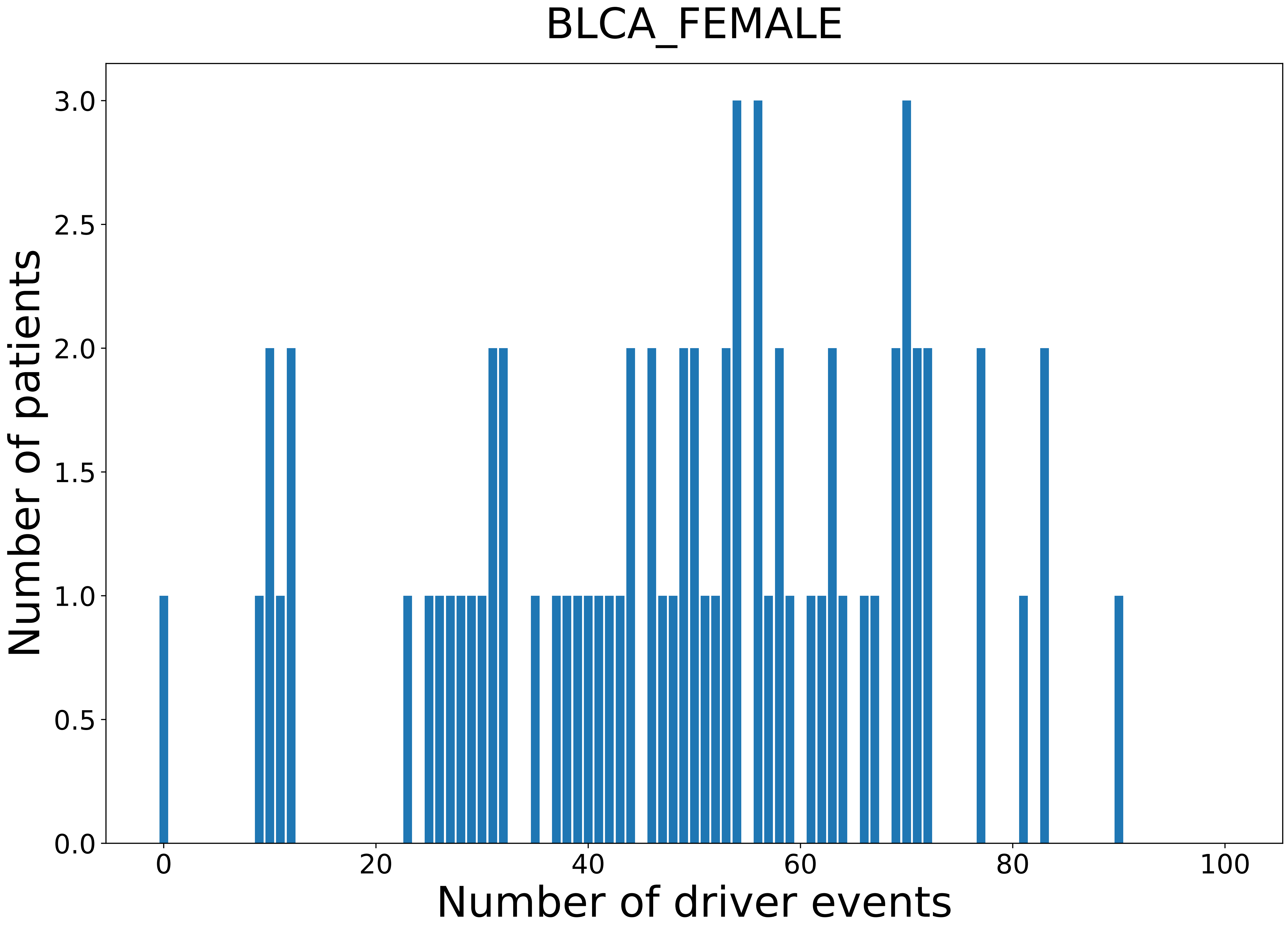

### 2021_8_22_13_42_BLCA_MALE.pdf

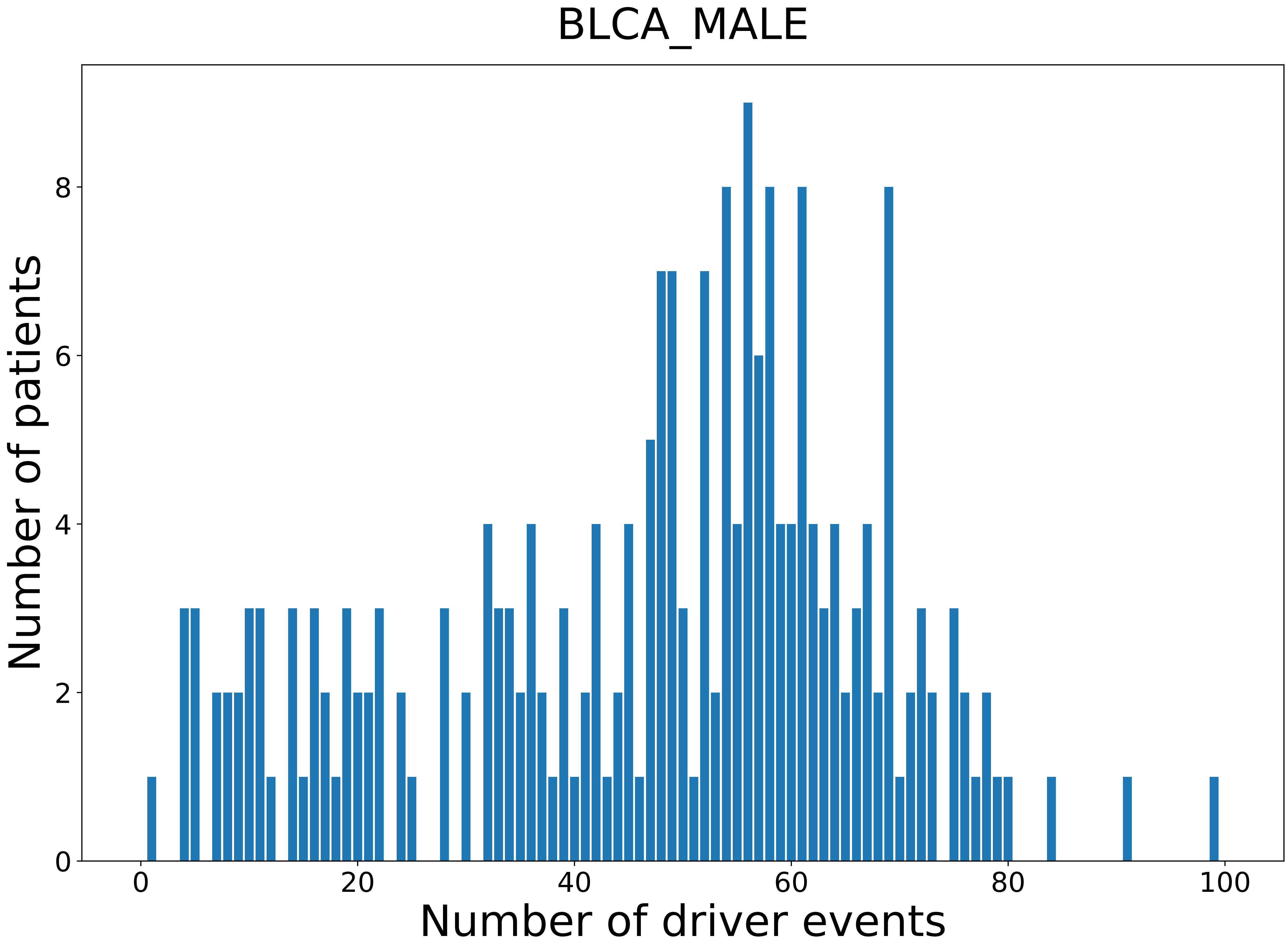

### 2021_8_22_13_42_BRCA.pdf

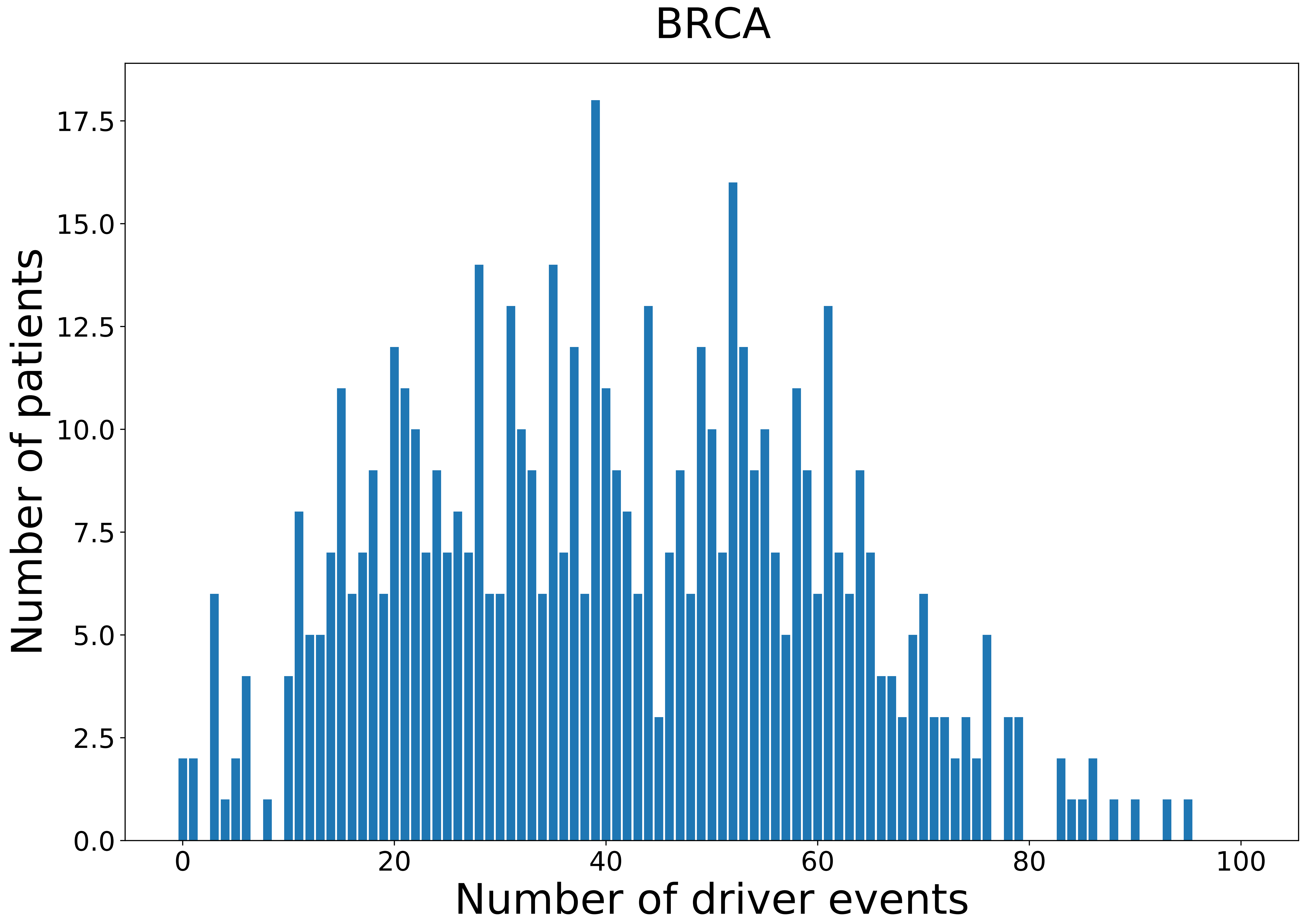

### 2021_8_22_13_42_BRCA_FEMALE.pdf

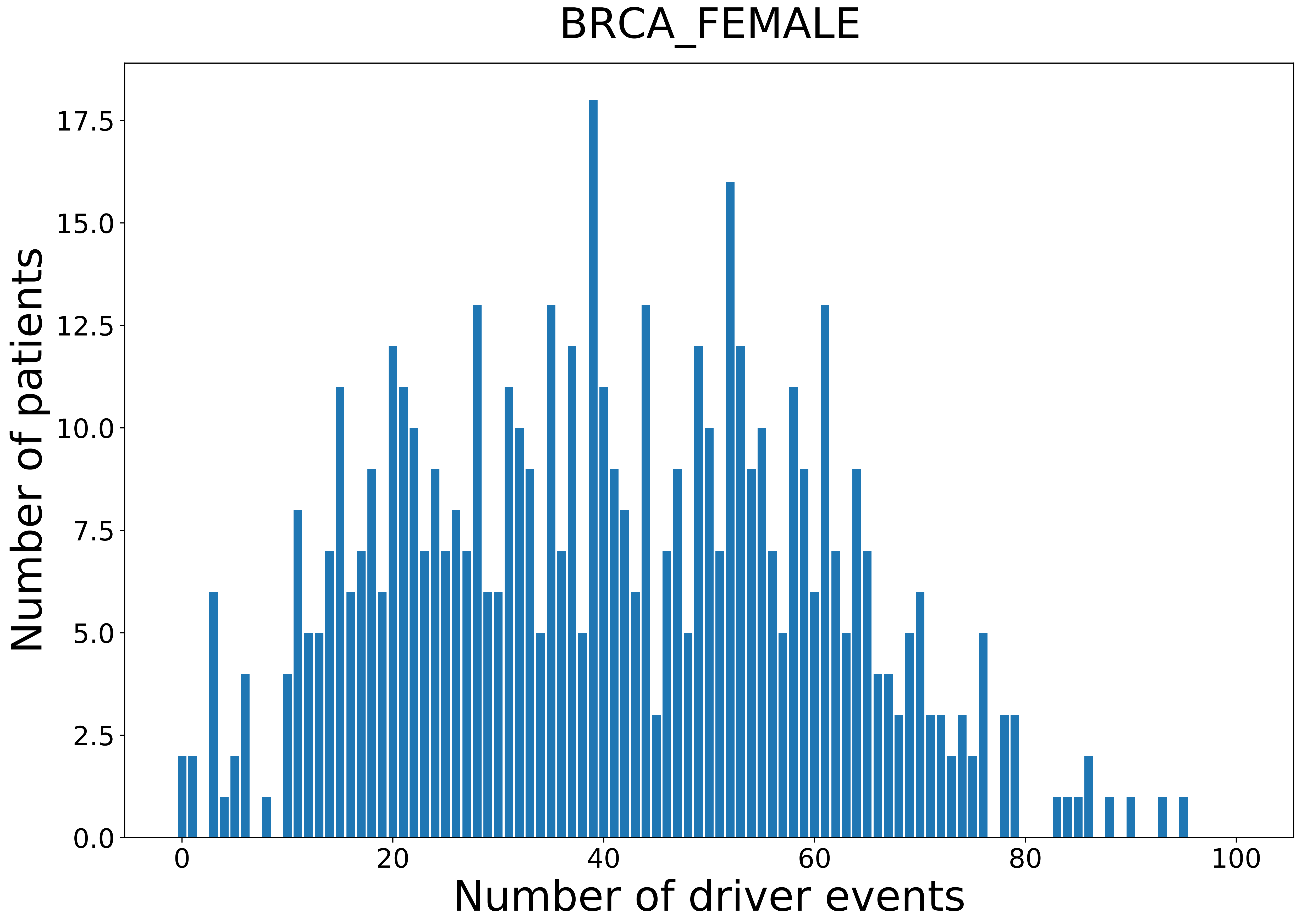

### 2021_8_22_13_42_CESC_FEMALE.pdf

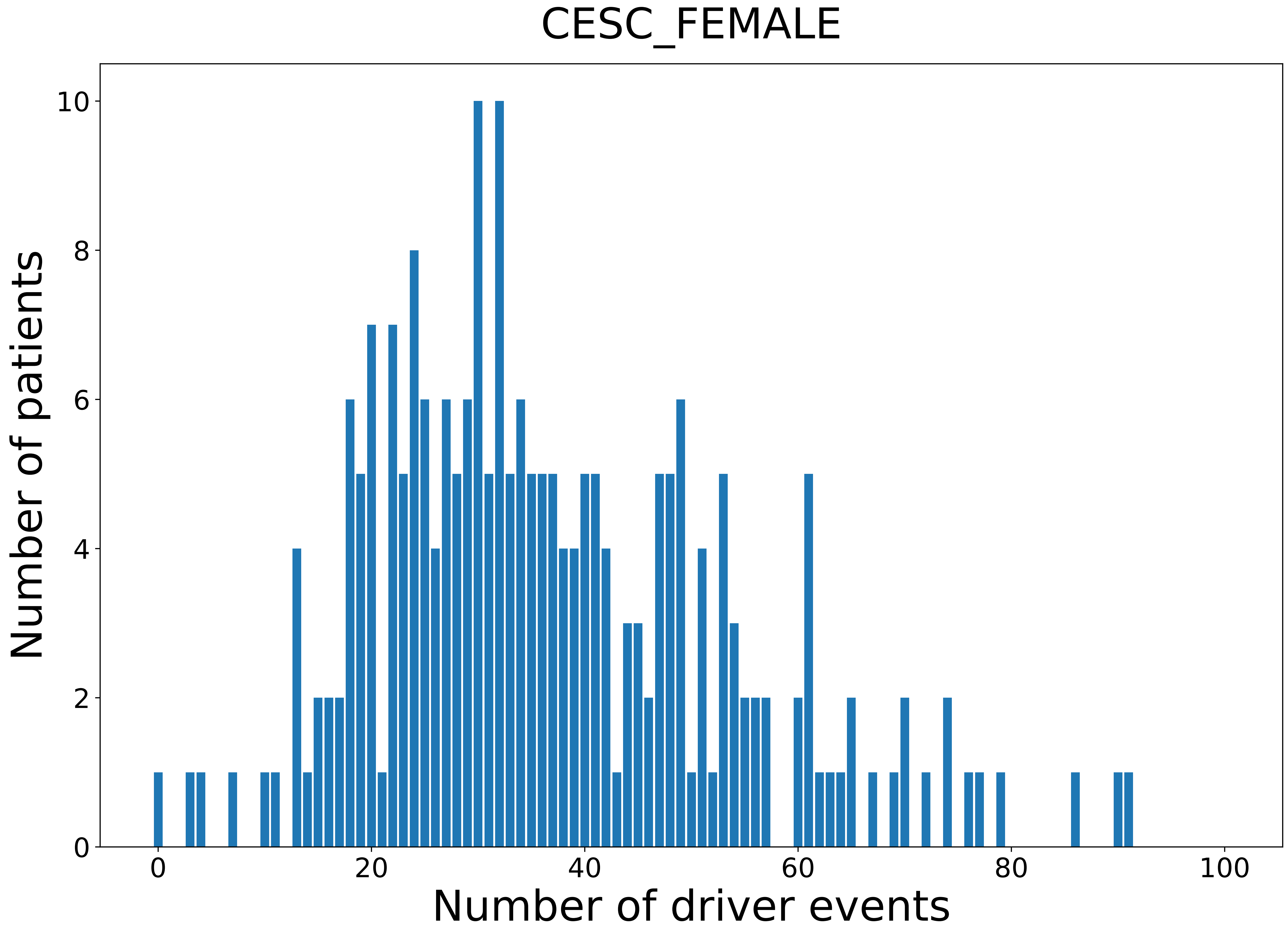

### 2021_8_22_13_42_CHOL.pdf

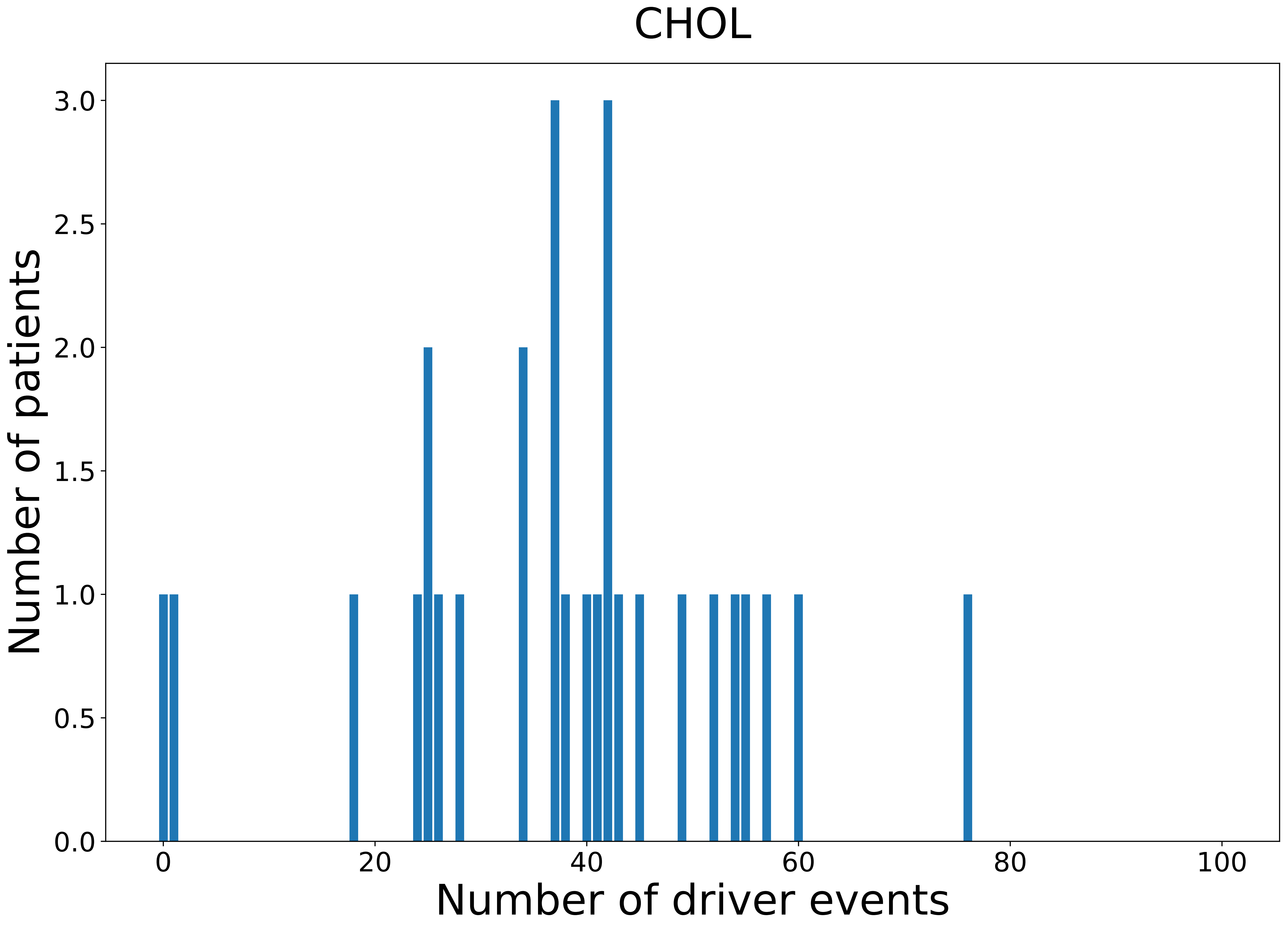

### 2021_8_22_13_42_CHOL_FEMALE.pdf

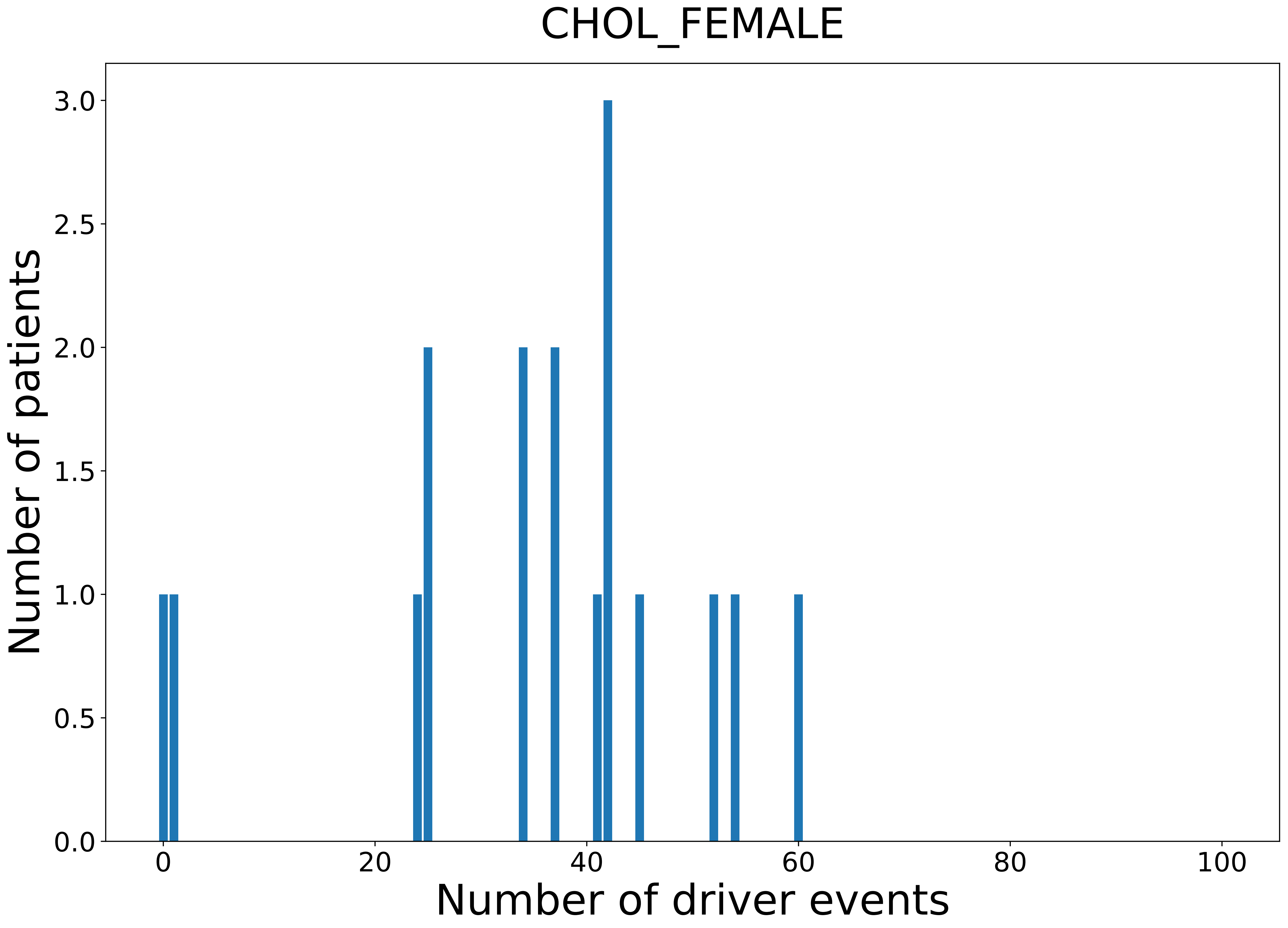

### 2021_8_22_13_42_CHOL_MALE.pdf

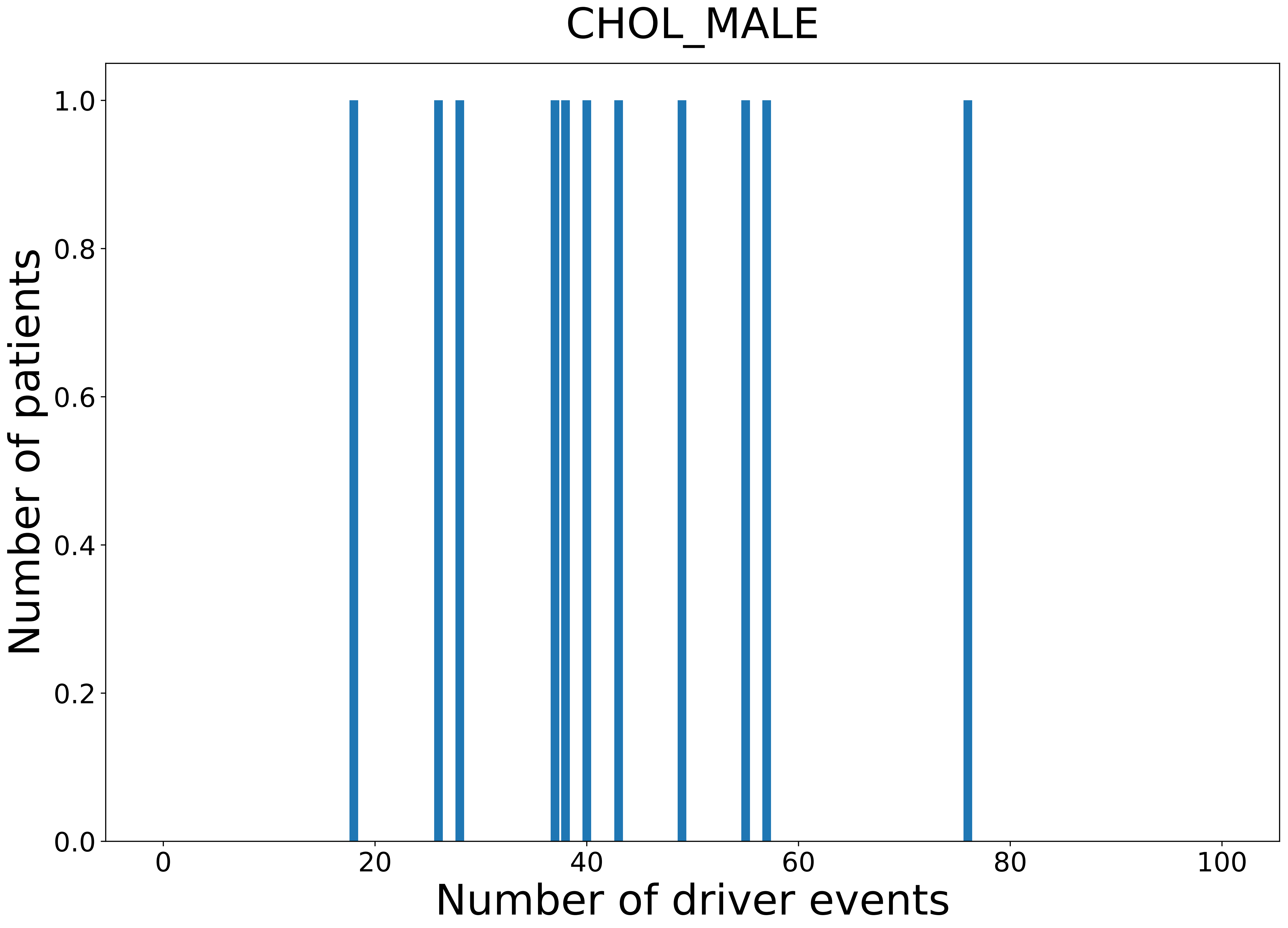

### 2021_8_22_13_42_COAD.pdf

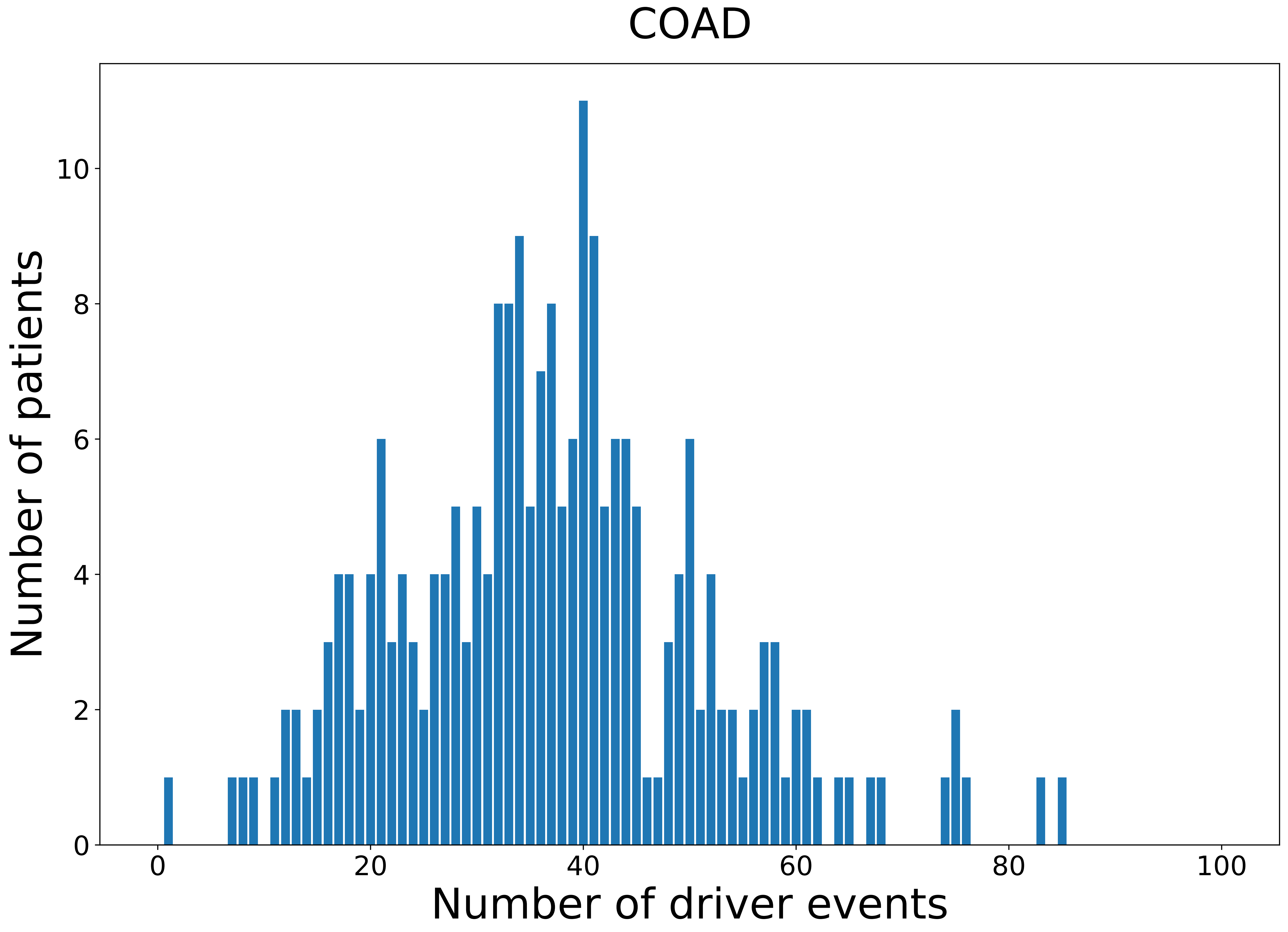

### 2021_8_22_13_42_COAD_FEMALE.pdf

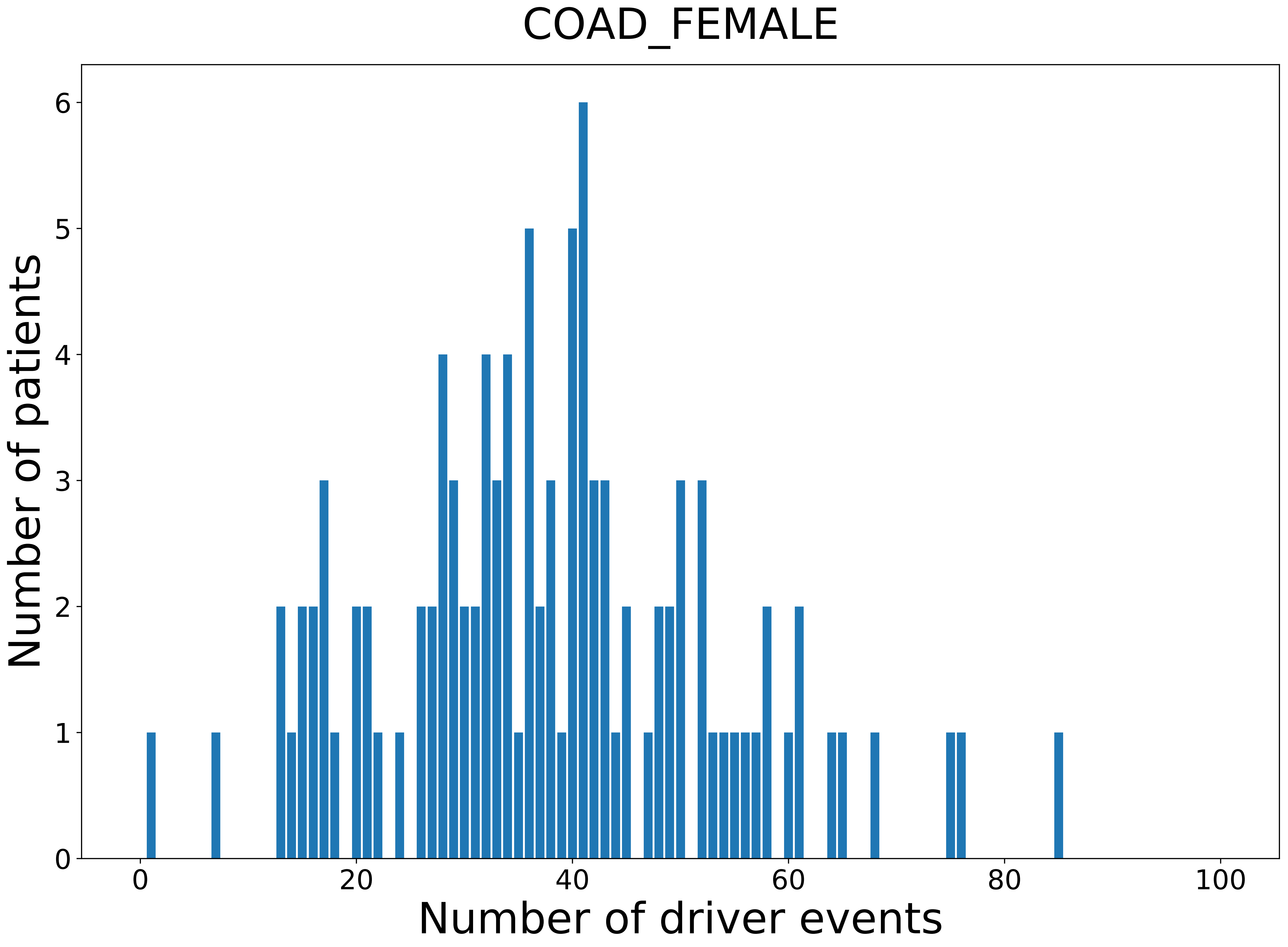

### 2021_8_22_13_42_COAD_MALE.pdf

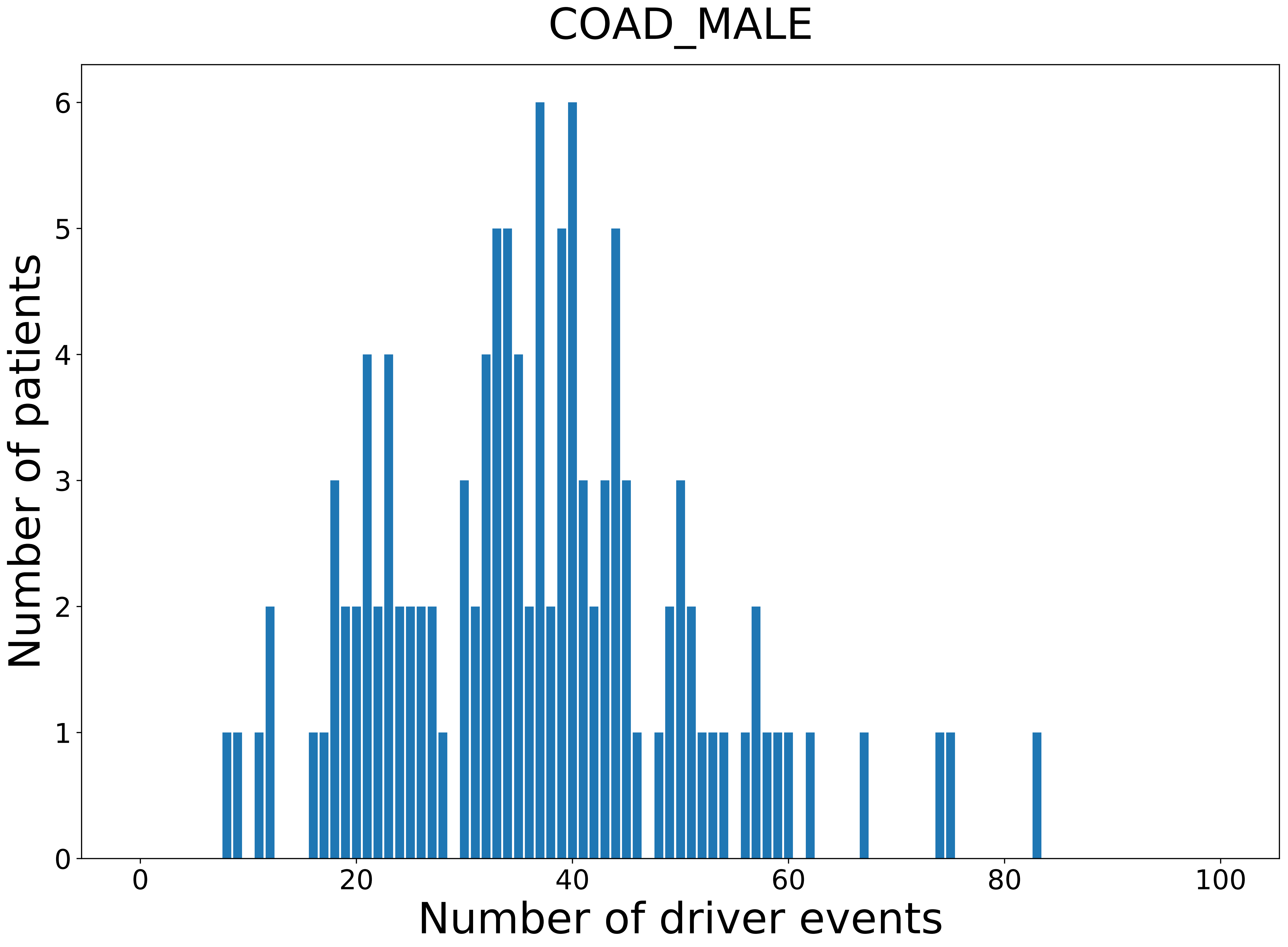

### 2021_8_22_13_42_distribution_age.pdf

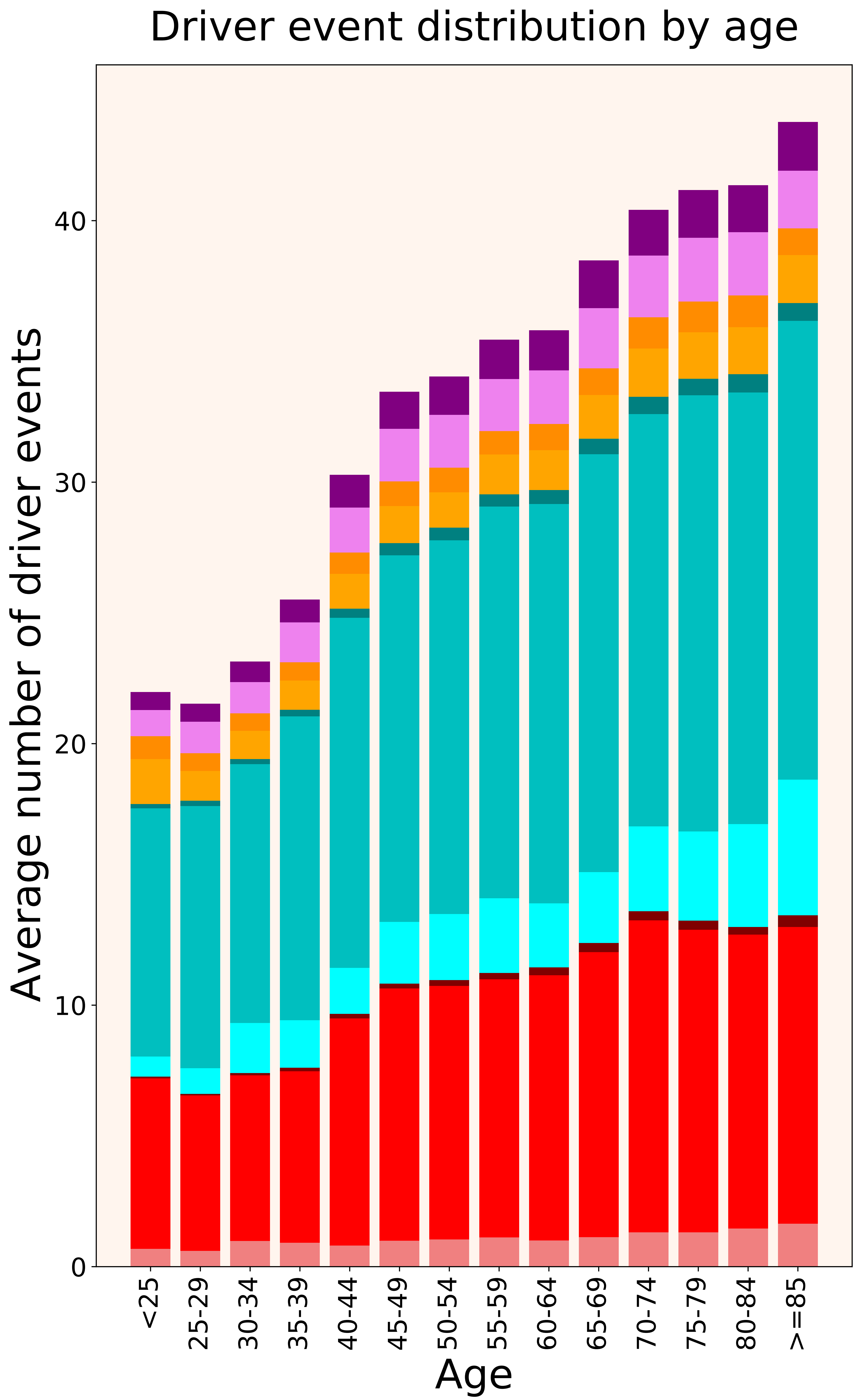

### 2021_8_22_13_42_distribution_age_females.pdf

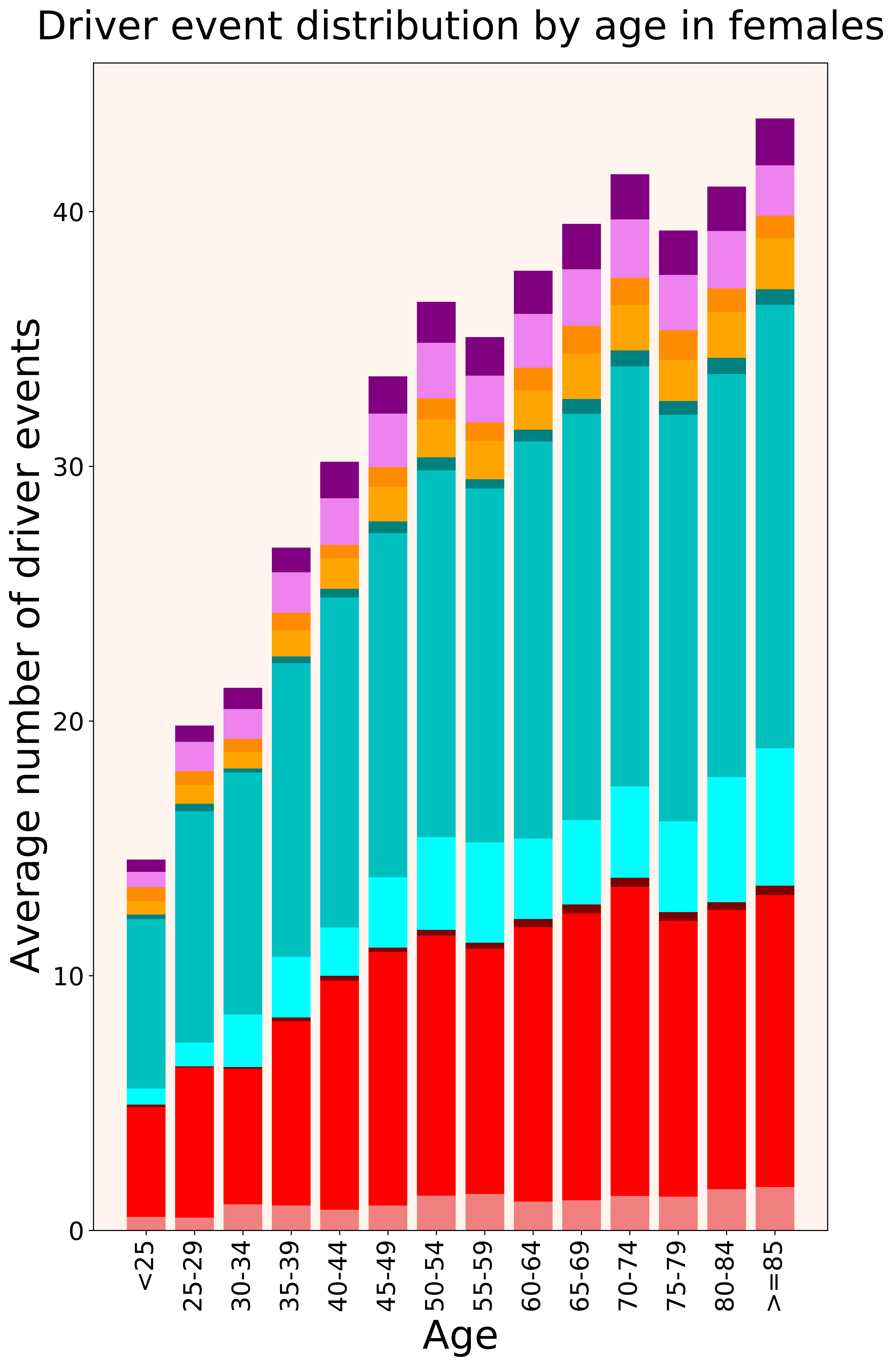

### 2021_8_22_13_42_distribution_age_males.pdf

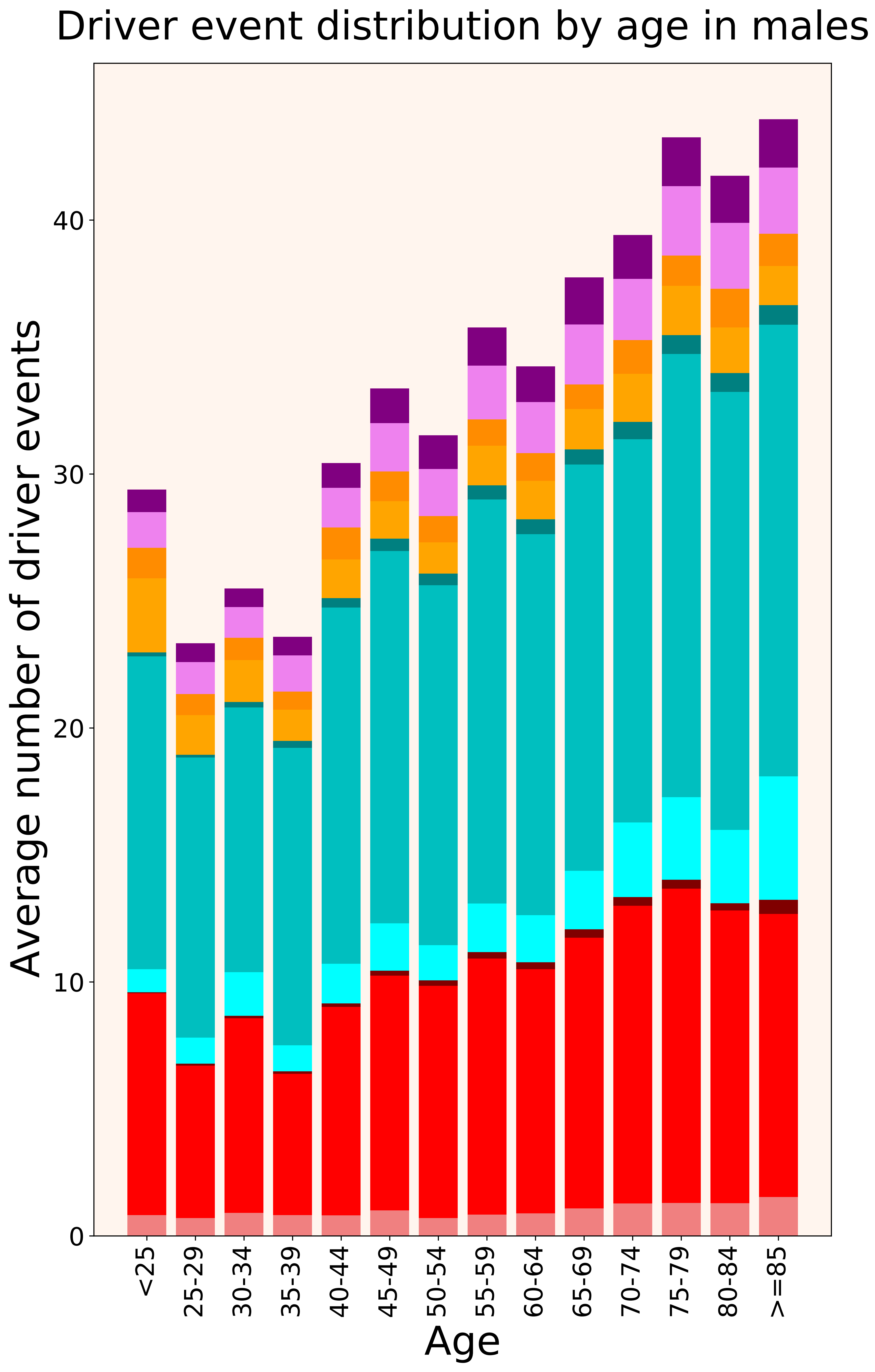

### 2021_8_22_13_42_distribution_cohorts.pdf

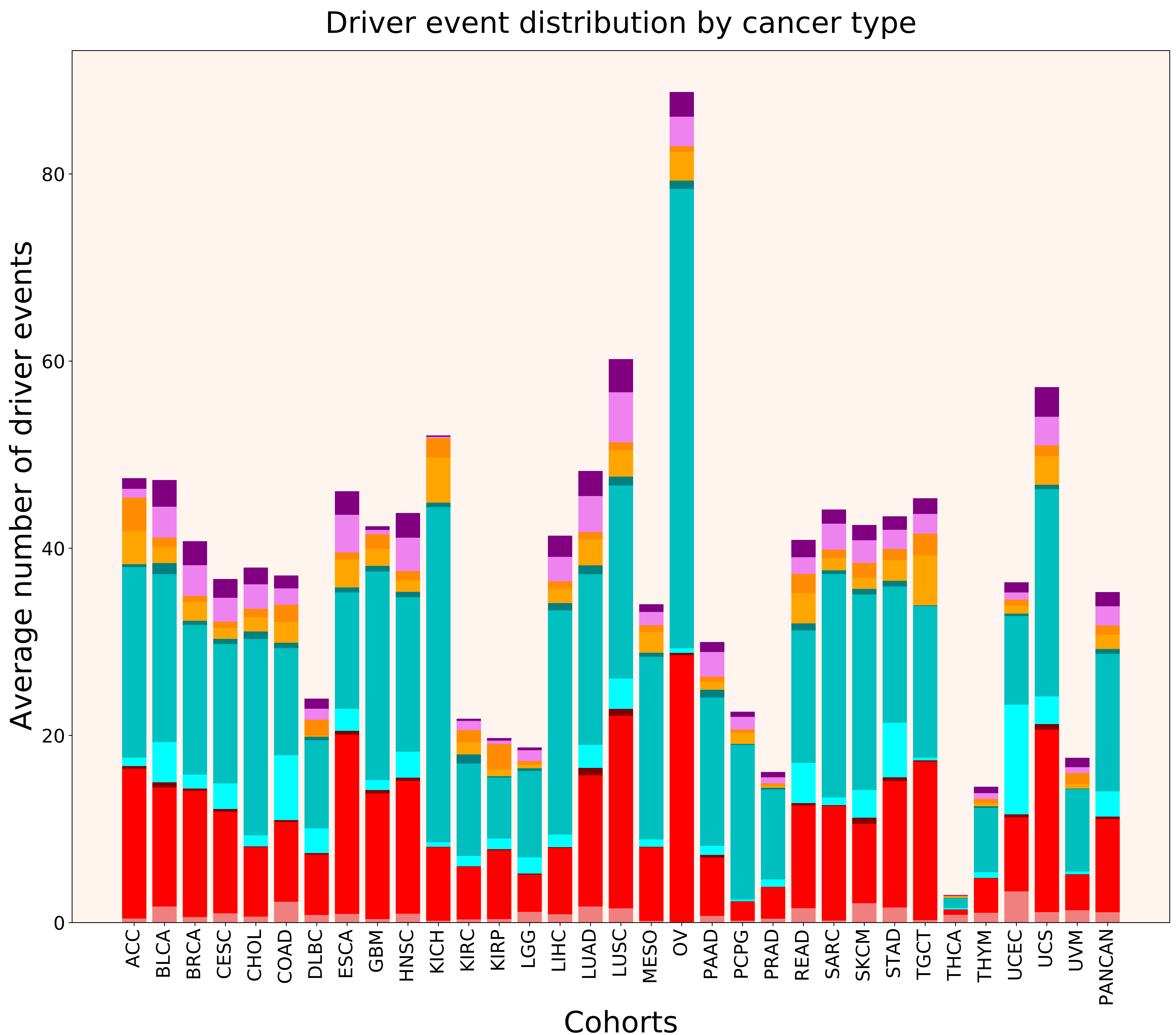

### 2021_8_22_13_42_distribution_cohorts_females.pdf

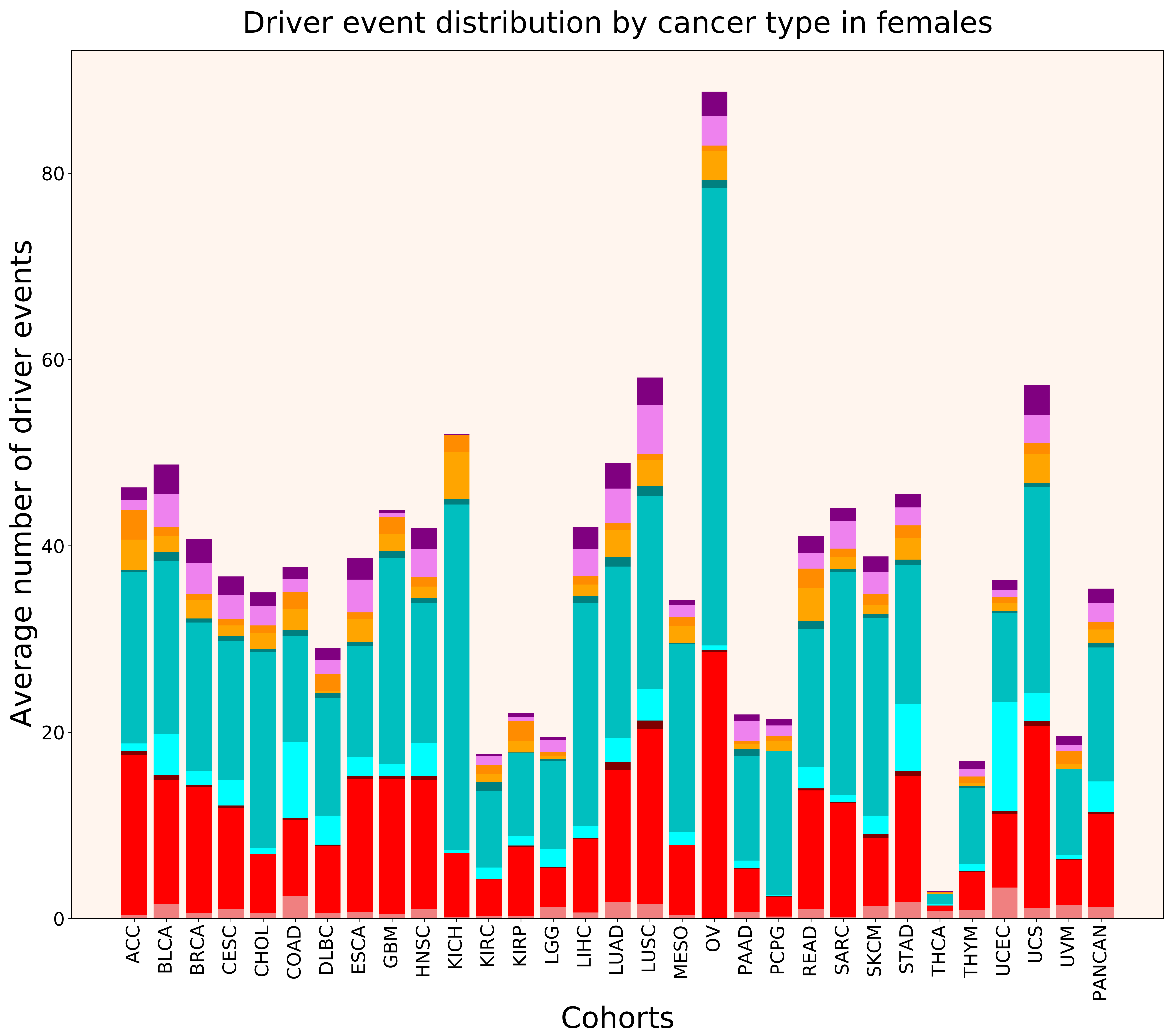

### 2021_8_22_13_42_distribution_cohorts_males.pdf

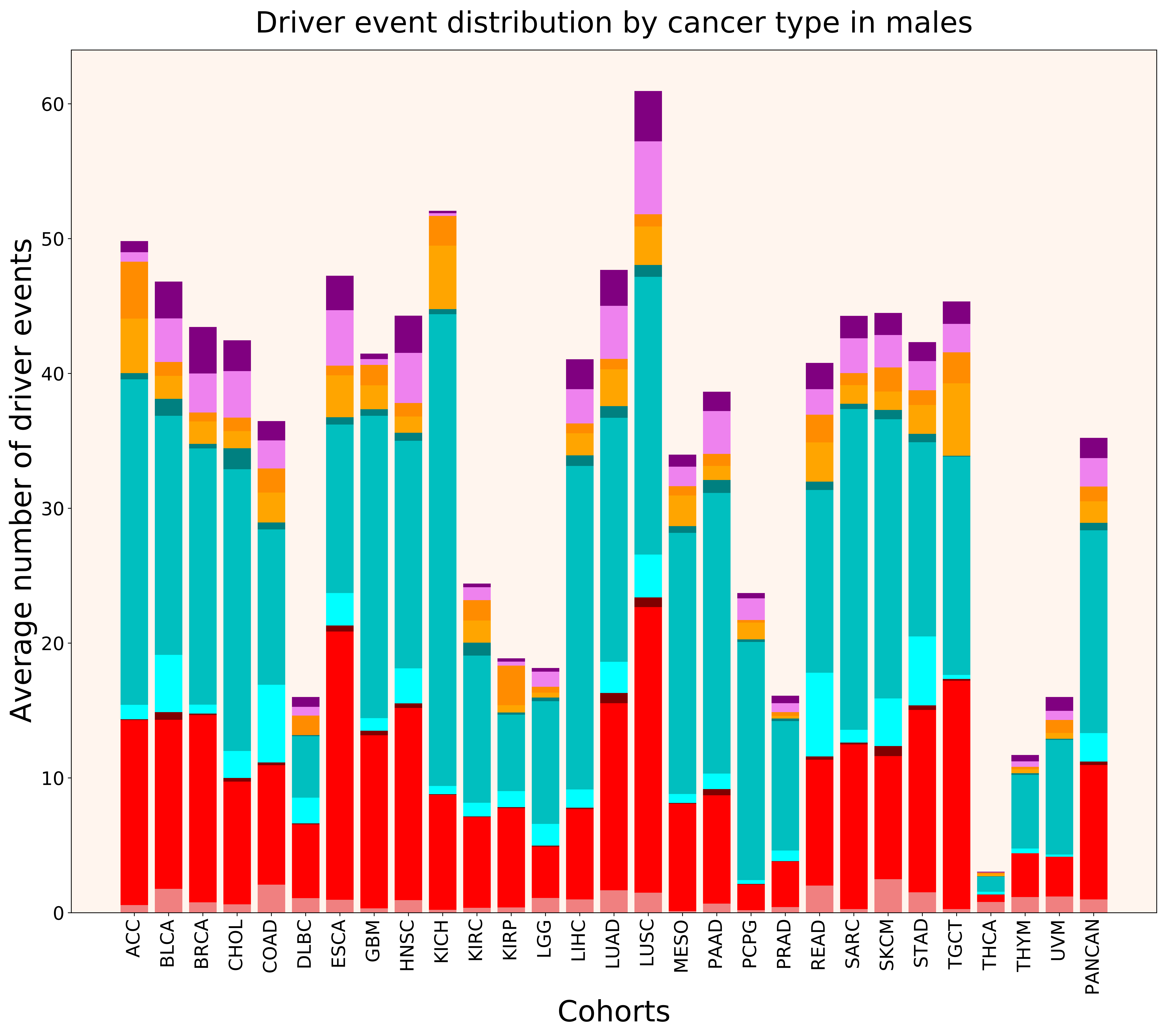

### 2021_8_22_13_42_distribution_events_detailed.pdf

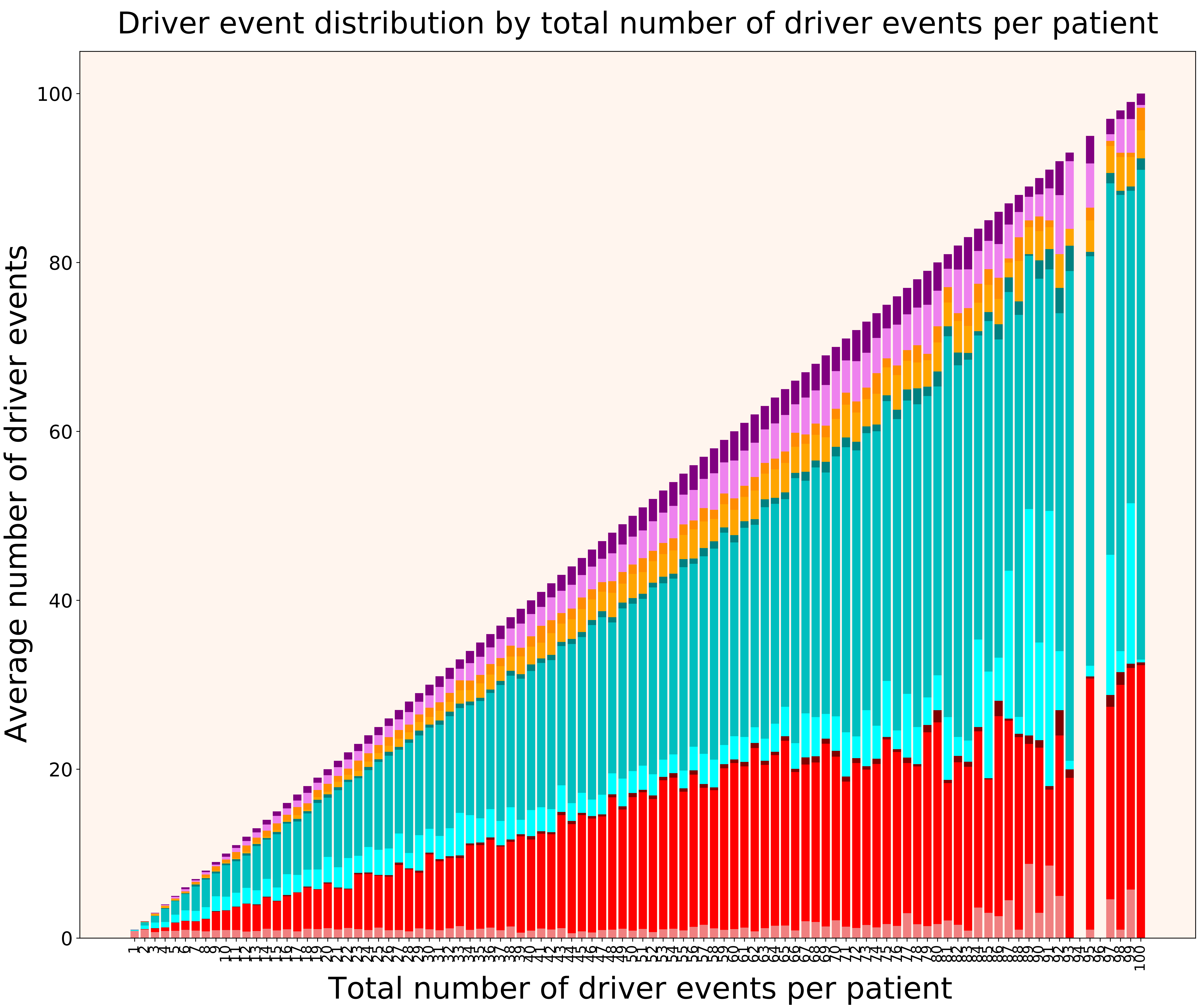

### 2021_8_22_13_42_distribution_events_detailed_females.pdf

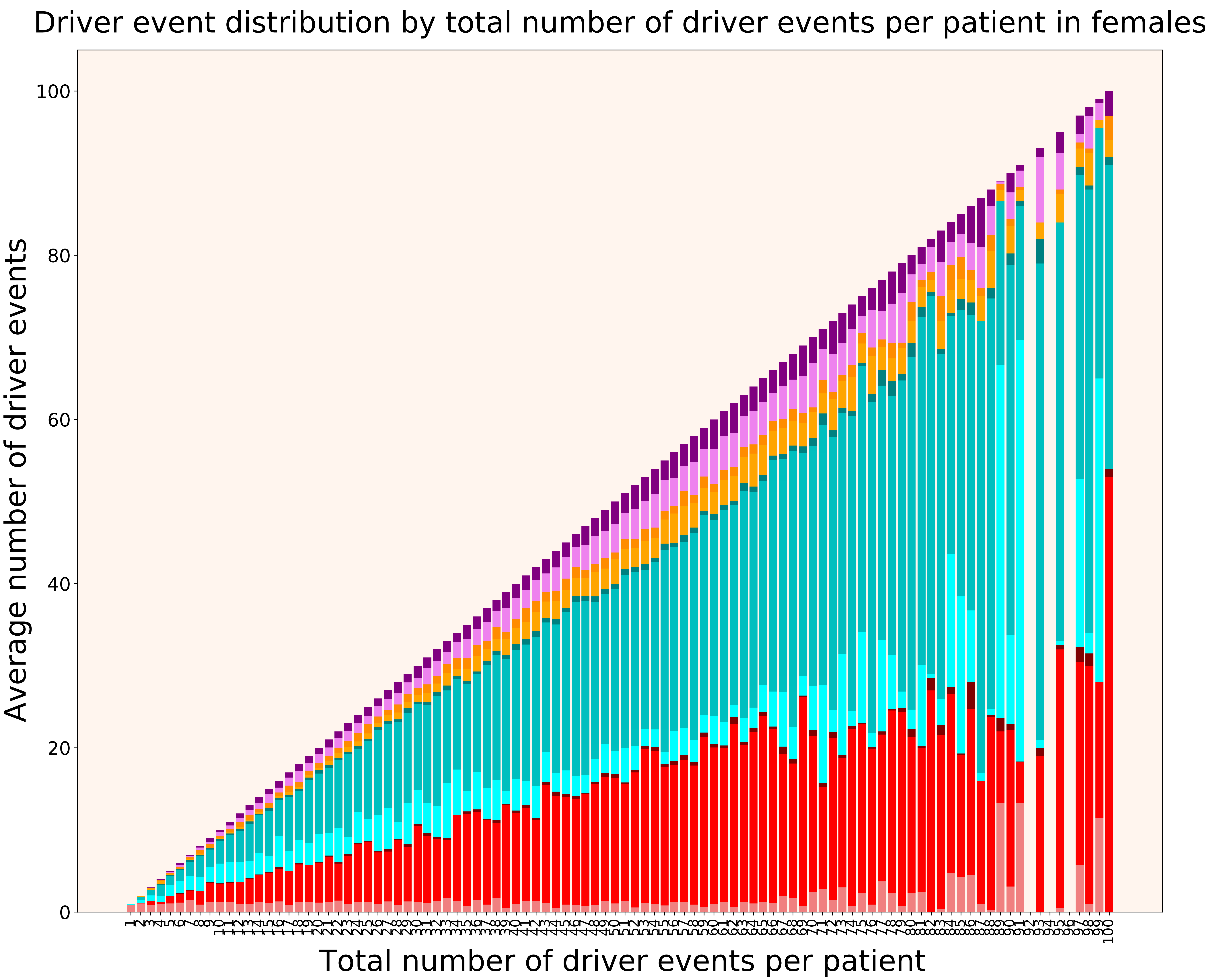

### 2021_8_22_13_42_distribution_events_detailed_males.pdf

#### Driver event distribution by total number of driver events per patient in males

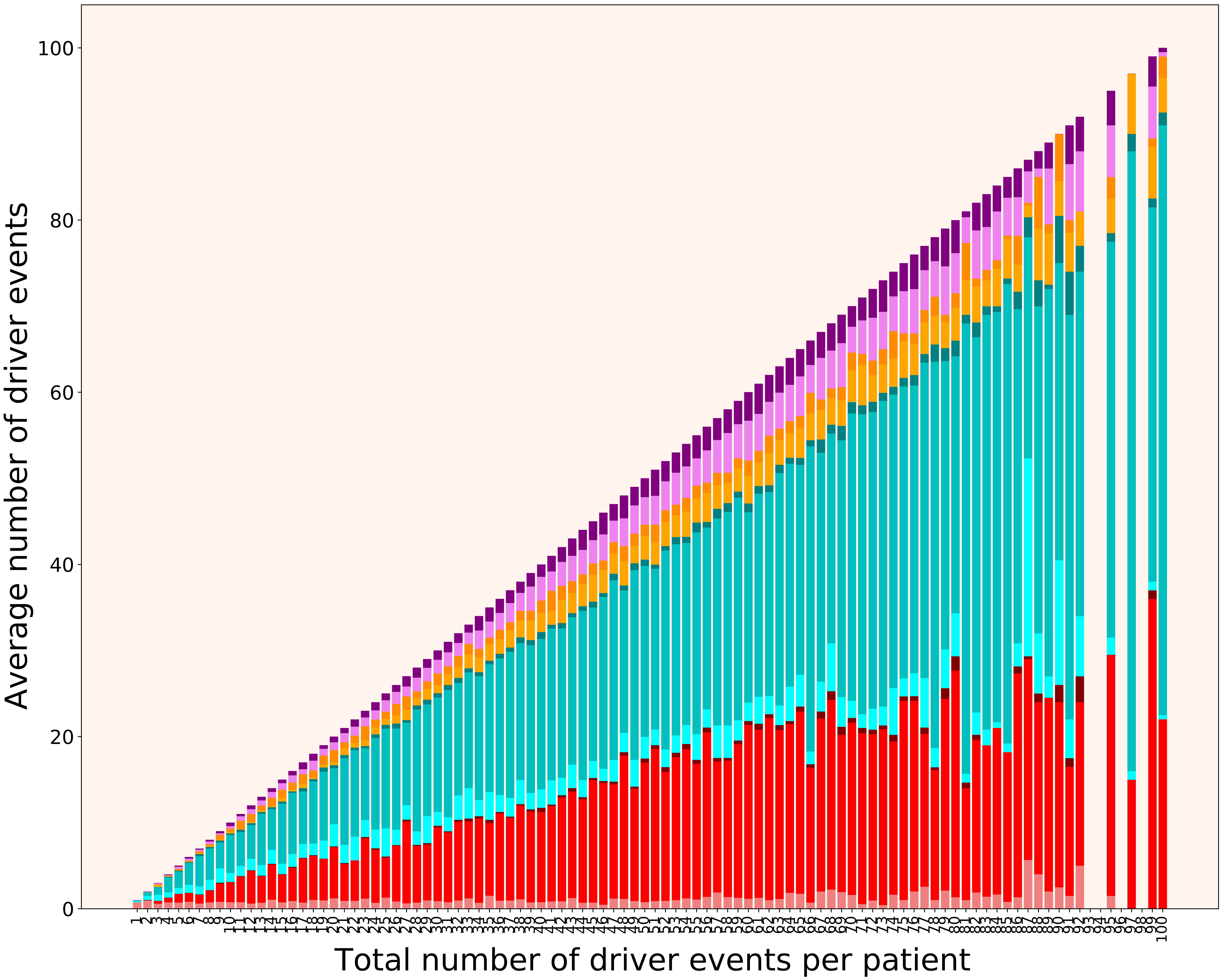

### 2021_8_22_13_42_distribution_gender.pdf

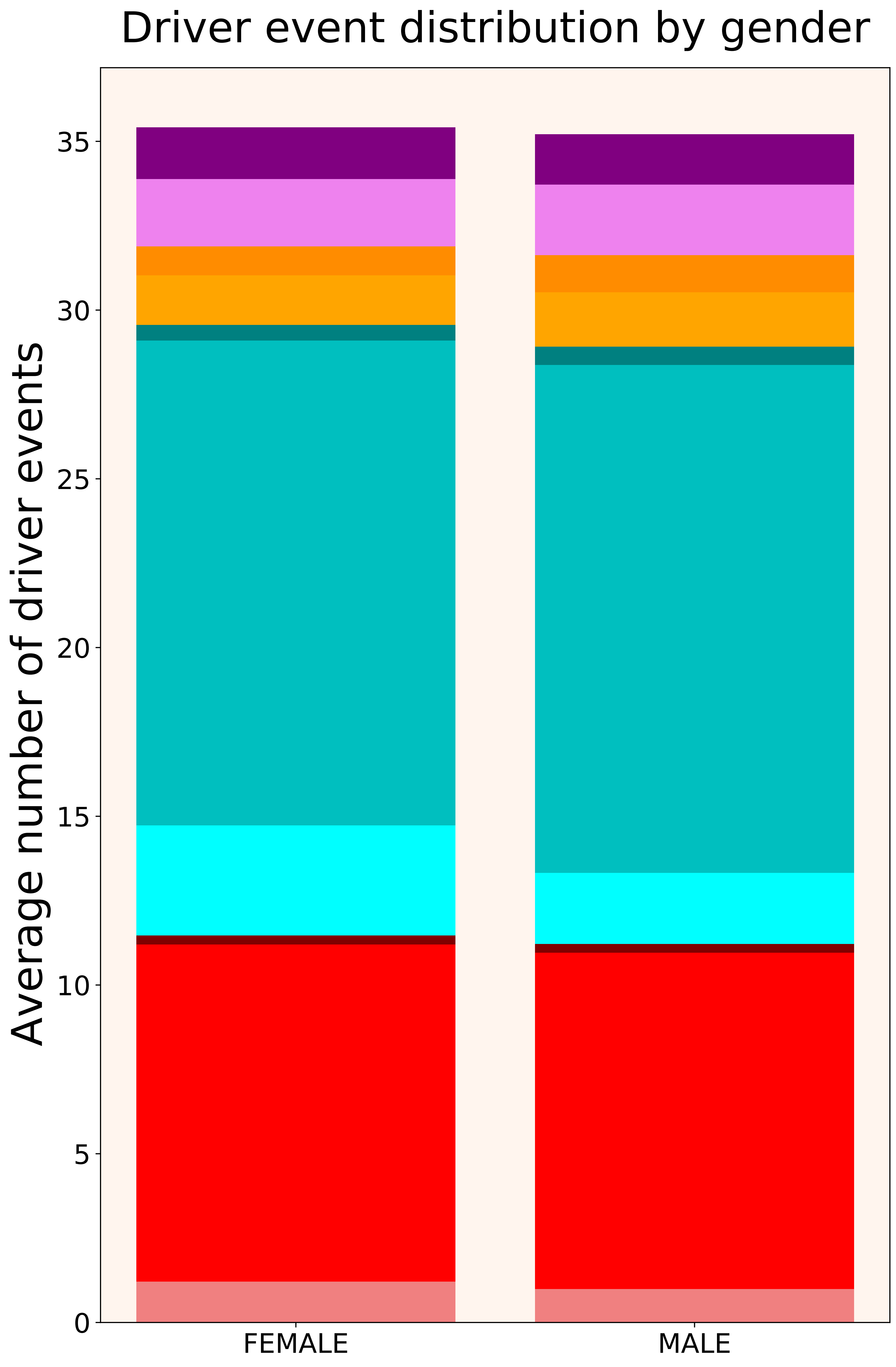

### 2021_8_22_13_42_distribution_stages.pdf

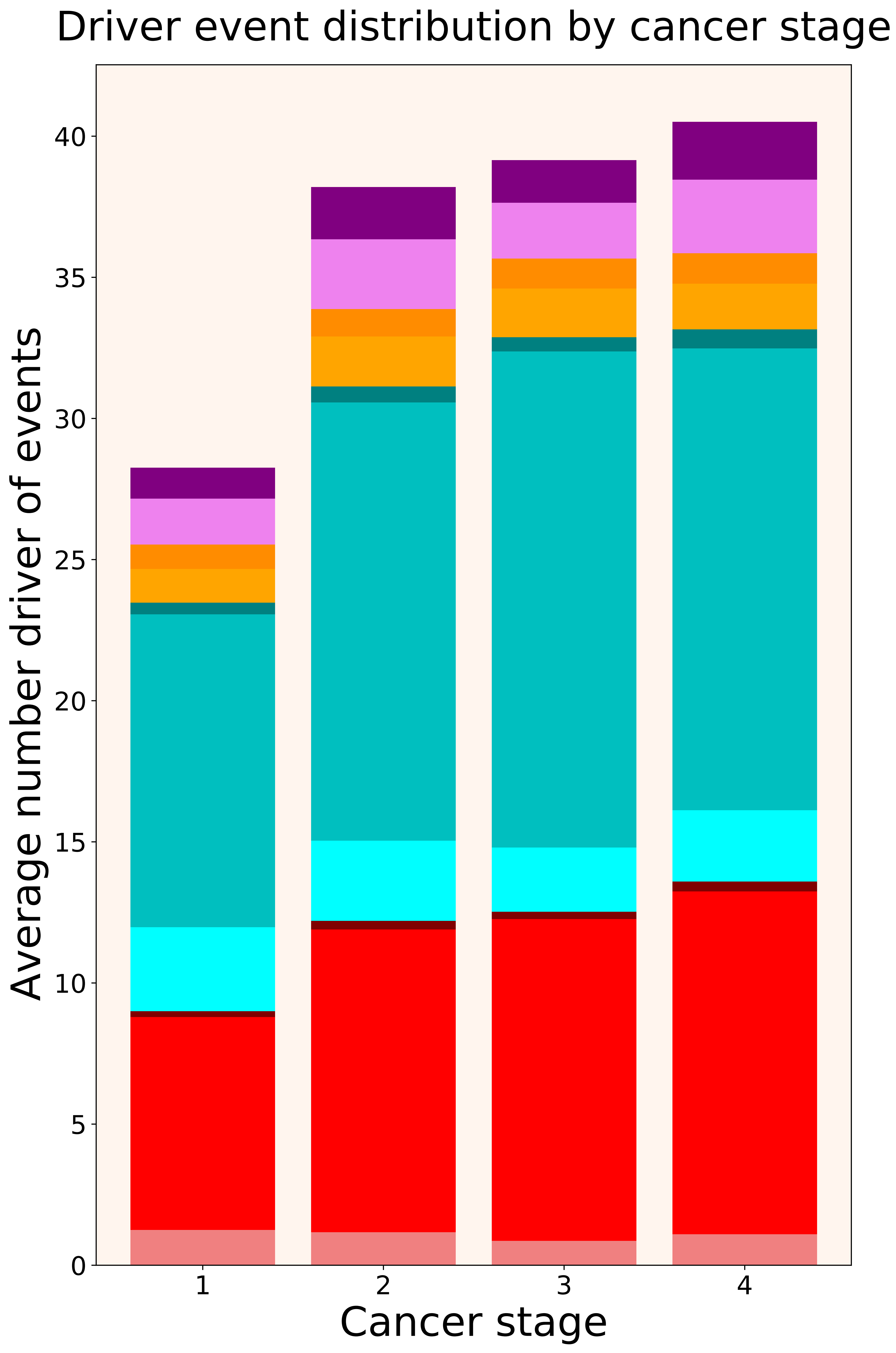

### 2021_8_22_13_42_distribution_stages_females.pdf

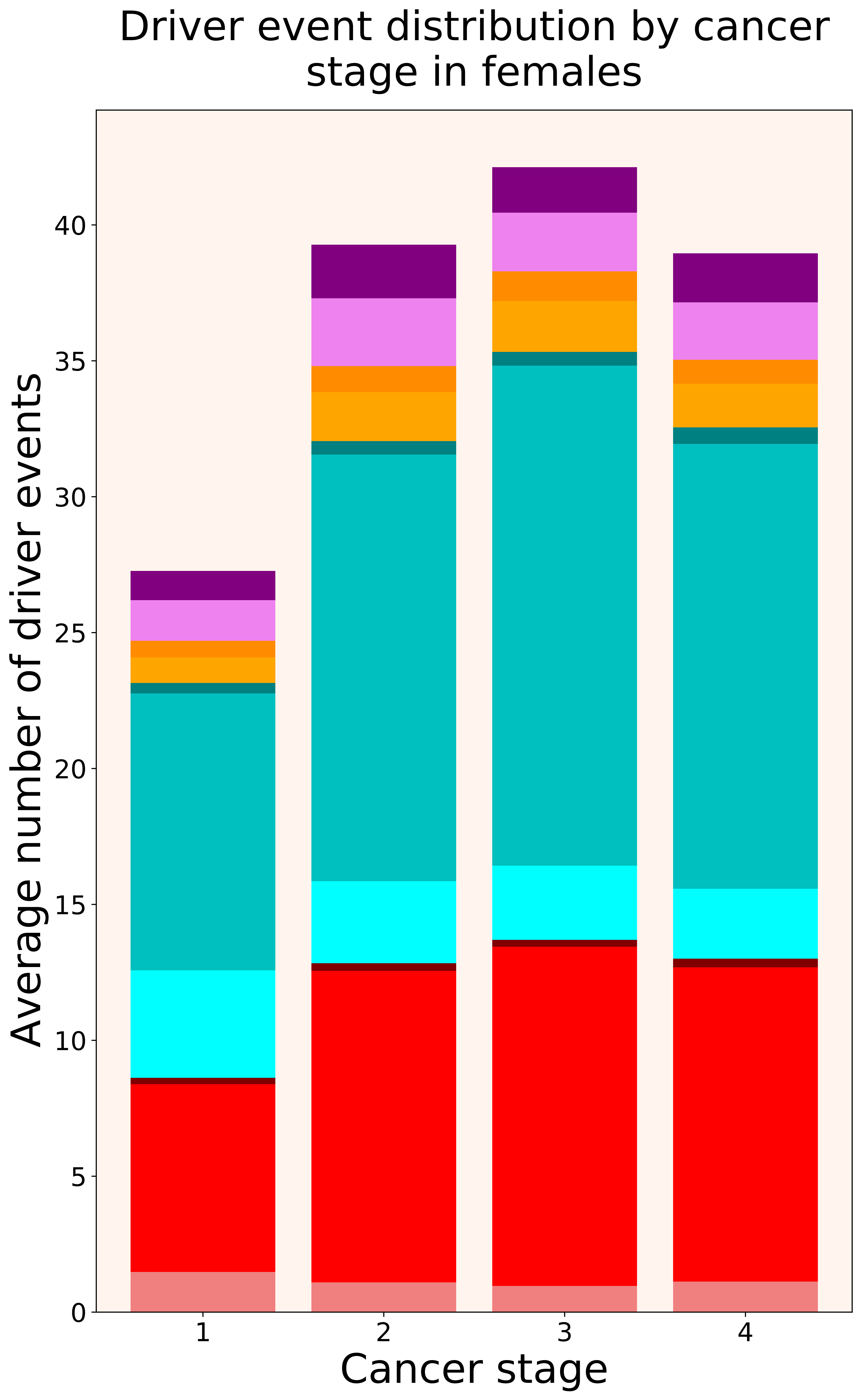

### 2021_8_22_13_42_distribution_stages_males.pdf

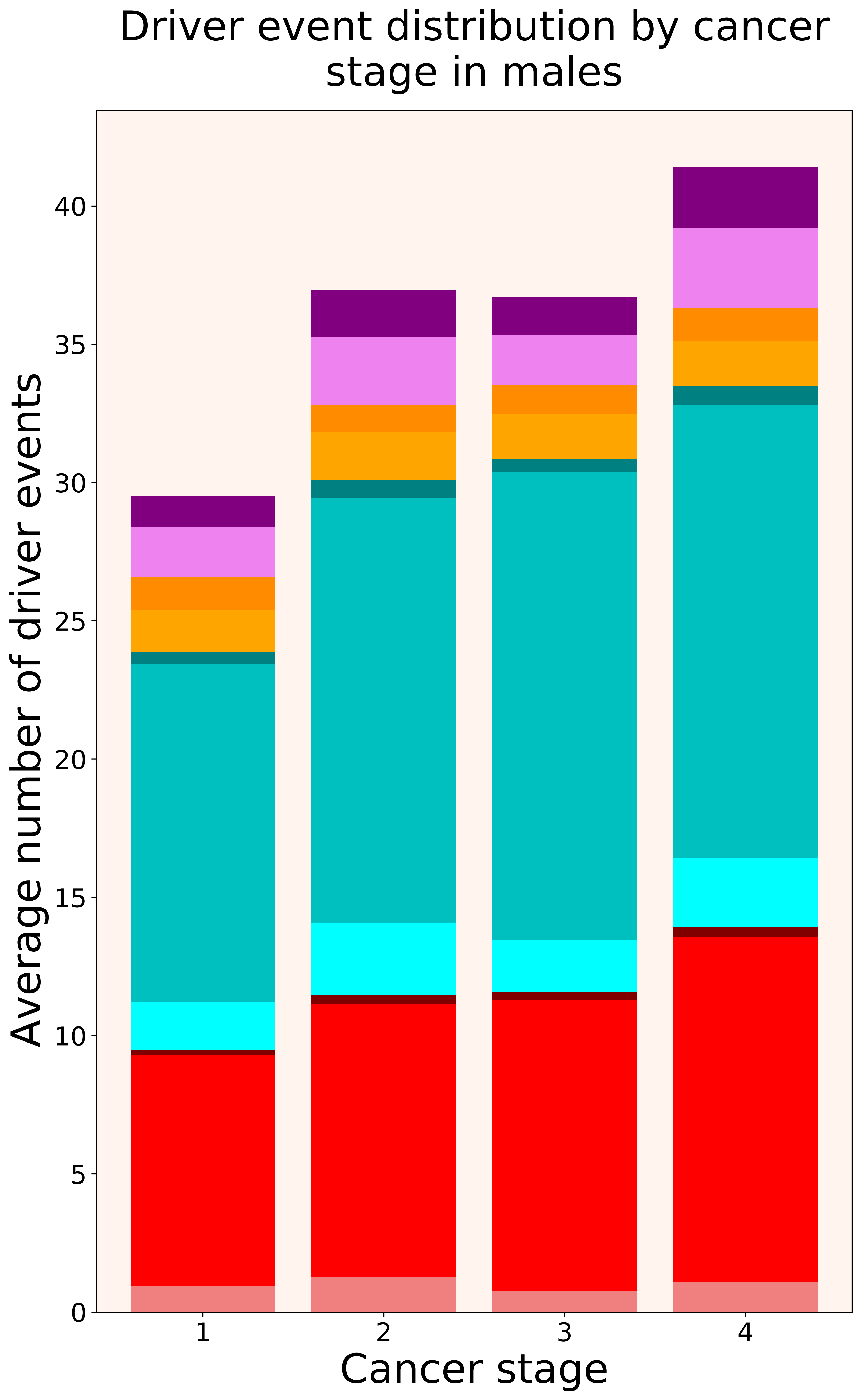

### 2021_8_22_13_42_DLBC.pdf

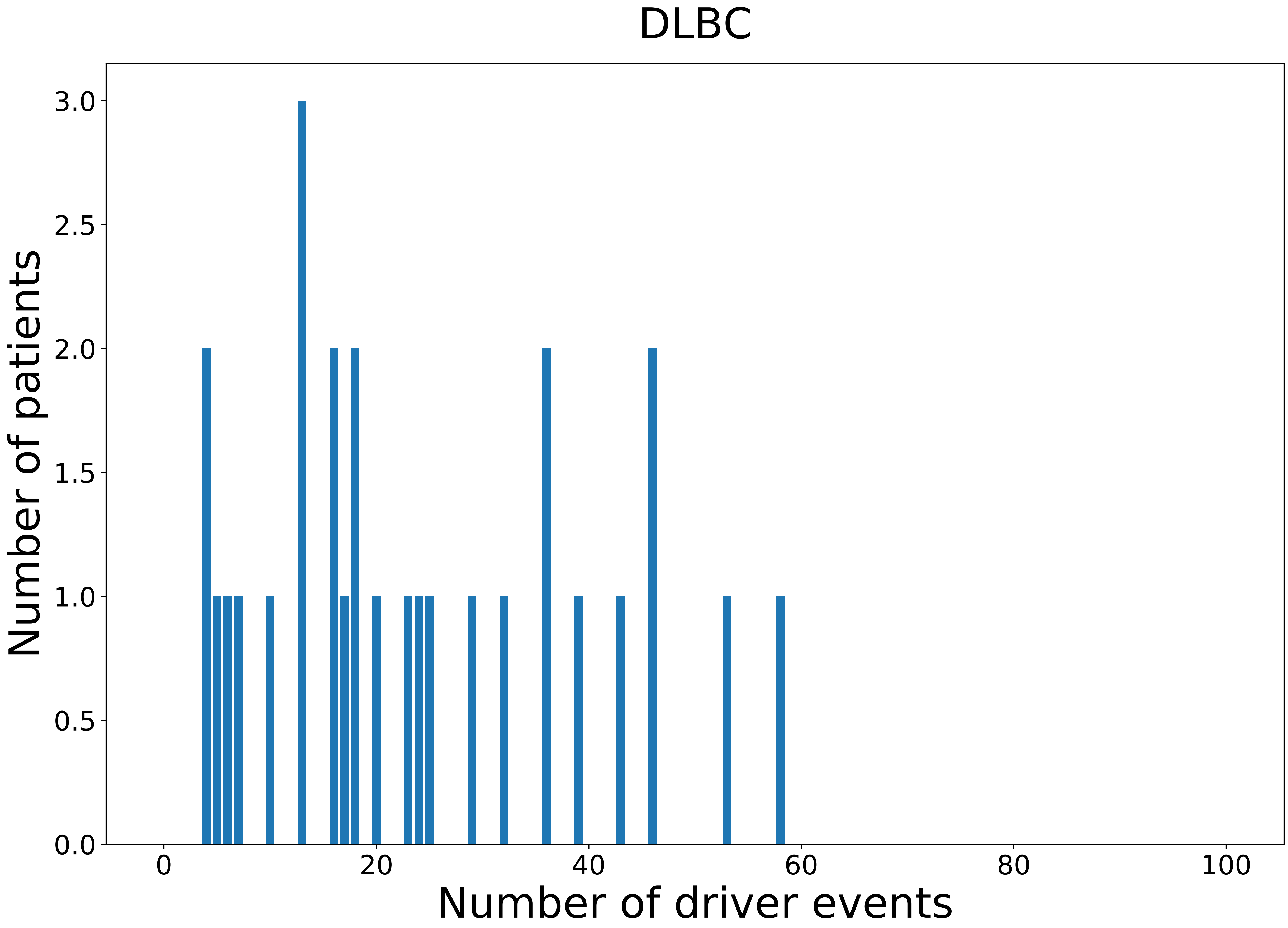

### 2021_8_22_13_42_DLBC_FEMALE.pdf

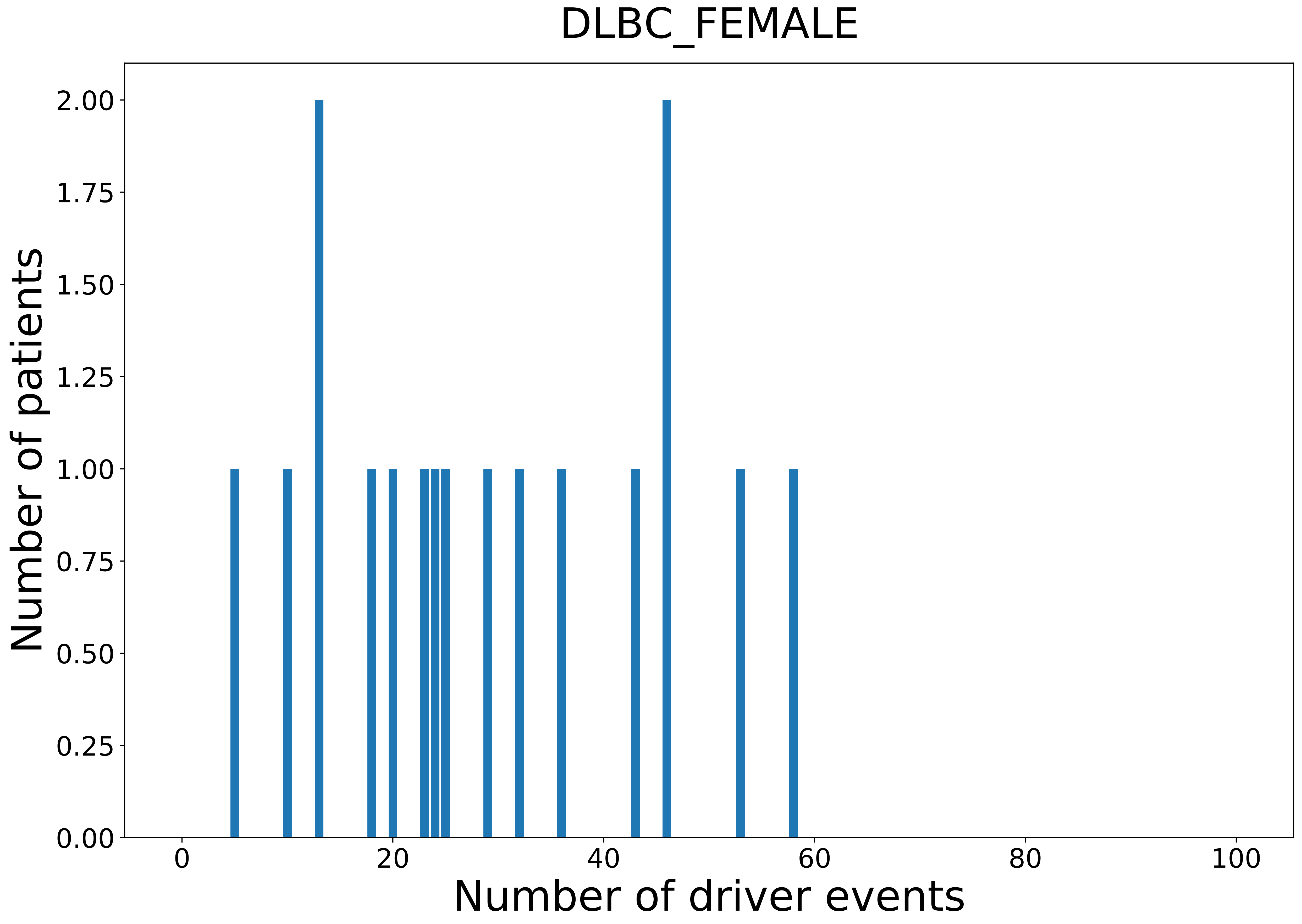

### 2021_8_22_13_42_DLBC_MALE.pdf

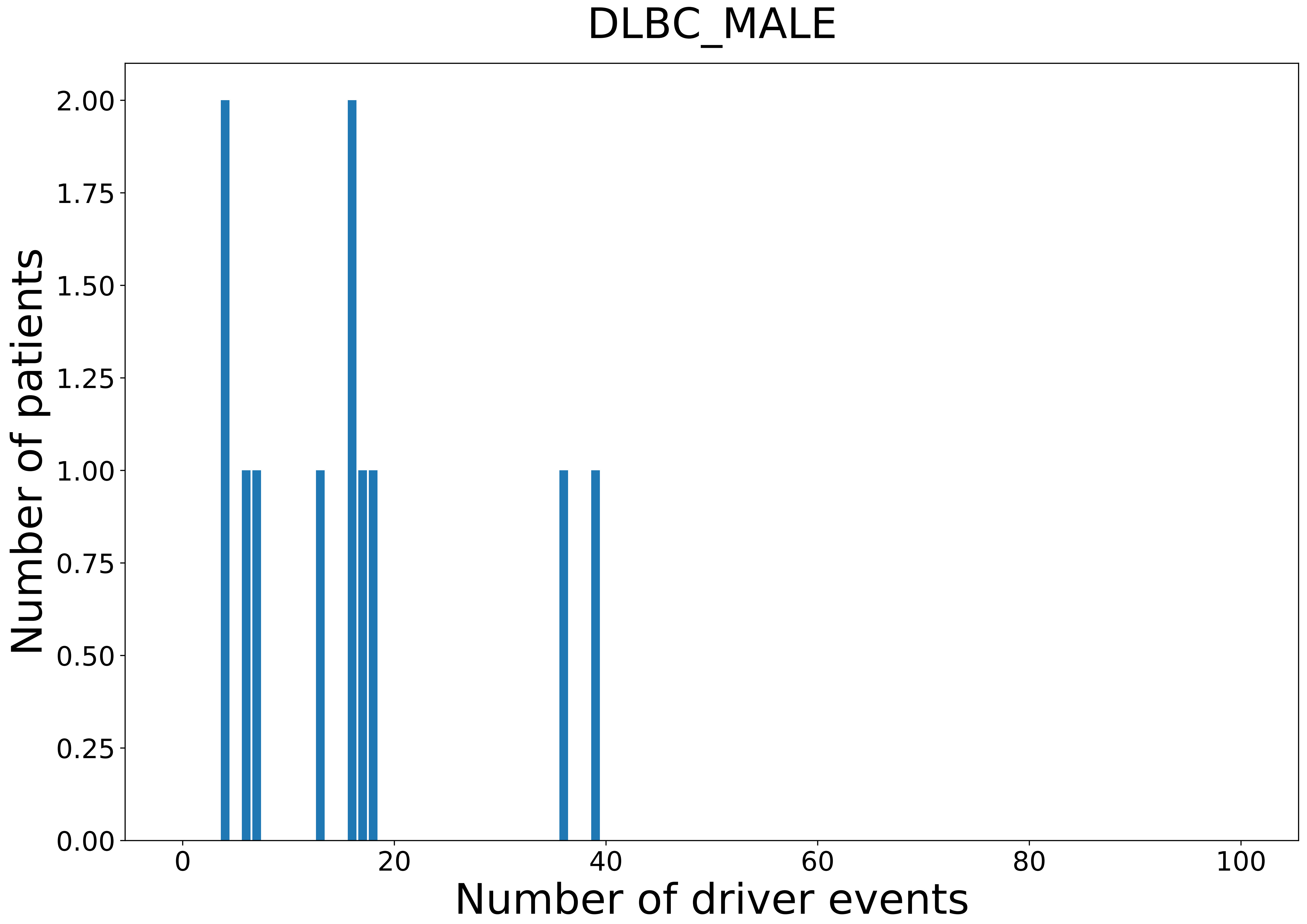

### 2021_8_22_13_42_ESCA.pdf

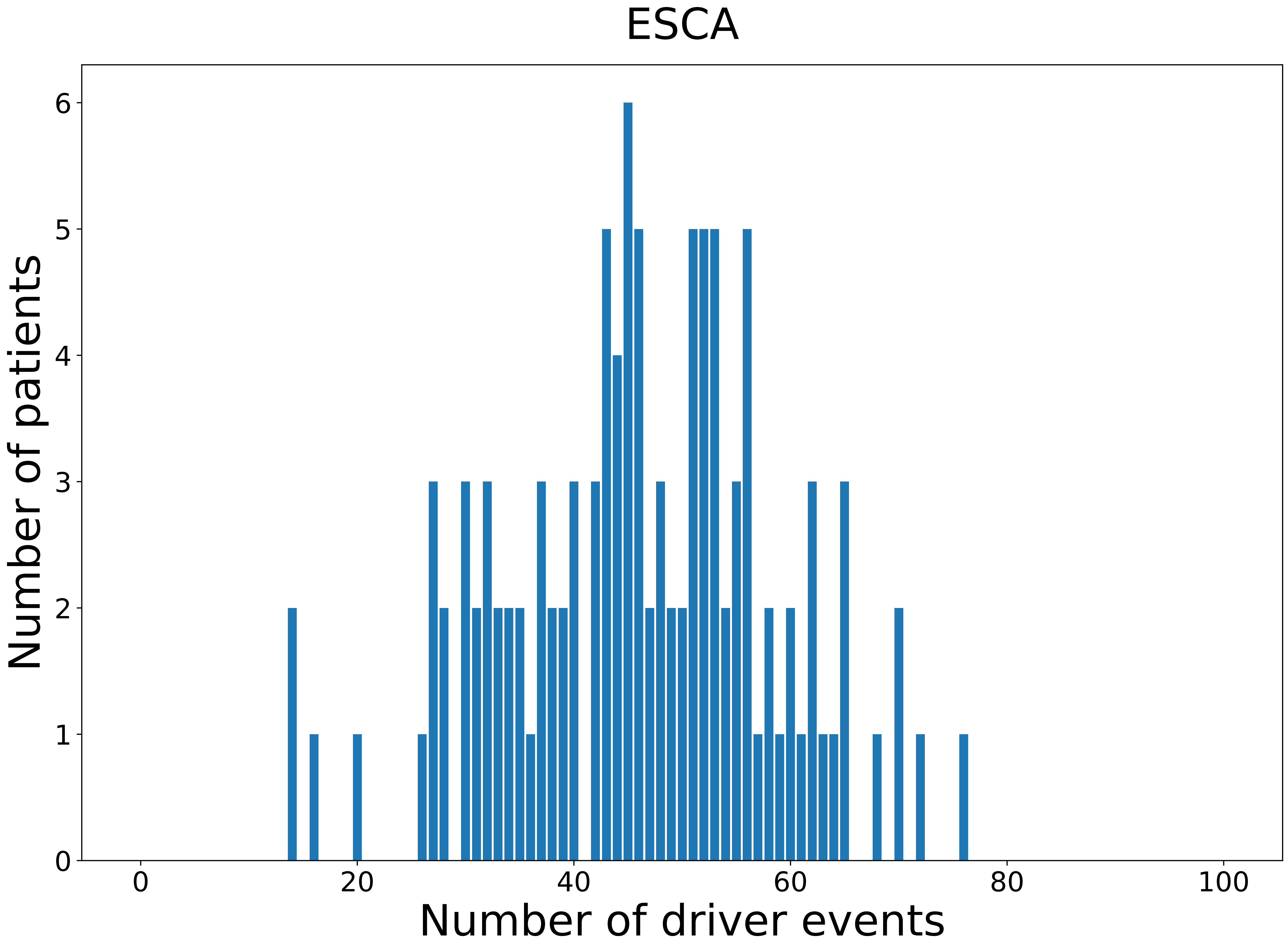

### 2021_8_22_13_42_ESCA_FEMALE.pdf

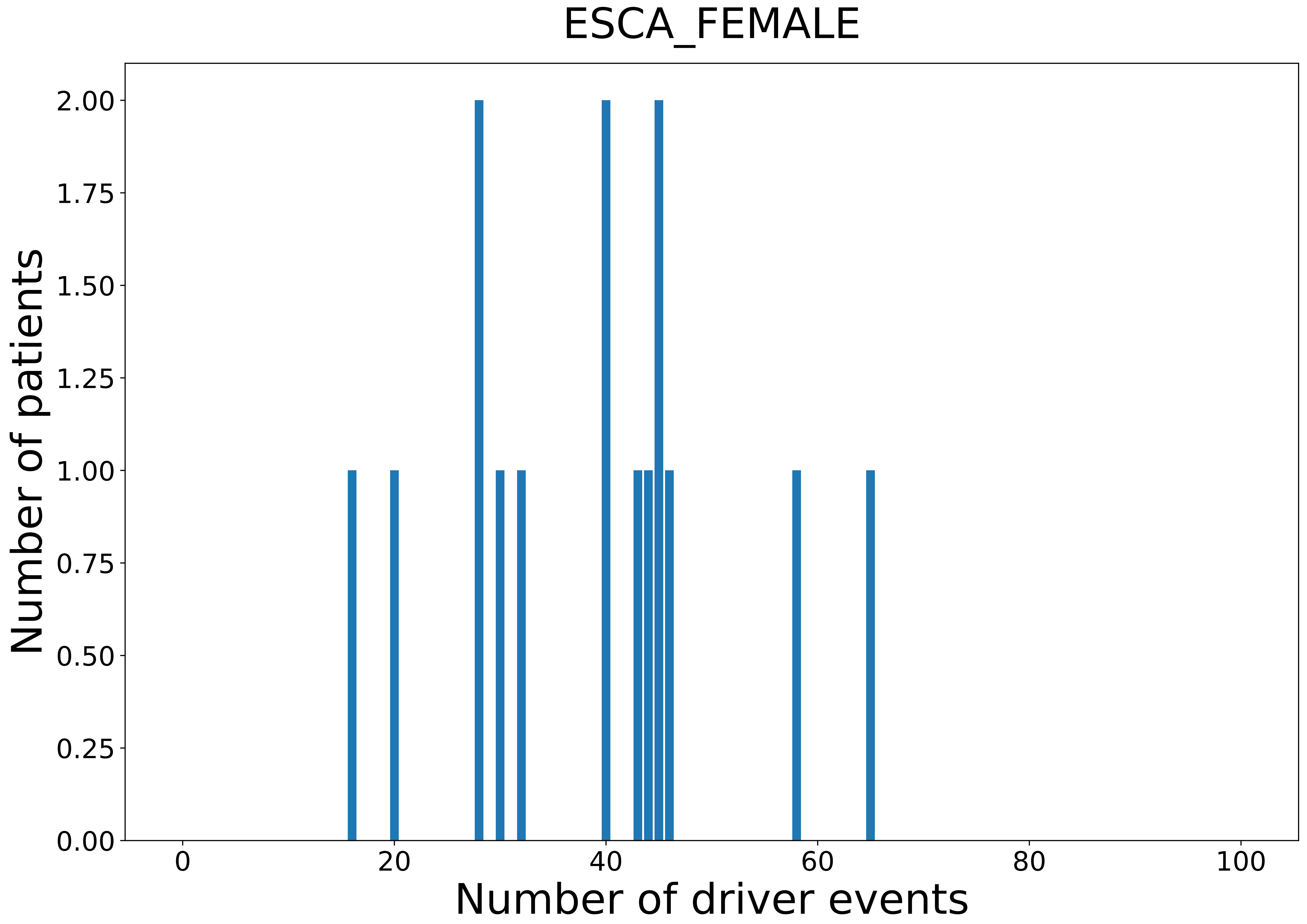

### 2021_8_22_13_42_ESCA_MALE.pdf

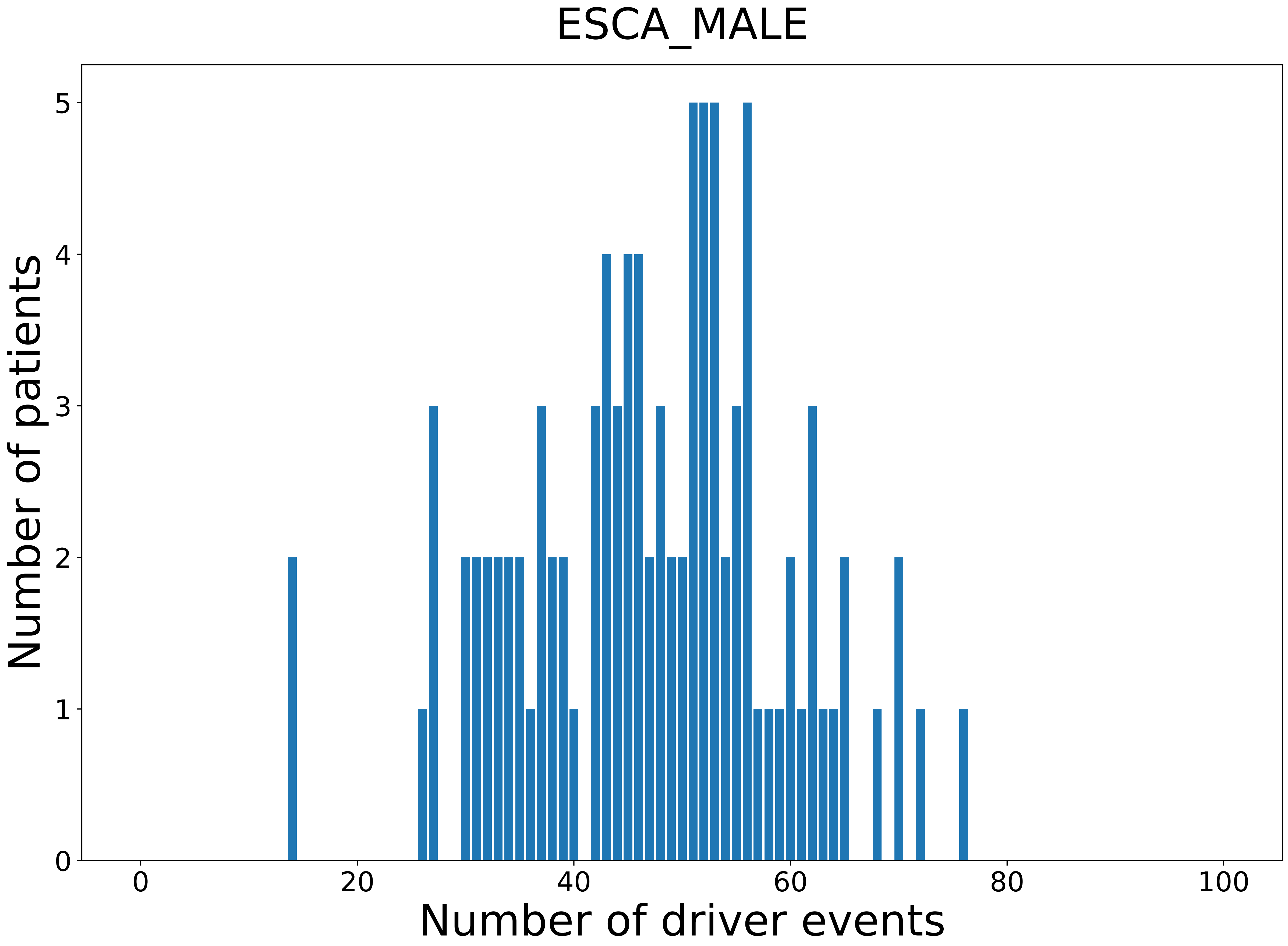

### 2021_8_22_13_42_GBM.pdf

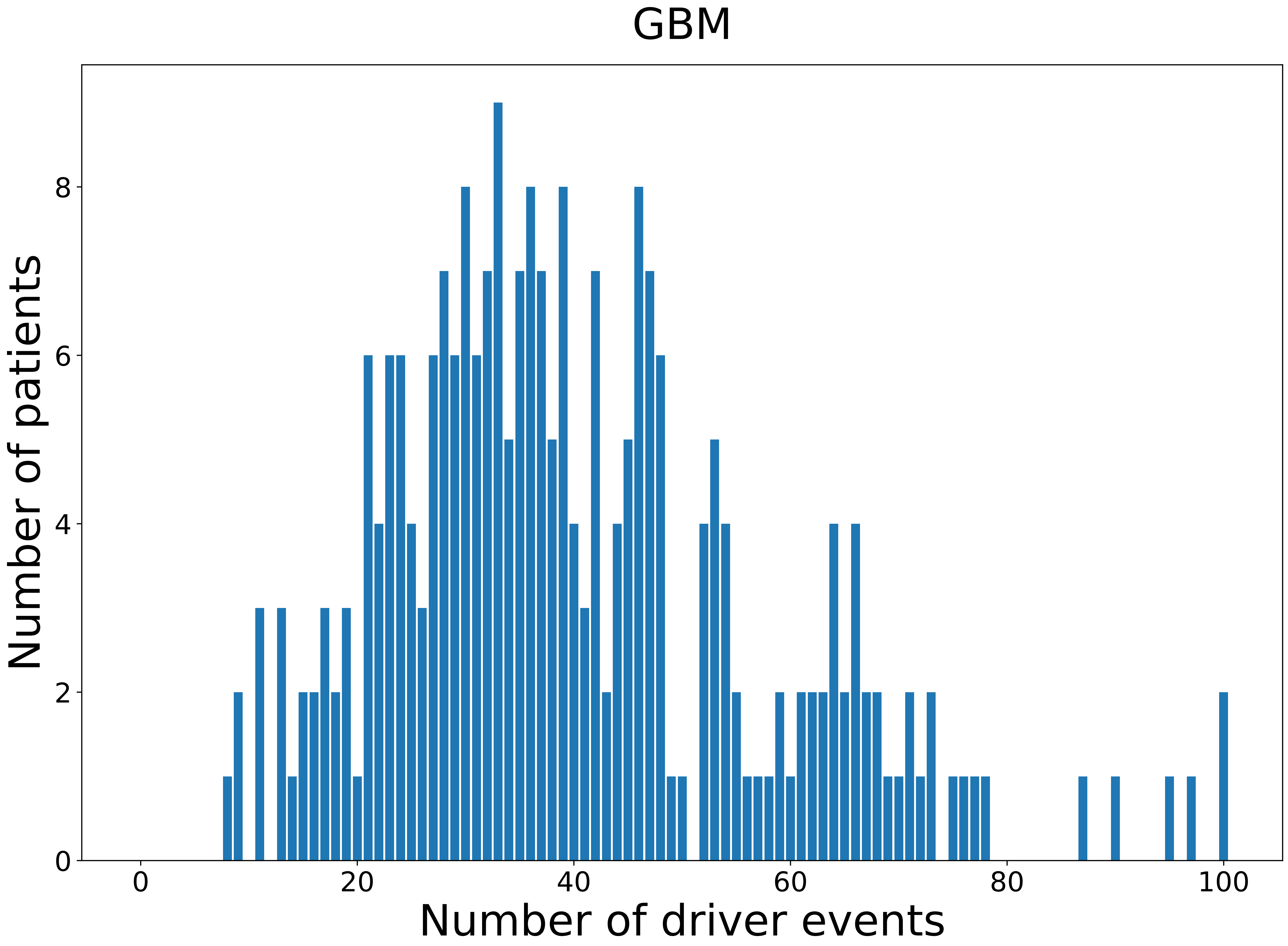

### 2021_8_22_13_42_GBM_FEMALE.pdf

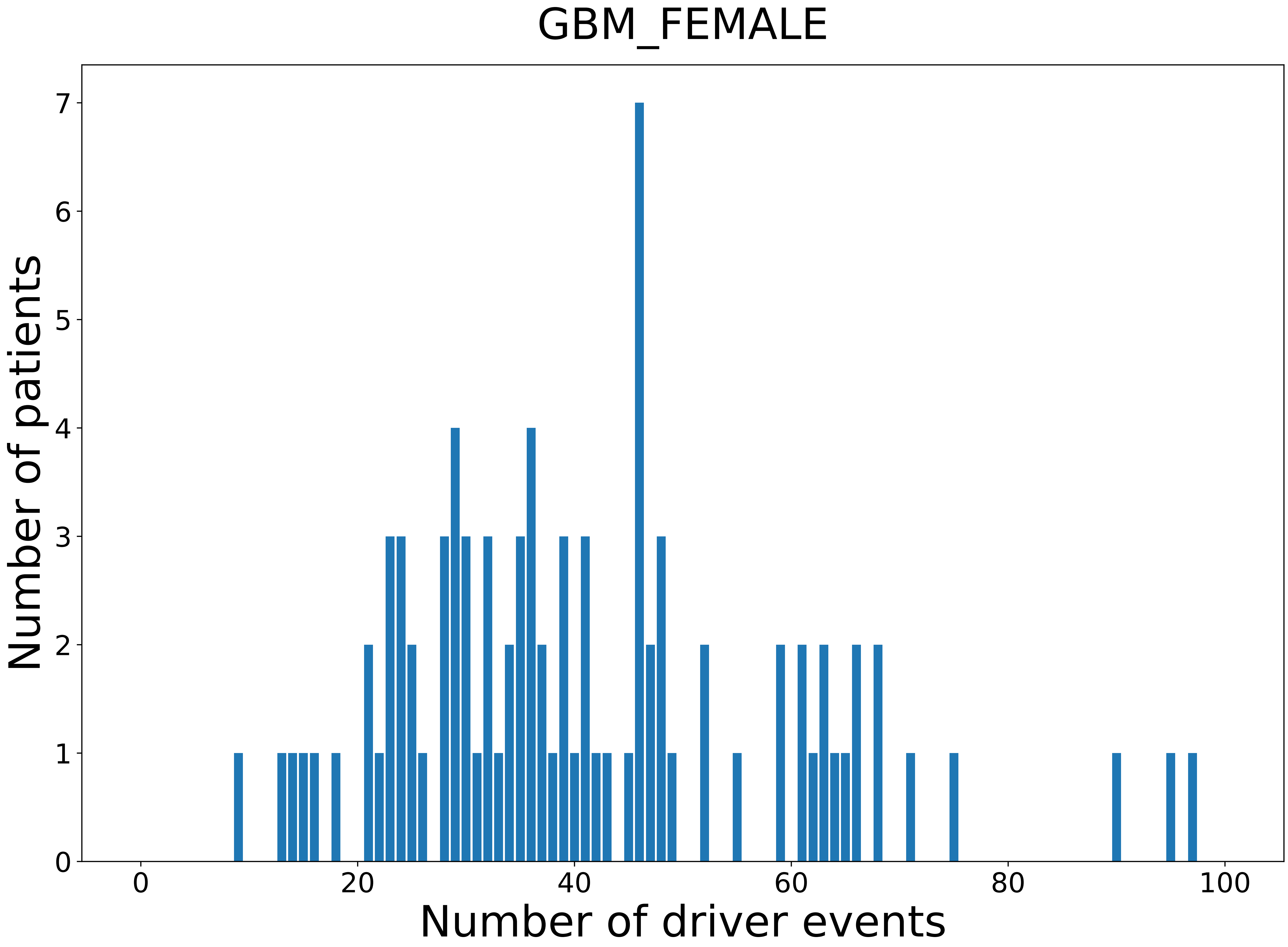

### 2021_8_22_13_42_GBM_MALE.pdf

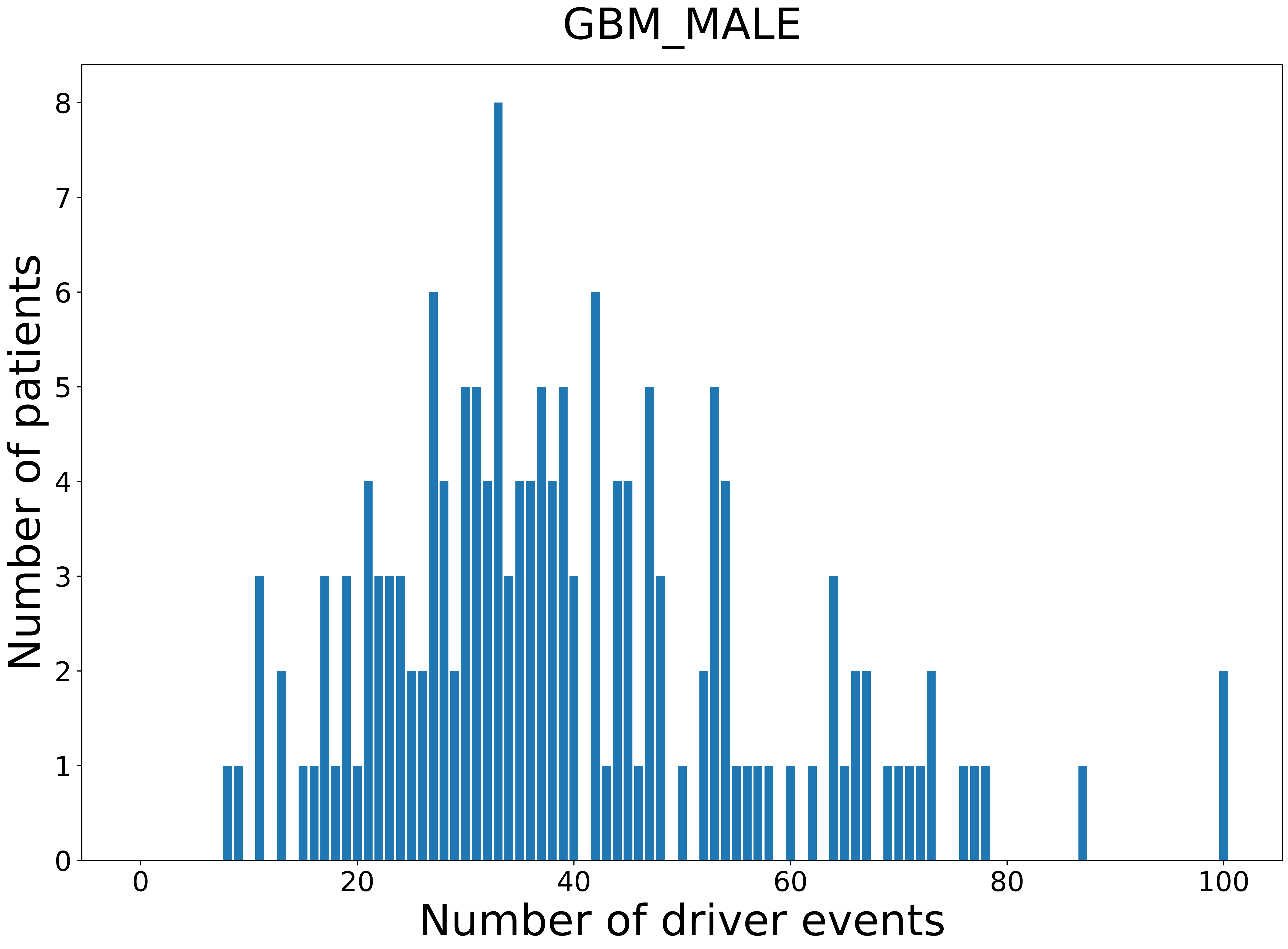

### 2021_8_22_13_42_HNSC.pdf

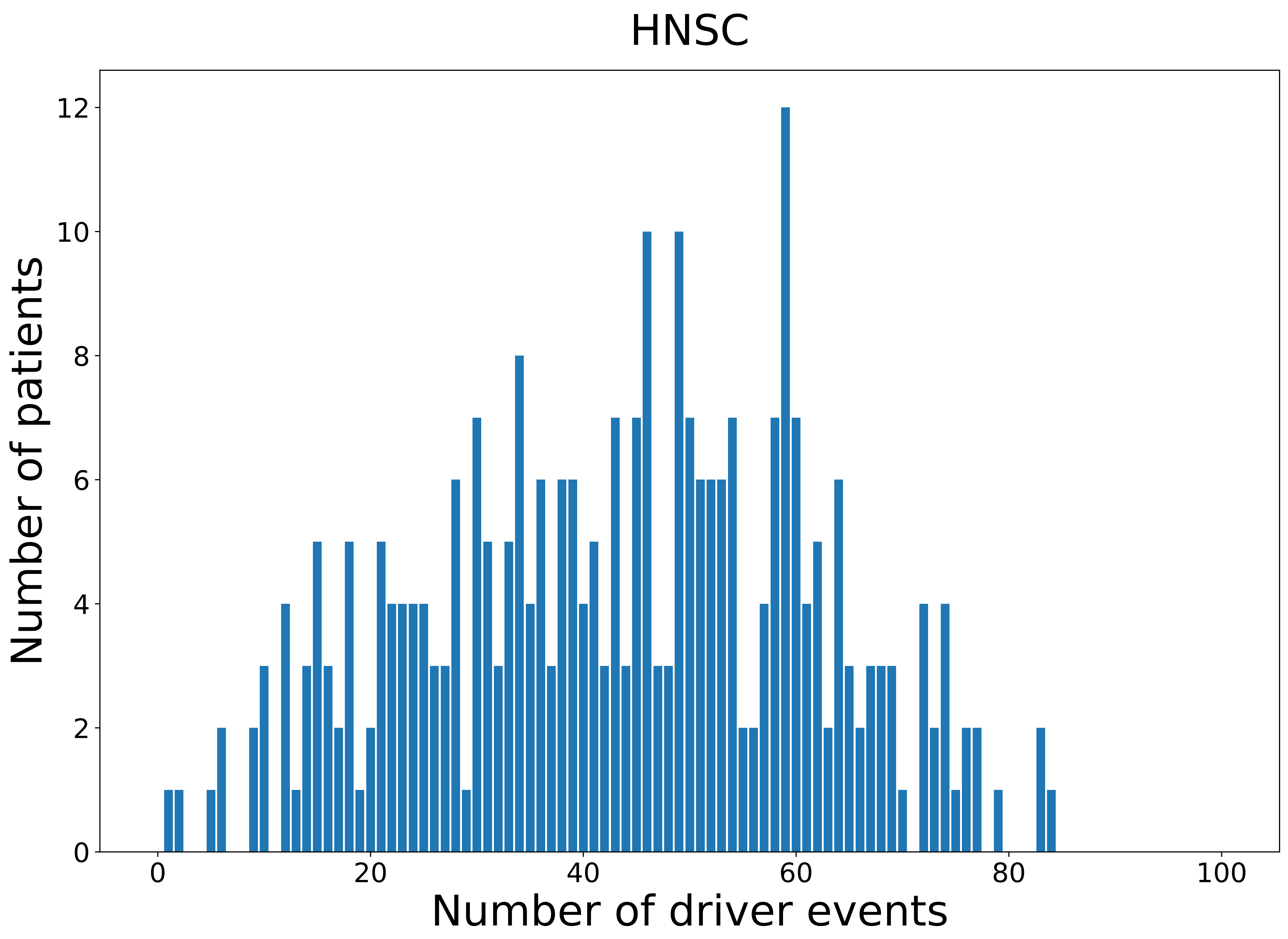

### 2021_8_22_13_42_HNSC_FEMALE.pdf

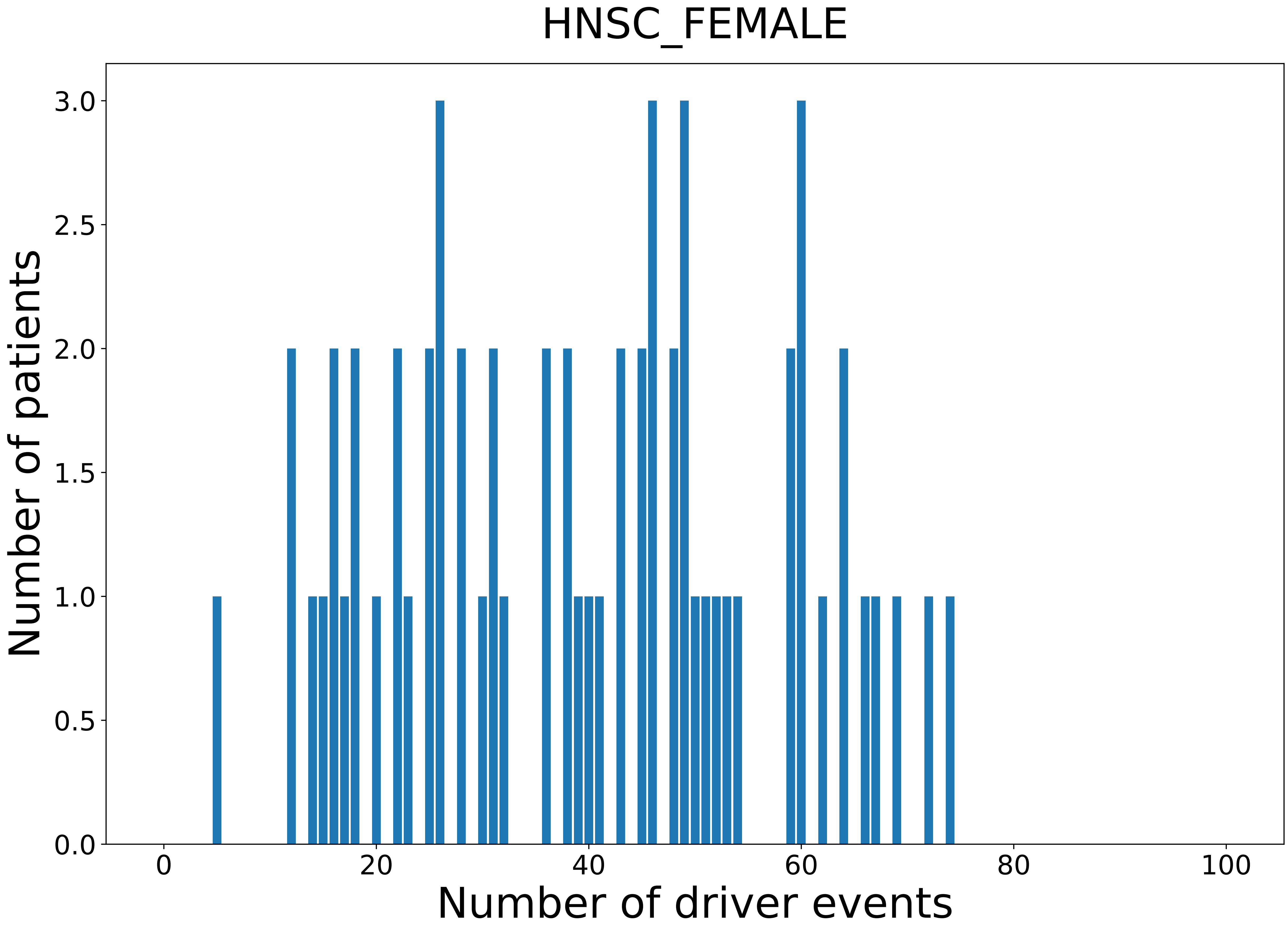

### 2021_8_22_13_42_HNSC_MALE.pdf

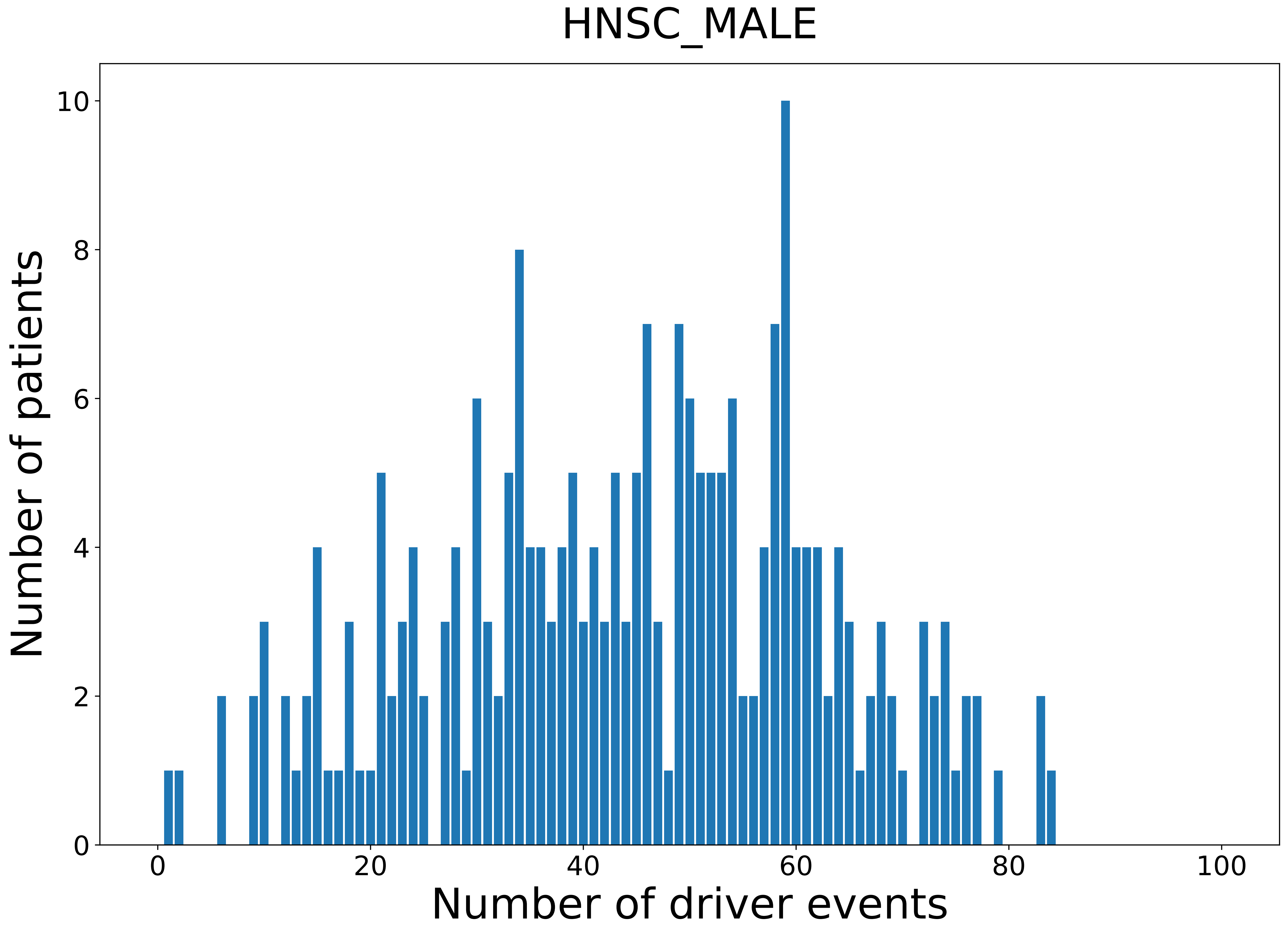

### 2021_8_22_13_42_KICH.pdf

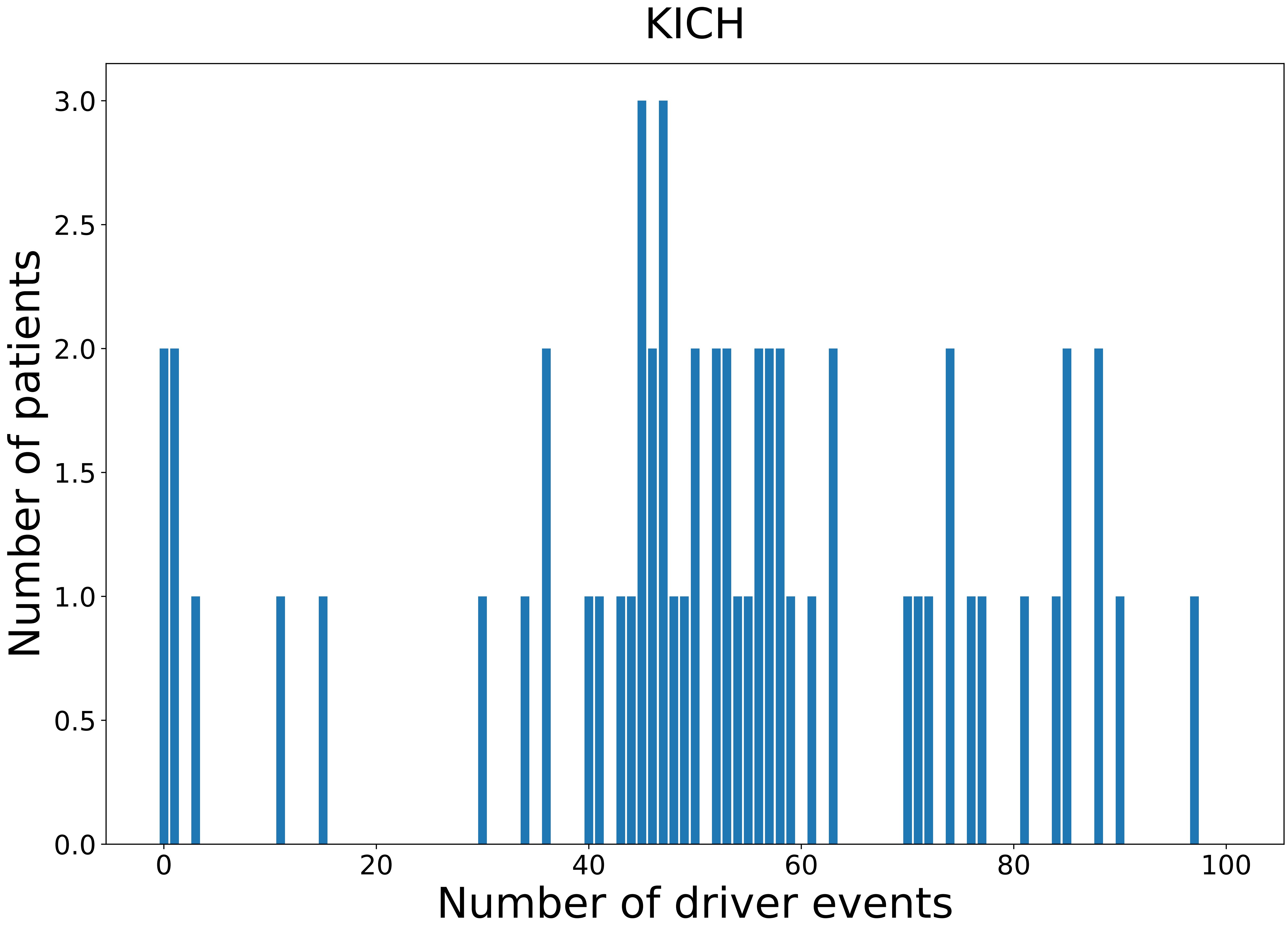

### 2021_8_22_13_42_KICH_FEMALE.pdf

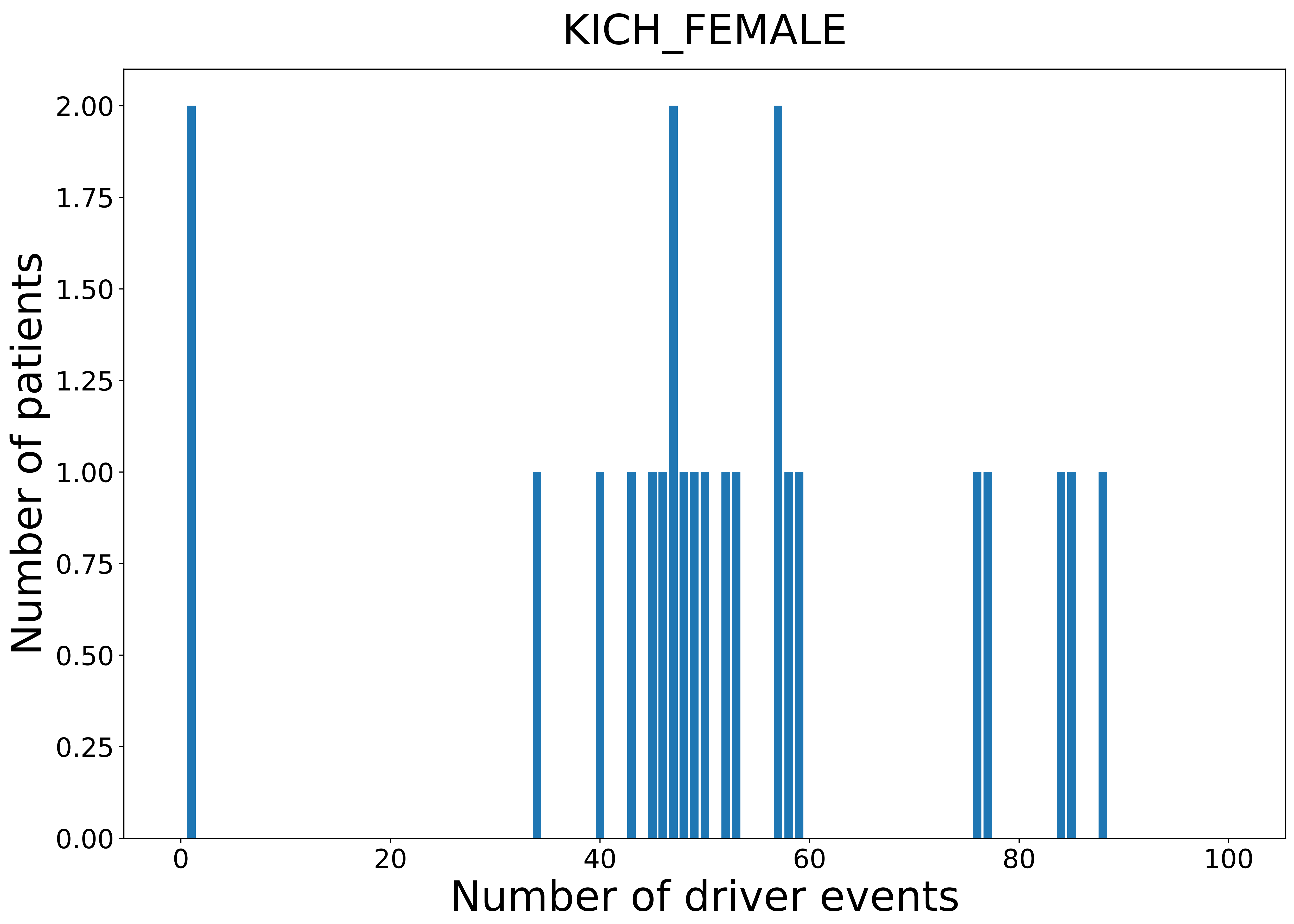

### 2021_8_22_13_42_KIRC.pdf

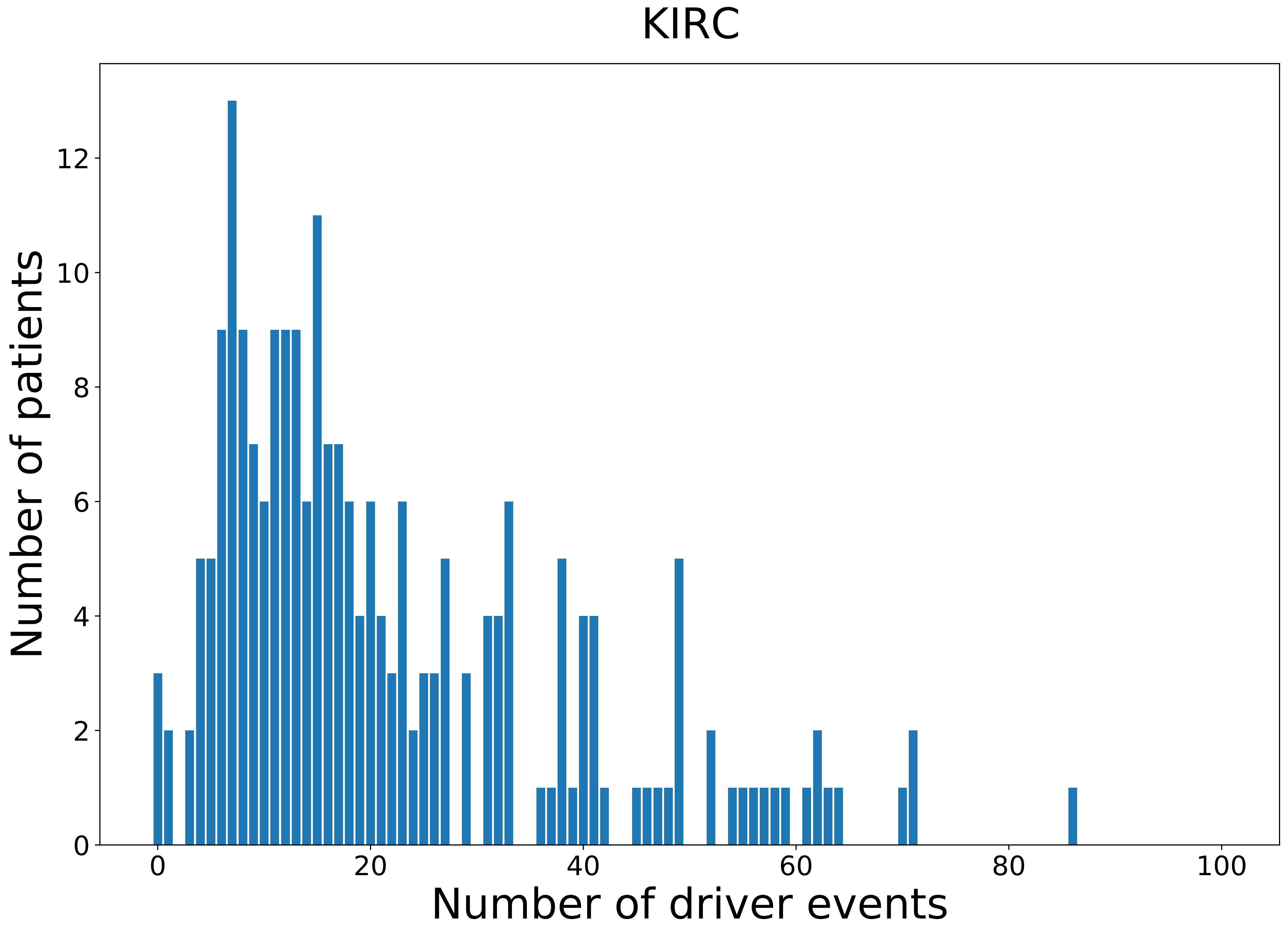

### 2021_8_22_13_42_KIRC_FEMALE.pdf

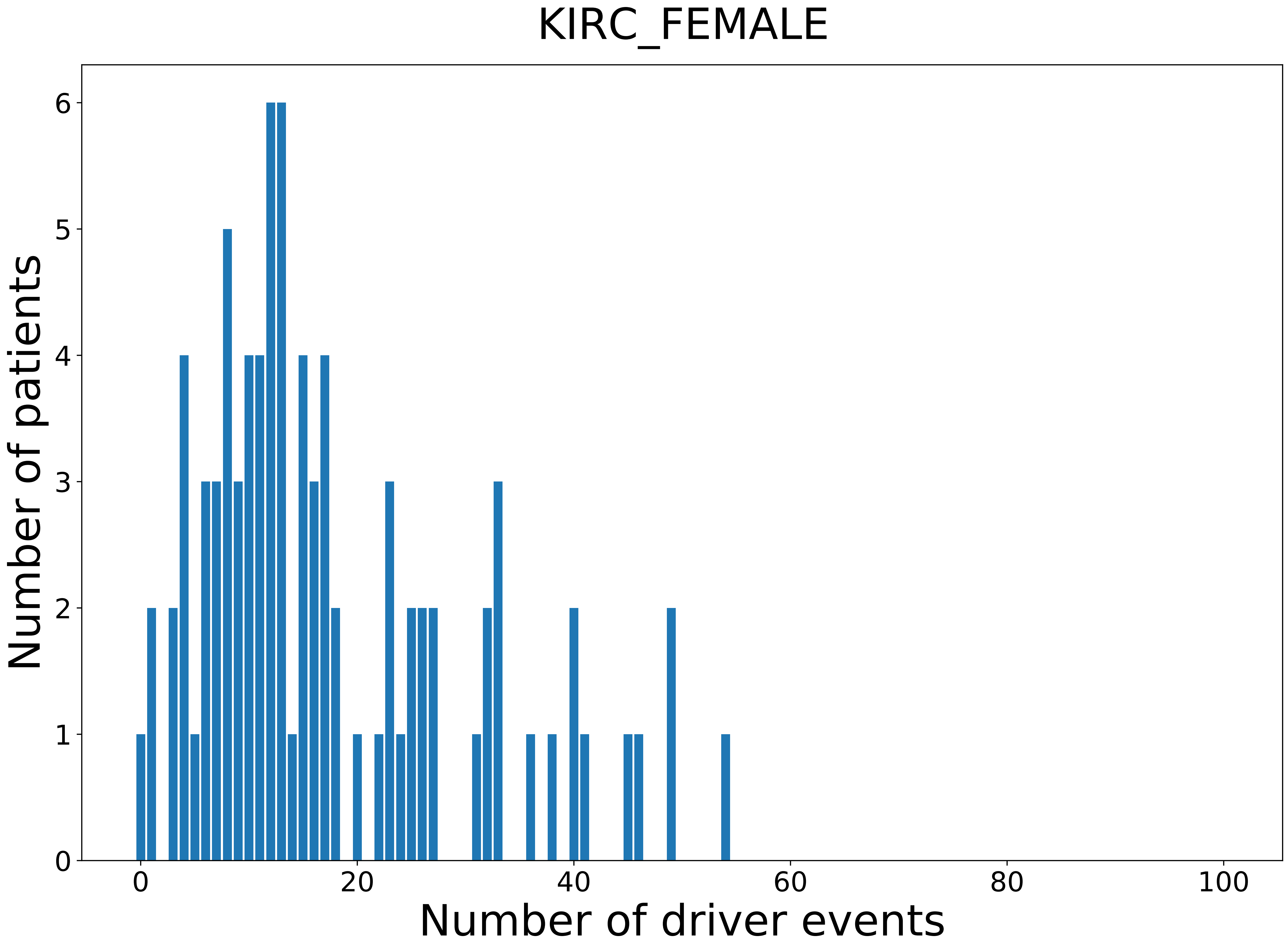

### 2021_8_22_13_42_KIRC_MALE.pdf

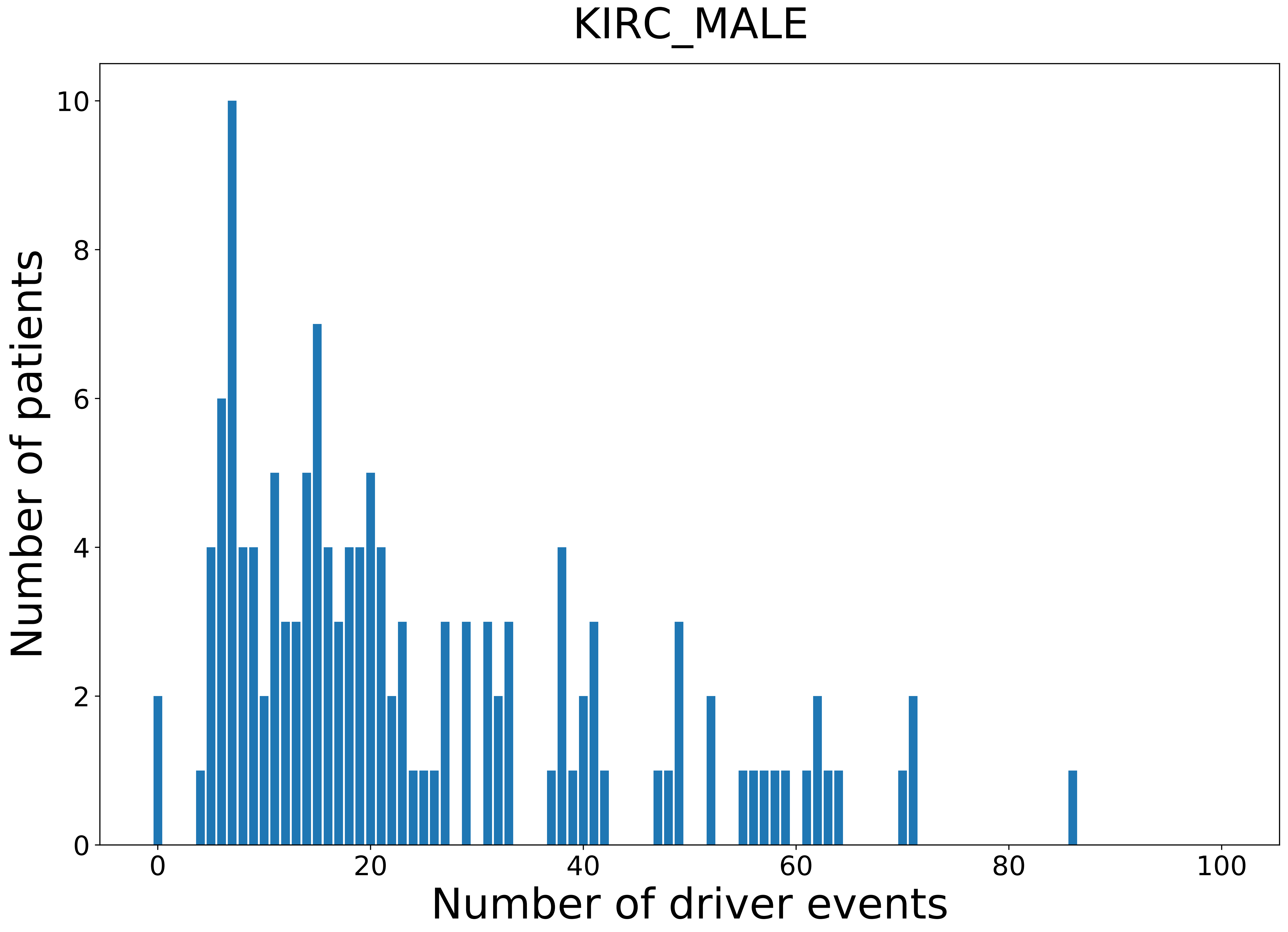

### 2021_8_22_13_42_KIRP.pdf

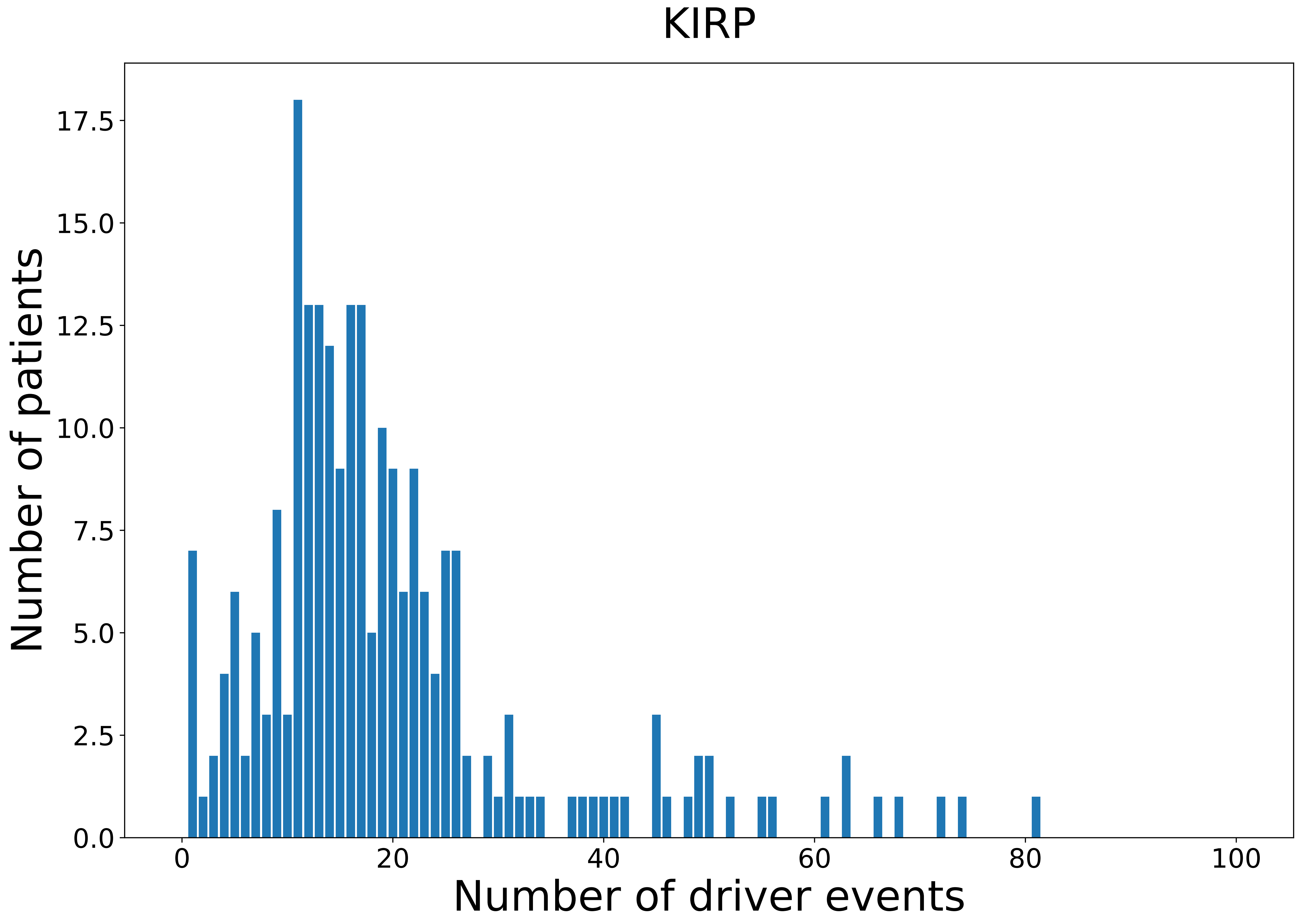

### 2021_8_22_13_42_KIRP_FEMALE.pdf

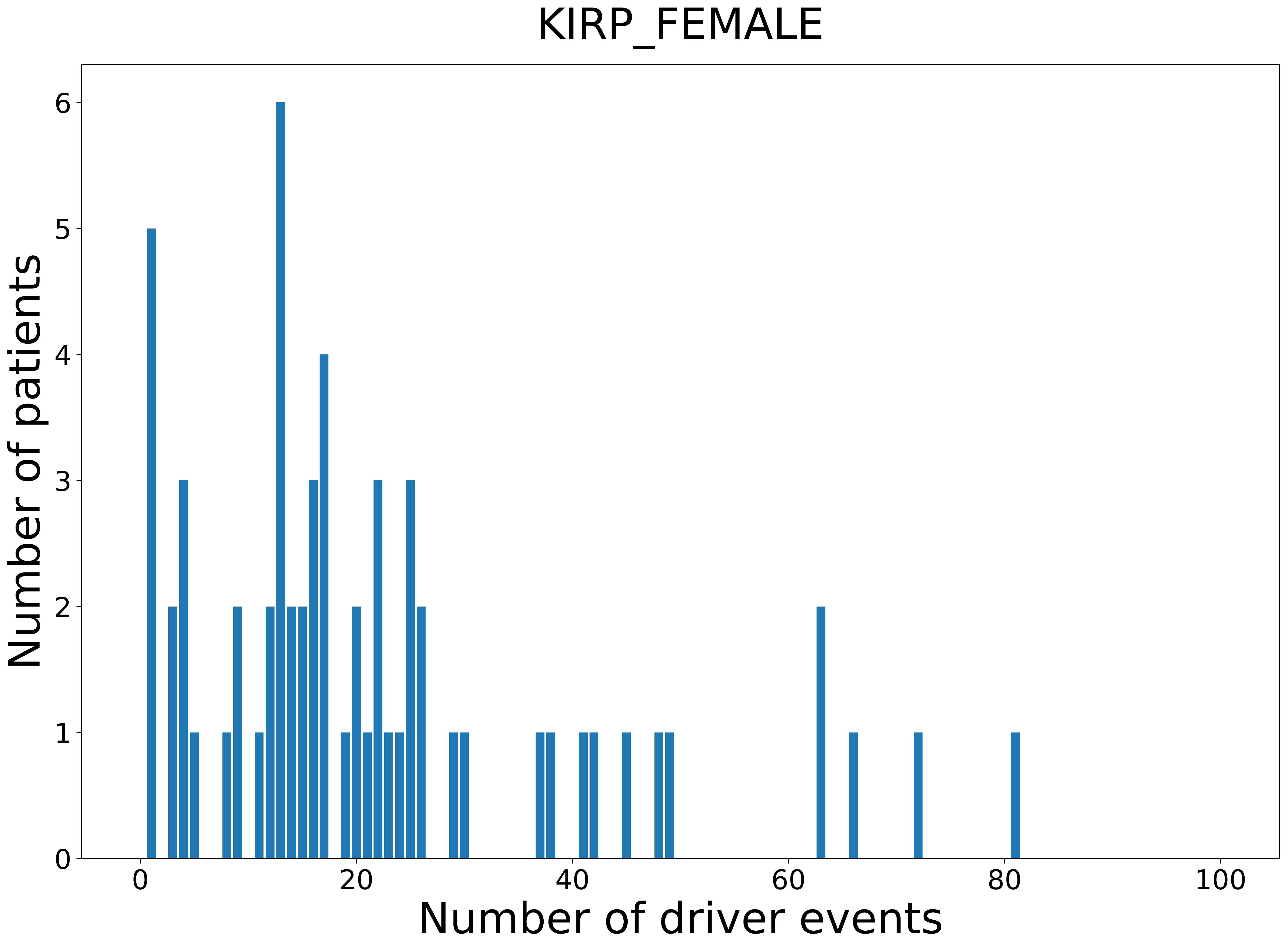

### 2021_8_22_13_42_KIRP_MALE.pdf

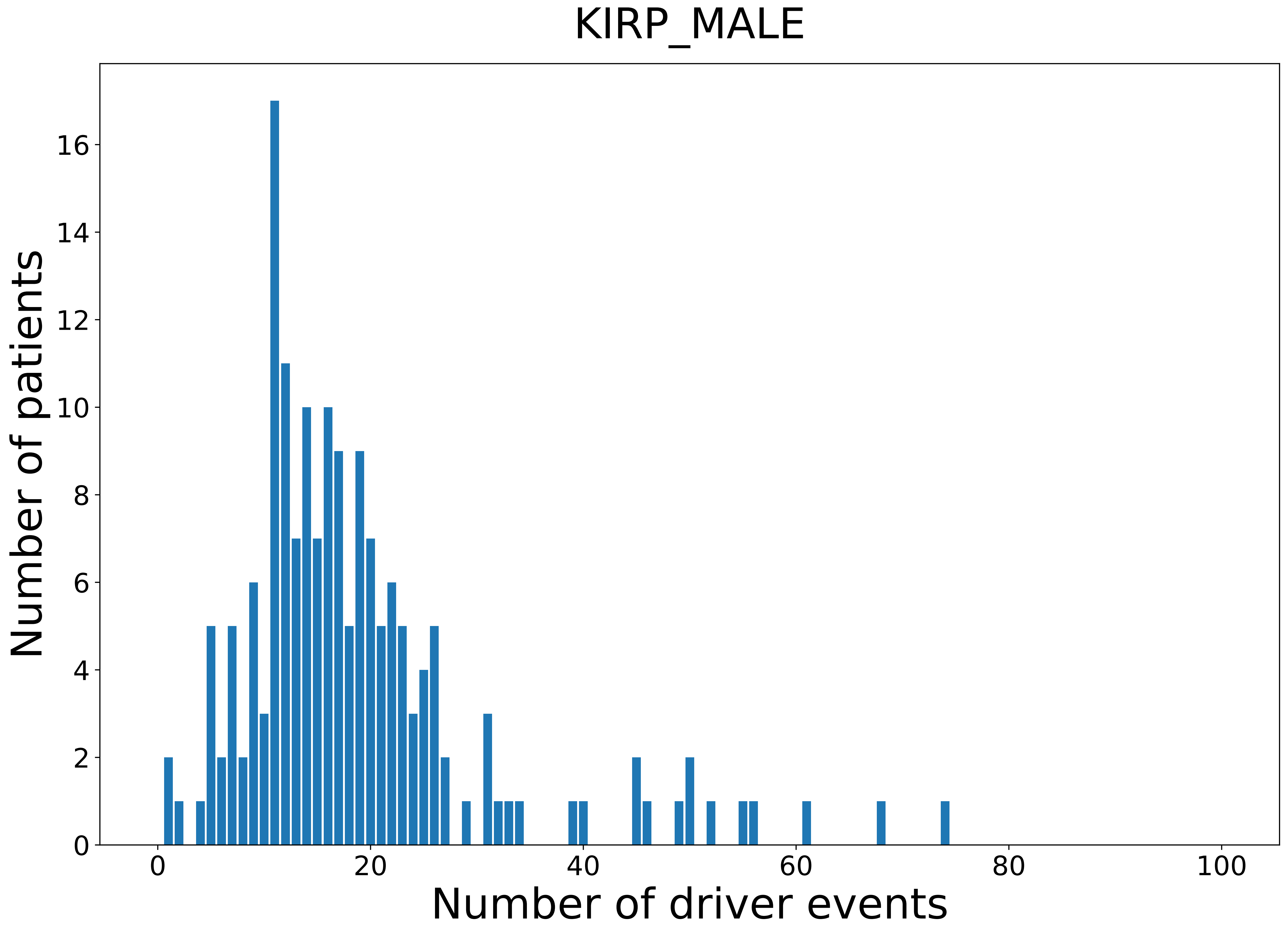

### 2021_8_22_13_42_LGG.pdf

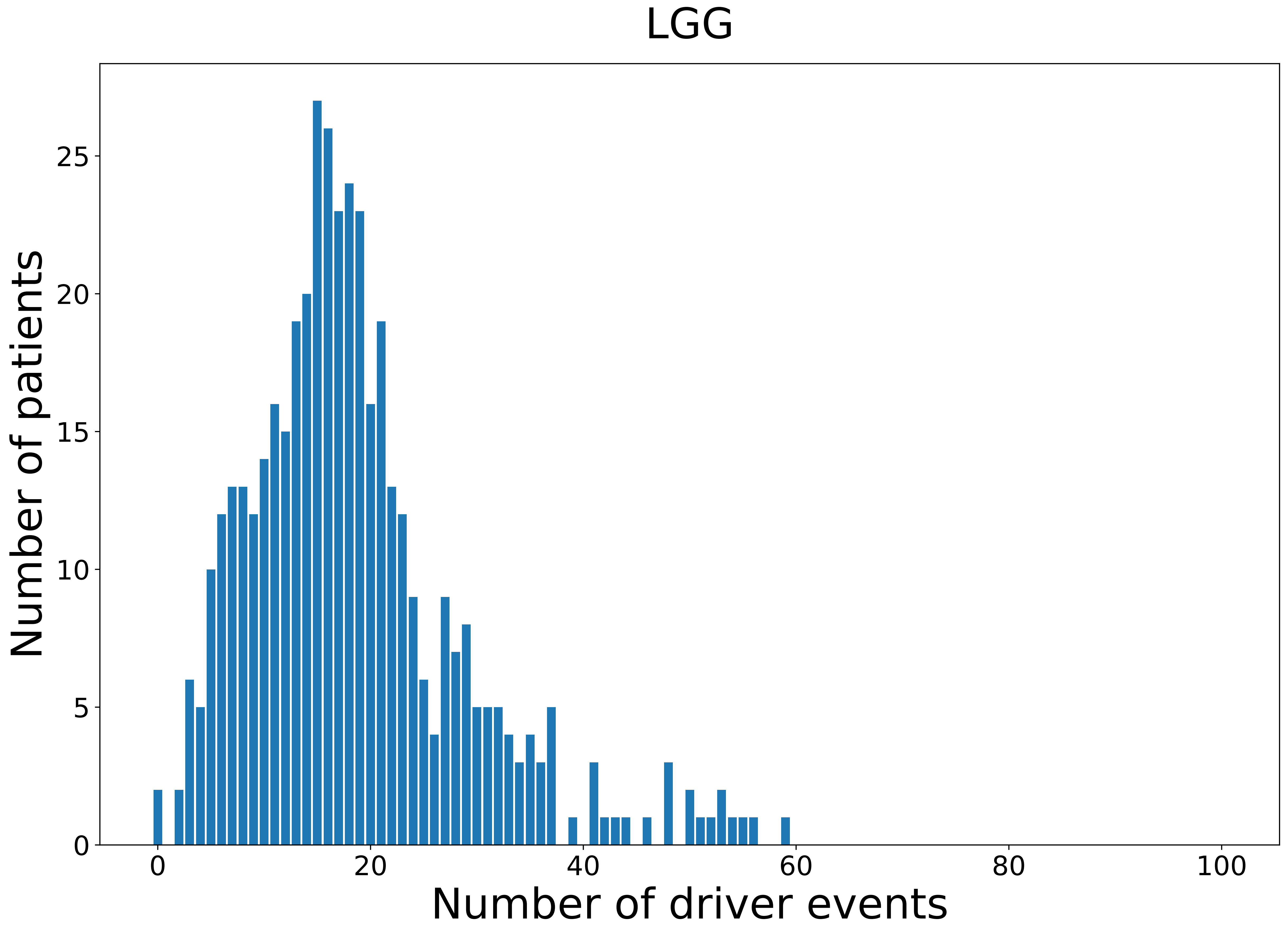

### 2021_8_22_13_42_LGG_FEMALE.pdf

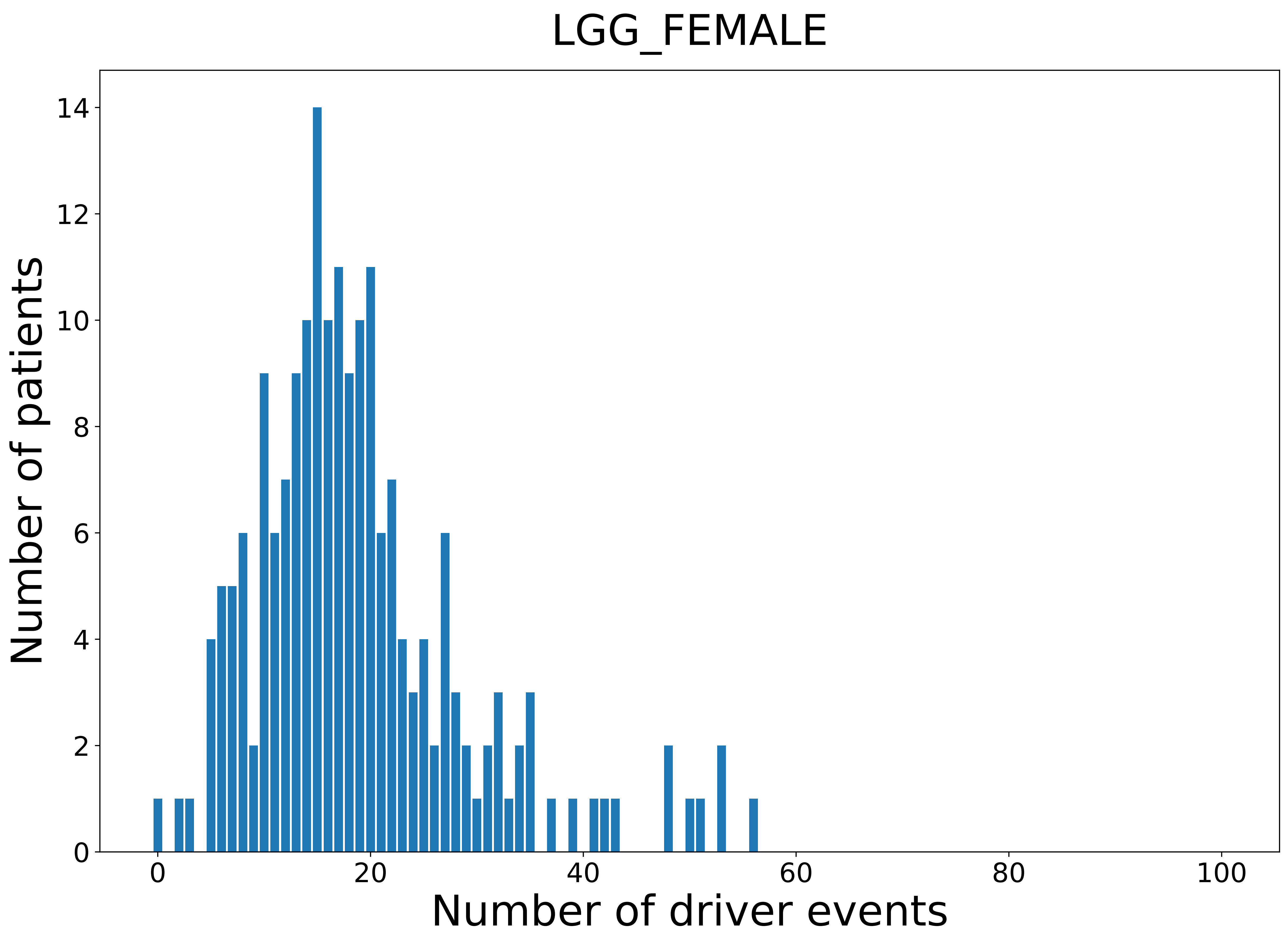

### 2021_8_22_13_42_LGG_MALE.pdf

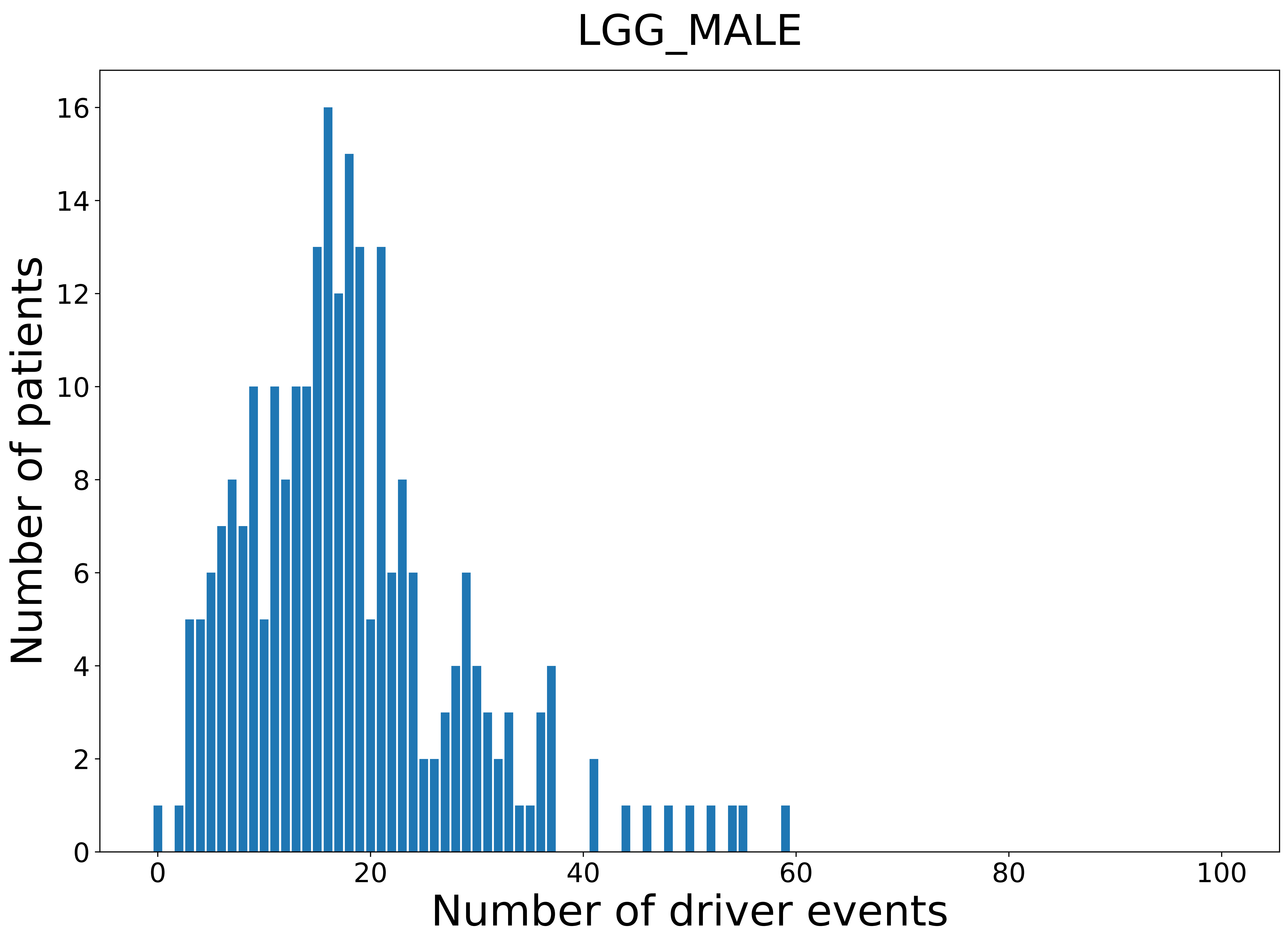

### 2021_8_22_13_42_LIHC.pdf

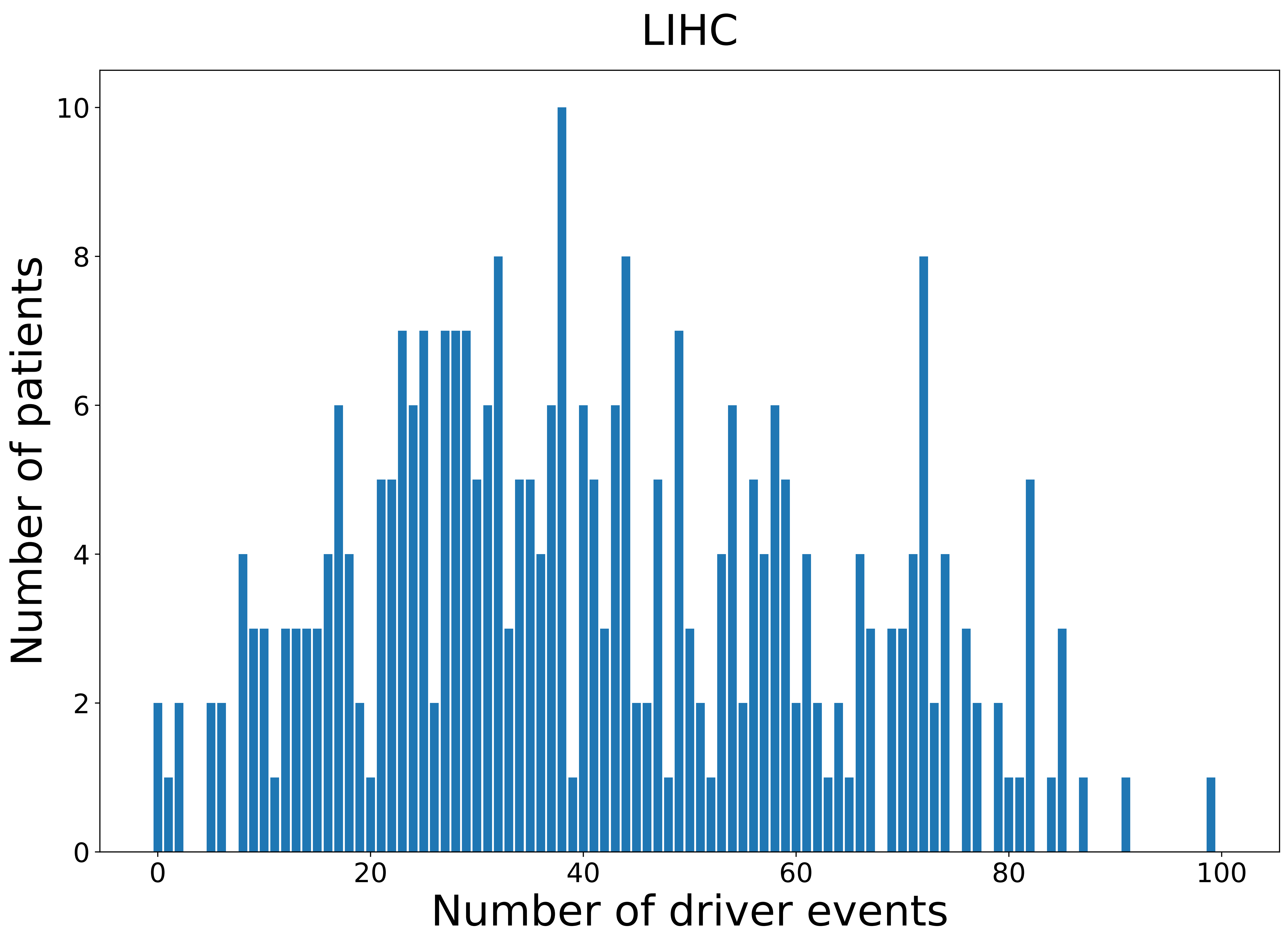

### 2021_8_22_13_42_LIHC_FEMALE.pdf

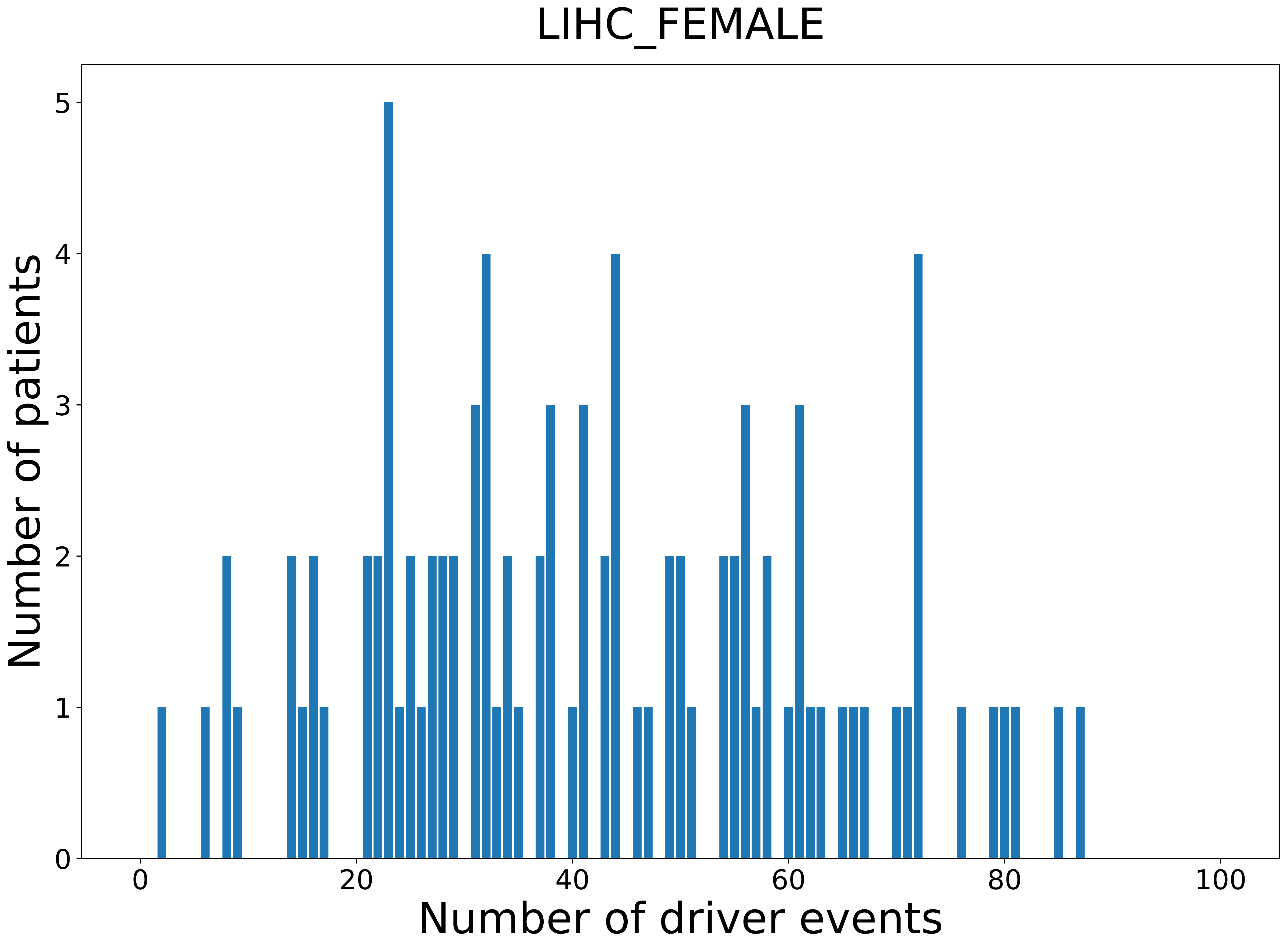

### 2021_8_22_13_42_LIHC_MALE.pdf

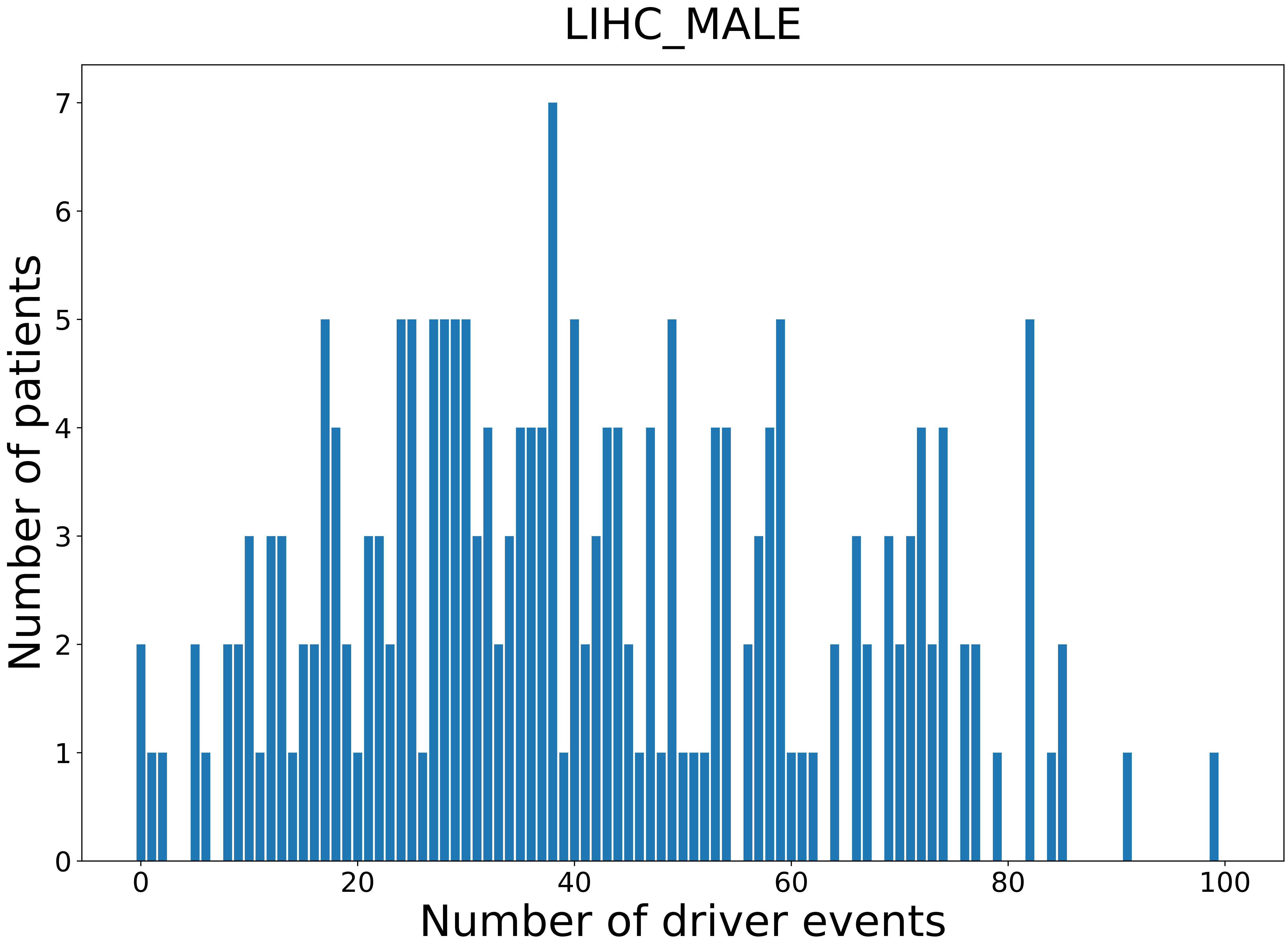

### 2021_8_22_13_42_LUAD.pdf

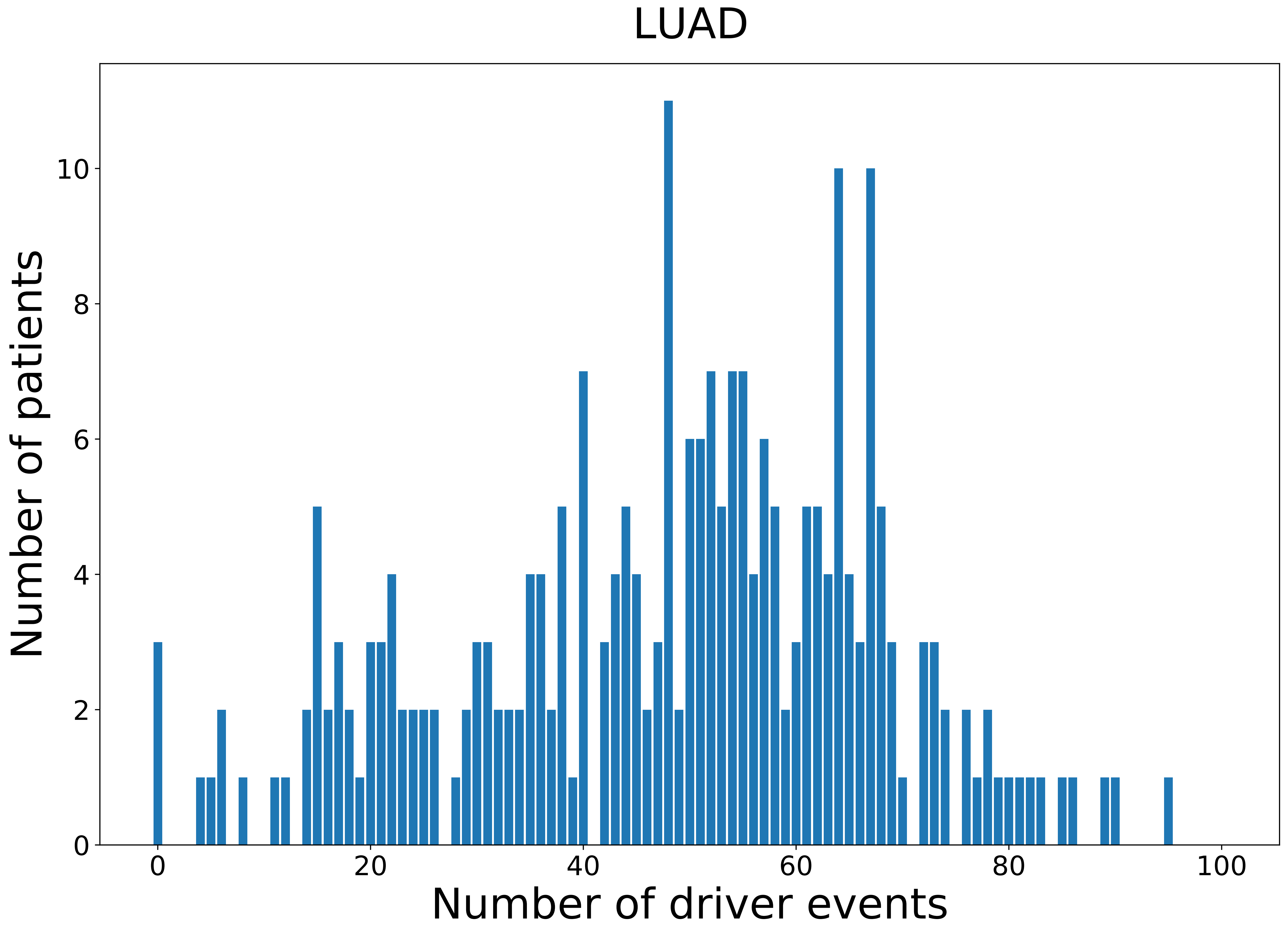

### 2021_8_22_13_42_LUAD_FEMALE.pdf

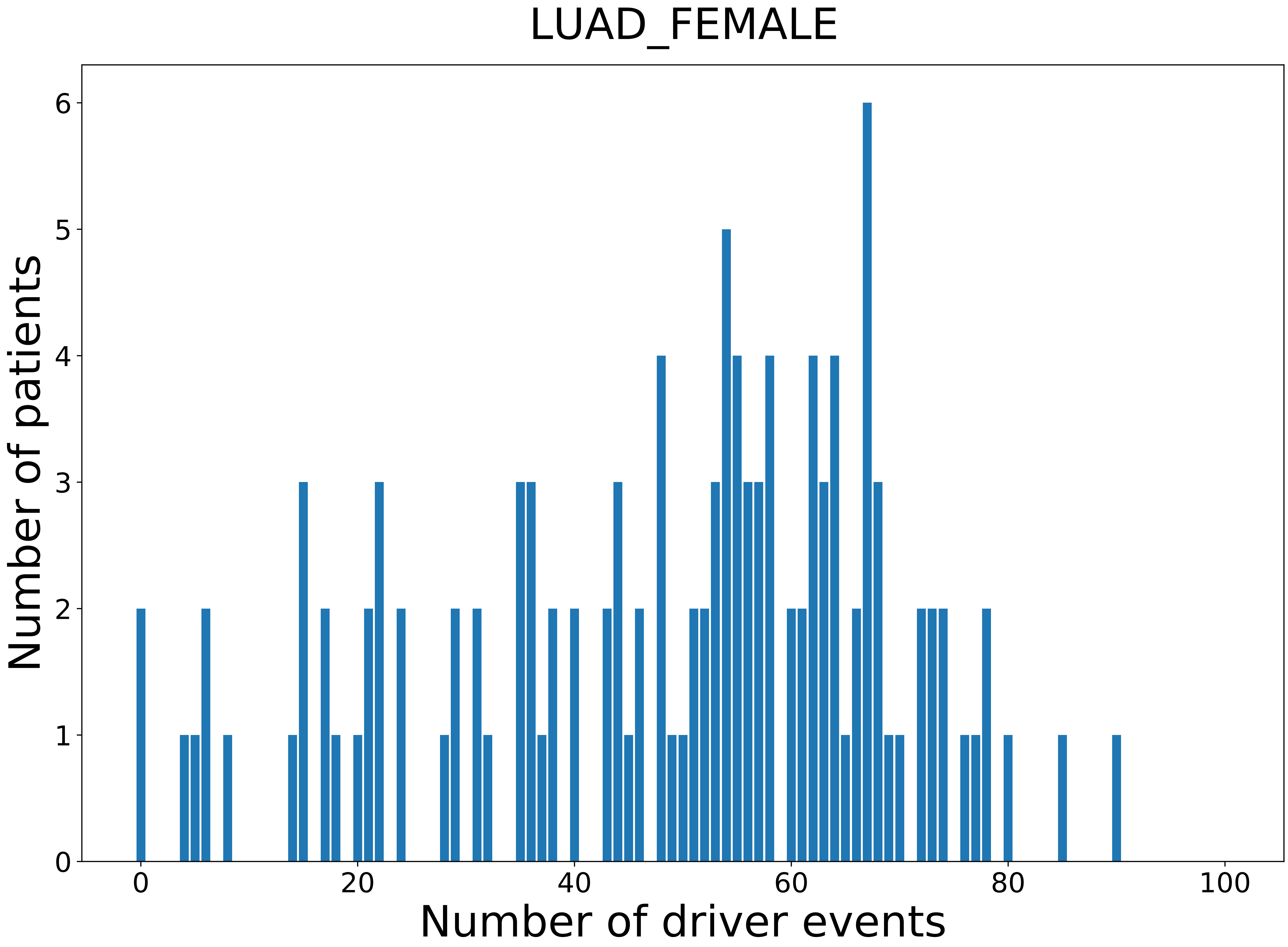

### 2021_8_22_13_42_LUAD_MALE.pdf

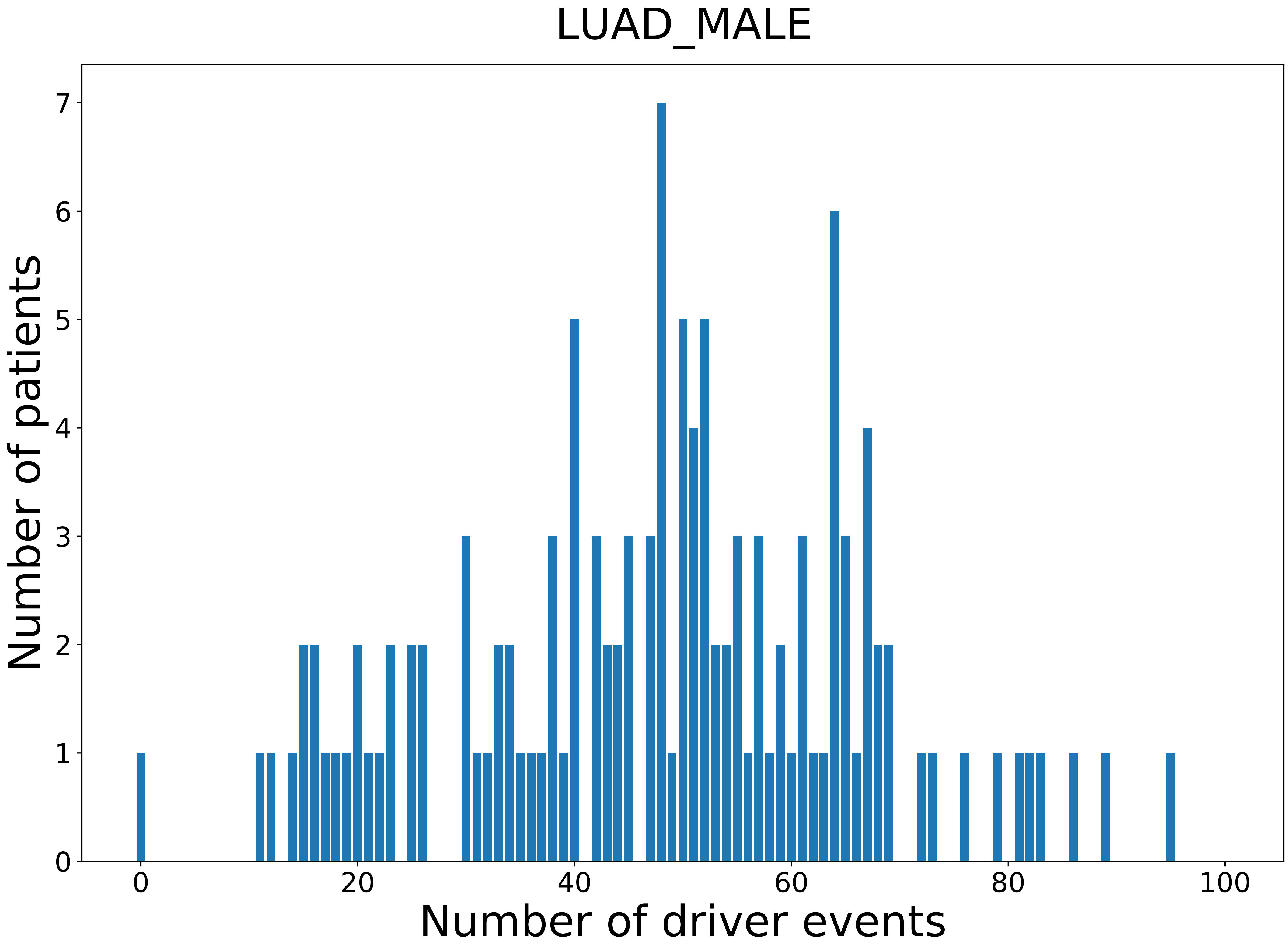

### 2021_8_22_13_42_LUSC.pdf

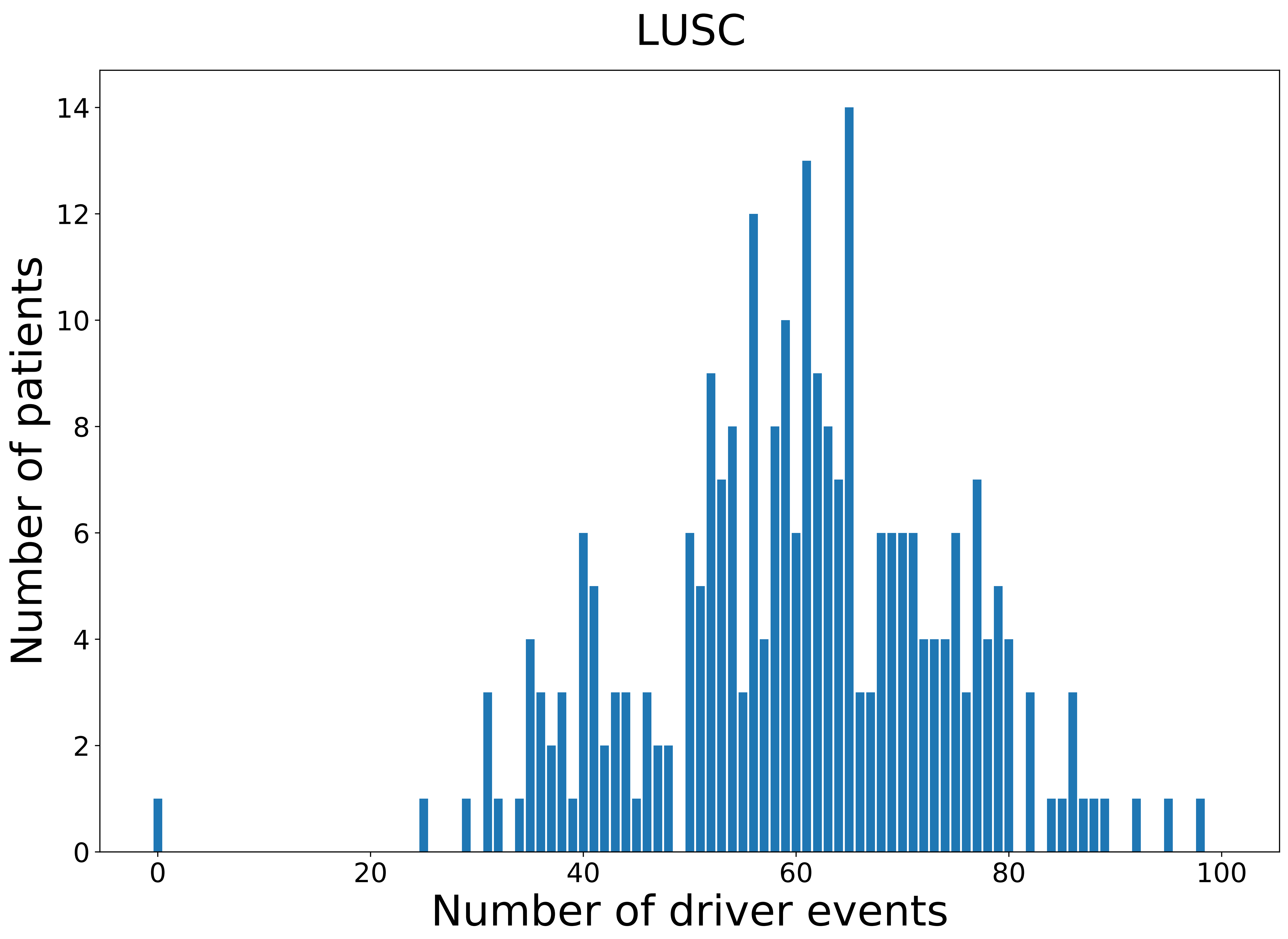

### 2021_8_22_13_42_LUSC_FEMALE.pdf

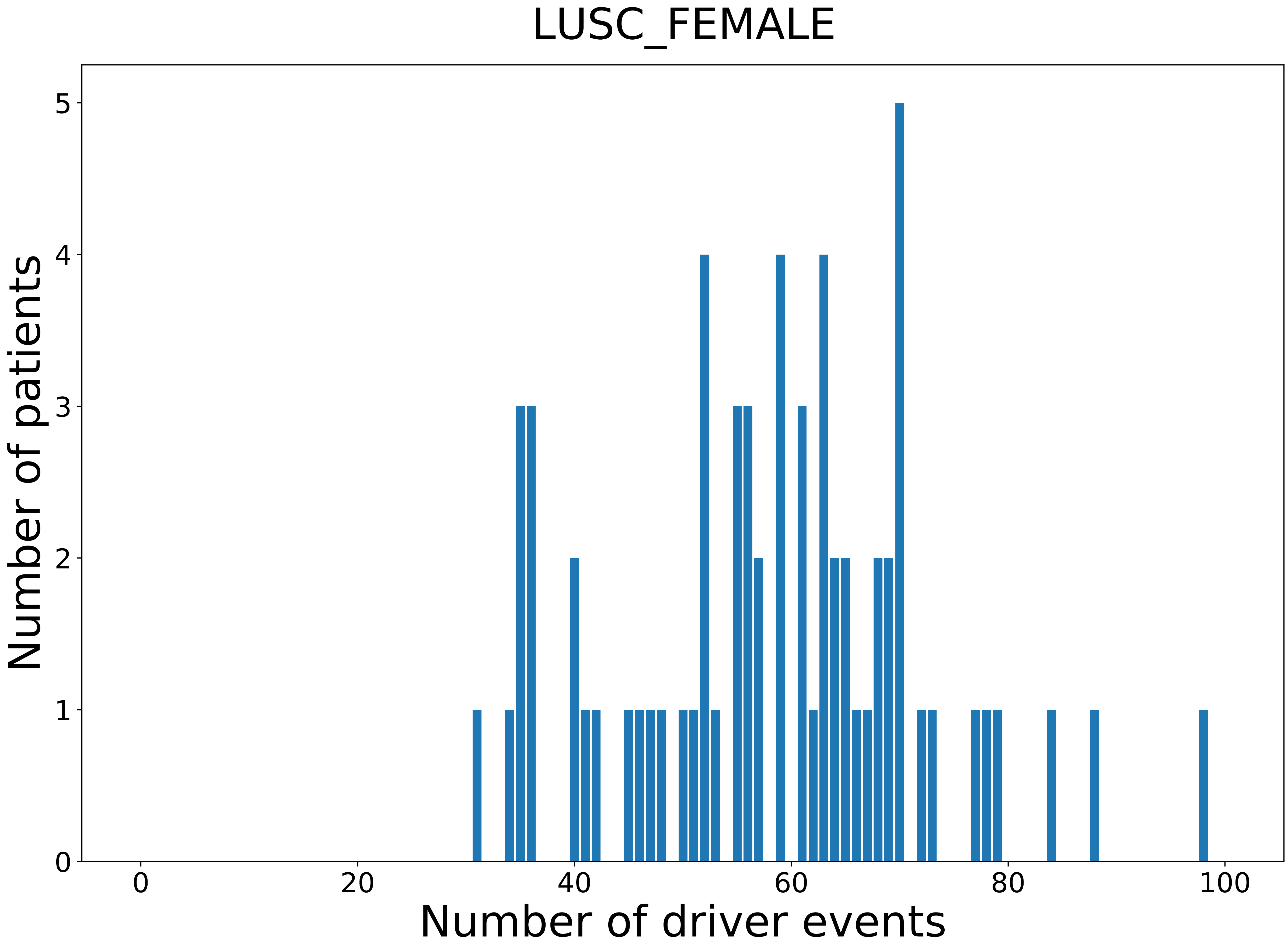

### 2021_8_22_13_42_LUSC_MALE.pdf

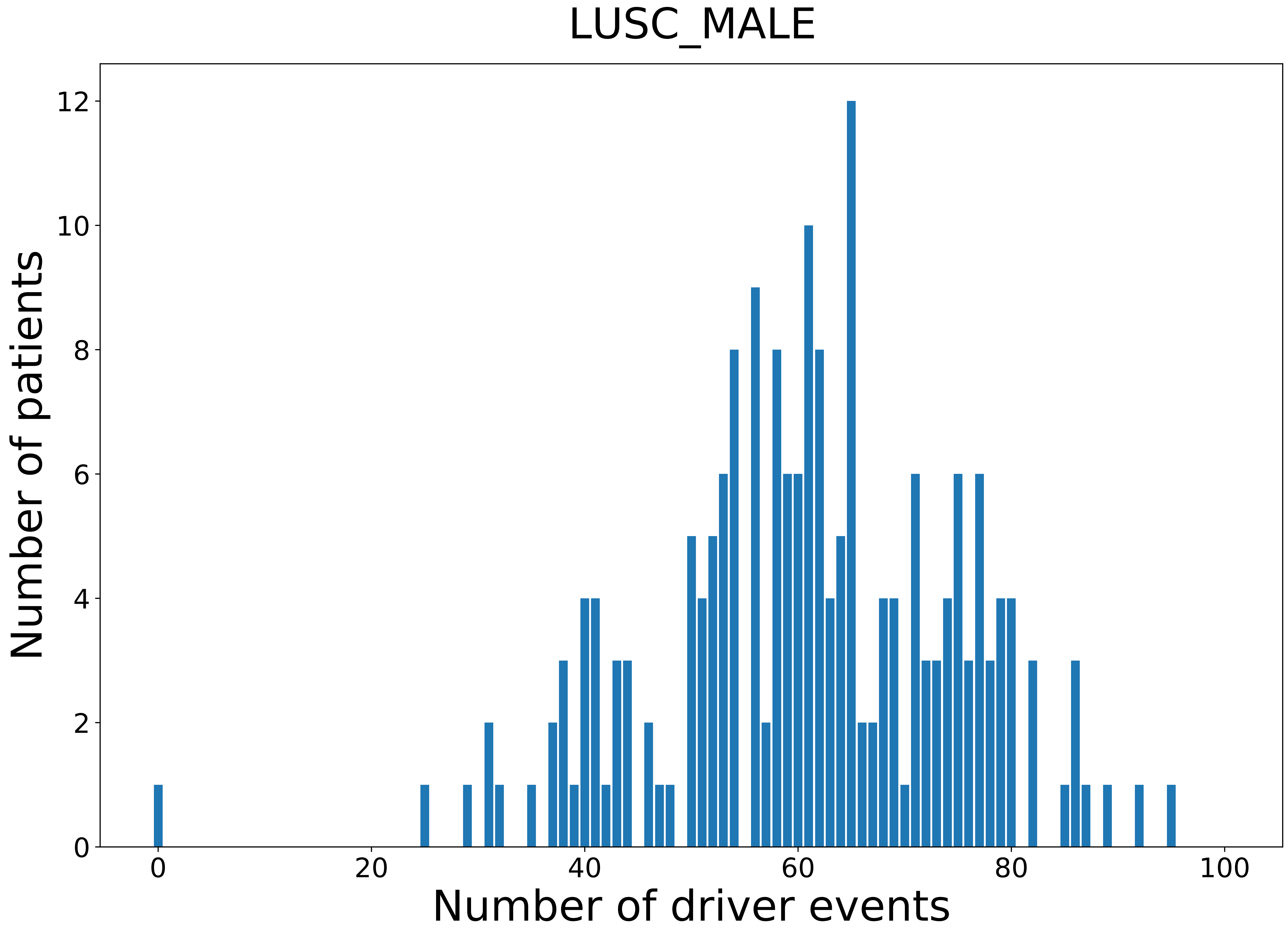

### 2021_8_22_13_42_MESO.pdf

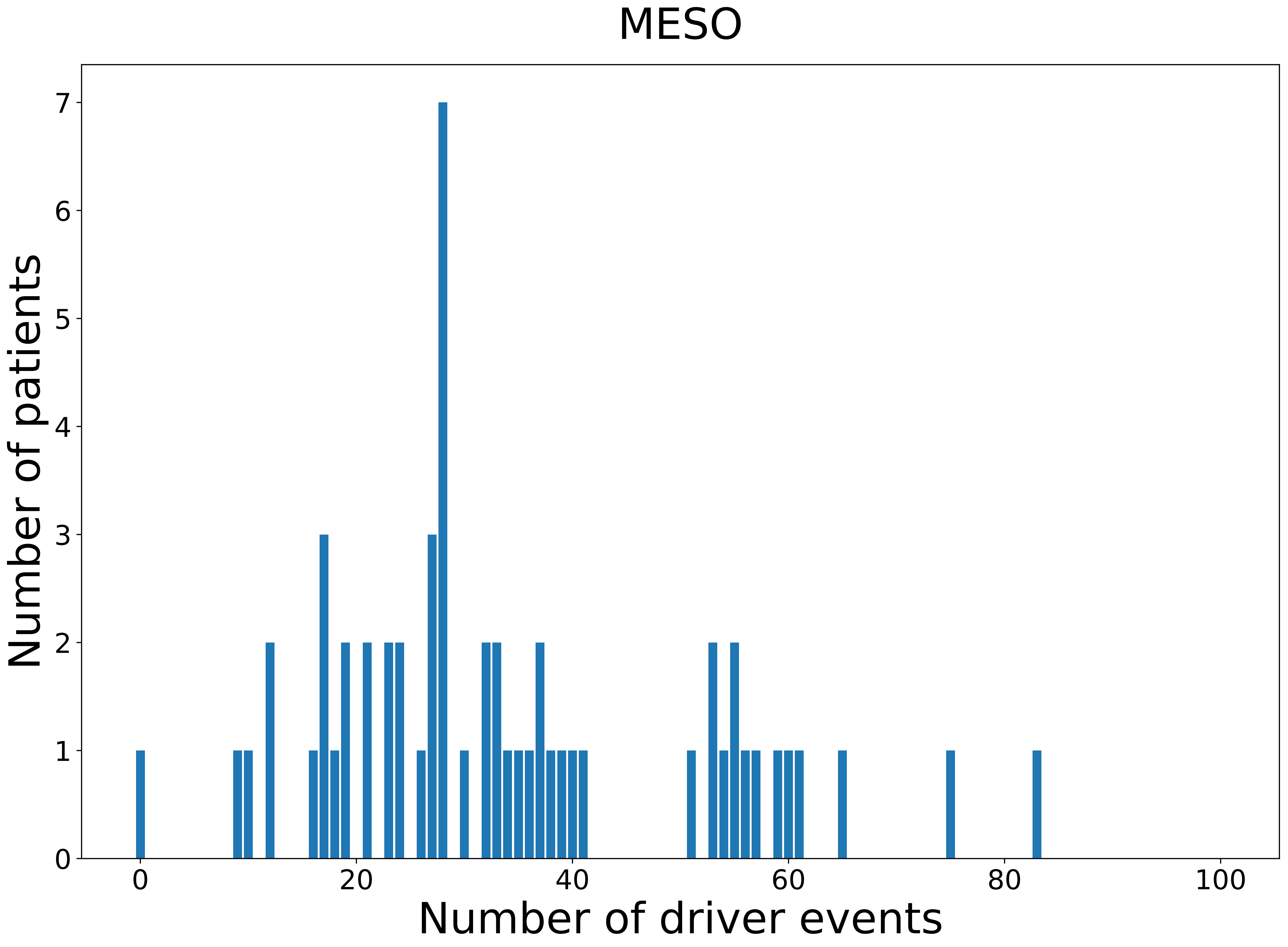

### 2021_8_22_13_42_MESO_FEMALE.pdf

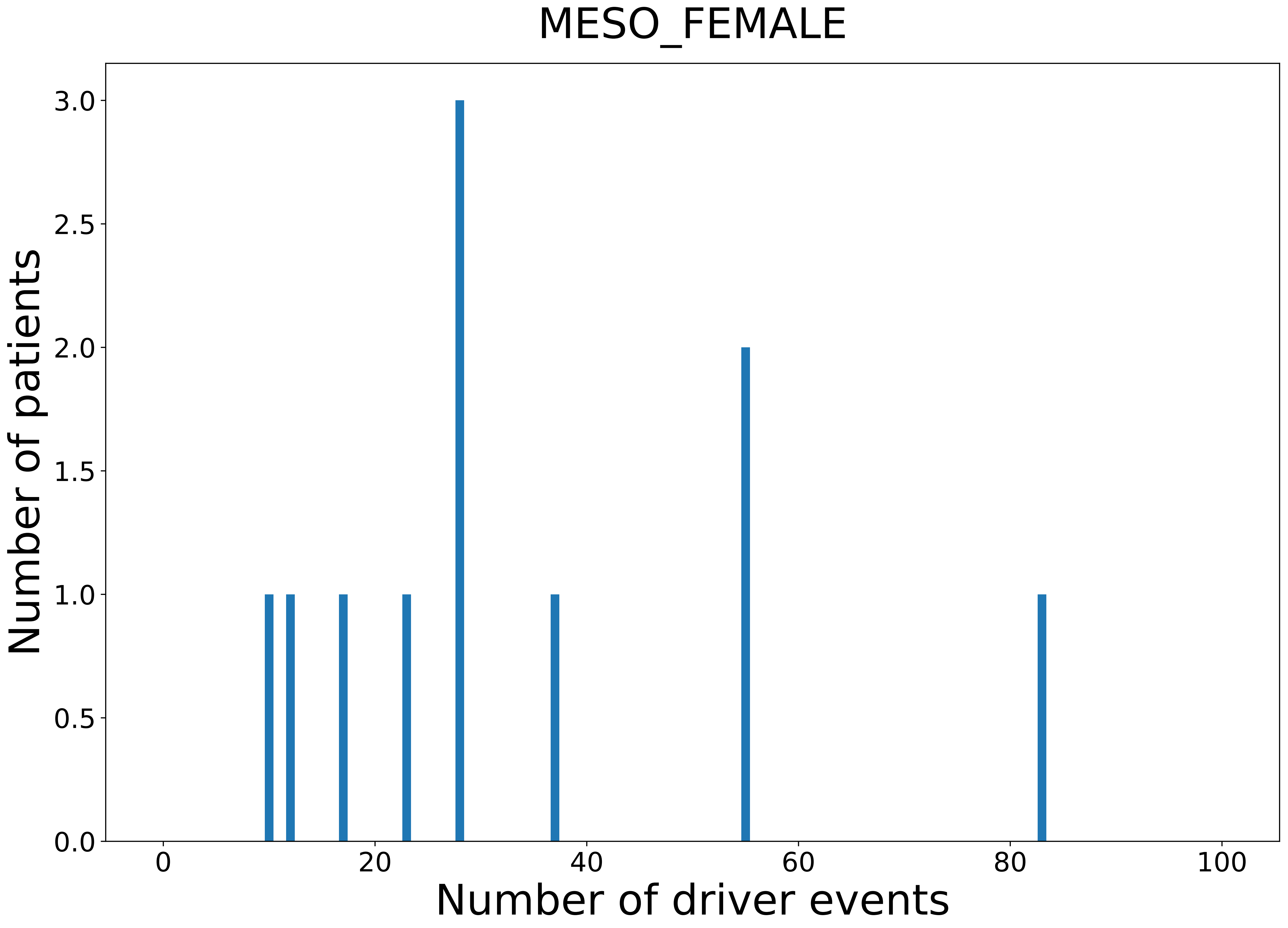

### 2021_8_22_13_42_MESO_MALE.pdf

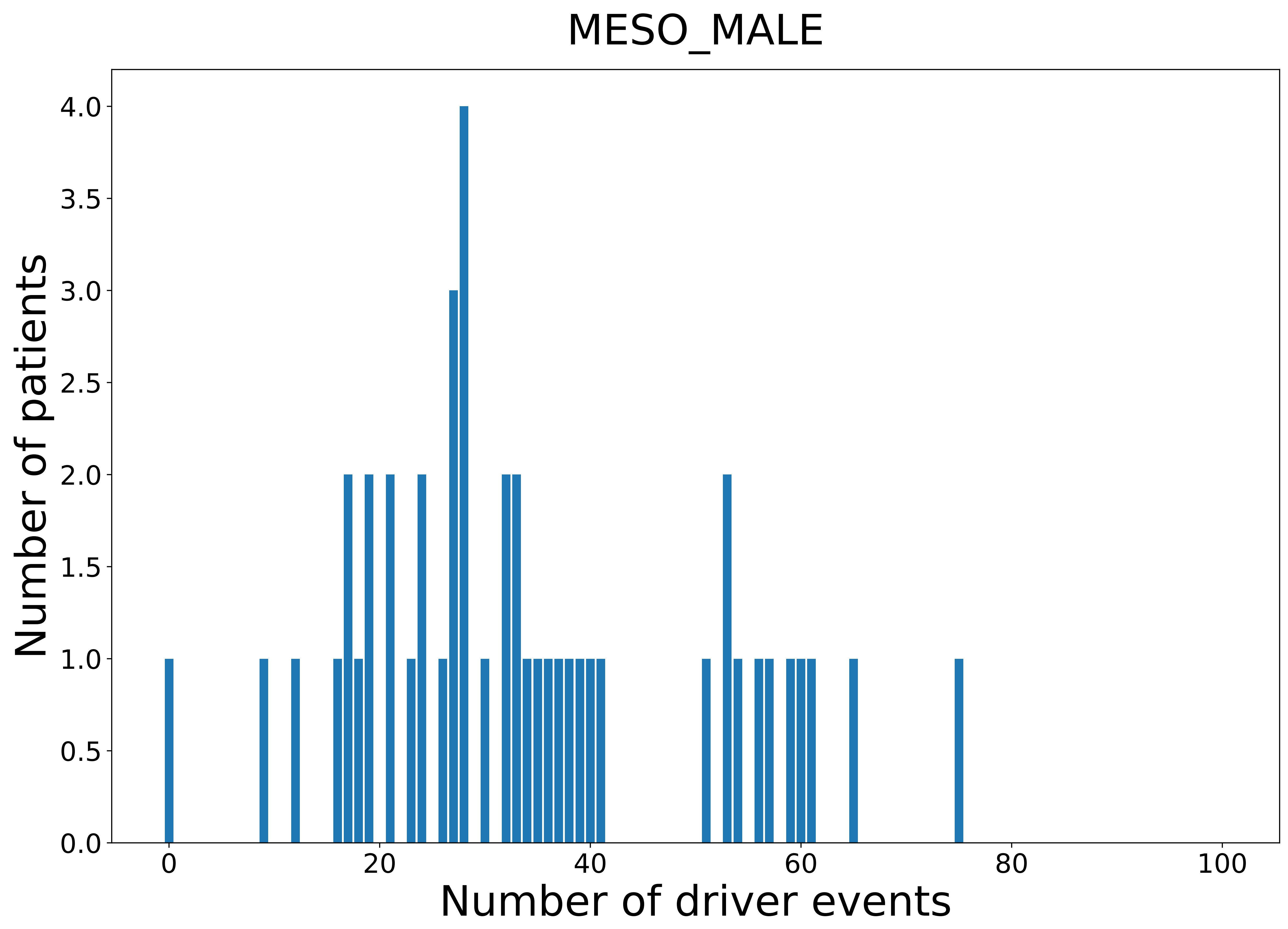

### 2021_8_22_13_42_OV.pdf

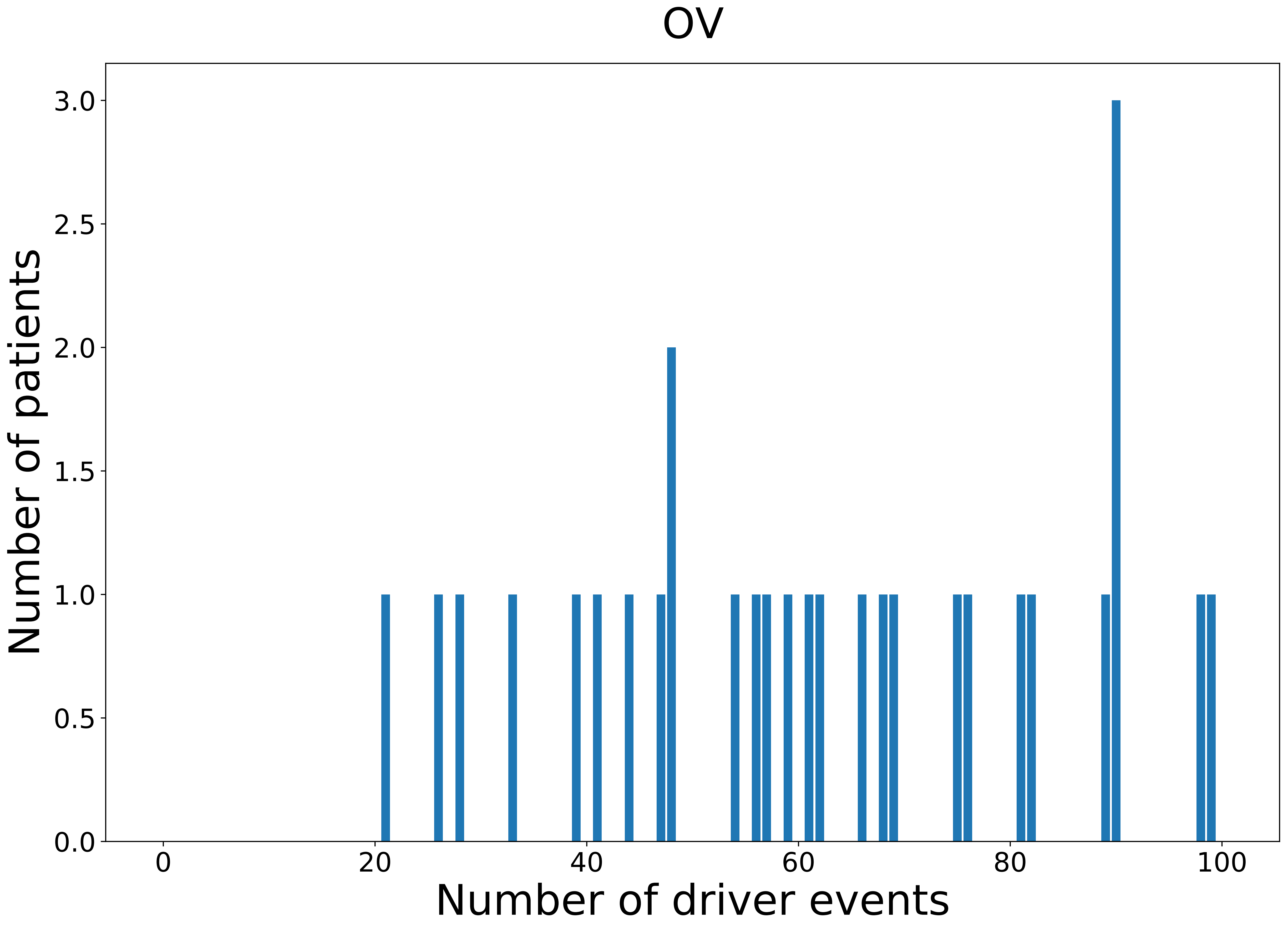

### 2021_8_22_13_42_OV_FEMALE.pdf

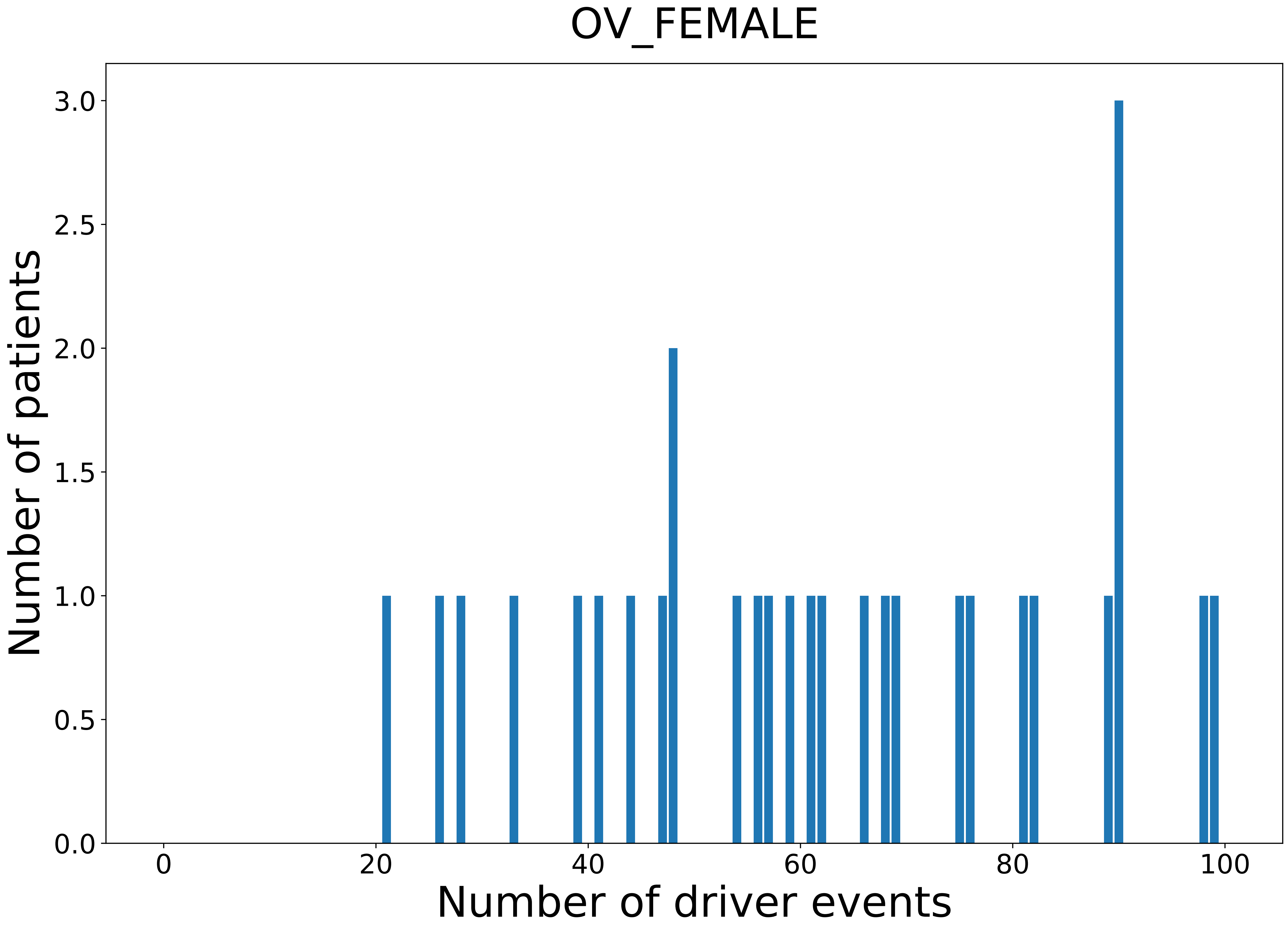

### 2021_8_22_13_42_PAAD.pdf

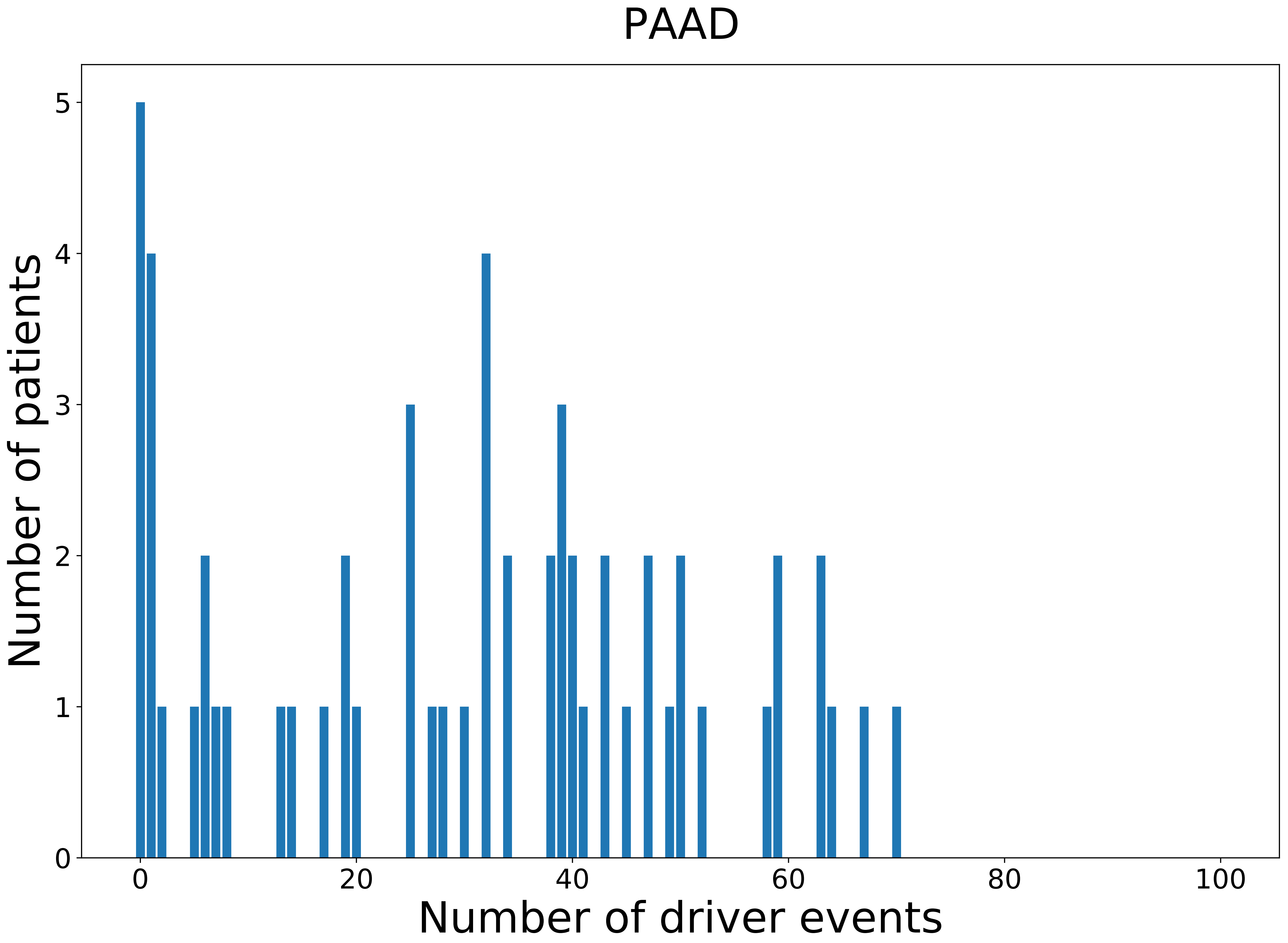

### 2021_8_22_13_42_PAAD_FEMALE.pdf

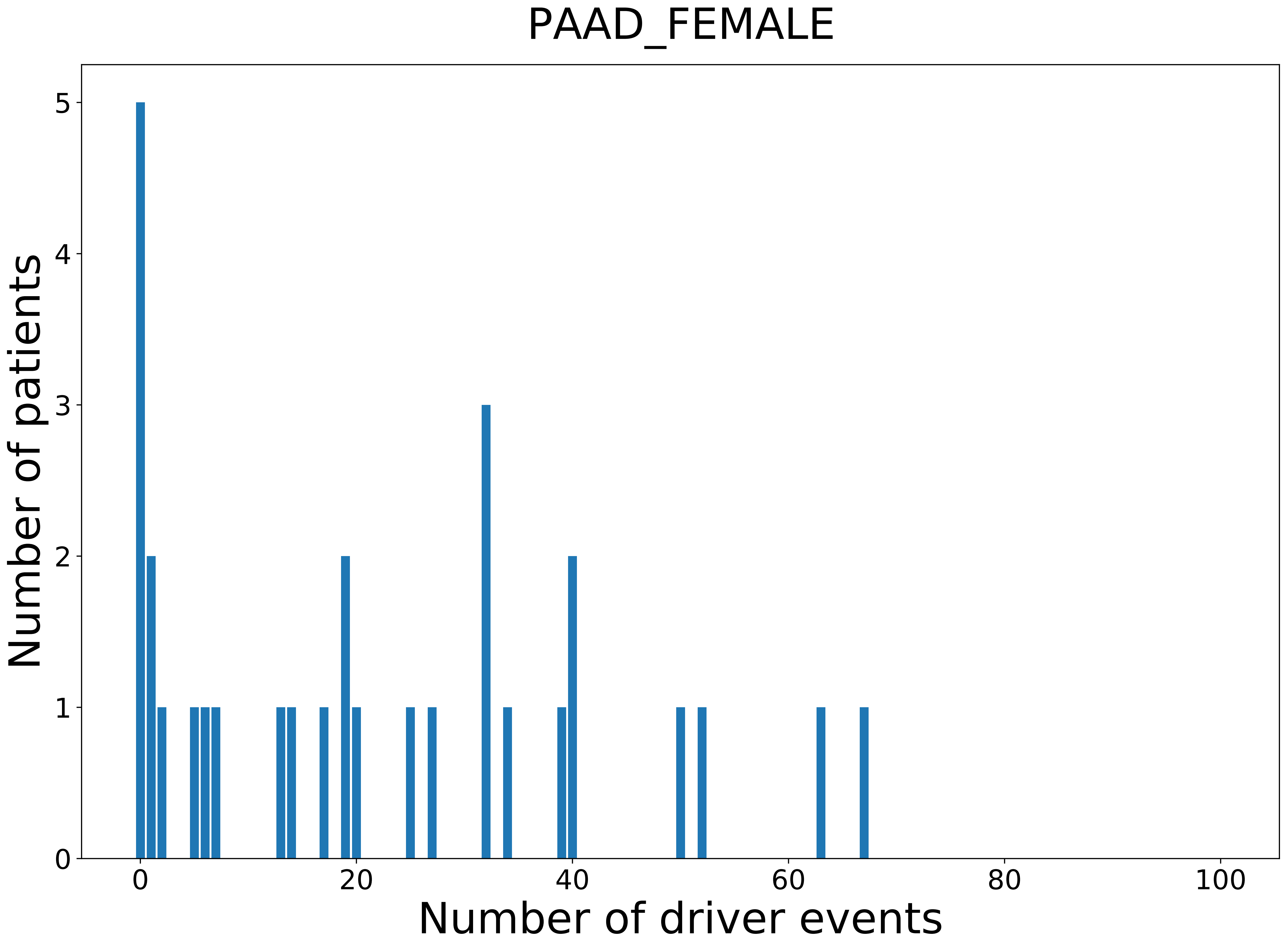

### 2021_8_22_13_42_PAAD_MALE.pdf

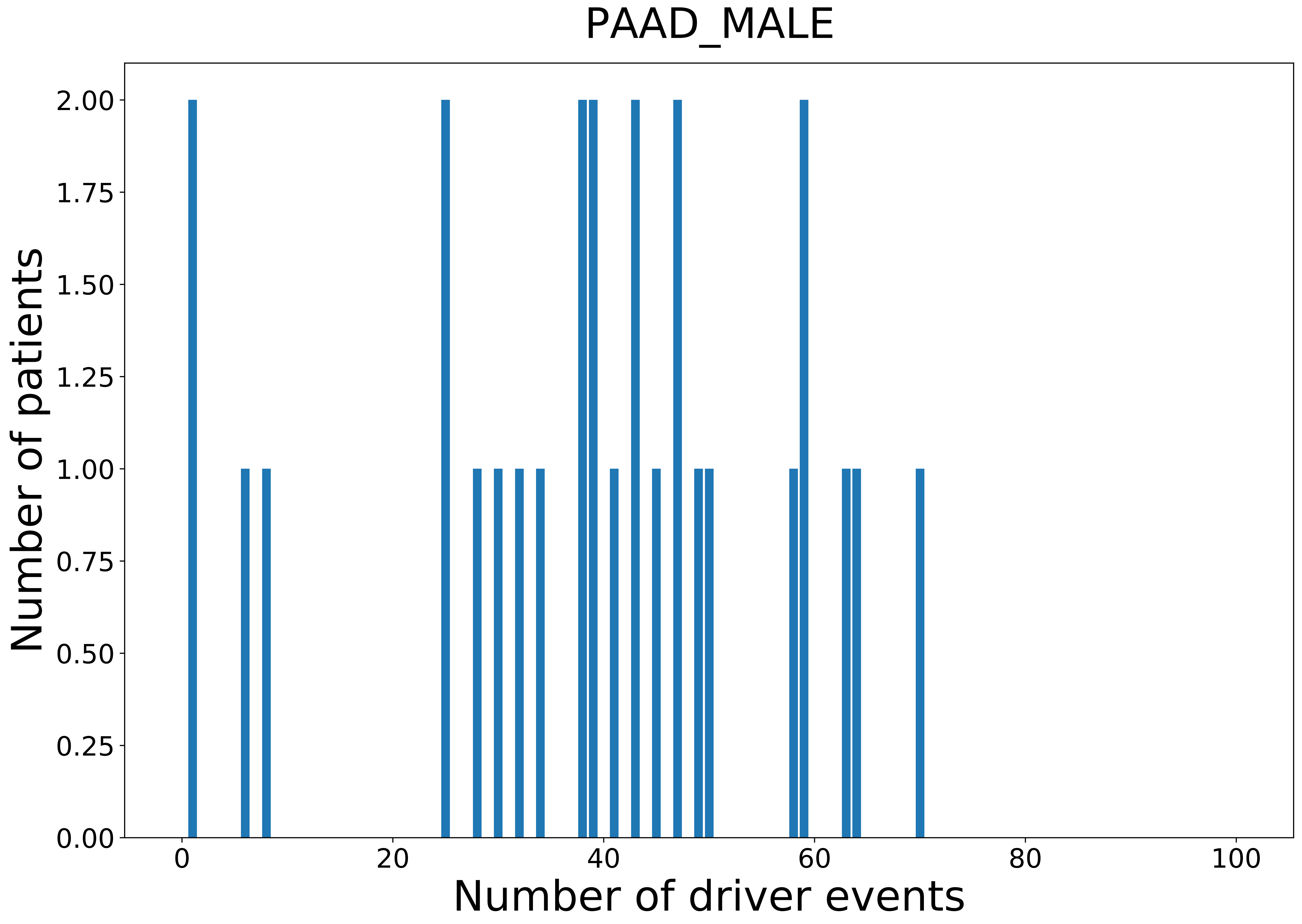

### 2021_8_22_13_42_PANCAN.pdf

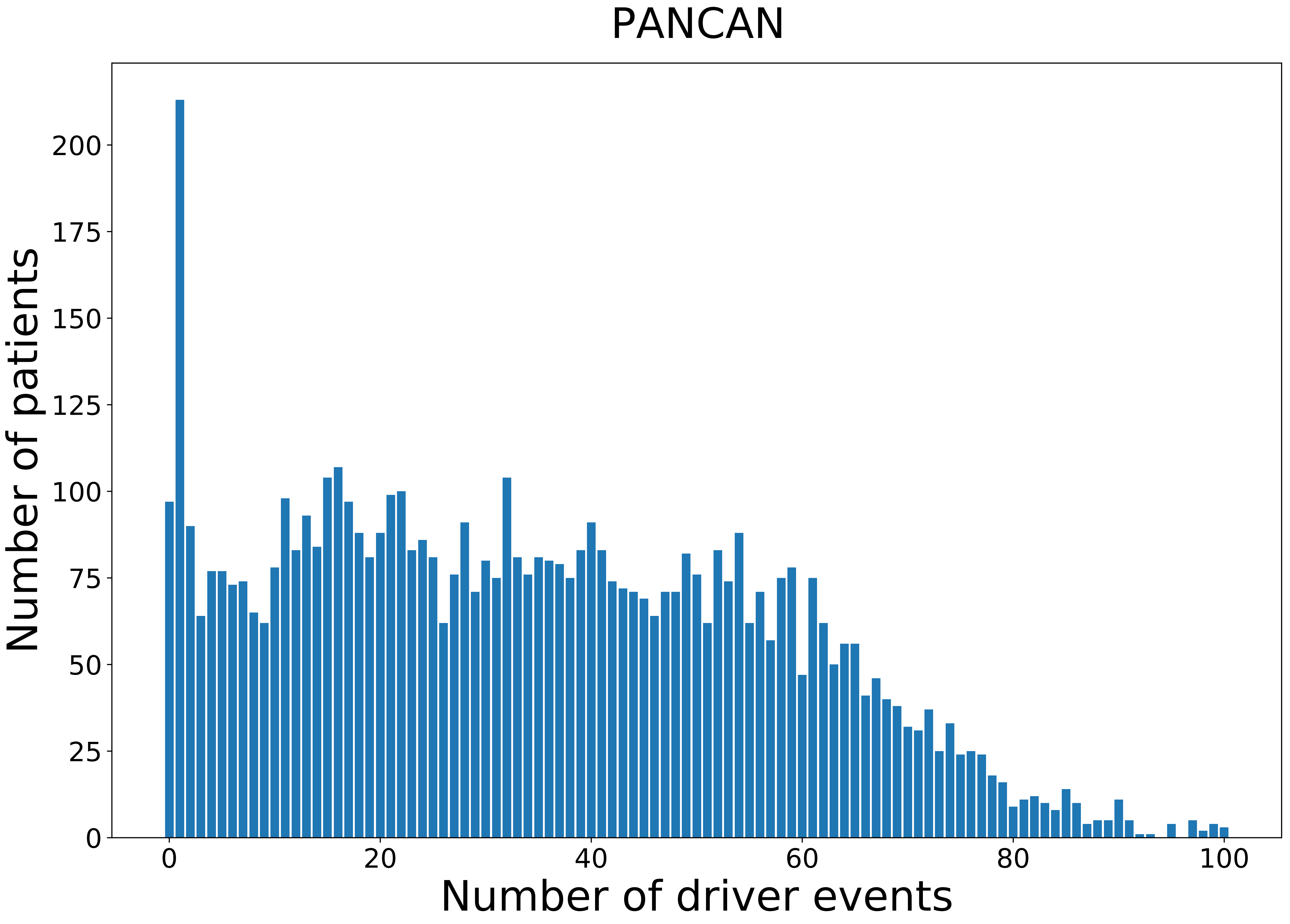

### 2021_8_22_13_42_PANCAN_FEMALE.pdf

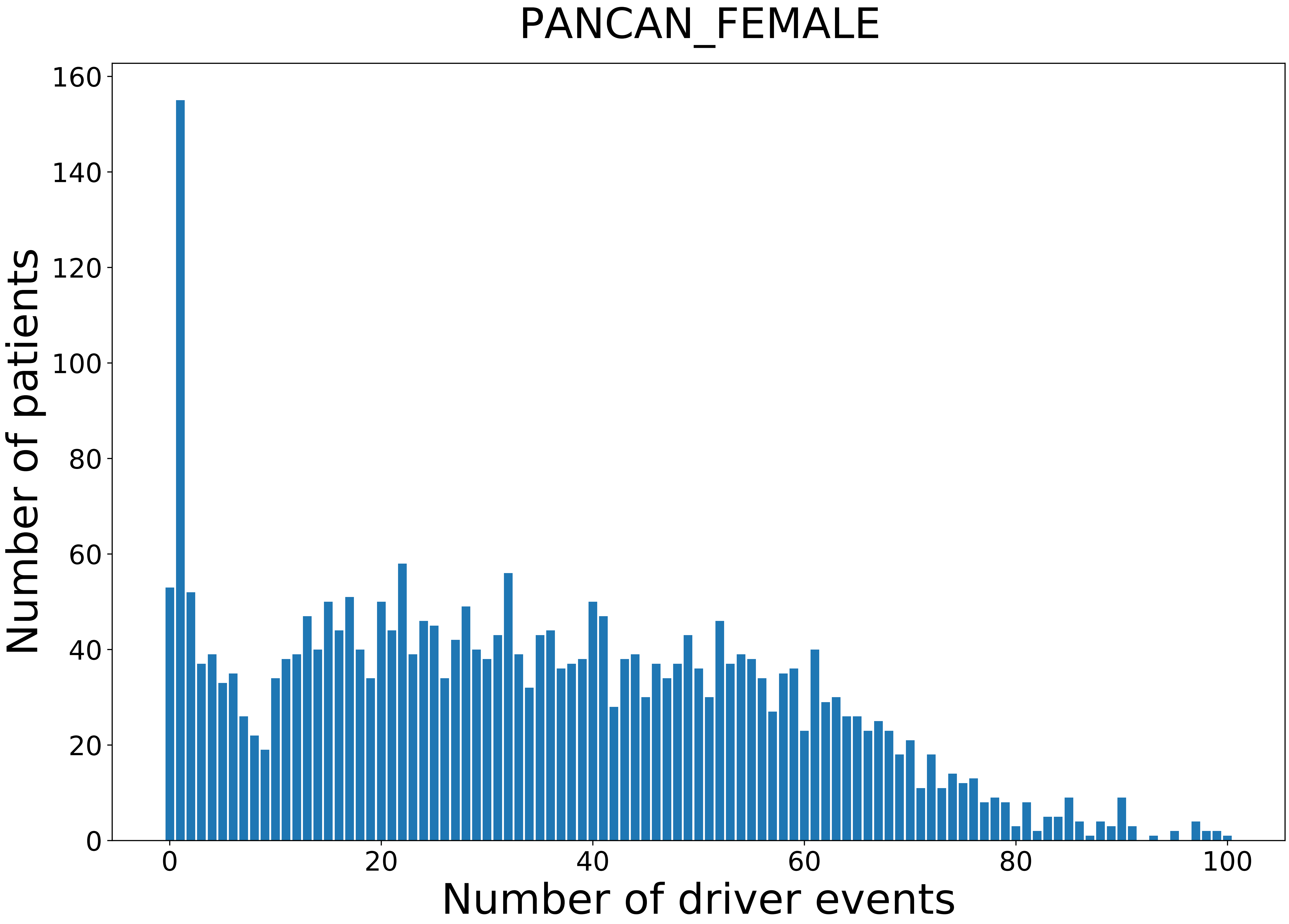

### 2021_8_22_13_42_PANCAN_MALE.pdf

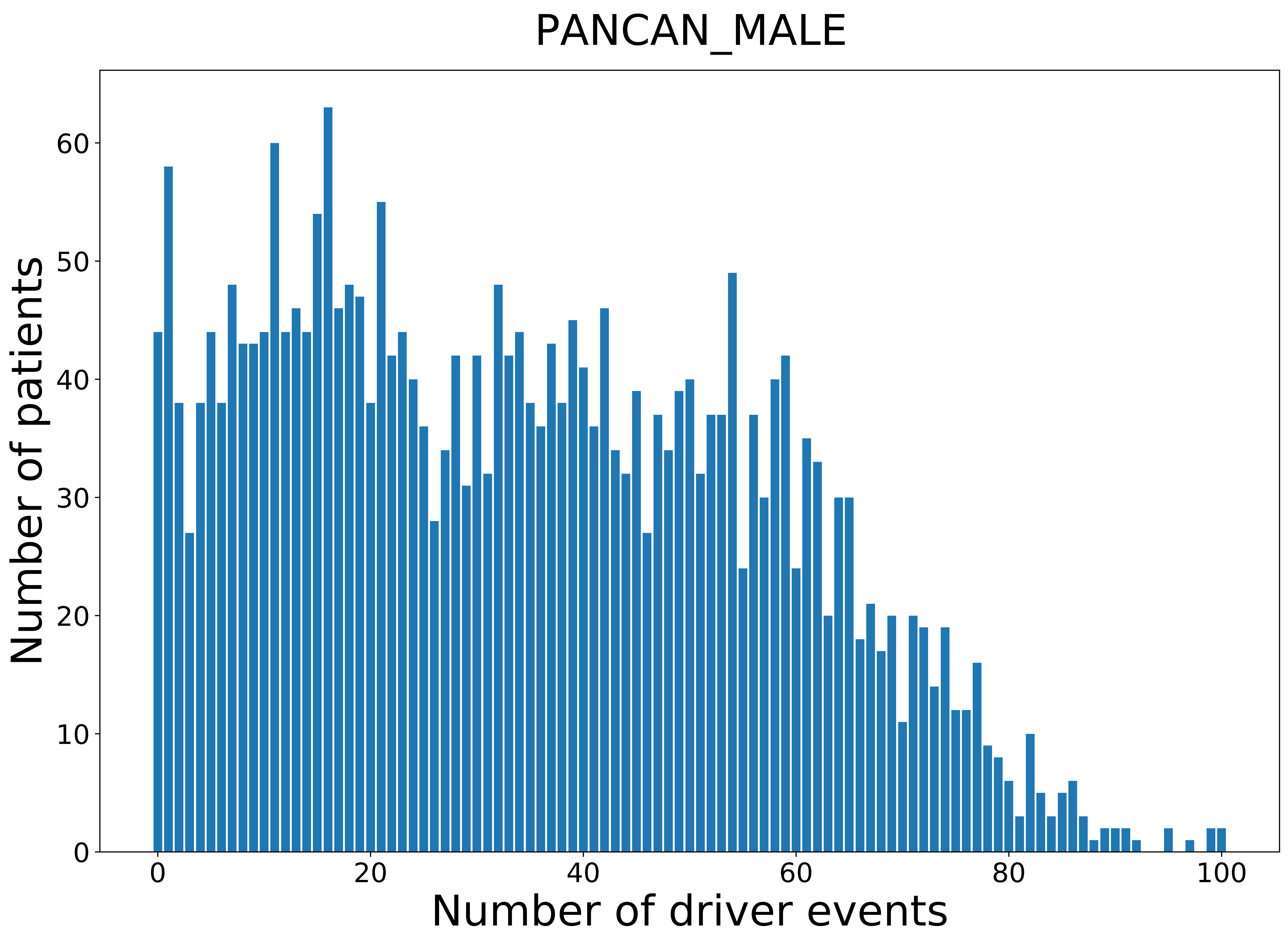

### 2021_8_22_13_42_PCPG.pdf

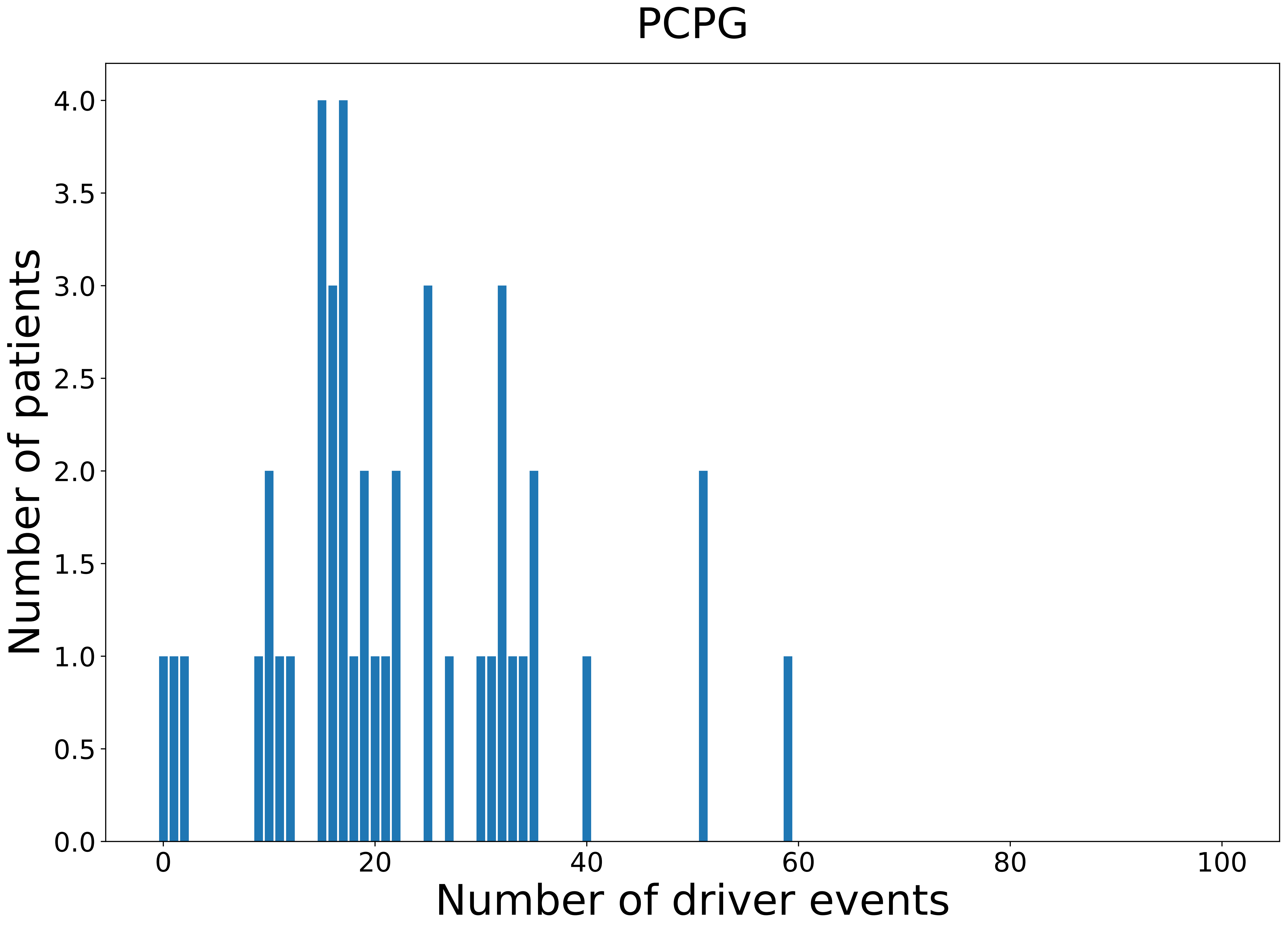

### 2021_8_22_13_42_PCPG_FEMALE.pdf

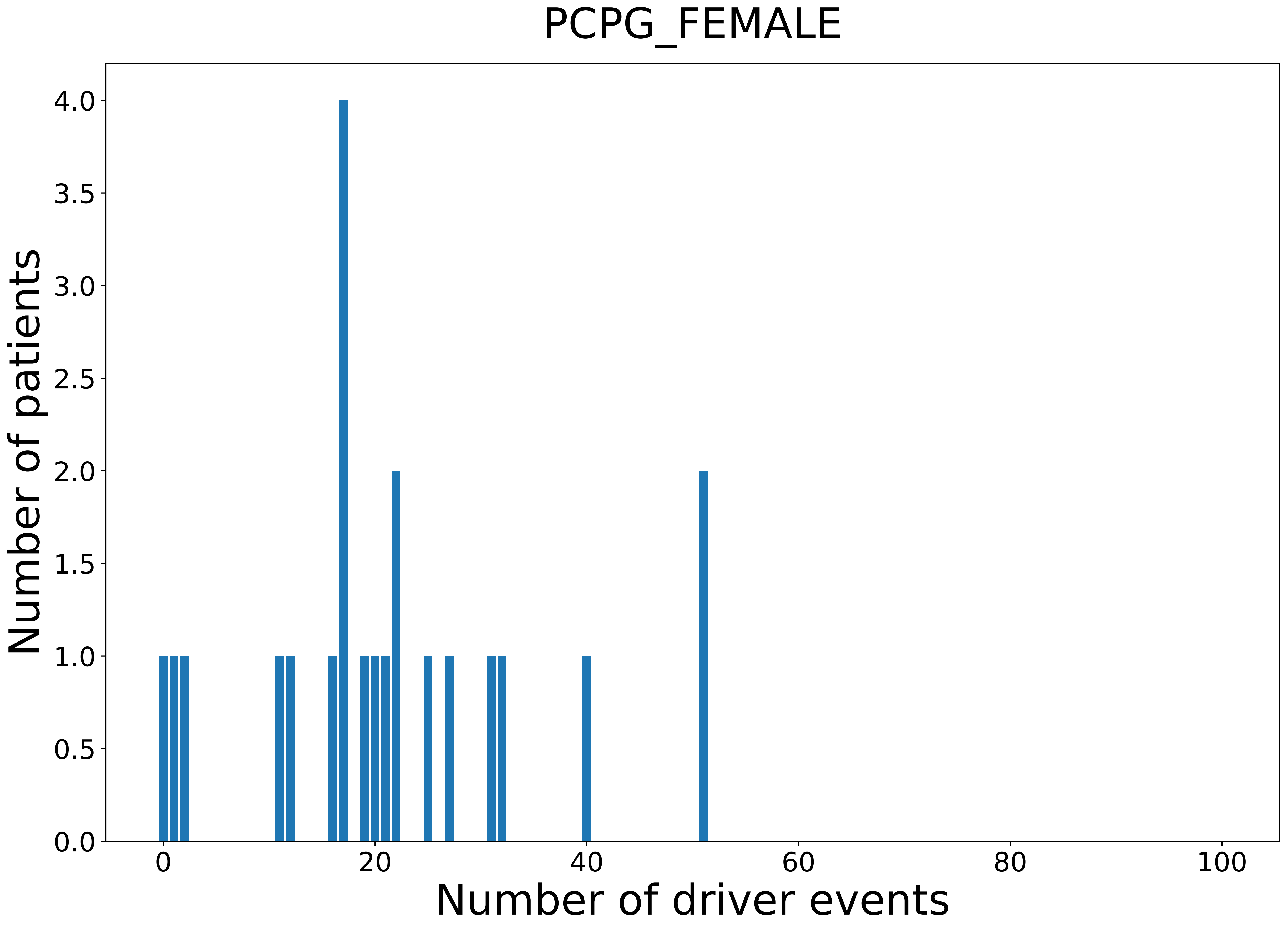

### 2021_8_22_13_42_PRAD.pdf

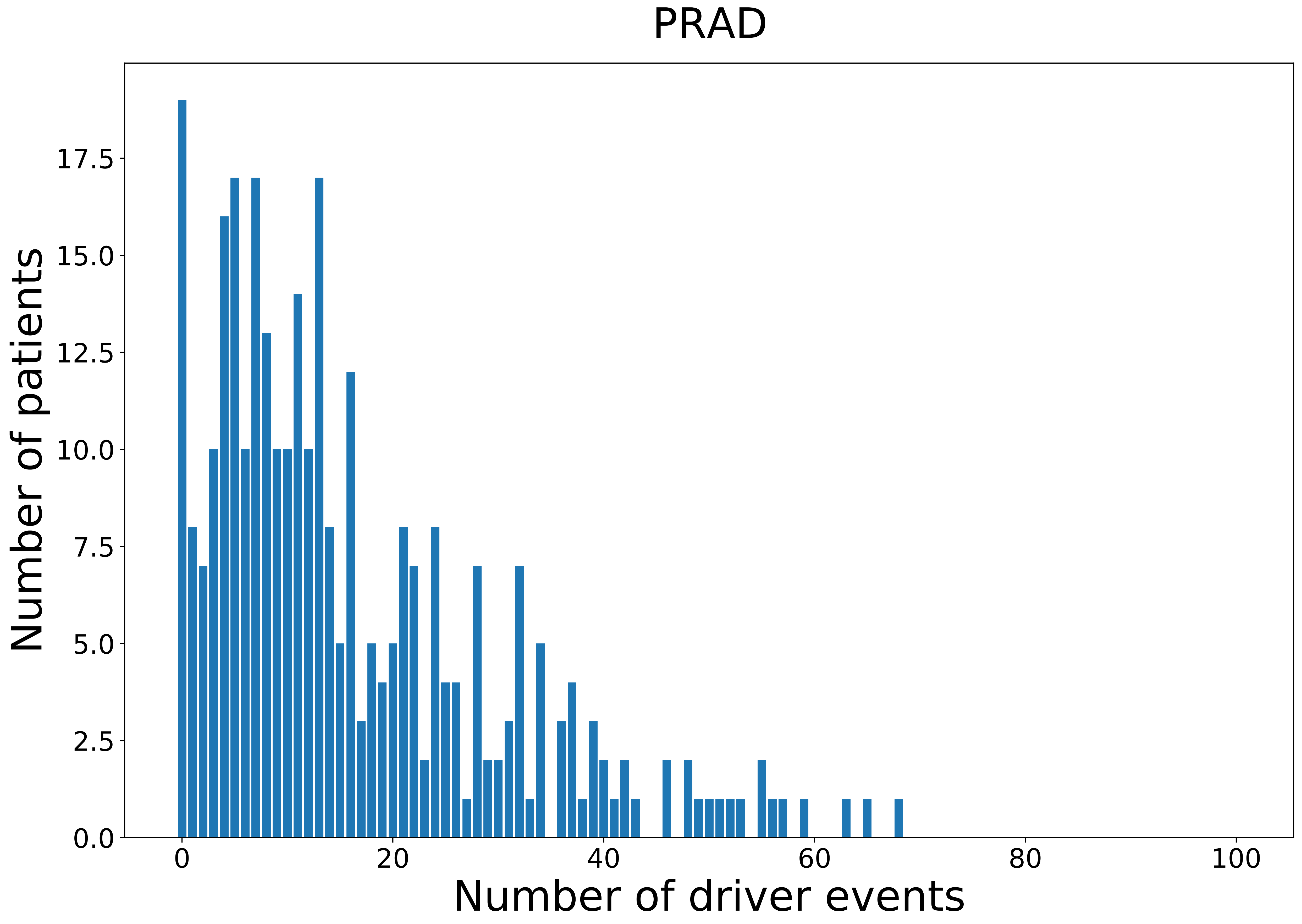

### 2021_8_22_13_42_PRAD_MALE.pdf

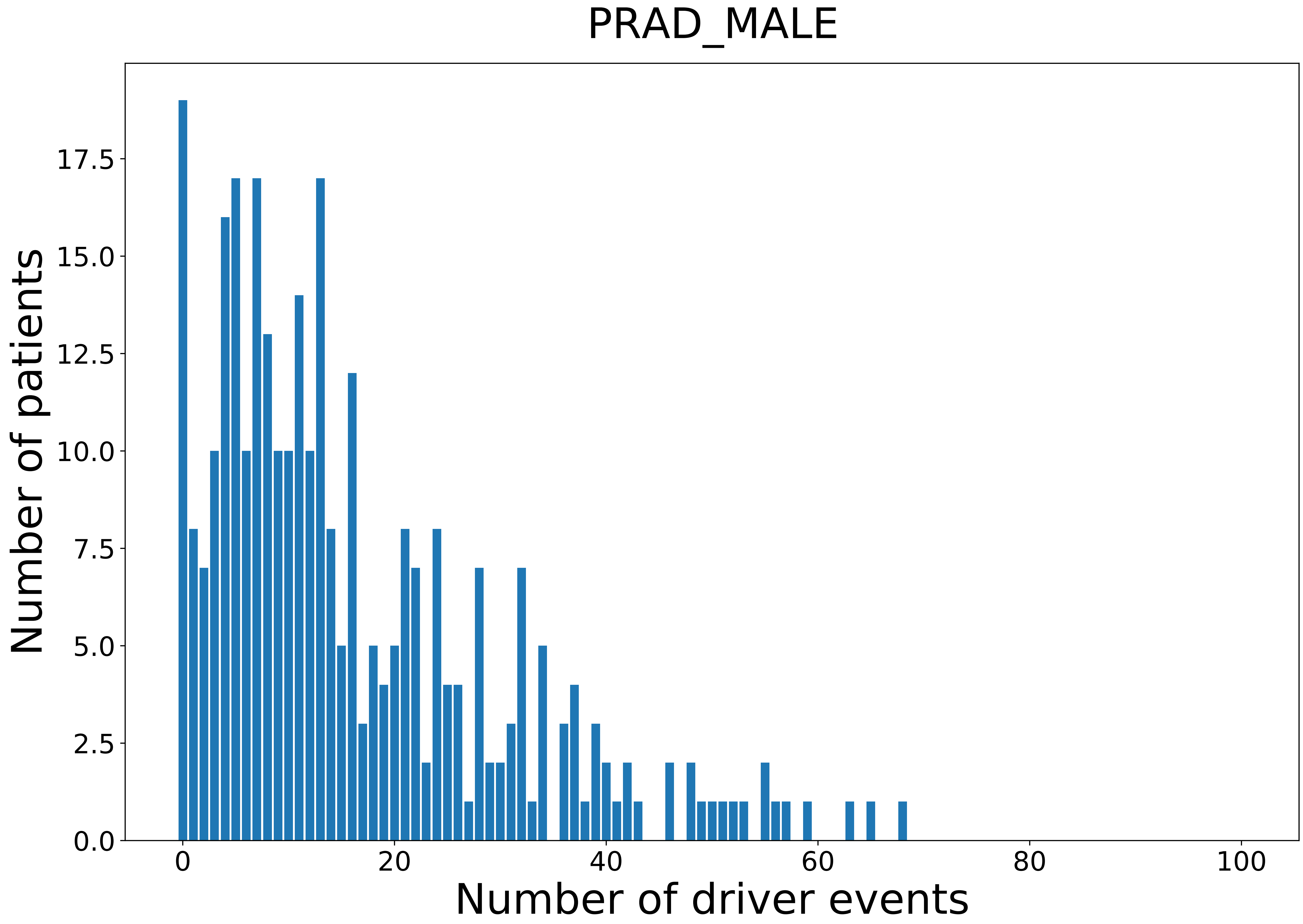

### 2021_8_22_13_42_READ.pdf

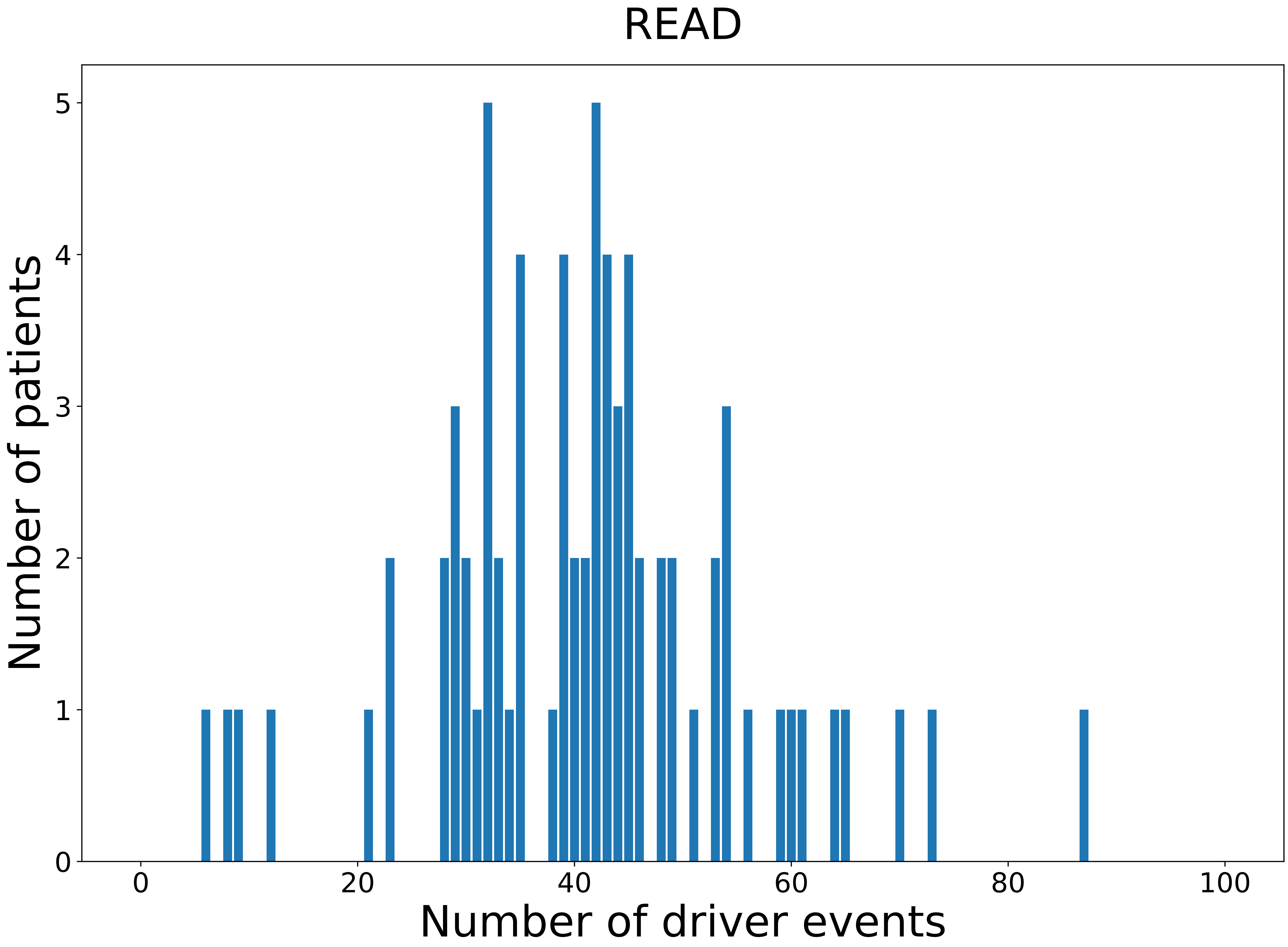

### 2021_8_22_13_42_READ_FEMALE.pdf

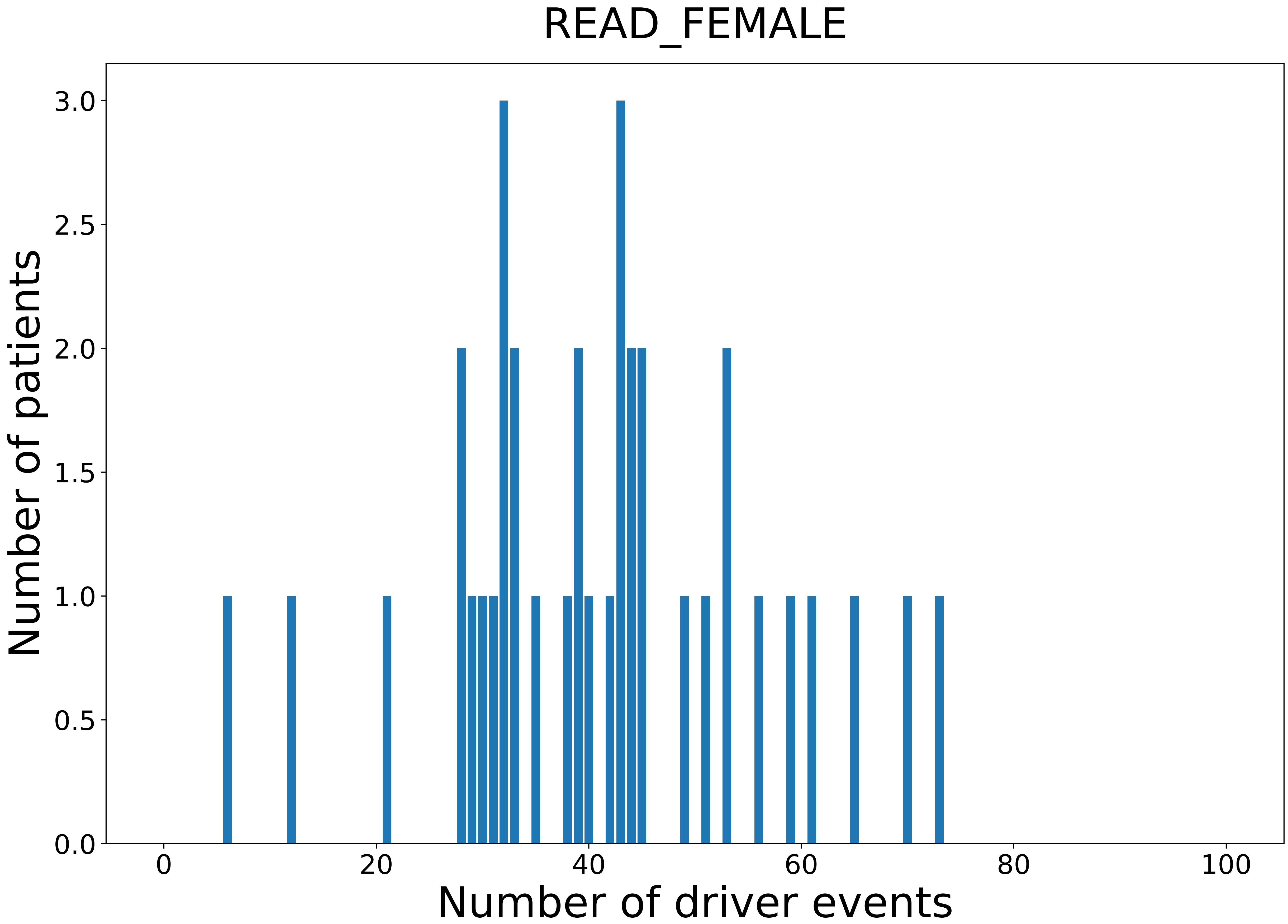

### 2021_8_22_13_42_SARC.pdf

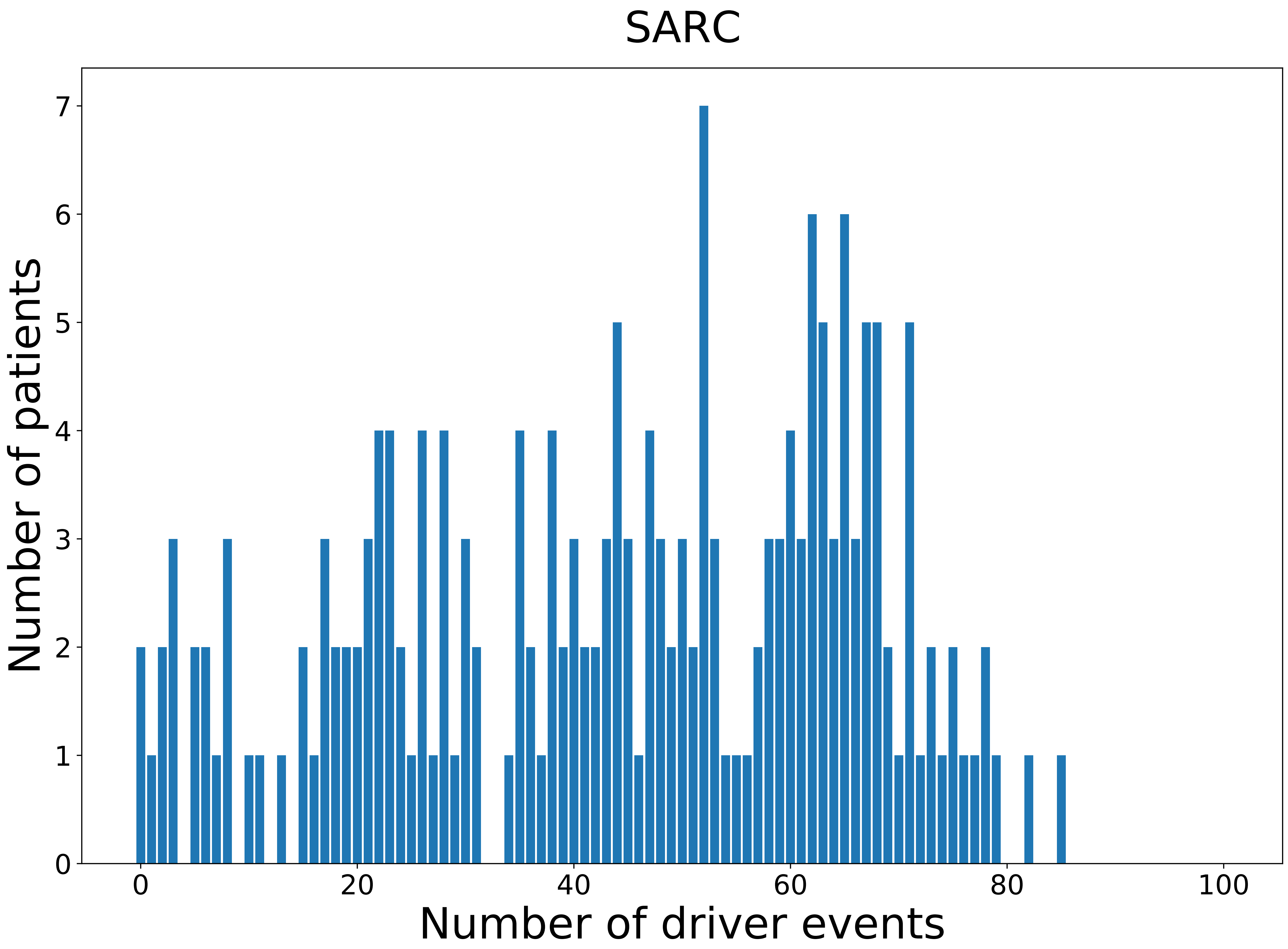

### 2021_8_22_13_42_SARC_FEMALE.pdf

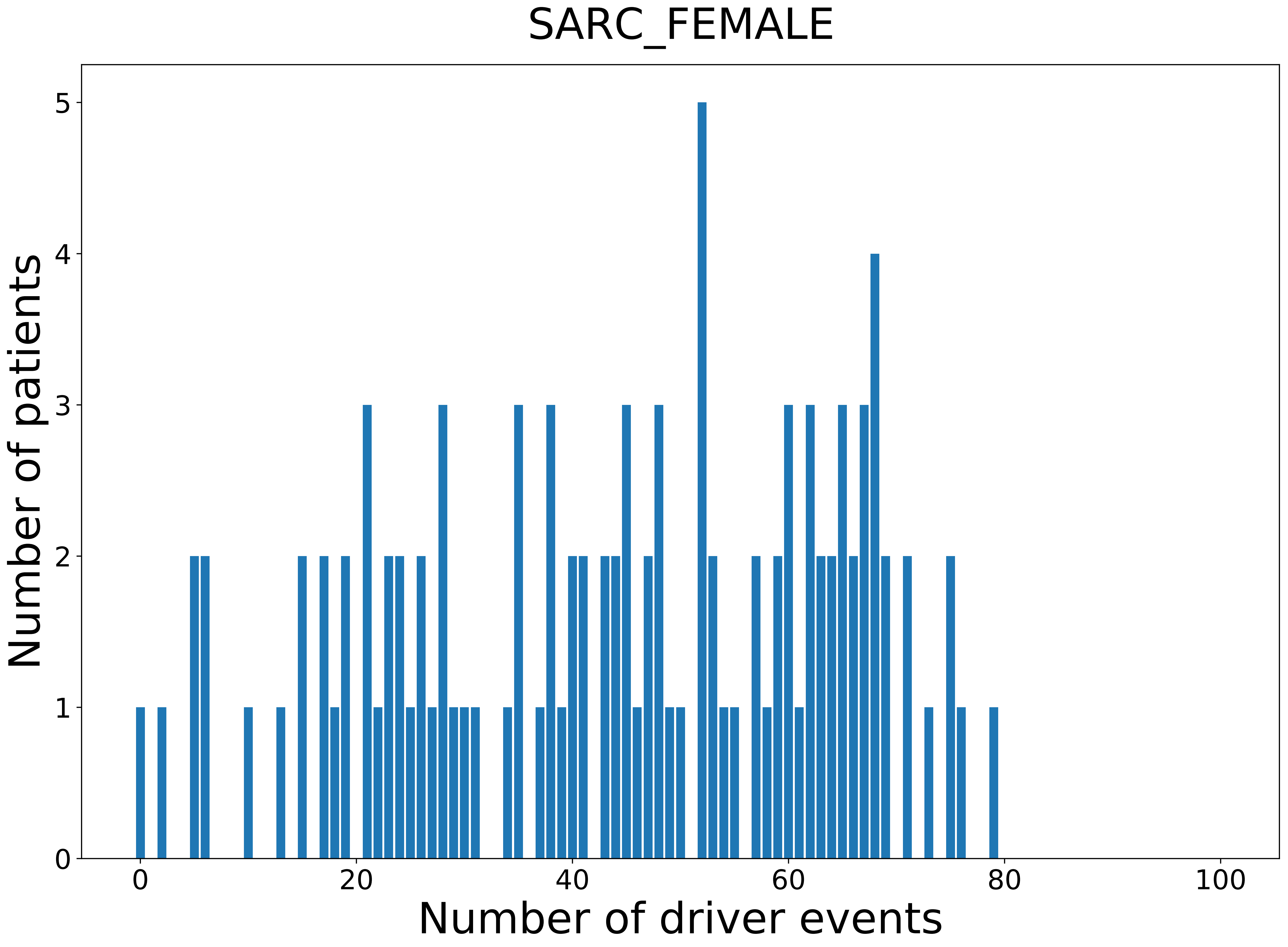

### 2021_8_22_13_42_SARC_MALE.pdf

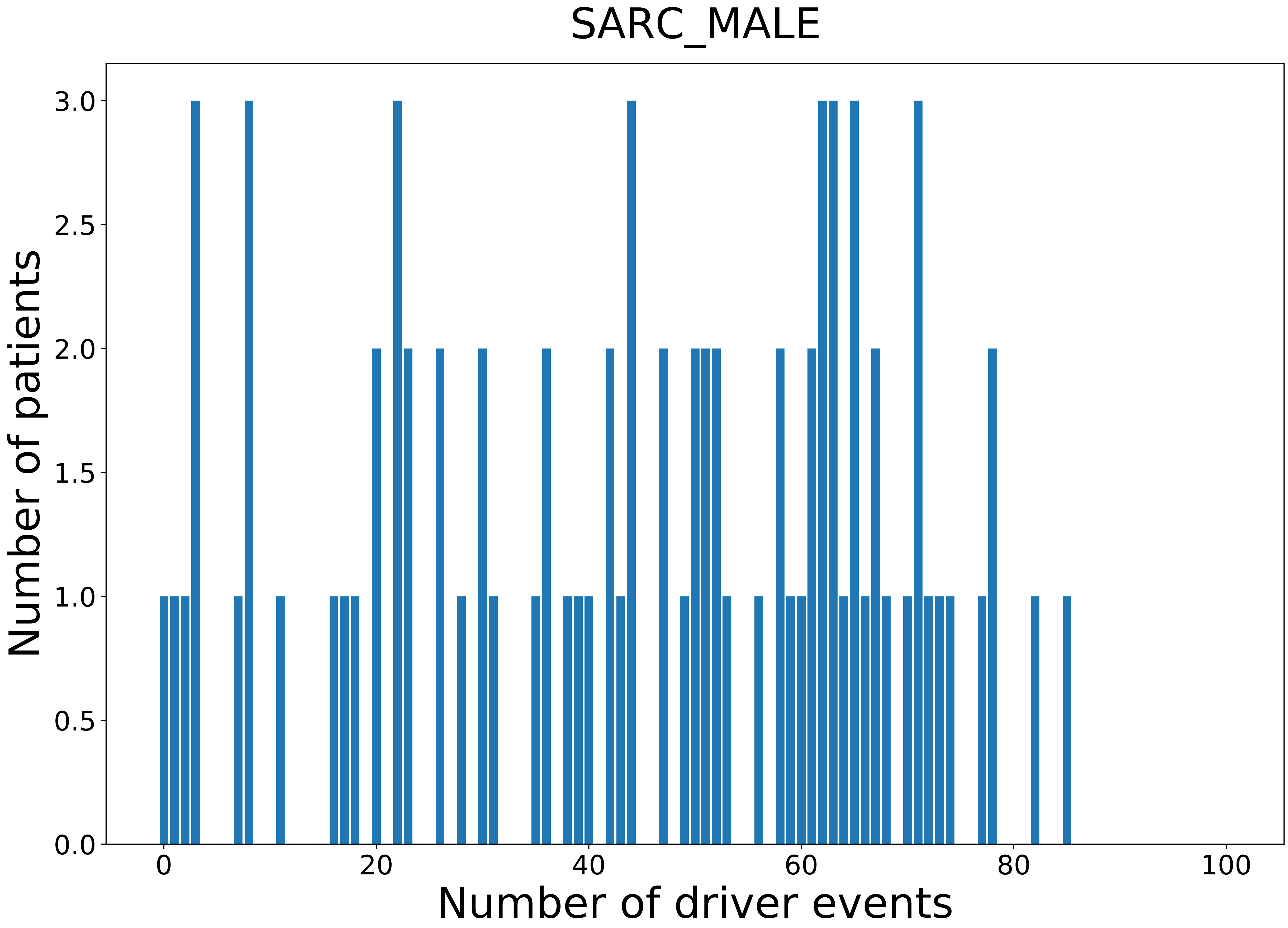

### 2021_8_22_13_42_SKCM.pdf

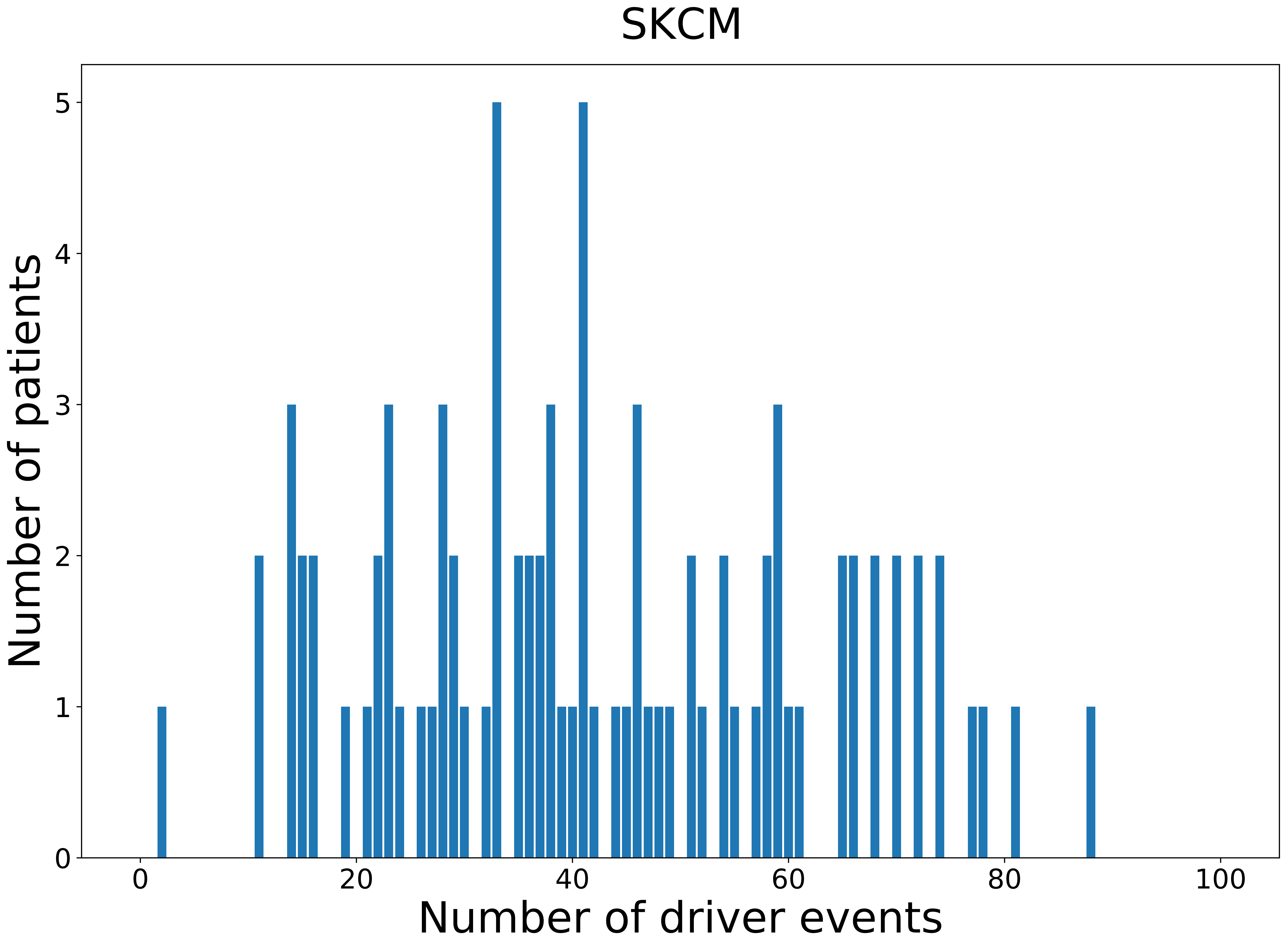

### 2021_8_22_13_42_SKCM_FEMALE.pdf

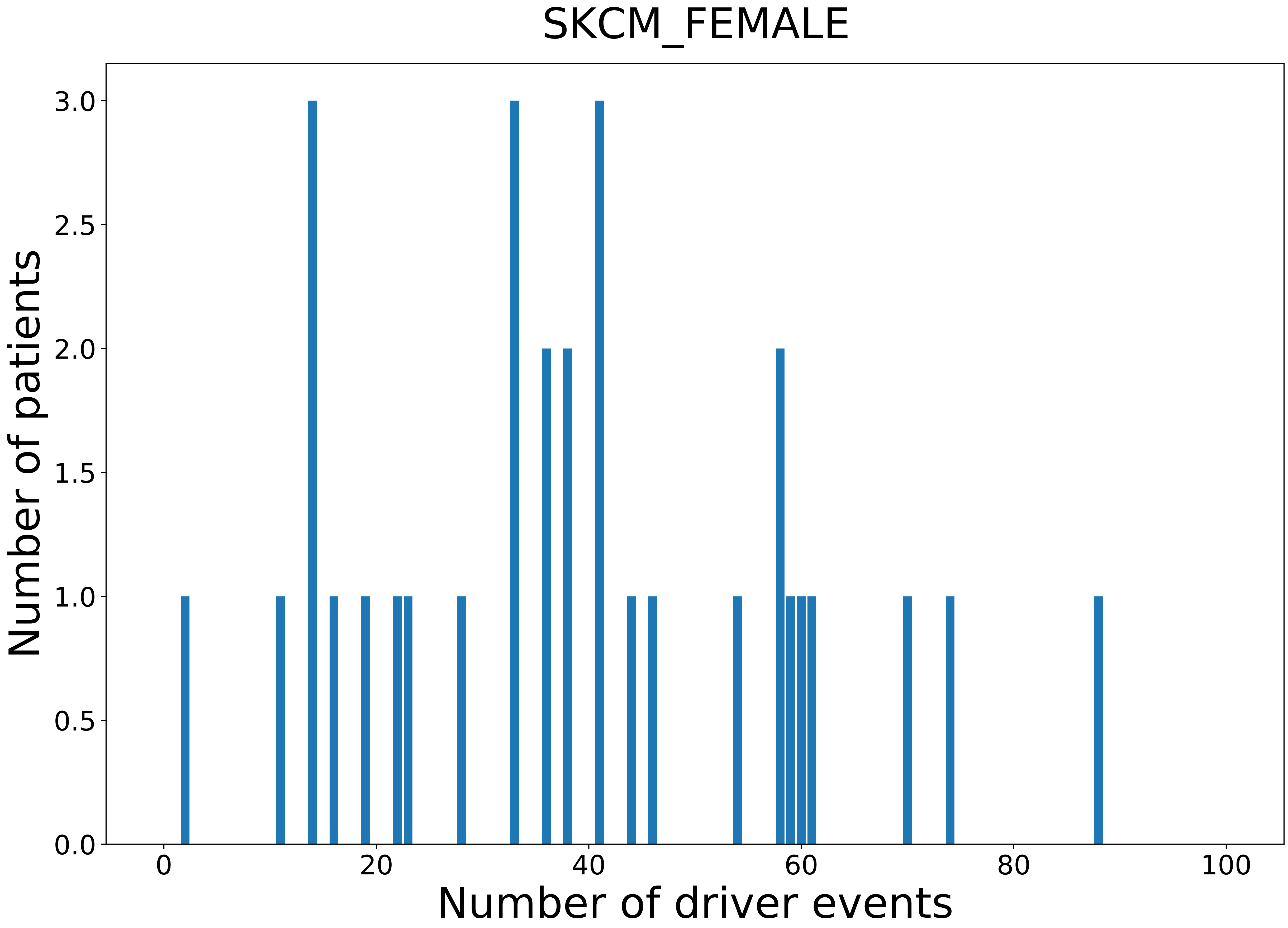

### 2021_8_22_13_42_SKCM_MALE.pdf

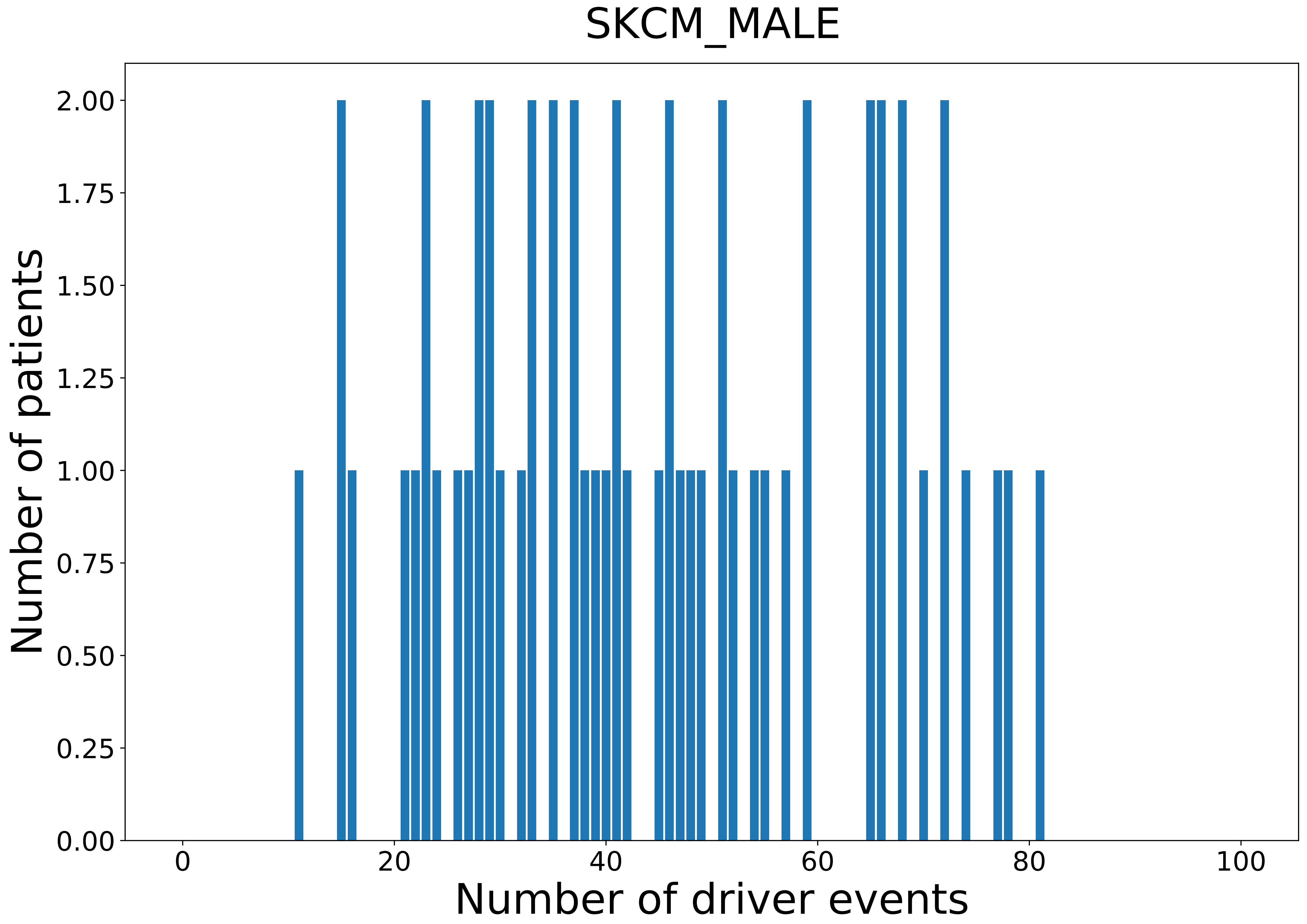

### 2021_8_22_13_42_STAD.pdf

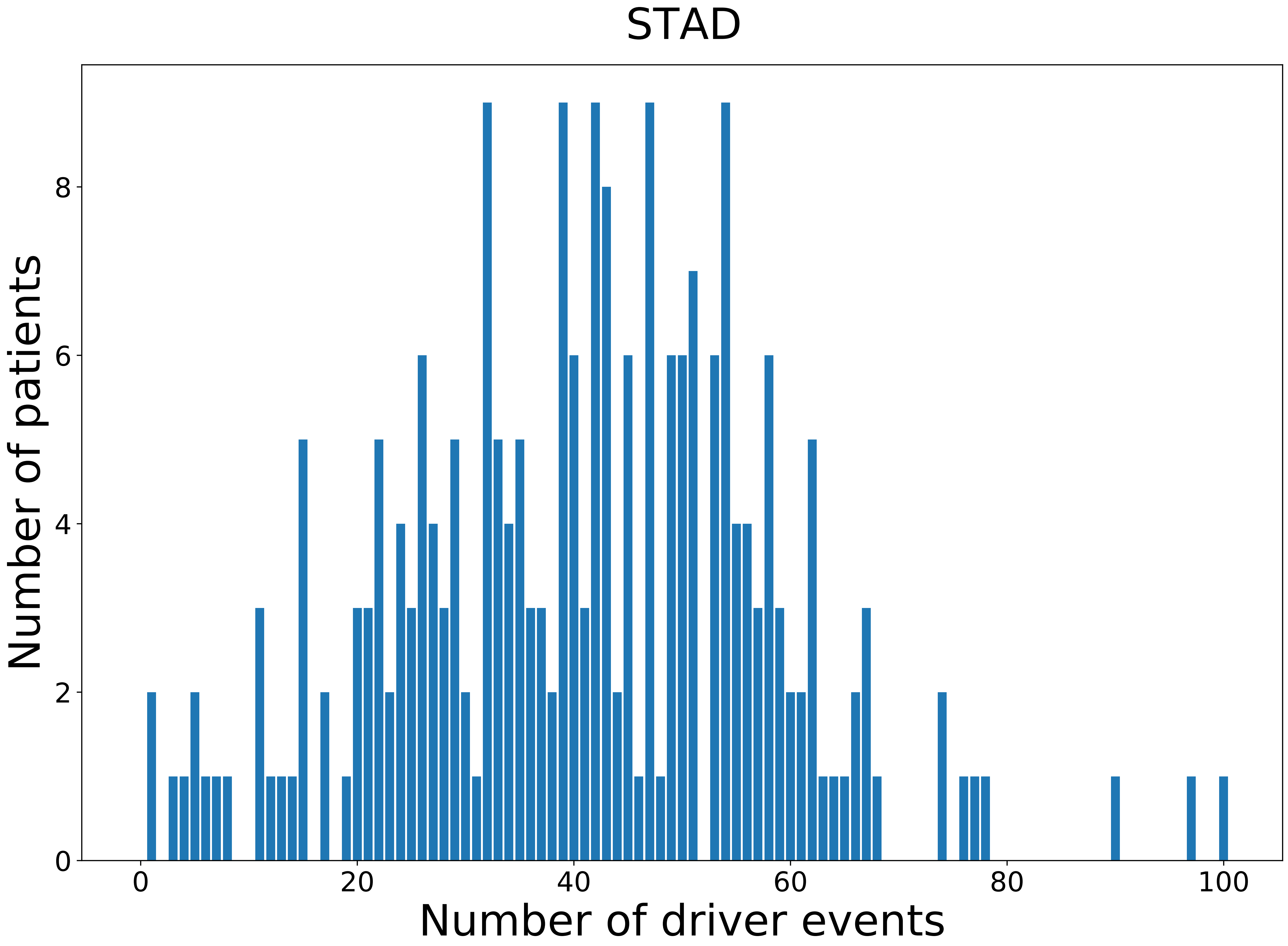

### 2021_8_22_13_42_STAD_MALE.pdf

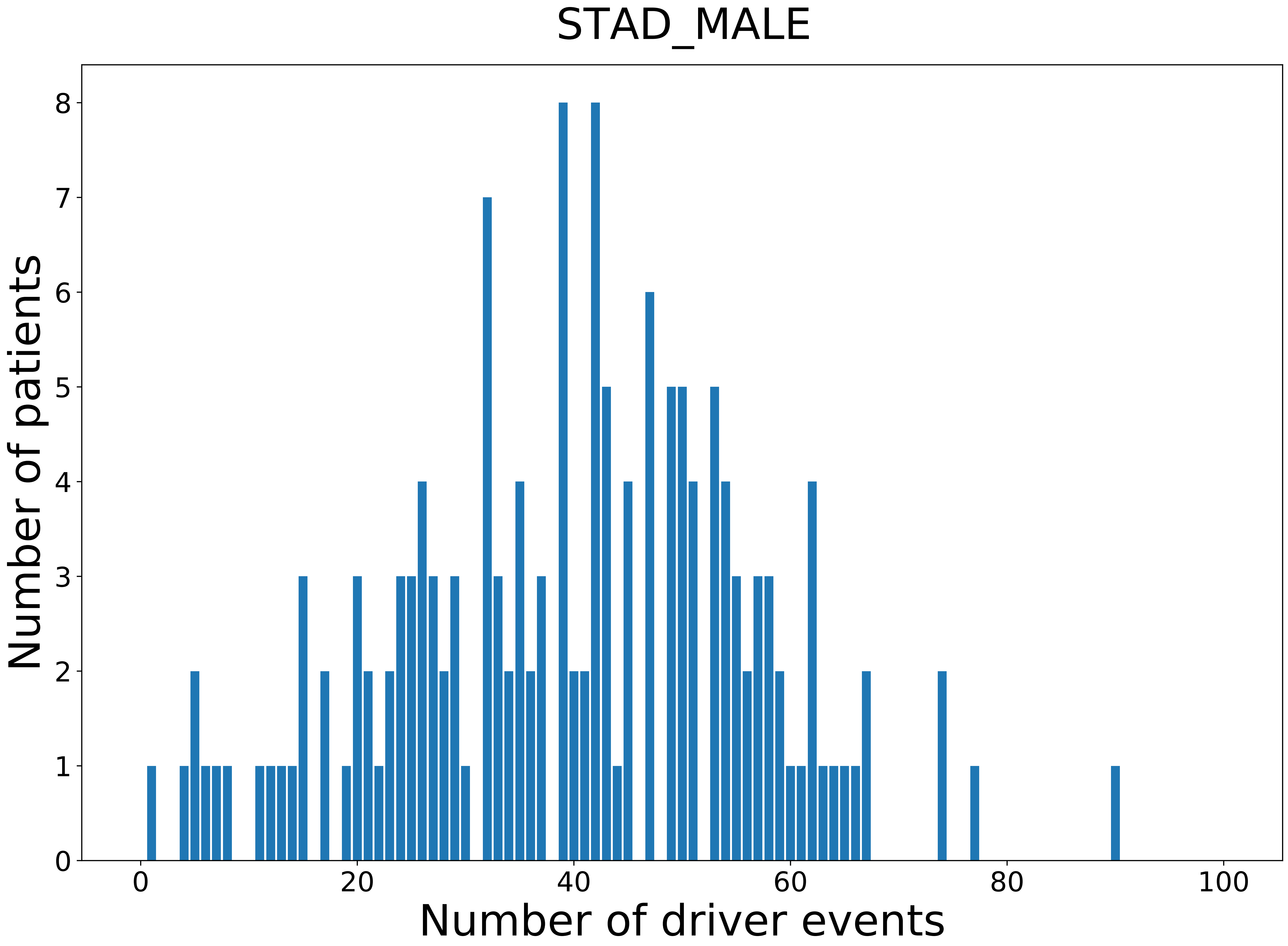

### 2021_8_22_13_42_TGCT.pdf

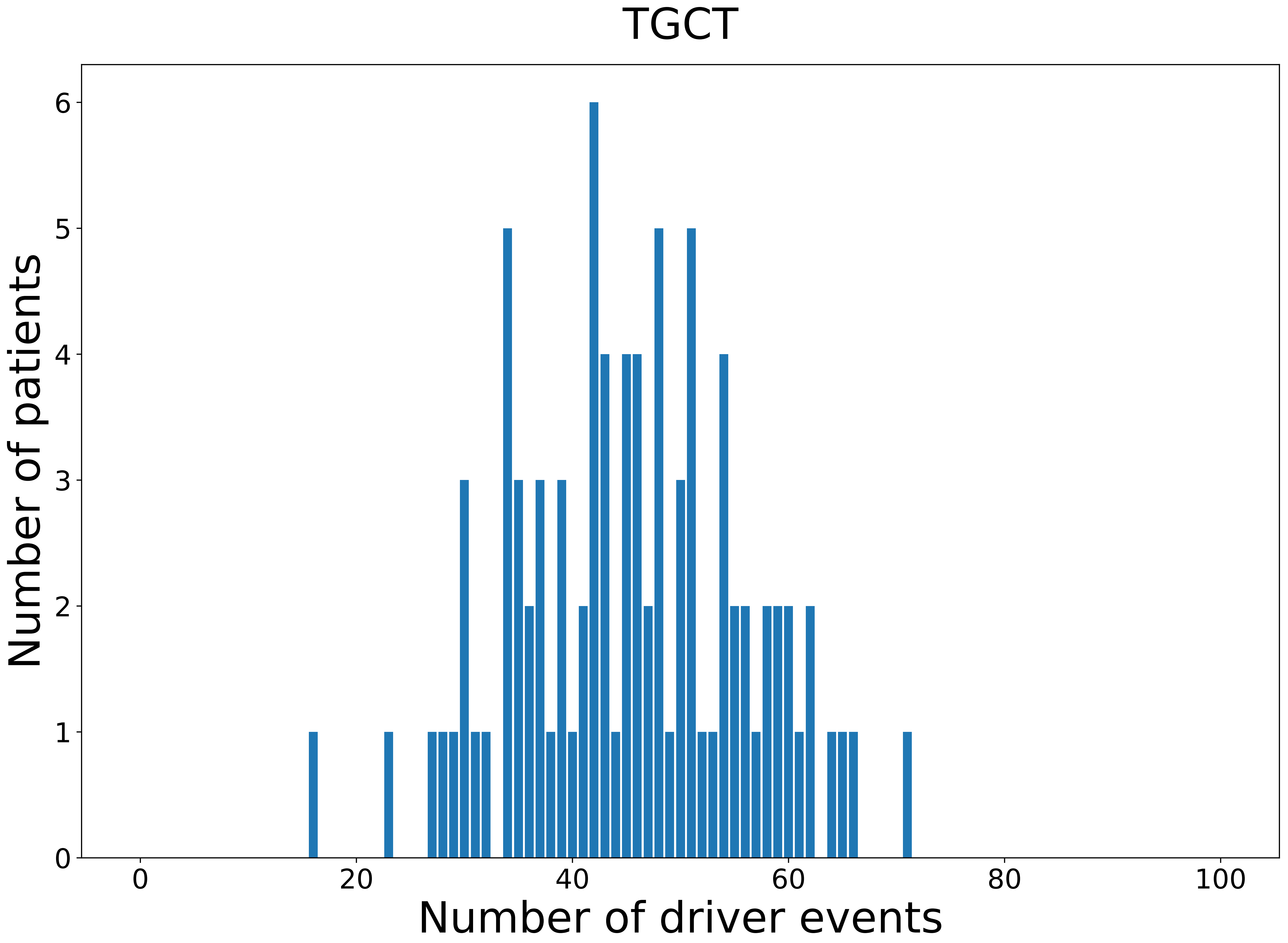

### 2021_8_22_13_42_TGCT_MALE.pdf

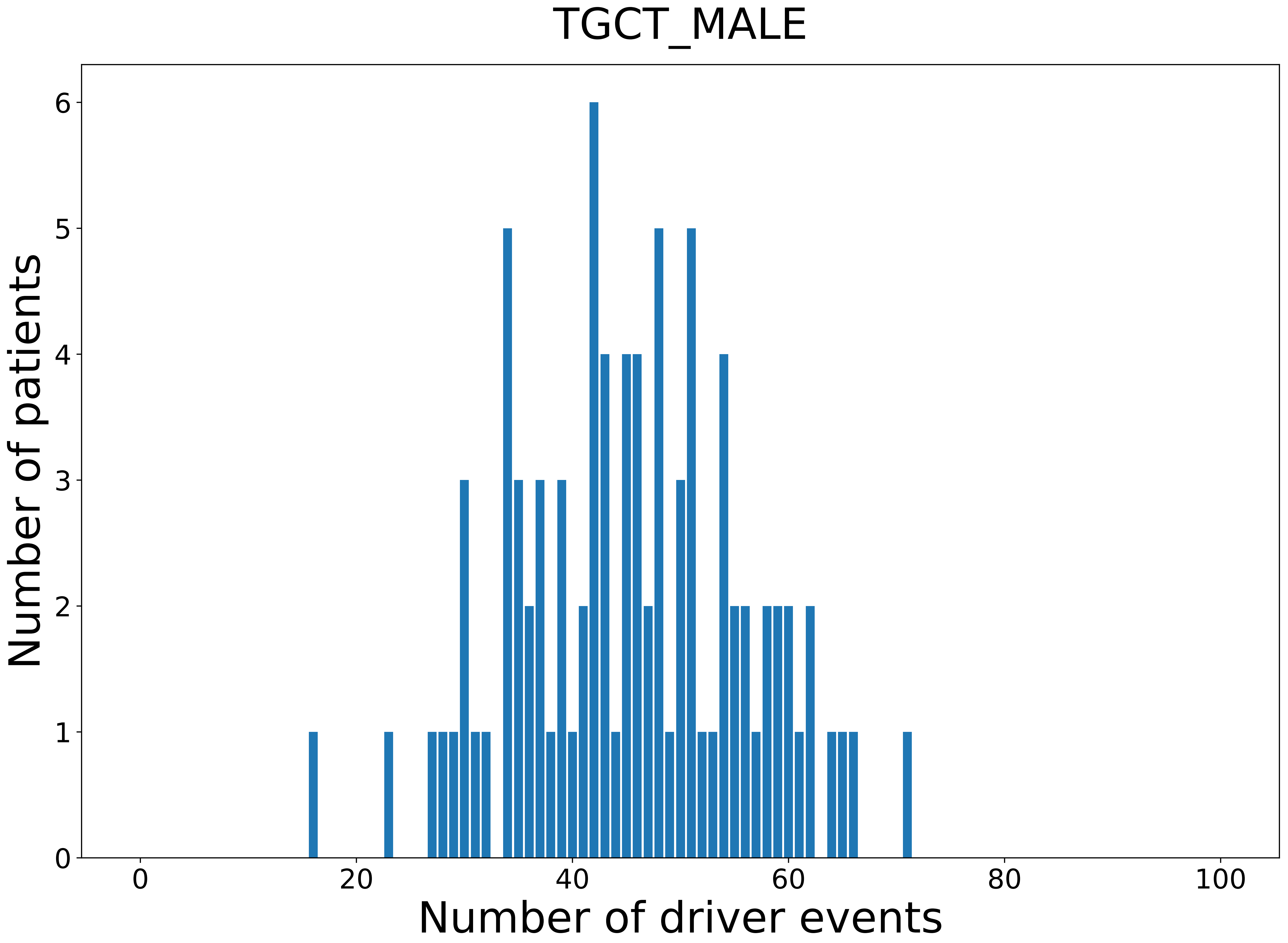

### 2021_8_22_13_42_THCA.pdf

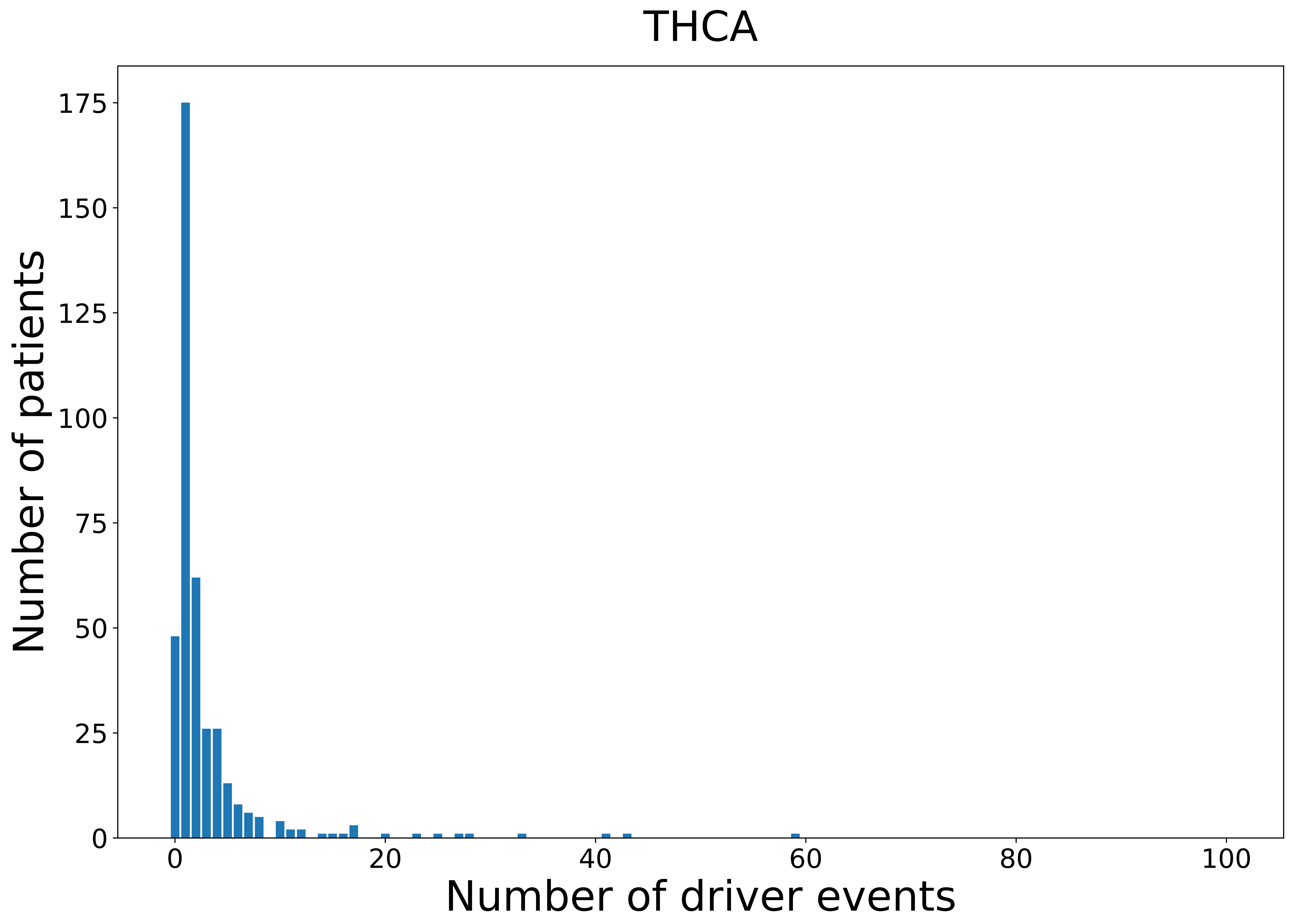

### 2021_8_22_13_42_THCA_FEMALE.pdf

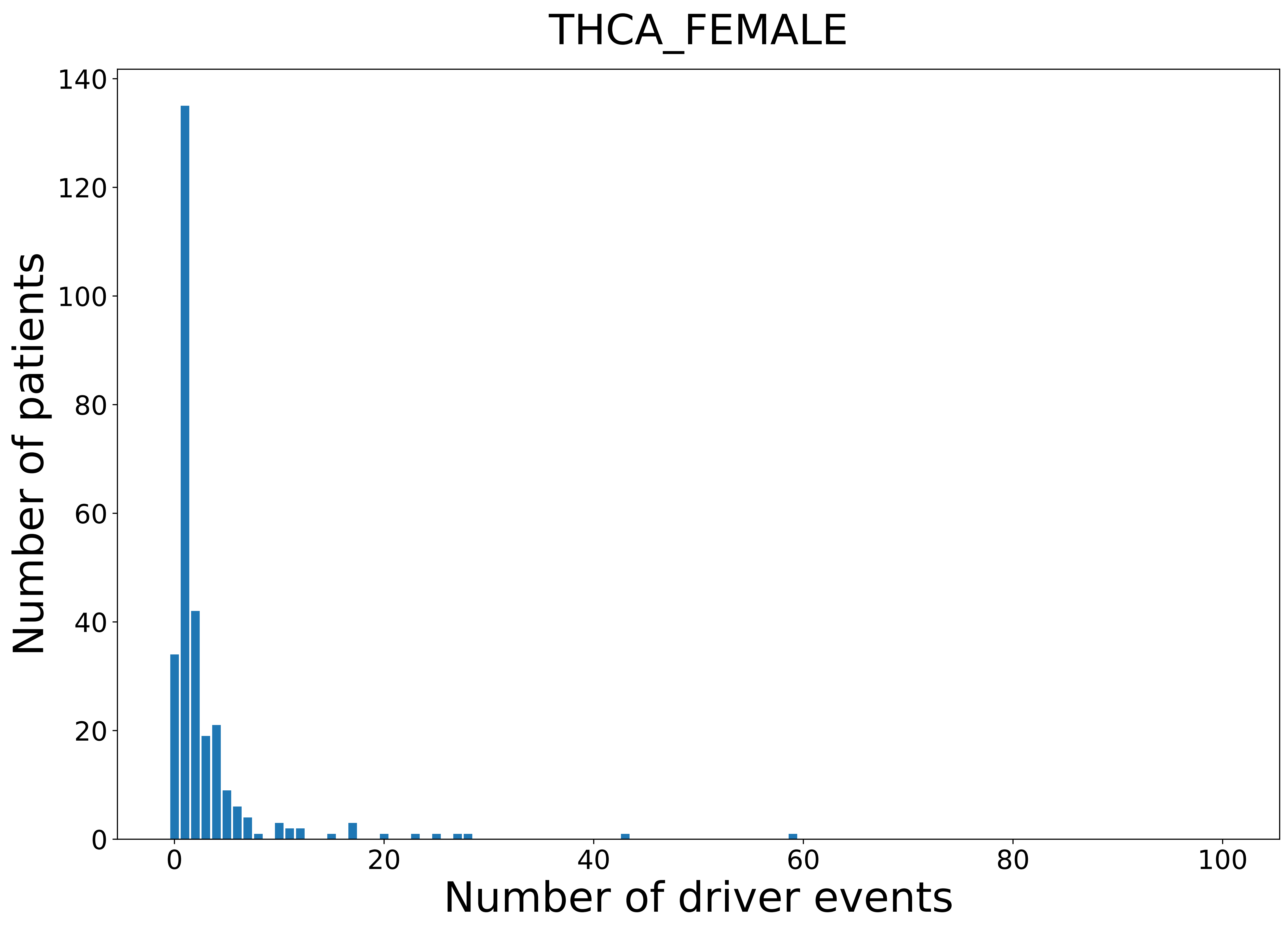

### 2021_8_22_13_42_THYM.pdf

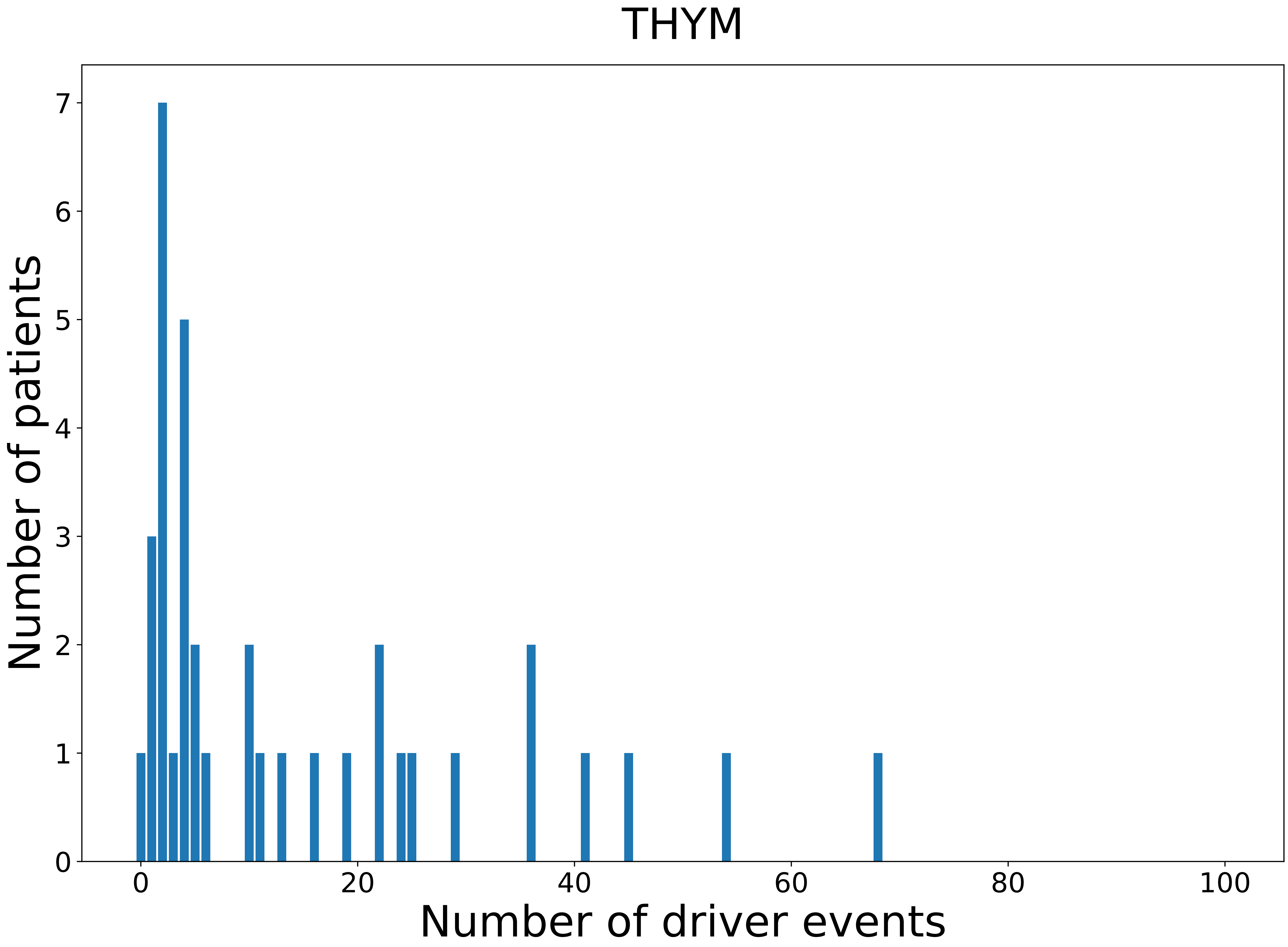

### 2021_8_22_13_42_THYM_FEMALE.pdf

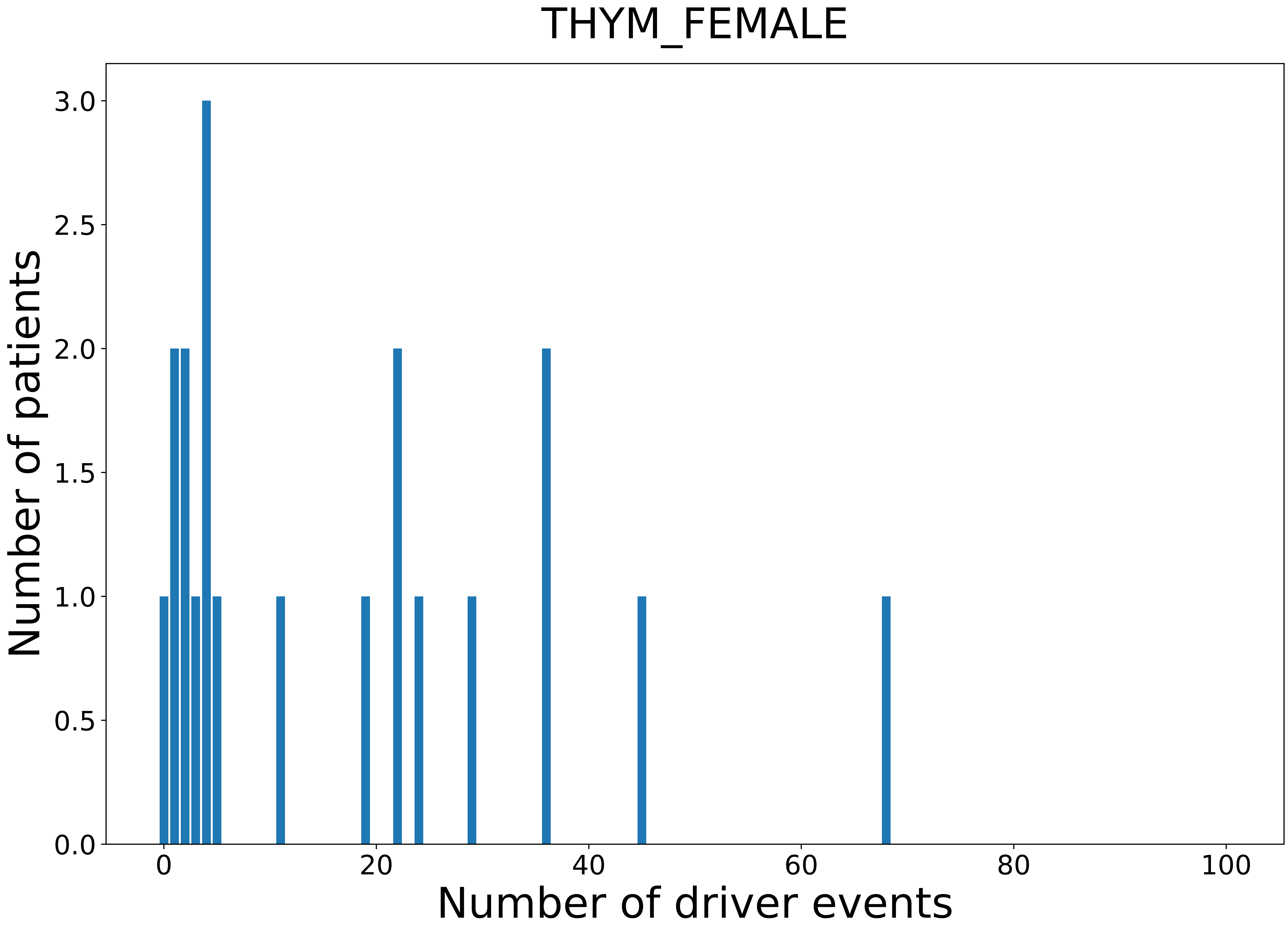

### 2021_8_22_13_42_THYM_MALE.pdf

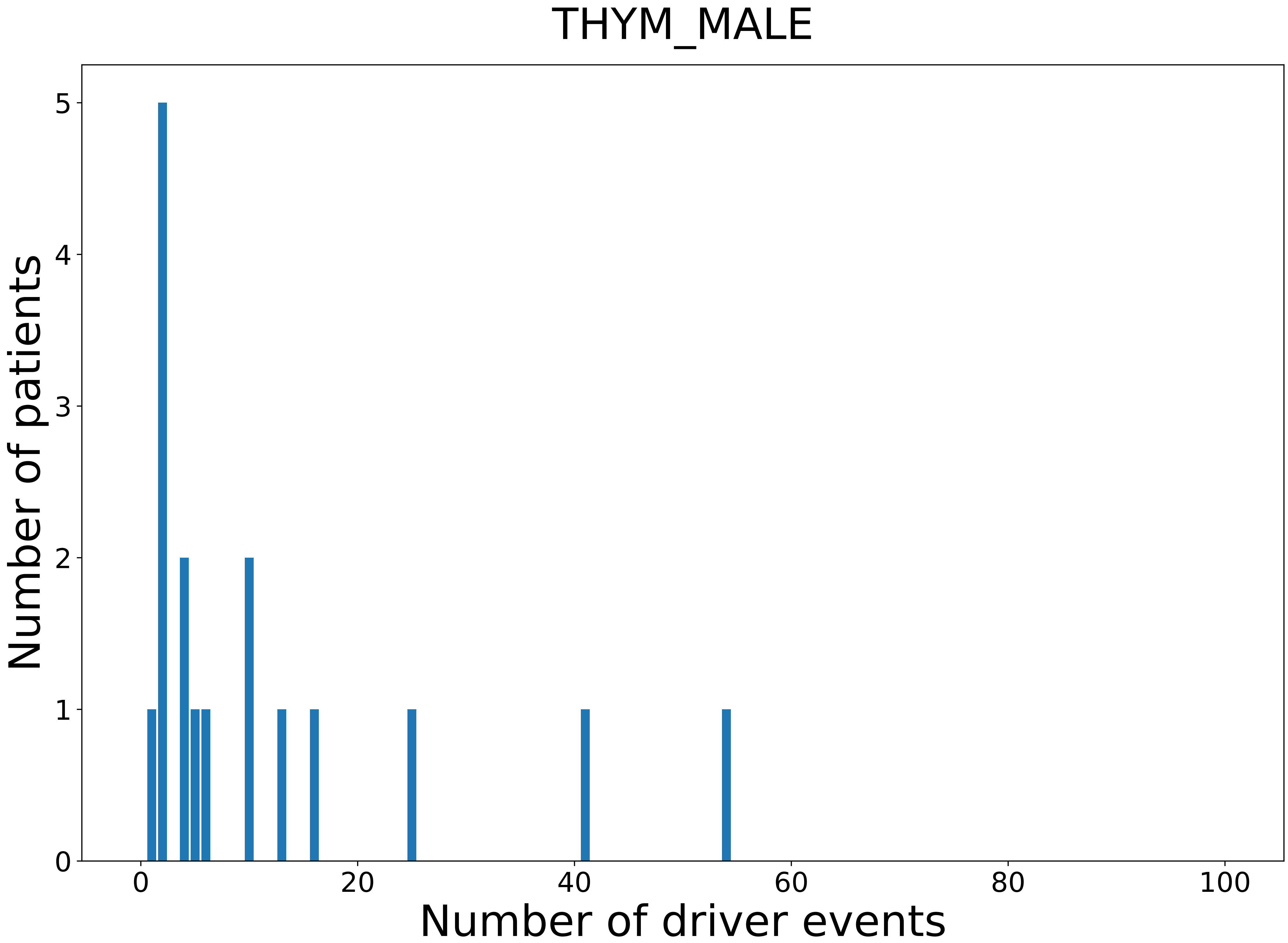

### 2021_8_22_13_42_UCEC.pdf

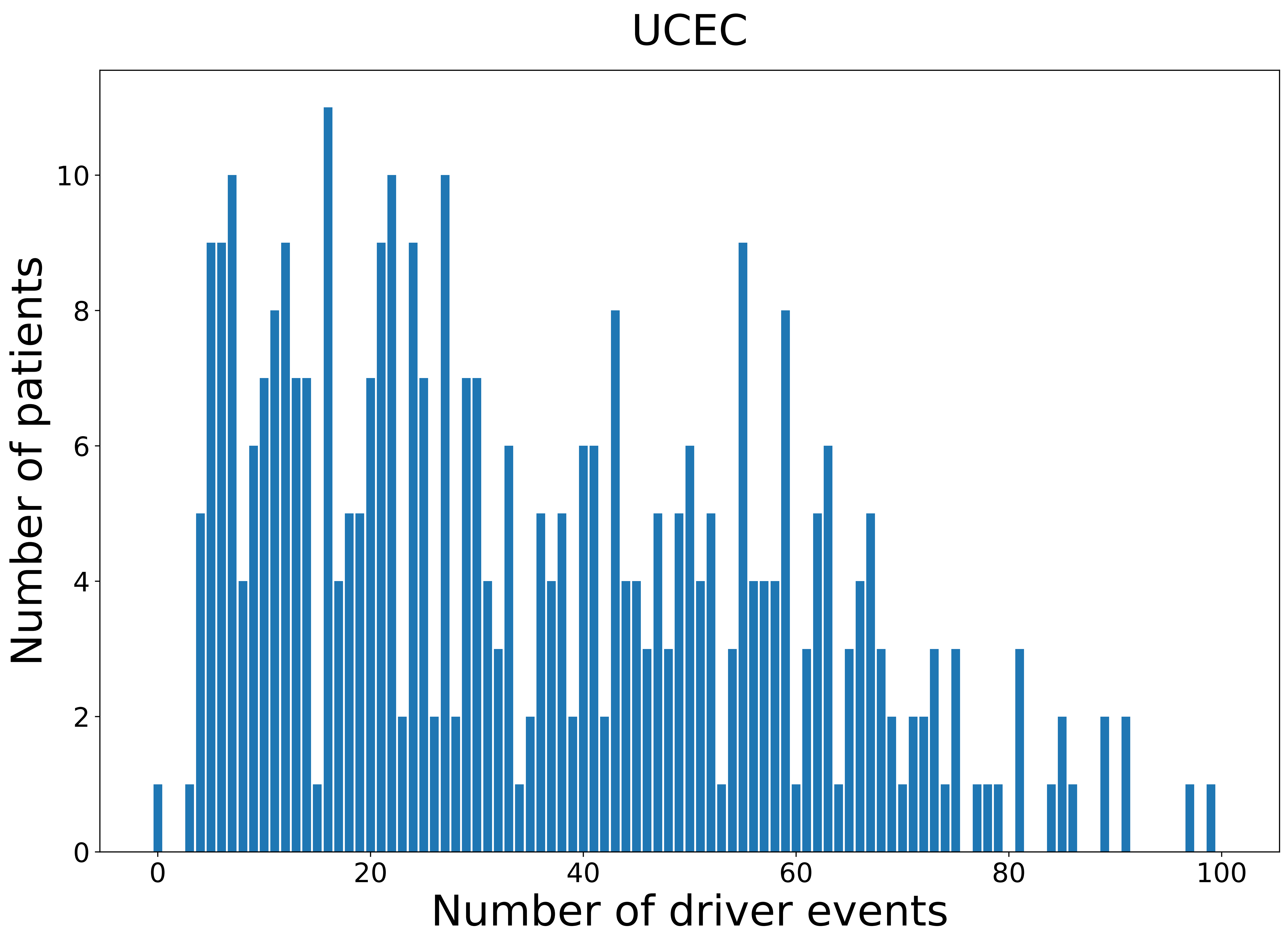

### 2021_8_22_13_42_UCEC_FEMALE.pdf

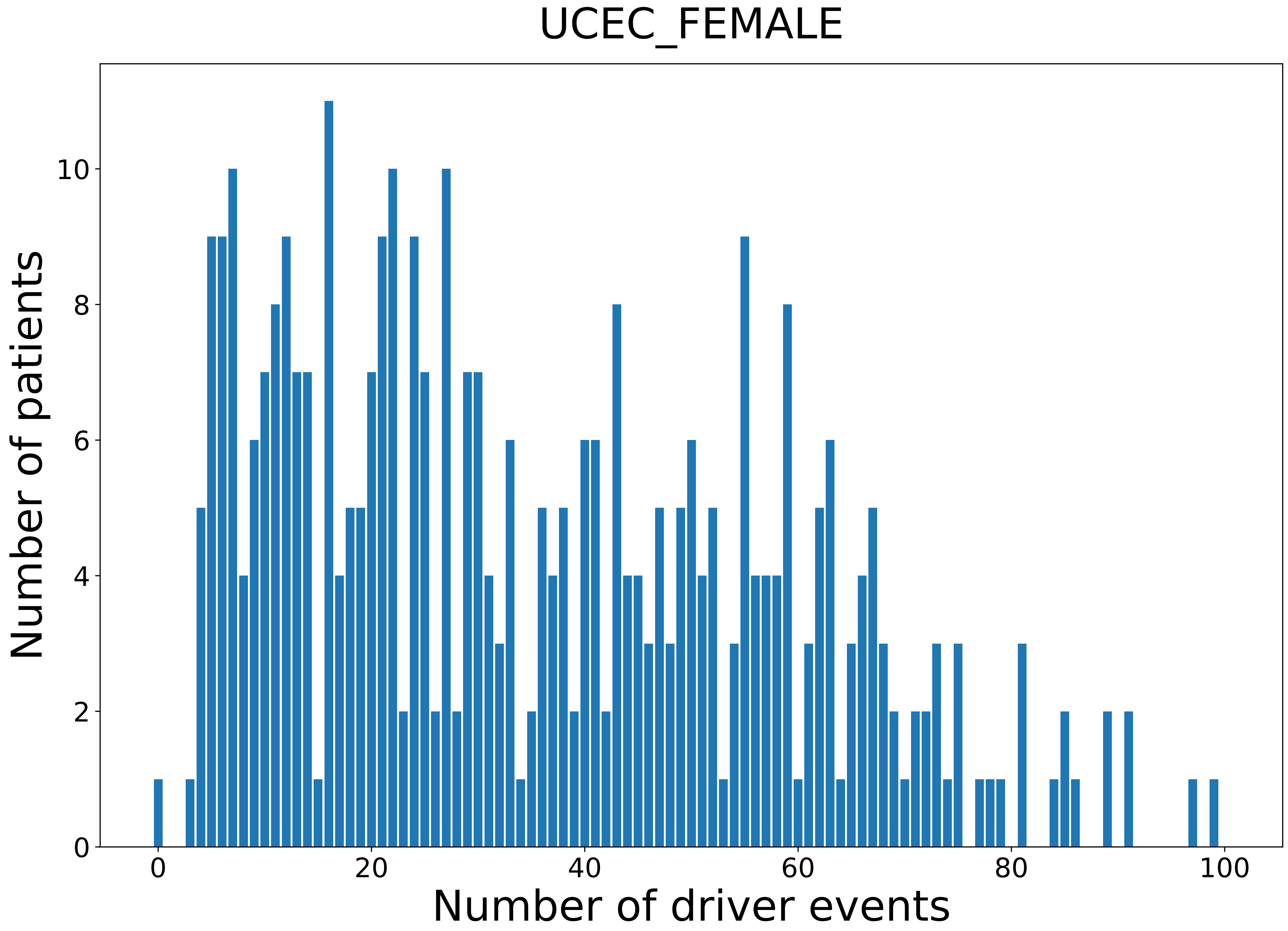

### 2021_8_22_13_42_UCS.pdf

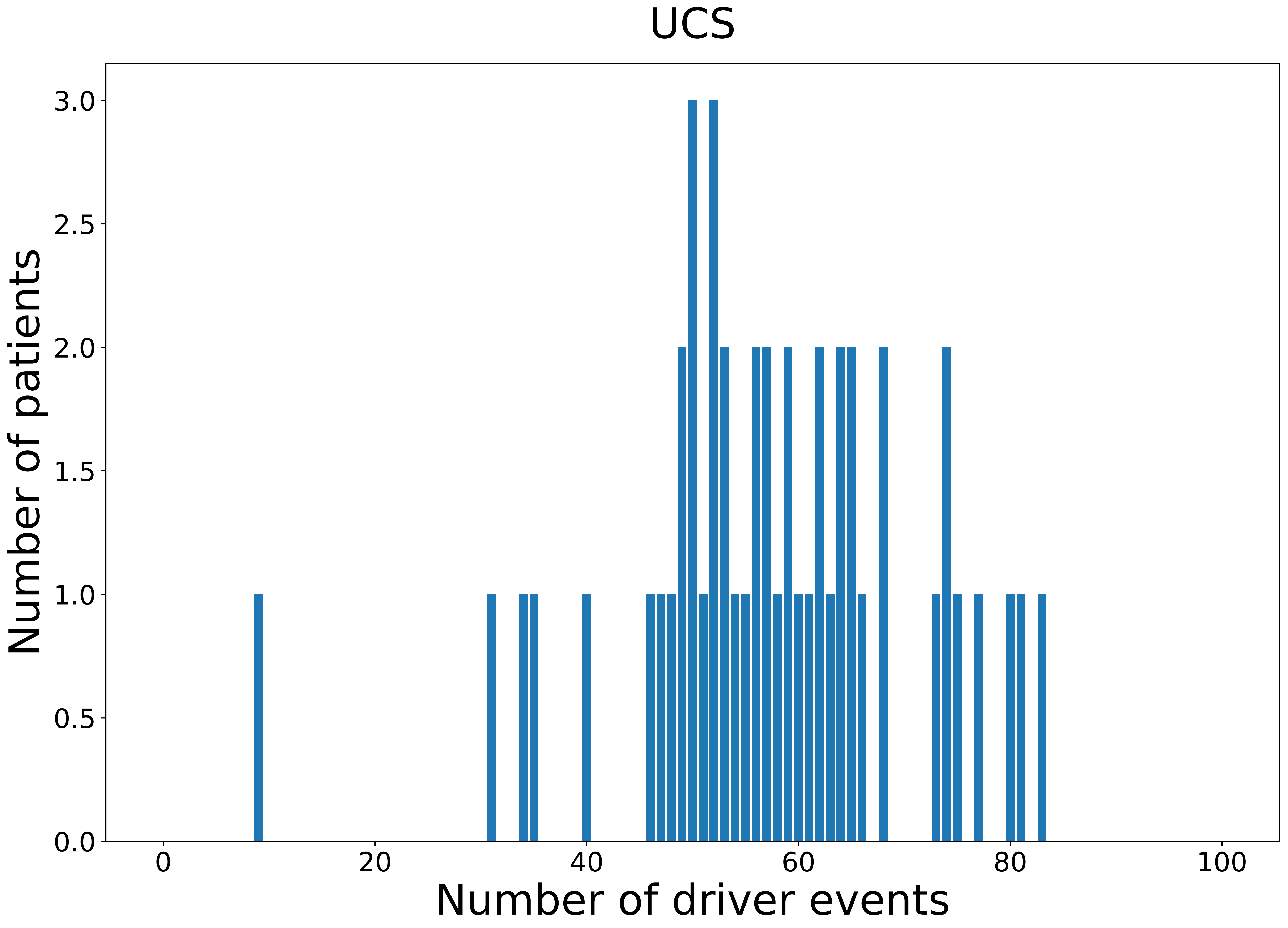

### 2021_8_22_13_42_UCS_FEMALE.pdf

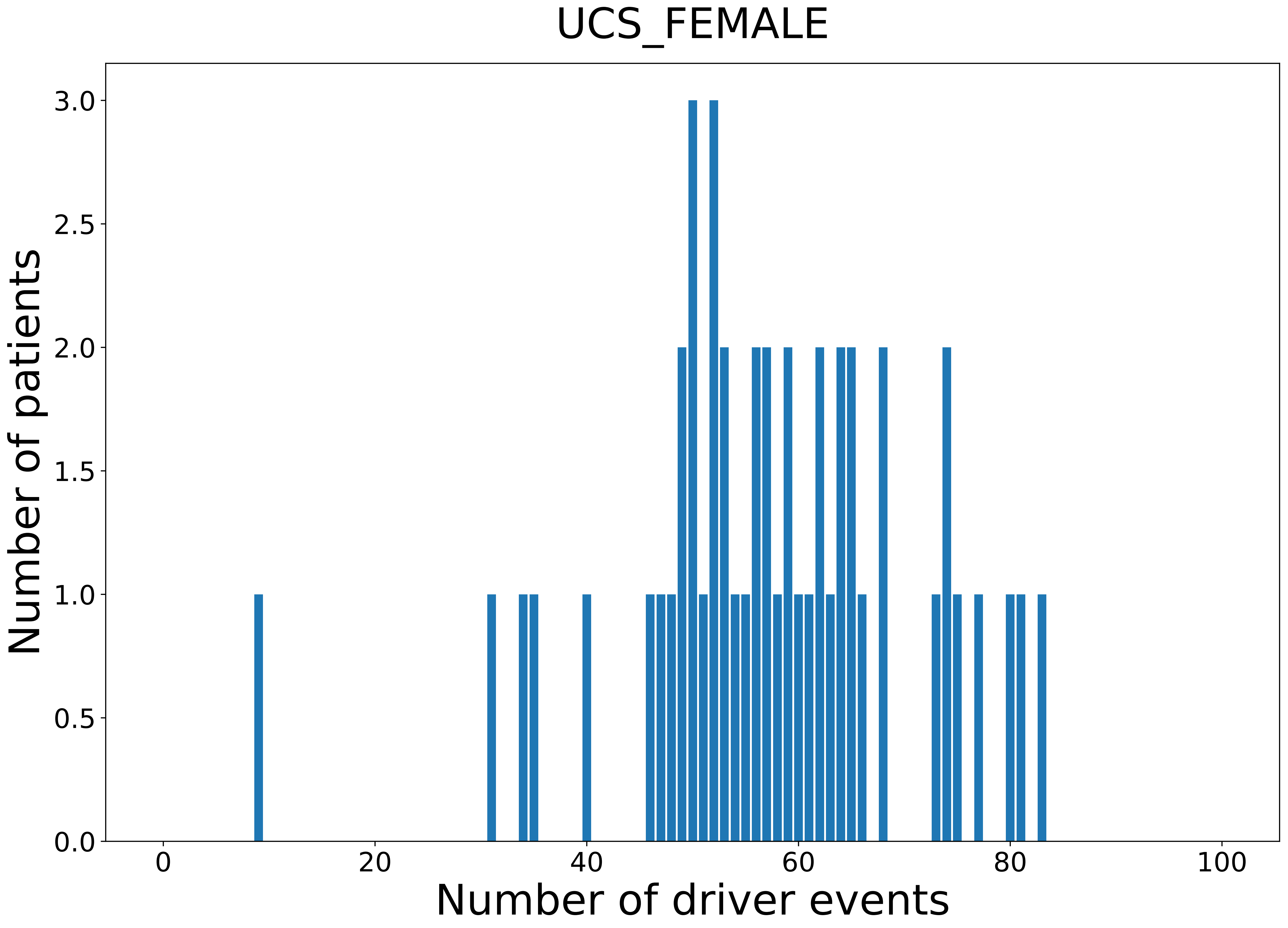

### 2021_8_22_13_42_UVM.pdf

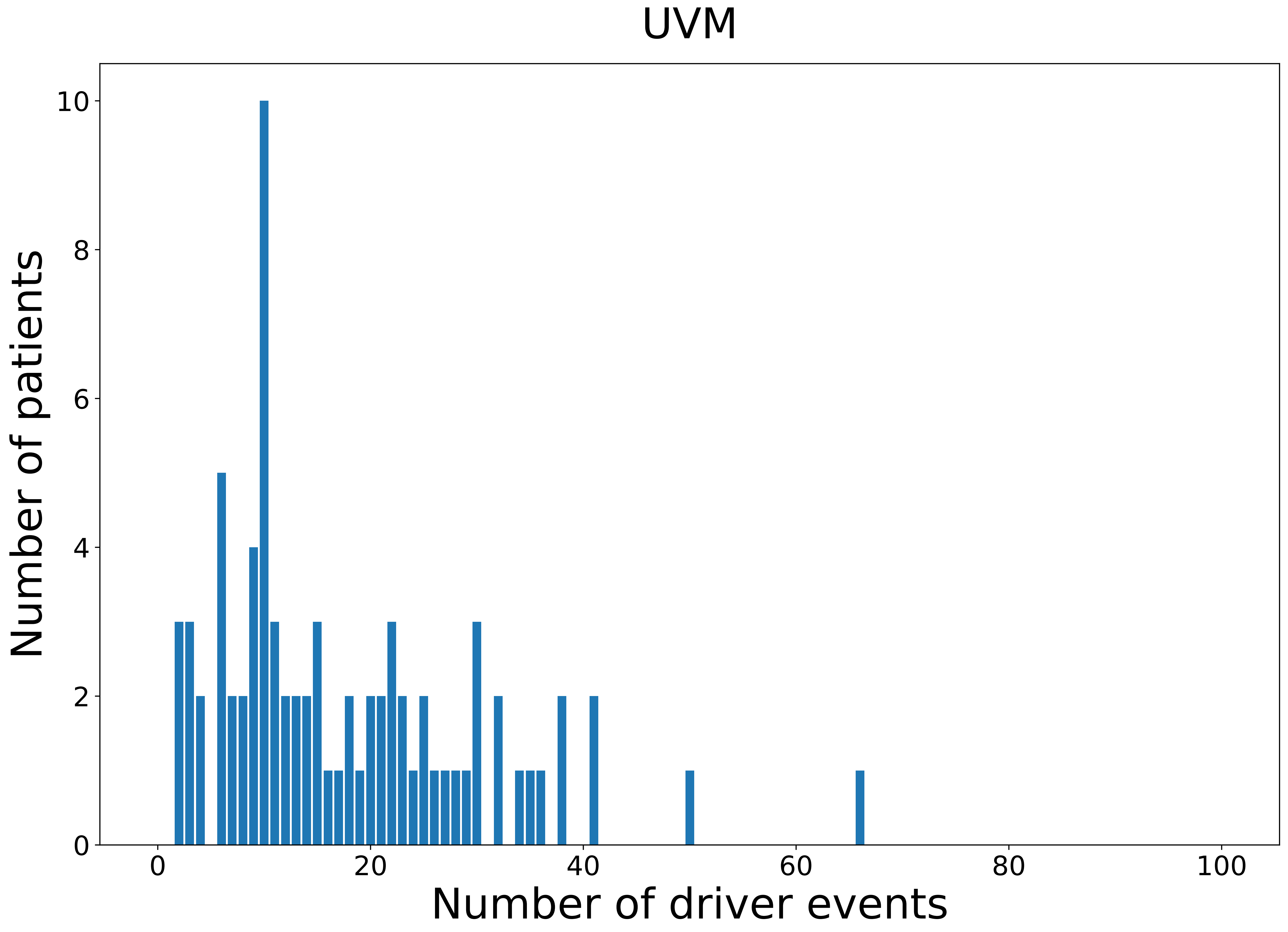

### 2021_8_22_13_42_UVM_FEMALE.pdf

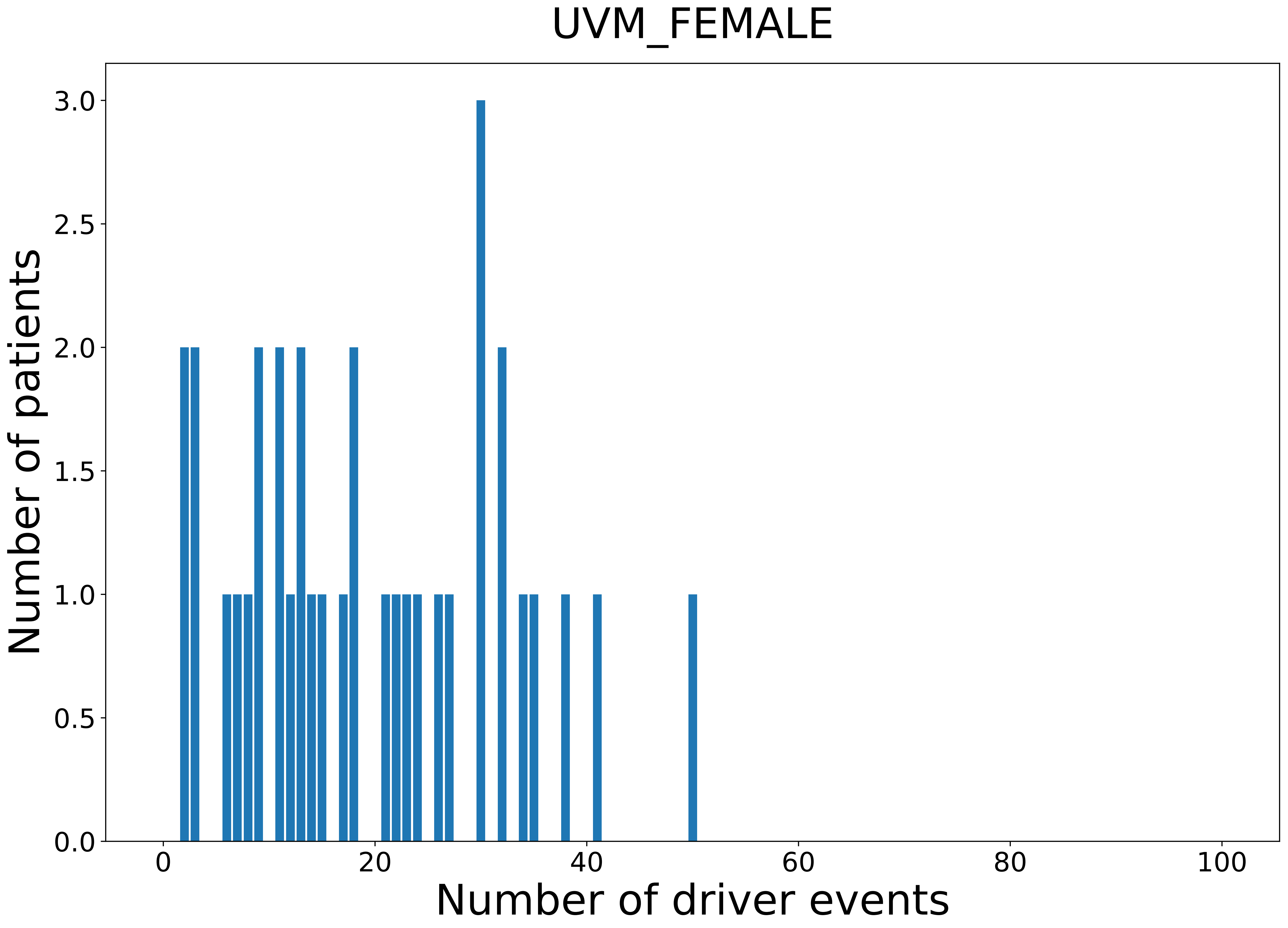

### 2021_8_22_13_42_UVM_MALE.pdf

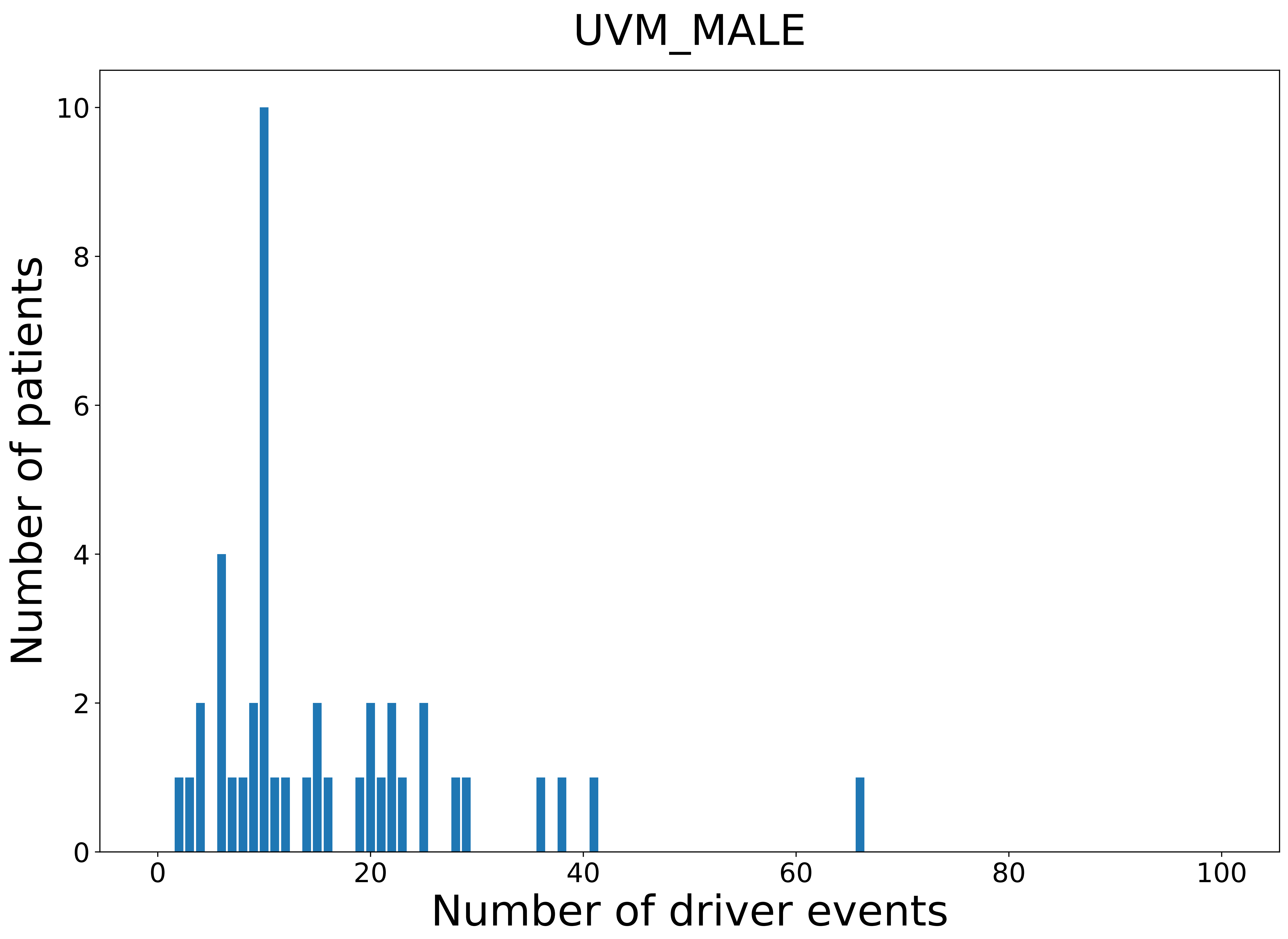

### legeng.png

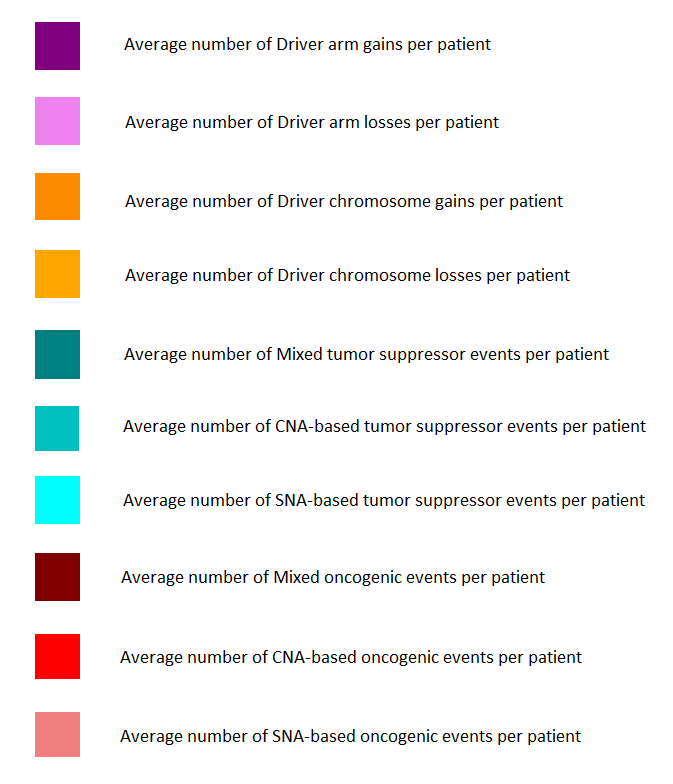
