## Supplementary figures and images for "Comprehensive patient-level classification and quantification of driver events in TCGA PanCanAtlas cohorts"

### 2021_8_22_13_29_ACC.pdf

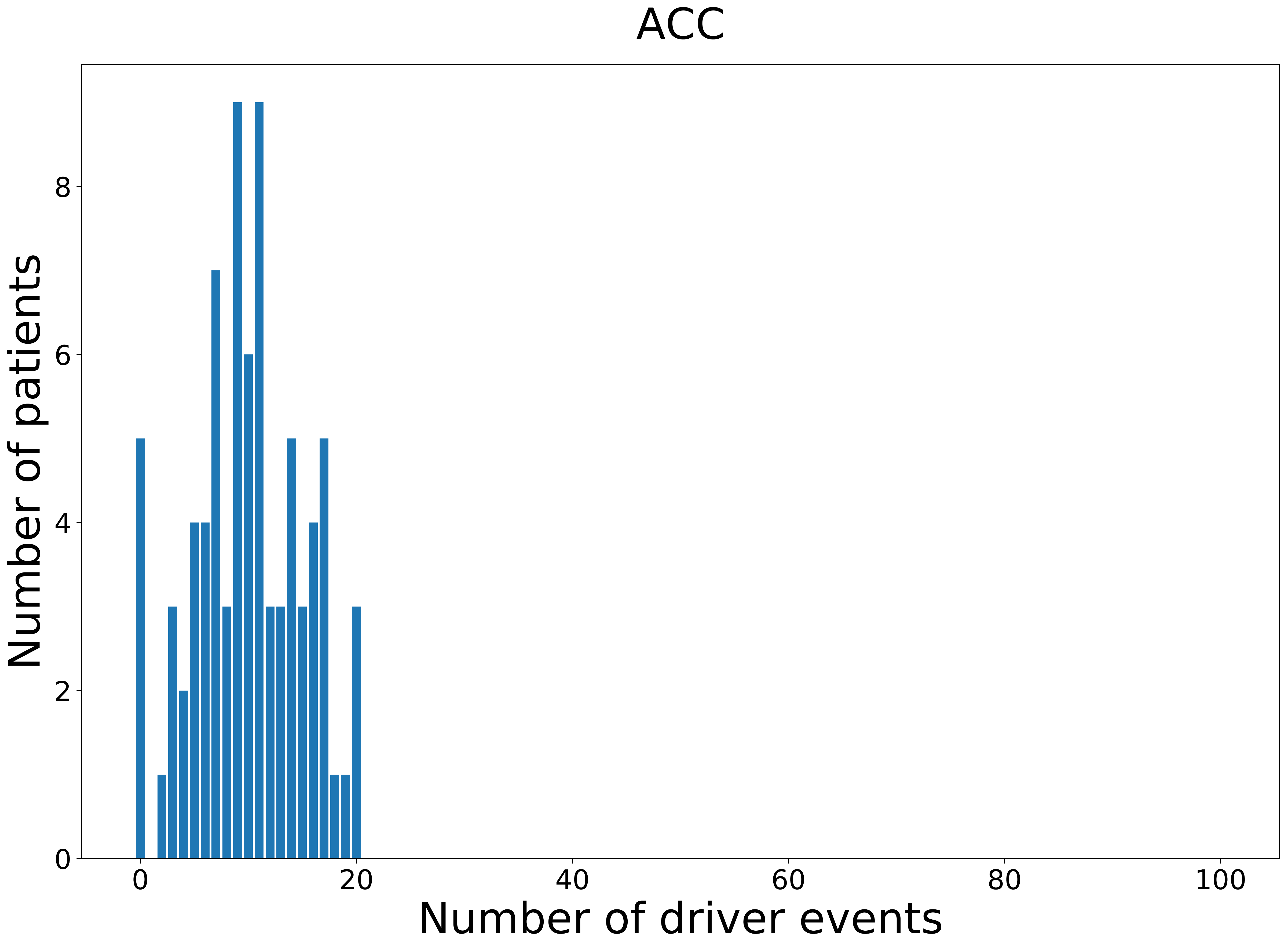

### 2021_8_22_13_29_COAD_FEMALE.pdf

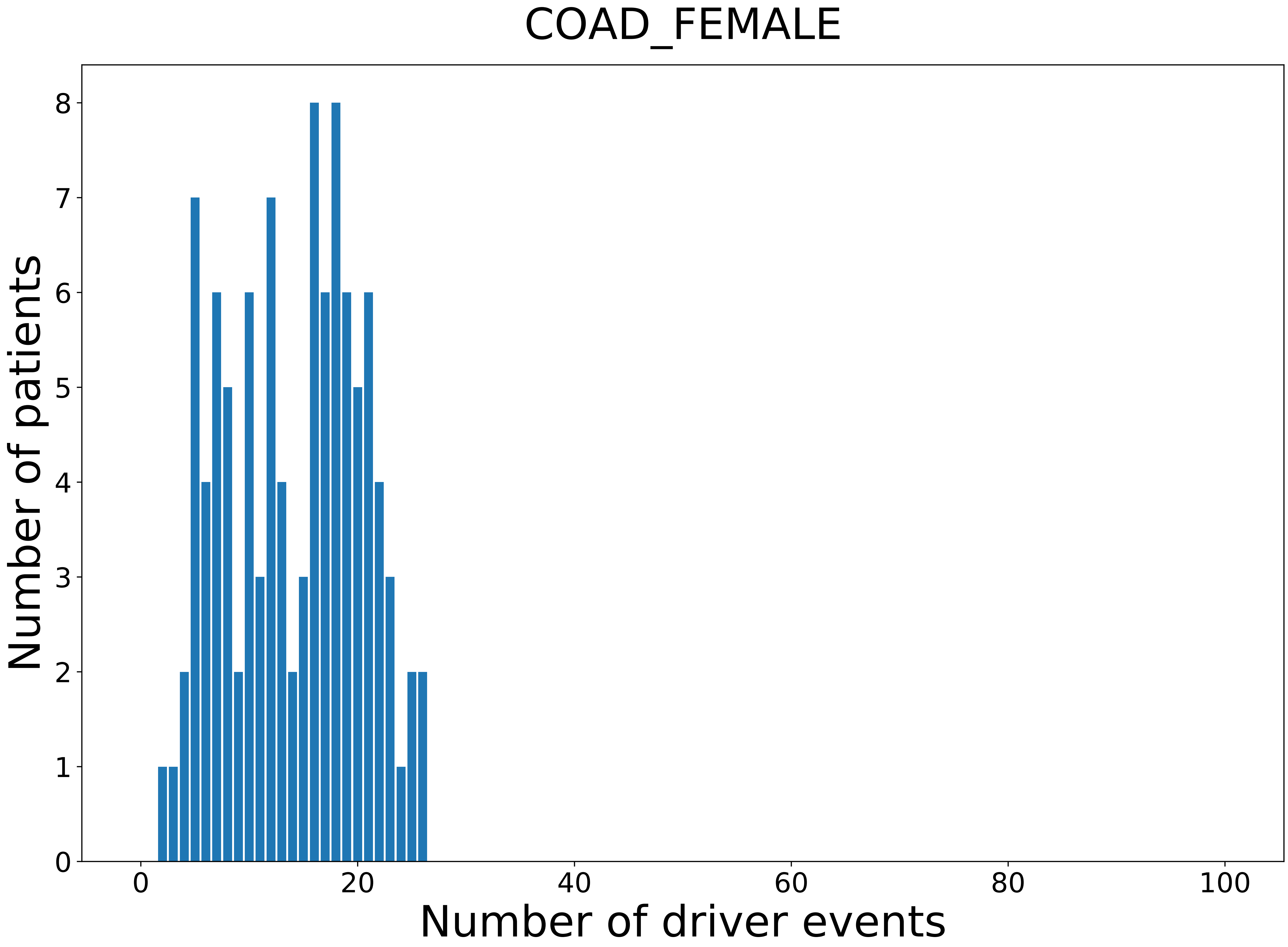

### 2021_8_22_13_29_KIRC_MALE.pdf

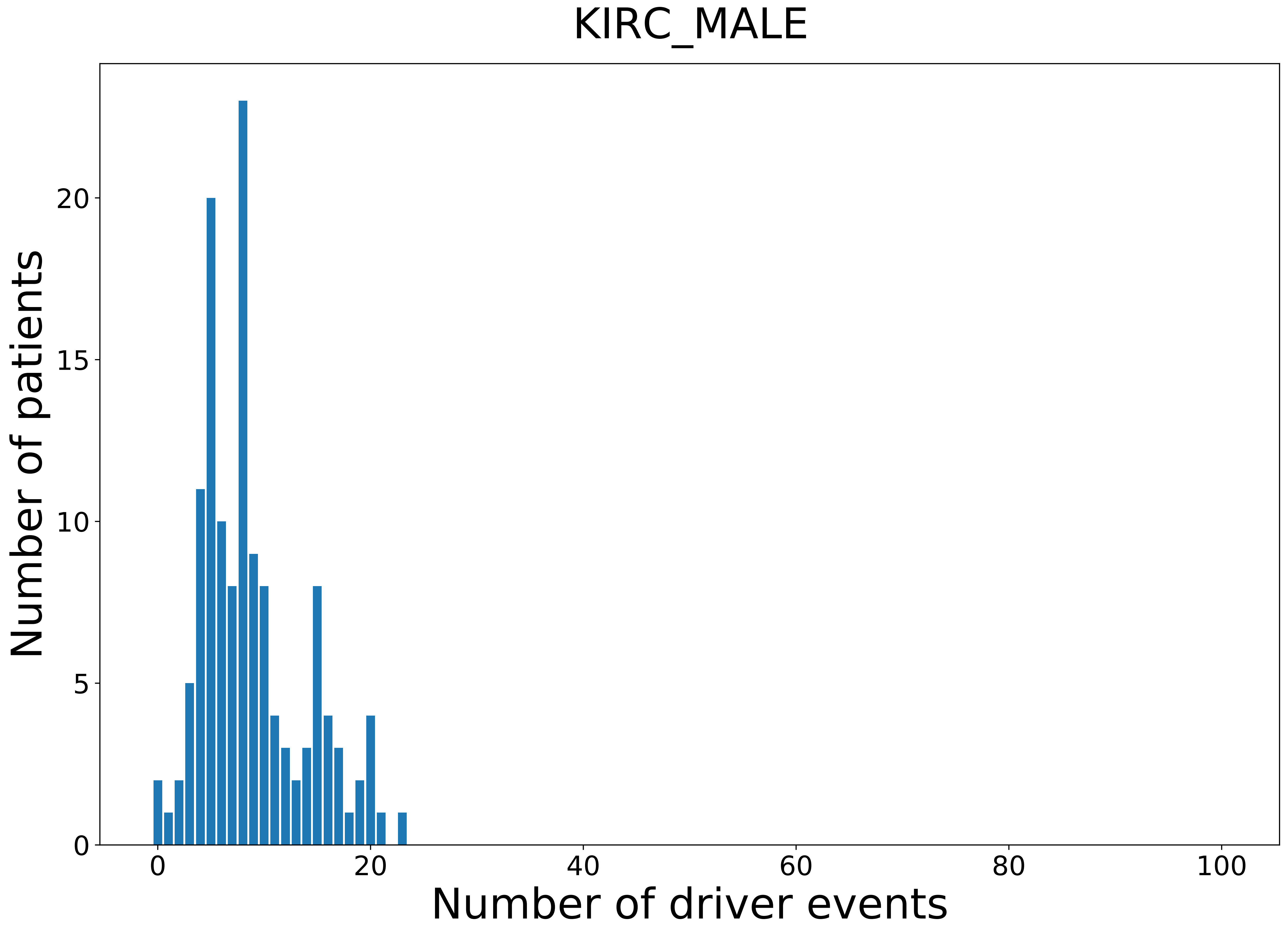

### 2021_8_22_13_29_LGG.pdf

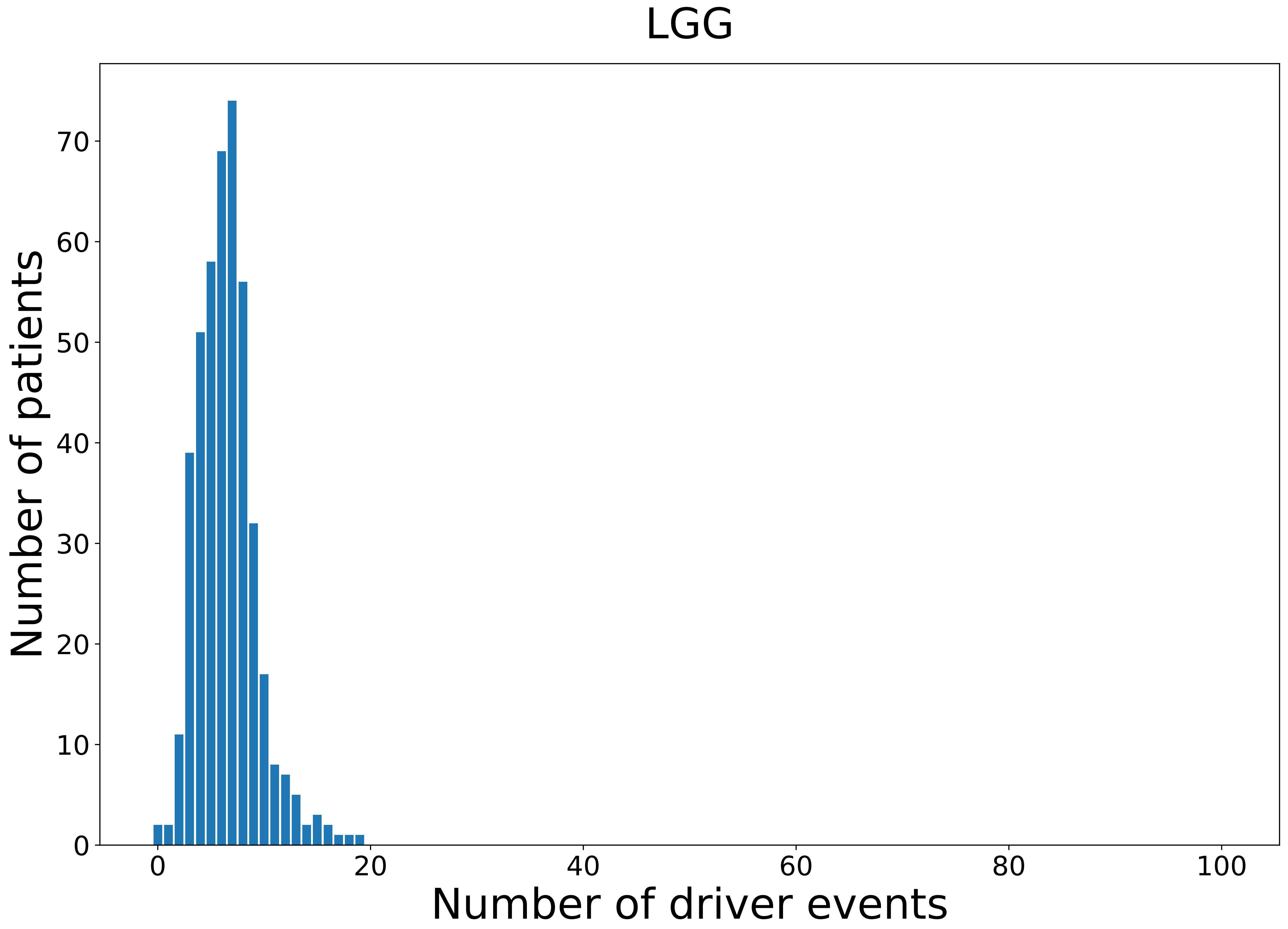

### 2021_8_22_13_29_LUSC.pdf

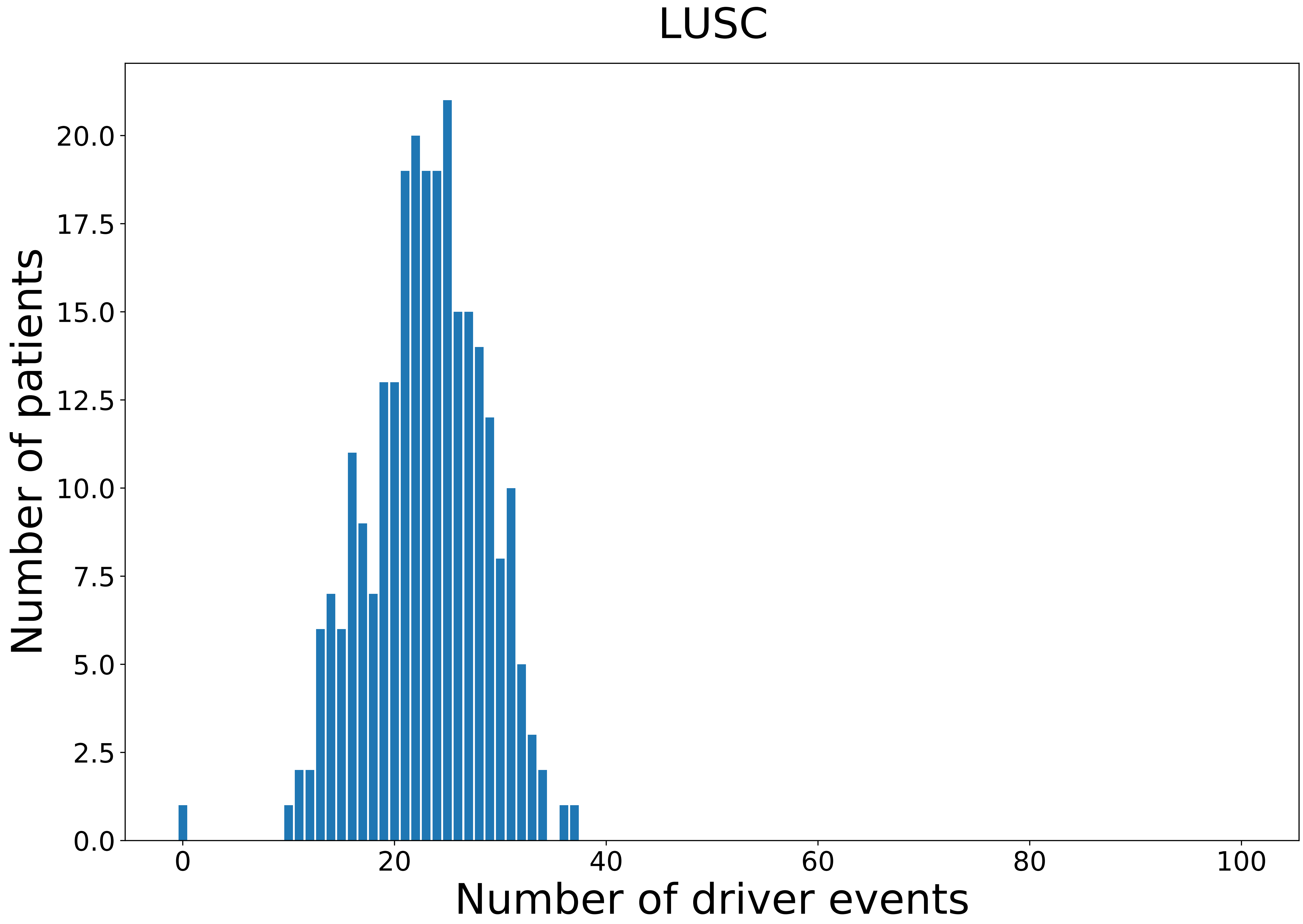

### 2021_8_22_13_29_PCPG.pdf

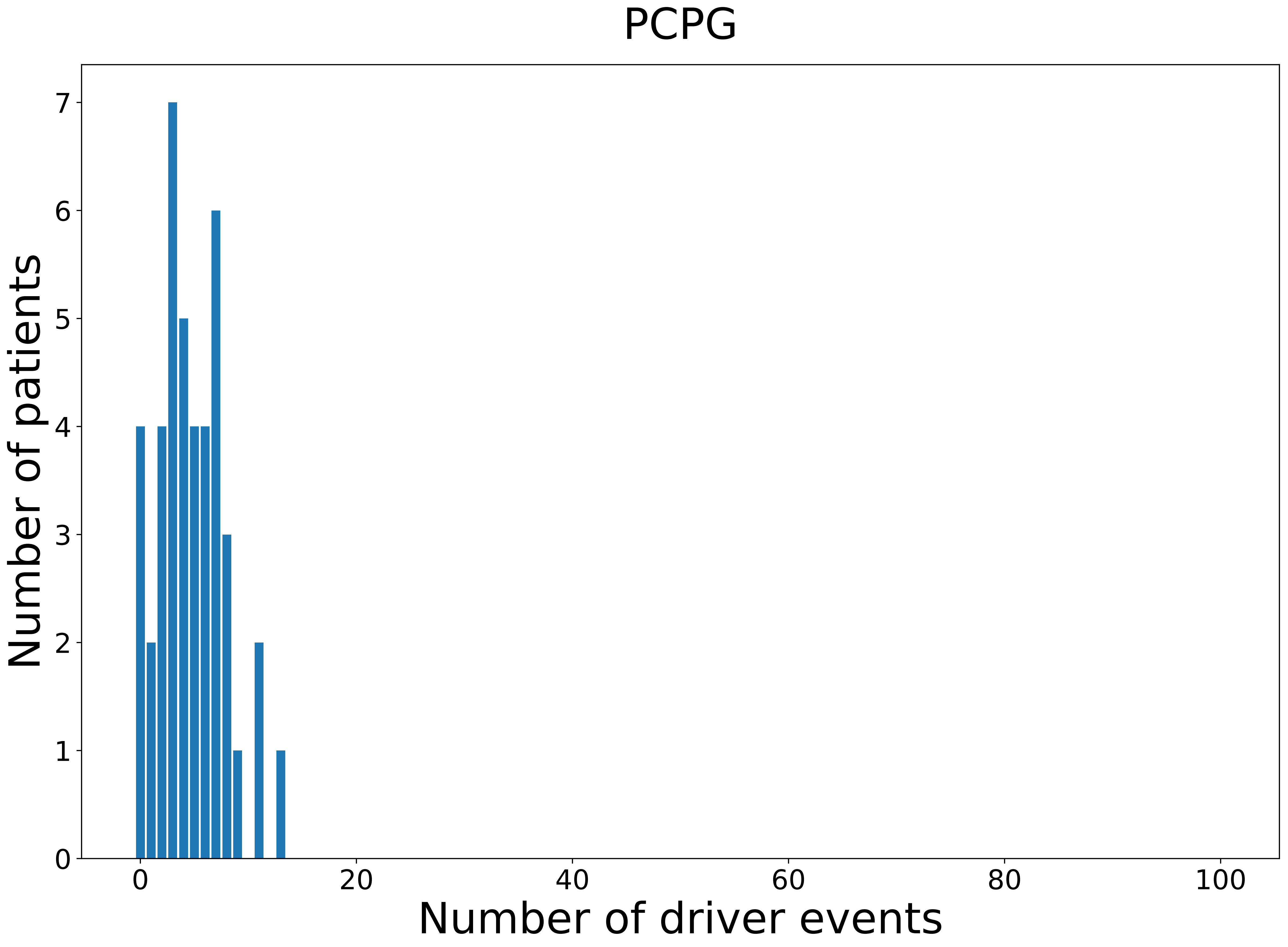

### 2021_8_22_13_29_SARC_FEMALE.pdf

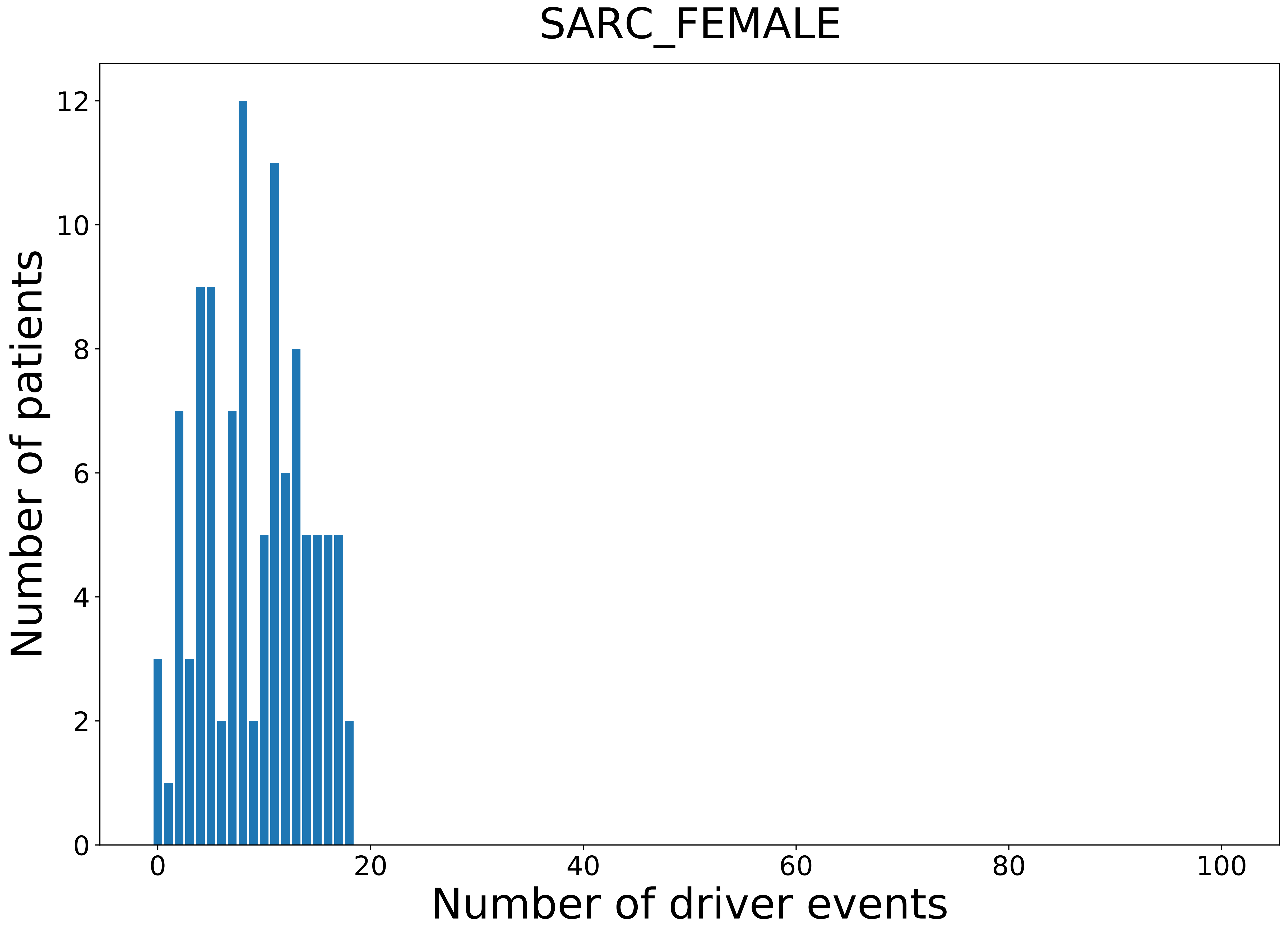

### 2021_8_22_13_29_STAD_FEMALE.pdf

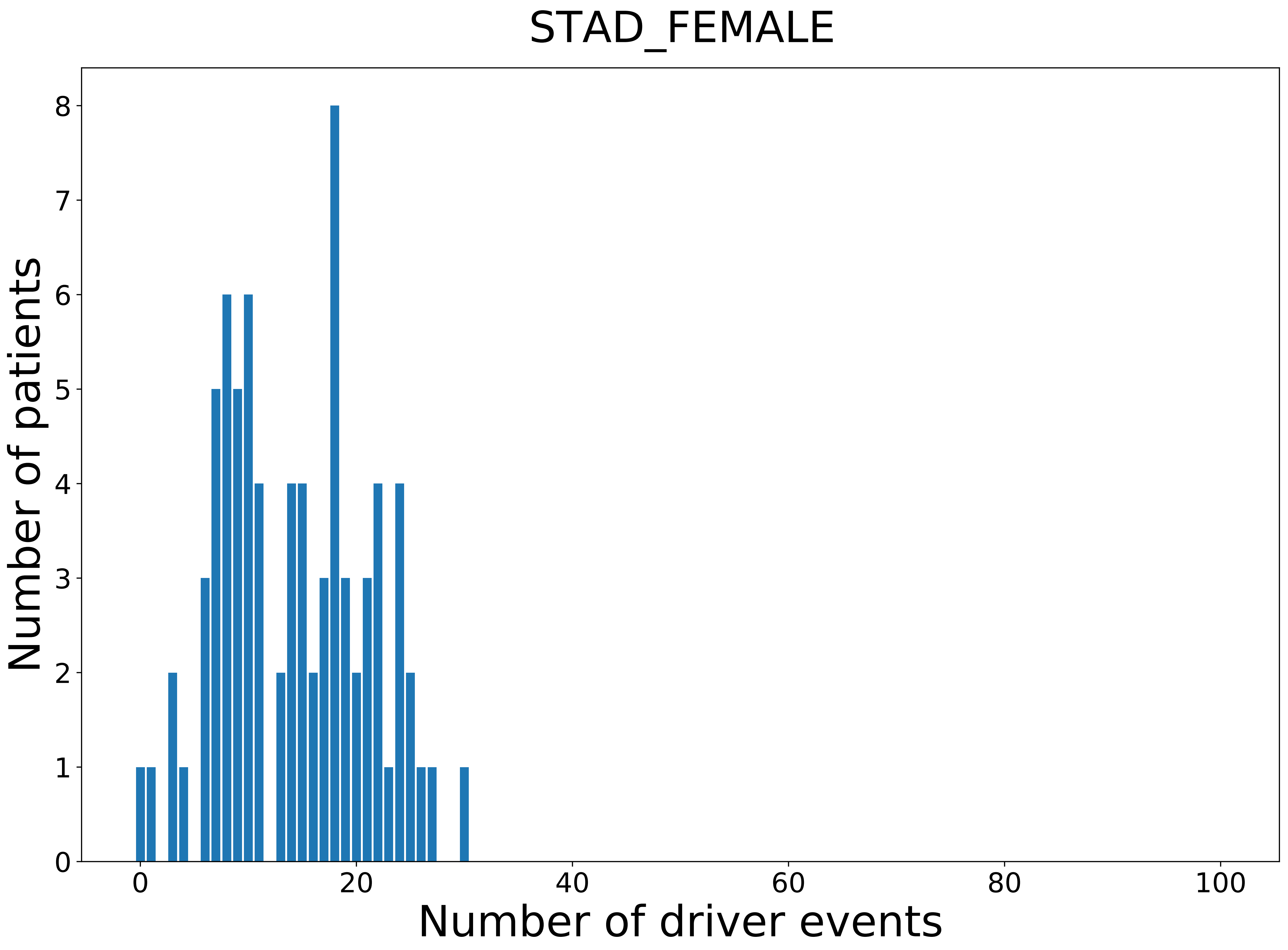

### 2021_8_22_13_29_TGCT.pdf

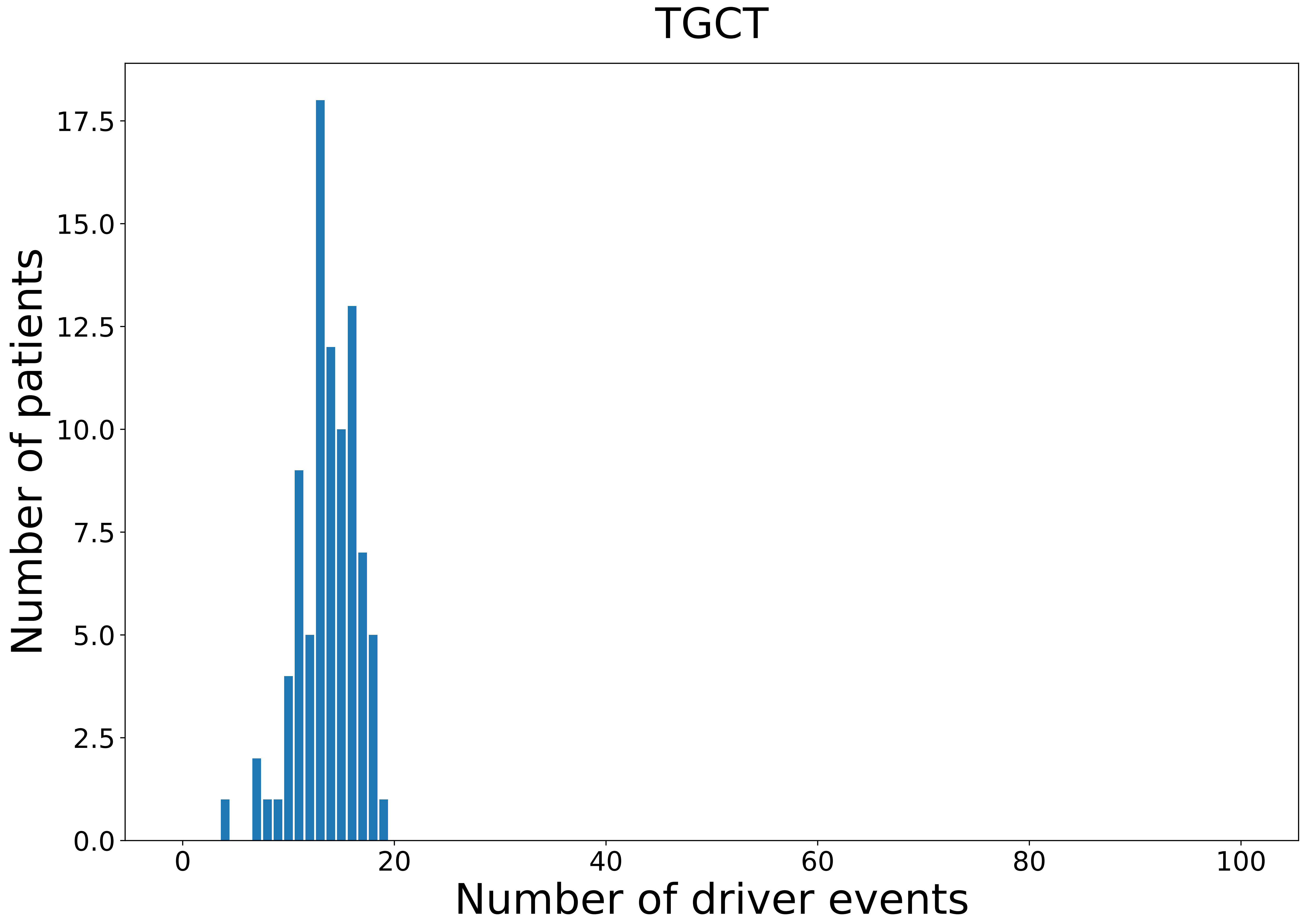

### 2021_8_22_13_29_UVM_FEMALE.pdf

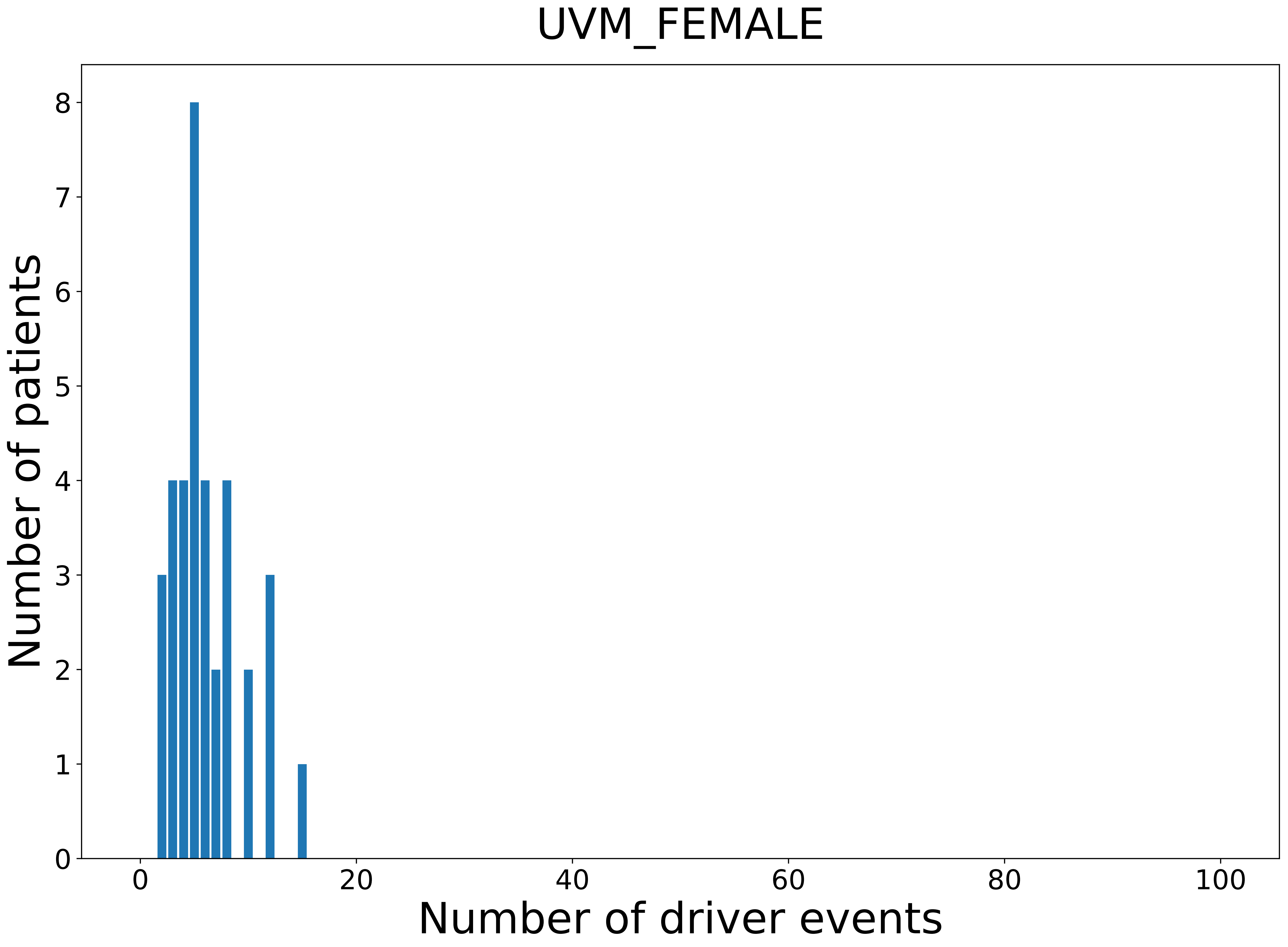

### 2021_8_22_13_29_UVM_MALE.pdf

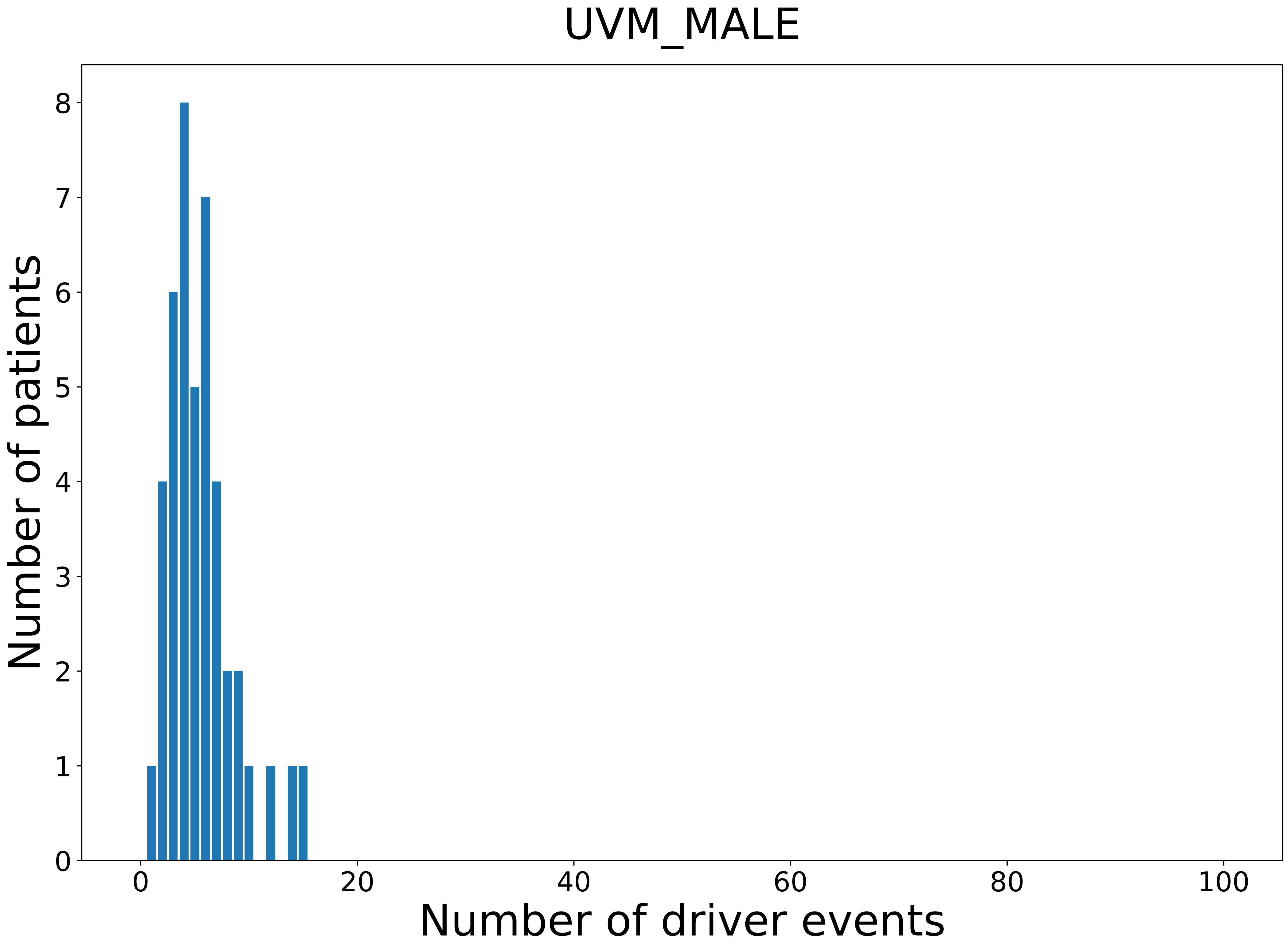
