## Supplementary material for "Comprehensive patient-level classification and quantification of driver events in TCGA PanCanAtlas cohorts": S5 Files: GECNAV description.pdf

GECNAV (Gene Expression-based CNA Validator) is a Python 3.7 software package that validates CNAs using gene and miRNA expression data from the TCGA PanCanAtlas (https://gdc.cancer.gov/about-data/publications/PanCan-CellOfOrigin). Low quality samples and metastatic samples are filtered out. CNA validation is based on comparing the CNA status of a given gene in a given patient to expression of this gene in this patient relative to the median expression of this gene across all patients. miRNA genes are also included into the verification process. The pipeline can be executed fully automatically in less than one hour on a modern PC (Linux, Windows or MacOS).

This package has been developed by

Aleksey V. Belikov, Dr.rer.nat.
https://github.com/belikov-av
Laboratory of Innovative Medicine, School of Biological and Medical Physics, Moscow Institute of Physics and Technology
Roles: concept, pipeline, supervision

and

Alexey D. Vyatkin
https://github.com/VyatkinAlexey
Skoltech
Roles: programming

A detailed description of pipeline steps can be found in the file GECNAV pipeline.pdf

Instructions for executing the code can be found in the file Instructions.txt
