## Supplementary material for "Comprehensive patient-level classification and quantification of driver events in TCGA PanCanAtlas cohorts": S5 Files: GECNAV instructions.txt

This script requires Python 3.7 and internet connection

Always open terminal from the folder GECNAV

First, run the following command to install required libraries:
`pip3 install --user -r requirements.txt`

If you can execute Python3 with command `python`, you may use `python`
command instead of `python3` in all code snippets below.

To run the whole pipeline: `python3 run_all.py`

To run a particular step: `python3 run_one_step.py N`,
where N is the step number

In order to get brief description for each step, one may run the command:
`python3 step_{step_number}.py --help`.

All downloaded data, intermediate and final results are located
by default in the folder "data".
