## Supplementary material for "Comprehensive patient-level classification and quantification of driver events in TCGA PanCanAtlas cohorts": S5 Files: GECNAV pipeline.pdf

1) Go to https://gdc.cancer.gov/about-data/publications/PanCan-CellOfOrigin
2) Download the file Analyte level annotations - merged_sample_quality_annotations.tsv
3) Using information in the column **aliquot_barcode**, delete all samples named TCGA-XX-XXXX-YYX-XXX-XXXX-XX, where YY – any number, except 01, 03 and 09 (see https://docs.gdc.cancer.gov/Encyclopedia/pages/TCGA_Barcode/ and table https://gdc.cancer.gov/resources-tcga-users/tcga-code-tables/sample-type-codes), and also, using information in the column **Do_not_use**, delete all samples with False value, and save the resulting file as **merged_sample_quality_annotations_do_not_use.tsv**.
4) Download the file gzipped ISAR-corrected GISTIC2.0 all_thresholded.by_genes file - ISAR_GISTIC.all_thresholded.by_genes.txt, extract to **ISAR_GISTIC.all_thresholded.by_genes.txt** and rename to **ISAR_GISTIC.all_thresholded.by_genes.tsv**
5) Delete from the file **ISAR_GISTIC.all_thresholded.by_genes.tsv** all samples named TCGA-XX-XXXX-YYX-XXX-XXXX-XX, where YY – any number, except 01, 03 и 09; and all samples present in the file **merged_sample_quality_annotations_do_not_use.tsv,** and save the resulting file as **ISAR_GISTIC.all_thresholded.by_genes_primary_whitelisted.tsv**
6) Download the file RNA batch corrected matrix - EBPlusPlusAdjustPANCAN_IlluminaHiSeq_RNASeqV2.geneExp.tsv
7) Delete from the file **EBPlusPlusAdjustPANCAN_IlluminaHiSeq_RNASeqV2-v2.geneExp.tsv** all samples named TCGA-XX-XXXX-YYX-XXX-XXXX-XX, where YY – any number, except 01, 03 и 09; and all samples present in the file **merged_sample_quality_annotations_do_not_use.tsv,** and save the resulting file as **EBPlusPlusAdjustPANCAN_IlluminaHiSeq_RNASeqV2-v2.geneExp_primary_whitelisted.tsv**
8) Download the file miRNA batch corrected matrix - pancanMiRs_EBadjOnProtocolPlatformWithoutRepsWithUnCorrectMiRs_08_04_16.csv and convert to **pancanMiRs_EBadjOnProtocolPlatformWithoutRepsWithUnCorrectMiRs_08_04_16.tsv**
9) Delete from the file **pancanMiRs_EBadjOnProtocolPlatformWithoutRepsWithUnCorrectMiRs_08_04_16.tsv** all samples named TCGA-XX-XXXX-YYX-XXX-XXXX-XX, where YY – any number, except 01, 03 и 09; and all samples present in the file **merged_sample_quality_annotations_do_not_use.tsv,** and save the resulting file as **pancanMiRs_EBadjOnProtocolPlatformWithoutRepsWithUnCorrectMiRs_08_04_16_primary_whitelisted.tsv**
10) Using the file **EBPlusPlusAdjustPANCAN_IlluminaHiSeq_RNASeqV2-v2.geneExp_primary_whitelisted.tsv**, determine the median expression level for each gene across patients. If the expression for a given gene in a given patient is below 0.05x median value, replace it with "-2", if between 0.05x and 0.75x median value, replace it with "-1", if between 1.25x and 1.75x median value, replace with "1", if above 1.75x median value, replace with "2", otherwise replace with "0". Save the file as **EBPlusPlusAdjustPANCAN_IlluminaHiSeq_RNASeqV2-v2.geneExp_primary_whitelisted_median.tsv**
11) Using the file **pancanMiRs_EBadjOnProtocolPlatformWithoutRepsWithUnCorrectMiRs_08_04_16_primary_whitelisted.tsv**, determine the median expression level for each miRNA across patients. If the expression for a given miRNA in a given patient is below 0.05x median value, replace it with "-2", if between 0.05x and 0.75x median value, replace it with "-1", if between 1.25x and 1.75x median value, replace with "1", if above 1.75x median value, replace with "2", otherwise replace with "0". Save the file as

**pancanMiRs_EBadjOnProtocolPlatformWithoutRepsWithUnCorrectMiRs_08_04_16_primary_whitelisted_median.tsv**

12) Process the file **ISAR_GISTIC.all_thresholded.by_genes_primary_whitelisted.tsv** according to the following table:

| | | **Gene CNA status in a given patient** ISAR_GISTIC.all_thresholded.by_genes_primary_whitelisted.tsv | | | | |
|---|---|---|---|---|---|---|
| | | **-2** | **-1** | **0** | **1** | **2** |
| **Gene expression status in the same patient** EBPlusPlusAdjustPANCAN_IlluminaHiSeq_RNASeqV2-v2.geneExp_primary_whitelisted_median.tsv | **2** | 0 | 0 | 0 | 2 | 2 |
| | **1** | 0 | 0 | 0 | 1 | 1 |
| | **0** | 0 | 0 | 0 | 0 | 0 |
| | **-1** | -1 | -1 | 0 | 0 | 0 |
| | **-2** | -2 | -2 | 0 | 0 | 0 |
| **If gene not found look in the miRNA file:** | | | | | | |
| **miRNA expression status in the same patient** pancanMiRs_EBadjOnProtocolPlatformWithoutRepsWithUnCorrectMiRs_08_04_16_primary_whitelisted_median.tsv | **2** | 0 | 0 | 0 | 2 | 2 |
| | **1** | 0 | 0 | 0 | 1 | 1 |
| | **0** | 0 | 0 | 0 | 0 | 0 |
| | **-1** | -1 | -1 | 0 | 0 | 0 |
| | **-2** | -2 | -2 | 0 | 0 | 0 |
| **If still not found then:** | | -2 | -1 | 0 | 1 | 2 |

Save the results as
**ISAR_GISTIC.all_thresholded.by_genes_primary_whitelisted_RNAfiltered.tsv**
