## Supplementary material for "Comprehensive patient-level classification and quantification of driver events in TCGA PanCanAtlas cohorts": S5 Files: run_all.py

```
# -*- coding: utf-8 -*-

import os

from step_2 import step2
from step_3 import step3
from step_4 import step4
from step_5 import step5
from step_6 import step6
from step_7 import step7
from step_8 import step8
from step_9 import step9
from step_10 import step10
from step_11 import step11
from step_12 import step12

for i in range(2, 12+1):
    step_to_call = locals()[f'step{i}']
    step_to_call()
```
