## Supplementary material for "Comprehensive patient-level classification and quantification of driver events in TCGA PanCanAtlas cohorts": S5 Files: step_2.py

```
# -*- coding: utf-8 -*-

from requests import get
from utils.utils import download, check_hash

import pandas as pd
import os
import argparse

def step2(save_folder: str = 'data'):

    print('Step 2')

    if not os.path.isdir(save_folder):
        os.makedirs(save_folder)

    output_file_path = os.path.join(save_folder, 'merged_sample_quality_annotations.tsv')

    if not os.path.isfile(output_file_path):
        # Downloading
        download_link = 'https://api.gdc.cancer.gov/data/1a7d7be8-675d-4e60-a105-19d4121bdebf'
        download(link=download_link, destination=output_file_path)

    # Hash sum check
    hash_pass = check_hash(output_file_path)
    if not hash_pass:
        raise Exception(f'Hash sum validation for {output_file_path} was not passed, check the file or download it again')

    print('OK')


if __name__ == '__main__':
    parser = argparse.ArgumentParser(description='This script downloads merged_sample_quality_annotations.tsv file ' + '\n' +
                                                 'and saves it to the data/ folder by default')
    parser.add_argument('-s', '--save_folder', type=str, help='full path to folder to save downloaded files', default='data')
    args = parser.parse_args()

    step2(args.save_folder)
```
