## Supplementary material for "Comprehensive patient-level classification and quantification of driver events in TCGA PanCanAtlas cohorts": S5 Files: step_3.py

```
# -*- coding: utf-8 -*-

import pandas as pd
import os
import argparse

from utils.utils import check_filesize


def get_sample_code(tcga_id):
    return tcga_id.split('-')[3][:2]

def step3(input_dir: str = 'data', output_folder_path: str = 'data'):

    print('Step 3')

    input_path = os.path.join(input_dir, 'merged_sample_quality_annotations.tsv')

    output_file_path = os.path.join(output_folder_path, 'merged_sample_quality_annotations_do_not_use.tsv')

    if not os.path.isfile(output_file_path):

        sample_annot_df = pd.read_csv(input_path, sep='\t')

        first_condition = sample_annot_df['aliquot_barcode'].apply(get_sample_code).isin(['01', '03', '09'])
        second_condition = sample_annot_df['Do_not_use'] == True

        sample_annot_df[first_condition & second_condition].to_csv(output_file_path,
                                                                   header=True,
                                                                   index=False,
                                                                   sep='\t')
    # Check for filesize
    size_pass = check_filesize(output_file_path)
    if not size_pass:
        raise Exception(f'file: {output_file_path} has wrong size, please check input file and do this step again')

    print('OK')


if __name__ == '__main__':
    parser = argparse.ArgumentParser(description='This script takes file merged_sample_quality_annotations.tsv,' +
                                                 '\n' + 'deletes all the aliqoutes but 01, 03, 09 (column aliquot_barсode),' +
                                                 '\n' + 'deletes all the aliqoutes with Do_not_use=False value.' +
                                                 '\n' + 'If output folder does not exist, script will create it.')
    parser.add_argument('-d', '--input_dir', type=str, help='full path to input folder',
                        default='data')
    parser.add_argument('-o', '--output_folder', type=str, help='full path to output folder', default='data')
    args = parser.parse_args()

    step3(args.input_dir , args.output_folder)
```
