## Supplementary material for "Comprehensive patient-level classification and quantification of driver events in TCGA PanCanAtlas cohorts": S5 Files: step_4.py

```
# -*- coding: utf-8 -*-
import os
import shutil

import pandas as pd
import argparse
import gzip

from utils.utils import download, check_hash

def step4(save_folder: str = 'data'):

    print('Step 4')

    if not os.path.isdir(save_folder):
        os.makedirs(save_folder)

    archive_file_path = os.path.join(save_folder, 'ISAR_GISTIC.all_thresholded.by_genes.txt.gz')
    output_file_path = archive_file_path.replace('.txt.gz', '.tsv')

    # download
    if not os.path.isfile(output_file_path):
        # download file
        download_link = 'https://api.gdc.cancer.gov/data/a9dae2ab-9462-4f9a-9730-5bad520ab2d7'
        download(download_link, archive_file_path)

        # unzip file
        with gzip.open(archive_file_path, 'rb') as f_in:
            with open(output_file_path.replace('.tsv', '.txt'), 'wb') as f_out:
                shutil.copyfileobj(f_in, f_out)
        # remove archive
        os.remove(archive_file_path)

        # rename to .tsv
        os.rename(output_file_path.replace('.tsv', '.txt'), output_file_path)

    # Check hashsum
    hash_pass = check_hash(output_file_path)
    if not hash_pass:
        raise Exception(f'Hash sum validation for {output_file_path} was not passed, check the file or download it again')

    print('OK')


if __name__ == '__main__':
    parser = argparse.ArgumentParser(description='This script downloads ISAR_GISTIC.all_thresholded.by_genes.txt file, ' + '\n' +
                                                 'changes its extention to ".tsv" and saves it to the data/ folder by default')
    parser.add_argument('-s', '--save_folder', type=str, help='full path to folder to save downloaded files', default='data')
    args = parser.parse_args()

    step4(args.save_folder)
```
