## Supplementary material for "Comprehensive patient-level classification and quantification of driver events in TCGA PanCanAtlas cohorts": S5 Files: step_7.py

```
# -*- coding: utf-8 -*-

import argparse
import os
import pandas as pd

from utils.utils import check_filesize


def get_sample_code(tcga_id):
    return tcga_id.split('-')[3][:2]


def get_index(header, column_name):
    header = header.split('\t')
    for i in range(len(header)):
        if header[i] == column_name:
            return i
    return -1


def step7(input_dir: str = 'data', output_folder_path: str = 'data'):

    print('Step 7')

    input_path = os.path.join(input_dir, 'EBPlusPlusAdjustPANCAN_IlluminaHiSeq_RNASeqV2-v2.geneExp.tsv')
    banned_samples_path = os.path.join(input_dir, 'merged_sample_quality_annotations_do_not_use.tsv')
    output_file_path = os.path.join(output_folder_path,
                                    'EBPlusPlusAdjustPANCAN_IlluminaHiSeq_RNASeqV2-v2.geneExp_primary_whitelisted.tsv')

    if not os.path.isdir(output_folder_path):
        os.makedirs(output_folder_path)

    if not os.path.isfile(output_file_path):

        # get banned samples set
        banned_samples = set(pd.read_csv(banned_samples_path,
                                         sep='\t',
                                         usecols=['aliquot_barcode'])['aliquot_barcode'])

        with open(input_path, 'r') as input_rna_file:
            header = input_rna_file.readline()[:-1].split('\t')
        header = [column.replace('"', '') for column in header]

        all_samples = set(header[1:])
        print('There were {} samples'.format(len(all_samples)))

        new_columns = [header[i] for i in range(1)]

        selected_samples = set()
        for barcode in header[1:]:
            first_condition = get_sample_code(barcode) in ['01', '03', '09']
            second_condition = barcode not in banned_samples
            if first_condition and second_condition:
                new_columns.append(barcode)
                selected_samples.add(barcode)
        print('{} samples were selected'.format(len(selected_samples)))

        new_columns_indexes = []
        for ix, column in enumerate(header):
            if column in new_columns:
                new_columns_indexes.append(ix)

        with open(input_path, 'r') as input_rna_file, open(output_file_path, 'w') as output_rna_file:
            output_rna_file.write('\t'.join(new_columns) + '\n')
            header = input_rna_file.readline()
            content_line = input_rna_file.readline()
            while content_line:
                content_line = content_line.split('\t')
                if content_line[new_columns_indexes[-1]][-1] == '\n':
                    output_rna_file.write('\t'.join([content_line[ix] for ix in new_columns_indexes]))
                else:
                    output_rna_file.write('\t'.join([content_line[ix] for ix in new_columns_indexes]) + '\n')
                content_line = input_rna_file.readline()

    # Check for filesize
    size_pass = check_filesize(output_file_path)
    if not size_pass:
        raise Exception(f'file: {output_file_path} has wrong size, please check input file and do this step again')

    print('OK')


if __name__ == '__main__':
    parser = argparse.ArgumentParser(description='This script takes ' +
                                                 'EBPlusPlusAdjustPANCAN_IlluminaHiSeq_RNASeqV2-v2.geneExp.tsv file,' + '\n' +
                                                 'keeps only 01, 03, 09 samples (column Tumor_Sample_Barcode),' +
                                                 '\n' + 'deletes all the aliquotes which mentioned in file: ' +
                                                 'merged_sample_quality_annotations_do_not_use.tsv (column aliquot_barcode).'
                                                 + '\n' + 'If output folder does not exist, script will create it.')
    parser.add_argument('-i', '--input_dir', type=str, help='full path to input folder', default='data')
    parser.add_argument('-o', '--output_folder', type=str, help='full path to output folder', default='data')

    args = parser.parse_args()

    step7(args.input_dir , args.output_folder)
```
