## Supplementary material for "Comprehensive patient-level classification and quantification of driver events in TCGA PanCanAtlas cohorts": S5 Files: step_8.py

```
# -*- coding: utf-8 -*-

import pandas as pd
import os
import argparse

from utils.utils import download, check_hash

def step8(save_folder: str = 'data'):

    print('Step 8')

    if not os.path.isdir(save_folder):
        os.makedirs(save_folder)

    output_file_path = os.path.join(save_folder, 'pancanMiRs_EBadjOnProtocolPlatformWithoutRepsWithUnCorrectMiRs_08_04_16.tsv')
    if not os.path.isfile(output_file_path):
        download_link = 'https://api.gdc.cancer.gov/data/1c6174d9-8ffb-466e-b5ee-07b204c15cf8'
        download(download_link, output_file_path.replace('.tsv', '.csv'))

        # Convert to .tsv
        df = pd.read_csv(output_file_path.replace('.tsv', '.csv'))
        df.to_csv(output_file_path, sep='\t', index=False)

        # Remove .csv file
        os.remove(output_file_path.replace('.tsv', '.csv'))

    # check hashsum
    hash_pass = check_hash(output_file_path)
    if not hash_pass:
        raise Exception(f'Hash sum validation for {output_file_path} was not passed, check the file or download it again')

    print('OK')


if __name__ == '__main__':
    parser = argparse.ArgumentParser(description=\
        'This script downloads pancanMiRs_EBadjOnProtocolPlatformWithoutRepsWithUnCorrectMiRs_08_04_16.csv file, ' + '\n' + \
        'converts to ".tsv" and saves it to the data/ folder by default')
    parser.add_argument('-s', '--save_folder', type=str, help='full path to folder to save downloaded files', default='data')
    args = parser.parse_args()

    step8(args.save_folder)
```
