## Supplementary material for "Comprehensive patient-level classification and quantification of driver events in TCGA PanCanAtlas cohorts": S5 Files: step_10.py

```
# -*- coding: utf-8 -*-

import argparse
import os
from tqdm import tqdm
import numpy as np

from utils.utils import check_filesize


def cut_expressions(expression, median_value):
    if np.isnan(expression):
        return 0
    else:
        if expression < median_value * 0.05: # almost zero, double deletion
            return -2
        elif expression < median_value * 0.75: # significantly lower, single deletion
            return -1
        elif expression > median_value * 1.75: # almost double, double amplification
            return 2
        elif expression > median_value * 1.25: # significantly bigger, single amplification
            return 1
        else: # no event
            return 0


def step10(input_dir: str = 'data', output_folder_path: str = 'data'):

    print('Step 10')

    input_path = os.path.join(input_dir, 'EBPlusPlusAdjustPANCAN_IlluminaHiSeq_RNASeqV2-v2.geneExp_primary_whitelisted.tsv')

    if not os.path.isdir(output_folder_path):
        os.makedirs(output_folder_path)

    output_path = os.path.join(output_folder_path,
                               'EBPlusPlusAdjustPANCAN_IlluminaHiSeq_RNASeqV2-v2.geneExp_primary_whitelisted_median.tsv')

    if not os.path.isfile(output_path):

        print('We will read the file in a memory efficient way, line by line')
        print('There are 20531 strings in the file, so 20531 iterations required')

        with open(input_path, 'r') as input_file, open(output_path, 'w') as output_file:
            header = input_file.readline()
            output_file.write(header)
            for line in tqdm(input_file):
                gene_name = line.split('\t')[0]
                line = line.replace('NA', 'nan')
                line = line.replace('\n', '')
                numbers = np.array([float(float_number) for float_number in line.split('\t')[1:]])
                median_value = np.median(numbers)
                vect_expression_flag = np.vectorize(lambda x: cut_expressions(x, median_value), otypes='f')
                numbers_flags = vect_expression_flag(numbers)

                output_file.write('\t'.join([gene_name] + [str(int(x)) for x in numbers_flags]) + '\n')

    # Check for filesize
    size_pass = check_filesize(output_path)
    if not size_pass:
        raise Exception(f'file: {output_path} has wrong size, please check input file and do this step again')

    print('OK')


if __name__ == '__main__':
    parser = argparse.ArgumentParser(description='This script takes ' +
                                                 'EBPlusPlusAdjustPANCAN_IlluminaHiSeq_RNASeqV2-v2.geneExp_primary_whitelisted.tsv ' + \
                                                 'file,' + '\n' + 'and and does cutoffs' + '\n' + \
                                                 'If output folder does not exist, script will create it.')
    parser.add_argument('-i', '--input_dir', type=str, help='full path to folder', default='data')
    parser.add_argument('-o', '--output_folder', type=str, help='full path to output folder', default='data')

    args = parser.parse_args()

    step10(args.input_dir , args.output_folder)
```
