## Supplementary material for "Comprehensive patient-level classification and quantification of driver events in TCGA PanCanAtlas cohorts": S5 Files: step_12.py

```
# -*- coding: utf-8 -*-
import os
import argparse
from collections import Counter
import shutil
import gzip

import pandas as pd
import numpy as np
from tqdm import tqdm

from utils.utils import check_filesize, check_hash, download_ftp


def to_patient(barcode):
    return '-'.join(barcode.split('-')[:3])

'''
def filter_val(cna_val, other_val):
    if cna_val * other_val > 0:
        if np.abs(cna_val) == 1 or np.abs(other_val) == 1:
            result = 1 * np.sign(cna_val)
        else:
            result = 2 * np.sign(cna_val)
    else:
        result = 0
    return np.int16(result)
'''

def filter_val(cna_val, other_val):
    if cna_val * other_val > 0:
        if np.abs(other_val) == 1:
            result = 1 * np.sign(cna_val)
        else:
            result = 2 * np.sign(cna_val)
    else:
        result = 0
    return np.int16(result)


def decide_by_rows(row1, row2):
    row1 = np.array(row1)
    row2 = np.array(row2)
    return np.max([row1, row2], axis=0)


def find_dtypes(df_path):
    with open(df_path, 'r') as df_file:
        first_line = df_file.readline().replace('\n', '')
    column_names = first_line.split('\t')
    dtype_dict = {}
    for column in column_names:
        if 'TCGA' in column:
            dtype_dict[column] = np.int16
        else:
            dtype_dict[column] = str
    return dtype_dict


def step12(input_dir: str = 'data', output_folder_path: str = 'data'):

    print('Step 12')

    cna_path = os.path.join(input_dir, 'ISAR_GISTIC.all_thresholded.by_genes_primary_whitelisted.tsv')
    rna_path = os.path.join(input_dir, 'EBPlusPlusAdjustPANCAN_IlluminaHiSeq_RNASeqV2-v2.geneExp_primary_whitelisted_median.tsv')
    mi_rna_path = os.path.join(input_dir, 'pancanMiRs_EBadjOnProtocolPlatformWithoutRepsWithUnCorrectMiRs_08_04_16_primary_whitelisted_median.tsv')
    gene_annot_path = os.path.join(input_dir, 'Homo_sapiens.gene_info')

    if not os.path.isfile(gene_annot_path):
        download_link = 'ftp://ftp.ncbi.nih.gov/gene/DATA/GENE_INFO/Mammalia/Homo_sapiens.gene_info.gz'
        download_ftp(download_link, gene_annot_path+'.gz')

        # unzip file
        with gzip.open(gene_annot_path+'.gz', 'rb') as f_in:
            with open(gene_annot_path, 'wb') as f_out:
                shutil.copyfileobj(f_in, f_out)

        # remove archive
        os.remove(gene_annot_path+'.gz')

    # Check size
    size_pass = check_filesize(gene_annot_path)
    if not size_pass:
        raise Exception(f'file: {gene_annot_path} has wrong size, please check input file and do this step again')

    output_file_path = os.path.join(output_folder_path, 'ISAR_GISTIC.all_thresholded.by_genes_primary_whitelisted_RNAfiltered.tsv')

    if not os.path.isdir(output_folder_path):
        os.makedirs(output_folder_path)

    if not os.path.isfile(output_file_path):

        vect_filter_val = np.vectorize(filter_val)

        print('Reading gene annotation table')
        required_columns = ['GeneID', 'Symbol', 'Synonyms']
        gene_corresp_df = pd.read_csv(gene_annot_path, sep='\t', usecols=required_columns)

        print('CNA file reading and preprocessing')
        cna_dtypes = find_dtypes(cna_path)
        cna_df = pd.read_csv(cna_path, sep='\t', header=0, dtype=cna_dtypes)
        cna_df.index = np.array(cna_df['Locus ID'], dtype=int)
        cna_df.drop(columns=['Locus ID'], inplace=True)
        cna_df.rename(columns={'Gene Symbol': 'gene_name'})
        num_columns_cna = len(cna_df.columns)
        cna_df.columns = list(map(to_patient, list(cna_df.columns)))
        cna_df.index.name = 'gene_id'

        # not unique genes deduplication
        cna_df = cna_df.loc[cna_df.index.drop_duplicates(keep=False)]

        print('RNA file reading and preprocessing')
        rna_dtypes = find_dtypes(rna_path)
        rna_df = pd.read_csv(rna_path, sep='\t', header=0, dtype=rna_dtypes)
        rna_df.columns = list(map(to_patient, list(rna_df.columns)))
        rna_df.insert(loc=0, column='gene_name', value=[s.split('|')[0].upper() for s in rna_df['gene_id']])
        rna_df['gene_id'] = [int(s.split('|')[1].upper()) for s in rna_df['gene_id']]
        rna_df.index = rna_df['gene_id']
        rna_df.index.name = 'gene_id'
        rna_df.drop(columns=['gene_id'], inplace=True)

        # not unique patients investigation
        rna_counter = Counter(list(rna_df.columns))
        not_unique_patients = [patient for patient, count in rna_counter.items() if count > 1]
        # remove those patients
        rna_df.drop(columns=not_unique_patients, inplace=True)

        # not unique genes deduplication
        rna_df = rna_df.loc[rna_df.index.drop_duplicates(keep=False)]

        # we choose common patients and genes, and apply rule to them
        print('CNA - RNA filtering')
        CNA_RNA_patients = sorted(list(set(rna_df.columns).intersection(set(cna_df.columns))))
        CNA_RNA_genes = sorted(list(set(rna_df.index).intersection(set(cna_df.index))))

        #cna_df.loc[CNA_RNA_genes, CNA_RNA_patients] = vect_filter_val(cna_df.loc[CNA_RNA_genes, CNA_RNA_patients],
        #                                                              rna_df.loc[CNA_RNA_genes, CNA_RNA_patients])
        cna_df.loc[CNA_RNA_genes, CNA_RNA_patients] = cna_df.loc[CNA_RNA_genes, CNA_RNA_patients]\
            .combine(rna_df.loc[CNA_RNA_genes, CNA_RNA_patients], vect_filter_val)

        print('miRNA file reading and preprocessing')
        mi_rna_dtypes = find_dtypes(mi_rna_path)
        mi_rna_df = pd.read_csv(mi_rna_path, sep='\t', header=0, dtype=mi_rna_dtypes)
        mi_rna_df.columns = list(map(to_patient, list(mi_rna_df.columns)))
        mi_rna_df['gene_id'] = [x.replace('hsa-', '') for x in mi_rna_df['gene_id']]
        mi_rna_df = mi_rna_df.rename(columns={'gene_id': 'gene_name'})
        mi_rna_df = mi_rna_df.sort_values(by='gene_name')
        mi_rna_df.index = mi_rna_df['gene_name']
        mi_rna_df.drop(columns=['gene_name'], inplace=True)

        # find 3-prime, 5-prime pairs
        sorted_gene_names = list(mi_rna_df.index)
        corresp_3_to_5 = dict()
        for row_ix in range(len(sorted_gene_names) - 1):
            if sorted_gene_names[row_ix].replace('3p', '5p') == sorted_gene_names[row_ix + 1]:
                corresp_3_to_5[sorted_gene_names[row_ix]] = sorted_gene_names[row_ix + 1]
        # choose max among them
        for gene_name_3p in corresp_3_to_5:
            gene_name_5p = corresp_3_to_5[gene_name_3p]
            mi_rna_df.loc[gene_name_3p] = decide_by_rows(mi_rna_df.loc[gene_name_3p].values,
                                                         mi_rna_df.loc[gene_name_5p].values)
        # delete unnecessary 5-prime
        indexes_to_delete = list(corresp_3_to_5.values())
        mi_rna_df.drop(index=indexes_to_delete, inplace=True)
        mi_rna_df.index = [x.replace('-3p', '').replace('-5p', '') for x in mi_rna_df.index]

        #finally match genes
        corresp_ids = []
        for query_gene in mi_rna_df.index:
            found = False
            for ref_ix, ref_gene_name in enumerate(gene_corresp_df['Symbol']):
                ref_gene_name = ref_gene_name.lower()
                query_gene = query_gene.replace('-', '').lower()
                if 'mir' + query_gene == ref_gene_name or query_gene == ref_gene_name:
                    corresp_ids.append(gene_corresp_df['GeneID'][ref_ix])
                    found = True
            if not found:
                corresp_ids.append(None)

        mi_rna_df = mi_rna_df.reset_index().rename(columns={'index': 'gene_name'})
        mi_rna_df.insert(loc=0, column='gene_id', value=corresp_ids)
        mi_rna_df = mi_rna_df[mi_rna_df['gene_id'].notna()]
        mi_rna_df.index = mi_rna_df['gene_id'].astype(int)
        mi_rna_df = mi_rna_df.drop(columns=['gene_id'])

        # not unique patients investigation
        mi_rna_counter = Counter(list(mi_rna_df.columns))
        not_unique_patients = [patient for patient, count in mi_rna_counter.items() if count > 1]
        # remove them
        mi_rna_df.drop(columns=not_unique_patients, inplace=True)

        # not unique genes deduplication
        mi_rna_df = mi_rna_df.loc[mi_rna_df.index.drop_duplicates(keep=False)]


        print('CNA - miRNA filtering')
        CNA_miRNA_patients = sorted(list(set(mi_rna_df.columns).intersection(set(cna_df.columns))))
        CNA_miRNA_genes = sorted(list(set(mi_rna_df.index).intersection(set(cna_df.index))))

        #cna_df.loc[CNA_miRNA_genes, CNA_miRNA_patients] = vect_filter_val(cna_df.loc[CNA_miRNA_genes, CNA_miRNA_patients],
        #                                                              mi_rna_df.loc[CNA_miRNA_genes, CNA_miRNA_patients])
        cna_df.loc[CNA_miRNA_genes, CNA_miRNA_patients] = cna_df.loc[CNA_miRNA_genes, CNA_miRNA_patients]\
            .combine(mi_rna_df.loc[CNA_miRNA_genes, CNA_miRNA_patients], vect_filter_val)

        print('saving the results')
        cna_df.to_csv(output_file_path, sep = '\t', header=True, index=True)

    # Check for filesize
    size_pass = check_filesize(output_file_path)
    if not size_pass:
        raise Exception(f'file: {output_file_path} has wrong size, please check input file and do this step again')

    print('OK')


if __name__ == '__main__':
    parser = argparse.ArgumentParser(description='This script takes ' +
                                                 'ISAR_GISTIC.all_thresholded.by_genes_primary_whitelisted.tsv file (CNA),' + '\n' + \
                                                 'EBPlusPlusAdjustPANCAN_IlluminaHiSeq_RNASeqV2-v2.geneExp_primary_whitelisted_quartiles.tsv file (RNA),' + '\n' + \
                                                 'pancanMiRs_EBadjOnProtocolPlatformWithoutRepsWithUnCorrectMiRs_08_04_16_primary_whitelisted_quartiles.tsv file (miRNA),' \
                                                 + '\n' + 'and basically filters CNA file where possible.' + '\n' + \
                                                 'If output folder does not exist, script will create it.')
    parser.add_argument('-i', '--input_dir', type=str, help='full path to input folder', default='data')
    parser.add_argument('-o', '--output_folder', type=str, help='full path to output folder', default='data')

    args = parser.parse_args()

    step12(args.input_dir , args.output_folder)
```
