## Supplementary material for "Comprehensive patient-level classification and quantification of driver events in TCGA PanCanAtlas cohorts": S5 Files: utils.py

```
import hashlib
import requests
import urllib.request
import sys
import os


def download(link, destination):
	with open(destination, "wb") as f:
		response = requests.get(link, stream=True)
		total_length = response.headers.get('content-length')

		if total_length is None: # no content length header
			f.write(response.content)
		else:
			dl = 0
			total_length = int(total_length)
			for data in response.iter_content(chunk_size=4096):
				dl += len(data)
				f.write(data)
				done = int(50 * dl / total_length)
				sys.stdout.write("\r[%s%s]" % ('=' * done, ' ' * (50-done)) )
				sys.stdout.flush()
	print("\n")


def download_ftp(link, destination):

	with urllib.request.urlopen(link) as response, open(destination, 'wb') as out_file:
	    data = response.read() # a `bytes` object
	    out_file.write(data)


def check_hash(file_path):
	# Read reference hash
	hash_dict = {}
	with open('utils/hashes.txt', 'r') as hash_file:
		for line_ix, line in enumerate(hash_file):
			if line_ix == 0:
				continue
			line = line.replace('\n', '')
			filename, md5_value, sha1_value = line.split()
			hash_dict[filename] = (md5_value, sha1_value)

	# Calculate hash from the file itself
	BUF_SIZE = 65536
	with open(file_path, 'rb') as f:
		md5 = hashlib.md5()
		sha1 = hashlib.sha1()
		while True:
			data = f.read(BUF_SIZE)
			if not data:
				break
			md5.update(data)
			sha1.update(data)
	current_filename = os.path.basename(file_path)

	if hash_dict[current_filename] == (md5.hexdigest(),sha1.hexdigest()):
		return True
	else:
		return False


def check_filesize(file_path):

	# Read reference size
	sizes_dict = {}
	with open('utils/sizes.txt', 'r') as sizes_file:
		for line_ix, line in enumerate(sizes_file):
			if line_ix == 0:
				continue
			line = line.replace('\n', '')
			filename, size = line.split()
			sizes_dict[filename] = int(size)
	reference_size = sizes_dict[os.path.basename(file_path)]

	# Find actual file size
	actual_file_size = os.stat(file_path).st_size # in bytes

	# Compare if filesize within interval
	percent_delta = 5 # delta in size measurement
	lower_bound = (1 - percent_delta / 100) * reference_size
	upper_bound = (1 + percent_delta / 100) * reference_size

	if (actual_file_size > lower_bound) and (actual_file_size < upper_bound):
		return True
	else:
		return False
```
