## Supplementary material for "Comprehensive patient-level classification and quantification of driver events in TCGA PanCanAtlas cohorts": S6 Files: ANDRIF description.pdf

ANDRIF (ANeuploidy DRIver Finder) is a Python 3.7 software package that predicts cancer driver aneuploidies (i.e. chromosomal arm or full chromosome gains or losses) from the TCGA PanCanAtlas aneuploidy data (https://gdc.cancer.gov/about-data/publications/PanCan-CellOfOrigin). Low quality samples and metastatic samples are filtered out. Driver prediction is based on calculating the average alteration status for each arm or chromosome in each cancer type. Bootstrapping is used to obtain the realistic distribution of the average alteration statuses under the null hypothesis. Benjamini–Hochberg procedure is performed to keep the false discovery rate under 5%. The pipeline can be executed fully automatically in less than 30 minutes on a modern PC (Linux, Windows or MacOS).

This package has been developed by

Aleksey V. Belikov, Dr.rer.nat.
https://github.com/belikov-av
Laboratory of Innovative Medicine, School of Biological and Medical Physics, Moscow Institute of Physics and Technology
Roles: concept, pipeline, supervision

and

Alexey D. Vyatkin
https://github.com/VyatkinAlexey
Skoltech
Roles: programming

A detailed description of pipeline steps can be found in the file ANDRIF pipeline.pdf

Instructions for executing the code can be found in the file Instructions.txt
