## Supplementary material for "Comprehensive patient-level classification and quantification of driver events in TCGA PanCanAtlas cohorts": S6 Files: ANDRIF pipeline.pdf

1. Go to https://gdc.cancer.gov/about-data/publications/PanCan-CellOfOrigin
2. Download the file Analyte level annotations - merged_sample_quality_annotations.tsv
3. Using information in the column **aliquot_barcode**, delete all aliquots named TCGA-XX-XXXX-YYX-XXX-XXXX-XX, where YY – any number, except 01, 03 and 09 (see https://docs.gdc.cancer.gov/Encyclopedia/pages/TCGA_Barcode/ and table https://gdc.cancer.gov/resources-tcga-users/tcga-code-tables/sample-type-codes), and also, using information in the column **Do_not_use**, delete all aliquots with False value, and save the resulting file as **merged_sample_quality_annotations_do_not_use.tsv**.
4. Download the file ABSOLUTE purity/ploidy - TCGA_mastercalls.abs_tables_JSedit.fixed.txt
5. Remove samples with identifiers TCGA-XX-XXXX-YY, where YY – any number, except 01, 03 и 09; with **Cancer DNA fraction** <0.5 or unknown; with **Subclonal genome fraction** >0.5 or unknown; and with identifiers from **merged_sample_quality_annotations_do_not_use.tsv**; and save file as **TCGA_mastercalls.abs_tables_JSedit.fixed_primary_whitelisted.tsv**
6. Download the file Aneuploidy scores and arm calls - PANCAN_ArmCallsAndAneuploidyScore_092817.txt
7. Remove from the file **PANCAN_ArmCallsAndAneuploidyScore_092817.txt** all samples not present in **TCGA_mastercalls.abs_tables_JSedit.fixed_primary_whitelisted.tsv**, and save as **Primary_whitelisted_arms.tsv**
8. Calculate the values for the whole chromosomes using these rules:

| | | | | | | | | |
|---|---|---|---|---|---|---|---|---|
| **IF p-arm status is** | <empty cell> | 0 OR 1 OR -1 | 0 | 1 OR -1 | 1 | -1 | 1 | -1 |
| **AND q-arm Status is** | <empty cell> OR 0 OR 1 OR -1 | <empty cell> | 0 OR 1 OR -1 | 0 | -1 | 1 | 1 | -1 |
| **THEN chromosome status is** | <empty cell> | <empty cell> | 0 | 0 | 0 | 0 | 1 | -1 |

   For one-arm chromosomes (13, 14, 15, 21, 22), their status equals the status of the arm. Save the file as **Primary_whitelisted_chromosomes.tsv**
9. Using the file **Primary_whitelisted_arms.tsv**, for each cancer **Type** calculate the average alteration status of each chromosomal arm (1p, 1q, 2p, 2q, 3p, 3q, 4p, 4q, 5p, 5q, 6p, 6q, 7p, 7q, 8p, 8q, 9p, 9q, 10p, 10q, 11p, 11q, 12p, 12q, 13q, 14q, 15q, 16p, 16q, 17p, 17q, 18p, 18q, 19p, 19q, 20p, 20q, 21q, 22q). Ignore empty cells both in the numerator and in the denominator, when calculating the averages. Save the results in the table **Arm_averages.tsv**
10. Using the file **Primary_whitelisted_chromosomes.tsv**, for each cancer **Type** calculate the average alteration status of each chromosome (1, 2, 3, 4, 5, 6, 7, 8, 9, 10, 11, 12, 13, 14, 15, 16, 17, 18, 19, 20, 21, 22). Ignore empty cells both in the numerator and in the denominator, when calculating the averages. Save the results in the table **Chromosome_averages.tsv**
11. By drawing statuses randomly with replacement (bootstrapping) from *any* cell of **Primary_whitelisted_arms.tsv**, for each cancer type generate the number of statuses corresponding to the number of patients in that cancer type and calculate their average (ignoring empty cells when calculating the average). Repeat the procedure 10000 times, calculate the median for each cancer type and save the results as **Bootstrapped_arm_averages.tsv**
12. By drawing statuses randomly with replacement (bootstrapping) from *any* cell of **Primary_whitelisted_chromosomes.tsv**, for each cancer type generate the number of statuses corresponding to the number of patients in that cancer type and calculate their average (ignoring

empty cells when calculating the average). Repeat the procedure 10000 times, calculate the median for each cancer type and save the results as **Bootstrapped_chromosome_averages.tsv**

13. For each cancer type, calculate P-value for each arm alteration status. To do this, first compare the alteration status for a given cancer type and a given arm in **Arm_averages.tsv** to the median bootstrapped arm alteration status for this cancer type in **Bootstrapped_arm_averages.tsv.** If the status in **Arm_averages.tsv** is higher than zero AND the median in **Bootstrapped_arm_averages.tsv,** count how many statuses for this cancer type in **Bootstrapped_arm_averages.tsv** are higher than the status in **Arm_averages.tsv,** and divide this number by 5000. If the status in **Arm_averages.tsv** is lower than zero AND the median in **Bootstrapped_arm_averages.tsv,** count how many statuses for this cancer type in **Bootstrapped_arm_averages.tsv** are lower than the status in **Arm_averages.tsv,** divide this number by 5000, and add minus to indicate arm loss. Ignore other values (leave cells empty). Save the file as **Arm_Pvalues_cohorts.tsv**
14. For each cancer type, calculate P-value for each chromosome alteration status. To do this, first compare the alteration status for a given cancer type and a given chromosome in **Chromosome_averages.tsv** to the median bootstrapped chromosome alteration status for this cancer type in **Bootstrapped_chromosome_averages.tsv.** If the status in **Chromosome_averages.tsv** is higher than zero AND the median in **Bootstrapped_chromosome_averages.tsv,** count how many statuses for this cancer type in **Bootstrapped_chromosome_averages.tsv** are higher than the status in **Chromosome_averages.tsv,** and divide this number by 5000. If the status in **Chromosome_averages.tsv** is lower than zero AND the median in **Bootstrapped_chromosome_averages.tsv,** count how many statuses for this cancer type in **Bootstrapped_chromosome_averages.tsv** are lower than the status in **Chromosome_averages.tsv,** divide this number by 5000 and add minus to indicate chromosome loss. Ignore other values (leave cells empty). Save the file as **Chromosome_Pvalues_cohorts.tsv**
15. For each cancer type, apply Benjamini–Hochberg procedure with FDR=5% to P-values in **Arm_Pvalues_cohorts.tsv** and replace those which pass with **DAG** (Driver arm gain) or **DAL** (Driver arm loss) if with minus. Make other cells empty and save the results as **Arm_drivers_FDR5_cohorts.tsv**
16. For each cancer type, apply Benjamini–Hochberg procedure with FDR=5% to P-values in **Chromosome_Pvalues_cohorts.tsv** and replace those which pass with **DCG** (Driver chromosome gain) or **DCL** (Driver chromosome loss) if with minus. Make other cells empty and save the results as **Chromosome_drivers_FDR5_cohorts.tsv**
17. Using the file **Primary_whitelisted_chromosomes.tsv** and referring to the file **Chromosome_drivers_FDR5_cohorts.tsv**, classify alterations according to this table:

| **Status in a given patient** Primary_whitelisted_chromosomes.tsv | | **Empty cell** | **-1** | **0** | **1** |
|---|---|---|---|---|---|
| **Average alteration status of a given chromosome in the same cancer type** Chromosome_drivers_FDR5_cohorts.tsv | **DCL** | Empty cell | **DCL** | Empty cell | Empty cell |
| | **DCG** | Empty cell | Empty cell | Empty cell | **DCG** |
| | Empty cell | Empty cell | Empty cell | Empty cell | Empty cell |

Calculate the total number of DCLs, DCGs and TCDs (total chromosome drivers). Remove patients with 0 TCDs. Save the results as **Chromosome_drivers_FDR5.tsv**

18. Use the file **Primary_whitelisted_arms.tsv** and referring to the files **Arm_drivers_FDR5_cohorts.tsv** and **Chromosome_drivers_FDR5.tsv**, classify alterations according to this table:

<table>
<tr><th colspan="2">Status in a given patient<br>Primary_whitelisted_arms.tsv</th><th>Empty cell</th><th>-1</th><th>0</th><th>1</th></tr>
<tr><td rowspan="3">Average alteration status of a given arm in the same cancer type<br>Arm_drivers_FDR5_cohorts.tsv</td><td>DAL</td><td>Empty cell</td><td>DAL*</td><td>Empty cell</td><td>Empty cell</td></tr>
<tr><td>DAG</td><td>Empty cell</td><td>Empty cell</td><td>Empty cell</td><td>DAG**</td></tr>
<tr><td>Empty cell</td><td>Empty cell</td><td>Empty cell</td><td>Empty cell</td><td>Empty cell</td></tr>
<tr><td colspan="6">*If in Chromosome_drivers_FDR5.tsv the status of the corresponding chromosome for the same patient is DCL, then make empty cell</td></tr>
<tr><td colspan="6">**If in Chromosome_drivers_FDR5.tsv the status of the corresponding chromosome for the same patient is DCG, then make empty cell</td></tr>
</table>

Calculate the total number of DALs, DAGs and TADs (total arm drivers). Remove patients with 0 TADs. Save the results as **Arm_drivers_FDR5.tsv**
