## Supplementary material for "Comprehensive patient-level classification and quantification of driver events in TCGA PanCanAtlas cohorts": S6 Files: step_4.py

```
# -*- coding: utf-8 -*-
import os
import shutil

import pandas as pd
import argparse
import gzip

from utils.utils import download, check_hash


def step4(save_folder: str = 'data'):

    print('Step 4', end='\t')

    if not os.path.isdir(save_folder):
        os.makedirs(save_folder)

    output_file_path = os.path.join(save_folder, 'TCGA_mastercalls.abs_tables_JSedit.fixed.txt')

    # download
    if not os.path.isfile(output_file_path):
        # download file
        download_link = 'https://api.gdc.cancer.gov/data/4f277128-f793-4354-a13d-30cc7fe9f6b5'
        download(download_link, output_file_path)

    # Check hashsum
    hash_pass = check_hash(output_file_path)
    if not hash_pass:
        raise Exception(f'Hash sum validation for {output_file_path} was not passed, check the file or download it again')

    print('OK')

if __name__ == '__main__':
    parser = argparse.ArgumentParser(description='This script downloads TCGA_mastercalls.abs_tables_JSedit.fixed.txt file')
    parser.add_argument('-s', '--save_folder', type=str, help='full path to folder to save downloaded files', default='data')
    args = parser.parse_args()

    step4(args.save_folder)
```
