## Supplementary material for "Comprehensive patient-level classification and quantification of driver events in TCGA PanCanAtlas cohorts": S6 Files: step_5.py

```
# -*- coding: utf-8 -*-

import argparse
import os
import pandas as pd

from utils.utils import check_filesize


def get_sample_code(tcga_id):
    return tcga_id.split('-')[3][:2]


def get_index(header, column_name):
    header = header.split('\t')
    for i in range(len(header)):
        if header[i] == column_name:
            return i
    return -1


def step5(input_dir: str = 'data', output_folder_path: str = 'data'):

    print('Step 5', end='\t')

    input_path = os.path.join(input_dir, 'TCGA_mastercalls.abs_tables_JSedit.fixed.txt')
    banned_samples_path = os.path.join(input_dir, 'merged_sample_quality_annotations_do_not_use.tsv')
    output_file_path = os.path.join(output_folder_path, 'TCGA_mastercalls.abs_tables_JSedit.fixed_primary_whitelisted.tsv')

    if not os.path.isdir(output_folder_path):
        os.makedirs(output_folder_path)

    if not os.path.isfile(output_file_path):
        # get banned samples set
        banned_samples = set(pd.read_csv(banned_samples_path,
                                         sep='\t',
                                         usecols=['aliquot_barcode'])['aliquot_barcode'])

        with open(input_path, 'r') as input_file, open(output_file_path, 'w') as out_file:

            header = input_file.readline()
            out_file.write(header)

            tumor_sample_barcode_ix = get_index(header, 'array')
            cancer_fraction_ix = get_index(header, 'Cancer DNA fraction')
            subclonal_genome_fraction_ix = get_index(header, 'Subclonal genome fraction')
            content_line = input_file.readline()
            all_barcodes = set()
            selected_barcodes = set()
            while content_line:
                content_line = content_line.split('\t')
                full_barcode = content_line[tumor_sample_barcode_ix]
                all_barcodes.add(full_barcode)
                first_condition = get_sample_code(full_barcode) in ['01', '03', '09']
                if content_line[cancer_fraction_ix].strip() != '':
                    second_condition = float(content_line[cancer_fraction_ix].strip()) >= 0.5
                else:
                    second_condition = False
                if content_line[subclonal_genome_fraction_ix].strip() != '':
                    third_condition = float(content_line[subclonal_genome_fraction_ix].strip()) <= 0.5
                else:
                    third_condition = False

                fourth_condition = full_barcode not in banned_samples
                if first_condition and second_condition and third_condition and fourth_condition:
                    out_file.write('\t'.join(content_line))
                    selected_barcodes.add(full_barcode)

                content_line = input_file.readline()
        print('There were {} samples'.format(len(all_barcodes)))
        print('{} samples were selected'.format(len(selected_barcodes)))

    # Check for filesize
    size_pass = check_filesize(output_file_path)
    if not size_pass:
        raise Exception(f'file: {output_file_path} has wrong size, please check input file and do this step again')

    print('OK')

if __name__ == '__main__':

    parser = argparse.ArgumentParser(description='This script takes TCGA_mastercalls.abs_tables_JSedit.fixed.txt,' + '\n' +
                                                 'keeps only 01, 03, 09 samples (column Tumor_Sample_Barcode),' +
                                                 '\n' + 'keeps Cancer DNA fraction >= 0.5' +
                                                 '\n' + 'keeps Subclonal genome fraction <= 0.5'
                                                 '\n' + 'deletes all the aliquotes which mentioned in file: ' +
                                                 'merged_sample_quality_annotations_do_not_use.tsv (column aliquot_barcode)'
                                                 'if output folder does not exist, script will create it.')
    parser.add_argument('-i', '--input_dir', type=str, help='full path to input folder', default='data')
    parser.add_argument('-o', '--output_folder', type=str, help='full path to output folder', default='data')
    args = parser.parse_args()

    step5(args.input_dir, args.output_folder)
```
