## Supplementary material for "Comprehensive patient-level classification and quantification of driver events in TCGA PanCanAtlas cohorts": S6 Files: step_6.py

```
# -*- coding: utf-8 -*-

from requests import get
from utils.utils import download, check_hash

import pandas as pd
import os
import argparse


def step6(save_folder: str = 'data'):

    print('Step 6', end='\t')

    if not os.path.isdir(save_folder):
        os.makedirs(save_folder)

    output_file_path = os.path.join(save_folder, 'PANCAN_ArmCallsAndAneuploidyScore_092817.txt')

    if not os.path.isfile(output_file_path):
        # Downloading
        download_link = 'https://api.gdc.cancer.gov/data/4c35f34f-b0f3-4891-8794-4840dd748aad'
        download(link=download_link, destination=output_file_path)

    # Hash sum check
    hash_pass = check_hash(output_file_path)
    if not hash_pass:
        raise Exception(f'Hash sum validation for {output_file_path} was not passed, check the file or download it again')

    print('OK')

if __name__ == '__main__':

    parser = argparse.ArgumentParser(description='This script downloads PANCAN_ArmCallsAndAneuploidyScore_092817.txt file ' + '\n' +
                                                 'and saves it to the data/ folder by default')
    parser.add_argument('-s', '--save_folder', type=str, help='full path to folder to save downloaded files', default='data')
    args = parser.parse_args()

    step6(args.save_folder)
```
