## Supplementary material for "Comprehensive patient-level classification and quantification of driver events in TCGA PanCanAtlas cohorts": S6 Files: step_7.py

```
# -*- coding: utf-8 -*-

import argparse
import os
import pandas as pd

from utils.utils import check_filesize


def get_sample_code(tcga_id):
    return tcga_id.split('-')[3][:2]


def get_index(header, column_name):
    header = header.split('\t')
    for i in range(len(header)):
        if header[i] == column_name:
            return i
    return -1


def step7(input_dir: str = 'data', output_folder_path: str = 'data'):

    print('Step 7', end='\t')

    input_path = os.path.join(input_dir, 'PANCAN_ArmCallsAndAneuploidyScore_092817.txt')
    allowed_samples_path = os.path.join(input_dir, 'TCGA_mastercalls.abs_tables_JSedit.fixed_primary_whitelisted.tsv')
    output_file_path = os.path.join(output_folder_path, 'Primary_whitelisted_arms.tsv')

    if not os.path.isdir(output_folder_path):
        os.makedirs(output_folder_path)

    if not os.path.isfile(output_file_path):

        PANCAN_df = pd.read_csv(input_path, sep='\t')

        allowed_patients = set(list(pd.read_csv(allowed_samples_path, sep='\t', usecols=['array'])['array']))

        PANCAN_df = PANCAN_df[PANCAN_df['Sample'].isin(allowed_patients)]

        # rename chromosomes
        excluded_columns = ['13', '14', '15', '21', '22']
        for chr_number in excluded_columns:
            full_colname = str(chr_number) + ' ({}q)'.format(chr_number)
            PANCAN_df = PANCAN_df.rename(columns={full_colname: str(chr_number)+'q'})

        PANCAN_df.to_csv(output_file_path, sep='\t', header=True, index=None)

    # Check for filesize
    size_pass = check_filesize(output_file_path)
    if not size_pass:
        raise Exception(f'file: {output_file_path} has wrong size, please check input file and do this step again')

    print('OK')

if __name__ == '__main__':
    parser = argparse.ArgumentParser(description='This script takes PANCAN_ArmCallsAndAneuploidyScore_092817.txt,' + '\n' +
                                                 'keeps only patients which are presented in ' + '\n' +
                                                 'TCGA_mastercalls.abs_tables_JSedit.fixed_primary_whitelisted.tsv' +
                                                 '\n' + 'if output folder does not exist, script will create it.')
    parser.add_argument('-i', '--input_dir', type=str, help='full path to input folder', default='data')
    parser.add_argument('-o', '--output_folder', type=str, help='full path to output folder', default='data')

    args = parser.parse_args()

    step7(args.input_dir, args.output_folder)
```
