## Supplementary material for "Comprehensive patient-level classification and quantification of driver events in TCGA PanCanAtlas cohorts": S6 Files: step_8.py

```
# -*- coding: utf-8 -*-

import argparse
import os
import pandas as pd
import numpy as np

from utils.utils import check_filesize


def chr_status_creation(args):
    x = args[0]
    y = args[1]
    if np.isnan(x):
        return x
    elif np.isnan(y):
        return y
    elif x == 0:
        return 0
    elif y == 0:
        return 0
    elif x * y < 0:
        return 0
    elif x==1 and y==1 :
        return 1
    elif x==-1 and y==-1:
        return -1


def step8(input_dir: str = 'data', output_folder_path: str = 'data'):

    print('Step 8', end='\t')

    input_path = os.path.join(input_dir, 'Primary_whitelisted_arms.tsv')
    output_file_path = os.path.join(output_folder_path,
                                    'Primary_whitelisted_chromosomes.tsv')

    if not os.path.isdir(output_folder_path):
        os.makedirs(output_folder_path)

    if not os.path.isfile(output_file_path):

        PANCAN_df = pd.read_csv(input_path, sep='\t')

        excluded_columns = ['13', '14', '15', '21', '22']
        for chr_number in range(1, 23):
            if str(chr_number) in excluded_columns:
                full_colname = '{}q'.format(chr_number)
                PANCAN_df = PANCAN_df.rename(columns={full_colname: str(chr_number)})
            else:
                PANCAN_df['{}'.format(chr_number)] = \
                    PANCAN_df[['{}p'.format(chr_number), '{}q'.format(chr_number)]].apply(chr_status_creation, axis=1)
                PANCAN_df.drop(columns = ['{}p'.format(chr_number), '{}q'.format(chr_number)], inplace=True)
        # change to right order
        first_columns = list(PANCAN_df.columns)[:3]
        PANCAN_df = PANCAN_df[first_columns + [str(number) for number in range(1, 23)]]

        PANCAN_df.to_csv(output_file_path, sep='\t', header=True, index=None)

    # Check for filesize
    size_pass = check_filesize(output_file_path)
    if not size_pass:
        raise Exception(f'file: {output_file_path} has wrong size, please check input file and do this step again')

    print('OK')

if __name__ == '__main__':

    parser = argparse.ArgumentParser(description='This script takes ' +
                                                 'Primary_whitelisted_arms.tsv,'
                                                 + '\n' + 'and calculates chromosome status' +
                                                 '\n' + 'if output folder does not exist, script will create it.')
    parser.add_argument('-i', '--input_dir', type=str, help='full path to input folder', default='data')
    parser.add_argument('-o', '--output_folder', type=str, help='full path to output folder', default='data')

    args = parser.parse_args()

    step8(args.input_dir, args.output_folder)
```
