## Supplementary material for "Comprehensive patient-level classification and quantification of driver events in TCGA PanCanAtlas cohorts": S6 Files: step_9.py

```
# -*- coding: utf-8 -*-

import argparse
import os
import pandas as pd

from utils.utils import check_filesize


def step9(input_dir: str = 'data', output_folder_path: str = 'data'):

    print('Step 9', end='\t')

    input_path = os.path.join(input_dir, 'Primary_whitelisted_arms.tsv')
    output_file_path = os.path.join(output_folder_path, 'Arm_averages.tsv')

    if not os.path.isdir(output_folder_path):
        os.makedirs(output_folder_path)

    if not os.path.isfile(output_file_path):

        PANCAN_df = pd.read_csv(input_path, sep='\t')

        Aneuploidy_averages_df = PANCAN_df.groupby('Type').mean().drop(['Aneuploidy Score'], axis=1)

        Aneuploidy_averages_df = Aneuploidy_averages_df.reset_index()
        Aneuploidy_averages_df.to_csv(output_file_path, sep='\t', header=True, index=False)

    # Check for filesize
    size_pass = check_filesize(output_file_path)
    if not size_pass:
        raise Exception(f'file: {output_file_path} has wrong size, please check input file and do this step again')

    print('OK')

if __name__ == '__main__':

    parser = argparse.ArgumentParser(description='This script takes ' +
                                                 'Primary_whitelisted_arms.tsv,'
                                                 + '\n' + 'and calculates arm averages among cohorts' + '\n' +
                                                 'and save the result in Aneuploidy_averages.tsv file' +
                                                 '\n' + 'if output folder does not exist, script will create it.')
    parser.add_argument('-i', '--input_dir', type=str, help='full path to input folder', default='data')
    parser.add_argument('-o', '--output_folder', type=str, help='full path to output folder', default='data')

    args = parser.parse_args()

    step9(args.input_dir, args.output_folder)
```
