## Supplementary material for "Comprehensive patient-level classification and quantification of driver events in TCGA PanCanAtlas cohorts": S6 Files: step_11.py

```
# -*- coding: utf-8 -*-

import argparse
import os
import pandas as pd
import numpy as np
from tqdm import tqdm

from utils.utils import check_filesize


def find_bootstr_mean(cohort_id, coh2count, whole_df):
    to_sample_number = coh2count[cohort_id]
    row_ixs = np.random.choice(np.arange(whole_df.shape[0]), size=to_sample_number)
    # 2 because there are "Sample" and "Type" columns
    column_ixs = np.random.choice(np.arange(2, whole_df.shape[1]), size=to_sample_number)
    return whole_df.iloc[row_ixs, column_ixs].mean().mean()


def step11(input_dir: str = 'data', output_folder_path: str = 'data'):

    print('Step 11', end='\t')

    input_path = os.path.join(input_dir, 'Primary_whitelisted_arms.tsv')
    output_file_path = os.path.join(output_folder_path, 'Bootstrapped_arm_averages.tsv')

    if not os.path.isdir(output_folder_path):
        os.makedirs(output_folder_path)

    if not os.path.isfile(output_file_path):

        print('reading input data...')
        input_df = pd.read_csv(input_path, sep='\t')
        input_df.drop(columns=['Aneuploidy Score'], inplace=True)

        # let's find number of samples of each type
        temporary_df = input_df[['Sample', 'Type']].groupby('Type').count()
        cohort_counts_dict = dict(zip(list(temporary_df.index), temporary_df['Sample']))

        N_botstr = 10_000

        cohorts_list = list(cohort_counts_dict.keys())
        final_df = pd.DataFrame(columns=['Cancer_type'] + [str(x) for x in range(1, N_botstr+1)])
        final_df['Cancer_type'] = cohorts_list

        find_bootstr_mean_vect = np.vectorize(lambda x: find_bootstr_mean(x, cohort_counts_dict, input_df))

        print('generating bootstrapped averages...')
        for column_name in tqdm(range(1, N_botstr+1)):
            column_name = str(column_name)
            curr_bootstrapped = find_bootstr_mean_vect(cohorts_list)
            final_df[column_name] = curr_bootstrapped

        final_df['Median'] = final_df.iloc[:, 1:].median(axis=1)

        print('saving the results...')
        final_df.to_csv(output_file_path, sep='\t', header=True, index=False)

    # Check for filesize
    size_pass = check_filesize(output_file_path)
    if not size_pass:
        raise Exception(f'file: {output_file_path} has wrong size, please check input file and do this step again')

    print('OK')

if __name__ == '__main__':
    parser = argparse.ArgumentParser(description='This script takes ' +
                                                 'Primary_whitelisted_arms.tsv,'
                                                 + '\n' + 'and calculates bootstrapped arm averages among cohorts' + '\n' +
                                                 'and save the result in Bootstrapped_arm_averages.tsv file' +
                                                 '\n' + 'if output folder does not exist, script will create it.')
    parser.add_argument('-i', '--input_dir', type=str, help='full path to input folder', default='data')
    parser.add_argument('-o', '--output_folder', type=str, help='full path to output folder', default='data')

    args = parser.parse_args()

    step11(args.input_dir, args.output_folder)
```
