## Supplementary material for "Comprehensive patient-level classification and quantification of driver events in TCGA PanCanAtlas cohorts": S6 Files: step_13.py

```
# -*- coding: utf-8 -*-

import argparse
import os
import pandas as pd
import numpy as np

from utils.utils import check_filesize


def define_p_value(value, bottstr_row):
    # last value is median
    median = bottstr_row[-1]
    N = len(bottstr_row[:-1])
    if value > median:
        if value < 0: # value between median and 0
            return 'None'
        else:
            p_value = np.count_nonzero( bottstr_row[:-1] > value) * 2 / float(N)
    else: # value <= median
        if value > 0: # value between 0 and median
            return 'None'
        else:
            p_value = np.count_nonzero( bottstr_row[:-1] < value) * 2 / float(N)

    # for distinguish gain and loss
    if value > 0:
        return '{0:.4f}'.format(p_value)
    else:
        return '-{0:.4f}'.format(p_value)


def step13(input_dir: str = 'data', output_folder_path: str = 'data'):

    print('Step 13', end='\t')

    bootstr_averages_path = os.path.join(input_dir, 'Bootstrapped_arm_averages.tsv')
    averages_path = os.path.join(input_dir, 'Arm_averages.tsv')
    output_file_path = os.path.join(output_folder_path, 'Arm_Pvalues_cohorts.tsv')

    if not os.path.isdir(output_folder_path):
        os.makedirs(output_folder_path)

    if not os.path.isfile(output_file_path):

        bootstr_averages_df = pd.read_csv(bootstr_averages_path, sep='\t')
        averages_df = pd.read_csv(averages_path, sep='\t')

        p_values_df = pd.DataFrame(columns=averages_df.columns).rename(columns={'Type': 'Cancer_type'})
        p_values_df['Cancer_type'] = averages_df['Type']

        print('calculating p-values...')

        for row_ix, row in p_values_df.iterrows():
            define_p_value_vect = np.vectorize(lambda x:
                                               define_p_value(x, np.array(bootstr_averages_df.iloc[row_ix, 1:])))
            row[1:] = define_p_value_vect(averages_df.iloc[row_ix, 1:])

        p_values_df.to_csv(output_file_path, sep='\t', header=True, index=False)

    # Check for filesize
    size_pass = check_filesize(output_file_path)
    if not size_pass:
        raise Exception(f'file: {output_file_path} has wrong size, please check input file and do this step again')

    print('OK')

if __name__ == '__main__':
    parser = argparse.ArgumentParser(description='This script takes ' +
                                                 'Bootstrapped_arm_averages.tsv and Arm_averages.tsv,'
                                                 + '\n' + 'and calculates p-values among cohorts for each arm' + '\n' +
                                                 'and save the result in Arm_Pvalues_cohorts.tsv file' +
                                                 '\n' + 'if output folder does not exist, script will create it.')

    parser.add_argument('-i', '--input_dir', type=str, help='full path to input folder', default='data')
    parser.add_argument('-o', '--output_folder', type=str, help='full path to output folder', default='data')

    args = parser.parse_args()

    step13(args.input_dir, args.output_folder)
```
