## Supplementary material for "Comprehensive patient-level classification and quantification of driver events in TCGA PanCanAtlas cohorts": S6 Files: step_15.py

```
# -*- coding: utf-8 -*-

import argparse
import os
import pandas as pd
import numpy as np
from statsmodels.stats.multitest import fdrcorrection
import copy

from utils.utils import check_filesize

def step15(input_dir: str = 'data', output_folder_path: str = 'data'):

    print('Step 15', end='\t')

    p_vals_path = os.path.join(input_dir, 'Arm_Pvalues_cohorts.tsv')
    output_file_path = os.path.join(output_folder_path, 'Arm_drivers_FDR5_cohorts.tsv')

    if not os.path.isdir(output_folder_path):
        os.makedirs(output_folder_path)

    if not os.path.isfile(output_file_path):

        # read as str not to lose '-'
        p_vals_df = pd.read_csv(p_vals_path, sep='\t', dtype=str)

        # create dict
        col_names = list(p_vals_df.columns[1:])
        cohort_ids = list(p_vals_df['Cancer_type'])

        p_vals_dict = dict() # store p_val by column name, cohort_id

        bool_dict = dict() # store decisions whether to include or not

        pos_neg_dict = dict() # store sign

        for col_name in col_names:
            for cohort_id in cohort_ids:
                value = p_vals_df[p_vals_df['Cancer_type'] == cohort_id][col_name].iloc[0]
                if value == 'None':
                    # not to accept 'None' values
                    bool_dict[(col_name, cohort_id)] = False
                elif value[0] == '-':
                    p_vals_dict[(col_name, cohort_id)] = np.abs(float(value))
                    pos_neg_dict[(col_name, cohort_id)] = '-'
                else:
                    p_vals_dict[(col_name, cohort_id)] = float(value)
                    pos_neg_dict[(col_name, cohort_id)] = '+'

        # sort by value
        p_vals_dict = {k: v for k, v in sorted(p_vals_dict.items(), key=lambda item: item[1])}

        # do BH correction
        rejected, corrected_p_vals = fdrcorrection(list(p_vals_dict.values()), alpha=0.05)

        bool_dict.update( dict(zip(list(p_vals_dict.keys()), rejected)) )

        p_vals_df_copy = copy.deepcopy(p_vals_df)
        p_vals_df_copy.index = p_vals_df_copy['Cancer_type']

        for key in bool_dict:
            arm, cohort_id = key
            if bool_dict[key]:
                if pos_neg_dict[key] == '+':
                    p_vals_df_copy.loc[cohort_id, arm] = 'DAG'
                elif pos_neg_dict[key] == '-':
                    p_vals_df_copy.loc[cohort_id, arm] = 'DAL'
            else:
                p_vals_df_copy.loc[cohort_id, arm] = None

        p_vals_df_copy = p_vals_df_copy.drop(columns=['Cancer_type']).reset_index()

        p_vals_df_copy.to_csv(output_file_path, sep='\t', index=None)

    # Check for filesize
    size_pass = check_filesize(output_file_path)
    if not size_pass:
        raise Exception(f'file: {output_file_path} has wrong size, please check input file and do this step again')

    print('OK')

if __name__ == '__main__':
    parser = argparse.ArgumentParser(description='This script takes ' +
                                                 'Arm_Pvalues_cohorts.tsv, does BH correction on that'+ '\n' +
                                                 'and save the result in Arm_drivers_FDR5_cohorts.tsv file' +
                                                 '\n' + 'if output folder does not exist, script will create it.')

    parser.add_argument('-i', '--input_dir', type=str, help='full path to input folder', default='data')
    parser.add_argument('-o', '--output_folder', type=str, help='full path to output folder', default='data')

    args = parser.parse_args()

    step15(args.input_dir, args.output_folder)
```
