## Supplementary material for "Comprehensive patient-level classification and quantification of driver events in TCGA PanCanAtlas cohorts": S6 Files: step_18.py

```
# -*- coding: utf-8 -*-

import argparse
import os
import pandas as pd
import numpy as np
import copy
from tqdm import tqdm

from utils.utils import check_filesize


def define_alter_arm(drivers, primary, chrom):
    if drivers == 'DAL' and primary == -1. and chrom != 'DCL':
        return 'DAL'
    elif drivers == 'DAG' and primary == 1. and chrom != 'DCG':
        return 'DAG'
    else:
        return None


def step18(input_dir: str = 'data', output_folder_path: str = 'data'):

    print('Step 18', end='\t')

    primary_arms_path = os.path.join(input_dir, 'Primary_whitelisted_arms.tsv')
    chr_final_drivers_path = os.path.join(input_dir, 'Chromosome_drivers_FDR5.tsv')
    drivers_arms_path = os.path.join(input_dir, 'Arm_drivers_FDR5_cohorts.tsv')

    output_file_path = os.path.join(output_folder_path, 'Arm_drivers_FDR5.tsv')

    if not os.path.isdir(output_folder_path):
        os.makedirs(output_folder_path)

    if not os.path.isfile(output_file_path):

        primary_arms_df = pd.read_csv(primary_arms_path, sep='\t')
        chr_final_df = pd.read_csv(chr_final_drivers_path, sep='\t')
        drivers_arms_cohorts_df = pd.read_csv(drivers_arms_path, sep='\t')

        drivers_arms_cohorts_df.index = drivers_arms_cohorts_df['Cancer_type']
        drivers_arms_cohorts_df = drivers_arms_cohorts_df.drop(columns=['Cancer_type'])

        # transform chr array to fit the form
        new_columns = ['Sample'] + list('112233445566778899') + \
                      ['10', '10', '11', '11', '12', '12', '13', '14', '15', ] + \
                      ['16', '16', '17', '17', '18', '18', '19', '19', '20', '20', '21', '22']

        chr_final_df = chr_final_df[new_columns]
        primary_arms_df_copy = copy.deepcopy(primary_arms_df)

        print('this action could take 2-3 minutes')
        for ix, row in tqdm(primary_arms_df.iterrows()):
            cohort_id = row['Type']
            primary_arms_df_copy.iloc[ix, 3:] = list(map(lambda x, y, z: define_alter_arm(x, y, z),
                                                         drivers_arms_cohorts_df.loc[cohort_id],
                                                         primary_arms_df.iloc[ix, 3:],
                                                         chr_final_df.iloc[ix, 1:]))

        primary_arms_df_copy = primary_arms_df_copy.drop(columns=['Type', 'Aneuploidy Score'])

        primary_arms_df_copy['DAGs'] = (primary_arms_df_copy.iloc[:, 1:] == 'DAG').sum(axis=1)
        primary_arms_df_copy['DALs'] = (primary_arms_df_copy.iloc[:, 1:] == 'DAL').sum(axis=1)
        primary_arms_df_copy['TADs'] = primary_arms_df_copy['DAGs'] + primary_arms_df_copy['DALs']

        primary_arms_df_copy.to_csv(output_file_path, sep='\t', index=None)

    # Check for filesize
    size_pass = check_filesize(output_file_path)
    if not size_pass:
        raise Exception(f'file: {output_file_path} has wrong size, please check input file and do this step again')

    print('OK')

if __name__ == '__main__':
    parser = argparse.ArgumentParser(description='This script takes ' +
                                                 'Primary_whitelisted_arms.tsv, Chromosome_drivers_FDR5.tsv and ' + '\n' +
                                                 'Arm_drivers_FDR5_cohorts.tsv files,' + '\n' +
                                                 'defines driver events on arm level' + '\n' +
                                                 'and save the result in Arm_drivers_FDR5.tsv file' +
                                                 '\n' + 'if output folder does not exist, script will create it.')

    parser.add_argument('-i', '--input_dir', type=str, help='full path to input folder', default='data')
    parser.add_argument('-o', '--output_folder', type=str, help='full path to output folder', default='data')

    args = parser.parse_args()

    step18(args.input_dir, args.output_folder)
```
