## Supplementary material for "Comprehensive patient-level classification and quantification of driver events in TCGA PanCanAtlas cohorts": S7 Files: Instructions.txt

This script requires Python 3.7 and internet connection

Always open terminal from the folder SNADRIF

First, run the following command to install required libraries:
pip3 install -r requirements.txt

To run the whole pipeline: python3 run.py

To run a particular step: python3 run_one_step.py N
where N is the step number

All downloaded data, intermediate and final results are located in the folder "data"
