## Supplementary material for "Comprehensive patient-level classification and quantification of driver events in TCGA PanCanAtlas cohorts": S7 Files: run.py

```
import sys
import os
from step_2 import step2
from step_3 import step3
from step_4 import step4
from step_5 import step5
from step_6 import step6
from step_7 import step7
from step_8 import step8
from step_9 import step9
from step_10 import step10
from step_11 import step11
from step_12 import step12
from step_13 import step13
from step_14 import step14
from step_15 import step15
from step_16 import step16

print ("start analysis")
step2()
step3()
step4()
step5()
step6()
step7()
step8()
step9()
step10()
step11()
step12()
step13()
step14()
step15()
step16()

print ("completed")
```
