## Supplementary material for "Comprehensive patient-level classification and quantification of driver events in TCGA PanCanAtlas cohorts": S7 Files: run_one_step.py

```
import sys
import os
from step_2 import step2
from step_3 import step3
from step_4 import step4
from step_5 import step5
from step_6 import step6
from step_7 import step7
from step_8 import step8
from step_9 import step9
from step_10 import step10
from step_11 import step11
from step_12 import step12
from step_13 import step13
from step_14 import step14
from step_15 import step15
from step_16 import step16

print ("start analysis")
step = sys.argv[1]
steps = {2:step2,
	3:step3,
	4:step4,
	5:step5,
	6:step6,
	7:step7,
	8:step8,
	9:step9,
	10:step10,
	11:step11,
	12:step12,
	13:step13,
	14:step14,
	15:step15,
	16:step16,}
print (step)
steps[int(step)]()

print ("completed")
```
