## Supplementary material for "Comprehensive patient-level classification and quantification of driver events in TCGA PanCanAtlas cohorts": S7 Files: SNADRIF description.pdf

SNADRIF (SNA DRIver Finder) is a Python 3.7 software package that predicts cancer driver genes from the TCGA PanCanAtlas SNA data and classifies them into oncogenes and tumor suppressors. Driver prediction is based on calculating the ratio of functional (hyperactivating or inactivating) SNAs to passenger SNAs, whereas driver classification is based on calculating the ratio of hyperactivating SNAs to inactivating SNAs. Bootstrapping is used to calculate statistical significance and Benjamini–Hochberg procedure is used to keep false discovery rate under 5%. The pipeline can be executed fully automatically in less than two hours on a modern PC (Linux, Windows or MacOS).

This package has been developed by

Aleksey V. Belikov, Dr.rer.nat.
https://github.com/belikov-av
Laboratory of Innovative Medicine, School of Biological and Medical Physics, Moscow Institute of Physics and Technology
Roles: concept, pipeline, supervision

and

Danila V. Otnykov
https://github.com/dan-otn
Department of bioinformatics, School of Biological and Medical Physics, Moscow Institute of Physics and Technology
Roles: programming

**Pipeline overview**

First, the SNA file and the sample quality file are downloaded from https://gdc.cancer.gov/about-data/publications/PanCan-CellOfOrigin.
Next, low quality samples and metastatic samples are filtered out.
Then, SNAs are classified into likely hyperactivating, likely inactivating, likely passenger and unclear.

| **Variant_Classification** | **Possible effect** |
|---|---|
| De_novo_Start_InFrame | hyperactivating |
| De_novo_Start_OutOfFrame | passenger |
| Frame_Shift_Del | inactivating |
| Frame_Shift_Ins | inactivating |
| IGR | unclear |
| In_Frame_Del | hyperactivating |
| In_Frame_Ins | hyperactivating |
| Intron | unclear |
| Missense_Mutation | hyperactivating |
| Nonsense_Mutation | inactivating |
| Nonstop_Mutation | inactivating |

| | |
|---|---|
| RNA | unclear |
| Silent | passenger |
| Splice_Site | unclear |
| Targeted_Region | unclear |
| Translation_Start_Site | inactivating |
| 3'Flank | unclear |
| 3'UTR | unclear |
| 5'Flank | unclear |
| 5'UTR | unclear |

The number of SNAs of each type is counted for each gene and two indices are calculated:

a) NSEI - Nonsynonymous SNA Enrichment Index

$\boldsymbol{NSEI} = \textbf{Number of hyperactivating SNAs} + \textbf{Number of inactivating SNAs} + \textbf{1} / \textbf{Number of passenger SNAs} + \textbf{1}$

b) HISR - Hyperactivating to Inactivating SNA Ratio

$\boldsymbol{HISR} = \textbf{Number of hyperactivating SNAs} + \textbf{1} / \textbf{Number of inactivating SNAs} + \textbf{1}$

Next, the gene-patient matrix is constructed, recording various combinations of hyperactivating, inactivating, passenger and unclear SNAs. Bootstrapping from this matrix is then used to obtain the realistic distribution of NSEI under the null hypothesis. P-values are then determined and Benjamini–Hochberg procedure performed to keep FDR under 5%.

Finally, the resulting driver genes are classified into oncogenes (HISR>5) or tumor suppressors (HISR<5) based on empirically determined HISR threshold.

A detailed description of pipeline steps can be found in the file SNADRIF pipeline.pdf

Instructions for executing the code can be found in the file Instructions.txt
