## Supplementary material for "Comprehensive patient-level classification and quantification of driver events in TCGA PanCanAtlas cohorts": S7 Files: SNADRIF pipeline.pdf

1) Go to https://gdc.cancer.gov/about-data/publications/PanCan-CellOfOrigin
2) Download the file Analyte level annotations - merged_sample_quality_annotations.tsv
3) Using information in the column **aliquot_barcode**, delete all aliquots named TCGA-XX-XXXX-YYX-XXX-XXXX-XX, where YY – any number, except 01, 03 and 09 (see https://docs.gdc.cancer.gov/Encyclopedia/pages/TCGA_Barcode/ and table https://gdc.cancer.gov/resources-tcga-users/tcga-code-tables/sample-type-codes), and also, using information in the column **Do_not_use**, delete all aliquots with False value, and save the resulting file as **merged_sample_quality_annotations_do_not_use.tsv**
4) Download the file Public mutation annotation file - mc3.v0.2.8.PUBLIC.maf.gz, extract to **mc3.v0.2.8.PUBLIC.maf** and rename to **mc3.v0.2.8.PUBLIC.tsv**
5) Using information in the column **Tumor_Sample_Barcode**, delete from the file **mc3.v0.2.8.PUBLIC.tsv** all aliquots named TCGA-XX-XXXX-YYX-XXX-XXXX-XX, where YY – any number, except 01, 03 и 09; all aliquots that do not have the PASS value in the column **FILTER**; and all aliquots present in the file **merged_sample_quality_annotations_do_not_use.tsv,** and save the resulting file as **mc3.v0.2.8.PUBLIC_primary_whitelisted.tsv**
6) In the file **mc3.v0.2.8.PUBLIC_primary_whitelisted.tsv**, replace zeros in the column **Entrez_Gene_Id** with actual Entrez gene IDs, determined from the corresponding ENSEMBL gene IDs in the column **Gene** using external database (ftp://ftp.ncbi.nih.gov/gene/DATA/GENE_INFO/Mammalia/Homo_sapiens.gene_info.gz), and save the file as **mc3.v0.2.8.PUBLIC_primary_whitelisted_Entrez.tsv**
7) Using the file **mc3.v0.2.8.PUBLIC_primary_whitelisted_Entrez.tsv** classify all SNAs according to the column **Variant_Classification** and the following table:

| **Variant_Classification** | **Possible effect** |
|---|---|
| De_novo_Start_InFrame | hyperactivating |
| De_novo_Start_OutOfFrame | passenger |
| Frame_Shift_Del | inactivating |
| Frame_Shift_Ins | inactivating |
| IGR | unclear |
| In_Frame_Del | hyperactivating |
| In_Frame_Ins | hyperactivating |
| Intron | unclear |
| Missense_Mutation | hyperactivating |
| Nonsense_Mutation | inactivating |
| Nonstop_Mutation | inactivating |
| RNA | unclear |
| Silent | passenger |
| Splice_Site | unclear |
| Targeted_Region | unclear |
| Translation_Start_Site | inactivating |
| 3'Flank | unclear |
| 3'UTR | unclear |
| 5'Flank | unclear |
| 5'UTR | unclear |

Save the classification results as the file **SNA_classification_patients.tsv,** with columns **Tumor_Sample_Barcode, Hugo_Symbol, Entrez_Gene_Id, Gene, Number of hyperactivating SNAs, Number of inactivating SNAs, Number of SNAs with unclear role, Number of passenger SNAs.**

8) Using the file **SNA_classification_patients.tsv,** for each gene calculate the sum of all alterations in all patients. Remove genes that contain only SNAs with unclear role (noncoding genes) and save the results as **SNA_classification_genes.tsv** with columns **Hugo_Symbol, Entrez_Gene_Id, Gene, Number of hyperactivating SNAs, Number of inactivating SNAs, Number of SNAs with unclear role, Number of passenger SNAs.** Also remove noncoding genes from **SNA_classification_patients.tsv**
9) Calculate the "nonsynonymous SNA enrichment index" as

$$NSEI = \frac{\textbf{Number of hyperactivating SNAs} + \textbf{Number of inactivating SNAs} + 1}{\textbf{Number of passenger SNAs} + 1}$$

and the "hyperactivating to inactivating SNA ratio" as

$$HISR = \frac{\textbf{Number of hyperactivating SNAs} + 1}{\textbf{Number of inactivating SNAs} + 1}$$

using the file **SNA_classification_genes.tsv** and add it as additional columns to that file. Remove genes for which the sum of hyperactivating, inactivating and passenger SNAs is less than 10 (to ensure sufficient precision of NSEI and HISR calculation) and save it as **SNA_classification_genes_NSEI_HISR.tsv.**
10) Using the file **SNA_classification_patients.tsv,** construct the gene-patient matrix **SNA_matrix.tsv** with columns **Hugo_Symbol, Entrez_Gene_Id, Gene** and individual Tumor Sample Barcodes**,** encoding the Number of hyperactivating SNAs, Number of inactivating SNAs, Number of SNAs with unclear role and Number of passenger SNAs as one number separated by dots (e.g. 2.0.1.1). If data for a given gene is absent in a given patient, encode as 0.0.0.0
11) By drawing statuses randomly with replacement (bootstrapping) 10000 times from *any* cell of **SNA_matrix.tsv**, fill the table **SNA_matrix_bootstrapped.tsv** with columns **Iteration** and individual Tumor Sample Barcodes
12) Calculate the sums of statuses in **SNA_matrix_bootstrapped.tsv** for each iteration separately, calculate the corresponding NSEI and HISR indices (see step 9). Calculate null hypothesis P-value for each iteration as the number of NSEI values higher than a given iteration's NSEI value and divided by 10000. Save the results as **SNA_bootstrapped_NSEI_HISR.tsv** with columns **Iteration, Number of hyperactivating SNAs, Number of inactivating SNAs, Number of SNAs with unclear role, Number of passenger SNAs, NSEI, HISR, P value.** Plot a histogram with the distribution of P values (x axis – P values with 0.05 precision, y axis – the number of occurrences of a given value) under the null hypothesis.
13) Calculate P-value for each gene as the number of NSEI values in **SNA_bootstrapped_NSEI_HISR.tsv** higher than its NSEI value in **SNA_classification_genes_NSEI_HISR.tsv** and divided by 10000. Add it as an additional column to **SNA_classification_genes_NSEI_HISR.tsv** and save the file as **SNA_classification_genes_NSEI_HISR_Pvalues.tsv**

14) Apply Benjamini–Hochberg procedure with FDR(Q)=5% to P-values in **SNA_classification_genes_NSEI_HISR_Pvalues.tsv**, remove the genes that do not pass and save the rest as **SNA_driver_gene_list_FDR5.tsv**
15) Using the file **SNA_classification_genes_NSEI_HISR_Pvalues.tsv,** plot histograms with the distribution of P values (x axis – P values with 0.05 precision, y axis – the number of occurrences of a given value), NSEI values (x axis – NSEI values with 0.5 precision, y axis – the number of occurrences of a given value) and HISR values (x axis – HISR values with 0.5 precision, y axis – the number of occurrences of a given value).
16) Using the file **SNA_driver_gene_list_FDR5.tsv,** classify driver genes into oncogenes (OG; HISR>5) and tumor suppressor genes (TSG; HISR<5), add the results as an additional column and save the file as **SNA_driver_gene_list_FDR5_OG_TSG.tsv**
