## Supplementary material for "Comprehensive patient-level classification and quantification of driver events in TCGA PanCanAtlas cohorts": S7 Files: step_3.py

```
import pandas as pd
import os


def get_sample_code(tcga_id):
	return tcga_id.split('-')[3][:2]


def step3():
	print ('step 3 start')
	try:
		ff = 'data/merged_sample_quality_annotations_do_not_use.tsv'
		if os.path.exists(ff) and os.path.getsize(ff) > 400000:
			pass
		else:
			input_path = 'data/merged_sample_quality_annotations.tsv'
			output_folder_path = 'data/'

			output_file_path = os.path.join(output_folder_path, 'merged_sample_quality_annotations_do_not_use.tsv')

			sample_annot_df = pd.read_csv(input_path, sep='\t')

			first_condition = sample_annot_df['aliquot_barcode'].apply(get_sample_code).isin(['01', '03', '09'])
			second_condition = sample_annot_df['Do_not_use'] == True

			sample_annot_df[first_condition & second_condition].to_csv(output_file_path, header=True, index=False, sep='\t')
	except:
		print ('check your file in data/ to execute script')
```
