## Supplementary material for "Comprehensive patient-level classification and quantification of driver events in TCGA PanCanAtlas cohorts": S7 Files: step_4.py

```
import os
#from plumbum.cmd import gunzip, mv
import gzip
import shutil

import requests
import sys
import hashlib

def check_hash(file_path):
	d1 = {'mc3.v0.2.8.PUBLIC.maf.gz':('639ad8f8386e98dacc22e439188aa8fa','a3482bbdd4a27991fe1ba5a1cca11c2a9237ad56'),
	'merged_sample_quality_annotations.tsv':('05ddd2270fb1fb24fbdc2fe9bf7384e5','a70a070954813a905d3155fd1a0bab335c10ec88')}

	BUF_SIZE = 65536 
	with open(file_path, 'rb') as f:
		md5 = hashlib.md5()
		sha1 = hashlib.sha1()
		while True:
			data = f.read(BUF_SIZE)
			if not data:
				break
			md5.update(data)
			sha1.update(data)

	tf = file_path.split('/')[-1]
	if d1[tf] == (md5.hexdigest(),sha1.hexdigest()):
		return True
	else:
		return False
def download(link,file_name):
	with open(file_name, "wb") as f:
		#print "Downloading %s" % file_name
		response = requests.get(link, stream=True)
		total_length = response.headers.get('content-length')

		if total_length is None: # no content length header
			f.write(response.content)
		else:
			dl = 0
			total_length = int(total_length)
			for data in response.iter_content(chunk_size=4096):
				dl += len(data)
				f.write(data)
				done = int(50 * dl / total_length)
				sys.stdout.write("\r[%s%s]" % ('=' * done, ' ' * (50-done)) )    
				sys.stdout.flush()
	print("\n")

def step4():
	print ('step 4 start')

	### downloading

	input_path = 'data/mc3.v0.2.8.PUBLIC.maf.gz'
	url = 'https://api.gdc.cancer.gov/data/1c8cfe5f-e52d-41ba-94da-f15ea1337efc'
	output_path = 'data/mc3.v0.2.8.PUBLIC.tsv'
	try:
		
		if os.path.exists(input_path) and check_hash(input_path):
			if os.path.exists(output_path) and os.path.getsize(output_path) > 3700000000:
				pass
			else:
				with gzip.open(input_path, 'rb') as f_in:
					with open(output_path, 'wb') as f_out:
						shutil.copyfileobj(f_in, f_out)
		else:
			download(url, input_path)
			with gzip.open(input_path, 'rb') as f_in:
				with open(output_path, 'wb') as f_out:
					shutil.copyfileobj(f_in, f_out)
	except:
		print ('cant download file: '+ url)
```
