## Supplementary material for "Comprehensive patient-level classification and quantification of driver events in TCGA PanCanAtlas cohorts": S7 Files: step_5.py

```
import os
import pandas as pd

def get_sample_code(tcga_id):
	return tcga_id.split('-')[3][:2]


def get_index(header, column_name):
	header = header.split('\t')
	for i in range(len(header)):
		if header[i] == column_name:
			return i
	return -1


def step5():
	print ('step 5 start')
	try:
		ff = 'data/mc3.v0.2.8.PUBLIC_primary_whitelisted.tsv'
		if os.path.exists(ff) and os.path.getsize(ff) >2600000000:
			
			pass
		else:
			input_path = 'data/mc3.v0.2.8.PUBLIC.tsv'
			banned_samples_path = 'data/merged_sample_quality_annotations_do_not_use.tsv'
			output_folder_path = 'data'
			output_file_path = os.path.join(output_folder_path, 'mc3.v0.2.8.PUBLIC_primary_whitelisted.tsv')

			# get banned samples set
			banned_samples = set(pd.read_csv(banned_samples_path,
												sep='\t',
												usecols=['aliquot_barcode'])['aliquot_barcode'])

			with open(input_path, 'r') as maf_file, open(output_file_path, 'w') as out_maf_file:

				header = maf_file.readline()
				out_maf_file.write(header)

				tumor_sample_barcode_ix = get_index(header, 'Tumor_Sample_Barcode')
				filter_ix = get_index(header, 'FILTER')

				content_line = maf_file.readline()
				all_barcodes = set()
				selected_barcodes = set()
				while content_line:
					content_line = content_line.split('\t')
					full_barcode = content_line[tumor_sample_barcode_ix]
					all_barcodes.add(full_barcode)
					first_condition = get_sample_code(full_barcode) in ['01', '03', '09']
					second_condition = content_line[filter_ix] == 'PASS'
					third_condition = full_barcode not in banned_samples
					if not third_condition:
						print(full_barcode)
					if first_condition and second_condition and third_condition:
						out_maf_file.write('\t'.join(content_line))
						selected_barcodes.add(full_barcode)

					content_line = maf_file.readline()
	except:
		print ('check your file in data/ to execute script')
```
