## Supplementary material for "Comprehensive patient-level classification and quantification of driver events in TCGA PanCanAtlas cohorts": S7 Files: step_6.py

```
#STEP 6
import os

def step6():
	print ('step 6 start')
	try:
		ff = "data/mc3.v0.2.8.PUBLIC_primary_whitelisted_Entrez.tsv"
		if os.path.exists(ff) and os.path.getsize(ff) >2600917674 :
			pass
		else:
			annot = dict()
			a2_not_syn = dict()
			a1 =dict()
			a2 =dict()
			with open("data/Homo_sapiens.gene_info",'r') as an:
				#print(an.readline().split("\t"))
				for line in an:
					ens = line.split("\t")[5]
					if "nsembl" in ens:
						ens_id = ens.split("|")
						for ii in ens_id:
							if "nsembl" in ii:
								id = ii.split(":")[-1]
								annot[id] = line.split("\t")[1]
					synonims = line.split("\t")[4]
					synonims = synonims.split('|')
					for i in synonims:
						a2[i] = line.split("\t")[1]
					a2_not_syn[line.split("\t")[2]] = line.split("\t")[1]
					#a1[line.split("\t")[6]] = line.split("\t")[5]
					#a2[line.split("\t")[1]] = line.split("\t")[6]

			with open("data/mc3.v0.2.8.PUBLIC_primary_whitelisted.tsv",'r') as sna:
				with open("data/mc3.v0.2.8.PUBLIC_primary_whitelisted_Entrez.tsv",'w') as out:
					head = sna.readline()
					out.write(head)
					for line in sna:
						a =line.split("\t")
						if a[47] in annot:#ensembl
							a[1] = annot[a[47]]
							out.write("\t".join(a))
						else:
							if a[0] in a2_not_syn:#name
								a[1] = a2_not_syn[a[0]]
								out.write("\t".join(a))
							elif a[0] in a2:#syn name
								a[1] = a2[a[0]]
								out.write("\t".join(a))
							else:
								out.write("\t".join(a))
	except:
		print ('check your file in data/ to execute script')
```
