## Supplementary material for "Comprehensive patient-level classification and quantification of driver events in TCGA PanCanAtlas cohorts": S7 Files: step_7.py

```
#STEP 7
import pandas as pd
import os
import numpy as np
from collections import defaultdict,Counter
def step7():
	print ('step 7 start')
	try:
		ff = "data/SNA_classification_patients.tsv"
		if os.path.exists(ff) and os.path.getsize(ff) >100000000:
			pass
		else:
			d = defaultdict(list)
			possible_eff = {"3'Flank":"unclear",
				 'De_novo_Start_OutOfFrame':"passenger",
				 'De_novo_Start_InFrame':'hyperactivating',
				 "3'UTR":"unclear",
				 "5'Flank":"unclear",
				 "5'UTR":"unclear",
				 'Frame_Shift_Del':'inactivating',
				 'Frame_Shift_Ins':'inactivating',
				 'In_Frame_Del':'hyperactivating',
				 'In_Frame_Ins':'hyperactivating',
				 'IGR':"unclear",
				 'Targeted_Region':"unclear",
				 'Intron':"unclear",
				 'Missense_Mutation':'hyperactivating',
				 'Nonsense_Mutation':'inactivating',
				 'Nonstop_Mutation':'inactivating',
				 'RNA':"unclear",
				 'Silent':'passenger', 
				 'Splice_Site':"unclear",
				 'Translation_Start_Site':'inactivating'}
			with open("data/mc3.v0.2.8.PUBLIC_primary_whitelisted_Entrez.tsv",'r')as sna:
				sna.readline()
				for line in sna:
					d[(line.split("\t")[15],line.split("\t")[0],
					   line.split("\t")[1],line.split("\t")[47])].append(possible_eff[line.split("\t")[8]])
			with open("data/SNA_classification_patients.tsv",'w') as out:
				out.write("\t".join(['Tumor_Sample_Barcode', 'Hugo_Symbol', 'Entrez_Gene_Id', 'Gene',
									'Number of hyperactivating SNAs','Number of inactivating SNAs',
									 'Number of SNAs with unclear role','Number of passenger SNAs'])+"\n")
				for k in d:
					s = ''
					if 'hyperactivating' in Counter(d[k]).keys():
						s =s + str(Counter(d[k])['hyperactivating'])+"\t"
					else :
						s = s +str(0)+"\t"
					if 'inactivating' in Counter(d[k]).keys():
						s = s + str(Counter(d[k])['inactivating'])+"\t"
					else :
						s = s +str(0)+"\t"    
					if 'unclear' in Counter(d[k]).keys():
						s = s + str(Counter(d[k])["unclear"])+"\t"
					else :
						s = s +str(0)+"\t"
					if 'passenger' in Counter(d[k]).keys():
						s = s + str(Counter(d[k])['passenger'])+"\n"
					else :
						s = s +str(0)+"\n"
					out.write(k[0]+"\t"+k[1]+"\t"+k[2]+"\t"+k[3]+"\t"+s)
	except:
		print ('check your file in data/ to execute script')
```
