## Supplementary material for "Comprehensive patient-level classification and quantification of driver events in TCGA PanCanAtlas cohorts": S7 Files: step_8.py

```
#STEP 8
from collections import defaultdict,Counter
#from utils.utils import check_hash
import os
def step8():
	print ('step 8 start')
	try:
		ff = "data/SNA_classification_genes.tsv"
		if os.path.exists(ff) and os.path.getsize(ff) > 800000 :
			pass
		else:
			hyper = defaultdict(int)
			inact = defaultdict(int)
			unclear = defaultdict(int)
			passenger = defaultdict(int)
			with open("data/SNA_classification_patients.tsv",'r') as f:
				patients = []
				patients.append(f.readline())
				for line in f:
					patients.append(line)
					a = line.split('\t')
					key = "\t".join(a[1:4])
					hyper[key]+=int(a[4])
					inact[key]+=int(a[5])
					unclear[key]+=int(a[6])
					passenger[key]+=int(a[7])
			h = ['Hugo_Symbol', 'Entrez_Gene_Id', 'Gene',
									'Number of hyperactivating SNAs','Number of inactivating SNAs',
									 'Number of SNAs with unclear role','Number of passenger SNAs']
			h = "\t".join(h)
			with open("data/SNA_classification_genes.tsv",'w') as o:
				o.write(h+'\n')
				unclear_genes_list = set()
				for a in hyper.keys():
					if  str(hyper[a])!= '0' or  str(inact[a])!= '0' or  str(passenger[a])!= '0':
						o.write(a + "\t")
						o.write(str(hyper[a]) + "\t")
						o.write(str(inact[a]) + "\t")
						o.write(str(unclear[a]) + "\t")
						o.write(str(passenger[a]) + "\n")
					else:
						unclear_genes_list.add(a)### список генов, где только unclear
			with open("data/SNA_classification_patients.tsv",'w') as f:
				for line in patients:
					genee = line.split('\t')
					geneee = "\t".join(genee[1:4])
					if geneee not in unclear_genes_list:
						f.write(line)
	except:
		print ('check your file in data/ to execute script')
```
