## Supplementary material for "Comprehensive patient-level classification and quantification of driver events in TCGA PanCanAtlas cohorts": S7 Files: step_9.py

```
#STEP 9
import os
#from utils.utils import check_hash
def step9():
	print ('step 9 start')
	try:
		ff = "data/SNA_classification_genes_NSEI_HISR.tsv"
		if os.path.exists(ff) and os.path.getsize(ff) >1300000:
			pass
		else:
			with open("data/SNA_classification_genes.tsv",'r') as o:
				with open("data/SNA_classification_genes_NSEI_HISR.tsv",'w') as f:
					head = o.readline()[:-1]+"\t"+"NSEI" +"\t"+"HISR"+"\n"
					f.write(head)
					for line in o:
						a = line.split("\t")
						nsei = (int(a[-3]) + int(a[-4]) + 1)/(int(a[-1])+1)
						hisr = (int(a[-4]) + 1)/(int(a[-3])+1)
						if int(a[-1])+int(a[-3])+int(a[-4])>=10:
							f.write(line[:-1]+"\t"+str(nsei)+"\t"+str(hisr)+"\n")
	except:
		print ('check your file in data/ to execute script')
```
