## Supplementary material for "Comprehensive patient-level classification and quantification of driver events in TCGA PanCanAtlas cohorts": S7 Files: step_10.py

```
#STEP 10
from collections import defaultdict
from tqdm import tqdm
import os.path
def step10():
	print ('step 10 start')
	try:
		ff = "data/SNA_matrix.tsv"
		if os.path.exists(ff) and os.path.getsize(ff) >140000000:
			pass
		else:
			#making header\
			header = ""
			brcd = set()
			with open("data/SNA_classification_patients.tsv",'r') as f:
				f.readline()
				for line in f:
					brcd.add(line.split("\t")[0])

			header = "Hugo_Simbol"+"\t"+"Entrez_Gene_Id"+"\t"+"Gene"+"\t" + "\t".join(brcd) + "\n"
			with open("data/SNA_matrix.tsv",'w') as out:  #!!!!
				out.write(header)
			#making genes set

			genes = set()
			#d = defaultdict(str) # dict of genes : barcode
			with open("data/SNA_classification_patients.tsv",'r') as f:
				f.readline()
				for line in f:
					a = line.split("\t")
					key = a[1] +"\t"+a[2] +"\t"+a[3]
					#d[key] =
					genes.add(key)
			genes = list(genes) 
			g_d = {}
			with open("data/SNA_classification_patients.tsv",'r') as f:
					d = defaultdict(str)
					for line in  f:
						a = line.split("\t")
						key = a[1] +"\t"+a[2] +"\t"+a[3]
						if key not in g_d.keys():
							g_d[key] = defaultdict(str)
						else:
							if a[-4] + "." +a[-3] + "." +a[-2] + "." +a[-1][:-1] == '':
								g_d[key][a[0]]= '0.0.0.0'
							else:
								g_d[key][a[0]] = a[-4] + "." +a[-3] + "." +a[-2] + "." +a[-1][:-1]
			#print(g_d[list(g_d.keys())[1]])
			fl = 0
			for i in tqdm(genes):
				fl +=1
				#with open("data/SNA_classification_patients.tsv",'r') as f:
				#	d = defaultdict(str)
				#	for line in  f:
				#		a = line.split("\t")
				#		key = a[1] +"\t"+a[2] +"\t"+a[3]
				#		if i == key:
							#print (key,i)
				#			if a[-4] + "." +a[-3] + "." +a[-2] + "." +a[-1][:-1] == '':
				#				d[a[0]]= '0.0.0.0'
				#			else:
				#				d[a[0]] = a[-4] + "." +a[-3] + "." +a[-2] + "." +a[-1][:-1]
				#print (genes)
				with open("data/SNA_matrix.tsv",'a') as out:  #!!!!
					out.write(i)
					for k in brcd:
						if g_d[i][k] == '':
							out.write("\t"+"0.0.0.0")
						else:
							out.write("\t"+g_d[i][k])
					out.write("\n")
	except:
		print ('check your file in data/ to execute script')
```
