## Supplementary material for "Comprehensive patient-level classification and quantification of driver events in TCGA PanCanAtlas cohorts": S7 Files: step_11.py

```
#STEP 11
import random
import os
from collections import defaultdict
import numpy
from scipy import stats
from tqdm import tqdm

from collections import defaultdict
def step11():
	print ('step 11 start')
	try:
		ff = "data/SNA_matrix_bootstrapped.tsv"
		if os.path.exists(ff) and os.path.getsize(ff)> 5*(10**8):
			pass
		else:
			patients = set()
			mut = defaultdict(int)
			with open("data/SNA_matrix.tsv",'r') as f:
				patients = f.readline().split("\t")[3:]
				for line in f:
					line = line[:-1]
					for i in line.split("\t")[3:]:
						mut[i]+=1
			with open("for_bootstrap",'w') as ff:
				for i in mut.keys():
					ff.write(i+"\t"+str(mut[i])+"\n")

			mutat = []
			p = []
			with open("for_bootstrap",'r') as f:
				for line in f:
					a = line.split("\t")
					if '.' in a[0]:
						mutat.append(a[0])
						p.append(int(a[1]))

			mutat = []
			p = []
			with open("for_bootstrap",'r') as f:
				for line in f:
					a = line.split("\t")
					if '.' in a[0]:
						mutat.append(a[0])
						p.append(int(a[1]))

			w = sum(p)
			new_p = list(map(lambda x :x/w,p))
			new_mut = []
			d={}
			k = 0
			for i in mutat:
				k+=1
				new_mut.append(k)
				d[k]=i


			with open("data/SNA_matrix_bootstrapped.tsv",'w') as f:
				f.write("Iteration"+"\t"+"\t".join(list(patients))+"\n")
				a = stats.rv_discrete( values=(new_mut,new_p))

				for i in tqdm(range(1,10001)):
					f.write(str(i))
					for i in range(len(patients)):
						f.write("\t"+str(d[a.rvs()]))
						
					f.write("\n")
	except:
		print ('check your file in data/ to execute script')
	try:
		os.remove('for_bootstrap')
	except:
		pass
```
