## Supplementary material for "Comprehensive patient-level classification and quantification of driver events in TCGA PanCanAtlas cohorts": S7 Files: step_12.py

```
#STEP 12
import os
import numpy as np
import matplotlib.pyplot as plt
def step12():
	print ('step 12 start')
	try:
		ff = "data/SNA_bootstrapped_NSEI_HISR.tsv"
		if os.path.exists(ff)and os.path.getsize(ff) > 300000:
			pass
		else:
			#SNA_matrix_bootstrapped.tsv  second row is empty
			header = "Iteration"+"\t"+ 'Number of hyperactivating SNAs' +"\t"+'Number of inactivating SNAs'+"\t"+'Number of SNAs with unclear role'+"\t"+'Number of passenger SNAs'+"\t"+"NSEI"+"\t"+"HISR"+"\n"
			with open("data/SNA_matrix_bootstrapped.tsv",'r') as f:
				nsei_set = []
				nsei_all = []
				with open("data/SNA_bootstrapped_NSEI_HISR.tsv",'w') as out:
					out.write(header)
					#nsei_set.append(header)
					k = 0
					for line in f:# number of iterations
						k+=1
						if k >2:
							out.write(line.split("\t")[0]+"\t")
							a = line.split("\t")[1:]
							h = 0
							i = 0
							u = 0
							p = 0
							for b in a: # b is muts for every patient
								h+=int(b.split(".")[0])
								i+=int(b.split(".")[1])
								u+=int(b.split(".")[2])
								p+=int(b.split(".")[3])
							out.write("\t".join([str(h),str(i),str(u),str(p)])+"\t")
							nsei = (h+i+1)/(p+1)
							hisr = (h+1)/(i+1)
							out.write("\t".join([str(nsei),str(hisr)])+"\n")
							nsei_all.append(nsei)
							nsei_set.append(line.split("\t")[0]+"\t"+"\t".join([str(h),str(i),str(u),str(p)])+"\t"+"\t".join([str(nsei),str(hisr)]))
			with open("data/SNA_bootstrapped_NSEI_HISR.tsv",'w') as out:
				out.write(header[:-1]+"\t"+"p value"+"\n")
				all_pval = []
				for line in nsei_set:
					out.write(line)
					p_val = 0
					nsei_curr = float(line.split("\t")[-2])
					for n in nsei_all:
						if n>nsei_curr:
							p_val+=1
					out.write("\t"+str(p_val/10000)+"\n")
					all_pval.append(p_val/10000)
			np_nsei = np.array(all_pval)
			#print ((max(np_hist)-min(np_hist))/0.1)
			hist,bin_edges = np.histogram(np_nsei,bins = int(round((max(np_nsei)-min(np_nsei))/0.05)))

			#PLOTTING 1
			plt.figure(figsize=[16,12])
			plt.bar(bin_edges[:-1], hist, width = 0.05, color='#0504aa',alpha=0.7)
			plt.xlim(0,1)
			plt.grid(axis='y', alpha=0.75)
			plt.xlabel('P values',fontsize=15)
			plt.ylabel('Frequency',fontsize=15)
			plt.xticks(fontsize=15)
			plt.yticks(fontsize=15)
			plt.title('P values Histogram',fontsize=15)
			#plt.axvline(x=hisr_med,color='r',linestyle='--')
			#plt.text(100,13,'HISR median = {}'.format(hisr_med), fontsize=16 )
			plt.savefig("data/histogram_p_values_bootstrap.pdf",dpi = 600)
	except:
		print ('check your file in data/ to execute script')
```
