## Supplementary material for "Comprehensive patient-level classification and quantification of driver events in TCGA PanCanAtlas cohorts": S7 Files: step_13.py

```
#STEP 13
import os
def step13():
	print ('step 13 start')
	try:
		ff = "data/SNA_classification_genes_NSEI_HISR_Pvalues.tsv"
		if os.path.exists(ff) and os.path.getsize(ff) > 300000:
			pass
		else:
			#make nsei sorted list from bootstrap
			nsei = []
			with open("data/SNA_bootstrapped_NSEI_HISR.tsv",'r') as f:
				f.readline()
				for line in f:
					nsei.append(float(line.split()[-3]))
			nsei = sorted(nsei)
			with open("data/SNA_classification_genes_NSEI_HISR.tsv",'r') as f:
				with open("data/SNA_classification_genes_NSEI_HISR_Pvalues.tsv",'w') as out:
					header = f.readline().split("\t")
					header[-1] = "HISR"
					header.append("P-value NSEI")
					header = "\t".join(header)+"\n"
					out.write(header)
					for line in f:
						a = line.split("\t")
					
						nsei_gene = float(a[-2])
						p = 0
						for n in nsei:
							if n > nsei_gene:
								p+=1
						p = p/10000
						a[-1] = float(a[-1])
						a[-1] = str(a[-1])
						a.append(str(p)+"\n")
						out.write("\t".join(a))
	except:
		print ('check your file in data/ to execute script')
```
