## Supplementary material for "Comprehensive patient-level classification and quantification of driver events in TCGA PanCanAtlas cohorts": S7 Files: step_14.py

```
#STEP 14
# https://github.com/puolival/multipy is used
#from statsmodels.stats.tests.test_multi import fdrcorrection
from multipy.fdr import lsu
from multipy.data import neuhaus
import numpy as np
import os
def step14():
	print ('step 14 start')
	try:
		ff = "data/SNA_driver_gene_list_FDR5.tsv"
		if os.path.exists(ff) and os.path.getsize(ff) > 1024:
			pass
		else:
			p = []
			genes = []
			with open("data/SNA_classification_genes_NSEI_HISR_Pvalues.tsv",'r') as f:
				f.readline()
				for line in f:
					genes.append(line.split("\t")[0])
					p.append(float(line.split("\t")[-1]))
			p = np.array(p)
			#p_fdr = fdrcorrection(p, alpha=0.05)

			p1 = lsu(p, q=0.05)
			true = []
			for i in range (len(p)):
				if p1[i] == True:
					true.append(genes[i])

			with open("data/SNA_classification_genes_NSEI_HISR_Pvalues.tsv",'r') as f:
				with open("data/SNA_driver_gene_list_FDR5.tsv",'w') as out:
					out.write(f.readline())
					for line in f:
						if line.split("\t")[0] in true:
							out.write(line)
	except:
		print ('check your file in data/ to execute script')
```
