## Supplementary material for "Comprehensive patient-level classification and quantification of driver events in TCGA PanCanAtlas cohorts": S7 Files: step_15.py

```
#STEP 15
import numpy as np
import matplotlib.pyplot as plt
import os
def step15():
	print ('step 15 start')
	try:
		ff = "data/histogram_fdr.pdf"
		if os.path.exists(ff) and os.path.getsize(ff)>100:
			pass
		else:
			hisr = []
			nsei = []
			pvalues = []
			with open("data/SNA_classification_genes_NSEI_HISR_Pvalues.tsv",'r') as f:
				f.readline()
				for line in f:
					hisr.append(float(line.split("\t")[-2]))
					nsei.append(float(line.split("\t")[-3]))
					pvalues.append(float(line.split("\t")[-1]))
			hisr_med = np.median (hisr)

			np_hist = np.array(hisr)
			#print ((max(np_hist)-min(np_hist))/0.1)
			hist,bin_edges = np.histogram(np_hist,bins = int(round((max(np_hist)-min(np_hist))/0.5)))

			#PLOTTING 1
			plt.figure(figsize=[16,12])
			plt.bar(bin_edges[:-1], hist, width = 0.5, color='#0504aa',alpha=0.7)
			plt.xlim(min(bin_edges)-5, max(bin_edges)+5)
			plt.grid(axis='y', alpha=0.75)
			plt.xlabel('HISR values',fontsize=15)
			plt.ylabel('Frequency',fontsize=15)
			plt.xticks(fontsize=15)
			plt.yticks(fontsize=15)
			plt.title('HISR Histogram',fontsize=15)
			#plt.axvline(x=hisr_med,color='r',linestyle='--')
			#plt.text(100,13,'HISR median = {}'.format(hisr_med), fontsize=16 )
			plt.savefig("data/histogram_HISR.pdf",dpi = 600)
			
			np_hist = np.array(nsei)
			#print ((max(np_hist)-min(np_hist))/0.1)
			hist,bin_edges = np.histogram(np_hist,bins = int(round((max(np_hist)-min(np_hist))/0.5)))

			#PLOTTING 1
			plt.figure(figsize=[16,12])
			plt.bar(bin_edges[:-1], hist, width = 0.5, color='#0504aa',alpha=0.7)
			plt.xlim(min(bin_edges)-5, max(bin_edges)+5)
			plt.grid(axis='y', alpha=0.75)
			plt.xlabel('NSEI values',fontsize=15)
			plt.ylabel('Frequency',fontsize=15)
			plt.xticks(fontsize=15)
			plt.yticks(fontsize=15)
			plt.title('NSEI Histogram',fontsize=15)
			#plt.axvline(x=hisr_med,color='r',linestyle='--')
			#plt.text(100,13,'NSEI median = {}'.format(hisr_med), fontsize=16 )
			plt.savefig("data/histogram_NSEI.pdf",dpi = 600)
			
			np_hist = np.array(pvalues)
			#print ((max(np_hist)-min(np_hist))/0.1)
			hist,bin_edges = np.histogram(np_hist,bins = int(round((max(np_hist)-min(np_hist))/0.05)))

			#PLOTTING 1
			plt.figure(figsize=[16,12])
			plt.bar(bin_edges[:-1], hist, width = 0.05, color='#0504aa',alpha=0.7)
			plt.xlim(0, 1)
			plt.grid(axis='y', alpha=0.75)
			plt.xlabel('P values',fontsize=15)
			plt.ylabel('Frequency',fontsize=15)
			plt.xticks(fontsize=15)
			plt.yticks(fontsize=15)
			plt.title('P values Histogram',fontsize=15)
			#plt.axvline(x=hisr_med,color='r',linestyle='--')
			#plt.text(100,13,'HISR median = {}'.format(hisr_med), fontsize=16 )
			plt.savefig("data/histogram_P_values.pdf",dpi = 600)
	except:
		print ('check your file in data/ to execute script')
```
