## Supplementary material for "Comprehensive patient-level classification and quantification of driver events in TCGA PanCanAtlas cohorts": S7 Files: step_16.py

```
#STEP 16
import numpy as np
import os
def step16():
	print ('step 16 start')
	try:
		ff = "data/SNA_driver_gene_list_FDR5_OG_TSG.tsv"
		if os.path.exists(ff) and os.path.getsize(ff)> 1024:
			pass
		else:
			hisr = []
			with open("data/SNA_driver_gene_list_FDR5.tsv",'r') as f:
				f.readline()
				for line in f:
					hisr.append(float(line.split("\t")[-2]))
			hisr_med = np.median (hisr)
			with open("data/SNA_driver_gene_list_FDR5.tsv",'r') as f:
				with open("data/SNA_driver_gene_list_FDR5_OG_TSG.tsv",'w') as out:
					out.write(f.readline()[:-1]+"\t"+"type"+"\n")
					for line in f:
						if float(line.split("\t")[-2])>5:
							out.write(line[:-1]+"\t"+"OG"+"\n")
						else:
							out.write(line[:-1]+"\t"+"TSG"+"\n")
	except:
		print ('check your file in data/ to execute script')
```
