## Supplementary material for "Comprehensive patient-level classification and quantification of driver events in TCGA PanCanAtlas cohorts": S8 Files: Instructions.txt

I.PREPARATION

1) To obtain the necessary starting files, run SNADRIF, GECNAV and ANDRIF pipelines before executing PALDRIC GENE pipeline.

Place the following files from the above-mentioned pipelines into the folder "PALDRIC_GENE/data":

mc3.v0.2.8.PUBLIC_primary_whitelisted_Entrez.tsv
SNA_classification_patients.tsv
SNA_classification_genes_NSEI_HISR.tsv
ISAR_GISTIC.all_thresholded.by_genes_primary_whitelisted.tsv
ISAR_GISTIC.all_thresholded.by_genes_primary_whitelisted_RNAfiltered.tsv
Primary_whitelisted_arms.tsv
Chromosome_drivers_FDR5.tsv
Arm_drivers_FDR5.tsv


2) Manually convert the outputs of third-party driver prediction algorithms into one of the following formats:

HUGO_symbol Ensembl_Transcript_ID amino_acid_substitution TCGA_cancer_cohort => for algorithms outputting lists of driver mutations

or

HUGO_symbol TCGA_cancer_cohort => for algorithms outputting lists of driver genes

Fill TCGA_cancer_cohort column with PANCAN if the analysis was not performed on separate cohorts.

Place the resulting files into the folder "PALDRIC_GENE/algorithms/algos".


II. EXECUTION


This script requires Python 3.7 and internet connection.

Always open terminal from the folder PALDRIC_GENE.

1) First, run the following command to install required libraries: 

pip3 install -r requirements.txt

2) Then run: 

python3 process_algo_outcome.py

Cohort-level driver gene (or driver mutation) lists with Entrez Gene IDs are automatically created and placed into the folder "PALDRIC_GENE/algorithms/with_entrez"

Patient-level driver gene (or driver mutation) lists with Entrez Gene IDs are automatically created and placed into the folder "PALDRIC_GENE/algorithms/result"

File endings indicate whether a given cohort-to-patient conversion was performed on the TCGA PanCanAtlas SNA data (_output_SNA.tsv), on the TCGA PanCanAtlas CNA data (_output_CNA.tsv) or on both datasets (_output.tsv).

To run a particular step (optional): 

python3 run_one_step.py N

where N is the step number


3) Finally, choose the file in the folder "PALDRIC_GENE/algorithms/result" that you want to analyze further and run the following command:

python3 pipeline.py AlgorithmName_output.tsv

A folder with the results will be automatically created under the name of the input file.

If you would like to combine driver gene lists from several algorithms, run: 

python3 pipeline.py 1 AlgorithmName1_output.tsv AlgorithmName2_output.tsv AlgorithmName3_output.tsv

You can combine as many algorithms as you want. Duplicate patient-gene pairs will be automatically removed. 


If you would like to find an overlap of driver gene lists from several algorithms, run: 

python3 pipeline.py N AlgorithmName1_output.tsv AlgorithmName2_output.tsv AlgorithmName3_output.tsv

where N should be replaced with a minimal number of algorithms that should predict the same gene in the same cancer type.

A folder with the results will be automatically created under the name composed of the date and time of the analysis. The file Algorithms.txt with the names of the files chosen for the analysis and the chosen degree of overlap will be added to that folder.


Examples of the commands:


python3 pipeline.py 2 2020plus_FDR5_COHORTS_output.tsv CHASMplus_FDR1_COHORTS_output.tsv CompositeDriver_FDR5_COHORTS_output.tsv dNdScv_TCGA_intogen_FDR5_COHORTS_output.tsv  DriverNet_FDR5_COHORTS_output.tsv HotMAPS_FDR5_COHORTS_output.tsv OncodriveCLUSTL_TCGA_intogen_FDR5_COHORTS_output.tsv OncodriveFML_TCGA_intogen_FDR5_COHORTS_output.tsv Bailey_official_COHORTS_output.tsv COSMIC_TIER1_SOMATIC_COHORTS_output.tsv


python3 pipeline.py 2 2020plus_FDR5_PANCAN_output.tsv CHASMplus_FDR1_PANCAN_output.tsv CompositeDriver_FDR5_PANCAN_output.tsv HotMAPS_FDR5_PANCAN_output.tsv OncodriveFML_FDR5_PANCAN_output.tsv SNADRIF_FDR5_PANCAN_output.tsv Bailey_official_PANCAN_output.tsv COSMIC_TIER1_SOMATIC_PANCAN_output.tsv
