## Supplementary material for "Comprehensive patient-level classification and quantification of driver events in TCGA PanCanAtlas cohorts": S8 Files: PALDRIC description.pdf

PAtient-Level DRIver Classifier is a Python 3.7 software package that translates cohort-level lists of driver genes or mutations from third-party algorithms to the patient level, classifies driver events according to the molecular causes and functional consequences, and presents comprehensive statistics on various kinds of driver events in various demographic and clinical groups of patients. PALDRIC allows to natively combine outputs of various third-party algorithms to investigate optimal combinations. PALDRIC is tightly integrated with SNADRIF (https://github.com/belikov-av/SNADRIF), GECNAV (https://github.com/belikov-av/GECNAV) and ANDRIF (https://github.com/belikov-av/ANDRIF) packages and requires their execution to obtain the necessary starting files. While conversion of the output files from third-party algorithms to a standard format necessary for ingestion has to be performed manually, the rest of the pipeline can be executed fully automatically in less than a minute per algorithm on a modern PC (Linux, Windows or MacOS).

This package has been developed by

Aleksey V. Belikov, Dr.rer.nat.
https://github.com/belikov-av
Laboratory of Innovative Medicine, School of Biological and Medical Physics, Moscow Institute of Physics and Technology
Roles: concept, pipeline, supervision

and

Danila V. Otnyukov
https://github.com/dan-otn
Department of bioinformatics, School of Biological and Medical Physics, Moscow Institute of Physics and Technology
Roles: programming

**Pipeline overview**

First, manually convert cohort-level lists of driver genes (or mutations) from third-party algorithms into standard format: HUGO symbol, (Ensembl Transcript ID, mutation), cohort, removing all entries with q-value >0.05.
Then, PALDRIC converts these standardized files into patient-level files using TCGA PanCanAtlas SNA and CNA data processed previously via SNADRIF and GECNAV packages.
You can then choose to analyze these processed outputs of third-party algorithms either individually or in any desirable combinations. You can also choose a degree of overlap between algorithms, depending on whether you would like to minimize false negatives or false positives.
Combining these results, PanCanAtlas clinical and demographic data, as well as data on chromosome and chromosomal arm gains and losses processed via ANDRIF package, PALDRIC automatically creates a comprehensive assortment of analyses. They include the number of patients with each integer total number of driver events (0,1,…,99, 100) for each cancer type (ACC,…, UVM, PANCAN); the average number of various types of driver events in patients with each integer total number of driver events (1,2,…,99, 100); the average number of various types of driver events in each cancer type (ACC,…, UVM, PANCAN); the average number of various types of driver events for each tumor stage (1,2,3,4); the average number of various types of

driver events for each age group (<25, 25-29,…,≥85). Each type of analysis is performed for the whole population, as well as for males and females separately. Corresponding multicolor cumulative histograms are plotted.

A detailed description of pipeline steps can be found in the file PALDRIC pipeline.pdf

Instructions for executing the code can be found in the file Instructions.txt
