## Supplementary material for "Comprehensive patient-level classification and quantification of driver events in TCGA PanCanAtlas cohorts": S8 Files: PALDRIC pipeline.pdf

**To obtain the necessary files, run SNADRIF, GECNAV and ANDRIF pipelines before executing PALDRIC pipeline.**

1) Go to https://gdc.cancer.gov/node/905/
2) Download Clinical with Follow-up - clinical_PANCAN_patient_with_followup.tsv
3) Remove from **clinical_PANCAN_patient_with_followup.tsv** all patients with **icd_o_3_histology** different from XXXX/3 (primary malignant neoplasm) and all patients not present (at the level TCGA-XX-XXXX) simultaneously in **mc3.v0.2.8.PUBLIC_primary_whitelisted_Entrez.tsv**, **ISAR_GISTIC.all_thresholded.by_genes_primary_whitelisted.tsv** and **Primary_whitelisted_arms.tsv** and save the resulting file as **clinical_PANCAN_patient_with_followup_primary_whitelisted.tsv**
4) A) Manually convert the outputs of third-party driver *mutation* prediction algorithms to tsv files with columns **HUGO symbol, Ensembl Transcript ID**, **mutation**, **cohort**, removing all results with q-value >0.05
   B) Manually convert the outputs of third-party driver *gene* prediction algorithms to tsv files with columns **HUGO symbol, cohort**, removing all results with q-value >0.05
5) Find Entrez Gene IDs using HUGO symbols and external database ftp://ftp.ncbi.nih.gov/gene/DATA/GENE_INFO/Mammalia/Homo_sapiens.gene_info.gz and update the file
6) A) For lists of driver *mutations*, remove all entries from **mc3.v0.2.8.PUBLIC_primary_whitelisted_Entrez.tsv** except those that satisfy the following conditions simultaneously: **Transcript_ID** matches **Ensembl Transcript ID** in the driver list; nucleotide/amino acid substitution matches the one in the driver list; cancer type (identified by matching **Tumor_Sample_Barcode** with **bcr_patient_barcode** and **acronym** in **clinical_PANCAN_patient_with_followup.tsv**) matches **cohort** in the driver list or the driver list is for pancancer analysis; **Variant_Classification** column contains one of the following values: De_novo_Start_InFrame, Frame_Shift_Del, Frame_Shift_Ins, In_Frame_Del, In_Frame_Ins, Missense_Mutation, Nonsense_Mutation, Nonstop_Mutation, Translation_Start_Site.
   Save the results as **AlgorithmName_output_SNA.tsv** with columns **TCGA Barcode, HUGO Symbol, Entrez Gene ID**
   B) For lists of driver *genes*, remove all entries from **mc3.v0.2.8.PUBLIC_primary_whitelisted_Entrez.tsv** except those that satisfy the following conditions simultaneously: **Entrez_Gene_ID** matches **Entrez Gene ID** in the driver list; cancer type (identified by matching **Tumor_Sample_Barcode** with **bcr_patient_barcode** and **acronym** in **clinical_PANCAN_patient_with_followup.tsv**) matches **cohort** in the driver list or the driver list is for pancancer analysis; **Variant_Classification** column contains one of the following values: De_novo_Start_InFrame, Frame_Shift_Del, Frame_Shift_Ins, In_Frame_Del, In_Frame_Ins, Missense_Mutation, Nonsense_Mutation, Nonstop_Mutation, Translation_Start_Site.
   Save the results as **AlgorithmName_output_SNA.tsv** with columns **TCGA Barcode, HUGO Symbol, Entrez Gene ID**

7) Remove all entries from **ISAR_GISTIC.all_thresholded.by_genes_primary_whitelisted.tsv** except those that satisfy the following conditions simultaneously: **Locus ID** matches **Entrez Gene ID** in the driver list; cancer type (identified by matching Tumor Sample Barcode with **bcr_patient_barcode** and **acronym** in **clinical_PANCAN_patient_with_followup.tsv**) matches **cohort** in the driver list or the driver list is for pancancer analysis; CNA values are 2,1, -1 or -2. Convert these data from the matrix to a list format (with columns **TCGA Barcode, HUGO Symbol, Entrez Gene ID**) and save as **AlgorithmName_output_CNA.tsv.**
8) Combine **AlgorithmName_output_SNA.tsv** and **AlgorithmName_output_CNA.tsv,** remove duplicate **TCGA Barcode-Entrez Gene ID** pairs, and save as **AlgorithmName_output.tsv**
9) Choose desired **AlgorithmName_output.tsv** files and fill the columns **TCGA Barcode, HUGO Symbol** and **Entrez Gene ID** of **AnalysisName_genes_level0.tsv,** removing duplicate **TCGA Barcode-Entrez Gene ID** pairs and patients not present in **clinical_PANCAN_patient_with_followup_primary_whitelisted.tsv.** If overlap was chosen by the user, remove all **TCGA Barcode-Entrez Gene ID** pairs present in fewer chosen **AlgorithmName_output.tsv** files than the user-chosen overlap number.
10) Use data from **SNA_classification_patients.tsv** to fill the columns **Number of hyperactivating SNAs** and **Number of inactivating SNAs** in **AnalysisName_genes_level0.tsv;** if a given **TCGA Barcode-Entrez Gene ID** pair is absent in **SNA_classification_patients.tsv,** write zeros. Use data from **SNA_classification_genes_NSEI_HISR.tsv** to fill the **HISR** column**;** if a given **Entrez Gene ID** is absent in **SNA_classification_genes_NSEI_HISR.tsv,** leave the cell empty. Use data from **ISAR_GISTIC.all_thresholded.by_genes_primary_whitelisted_RNAfiltered.tsv** to fill the **CNA status** column**;** if a given **TCGA Barcode-Entrez Gene ID** pair is absent in **ISAR_GISTIC.all_thresholded.by_genes_primary_whitelisted_RNAfiltered.tsv** write zero. Save the results as **AnalysisName_genes_level1.tsv**
11) Use data from **AnalysisName_genes_level1.tsv** to classify driver alterations according to the following table:

| Driver type | Number of hyperactivating SNAs + inactivating SNAs | Number of inactivating SNAs | HISR | CNA status | Count as … driver event(s) |
|---|---|---|---|---|---|
| SNA-based oncogene | ≥1 | 0 | >5 | 0 | 1 |
| CNA-based oncogene | 0 | 0 | >5 | 1 or 2 | 1 |
| Mixed oncogene | ≥1 | 0 | >5 | 1 or 2 | 1 |
| SNA-based tumor suppressor | ≥1 | ≥0 | ≤5 | 0 | 1 |
| CNA-based tumor suppressor | 0 | 0 | ≤5 | -1 or -2 | 1 |
| Mixed tumor suppressor | ≥1 | ≥0 | ≤5 | -1 or -2 | 1 |
| Passenger | 0 | 0 | | 0 | 0 |
| Low-probability driver | All the rest | | | | 0 |

and fill the columns **CNA status, Driver type** and **Count as … driver event(s)** of **AnalysisName_genes_level1.tsv**, saving it as **AnalysisName_genes_level2.tsv**
12) Use data from **AnalysisName_genes_level2.tsv, Chromosome_drivers_FDR5.tsv** и **Arm_drivers_FDR5.tsv**, to count for each patient the number of driver events of various classes (**Number of SNA-based oncogenic events, Number of CNA-based oncogenic events, Number of Mixed oncogenic events, Number of SNA-based tumor suppressor events, Number of CNA-based tumor suppressor events, Number of Mixed tumor suppressor events. Number of Driver chromosome losses, Number of**

**Driver chromosome gains, Number of Driver arm losses, Number of Driver arm gains, Total number of driver events**), counting each tumor suppressor as 2 events (see table above)**.** Use data from **clinical_PANCAN_patient_with_followup_primary_whitelisted.tsv** to fill the columns **Cancer type (acronym), Gender (gender), Age (age_at_initial_pathologic_diagnosis),** and **Tumor stage** (**pathologic_stage,** if data absent then **clinical_stage**, if data absent then **pathologic_T**, if data absent then **clinical_T**, convert to Arabic number**)** and add patients without any identified drivers (i.e. whose TCGA barcodes are absent in **AnalysisName_genes_level2.tsv, Chromosome_drivers_FDR5.tsv** and **Arm_drivers_FDR5.tsv,** but present in **clinical_PANCAN_patient_with_followup_primary_whitelisted.tsv**) writing zero values for them and saving the results as **AnalysisName_patients.tsv**

13) Use data from **AnalysisName_patients.tsv** to count the number of patients with each integer total number of driver events (0,1,…,99, 100) for each cancer type, also for males and females separately, and save as **AnalysisName_distribution_events.tsv, AnalysisName_distribution_events_males.tsv** and **AnalysisName_distribution_events_females.tsv**. For each cancer type-gender combination plot a histogram.
14) Use data from **AnalysisName_patients.tsv** to count the average number of various types of driver events in patients with each integer total number of driver events (1,2,…, 99, 100), also for males and females separately, and save as **AnalysisName_distribution_events_detailed.tsv, AnalysisName_distribution_events_detailed_males.tsv** and **AnalysisName_distribution_events_detailed_females.tsv**. For each file plot a multicolor cumulative histogram “Driver event distribution by total number of driver events per patient”.
15) Use data from **AnalysisName_patients.tsv** to count the average number of various types of driver events in each cancer type (ACC,…, UVM, PANCAN), also for males and females separately, and save as **AnalysisName_distribution_cohorts.tsv, AnalysisName_distribution_cohorts_males.tsv** and **AnalysisName_distribution_cohorts_females.tsv**. For each file plot a multicolor cumulative histogram “Driver event distribution by cancer type”
16) Use data from **AnalysisName_patients.tsv** to count the average number of various types of driver events for males and females separately, and save as **AnalysisName_distribution_gender.tsv.** Plot a multicolor cumulative histogram “Driver event distribution by gender”
17) Use data from **AnalysisName_patients.tsv** to count the average number of various types of driver events for each tumor stage (1,2,3,4), also for males and females separately, and save as **AnalysisName_distribution_stage.tsv, AnalysisName_distribution_stage_males.tsv** and **AnalysisName_distribution_stage_females.tsv**. For each file plot a multicolor cumulative histogram “Driver event distribution by cancer stage”
18) Use data from **AnalysisName_patients.tsv** to count the average number of various types of driver events for each age group (<25, 25-29,…,≥85), also for males and females separately, and save as **AnalysisName_distribution_age.tsv, AnalysisName_distribution_age_males.tsv** and **AnalysisName_distribution_age_females.tsv**. For each file plot a multicolor cumulative histogram “Driver event distribution by age”
