## Supplementary material for "Comprehensive patient-level classification and quantification of driver events in TCGA PanCanAtlas cohorts": S8 Files: pipeline.py

```
import sys
import os
import pandas as pd
from datetime import datetime
from collections import Counter
from step_9 import step9
from step_10 import step10
from step_11 import step11
from step_12 import step12
from step_13 import step13
from step_14 import step14
from step_15 import step15
from step_16 import step16
from step_17 import step17
from step_18 import step18


print ("start analysis")
overlap = int(sys.argv[1])
algos = sys.argv[2:]
print ('combine data:')
algs = []
a = []
for i in algos:
	print (i[:-4])
	if len(algos)>1:
		a.append(i.split("_")[0])
	else:
		a.append(i[:-4])
	algs.append(pd.read_csv('algorithms/result/'+i, sep='\t').drop_duplicates())

result = pd.concat(algs)
## make overlap
result['overlap_pair'] = result['TCGA barcode']+"/"+result['Entrez id'].apply(str)
cc = Counter(result['overlap_pair'])
overlapped = [k for k in cc if cc[k]>=overlap]
result = result.loc[result['overlap_pair'].isin(overlapped),['TCGA barcode','HUGO symbol','Entrez id']]


result = result.drop_duplicates()

d = datetime.now()
d = [d.year,d.month,d.day,d.hour,d.minute]
d = [str(i) for i in d]
if len(algos)>1:
	name ="_".join(d)
else:
	name = "_".join(a)


try:
	os.mkdir(name)
except:
	pass
with open(name+'/algorithms.txt','w') as f:
	f.write('overlap = '+ str(overlap)+"\n")
	for i in algos:
		f.write(i+"\n")

if len(algos)>1:
	try:
		os.mkdir('algorithms/result/combined/')
	except:
		pass
	result.to_csv('algorithms/result/combined/'+name+".tsv", sep='\t', index=False)
	result.to_csv('algorithms/result/'+name+".tsv", sep='\t', index=False)
	print ("Saved as",name+".tsv")
else:
	result.to_csv('algorithms/result/'+name+".tsv", sep='\t', index=False)
	print ("Saved as",name+".tsv")

algo = name+".tsv"

step9(algo)
step10(algo)
step11(algo)
step12(algo)
step13(algo)
step14(algo)
step15(algo)
step16(algo)
step17(algo)
step18(algo)

if len(algos)>1:
	os.remove('algorithms/result/'+name+".tsv")
print ("completed")
```
