## Supplementary material for "Comprehensive patient-level classification and quantification of driver events in TCGA PanCanAtlas cohorts": S8 Files: process_algo_outcome.py

```
import sys
import os
import pandas as pd
from step_2 import step2
from step_3 import step3
from step_4 import step4
from step_5 import step5
from step_6 import step6
from step_7 import step7
from step_8 import step8

print('start proccesing')
step2()
step3()
step4()
step5()
step6()
step7()
step8()


print ("completed")
```
