## Supplementary material for "Comprehensive patient-level classification and quantification of driver events in TCGA PanCanAtlas cohorts": S8 Files: run_one_step.py

```
import sys
import os
import pandas as pd
from collections import Counter
from step_2 import step2
from step_3 import step3
from step_4 import step4
from step_5 import step5
from step_6 import step6
from step_7 import step7
from step_8 import step8
from step_9 import step9
from step_10 import step10
from step_11 import step11
from step_12 import step12
from step_13 import step13
from step_14 import step14
from step_15 import step15
from step_16 import step16
from step_17 import step17
from step_18 import step18

print ("start analysis")
step = sys.argv[1]
steps = {2:step2,
	3:step3,
	4:step4,
	5:step5,
	6:step6,
	7:step7,
	8:step8,
	9:step9,
	10:step10,
	11:step11,
	12:step12,
	13:step13,
	14:step14,
	15:step15,
	16:step16,
	17:step17,
	18:step18,}
print (step)
if int(step)<10:
	steps[int(step)]()
else:
	algos = sys.argv[3:]
	overlap = int(sys.argv[1])
	print ('Combine data:')
	algs = []
	a = []
	for i in algos:
		print (i[:-4])
		if len(algos)>1:
			a.append(i.split("_")[0])
		else:
			a.append(i[:-4])
		algs.append(pd.read_csv('algorithms/result/'+i, sep='\t').drop_duplicates())
	result = pd.concat(algs)
	## make overlap
	result['overlap_pair'] = result['TCGA barcode']+"/"+result['Entrez id'].apply(str)
	cc = Counter(result['overlap_pair'])
	overlapped = [k for k in cc if cc[k]>=overlap]
	result = result.loc[result['overlap_pair'].isin(overlapped),['TCGA barcode','HUGO symbol','Entrez id']]
	result = result.drop_duplicates()
	name = "_".join(a)
	if len(algos)>1:
		try:
			os.mkdir('algorithms/result/combined/')
		except:
			pass
		result.to_csv('algorithms/result/'+name+".tsv", sep='\t', index=False)
		result.to_csv('algorithms/result/combined/'+name+".tsv", sep='\t', index=False)
		#print ("Saved as",name+".tsv")

	algo = name+".tsv"
	try:
		steps[int(step)](algo)
	except:
		print('Check your datafiles from previous steps')
	if len(algos)>1:
		try:
			os.remove('algorithms/result/'+name+".tsv")
		except:
			pass
print ("completed")
```
