## Supplementary material for "Comprehensive patient-level classification and quantification of driver events in TCGA PanCanAtlas cohorts": S8 Files: step_2.py

```
import os.path
import sys
import requests
import hashlib

def check_hash(file_path):
	d1 = {'clinical_PANCAN_patient_with_followup.tsv':('ffcb35edda305dd8d615497f9214eb92','216f5eb4216113da10654ff4fb5f44454eadb078'),
	'merged_sample_quality_annotations.tsv':('05ddd2270fb1fb24fbdc2fe9bf7384e5','a70a070954813a905d3155fd1a0bab335c10ec88')}

	BUF_SIZE = 65536 
	with open(file_path, 'rb') as f:
		md5 = hashlib.md5()
		sha1 = hashlib.sha1()
		while True:
			data = f.read(BUF_SIZE)
			if not data:
				break
			md5.update(data)
			sha1.update(data)

	tf = file_path.split('/')[-1]

	if d1[tf] == (md5.hexdigest(),sha1.hexdigest()):
		return True
	else:
		return False
def download(link,file_name):
	with open(file_name, "wb") as f:
		#print "Downloading %s" % file_name
		response = requests.get(link, stream=True)
		total_length = response.headers.get('content-length')

		if total_length is None: # no content length header
			f.write(response.content)
		else:
			dl = 0
			total_length = int(total_length)
			for data in response.iter_content(chunk_size=4096):
				dl += len(data)
				f.write(data)
				done = int(50 * dl / total_length)
				sys.stdout.write("\r[%s%s]" % ('=' * done, ' ' * (50-done)) )    
				sys.stdout.flush()
	print("\n")
		
def step2():
	print ('step 2 start')
	url = 'http://api.gdc.cancer.gov/data/0fc78496-818b-4896-bd83-52db1f533c5c'
	file = 'data/clinical_PANCAN_patient_with_followup.tsv'

	try:
		if os.path.exists(file) and check_hash(file):
			pass
		else:
			download(url, file)
	except:
		print ('cant download file: '+ url)
	print('step 2 end')
```
