## Supplementary material for "Comprehensive patient-level classification and quantification of driver events in TCGA PanCanAtlas cohorts": S8 Files: step_3.py

```
import os
import pandas as pd
import numpy as np

def sample_to_Patient(sample):
	return '-'.join(sample.split('-')[:3])

def icd_cond(s):
	if s == '[Not Available]':
		return False
	else:
		return s.split('/')[1] == '3'
def step3():
	print ("step 3 start")
	vect_sam2pat = np.vectorize(sample_to_Patient)
	input_path = 'data/clinical_PANCAN_patient_with_followup.tsv'
	mc3_path = 'data/mc3.v0.2.8.PUBLIC_primary_whitelisted_Entrez.tsv'
	cna_path = 'data/ISAR_GISTIC.all_thresholded.by_genes_primary_whitelisted.tsv'
	aneupl_path = "data/Primary_whitelisted_arms.tsv"
	#'PANCAN_ArmCallsAndAneuploidyScore_092817_primary_whitelisted.tsv'
	output_file_path = 'data/clinical_PANCAN_patient_with_followup_primary_whitelisted.tsv'


	input_file = pd.read_csv(input_path, sep='\t', encoding = 'unicode_escape',low_memory=False)

		# reading snv patients
	mc3_patients = list(pd.read_csv(mc3_path, sep='\t', usecols=['Tumor_Sample_Barcode'])['Tumor_Sample_Barcode'] \
							.unique())
	mc3_patients = vect_sam2pat(mc3_patients)

		# reading cna patients
	cna_patients_ix = 3
	with open(cna_path, 'r') as cna_file:
		cna_patients = cna_file.readline()[:-1].split('\t')[cna_patients_ix:]
	cna_patients = vect_sam2pat(cna_patients)

		# reading aneuploidy_patients!!!!!!!!!!!!
	aneuploidy_patients = list(pd.read_csv(aneupl_path, sep='\t', usecols=['Sample'])['Sample'])
	aneuploidy_patients = vect_sam2pat(aneuploidy_patients)

		# filter inproper 'icd_o_3_histology'
	input_file['icd_o_3_histology_status'] = input_file['icd_o_3_histology'].apply(icd_cond)
	input_file = input_file[input_file['icd_o_3_histology_status']]
	input_file = input_file.reset_index().drop(columns=['index', 'icd_o_3_histology_status'])

		# filter inproper patients
	patients_intersection = set(input_file['bcr_patient_barcode'])
	patients_intersection = patients_intersection.intersection(set(mc3_patients),
																   set(cna_patients),
																   set(aneuploidy_patients))
	input_file = input_file[input_file['bcr_patient_barcode'].isin(patients_intersection)]

	input_file.to_csv(output_file_path, sep='\t', index=None)
	print ('step 3 end')
```
