## Supplementary material for "Comprehensive patient-level classification and quantification of driver events in TCGA PanCanAtlas cohorts": S8 Files: step_4.py

```
def step4():
	header4 = ['HUGO symbol', 'Entrez Gene ID', 'cohort']
	header1 = ['HUGO symbol', 'Entrez Gene ID', 'Ensembl Transcript ID', 'mutation', 'cohort']
	print('step 4 start')
	possible_eff = {"3'Flank":"0",
	 'De_novo_Start_OutOfFrame':"0",
	  'De_novo_Start_InFrame':'1',
	 "3'UTR":"0",
	 "5'Flank":"0",
	 "5'UTR":"0",
	 'Frame_Shift_Del':'1',
	 'Frame_Shift_Ins':'1',
	 'In_Frame_Del':'1',
	 'In_Frame_Ins':'1',
	 'IGR':"0",
	 'Targeted_Region':"0",
	 'Intron':"0",
	 'Missense_Mutation':'1',
	 'Nonsense_Mutation':'1',
	 'Nonstop_Mutation':'1',
	 'RNA':"0",
	 'Silent':'0', 
	 'Splice_Site':"0",
	 'Translation_Start_Site':'1'}
```
