## Supplementary material for "Comprehensive patient-level classification and quantification of driver events in TCGA PanCanAtlas cohorts": S8 Files: step_5.py

```
import pandas as pd
import os
def step5():
	print('step 5 start')
	algos = os.listdir('algorithms/algos/')
	algos = [a for a in algos if ".txt" in a or ".tsv" in a]
	header4 = ['HUGO symbol', 'Entrez Gene ID', 'cohort']
	header1 = ['HUGO symbol', 'Entrez Gene ID', 'Ensembl Transcript ID', 'mutation', 'cohort']

	possible_eff = {"3'Flank":"0",
	 'De_novo_Start_OutOfFrame':"0",
	  'De_novo_Start_InFrame':'1',
	 "3'UTR":"0",
	 "5'Flank":"0",
	 "5'UTR":"0",
	 'Frame_Shift_Del':'1',
	 'Frame_Shift_Ins':'1',
	 'In_Frame_Del':'1',
	 'In_Frame_Ins':'1',
	 'IGR':"0",
	 'Targeted_Region':"0",
	 'Intron':"0",
	 'Missense_Mutation':'1',
	 'Nonsense_Mutation':'1',
	 'Nonstop_Mutation':'1',
	 'RNA':"0",
	 'Silent':'0', 
	 'Splice_Site':"0",
	 'Translation_Start_Site':'1'}
	annot = dict()
	a2 =dict()
	a2_not_syn =dict()
	with open("data/Homo_sapiens.gene_info",'r') as an:
	#print(an.readline().split("\t"))
		for line in an:
			#annot[line.split("\t")[5]] = line.split("\t")[6]
			synonims = line.split("\t")[4]
			synonims = synonims.split('|')
			for i in synonims:
				a2[i] = line.split("\t")[1]
			a2_not_syn[line.split("\t")[2]] = line.split("\t")[1]
			
	for a in algos:
		with open('algorithms/algos/'+a,'r') as file:
			file.readline()
			check = file.readline().split('\t')
		if len(check) == 2:
			type_file = 1
		else:
			if '.' not in check[-2]:
				type_file = 2
			else:
				type_file = 3
		if type_file == 1:
			df = pd.read_csv('algorithms/algos/'+a,sep="\t",header=None,names=[header4[0],header4[-1]])
			for i in range(len(df)):
				if df.iloc[i][0] in a2_not_syn.keys():
					df.loc[i,'Entrez Gene ID'] = a2_not_syn[df.iloc[i][0]]
				elif df.iloc[i][0] in a2.keys():
					df.loc[i,'Entrez Gene ID'] = a2[df.iloc[i][0]]
				else:
					df.loc[i,'Entrez Gene ID'] = None
			df = df[header4]
			df = df.dropna()
			df.to_csv('algorithms/with_entrez/'+a.split('.')[0]+'.tsv', header=True, index=False, sep='\t')
			
		elif type_file == 2:
			df = pd.read_csv('algorithms/algos/'+a,sep="\t",header=None,names=[header1[0],header1[2],header1[3],header1[4]])
			for i in range(len(df)):
				if df.iloc[i][0] in a2_not_syn.keys():
					df.loc[i,'Entrez Gene ID'] = a2_not_syn[df.iloc[i][0]]
				elif df.iloc[i][0] in a2.keys():
					df.loc[i,'Entrez Gene ID'] = a2[df.iloc[i][0]]
				else:
					df.loc[i,'Entrez Gene ID'] = None
			df = df[header1]
			df = df.dropna()
			df.to_csv('algorithms/with_entrez/'+a.split('.')[0]+'.tsv', header=True, index=False, sep='\t')
		
		elif type_file == 3:
			df = pd.read_csv('algorithms/algos/'+a,sep="\t",header=None,names=[header1[0],header1[2],header1[3],header1[4]])
			for i in range(len(df)):
				if df.iloc[i][0] in a2_not_syn.keys():
					df.loc[i,'Entrez Gene ID'] = a2_not_syn[df.iloc[i][0]]
				elif df.iloc[i][0] in a2.keys():
					df.loc[i,'Entrez Gene ID'] = a2[df.iloc[i][0]]
				else:
					df.loc[i,'Entrez Gene ID'] = None
			df = df[header1]
			df = df.dropna()
			df.to_csv('algorithms/with_entrez/'+a.split('.')[0]+'.tsv', header=True, index=False, sep='\t')

	print('step 5 end')
```
