## Supplementary material for "Comprehensive patient-level classification and quantification of driver events in TCGA PanCanAtlas cohorts": S8 Files: step_6.py

```
from collections import defaultdict
import os
def step6():
	algos = os.listdir('algorithms/algos')
	algos = [a for a in algos if ".txt" in a or ".tsv" in a]
	algos2 = os.listdir('algorithms/with_entrez/')
	algos2 = [a for a in algos2 if ".txt" in a or ".tsv" in a]
	#algos.extend(algos2)
	#algos = set(algos)
	header4 = ['HUGO symbol', 'Entrez Gene ID', 'cohort']
	header1 = ['HUGO symbol', 'Entrez Gene ID', 'Ensembl Transcript ID', 'mutation', 'cohort']
	print('step 6 start')
	possible_eff = {"3'Flank":"0",
	 'De_novo_Start_OutOfFrame':"0",
	  'De_novo_Start_InFrame':'1',
	 "3'UTR":"0",
	 "5'Flank":"0",
	 "5'UTR":"0",
	 'Frame_Shift_Del':'1',
	 'Frame_Shift_Ins':'1',
	 'In_Frame_Del':'1',
	 'In_Frame_Ins':'1',
	 'IGR':"0",
	 'Targeted_Region':"0",
	 'Intron':"0",
	 'Missense_Mutation':'1',
	 'Nonsense_Mutation':'1',
	 'Nonstop_Mutation':'1',
	 'RNA':"0",
	 'Silent':'0', 
	 'Splice_Site':"0",
	 'Translation_Start_Site':'1'}
	d =  {}#1,2
	with open('data/clinical_PANCAN_patient_with_followup.tsv','r',encoding = 'unicode_escape') as f:
		for line in f:
			try:
				d[line.split('\t')[1]] = line.split('\t')[2]
			except:
				pass

	for a in algos2:
		with open('algorithms/with_entrez/'+a,'r') as file:
			file.readline()
			check = file.readline().split('\t')
		if len(check) == 3:
			type_file = 1
		else:
			if '.' not in check[-2]:
				type_file = 2
			else:
				type_file = 3
		if type_file == 1:
			print(a)
			#a = a.split('.')[0]+'.tsv'
			cohort = defaultdict(list)
			entrezes = {}
			#enst = defaultdict(list)
			#poly = defaultdict(list)
			with open('algorithms/with_entrez/'+a,'r') as f:
				f.readline()
				for line in f:
					entrez = line.split("\t")[1]
					entrezes[line.split("\t")[0]] = entrez
					#enst = line.split("\t")[2]
					#cohort[entrez].append(line.split("\t")[-1][:-1])
					cohort[entrez].append(line.split("\t")[-1][:-1])

			with open("data/mc3.v0.2.8.PUBLIC_primary_whitelisted_Entrez.tsv",'r') as f:
				with open("algorithms/result/"+a[:-4]+"_output_SNA.tsv",'w') as out:
					f.readline()
					out.write("TCGA barcode"+"\t"+"HUGO symbol"+"\t"+"Entrez id"+"\n")
					for line in f:
						fl = 0
						a = line.split("\t")
						if a[1] in cohort.keys() :

							if a !=[''] and possible_eff[a[8]] == '1': 
								if a[15][:12] in d.keys():
									if  d[a[15][:12]] in cohort[a[1]]: ###acronym
										fl = 1
									if 'PANCAN' in cohort[a[1]]:
										fl = 1

						if fl == 1:
							try:
								out.write(a[15]+"\t"+a[0]+"\t"+entrezes[a[0]]+"\n")
							except:
								pass
		elif type_file == 2:
			print(a)
			#a = a.split('.')[0]+'.tsv'
			cohort = defaultdict(list)
			entrezez = {}
			enst = defaultdict(list)
			poly = defaultdict(list)
			with open('algorithms/with_entrez/'+a,'r') as f:
				f.readline()
				for line in f:
					entrez = line.split("\t")[1]
					entrezes[line.split("\t")[0]] = entrez
					enst = line.split("\t")[2]
					cohort[enst].append(line.split("\t")[-1][:-1])
					poly[enst].append(line.split("\t")[-2])
					#cohort[entrez].append(line.split("\t")[-1][:-1])

		#a[53]  in poly[a[0]]   mutations only 

			with open("data/mc3.v0.2.8.PUBLIC_primary_whitelisted_Entrez.tsv",'r') as f:
				with open("algorithms/result/"+a[:-4]+"_output_SNA.tsv",'w') as out:
					f.readline()
					out.write("TCGA barcode"+"\t"+"HUGO symbol"+"\t"+"Entrez id"+"\n")
					for line in f:
						fl = 0
						a = line.split("\t")
						if a[37] in cohort.keys() :

							if a !=[''] and possible_eff[a[8]] == '1'and a[53]  in poly[a[37]]: 
								if a[15][:12] in d.keys():
									if  d[a[15][:12]] in cohort[a[37]]: ###acronym
										fl = 1
									if 'PANCAN' in cohort[a[37]]:
										fl = 1

						if fl == 1:
							try:
								out.write(a[15]+"\t"+a[0]+"\t"+entrezes[a[0]]+"\n")
							except:
								pass
		elif type_file ==3:
			print(a)
			#a = a.split('.')[0]+'.tsv'
			cohort = defaultdict(list)
			entrezes = {}
			enst = defaultdict(list)
			poly = defaultdict(list)
			with open('algorithms/with_entrez/'+a,'r') as f:
				f.readline()
				for line in f:
					entrez = line.split("\t")[1]
					enst = line.split("\t")[2]
					entrezes[line.split("\t")[0]] = entrez
					cohort[enst].append(line.split("\t")[-1][:-1])
					poly[enst].append(line.split("\t")[-2])
					#cohort[entrez].append(line.split("\t")[-1][:-1])

		#a[53]  in poly[a[0]]   mutations only 

			with open("data/mc3.v0.2.8.PUBLIC_primary_whitelisted_Entrez.tsv",'r') as f:
				with open("algorithms/result/"+a[:-4]+"_output_SNA.tsv",'w') as out:
					f.readline()
					out.write("TCGA barcode"+"\t"+"HUGO symbol"+"\t"+"Entrez id"+"\n")
					for line in f:
						fl = 0
						a = line.split("\t")
						if a[37] in cohort.keys() :

							if a !=[''] and possible_eff[a[8]] == '1'and a[36]  in poly[a[37]]: 
								if a[15][:12] in d.keys():
									if  d[a[15][:12]] in cohort[a[37]]: ###acronym
										fl = 1
									if 'PANCAN' in cohort[a[37]]:
										fl = 1

						if fl == 1:
							try:
								out.write(a[15]+"\t"+a[0]+"\t"+entrezes[a[0]]+"\n")
							except:
								pass

	print('step 6 end')
```
