## Supplementary material for "Comprehensive patient-level classification and quantification of driver events in TCGA PanCanAtlas cohorts": S8 Files: step_7.py

```
from collections import defaultdict
import os
def step7():
	print ('step 7 start')
	d =  {}#1,2
	with open('data/clinical_PANCAN_patient_with_followup.tsv','r',encoding = 'unicode_escape') as f:
		for line in f:
			try:
				d[line.split('\t')[1]] = line.split('\t')[2]
			except:
				pass
	algos = os.listdir('algorithms/with_entrez/')
	algos = [a for a in algos if ".txt" in a or ".tsv" in a]
	for alg in algos:
		print(alg)
		cohort = defaultdict(list)
		with open('algorithms/with_entrez/'+alg,'r') as f:
			f.readline()
			for line in f:
				entrez = line.split("\t")[1]
				cohort[entrez].append(line.split("\t")[-1][:-1])

		with open("data/ISAR_GISTIC.all_thresholded.by_genes_primary_whitelisted.tsv",'r') as f:
			with open("algorithms/result/"+alg.split('.')[0]+"_output_CNA.tsv",'w') as out:
				out.write("TCGA barcode"+"\t"+"HUGO symbol"+"\t"+"Entrez id"+"\n")
				#bb = f.readline().split("\t")
				brcd =f.readline().split("\t")[3:]
				for line in f:
					a = line.split("\t")
					pos = a[3:]
					fl = 0
					if a[1] in cohort.keys():
						for i in range(len(pos)):
							fl = 0
							if pos[i] =='1' or pos[i] == '-1' or pos[i] =='2' or pos[i] == '-2':
								if brcd[i][:12] in d.keys():
									if d[brcd[i][:12]] in cohort[a[1]]:
										fl = 1
									if "PANCAN" in cohort[a[1]]:
										fl = 1
							if fl == 1 and a[1] in cohort.keys():

								out.write(brcd[i]+"\t"+a[0]+"\t"+a[1]+"\n")
	print ('step 7 start')
```
