## Supplementary material for "Comprehensive patient-level classification and quantification of driver events in TCGA PanCanAtlas cohorts": S8 Files: step_8.py

```
import os
def step8():
	print ("step 8 start")
	
	algos = os.listdir('algorithms/with_entrez/')
	algos = [a for a in algos if ".txt" in a or ".tsv" in a]
	for alg in algos:
		with open("algorithms/result/"+alg[:-4]+"_output_SNA.tsv",'r') as f:
			f.readline()
			alls = set()
			for line in f:
				alls.add(line)
		with open("algorithms/result/"+alg[:-4]+"_output_CNA.tsv",'r') as f:
			h = f.readline()
			with open("algorithms/result/"+alg[:-4]+"_output.tsv",'w') as out:
				out.write(h)
				for line in f:
					if line not in alls:
						out.write(line)
				for i in alls:
					out.write(i)
	import pandas as pd
	a = pd.read_csv("algorithms/result/"+alg[:-4]+"_output.tsv", sep='\t')
	a = a.drop_duplicates()
	a.to_csv("algorithms/result/"+alg[:-4]+"_output.tsv", sep='\t', index=False)
	print ("step 8 end")
```
