## Supplementary material for "Comprehensive patient-level classification and quantification of driver events in TCGA PanCanAtlas cohorts": S8 Files: step_9.py

```
import pandas as pd
import os
from utils.utils import barcode_to_id

def step9(algo = 'Bayley'):
	print ("step 9 start")
	algo_name = algo[:-4]
	import os
	try:
		os.mkdir(algo_name)
	except:
		pass
	try:
		os.mkdir(algo_name+'/data/')
	except:
		pass
	alg_analysis_path = 'algorithms/result/'+algo
	clinical_PANCAN_path = 'data/clinical_PANCAN_patient_with_followup_primary_whitelisted.tsv'
	output_df = algo[:-4]+'/data/'+algo[:-4] +'_genes_level0.tsv'
	
	alg_analysis_df = pd.read_csv(alg_analysis_path, sep='\t')
	clin_pancan_df = pd.read_csv(clinical_PANCAN_path, sep='\t', usecols=['bcr_patient_barcode', 'acronym'])
	alg_analysis_df['TCGA barcode'] = [barcode_to_id(uuid) for uuid in alg_analysis_df['TCGA barcode']]
	clin_pancan_patients = set(list(clin_pancan_df['bcr_patient_barcode']))
	alg_analysis_df = alg_analysis_df[alg_analysis_df['TCGA barcode'].isin(clin_pancan_patients)]
	alg_analysis_df.reset_index(inplace=True, drop=True)
	alg_analysis_df['Number of hyperactivating SNAs'] = ''
	alg_analysis_df['Number of inactivating SNAs'] = ''
	alg_analysis_df['HISR'] = ''
	alg_analysis_df['CNA status'] = ''
	alg_analysis_df['Driver type'] = ''
	alg_analysis_df['Count as driver events'] = ''
	alg_analysis_df = alg_analysis_df.drop_duplicates()
	alg_analysis_df.to_csv(output_df, sep='\t', index=False)
	print ("step 9 end")
```
