## Supplementary material for "Comprehensive patient-level classification and quantification of driver events in TCGA PanCanAtlas cohorts": S8 Files: step_10.py

```
import os
from utils.utils import barcode_to_id

def step10(algo):
	print ("step 10 start")
	algo_name = algo[:-4]
	try:
		os.mkdir(algo_name)
	except:
		pass
	input_file = algo[:-4] +'/data/'+algo[:-4] + '_genes_level0.tsv'
	sna_class = 'data/SNA_classification_patients.tsv'
	hisr_file = 'data/SNA_classification_genes_NSEI_HISR.tsv'
	cna_file = 'data/ISAR_GISTIC.all_thresholded.by_genes_primary_whitelisted_RNAfiltered.tsv'
	output_file = algo[:-4] +'/data/'+algo[:-4] + '_genes_level1.tsv'
	flag = 0
	cna = {}
	if 'for10step-1.tsv' in list(os.listdir('data/')) and 'for10step-2.tsv' in list(os.listdir('data/')): 
		flag = 1
	if flag == 0:
		with open (cna_file,'r') as inpt:
			with open('data/for10step-1.tsv','w') as out:
				brcds = inpt.readline().split("\t")[3:]
				brcds[-1] = brcds[-1][:-1]
				for line in inpt:
					l = line.split("\t")
					l[-1] = l[-1][:-1]
					entrez = l[0]
					hugo = l[1]
					statuses=l[3:]
					for i in range(len(statuses)):
						if statuses[i]=='-1':
							out.write(brcds[i]+"\t"+entrez+"\t"+hugo+"\t"+statuses[i]+"\n")
		with open (cna_file,'r') as inpt:
			with open('data/for10step1.tsv','w') as out:
				brcds = inpt.readline().split("\t")[3:]
				brcds[-1] = brcds[-1][:-1]
				for line in inpt:
					l = line.split("\t")
					l[-1] = l[-1][:-1]
					entrez = l[0]
					hugo = l[1]
					statuses=l[3:]
					for i in range(len(statuses)):
						if statuses[i]=='1':
							out.write(brcds[i]+"\t"+entrez+"\t"+hugo+"\t"+statuses[i]+"\n")
		with open (cna_file,'r') as inpt:
			with open('data/for10step2.tsv','w') as out:
				brcds = inpt.readline().split("\t")[3:]
				brcds[-1] = brcds[-1][:-1]
				for line in inpt:
					l = line.split("\t")
					l[-1] = l[-1][:-1]
					entrez = l[0]
					hugo = l[1]
					statuses=l[3:]
					for i in range(len(statuses)):
						if statuses[i]=='2':
							out.write(brcds[i]+"\t"+entrez+"\t"+hugo+"\t"+statuses[i]+"\n")
						
		with open (cna_file,'r') as inpt:
			with open('data/for10step-2.tsv','w') as out:
				brcds = inpt.readline().split("\t")[3:]
				brcds[-1] = brcds[-1][:-1]
				for line in inpt:
					l = line.split("\t")
					l[-1] = l[-1][:-1]
					entrez = l[0]
					hugo = l[1]
					statuses=l[3:]
					for i in range(len(statuses)):
						if statuses[i]=='-2':
							out.write(brcds[i]+"\t"+entrez+"\t"+hugo+"\t"+statuses[i]+"\n")
	sna = {}
	with open(sna_class,'r') as inpt:
		inpt.readline().split("\t")
		for line in inpt:
			s = line.split("\t")
			#key = barcode_to_id(s[0])+s[1]+s[2]
			key = barcode_to_id(s[0])+s[2]###brcd + entrez: muts
			sna[key] = (s[4],s[5])
	print ("loaded 1st part of data")
	hisr = {} ## 
	with open(hisr_file,'r') as inpt:
		inpt.readline().split("\t")
		for line in inpt:
			s = line.split("\t")
			#hisr[(s[0],s[1])] = s[-1][:-1]
			hisr[s[1]] = s[-1][:-1]###entrez:hisr
		
	cna = {} ## 'Hugo_Symbol', 'Entrez_Gene_Id'
	with open('data/for10step1.tsv','r') as inpt:
		for line in inpt:
			s = line.split("\t")
			key = s[0]+s[1]
        
			cna[key] = s[-1][:-1]
	with open(input_file,'r') as inpt:
		with open(output_file,'w') as outpt:
			outpt.write(inpt.readline())
			for line in inpt:
				s = line.split("\t")## brcd hugo entrez
				current = s[0]+"\t"+s[1]+"\t"+s[2]
				s[0] = barcode_to_id(s[0])
				#key  = s[0]+s[1]+s[2]#[:-1]
				key  = s[0]+s[2]#[:-1]
				key_cna = s[0]+s[2]
				if key in sna.keys():
					current += "\t"+ sna[key][0]
					current += "\t"+ sna[key][1]
				else:
					current += "\t"+ '0'
					current += "\t"+ '0'
				if s[2] in hisr.keys():
					current += "\t"+ hisr[s[2]]
				else:
					current += "\t"
				if key_cna in cna.keys():
					current += "\t"+ cna[key_cna]  
				else:
					current += "\t"+ '0'
				current += "\n"
				outpt.write(current)
	print ("loaded 2nd part of data")
	cna = {} ## 'Hugo_Symbol', 'Entrez_Gene_Id'
	with open('data/for10step-1.tsv','r') as inpt:
		for line in inpt:
			s = line.split("\t")
			key = s[0]+s[1]

			cna[key] = s[-1][:-1]
	with open(output_file,'r') as inpt:
		with open('data/temp.tsv','w') as outpt:
			outpt.write(inpt.readline())
			for line in inpt:
				s = line.split("\t")## brcd hugo entrez
				key_cna = s[0]+s[2]
				if key_cna in cna.keys():
					s[-1] = cna[key_cna]  +"\n"
				current = "\t".join(s)
				outpt.write(current)
	cna = {} ## 'Hugo_Symbol', 'Entrez_Gene_Id'
	#print ("started analysis")
	with open('data/for10step2.tsv','r') as inpt:
		for line in inpt:
			s = line.split("\t")
			key = s[0]+s[1]
			cna[key] = s[-1][:-1]
	with open('data/temp.tsv','r') as inpt:
		with open('data/temp2.tsv','w') as outpt:
			outpt.write(inpt.readline())
			for line in inpt:
				s = line.split("\t")## brcd hugo entrez
				key_cna = s[0]+s[2]
				if key_cna in cna.keys():
					s[-1] = cna[key_cna]  +"\n"
				current = "\t".join(s)
				outpt.write(current)
	cna = {} ## 'Hugo_Symbol', 'Entrez_Gene_Id'
	with open('data/for10step-2.tsv','r') as inpt:
		for line in inpt:
			s = line.split("\t")
			key = s[0]+s[1]
			cna[key] = s[-1][:-1]
	#print ("step 6 almost done")
	with open('data/temp2.tsv','r') as inpt:
		with open(output_file,'w') as outpt:
			outpt.write(inpt.readline())
			for line in inpt:
				s = line.split("\t")## brcd hugo entrez
				key_cna = s[0]+s[2]
				if key_cna in cna.keys():
					s[-1] = cna[key_cna]  +"\n"
				current = "\t".join(s)
				outpt.write(current)
	os.remove('data/temp.tsv')
	os.remove('data/temp2.tsv')
	print ("step 10 end")
```
