## Supplementary material for "Comprehensive patient-level classification and quantification of driver events in TCGA PanCanAtlas cohorts": S8 Files: step_11.py

```
import os

def step11(algo):
	print ("step 11 start")
	input_file = algo[:-4] +'/data/'+ algo[:-4] +'_genes_level1.tsv'
	algo_name = algo[:-4]
	try:
		os.mkdir(algo_name)
	except:
		pass
	output_file = algo[:-4] +'/data/'+algo[:-4] + '_genes_level2.tsv'
	Key_driver = [r'Driver type',r'Number of hyperactivating SNAs + Number of inactivating SNAs',r'Number of inactivating SNAs',
	r'HISR',r'CNA status',r'Count as ... driver event(s)']
	with open (input_file,'r') as inpt:
		with open(output_file,'w') as o:
			o.write("\t".join(inpt.readline().split("\t")[:-1])+"\n")
			for i in inpt:
				s = i.split("\t")
				s[-1] = s[-1][:-1]
				if s[5]!='':
					if int(s[3]) + int(s[4])>= 1 and int(s[4]) == 0 and float(s[5]) > 5 and int(s[6]) ==0:
						s.append("SNA-based oncogene")
						s.append("1")
					elif int(s[3]) + int(s[4]) == 0 and int(s[4]) == 0 and float(s[5]) > 5 and int(s[6]) in [1,2]:
						s.append("CNA-based oncogene")
						s.append("1")
					elif int(s[3]) + int(s[4]) >= 1 and int(s[4]) == 0 and float(s[5]) > 5 and int(s[6]) in [1,2]:
						s.append("Mixed oncogene")
						s.append("1")
					elif int(s[3]) + int(s[4]) >= 1 and int(s[4]) >= 0 and float(s[5]) <= 5 and int(s[6]) ==0:
						s.append("SNA-based tumor suppressor")
						s.append("1")
					elif int(s[3]) + int(s[4]) == 0 and int(s[4]) == 0 and float(s[5]) <= 5 and int(s[6]) in [-1,-2]:
						s.append("CNA-based tumor suppressor")
						s.append("1")
					elif int(s[3]) + int(s[4]) >= 1 and int(s[4]) >= 0 and float(s[5]) <= 5 and int(s[6]) in [-1,-2]:
						s.append("Mixed tumor suppressor")
						s.append("1")
					elif int(s[3]) + int(s[4]) == 0 and int(s[4]) == 0 and int(s[6]) ==0:
						s.append("Passenger")
						s.append("0")
					else:
						s.append("Low-probability driver")
						s.append("0")
				else:
					if int(s[3]) + int(s[4]) == 0 and int(s[4]) == 0 and int(s[6]) ==0:
						s.append("Passenger")
						s.append("0")
					else:
						s.append("Low-probability driver")
						s.append("0")
				o.write("\t".join(s)+"\n")
	print ("step 11 end") 
```
