## Supplementary material for "Comprehensive patient-level classification and quantification of driver events in TCGA PanCanAtlas cohorts": S8 Files: step_12.py

```
from collections import defaultdict
import os
def step12(algo):
	print ("step 12 start")
	input_file = algo[:-4] + '/data/'+algo[:-4] +'_genes_level2.tsv'
	output_file = algo[:-4] +'/data/'+algo[:-4] + '_patients.tsv'
	chromosome = 'data/Chromosome_drivers_FDR5.tsv'
	algo_name = algo[:-4]

	try:
		os.mkdir(algo_name)
	except:
		pass
	arm = 'data/Arm_drivers_FDR5.tsv'
	seer_file = 'data/clinical_PANCAN_patient_with_followup_primary_whitelisted.tsv'
	header =[ 'Full TCGA barcode','Cancer type','Gender',
        'Age','Tumor stage','Number of SNA-based oncogenic events','Number of CNA-based oncogenic events',
        'Number of Mixed oncogenic events','Number of SNA-based tumor suppressor events',
         'Number of CNA-based tumor suppressor events',
        'Number of Mixed tumor suppressor events',
         'Number of Driver chromosome losses','Number of Driver chromosome gains',
        'Number of Driver arm losses','Number of Driver arm gains','Total number of driver events']


	s1 = ['I or II NOS','Stage I', 'Stage IA', 'Stage IA1', 'Stage IB', 'Stage IB1', 'Stage IB2',
		'Stage IC','Stage IS','T1', 'T1a', 'T1a1', 'T1b', 'T1b1', 'T1b2', 'T1c']

	s2 = ['Stage II', 'Stage IIA', 'Stage IIA1', 'Stage IIA2', 'Stage IIB', 'Stage IIC','T2',
		'T2a', 'T2a1', 'T2a2', 'T2b', 'T2c']

	s3 = ['Stage III', 'Stage IIIA', 'Stage IIIB', 'Stage IIIC', 'Stage IIIC1','Stage IIIC2',
		'T3', 'T3a', 'T3b', 'T3c']

	s4 = ['Stage IV','Stage IVA','Stage IVB','Stage IVC','T4','T4a','T4b','T4c','T4d','T4e']

	level_seer = defaultdict()  
	allc = set()
	s36 = ['[Discrepancy]','[Not Applicable]','[Not Available]','[Unknown]']
	s38 = [ '[Not Applicable]','[Not Available]']
	s31 = ['[Discrepancy]','[Not Applicable]', '[Not Available]']
	s39 = ['[Not Applicable]', '[Not Available]', '[Unknown]']
	with open (seer_file,'r') as inpt:
		kk =inpt.readline().split("\t")
		for line in inpt:
			s = line.split("\t")
			allc.add(s[36])
			allc.add(s[38])
			allc.add(s[31])
			allc.add(s[39])
			if s[1] not in level_seer.keys():
				level_seer[s[1]] = [0,0,0,0]
			if s[36] not in s36:
				level_seer[s[1]][3] = s[36]
			elif s[38] not in s38:
				level_seer[s[1]][3] = s[38]
			elif s[31] not in s31:
				level_seer[s[1]][3] = s[31]
			else: 
				level_seer[s[1]][3] = s[39]
			level_seer[s[1]][0] = s[2]#+
			#level_seer[s[1]][1] = s[15]
			#level_seer[s[1]][2] = s[12]
			level_seer[s[1]][1] = s[3]#+
			level_seer[s[1]][2] = s[9]#+
			
			if level_seer[s[1]][3] in s1:
				level_seer[s[1]][3] = 'I'
			if level_seer[s[1]][3] in s2:
				level_seer[s[1]][3] = 'II'
			if level_seer[s[1]][3] in s3:
				level_seer[s[1]][3] = 'III'
			if level_seer[s[1]][3] in s4:
				level_seer[s[1]][3] = 'IV'

	Key_driver = [r'Driver type',r'Number of hyperactivating SNAs',r'Number of inactivating SNAs',
	r'HISR',r'CNA status',r'Count as ... driver event(s)']

	level2 = defaultdict()  
	with open (input_file,'r') as inpt:
		inpt.readline().split("\t")
		for i in inpt:
			s = i.split("\t")
			if s[0] not in level2.keys():
				level2[s[0]] = [0,0,0,0,0,0]
			if 'SNA-based oncoge' in s[-2]:
				level2[s[0]][0]+=int(s[-1])
			if 'CNA-based oncoge' in s[-2]:
				level2[s[0]][1]+=int(s[-1])
			if 'Mixed oncoge' in s[-2]:
				level2[s[0]][2]+=int(s[-1])
			if 'SNA-based tumor sup' in s[-2]:
				level2[s[0]][3]+=int(s[-1])
			if 'CNA-based tumor sup' in s[-2]:
				level2[s[0]][4]+=int(s[-1])
			if 'Mixed tumor supp' in s[-2]:
				level2[s[0]][5]+=int(s[-1])
	level_chromosome = defaultdict()
	with open (chromosome,'r') as inpt:
		inpt.readline()
		for i in inpt:
			s = i.split("\t")
			s[0] = s[0][:-3]
			if s[0] not in level_chromosome.keys():
				level_chromosome[s[0]] = [0,0] #dc gains,dc losses
			level_chromosome[s[0]][0]+=int(s[-2])
			level_chromosome[s[0]][1]+=int(s[-3])
	level_arm = defaultdict()
	with open (arm,'r') as inpt:
		inpt.readline()
		for i in inpt:
			s = i.split("\t")
			s[0] = s[0][:-3]
			if s[0] not in level_arm.keys():
				level_arm[s[0]] = [0,0] #dc gains,dc losses
			level_arm[s[0]][0]+=int(s[-2])
			level_arm[s[0]][1]+=int(s[-3])
	with open(output_file,'w') as out:
		k = 0
		out.write("\t".join(header)+"\n")
		for brcd in level_seer.keys():
			flag = 1
			line_type = "\t".join(level_seer[brcd])
			if brcd in level2.keys():
				l2 = level2[brcd]
				l2 = list(map(lambda x:str(x),l2))
				line_2 = "\t".join(l2) 
			else:
				k+=1
				l2 = ["0" for i in range(6)]
				line_2 = "\t".join(l2) 
			if brcd in level_arm.keys():
				line_arm = "\t".join([str(level_arm[brcd][0]),str(level_arm[brcd][1])])
			else:
				line_arm = "\t".join(['0','0'])
			if brcd in level_chromosome.keys():
				line_chromosome = "\t".join([str(level_chromosome[brcd][0]),str(level_chromosome[brcd][1])])
			else:
				line_chromosome = "\t".join(['0','0'])
			if brcd in level2.keys():
				l2 = 0
				ad_sum = int(line_chromosome.split("\t")[0])+int(line_chromosome.split("\t")[1])
				ad_sum += int(line_arm.split("\t")[0])+int(line_arm.split("\t")[1])
				ad_sum += int(line_2.split("\t")[0])+int(line_2.split("\t")[1])+int(line_2.split("\t")[2])
				ad_sum += int(line_2.split("\t")[3])+int(line_2.split("\t")[4])+int(line_2.split("\t")[5])
				line_total = str(l2+ad_sum)
			else:
				l2 ='0'
				line_total = l2 
			line =brcd +"\t"+ line_type+"\t"+line_2+"\t"+line_chromosome+"\t"+ line_arm+"\t"+line_total+"\n"
			if flag == 1: out.write(line)
			if flag == 0: pass#print (level_seer[brcd])
	print ("step 12 end")
```
