## Supplementary material for "Comprehensive patient-level classification and quantification of driver events in TCGA PanCanAtlas cohorts": S8 Files: step_13.py

```
from tqdm import tqdm
import os
import numpy as np
import matplotlib
matplotlib.rcParams.update({'figure.max_open_warning':0})
import matplotlib.pyplot as plt
import pandas as pd
from utils.utils import med, move, sumVect, cancers, genders

def step13(algo):
	print ("step 13 start")
	input_file = algo[:-4] + '/data/'+algo[:-4] +'_patients.tsv'
	algo_name = algo[:-4]

	try:
		os.mkdir(algo_name)
	except:
		pass
	try:
		os.mkdir(algo_name+"/patient distributions")
	except:
		pass

	header = ['Cancer type']


	for i in range(100):
		header.append('Number of patients with {} driver events'.format(i))
	header.append('Number of patients with >=100 driver events')
	header = "\t".join(header)+"\n"

	table = {}
	with open(input_file,'r') as inpt:
		inpt.readline().split("\t")[7]
		for line in inpt:
			s = line.split("\t")
			if s[1]+"_"+s[2] not in table.keys():
				table[s[1]+"_"+s[2]] = [0 for i in range(101)]
			if int(s[-1])<100:
				table[s[1]+"_"+s[2]][int(s[-1])]+=1
			elif int(s[-1])==100:
				table[s[1]+"_"+s[2]][-1]+=1
	for g in genders:
		with open(algo_name+'/data/'+algo_name+"_distribution_events_"+g+".tsv",'w') as out:
			out.write(header)
			total = [0 for i in range(101)]
			for c in cancers:
				if c+"_"+g in table.keys():
					out.write(c+"\t")
					out.write("\t".join(list(map(lambda x:str(x),table[c+"_"+g])))+"\n")
			### add PANCAN
					total =sumVect(table[c+"_"+g],total)
			out.write("PANCAN"+"\t")
			out.write("\t".join(list(map(lambda x:str(x),total)))+"\n")
			table["PANCAN"+"_"+g] = total

	for file in table.keys():
		if file not in ["BRCA_MALE"]:
			#data = move(table[file],2)[2:-2]
			data = table[file]
			x = range(len(data))
			#median = med(data)
			plt.figure(figsize=(15,10))
			ax = plt.gca()
			plt.title(file,pad = 18, fontdict = {'fontsize':30})
			ax.bar(x, data)
			plt.xlabel('Number of driver events', fontdict = {'fontsize':30})
			plt.ylabel('Number of patients', fontdict = {'fontsize':30})
			plt.xticks(fontsize = 19)
			plt.yticks(fontsize = 19)
			plt.savefig(algo_name+"/"+"/patient distributions/"+algo_name+"_"+file+".pdf",dpi = 600)

	table = {}
	with open(input_file,'r') as inpt:
		inpt.readline().split("\t")[7]
		for line in inpt:
			s = line.split("\t")
			if s[1] not in table.keys():
				table[s[1]] = [0 for i in range(101)]
			if int(s[-1])<100:
				table[s[1]][int(s[-1])]+=1
			elif int(s[-1])==100:
				table[s[1]][-1]+=1

	with open(algo_name+'/data/'+algo_name+"_distribution_events"+".tsv",'w') as out:
		out.write(header)
		total = [0 for i in range(101)]
		for c in cancers:
			if c in table.keys():
				out.write(c+"\t")
				out.write("\t".join(list(map(lambda x:str(x),table[c])))+"\n")
				total =sumVect(table[c],total)
		out.write("PANCAN"+"\t")
		out.write("\t".join(list(map(lambda x:str(x),total)))+"\n")
		table["PANCAN"] = total
	for file in table.keys():
		#data = move(table[file],2)[2:-2]
		data = table[file]
		x = range(len(data))
		plt.figure(figsize=(15,10))
		ax = plt.gca()
		plt.title(file, pad = 18,fontdict = {'fontsize':30})
		ax.bar(x, data)
		plt.xlabel('Number of driver events', fontdict = {'fontsize':30})
		plt.ylabel('Number of patients', fontdict = {'fontsize':30})
		plt.xticks(fontsize = 19)
		plt.yticks(fontsize = 19)
		plt.savefig(algo_name+"/"+"/patient distributions/"+algo_name+"_"+file+".pdf",dpi = 600)
	print ("step 13 end")
```
