## Supplementary material for "Comprehensive patient-level classification and quantification of driver events in TCGA PanCanAtlas cohorts": S8 Files: step_14.py

```
import os
import numpy as np
import matplotlib
import matplotlib.pyplot as plt
from utils.utils import  sumVect, cancers, genders, rows, sort
import shutil
def step14(algo):
	print ("step 14 start")
	input_file = algo[:-4] + '/data/'+algo[:-4] +'_patients.tsv'
	algo_name = algo[:-4]

	try:
		os.mkdir(algo_name)
	except:
		pass
	try:
		os.mkdir(algo_name+"/cumulative histograms")
	except:
		pass
	try:
		shutil.copyfile('legeng.png',algo_name+"/cumulative histograms/"+'legeng.png')
	except:
		pass
	try:
		shutil.copyfile('legend.png',algo_name+"/cumulative histograms/"+'legend.png')
	except:
		pass
	header = "Total number of driver events in a patient"+"\t"+"\t".join(rows)+"\n"
	total_events = {}
	male_events = {}
	female_events = {}
	with open(input_file,'r') as inpt:
		inpt.readline().split("\t")
		for line in inpt:
			s = line.split("\t")
			s[-1] = s[-1][:-1]
			key = s[-1]
			if int(key)>=1 and int(key)<=100:
				if key not in total_events.keys():
					total_events[key] = [0 for i in range(11)]
					total_events[key].append(set())
				total_events[key][-1].add(s[0]) #add patient
				for i in range(11):
					total_events[key][i] += int(s[5+i])
			###males
				if s[2] == "MALE":
					if key not in male_events.keys():
						male_events[key] = [0 for i in range(11)]
						male_events[key].append(set())
					male_events[key][-1].add(s[0]) #add patient
					for i in range(11):
						male_events[key][i] += int(s[5+i])
			###females
				if s[2] == "FEMALE":
					if key not in female_events.keys():
						female_events[key] = [0 for i in range(11)]
						female_events[key].append(set())
					female_events[key][-1].add(s[0]) #add patient
					for i in range(11):
						female_events[key][i] += int(s[5+i])

	with open(algo_name+'/data/'+algo_name+"_distribution_events_detailed.tsv",'w') as out:
		out.write(header)
		types_plot = {}
		for c in sort(total_events.keys()):
			out.write(c+"\t")
			num = len(total_events[c][-1])
			a = total_events[c][:-1]
			a = list(map(lambda x:x/num,a))
			types_plot[c] = a
			line = "\t".join(list(map(lambda x:str(x),a)))+"\n"
			out.write(line)

	for i in range (1,101):
		if str(i) not in types_plot.keys():
			types_plot[str(i)] = [0.0 for  i in range(11)]
	data = types_plot
	x = [str(i) for i in range (1,101)]
	fig, ax = plt.subplots()
	title = 'Driver event distribution by total number of driver events per patient'
	for i in x:
		ax.bar(i, data[i][0],color = 'lightcoral')
		bott = data[i][0]
		ax.bar(i, data[i][1], bottom = bott,color = 'red')
		bott+= data[i][1]
		ax.bar(i, data[i][2], bottom = bott,color = 'maroon')
		bott+= data[i][2]
		ax.bar(i, data[i][3], bottom = bott,color = 'cyan')
		bott+= data[i][3]
		ax.bar(i, data[i][4], bottom = bott,color = 'c')
		bott+= data[i][4]
		ax.bar(i, data[i][5], bottom = bott,color = 'teal')
		bott+= data[i][5]
		ax.bar(i, data[i][6], bottom = bott,color = 'orange')
		bott+= data[i][6]
		ax.bar(i, data[i][7], bottom = bott,color = 'darkorange')
		bott+= data[i][7]
		ax.bar(i, data[i][8], bottom = bott,color = 'violet')
		bott+= data[i][8]
		ax.bar(i, data[i][9], bottom = bott,color = 'purple')
	ax.set_ylabel('Average number of driver events', fontdict = {'fontsize':30})
	ax.set_xlabel('Total number of driver events per patient', fontdict = {'fontsize':30})
	ax.set_title(title, pad = 18,fontdict = {'fontsize':30})
	plt.xticks(rotation = 90,fontsize = 14)
	plt.yticks(fontsize = 19)
	fig.set_figwidth(20)
	fig.set_figheight(16)
	fig.set_facecolor('floralwhite')
	ax.set_facecolor('seashell')
	plt.savefig(algo_name+"/"+"/cumulative histograms/"+algo_name+"_"+"distribution_events_detailed"+".pdf",dpi = 600)

	with open(algo_name+'/data/'+algo_name+"_distribution_events_detailed_males.tsv",'w') as out:
		out.write(header)
		males_plot= {}
		for c in sort(male_events.keys()):
			out.write(c+"\t")
			num = len(male_events[c][-1])
			a = male_events[c][:-1]
			a = list(map(lambda x:x/num,a))
			males_plot[c] = a
			line = "\t".join(list(map(lambda x:str(x),a)))+"\n"
			out.write(line)
	for i in range (1,101):
		if str(i) not in males_plot.keys():
			males_plot[str(i)] = [0.0 for  i in range(11)]		
	data = males_plot
	x = [str(i) for i in range (1,101)]
	fig, ax = plt.subplots()
	title = 'Driver event distribution by total number of driver events per patient in males'
	for i in x:
		ax.bar(i, data[i][0],color = 'lightcoral')
		bott = data[i][0]
		ax.bar(i, data[i][1], bottom = bott,color = 'red')
		bott+= data[i][1]
		ax.bar(i, data[i][2], bottom = bott,color = 'maroon')
		bott+= data[i][2]
		ax.bar(i, data[i][3], bottom = bott,color = 'cyan')
		bott+= data[i][3]
		ax.bar(i, data[i][4], bottom = bott,color = 'c')
		bott+= data[i][4]
		ax.bar(i, data[i][5], bottom = bott,color = 'teal')
		bott+= data[i][5]
		ax.bar(i, data[i][6], bottom = bott,color = 'orange')
		bott+= data[i][6]
		ax.bar(i, data[i][7], bottom = bott,color = 'darkorange')
		bott+= data[i][7]
		ax.bar(i, data[i][8], bottom = bott,color = 'violet')
		bott+= data[i][8]
		ax.bar(i, data[i][9], bottom = bott,color = 'purple')
	ax.set_ylabel('Average number of driver events', fontdict = {'fontsize':30})
	ax.set_xlabel('Total number of driver events per patient', fontdict = {'fontsize':30})
	ax.set_title(title, pad = 18,fontdict = {'fontsize':30})
	plt.xticks(rotation = 90,fontsize = 14)
	plt.yticks(fontsize = 19)
	fig.set_figwidth(20)
	fig.set_figheight(16)
	fig.set_facecolor('floralwhite')
	ax.set_facecolor('seashell')
	plt.savefig(algo_name+"/"+"/cumulative histograms/"+algo_name+"_"+"distribution_events_detailed_males"+".pdf",dpi = 600)

	with open(algo_name+'/data/'+algo_name+"_distribution_events_detailed_females.tsv",'w') as out:
		out.write(header)
		females_plot = {}
		for c in sort(female_events.keys()):
			out.write(c+"\t")
			num = len(female_events[c][-1])
			a = female_events[c][:-1]
			a = list(map(lambda x:x/num,a))
			females_plot[c] = a
			line = "\t".join(list(map(lambda x:str(x),a)))+"\n"
			out.write(line)
	for i in range (1,101):
		if str(i) not in females_plot.keys():
			females_plot[str(i)] = [0.0 for  i in range(11)]
	data = females_plot
	x = [str(i) for i in range (1,101)]
	fig, ax = plt.subplots()
	title = 'Driver event distribution by total number of driver events per patient in females'
	for i in x:
		ax.bar(i, data[i][0],color = 'lightcoral')
		bott = data[i][0]
		ax.bar(i, data[i][1], bottom = bott,color = 'red')
		bott+= data[i][1]
		ax.bar(i, data[i][2], bottom = bott,color = 'maroon')
		bott+= data[i][2]
		ax.bar(i, data[i][3], bottom = bott,color = 'cyan')
		bott+= data[i][3]
		ax.bar(i, data[i][4], bottom = bott,color = 'c')
		bott+= data[i][4]
		ax.bar(i, data[i][5], bottom = bott,color = 'teal')
		bott+= data[i][5]
		ax.bar(i, data[i][6], bottom = bott,color = 'orange')
		bott+= data[i][6]
		ax.bar(i, data[i][7], bottom = bott,color = 'darkorange')
		bott+= data[i][7]
		ax.bar(i, data[i][8], bottom = bott,color = 'violet')
		bott+= data[i][8]
		ax.bar(i, data[i][9], bottom = bott,color = 'purple')
	ax.set_ylabel('Average number of driver events', fontdict = {'fontsize':30})
	ax.set_xlabel('Total number of driver events per patient', fontdict = {'fontsize':30})
	ax.set_title(title,pad = 18, fontdict = {'fontsize':30})
	plt.xticks(rotation = 90,fontsize = 14)
	plt.yticks(fontsize = 19)
	fig.set_figwidth(20)
	fig.set_figheight(16)
	fig.set_facecolor('floralwhite')
	ax.set_facecolor('seashell')
	plt.savefig(algo_name+"/"+"/cumulative histograms/"+algo_name+"_"+"distribution_events_detailed_females"+".pdf",dpi = 600)
	print ("step 14 end")
```
