## Supplementary material for "Comprehensive patient-level classification and quantification of driver events in TCGA PanCanAtlas cohorts": S8 Files: step_15.py

```
import numpy as np
import matplotlib.pyplot as plt
import os
import matplotlib
import scipy.stats
import pandas as pd
from utils.utils import  sumVect, cancers, genders, rows

def step15(algo):
	print ("step 15 start")
	input_file = algo[:-4] +'/data/'+algo[:-4] + '_patients.tsv'
	algo_name = algo[:-4]
	try:
		os.mkdir(algo_name)
	except:
		pass
	try:
		os.mkdir(algo_name+"/cumulative histograms")
	except:
		pass
		
	#try:
	#	os.mkdir(algo_name+"/erlang")
	#except:
	#	pass

	#intervals95 = {}
	#intr = pd.read_table("data/erlang_adult_r2_out_intervals_095.tsv",sep = "\t")
	#for i in intr.columns[1:]:
	#	mi = intr[i][1]
	#	ma = intr[i][2]
	#	if np.isnan(mi) :
	#		mi = intr[i][0]
	#	if np.isnan(ma):
	#		ma = intr[i][0]
	#	intervals95[i.upper()[:-1]] = (mi,ma,intr[i][0])

	#intervals99 = {}
	#intr = pd.read_table("data/erlang_adult_r2_out_intervals_099.tsv",sep = "\t")
	#for i in intr.columns[1:]:
	#	mi = intr[i][1]
	#	ma = intr[i][2]
	#	if np.isnan(mi) :
	#		mi = intr[i][0]
	#	if np.isnan(ma):
	#		ma = intr[i][0]
	#	intervals99[i.upper()[:-1]] = (mi,ma,intr[i][0])

	header = "Cancer type"+"\t"+"\t".join(rows)+"\n"

	total_events = {}
	male_events ={}
	female_events = {}


	with open(input_file,'r') as inpt:
		inpt.readline().split("\t")
		
		for line in inpt:
			s = line.split("\t")
			if s[1] not in total_events.keys():
				total_events[s[1]] = [0 for i in range(11)]
				total_events[s[1]].append(set())
			total_events[s[1]][-1].add(s[0]) #add patient
			for i in range(11):
				total_events[s[1]][i] += int(s[5+i])

			###males
			if s[2] == "MALE":
				if s[1] not in male_events.keys():
					male_events[s[1]] = [0 for i in range(11)]
					male_events[s[1]].append(set())
				male_events[s[1]][-1].add(s[0]) #add patient
				for i in range(11):
					male_events[s[1]][i] += int(s[5+i])
			###females    
			if s[2] == "FEMALE":
				if s[1] not in female_events.keys():
					female_events[s[1]] = [0 for i in range(11)]
					female_events[s[1]].append(set())
				female_events[s[1]][-1].add(s[0]) #add patient
				for i in range(11):
					female_events[s[1]][i] += int(s[5+i])
					
	total = [0 for i in range(11)]
	all_p = set()
	for i in total_events.keys():
		total =sumVect(total_events[i][:-1],total)
		for j in total_events[i][-1]:
			all_p.add(j)
	total_events['PANCAN'] = total
	total_events['PANCAN'].append(all_p)

	total = [0 for i in range(11)]
	all_p = set()
	for i in male_events.keys():
		total =sumVect(male_events[i][:-1],total)
		for j in male_events[i][-1]:
			all_p.add(j)
	male_events['PANCAN'] = total
	male_events['PANCAN'].append(all_p)
	total = [0 for i in range(11)]
	all_p = set()
	for i in female_events.keys():
		total =sumVect(female_events[i][:-1],total)
		for j in female_events[i][-1]:
			all_p.add(j)
	female_events['PANCAN'] = total
	female_events['PANCAN'].append(all_p)

	with open(algo_name+'/data/'+algo_name+"_distribution_cohorts.tsv",'w') as out:
		out.write(header)
		types_plot = {}
		for c in total_events.keys():
			out.write(c+"\t")
			num = len(total_events[c][-1])
			a = total_events[c][:-1]
			a = list(map(lambda x:x/num,a))
			types_plot[c] = a
			line = "\t".join(list(map(lambda x:str(x),a)))+"\n"
			out.write(line)

	x = total_events.keys()
	data = types_plot
	fig, ax = plt.subplots()
	title = 'Driver event distribution by cancer type'
	for i in x:
		ax.bar(i, data[i][0],color = 'lightcoral')
		bott = data[i][0]
		ax.bar(i, data[i][1], bottom = bott,color = 'red')
		bott+= data[i][1]
		ax.bar(i, data[i][2], bottom = bott,color = 'maroon')
		bott+= data[i][2]
		ax.bar(i, data[i][3], bottom = bott,color = 'cyan')
		bott+= data[i][3]
		ax.bar(i, data[i][4], bottom = bott,color = 'c')
		bott+= data[i][4]
		ax.bar(i, data[i][5], bottom = bott,color = 'teal')
		bott+= data[i][5]
		ax.bar(i, data[i][6], bottom = bott,color = 'orange')
		bott+= data[i][6]
		ax.bar(i, data[i][7], bottom = bott,color = 'darkorange')
		bott+= data[i][7]
		ax.bar(i, data[i][8], bottom = bott,color = 'violet')
		bott+= data[i][8]
		ax.bar(i, data[i][9], bottom = bott,color = 'purple')
	ax.set_ylabel('Average number of driver events', fontdict = {'fontsize':30})
	ax.set_xlabel('Cohorts', fontdict = {'fontsize':30})
	ax.set_title(title, pad = 18,fontdict = {'fontsize':30})
	plt.xticks(rotation = 90,fontsize = 19)
	plt.yticks(fontsize = 19)
	fig.set_figwidth(20)
	fig.set_figheight(16)
	fig.set_facecolor('floralwhite')
	ax.set_facecolor('seashell')
	plt.savefig(algo_name+"/"+"/cumulative histograms/"+algo_name+"_"+"distribution_cohorts"+".pdf",dpi = 600)

###MALES
	with open(algo_name+'/data/'+algo_name+"_distribution_cohorts_males.tsv",'w') as out:
		out.write(header)
		males_plot= {}
		for c in male_events.keys():
			out.write(c+"\t")
			num = len(male_events[c][-1])
			a = male_events[c][:-1]
			a = list(map(lambda x:x/num,a))
			males_plot[c] = a
			line = "\t".join(list(map(lambda x:str(x),a)))+"\n"
			out.write(line)
	x = male_events.keys()
	data = males_plot
	fig, ax = plt.subplots()
	title = 'Driver event distribution by cancer type in males'
	for i in x:
		ax.bar(i, data[i][0],color = 'lightcoral')
		bott = data[i][0]
		ax.bar(i, data[i][1], bottom = bott,color = 'red')
		bott+= data[i][1]
		ax.bar(i, data[i][2], bottom = bott,color = 'maroon')
		bott+= data[i][2]
		ax.bar(i, data[i][3], bottom = bott,color = 'cyan')
		bott+= data[i][3]
		ax.bar(i, data[i][4], bottom = bott,color = 'c')
		bott+= data[i][4]
		ax.bar(i, data[i][5], bottom = bott,color = 'teal')
		bott+= data[i][5]
		ax.bar(i, data[i][6], bottom = bott,color = 'orange')
		bott+= data[i][6]
		ax.bar(i, data[i][7], bottom = bott,color = 'darkorange')
		bott+= data[i][7]
		ax.bar(i, data[i][8], bottom = bott,color = 'violet')
		bott+= data[i][8]
		ax.bar(i, data[i][9], bottom = bott,color = 'purple')
	ax.set_ylabel('Average number of driver events', fontdict = {'fontsize':30})
	ax.set_xlabel('Cohorts', fontdict = {'fontsize':30})
	ax.set_title(title,pad = 18, fontdict = {'fontsize':30})
	plt.xticks(rotation = 90,fontsize = 19)
	plt.yticks(fontsize = 19)
	fig.set_figwidth(20)
	fig.set_figheight(16)
	fig.set_facecolor('floralwhite')
	ax.set_facecolor('seashell')
	plt.savefig(algo_name+"/"+"/cumulative histograms/"+algo_name+"_"+"distribution_cohorts_males"+".pdf",dpi = 600)

### FEMALES
	with open(algo_name+'/data/'+algo_name+"_distribution_cohorts_females.tsv",'w') as out:
		out.write(header)
		females_plot = {}
		for c in female_events.keys():
			out.write(c+"\t")
			num = len(female_events[c][-1])
			a = female_events[c][:-1]
			a = list(map(lambda x:x/num,a))
			females_plot[c] = a
			line = "\t".join(list(map(lambda x:str(x),a)))+"\n"
			out.write(line)

	x = female_events.keys()
	data = females_plot
	fig, ax = plt.subplots()
	title = 'Driver event distribution by cancer type in females'
	for i in x:
		ax.bar(i, data[i][0],color = 'lightcoral')
		bott = data[i][0]
		ax.bar(i, data[i][1], bottom = bott,color = 'red')
		bott+= data[i][1]
		ax.bar(i, data[i][2], bottom = bott,color = 'maroon')
		bott+= data[i][2]
		ax.bar(i, data[i][3], bottom = bott,color = 'cyan')
		bott+= data[i][3]
		ax.bar(i, data[i][4], bottom = bott,color = 'c')
		bott+= data[i][4]
		ax.bar(i, data[i][5], bottom = bott,color = 'teal')
		bott+= data[i][5]
		ax.bar(i, data[i][6], bottom = bott,color = 'orange')
		bott+= data[i][6]
		ax.bar(i, data[i][7], bottom = bott,color = 'darkorange')
		bott+= data[i][7]
		ax.bar(i, data[i][8], bottom = bott,color = 'violet')
		bott+= data[i][8]
		ax.bar(i, data[i][9], bottom = bott,color = 'purple')
	ax.set_ylabel('Average number of driver events', fontdict = {'fontsize':30})
	ax.set_xlabel('Cohorts', fontdict = {'fontsize':30})
	ax.set_title(title,pad = 18, fontdict = {'fontsize':30})
	plt.xticks(rotation = 90,fontsize = 19)
	plt.yticks(fontsize = 19)
	fig.set_figwidth(20)
	fig.set_figheight(16)
	fig.set_facecolor('floralwhite')
	ax.set_facecolor('seashell')
	plt.savefig(algo_name+"/"+"/cumulative histograms/"+algo_name+"_"+"distribution_cohorts_females"+".pdf",dpi = 600)


	print ("step 15 start")
```
