## Supplementary material for "Comprehensive patient-level classification and quantification of driver events in TCGA PanCanAtlas cohorts": S8 Files: step_16.py

```
from tqdm import tqdm
import os
import numpy as np
import matplotlib
import matplotlib.pyplot as plt
from utils.utils import med, sumVect, cancers, genders,rows

def step16(algo):
	print ("step 16 start")
	input_file = algo[:-4] +'/data/'+algo[:-4] + '_patients.tsv'
	algo_name = algo[:-4]
	try:
		os.mkdir(algo_name)
	except:
		pass
	try:
		os.mkdir(algo_name+"/cumulative histograms")
	except:
		pass

	header = "Gender"+"\t"+"\t".join(rows)+"\n"

	male_events ={}
	female_events = {}
	total_events = {}

	with open(input_file,'r') as inpt:
		inpt.readline().split("\t")
		for line in inpt:
			s = line.split("\t")
			if s[2] not in total_events.keys():
				total_events[s[2]] = [0 for i in range(11)]
				total_events[s[2]].append(set())
			total_events[s[2]][-1].add(s[0]) #add patient
			for i in range(11):
				total_events[s[2]][i] += int(s[5+i])

	with open(algo_name+'/data/'+algo_name+"_distribution_gender.tsv",'w') as out:
		out.write(header)
		genders_plot = {}
		for c in total_events.keys():
			out.write(c+"\t")
			num = len(total_events[c][-1])
			a = total_events[c][:-1]
			a = list(map(lambda x:x/num,a))
			genders_plot[c] = a
			line = "\t".join(list(map(lambda x:str(x),a)))+"\n"
			out.write(line)

	x = genders
	data = genders_plot
	fig, ax = plt.subplots()
	title = 'Driver event distribution by gender'
	for i in x:
		ax.bar(i, data[i][0],color = 'lightcoral')
		bott = data[i][0]
		ax.bar(i, data[i][1], bottom = bott,color = 'red')
		bott+= data[i][1]
		ax.bar(i, data[i][2], bottom = bott,color = 'maroon')
		bott+= data[i][2]
		ax.bar(i, data[i][3], bottom = bott,color = 'cyan')
		bott+= data[i][3]
		ax.bar(i, data[i][4], bottom = bott,color = 'c')
		bott+= data[i][4]
		ax.bar(i, data[i][5], bottom = bott,color = 'teal')
		bott+= data[i][5]
		ax.bar(i, data[i][6], bottom = bott,color = 'orange')
		bott+= data[i][6]
		ax.bar(i, data[i][7], bottom = bott,color = 'darkorange')
		bott+= data[i][7]
		ax.bar(i, data[i][8], bottom = bott,color = 'violet')
		bott+= data[i][8]
		ax.bar(i, data[i][9], bottom = bott,color = 'purple')
	ax.set_ylabel('Average number of driver events', fontdict = {'fontsize':30})
	ax.set_title(title,pad = 18, fontdict = {'fontsize':30})
	plt.yticks(fontsize = 19)
	plt.xticks(fontsize = 19)
	fig.set_figwidth(10)
	fig.set_figheight(16)
	fig.set_facecolor('floralwhite')
	ax.set_facecolor('seashell')
	plt.savefig(algo_name+"/"+"/cumulative histograms/"+algo_name+"_"+"distribution_gender"+".pdf",dpi = 600)
	print ("step 16 end")
```
