## Supplementary material for "Comprehensive patient-level classification and quantification of driver events in TCGA PanCanAtlas cohorts": S8 Files: step_18.py

```
import os
import numpy as np
import matplotlib
import matplotlib.pyplot as plt
from utils.utils import med, sumVect, cancers, genders,rows

def step18(algo):
	print ("step 18 start")
	input_file = algo[:-4] + '/data/'+algo[:-4] +'_patients.tsv'
	algo_name = algo[:-4]

	try:
		os.mkdir(algo_name)
	except:
		pass
	try:
		os.mkdir(algo_name+"/cumulative histograms")
	except:
		pass

	header = "Age group"+"\t"+"\t".join(rows)+"\n"

	ages = ['<25','25-29', '30-34','35-39', '40-44',
	 '45-49', '50-54','55-59','60-64','65-69','70-74','75-79', '80-84','>=85']
	ages1 = list(map(lambda x:int(x.split("-")[0]),ages[1:-1]))


	total_events = {}
	male_events ={}
	female_events = {}


	with open(input_file,'r') as inpt:
		inpt.readline().split("\t")[5]
		for line in inpt:
			s = line.split("\t")
			if s[3]!='[Not Available]':
				if int(s[3])>=85:
					key = '>=85'
				elif int(s[3])<25:
					key = '<25'
				else:
					age = int(s[3])//5
					age = age*5
					key = ages[ages1.index(age)+1]
				if key not in total_events.keys():
					total_events[key] = [0 for i in range(11)]
					total_events[key].append(set())
				total_events[key][-1].add(s[0]) #add patient
				for i in range(11):
					total_events[key][i] += int(s[5+i])

			###males
				if s[2] == "MALE":
					if key not in male_events.keys():
						male_events[key] = [0 for i in range(11)]
						male_events[key].append(set())
					male_events[key][-1].add(s[0]) #add patient
					for i in range(11):
						male_events[key][i] += int(s[5+i])
			###females
				if s[2] == "FEMALE":
					if key not in female_events.keys():
						female_events[key] = [0 for i in range(11)]
						female_events[key].append(set())
					female_events[key][-1].add(s[0]) #add patient
					for i in range(11):
						female_events[key][i] += int(s[5+i])

	total = ['<25','25-29','30-34', '35-39', '40-44', '45-49', '50-54', '55-59', '60-64', '65-69', '70-74', '75-79', '80-84', '>=85']
	male = total
	female = total

	with open(algo_name+'/data/'+algo_name+"_distribution_age.tsv",'w') as out:
		out.write(header)
		ages_plot = {}
		for c in total:
			out.write(c+"\t")
			num = len(total_events[c][-1])
			a = total_events[c][:-1]
			a = list(map(lambda x:x/num,a))
			ages_plot[c] = a
			line = "\t".join(list(map(lambda x:str(x),a)))+"\n"
			out.write(line)

	x = total
	data = ages_plot
	fig, ax = plt.subplots()
	title = 'Driver event distribution by age'
	for i in x:
		ax.bar(i, data[i][0],color = 'lightcoral')
		bott = data[i][0]
		ax.bar(i, data[i][1], bottom = bott,color = 'red')
		bott+= data[i][1]
		ax.bar(i, data[i][2], bottom = bott,color = 'maroon')
		bott+= data[i][2]
		ax.bar(i, data[i][3], bottom = bott,color = 'cyan')
		bott+= data[i][3]
		ax.bar(i, data[i][4], bottom = bott,color = 'c')
		bott+= data[i][4]
		ax.bar(i, data[i][5], bottom = bott,color = 'teal')
		bott+= data[i][5]
		ax.bar(i, data[i][6], bottom = bott,color = 'orange')
		bott+= data[i][6]
		ax.bar(i, data[i][7], bottom = bott,color = 'darkorange')
		bott+= data[i][7]
		ax.bar(i, data[i][8], bottom = bott,color = 'violet')
		bott+= data[i][8]
		ax.bar(i, data[i][9], bottom = bott,color = 'purple')
	ax.set_ylabel('Average number of driver events', fontdict = {'fontsize':30})
	ax.set_xlabel('Age', fontdict = {'fontsize':30})
	ax.set_title(title,pad = 18, fontdict = {'fontsize':30})
	plt.xticks(rotation = 90,fontsize = 19)
	plt.yticks(fontsize = 19)
	fig.set_figwidth(10)
	fig.set_figheight(16)
	fig.set_facecolor('floralwhite')
	ax.set_facecolor('seashell')
	plt.savefig(algo_name+"/"+"/cumulative histograms/"+algo_name+"_"+"distribution_age"+".pdf",dpi = 600)

	with open(algo_name+'/data/'+algo_name+"_distribution_age_males.tsv",'w') as out:
		out.write(header)
		males_plot = {}
		for c in male:
			out.write(c+"\t")
			num = len(male_events[c][-1])
			a = male_events[c][:-1]
			a = list(map(lambda x:x/num,a))
			males_plot[c] = a
			line = "\t".join(list(map(lambda x:str(x),a)))+"\n"
			out.write(line)
	x = total
	data = males_plot
	fig, ax = plt.subplots()
	title = 'Driver event distribution by age in males'
	for i in x:
		ax.bar(i, data[i][0],color = 'lightcoral')
		bott = data[i][0]
		ax.bar(i, data[i][1], bottom = bott,color = 'red')
		bott+= data[i][1]
		ax.bar(i, data[i][2], bottom = bott,color = 'maroon')
		bott+= data[i][2]
		ax.bar(i, data[i][3], bottom = bott,color = 'cyan')
		bott+= data[i][3]
		ax.bar(i, data[i][4], bottom = bott,color = 'c')
		bott+= data[i][4]
		ax.bar(i, data[i][5], bottom = bott,color = 'teal')
		bott+= data[i][5]
		ax.bar(i, data[i][6], bottom = bott,color = 'orange')
		bott+= data[i][6]
		ax.bar(i, data[i][7], bottom = bott,color = 'darkorange')
		bott+= data[i][7]
		ax.bar(i, data[i][8], bottom = bott,color = 'violet')
		bott+= data[i][8]
		ax.bar(i, data[i][9], bottom = bott,color = 'purple')
	ax.set_ylabel('Average number of driver events', fontdict = {'fontsize':30})
	ax.set_xlabel('Age', fontdict = {'fontsize':30})
	ax.set_title(title,pad = 18, fontdict = {'fontsize':30})
	plt.xticks(rotation = 90,fontsize = 19)
	plt.yticks(fontsize = 19)
	fig.set_figwidth(10)
	fig.set_figheight(16)
	fig.set_facecolor('floralwhite')
	ax.set_facecolor('seashell')
	plt.savefig(algo_name+"/"+"/cumulative histograms/"+algo_name+"_"+"distribution_age_males"+".pdf",dpi = 600)

	with open(algo_name+'/data/'+algo_name+"_distribution_age_females.tsv",'w') as out:
		out.write(header)
		females_plot = {}
		for c in female:
			out.write(c+"\t")
			num = len(female_events[c][-1])
			a = female_events[c][:-1]
			a = list(map(lambda x:x/num,a))
			females_plot[c] = a
			line = "\t".join(list(map(lambda x:str(x),a)))+"\n"
			out.write(line)
	x = total
	data = females_plot
	fig, ax = plt.subplots()
	title = 'Driver event distribution by age in females'
	for i in x:
		ax.bar(i, data[i][0],color = 'lightcoral')
		bott = data[i][0]
		ax.bar(i, data[i][1], bottom = bott,color = 'red')
		bott+= data[i][1]
		ax.bar(i, data[i][2], bottom = bott,color = 'maroon')
		bott+= data[i][2]
		ax.bar(i, data[i][3], bottom = bott,color = 'cyan')
		bott+= data[i][3]
		ax.bar(i, data[i][4], bottom = bott,color = 'c')
		bott+= data[i][4]
		ax.bar(i, data[i][5], bottom = bott,color = 'teal')
		bott+= data[i][5]
		ax.bar(i, data[i][6], bottom = bott,color = 'orange')
		bott+= data[i][6]
		ax.bar(i, data[i][7], bottom = bott,color = 'darkorange')
		bott+= data[i][7]
		ax.bar(i, data[i][8], bottom = bott,color = 'violet')
		bott+= data[i][8]
		ax.bar(i, data[i][9], bottom = bott,color = 'purple')
	ax.set_ylabel('Average number of driver events', fontdict = {'fontsize':30})
	ax.set_xlabel('Age', fontdict = {'fontsize':30})
	ax.set_title(title, pad = 18,fontdict = {'fontsize':30})
	plt.xticks(rotation = 90,fontsize = 19)
	plt.yticks(fontsize = 19)
	fig.set_figwidth(10)
	fig.set_figheight(16)
	fig.set_facecolor('floralwhite')
	ax.set_facecolor('seashell')
	plt.savefig(algo_name+"/"+"/cumulative histograms/"+algo_name+"_"+"distribution_age_females"+".pdf",dpi = 600)
	print ("step 18 end")
```
