## Supplementary material for "Comprehensive patient-level classification and quantification of driver events in TCGA PanCanAtlas cohorts": S8 Files: utils.py

```
def barcode_to_id(patient_barcode):
	return '-'.join(patient_barcode.split('-')[:3])

def sumVect(v1,v2):
	return list(map((lambda x: x[0]+x[1]),zip(v1,v2)))

def sort(keys):
	newkeys = []
	f = 0
	for i in keys:
		if i!= '>=50':
			newkeys.append(int(i))
		else: f = 1
	keys = sorted(newkeys)
	keys = list(map(lambda x:str(x),keys))
	if f == 0 :
		return keys
	else:
		keys.append(">=50")
		return(keys)


	total_events = {}
	male_events ={}
	female_events = {}

def med(d):
	s = 0
	for i in range(len(d)):
		s+=d[i]
		if s>=sum(d)/2:
			med = i
			break
	return med

def move(l,N):
	new = [0 for i in range(len(l))]
	for i in range(len(l)):
		new[i] = l[i]
	left = [0 for i in range(N)]
	right = [0 for i in range(N)]
	left.extend(new)
	left.extend(right)
	new2 = left
	new_ = [0 for i in range(len(l))]
	for i in range(len(l)):
		new_[i] = l[i]
	left_ = [0 for i in range(N)]
	right_ = [0 for i in range(N)]
	left_.extend(new_)
	left_.extend(right_)
	new3 = left_
	for i in range(N,len(new2)-N):
		for j in range(1,N+1):
			new2[i] +=(new3[i-j]+new3[i+j])
		new2[i] = new2[i]/(2*N+1)
	return new2

cancers = ['ACC', 'BLCA', 'BRCA', 'CESC', 'CHOL', 'COAD', 'DLBC', 'ESCA-AD', 'ESCA-SC', 'GBM', 'HNSC-OP', 
		   'HNSC-LA', 'HNSC', 'KICH', 'KIRC', 'KIRP', 'LGG', 'LIHC', 'LUAD', 'LUSC', 'MESO', 'OV', 'PAAD', 'PCPG', 
		   'PRAD', 'READ', 'SARC', 'SKCM', 'STAD', 'TGCT', 'THCA', 'THYM', 'UCEC', 'UCS', 'UVM', 'PANCAN']
genders = ['FEMALE', 'MALE']

rows = [
	'Average number of SNA-based oncogenic events per patient',
	'Average number of CNA-based oncogenic events per patient' ,
	'Average number of Mixed oncogenic events per patient' ,
	'Average number of SNA-based tumor suppressor events per patient' ,
	'Average number of CNA-based tumor suppressor events per patient' ,
	'Average number of Mixed tumor suppressor events per patient' ,
	'Average number of Driver chromosome losses per patient' ,
	'Average number of Driver chromosome gains per patient' ,
	'Average number of Driver arm losses per patient' ,
	'Average number of Driver arm gains per patient' ,
	'Average number of driver events of all classes per patient']
```
